# A wide-ranging *Pseudomonas aeruginosa* PeptideAtlas build: a useful proteomic resource for a versatile pathogen

**DOI:** 10.1101/2020.11.18.386490

**Authors:** J.A. Reales-Calderón, Z. Sun, V. Mascaraque, E. Pérez-Navarro, V. Vialás, E.W. Deutsch, RL Moritz, C. Gil, JL Martínez, G. Molero

## Abstract

*Pseudomonas aeruginosa* is an important opportunistic pathogen with high prevalence in nosocomial infections. This microorganism is a good model for understanding biological processes such as the quorum-sensing response, the metabolic integration of virulence, the mechanisms of global regulation of bacterial physiology, and the evolution of antibiotic resistance. Till now, *P. aeruginosa* proteomic data, although available in several on-line repositories, were dispersed and difficult to access. In the present work, proteomes of the PAO1 strain grown under very different conditions and from diverse cellular compartments have been analyzed and joined to build the *Pseudomonas* PeptideAtlas. This resource is a comprehensive mass spectrometry-derived peptide and inferred protein database with 71.3% coverage of the total predicted proteome of *P. aeruginosa* PAO1. This is the highest published coverage among the eight bacterial PeptideAtlas datasets currently available. The proteins in the *Pseudomonas* PeptideAtlas cover 84% of metabolic proteins, 71% of proteins involved in genetic information processing, 72% of proteins responsible for environmental information processing, more than 80% of proteins related to quorum sensing and biofilm formation, and 81% of proteins responsible for antimicrobial resistance. It exemplifies a necessary tool for targeted proteomics studies, system-wide observations, and cross-species observational studies. Here we describe how this resource was built and some of the physiologically important proteins of this pathogen.

**Significance:** *Pseudomonas aeruginosa* is among the most versatile bacterial pathogens. Studies of its proteome are very important as they can reveal virulence factors and mechanisms of antibiotic resistance. The construction of a proteomic resource such as the PeptideAtlas enables targeted proteomics studies, system-wide observations, and cross-species observational studies.

## 1. Introduction

*Pseudomonas aeruginosa* is an important opportunistic pathogen with high prevalence in nosocomial infections [1, 2]. In addition, this bacterial pathogen is the major cause of chronic infections in patients suffering obstructive pulmonary disease and cystic fibrosis (CF)[3–5]. It is able to survive, persist in, and colonize diverse environments [6], including soil and water. The phylogenetic analysis of clinical and environmental isolates of this organism show that the population is panmictic. There are no differentiated clinical and environmental lineages, and the success of infection relies more on the underlying situation of the patient than on the specific characteristics of a given clone that may have evolved towards virulence [7–9]. Differing from other more specific pathogens that have become virulent through the acquisition of genetic material [10], the virulence determinants of *P. aeruginosa* belong to the core genome of this microorganism [11]: they are present in all members of the species and have not been acquired by a specific lineage evolved to be infectious. Using similar mechanisms, *P. aeruginosa* is able to infect a large variety of hosts, including amoebae, plants, worms, insects, and mammals [12]. A highly versatile species, *P. aeruginosa* grows under aerobic and anaerobic conditions, over a wide temperature range, and can make use of several carbon sources [13]. Further, it presents a large repertoire of virulence determinants [13, 14]. The expression of several of them is regulated through a complex quorum-sensing network [15, 16], as well as a battery of secreted virulence factors (including toxins, siderophores and proteases) and vesicle-engulfed compounds [17–19]. One of the main problems in treating *P. aeruginosa* infections is the low susceptibility to antibiotics of this microorganism, being classified by WHO in the ESKAPE group of pathogens with higher risk of antibiotic resistance [14, 20]. This is due to the presence in its genome of genes encoding beta-lactamases and several multidrug efflux pumps [21], among other elements [22]. In addition*, P. aeruginosa* can evolve to acquire increasing levels of resistance through mutations or through the acquisition of antibiotic resistance genes *via* horizontal gene transfer, allowing the dissemination of multidrug resistant strains causing nosocomial infections worldwide [23].

The versatility of *P. aeruginosa* is supported by its large genome, ranging between 5 and 7 Mbp in different strains [6], together with a large number of regulatory elements (both transcriptional and posttranscriptional) that allow the organism to adapt its physiology when confronted with different physicochemical conditions and cues specific to each of the habitats it can colonize. All these features make this microorganism, in addition to an important pathogen, a good model for understanding several biological processes, including the quorum-sensing response [15, 16], metabolic integration of virulence [24, 25], mechanisms of global regulation of bacterial physiology [26, 27], and the evolution of antibiotic resistance [28–34], among others. For this reason, it has been proposed that *P. aeruginosa* should be considered a “lab rat”, similar to other model bacteria such as *Escherichia coli* and *Bacillus subtilis* [13].

The integrated analysis of *P. aeruginosa* under a systems biology umbrella requires the use of multi-omic techniques, and accurate databases are fundamental to this purpose. Among them, the on-line resource at the *Pseudomonas* Genomic Database (www.pseudomonas.com) [35] provides updated and well-curated information on the different sequenced strains of this microorganisms at the genomic level, while PAMDB, http://pseudomonas.umaryland.edu [36] includes metabolomic information. Given that post-transcriptional regulation plays a major role in the adaptation of this microorganism to its different habitats [37], and that the identification of factors that are secreted to the extracellular milieu or are parts of extracellular vesicles is fundamental to a full understanding of the virulence of this pathogen, proteomic studies are needed for a comprehensive analysis of *P. aeruginosa* physiology. However, proteomic data, although available in several on-line repositories, are dispersed and difficult to access. Thus, resources like PeptideAtlas, which compiles the available LC-MS/MS-based shotgun proteomics data, are necessary for targeted proteomics studies, system-wide observations, and cross-species observational studies [38]. To date, 43 PeptideAtlas builds are available, but only 8 of them are from bacteria (http://www.peptideatlas.org/builds/).

Here we present the *P. aeruginosa* PeptideAtlas. To build it, proteomes of *P. aeruginosa* strain PAO1 grown under very different conditions and from diverse cellular compartments have been obtained, analyzed, and joined with data from other authors, one of them including proteomes from clinical strains. The resulting on-line resource is a comprehensive mass spectrometry-derived peptide and inferred protein database with high coverage of the total predicted proteome from this bacterium.

## 2. Materials and Methods

### 2.1. Bacterial strains and growth conditions

The *P. aeruginosa* strain used by our group to develop the *Pseudomonas* PeptideAtlas was the widely used wild-type PAO1. All the growth conditions employed to generate new proteomes used to build the PeptideAtlas are represented in Figure 1 and Table 1, together with other additional datasets from our group and others deposited in PRIDE [39].

**Figure 1.**
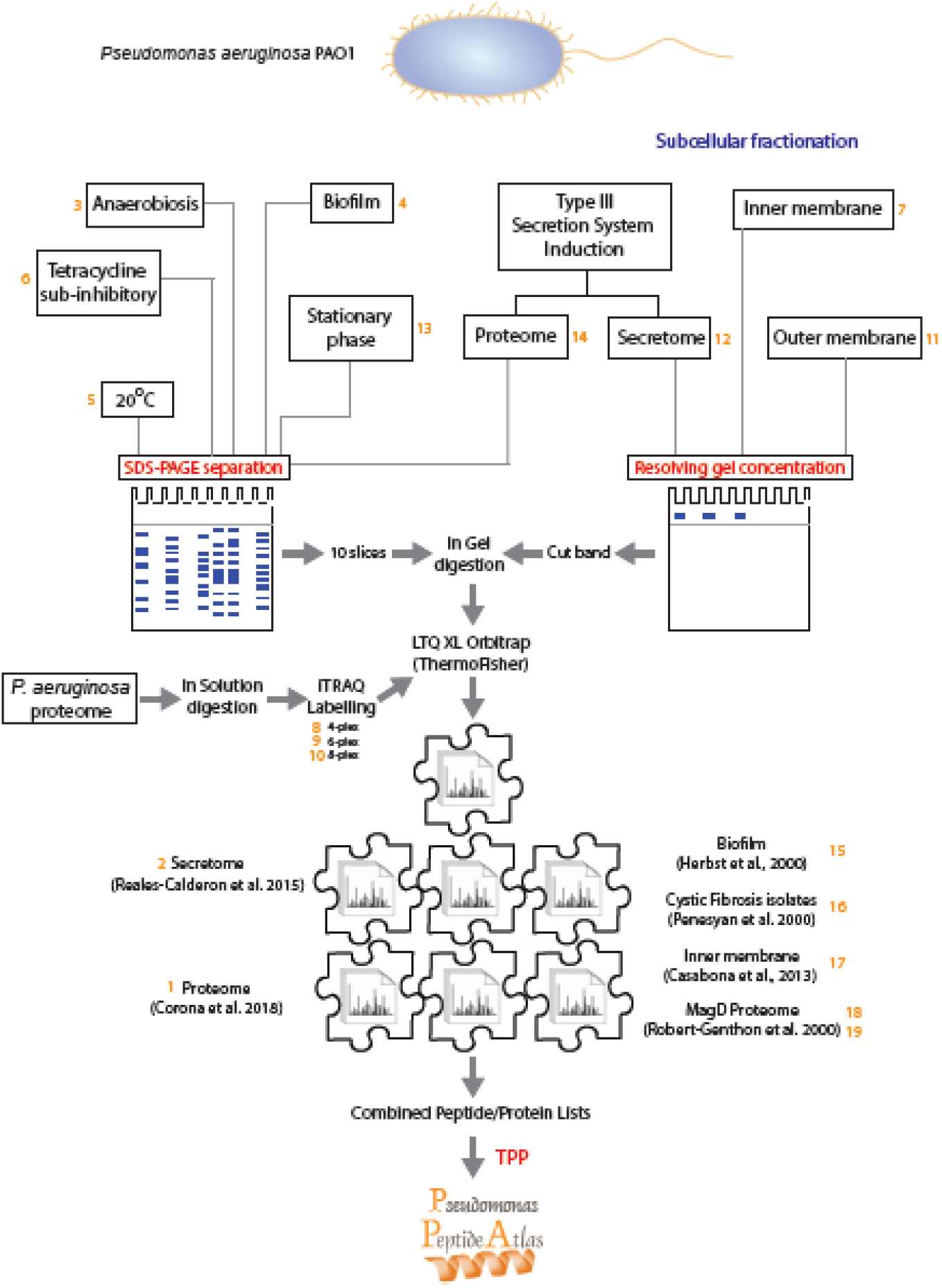
Diagram showing the different samples obtained and analyzed to built-in the *Pseudomonas aeruginosa* PeptideAtlas. The different samples are identified by the number that PeptideAtlas has assigned (Table 1). Cytosolic, sub-cellular and secreted protein fractions were obtained for *P. aeruginosa* PAO1 grown under different conditions. Proteins were exhaustively separated on SDS-PAGE and peptides by OFFGEL separation. The MS output results, together with additional MS datasets from other authors were all processed with the TPP tool suite.

**TABLE 1.** *Pseudomonas aeruginosa* proteomic samples included in this work.

| Sample number | Database name | Proteomic analysis | Runs | Number of identified proteins | Unique proteins | Pubmed ID | Repositories* |
| --- | --- | --- | --- | --- | --- | --- | --- |
| 1 | pao1_fcp001_proteome_4plex | iTRAQ/ LC-MS/MS | 30 | 2762 | 58 | 30429516 | PASS00809 |
| 2 | pao1_vescl_secretome_4plex | iTRAQ/ LC-MS/MS | 12 | 1265 | 13 | 26102536 | PASS00808 |
| 3 | pao1_anaerobiosis | LC-MS/MS | 5 | 2252 | 22 |  | PASS00804 |
| 4 | pao1_biofilm | LC-MS/MS | 5 | 1689 | 15 |  | PASS00803 |
| 5 | pao1_growth_20c | LC-MS/MS | 5 | 2313 | 12 |  | PASS00800 |
| 6 | pao1_growth_TCN | LC-MS/MS | 5 | 2465 | 18 |  | PASS00801 |
| 7 | pao1_inner_memb | LC-MS/MS | 5 | 1158 | 3 |  | PASS00805 |
| 8 | pao1_itraq_4plex | iTRAQ/ LC-MS/MS | 2 | 1568 | 3 |  | PASS00821 |
| 9 | pao1_itraq_6plex | iTRAQ/ LC-MS/MS | 2 | 1582 | 3 |  | PASS00822 |
| 10 | pao1_itraq_8plex | iTRAQ/ LC-MS/MS | 2 | 1413 | 2 |  | PASS00823 |
| 11 | pao1_outer_memb | LC-MS/MS | 1 | 1080 | 4 |  | PASS00806 |
| 12 | pao1_secretome_T3SS | LC-MS/MS | 1 | 473 | 6 |  | PASS00807 |
| 13 | pao1_stat_phase | LC-MS/MS | 5 | 2504 | 20 |  | PASS00799 |
| 14 | pao1_T3SS_induction | LC-MS/MS | 1 | 2486 | 12 |  | PASS00802 |
| 15 | PAO1_Fap | LC-MS/MS | 288 | 3698 | 0 | 25317949 | PXD001373 |
| 16 | Pseudomonas_Cystic_Fibrosis | DIA LC-MS/MS | 80 | 1787 | 223 | 26431321 | PXD002865 |
| 17 | PAO1_Inner_Membrane_Proteome | LC-MS/MS | 6 | 1656 | 30 | 23744604 | PXD000107 |
| 18 | PAO1_MagD-WT | LC-MS/MS | 2 | 1102 | 5 | 23919994 | PXD000189 |
| 19 | PAO1_delta-MagD | LC-MS/MS | 2 | 446 | 0 | 23919994 | PXD000189 |

\* Datasets available in PeptideAtlas Database (accession code PASS\_) and PRIDE Database (accession code PXD\_).

For the samples prepared for the present work, PAO1 was grown until exponential phase of growth (OD_600_ of 0.6) in LB Luria Bertani (LB) broth (10 g Peptone, 5 g Yeast Extract, 5 g Sodium Chloride) at 37°C with the following exceptions: sample 4: biofilm formed in p24 plates at 37°C during 48h; sample 5: at 20°C; sample 6: in LB agar; and sample 13: until OD_600_ of 4 to reach stationary phase of growth. Additives were used for the following samples: samples 12 and 14: 5 mM EGTA and 20 mM MgCl_2_ for stimulating Type III secretion [40] during 4h; and sample 6: Tetracycline sub-inhibitory concentration (0.5 mg/l) during 4h.

All the samples were grown under aerobic conditions except for sample 3, where anaerobiosis was achieved in LB agar plates placed in a generator for GENbox jar 2.5 l anaerobic environment (GENbox anaer, Biomerieux) during 48h at 37°C.

All the samples were total extracts (cytosolic mainly) except for samples 7 and 11, where subcellular fractionations were needed to purify inner and outer membranes, and the secretome obtained for sample 12.

### 2.2. Protein extraction and digestion

Pellets from *P. aeruginosa* PAO1 cells under the different conditions were washed and resuspended in 1 ml of PBS supplemented with 1/1000 Protease Inhibitor Cocktail (Complete, Mini, EDTA-free Protease Inhibitor Cocktail Tablets). PAO1 biofilm was washed with PBS and cells were collected to protein extraction. Cells from the different samples were lysed by sonication (Labsonic U) for 3 times, 1 min each, and centrifuged for 20 min at 6000g at 4°C. Cell debris was removed by centrifugation (14000*g*, 10 min, 4°C). Proteins were frozen at −80°C until used. Protein concentration was measured using Bradford assay (Biorad).

The secretome from the T3SS-induction sample was collected by centrifugation at 7000*g*, 10 min, 4°C and supernatants were filtered through a 0.2 μm pore size Nalgene Disposable Filter Unit (Thermo Fisher Scientific) to remove the remaining cells. Proteins were precipitated with methanol/chloroform. The pellet was washed 2 times with PBS and resuspended in 0.5 M Triethylammonium bicarbonate (TEAB) supplemented with 1/1000 Protease Inhibitor Cocktail.

In the case of the inner and outer membranes enriched fractions, cells were pelleted and washed with PBS and resuspended in 1.5 ml of 20 mM HEPES pH 8.0. Then, lysis was reached by sonication for 3 times, 1 min each, and centrifuged for 20 min at 6000g at 4°C. After this, supernatant was collected and ultracentrifuged at 100000g during 1h at 4°C. Then, supernatant was collected and pellet was resuspended in 1.5 ml 20 mM HEPES pH 8.0 and the suspension was ultracentrifuged at 100000 g during 30 at 4°C. Total membranes were pelleted and resuspended in 1.5 ml of 20 mM HEPES pH 8.0 + 15 μl TRITON X-100, ultracentrifuged at 100000g during 1h at 4°C. Outer membranes were located in the pellet and inner membranes were enriched in the supernatant. Inner membranes were collected. The pellet was washed in 1.5 ml of 20 mM HEPES pH 8.0, ultracentrifuged at 100000g during 1h at 4°C and pellet, enriched in outer membranes, was resuspended in 300 μl of 20 mM HEPES pH 8.0.

One hundred μg of the different PAO1 cytoplasmic extracts were loaded onto conventional SDS-PAGE 10% Bis-Tris gels (mini-protean TGX Stain-free precast Gels, BioRad). The gel was stained with Coomassie blue and each lane was cut into 10 bands. Gel slices were cut into 1 mm^3^ cubes, washed twice with water and dehydrated with 100% ACN (v/v).

One hundred μg of protein of subproteomes (outer and inner membranes and T3SS secretome) were loaded in a conventional SDS-PAGE gel (1mm-thick, 4% stacking, and 12% resolving). Then run was stopped as soon as the front entered into the resolving gel, so that the whole proteome became concentrated in the stacking/resolving gel interface. The unseparated protein bands were visualized by Coomassie staining, excised, cut into cubes (1mm^3^), washed twice with water and dehydrated with 100% ACN (v/v). All the gel slices were incubated with 10 mM DTT in 50 mM NH_4_HCO_3_ for 30 min at 56°C for protein reduction; alkylation was carried out with 50 mM IAA in 50 mM ammonium bicarbonate solution.

The gel pieces were washed with 50% ACN (v/v), and then washed again with 10 mM NH_4_HCO_3_, dehydrated with 100% ACN (v/v), and then dried in a vacuum concentrator. The gel pieces were rehydrated by adding sequence grade-modified trypsin (Roche) 1:20 in 50mM NH_4_HCO_3_ and incubated overnight at room temperature in the dark for protein digestion. Supernatants were transferred to clean tubes, and gel pieces were incubated in 50mM NH_4_HCO_3_ at 50 °C for 1 h. Then, remaining peptides were collected by incubation with 5% formic acid for 15 min and with 100% ACN for 15 min more. The extracts were combined, and the organic solvent was removed in a vacuum concentrator.

The tryptic eluted peptides were dried by speed-vacuum centrifugation and then desalted onto StageTip C18 Pipette tips (Thermo Scientific) until the mass spectrometric analysis. The 10 slices of the whole proteomes were combined and 5 fractions were used for the mass spectrometry analysis.

### 2.3. LC-MS/MS analysis

The MS system used was an LTQ XL Orbitrap (ThermoFisher) equipped with a nanoESI ion source. A total amount of 5 μg from each sample (volume of 20 μl) was loaded into the chromatographic system consisting in a C18 pre-concentration cartridge (Agilent Technologies) connected to a 360 cm long, 100 μm i.d. C18 column (NanoSeparations). The separation was done at 0.25 μl/min in a 360-min acetonitrile gradient from 3 to 40% (solvent A: 0.1% formic acid, solvent B: acetonitrile 0.1% formic acid). The HPLC system was composed of an Agilent 1200 capillary nano pump, a binary pump, a thermostated micro injector and a micro switch valve. The LTQ XL Orbitrap was operated in the positive ion mode with a spray voltage of 1.8 kV. The spectrometric analysis was performed in a data dependent mode, acquiring a full scan followed by 10 MS/MS scans of the 10 most intense signals detected in the MS scan from the global list. The full MS (range 400-1800) was acquired in the Orbitrap with a resolution of 60.000. The MS/MS spectra were done in the linear ion-trap.

### 2.4. Compiling of additional *P. aeruginosa* datasets

#### 2.4.1. Datasets from our group

In addition to the current proteomes analyzed, three different *P. aeruginosa* MS/MS datasets from our lab were compiled in order to contribute to the *P. aeruginosa* PeptideAtlas, extending the coverage in two more specific conditions, the secretome of PAO1 compared with the mutant FCP001 (EVs and EV-free secre tome) [24], and the comparison of the whole proteomes of PAO1 and mutant strain FCP001 grown until under exponential phase of growth [41].

A comparison of the labelling by iTRAQ 4-plex, 6-plex (6 of the 8-plex kit markers) and 8-plex (using 4, 6 or 8 biological replicates from the whole proteome of PAO1 until exponential phase growth). Briefly, 8 independent cultures of PAO001 (20 ml of each) were grown in LB medium to reach mid-exponential growth phase (OD_600_ of 0.6), moment at which the cells were pelleted by centrifugation and proteins were obtained in 1 ml of PBS supplemented with 1/1000 Protease Inhibitor Cocktail (Complete, Mini, EDTA-free Protease Inhibitor Cocktail Tablets) by sonication. Forty μg of protein from each condition was precipitated with methanol/chloroform and resuspended in 0.5 M Triethylammonium bicarbonate (TEAB) and after trypsin digestion, samples were labeled at room temperature for 2 h with a half unit of iTRAQ Reagent Multi-plex kit (AB SCIEX, Foster City, CA, USA). Finally, samples were combined and bRP-LC C18 fractionation of the iTRAQ labelled peptides was performed on the SmartLine (Knauer, Germany) HPLC system using the Waters, XBridge C18 column (100 × 2.1 mm, 5 μm particle). Thirty fractions were collected over the elution profile and pooled into 5 fractions. The peptide fractions were dried, desalted using a SEP-PAK C18 Cartridge (Waters) and stored at −20 °C until the LC−MS analysis.

A 1.5 μg aliquot of each peptide fraction was subjected to 2D-nano LC ESI-MSMS analysis using a nano liquid chromatography system (Eksigent Technologies nanoLC Ultra 1D plus, AB SCIEX, Foster City, CA) coupled to high speed Triple TOF 5600 mass spectrometer (AB SCIEX, Foster City, CA) with a Nanospray III Source. The loading pump delivered a solution of 0.1% formic acid in water at 2 μl/min. The nano-pump provided a flow-rate of 300 nl/min and was operated under gradient elution conditions, using 0.1% formic acid in water as mobile phase A, and 0.1% formic acid in acetonitrile as mobile phase B. Gradient elution was performed using a 120 minutes gradient ranging from 2% to 90% mobile phase B. Injection volume was 5 μl.

Data acquisition was performed with a TripleTOF 5600 System (AB SCIEX, Concord, ON). Data was acquired using an ionspray voltage floating (ISVF) 2800 V, curtain gas (CUR) 20, interface heater temperature (IHT) 150, ion source gas 1 (GS1) 20, declustering potential (DP) 85 V. All data was acquired using information-dependent acquisition (IDA) mode with Analyst TF 1.5 software (AB SCIEX, USA). For IDA parameters, 0.25 s MS survey scan in the mass range of 350–1250 Da were followed by 30 MS/MS scans of 150ms in the mass range of 100–1800 (total cycle time: 4.04 s). Switching criteria were set to ions greater than mass to charge ratio (m/z) 350 and smaller than m/z 1250 with charge state of 2–5 and an abundance threshold of more than 90 counts (cps). Former target ions were excluded for 20 s. IDA rolling collision energy (CE) parameters script was used for automatically controlling the CE.

#### 2.4.2. Other PRIDE *P. aeruginosa* datasets

Together with the experiments from our group, other published works that have been uploaded to the public repository PRIDE [42] have been included in order to increase the coverage of the *P. aeruginosa* PeptideAtlas. Included are datasets from Herbst *et al*. (PXD001373)[43]; Penesyan *et al*. (PXD002865)[44]; Casabona *et al*. (PXD000107)[45] and Robert-Genthon *et al.* (PXD000189)[46] (Table 1, Figure 1).

### 2.5. Post spectra acquisition processing

LC–MS/MS spectra files resulting from the different proteomes (Figure 1), in their native vendor-specific format, along with the meta data corresponding to each approach, were submitted to PeptideAtlas via the PeptideAtlas Submission System (PASS) on-line submission form with dataset identifications: PASS00799, PASS00800, PASS00801, PASS00802, PASS00803, PASS00804, PASS00805, PASS00806, PASS00807, PASS00808, PASS00809, PASS00821 and PASS00822, together with the already included in PRIDE previously cited. LC–MS/MS spectra files were converted to XML-based HUPO-PSI-adopted standard format for mass spectrometry output, mzML [47]. The protein sequence FASTA file was obtained from the *Pseudomonas* Genome Database (www.pseudomonas.com) [35]. Sequences were appended with a set of common contaminant proteins from the cRAP (common Repository of Adventitious Proteins) set from the GPM (http://www.thegpm.org/crap/) and decoy counterparts for every entry to add up a total of 11376 entries.

Then database searches were performed using three different search engines: Comet [48], an open-source, freely available version of SEQUEST [49], X!Tandem [50] with the k-score algorithm plugin [51], and OMSSA [52]. The search parameters were established depending on the type of experiment and instrument (see supplementary table S1 ‘database_search_parameters’ for a list of parameters).

Following sequence database searching, we used the TPP (Trans-Proteomic Pipeline) tool suite [53, 54] to validate the results. First, PeptideProphet [55] creates a discriminant search engine-independent score, models distributions of correctly and incorrectly assigned peptide spectrum matches (PSMs) and computes PSM posterior probabilities. Next, iProphet [56], was used to combine three search results, further refine the PSM-level probabilities and calculate distinct peptide-level probabilities using corroborating information from other PSMs in the dataset. ProteinProphet [57] then was used to further refine peptide probabilities based on the Number of Sibling Peptides (NSP) that each peptide shares within a protein; it also groups and reports proteins with a protein-level probability estimated from peptide-level probabilities. To assemble the *P. aeruginosa* PeptideAtlas, all individual iProphet files from the 13 compiled datasets were filtered at a variable probability threshold to reach a constant PSM-level FDR threshold of 0.0002 across all datasets.

The *Pseudomonas aeruginosa* PeptideAtlas is made available at https://db.systemsbiology.net/sbeams/cgi/PeptideAtlas/buildDetails?atlas_build_id=455.

Mayu [58] software, designed for large-scale protein FDR estimation, was used to report FDR values at different levels (PSM, unique peptides and protein-level) for the whole build based on a strategy that estimates the number of false positive protein identifications from the number of proteins containing false positive PSMs, including a correction for high proteome coverage.

### 2.5 In silico Functional Analyses

Gene Ontology classifications were performed using the *Pseudomonas* Genome Database (www.pseudomonas.com) [35] and KEGG PATHWAY Database (https://www.genome.jp/kegg/) was used for pathway analysis [59, 60].

## 3. Results and Discussion

### 3.1. Construction of the *P. aeruginosa* PeptideAtlas and strategies for exhaustive proteome characterization

The *Pseudomonas* PeptideAtlas has been built using data mainly obtained from the wild-type strain PAO1, one of the most widely used model strains, whose genome sequence is available [61]. We used a wide range of experimental growth conditions and different subcellular fractions to increase the proteome coverage as much as possible (Figure 1 and Table 1). Table 1 and Supplemental Table S2 show the number of proteins identified in the different samples. In Table S2, proteins are ordered by number in the *Pseudomonas* Genome Database (www.pseudomonas.com) [35], facilitating the visualization of the identified proteins in the different conditions in order to observe commonalities and differences between conditions. Uniquely identified proteins in each dataset are highlighted. Of the total 648 uniquely identified proteins, 164 belong to the datasets generated by our group. This shows the necessity of analyzing of proteomes produced under different environmental conditions.

Samples 1 and 2 were obtained from PAO1 and the mutant strain Fcp001 *(Δcrc*) grown until they reached the exponential phase of growth at 37°C. In the work by Corona *et al.* [41], the whole proteomes were studied (Sample 1), and in the case of Reales-Calderon *et al.* [24], the secretome (Sample 2). In both, proteins were labeled with iTRAQ and analyzed by LC-MS/MS. In sample 1, 2762 cytosolic proteins were identified in a total of 30 runs, and in sample 2, a total of 1265 proteins in the secretome and EVs were identified in 12 runs. The same growth conditions were used for samples 8, 9, and 10, allowing the identification of around 1500 proteins in each experiment. The difference between these samples was the iTRAQ labeling strategy, with 4, 6, or 8 plex used. Only two runs were done for each sample. As expected, the three samples differed in only a few proteins (Table S2). All these samples were grown until the exponential phase of growth, a condition widely used for microbiological studies, with the expectation that many proteins should be abundant because the microorganism is actively growing. However, the stationary phase of growth can unveil proteins that are important for the maintenance of the microorganism once the carbon source has been exhausted and secreted components are accumulating in the culture. This condition is represented by sample 13, which comprises 2504 proteins, including 18 not detected in the other samples.

For sample 3 anaerobic conditions were used because, during chronic infection of the CF lung, *P. aeruginosa* is able to grow and persist in a microaerobic to anaerobic environment [62]. This analysis rendered 2252 proteins, 20 of them unique to this sample. Growth as a biofilm is very important in *P. aeruginosa* virulence and antibiotic resistance [14], in its persistence in environments such as water supplies [63], and in CF patients [3]. Samples 4, 15, and 16 represent this kind of growth. In sample 4, 1689 proteins were identified, including 15 uniquely identified in this sample.

As *P. aeruginosa* is present in natural environments, we grew PAO1 at 20°C (sample 5). In 5 runs, 2313 proteins were identified, of which only 6 were unique to this condition. Because *P. aeruginosa* is quite resistant to antibiotics [14, 20] and it has been reported that subinhibitory antibiotic concentrations may modify *P. aeruginosa* physiology [64], sample 6, grown in tetracycline (TCN) was included, with 2465 proteins identified, including 13 unique to this condition. In addition, PAO1 was cultured in the medium that induces the type III secretion system (T3SS), as T3SS is a very important virulence factor, and the proteome and secretome were analyzed (samples 14 and 12, respectively). This condition yielded a total of more than 2800 identified proteins from the two samples, including 15 that were unique to them.

Cytosolic extracts are the most commonly studied fractions in proteomic analysis, but some proteins are under-represented in or missing from them, for instance membrane proteins [65]. With the goal of maximum coverage of the proteome, subcellular fractions enriched in membrane proteins (samples 7 and 11) were also analyzed, and 7 more proteins were identified.

Besides all the proteomic samples described, proteins identified in published works [24, 41, 43–46] were also included. This added 547 uniquely identified proteins to the PeptideAtlas. These protein databases included proteomes from samples enriched in subcellular fractions and also from strains different from PAO1 [44]. Sample 16, from Penesyan *et al.*, includes proteins from four clinical isolates. As could be expected, the proteomes of these freshly isolated strains differ from that of the PAO1 strain, as only 29.4% of the identified proteins were common to all 5 strains. Although this does not mean that the proteins are not present in all these strains, it is a sign of differences in the most abundant proteins between samples showing, as the authors highlight, the need for analyses of strains isolated from clinical or environmental samples [44]. Thus, sample 16 contributed 1787 proteins, of which 223 were uniquely identified. The dataset with the most proteins identified is sample 15 [43] with 3698, and it also comprises the largest number of unique proteins, 239.

A total of 5,078,535 tandem mass (MS/MS) spectra from the 19 different proteomic experiments and 459 individual runs using a wide variety of mass spectrometers (TripleTOF 5600, LTQ Orbitrap XL, LTQ-Orbitrap Velos, LTQ Orbitrap Velos pro and Q Exactive) were used to generate the *P. aeruginosa* PeptideAtlas. Almost 40% (2,021,521) of the generated spectra were matched to peptides with probability high enough to pass our threshold. From these, 13,059 peptides were identified in only one dataset. The most important big dataset was from the Fap PAO1 sample [43], with the highest number of runs (288). It contained 2,516,514 spectra, allowing the identification of 3698 proteins. The proteomic characterization of sample 1, a 4-plex iTRAQ, also yielded a high number of proteins, 2762. Overviews of the contributions of each dataset to the entirety of the PeptideAtlas build are depicted in Table 1 and in Figure 2.

**Fig. 2.**
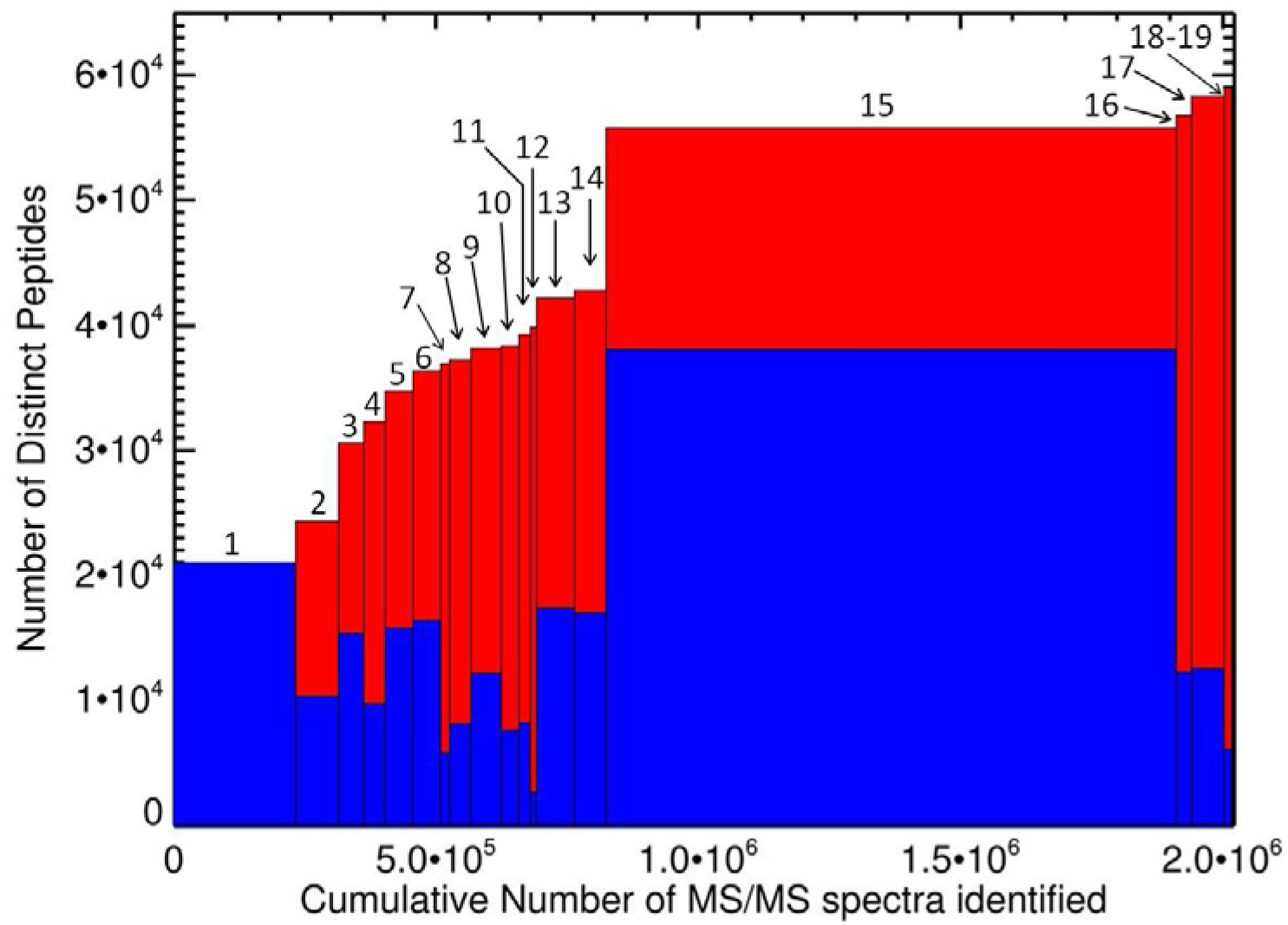
Contribution of the different constituent datasets of the *Pseudomonas* PeptideAtlas. Plot shows the number of peptides contributed by each experiment, and the cumulative number of distinct peptides for the build as of that experiment. For each dataset, the height of the blue bar represents the number of distinct peptides and the total height of bar represents the cumulative number of distinct peptides identified. The width of bars represents the contributions in terms of spectra identified. Numbers designate the samples as ordered in Table 1.

### 3.2. Functional and Gene ontology enrichment analyses of the proteins included in the *P. aeruginosa* PeptideAtlas

The proteomic analyses identified 4043 proteins from a total of 5668 proteins predicted to be encoded by the *P. aeruginosa* PAO1 genome, achieving 71.3% coverage of the predicted *Pseudomonas* proteome. Table 2 lists the more highly represented KEGG pathways, and those with coverage superior to 90% are highlighted. The *Pseudomonas* PeptideAtlas includes 84% of metabolic proteins, 71% of the proteins involved in genetic information processing, 72% of the proteins responsible for environmental information processing, more than 80% of the proteins related to quorum sensing and biofilm formation, and 81% of the proteins responsible for antimicrobial resistance.

**TABLE 2.** KEGG pathway enrichment analysis.

| KEGG identifiers and KEGG pathway maps | Identified proteins | Total protein in KEGG pathway maps | Percent of protein in Peptide Atlas* |
| --- | --- | --- | --- |
| **1. Metabolism** |  |  |  |
| **Global and overview maps** |  |  |  |
| 01200 Carbon metabolism | 117 | 129 | 90.70 |
| 01210 2-Oxocarboxylic acid metabolism | 26 | 30 | 86.67 |
| 01212 Fatty acid metabolism | 37 | 48 | 77.08 |
| 01230 Biosynthesis of amino acids | 131 | 145 | 90.34 |
| 01220 Degradation of aromatic compounds | 10 | 27 | 37.04 |
| **Carbohydrate metabolism** |  |  |  |
| 00010 Glycolysis / Gluconeogenesis | 32 | 35 | 91.43 |
| 00020 Citrate cycle (TCA cycle) | 30 | 31 | 96.77 |
| 00030 Pentose phosphate pathway | 24 | 27 | 88.89 |
| 00040 Pentose and glucuronate interconversions | 3 | 7 | 42.86 |
| 00051 Fructose and mannose metabolism | 12 | 19 | 63.16 |
| 00052 Galactose metabolism | 4 | 5 | 80.00 |
| 00053 Ascorbate and aldarate metabolism | 3 | 6 | 50.00 |
| 00500 Starch and sucrose metabolism | 12 | 16 | 75.00 |
| 00520 Amino sugar and nucleotide sugar metabolism | 33 | 37 | 89.19 |
| 00620 Pyruvate metabolism | 56 | 61 | 91.80 |
| 00630 Glyoxylate and dicarboxylate metabolism | 51 | 59 | 86.44 |
| 00640 Propanoate metabolism | 41 | 47 | 87.23 |
| 00650 Butanoate metabolism | 33 | 41 | 80.49 |
| 00660 C5-Branched dibasic acid metabolism | 9 | 14 | 64.29 |
| 00562 Inositol phosphate metabolism | 6 | 7 | 85.71 |
| **Energy metabolism** |  |  |  |
| 00190 Oxidative phosphorylation | 48 | 57 | 84.21 |
| 00680 Methane metabolism | 24 | 26 | 92.31 |
| 00910 Nitrogen metabolism | 27 | 35 | 77.14 |
| 00920 Sulfur metabolism | 31 | 49 | 63.27 |
| **Lipid metabolism** |  |  |  |
| 00061 Fatty acid biosynthesis | 23 | 26 | 88.46 |
| 00071 Fatty acid degradation | 22 | 32 | 68.75 |
| 00072 Synthesis and degradation of ketone bodies | 9 | 10 | 90.00 |
| 00561 Glycerolipid metabolism | 11 | 14 | 78.57 |
| 00564 Glycerophospholipid metabolism | 19 | 26 | 73.08 |
| 00565 Ether lipid metabolism | 2 | 3 | 66.67 |
| 00590 Arachidonic acid metabolism | 3 | 4 | 75.00 |
| 00592 alpha-Linolenic acid metabolism | 1 | 3 | 33.33 |
| 01040 Biosynthesis of unsaturated fatty acids | 3 | 3 | 100.00 |
| **Nucleotide metabolism** |  |  |  |
| 00230 Purine metabolism | 60 | 68 | 88.24 |
| 00240 Pyrimidine metabolism | 27 | 33 | 81.82 |
| **Amino acid metabolism** |  |  |  |
| 00250 Alanine, aspartate and glutamate metabolism | 34 | 37 | 91.89 |
| 00260 Glycine, serine and threonine metabolism | 45 | 49 | 91.84 |
| 00270 Cysteine and methionine metabolism | 37 | 41 | 90.24 |
| 00280 Valine, leucine and isoleucine degradation | 39 | 47 | 82.98 |
| 00290 Valine, leucine and isoleucine biosynthesis | 17 | 19 | 89.47 |
| 00300 Lysine biosynthesis | 14 | 15 | 93.33 |
| 00310 Lysine degradation | 19 | 21 | 90.48 |
| 00220 Arginine biosynthesis | 27 | 29 | 93.10 |
| 00330 Arginine and proline metabolism | 36 | 47 | 76.60 |
| 00340 Histidine metabolism | 22 | 24 | 91.67 |
| 00350 Tyrosine metabolism | 13 | 27 | 48.15 |
| 00360 Phenylalanine metabolism | 20 | 26 | 76.92 |
| 00380 Tryptophan metabolism | 28 | 30 | 93.33 |
| 00400 Phenylalanine, tyrosine and tryptophan biosynthesis | 26 | 30 | 86.67 |
| **Metabolism of other amino acids** |  |  |  |
| 00410 beta-Alanine metabolism | 18 | 26 | 69.23 |
| 00430 Taurine and hypotaurine metabolism | 6 | 8 | 75.00 |
| 00440 Phosphonate and phosphinate metabolism | 2 | 10 | 20.00 |
| 00450 Selenocompound metabolism | 8 | 9 | 88.89 |
| 00460 Cyanoamino acid metabolism | 10 | 11 | 90.91 |
| 00471 D-Glutamine and D-glutamate metabolism | 5 | 5 | 100.00 |
| 00472 D-Arginine and D-ornithine metabolism | 2 | 2 | 100.00 |
| 00473 D-Alanine metabolism | 4 | 4 | 100.00 |
| 00480 Glutathione metabolism | 27 | 29 | 93.10 |
| **Glycan biosynthesis and metabolism** |  |  |  |
| 00540 Lipopolysaccharide biosynthesis | 28 | 29 | 96.55 |
| 00550 Peptidoglycan biosynthesis | 19 | 20 | 95.00 |
| **Metabolism of cofactors and vitamins** |  |  |  |
| 00730 Thiamine metabolism | 12 | 13 | 92.31 |
| 00740 Riboflavin metabolism | 9 | 11 | 81.82 |
| 00750 Vitamin B6 metabolism | 8 | 9 | 88.89 |
| 00760 Nicotinate and nicotinamide metabolism | 21 | 22 | 95.45 |
| 00770 Pantothenate and CoA biosynthesis | 16 | 23 | 69.57 |
| 00780 Biotin metabolism | 18 | 21 | 85.71 |
| 00785 Lipoic acid metabolism | 2 | 3 | 66.67 |
| 00790 Folate biosynthesis | 29 | 32 | 90.63 |
| 00670 One carbon pool by folate | 17 | 18 | 94.44 |
| 00860 Porphyrin and chlorophyll metabolism | 37 | 45 | 82.22 |
| 00130 Ubiquinone and other terpenoid-quinone biosynthesis | 16 | 19 | 84.21 |
| **Metabolism of terpenoids and polyketides** |  |  |  |
| 00900 Terpenoid backbone biosynthesis | 15 | 16 | 93.75 |
| 00903 Limonene and pinene degradation | 7 | 7 | 100.00 |
| 00281 Geraniol degradation | 12 | 16 | 75.00 |
| 00523 Polyketide sugar unit biosynthesis | 5 | 5 | 100.00 |
| 01053 Biosynthesis of siderophore group nonribosomal peptides | 7 | 7 | 100.00 |
| **Biosynthesis of other secondary metabolites** |  |  |  |
| 00332 Carbapenem biosynthesis | 2 | 2 | 100.00 |
| 00261 Monobactam biosynthesis | 8 | 9 | 88.89 |
| 00521 Streptomycin biosynthesis | 8 | 8 | 100.00 |
| 00525 Acarbose and validamycin biosynthesis | 2 | 2 | 100.00 |
| 00401 Novobiocin biosynthesis | 4 | 5 | 80.00 |
| 00405 Phenazine biosynthesis | 24 | 24 | 100.00 |
| **Xenobiotics biodegradation and metabolism** |  |  |  |
| 00362 Benzoate degradation | 21 | 35 | 60.00 |
| 00625 Chloroalkane and chloroalkene degradation | 6 | 8 | 75.00 |
| 00361 Chlorocyclohexane and chlorobenzene degradation | 1 | 6 | 16.67 |
| 00633 Nitrotoluene degradation | 1 | 2 | 50.00 |
| 00643 Styrene degradation | 8 | 12 | 66.67 |
| 00930 Caprolactam degradation | 6 | 8 | 75.00 |
| 00626 Naphthalene degradation | 2 | 2 | 100.00 |
| **2. Genetic Information Processing** |  |  |  |
| **Transcription** |  |  |  |
| 03020 RNA polymerase | 4 | 4 | 100.00 |
| **Translation** |  |  |  |
| 03010 Ribosome | 54 | 68 | 79.41 |
| 00970 Aminoacyl-tRNA biosynthesis | 27 | 90 | 30.00 |
| **Folding, sorting and degradation** |  |  |  |
| 03060 Protein export | 16 | 18 | 88.89 |
| 04122 Sulfur relay system | 18 | 19 | 94.74 |
| 03018 RNA degradation | 16 | 17 | 94.12 |
| **Replication and repair** |  |  |  |
| 03030 DNA replication | 14 | 15 | 93.33 |
| 03410 Base excision repair | 12 | 13 | 92.31 |
| 03420 Nucleotide excision repair | 7 | 7 | 100.00 |
| 03430 Mismatch repair | 16 | 17 | 94.12 |
| 03440 Homologous recombination | 22 | 23 | 95.65 |
| **3. Environmental Information Processing** |  |  |  |
| **Membrane transport** |  |  |  |
| 02010 ABC transporters | 121 | 189 | 64.02 |
| 02060 Phosphotransferase system (PTS) | 5 | 5 | 100.00 |
| 03070 Bacterial secretion system | 60 | 92 | 65.22 |
| **Signal transduction** |  |  |  |
| 02020 Two-component system | 166 | 201 | 82.59 |
| **4. Cellular Processes** |  |  |  |
| **Cellular community - prokaryotes** |  |  |  |
| 02024 Quorum sensing | 79 | 94 | 84.04 |
| 02025 Biofilm formation - Pseudomonas aeruginosa | 92 | 113 | 81.42 |
| **Cell motility** |  |  |  |
| 02030 Bacterial chemotaxis | 45 | 45 | 100.00 |
| 02040 Flagellar assembly | 28 | 37 | 75.68 |
| **5. Human Diseases** |  |  |  |
| **Drug resistance: Antimicrobial** |  |  |  |
| 01501 beta-Lactam resistance | 21 | 27 | 77.78 |
| 01502 Vancomycin resistance | 7 | 8 | 87.50 |
| 01503 Cationic antimicrobial peptide (CAMP) resistance | 23 | 28 | 82.14 |

\*Functions highlighted are the ones represented in a percentage higher than 90% in *Pseudomonas* PeptideAtlas.

The contribution of each sample and the role of some of the proteins included in the PeptideAtlas in processes important for *P. aeruginosa* virulence or adaptation to the conditions studied are discussed below.

#### 3.2.1. Proteins relevant to biofilm formation and maintenance

As previously mentioned, samples 4, 15, and 16 were grown as biofilms. When comparing the proteins identified in strains from CF patients and the proteome of PAO1 biofilm (sample 4), we found 1154 proteins common to both proteomes. Of them, 23 are related to biofilm formation, 33 to the quorum-sensing response, and 19 to antibiotic resistance. In sample 4, of the 15 unique proteins, we found ErcS (PA1976), ErbR (PA1978), EraR (PA1980), and PqqC (PA1987), proteins involved in ethanol oxidation, to be highly expressed in biofilm cells and probably involved in the ciprofloxacin resistance of biofilms [66]. TspR (PA4857), required for T3SS expression, biofilm formation, and bacterial motility [67, 68], was also found.

In sample 15, in which functional amyloid was induced, 4 of the amyloids, FapE (PA1952), FapD (PA1953), FapC (PA1954), and FapB (PA1955) were uniquely identified [43]. It is also noteworthy, but not surprising, that the genes involved in alginate biosynthesis (PA3540 to PA 3551) were identified only in samples 15 and 16, as alginate is the main component of mucoid biofilms and of the *P. aeruginosa* biofilms encountered in CF patients [43, 69]. Another group of proteins involved in biofilm formation, the Pel extracellular polysaccharide biosynthetic proteins PelD (PA3061), PelC (PA3062), and PelB (PA3063), were identified only in sample 15. These genes are the back-up system for the extracellular polysaccharide biosynthetic process for single-species biofilm formation [70] when Psl polysaccharide biosynthesis fails. The genes for Psl biosynthesis are present in almost all the samples (PA2231– PA2245). We found other proteins related to biofilm formation only in sample 15. These included IcmF3 (PA2361); Arr (PA2818), which is essential for biofilm induction and aminoglycoside resistance [71]; and CupE3 (PA4650) and CupE5 (PA4652), part of the cupE gene cluster that encodes a chaperone usher pathway that is homologous to the *Acinetobacter baumanii* Csu pili gene cluster and involved in the early and late stages of biofilm formation [72].

CdrA is a cyclic-di-GMP-regulated adhesin that reinforces the extracellular matrix of *P. aeruginosa* biofilm [73]. In accord with its function, it was identified in both biofilm proteomes (samples 4 and 15) and is also common to both secretomes (samples 2 and 12), and is thus determined to be secreted. Mep72 is a metalloendopeptidase [74] common to secretomes and to samples 18 and 19, and it has been described as secreted in biofilms [75].

As previously mentioned, the study of the proteome under anaerobic conditions (sample 3), is also interesting because during chronic infection of the CF lung, *P. aeruginosa* is able to grow and persist in a microaerobic to anaerobic environment [62], thus suffering drastic physiological changes. Among the 20 unique proteins in this sample, we highlight PscU (PA1690), PopN (PA1698), ExsB (PA1712), and PscC (PA1716), related to T3SS, which is induced in these conditions [76]. AtvR (PA2899), identified only in sample 15, is an important regulator of the *Pseudomonas* adaptation to hypoxia and crucial for survival in the host [77].

#### 3.2.2. Proteins involved in secretion systems

Microorganisms often relate to their environments by secreting proteins, either by using classic secretion mechanisms, or through extracellular vesicles. Two secretomes with a large proportion of their proteins in common have been included in this study: a secretome obtained during exponential growth, with 12 unique proteins (sample 2), and a secretome sampled under the conditions for T3SS induction (sample 12), with 4 more. With respect to the cytoplasmic proteome under T3SS induction conditions (sample 14), proteins from glycine betaine catabolism (CdhR [PA5389], DgcA [PA5398], GbcB [PA5411], SoxD [PA5417], and SoxA [PA5418]) were detected only in this sample. The generation of glycine betaine via carnitine catabolism allows carnitine to induce the expression of the virulence factor hemolytic phospholipase C, PlcH [78], which induces a proinflammatory response, suppresses oxidative burst in neutrophils, degrades pulmonary surfactant, and increases endothelial cell death [79] during infection. Phospholipase C precursor (PlcH, PA1249) was detected as a unique protein in the secretome under T3SS induction conditions, which included other virulence factors, such as AprA (PA1249) [80] and the lipase LipA (PA2862) [81]. Thus, the analysis of the proteome under T3SS induction conditions has provided important contributions to the *P. aeruginosa* PeptideAtlas. Proteins involved in T3SS regulation, such as MexT (PA2492) [82], also involved in the regulation of the expression of the multidrug efflux pump MexEF-OprN [83] were identified only in sample 15.

Analysis of the inner and outer membranes of *P. aeruginosa* PAO1 yielded 25 more unique proteins in datasets 7, 11, and 17 (Table 1). Ten of them are related to transport through membranes or secretion systems. As an example, FleS (PA1098), identified in the outer-membrane proteome, is involved in the regulation of both mucin adhesion and motility [84]. The Tad proteins, involved in the Type II secretion system (PA2494–PA4303), have mostly been identified only in sample 15, and thus might be important for amyloid-induced biofilm growth.

The Type VI secretome and the proteins secreted are relevant to bacterial pathogenesis and cell survival in the host [85]. All of these proteins (PA0070–PA0091) were identified in the amyloid-induced biofilm (sample 15) and many of them in the rest of the samples, together with several VgrG proteins, including VgrG4a (PA3294), VgrG4b (PA3486), VgrG5 (PA5090), and VgrG6 (PA5266), that are secreted by this system [86].

#### 3.2.3. Proteins involved in bacterial response to antibiotics

The study of antibiotic resistance suggests new methods to fight it [87]. Besides, antibiotics can challenge bacterial physiology in some aspects going beyond resistance to these compounds [22].

Thus, the analysis of the proteome under a sub-inhibitory concentration of tetracycline was also included, and yielded 2465 proteins. Thirteen were unique, including GltR (PA3192), which is required for glucose transport and involved in the regulation of the expression of exotoxin A, a primary *P. aeruginosa* virulence factor [88]. Biofilms are more resistant to antibiotics, and proteins related to antibiotic resistance were uniquely identified in sample 15. These include PA5514, a probable beta lactamase; PoxB, regulated by AmpR, together with AmpC (PA4110) [89], identified in samples 15 and 16 and in the inner membrane proteome (sample 7); and CzcR (PA2523), which modulates antibiotic, Zn, Cd, and Co resistance [90].

#### 3.2.4. Proteins involved in iron acquisition

Iron is a key nutrient that is often in limiting concentrations [91], a situation of particular relevance in the case of infection because bacteria must compete with the host’s cells to acquire the iron required for growth. Consequently, siderophores and proteins involved in Fe transport are fundamental to bacterial survival.

FepB (PA4159), an iron-enterobactin transporter, and PfeA (PA2688), a ferric enterobactin receptor, were identified in the secretome of PAO1 (sample 2) [24]. As previously mentioned, PvdS (PA2426) has been identified in cells in stationary phase. PvdS is an RNA polymerase sigma-70 factor from the extracytoplasmic function sigma factor subfamily [92] that controls the synthesis of the siderophore pyoverdine. Its expression is regulated by FpvR (PA2388), a protein present in 6 datasets, and FpvA (PA2398), identified in PAO1 cytosolic extract (sample 9). An increased abundance of the pyoverdine synthesis machinery was observed in sample 15 [43]. In this sample there are several uniquely identified proteins related to iron, such as PvdR (PA2389) and PvdT (PA2390), which are involved in pyoverdine biosynthesis and secretion together with OmpQ (PA2391) [93]; FemA (PA1910), a ferric-mycobactin receptor; FvbA (PA4156), a receptor homologous to the one from *Vibrio cholerae* and able to use vibriobactin as a source of iron in the absence of siderophores [94]; FiuR (PA0471), involved in the uptake of ferrioxiamine B [95] with FiuI (PA0472), in sample 16; FvpG (PA2403) and the rest of the proteins of a putative ABC transporter complex (PA2404–PA2410); and FoxR (PA2467), part of the Fox cell-surface signaling (CSS) pathway employed by *P. aeruginosa* to sense and respond to the presence of ferrioxamine in the environment. FoxA (PA2466) is the receptor controlled by this system, and was identified only in samples 15 and 13 (a stationary phase sample) [96]. These results demonstrate the importance of iron capture and transport for *P. aeruginosa* biofilm growth.

Iron acquisition is also important for survival outside of hosts. The heme oxygenase Bph0 (PA4116), necessary to yield iron from heme [97], was identified in the sample grown at 20°C.

#### 3.2.5. Other important proteins

Another group of proteins important for *P. aeruginosa* virulence are the Alp proteins, which constitute a self-lysis pathway that enhances virulence in the host [98]. Only three Alp proteins were identified, two of them unique to samples 15 (AlpA, PA0907) and sample 2 (vesicle and exoproteome; AlpE [PA0911]).

## 4. Conclusions

This work describes the results of analyses of widely diverse *P. aeruginosa* proteomes. The datasets used to build the *P. aeruginosa* PeptideAtlas include the proteomes obtained from strain PAO1 grown under various conditions and from different subcellular fractions (cytosol, membranes, and secretomes). They were analyzed using different proteomic techniques of fractionation, labeling, and MS analysis. It also includes datasets from other authors, including a dataset comprising proteins from clinical *P. aeruginosa* strains. All this has allowed the widest coverage of *P. aeruginosa* PAO1 proteins and the highest published coverage by a bacterial PeptideAtlas, higher than those of *Leptospira interrogans* (65.9%) [99], *Halobacterium salinarum* (62.7%) [100], and *Streptococcus pyogenes* (55.7%) [101]. Only eight PeptideAtlas builds are from bacteria, including six derived from pathogens, highlighting the importance of the present work. It also a valuable resource for the selection of candidate proteotypic peptides for targeted proteomics experiments *via* selected reaction monitoring (SRM) or parallel reaction monitoring (PRM).

## 5. Acknowledgements

This study was supported by RTI2018-094004-B-100 from the Spanish Ministry of Science and Innovation, InGEMICS-CM B2017/BMD3691 from the Comunidad de Madrid, Spanish Network for Research in Infectious Diseases (REIPI RD16/0016/0011), and PRB3 (PT17/0019/0012) from the ISCIII. InGEMICSCM, REIPI, and PRB3 are co-financed by the European Development Regional Fund (ERDF) “A way to achieve Europe”. These results are lined up with the Human Infectious Diseases HPP initiative from the Human Proteome Project (HID-HPP). The proteomics analyses were performed in the Proteomics facilities of Centro Nacional de Biotecnología (CNB) and Complutense University of Madrid (UCM), both members of the ProteoRed-ISCIII network. This work was funded in part by the National Institutes of Health grants R01GM087221 (EWD/RLM) and R24GM127667(EWD), and by the National Science Foundation grant DBI-1933311 (EWD) and Award Number 1920268 (RLM).

## Legend to Figures and Tables

Table S1. Database search parameters.

Table S2. Proteins identified in each sample. Unique proteins in each sample are highlighted.

## References

[1] A. Oliver, X. Mulet, C. Lopez-Causape, C. Juan, The increasing threat of Pseudomonas aeruginosa high-risk clones, Drug resistance updates: reviews and commentaries in antimicrobial and anticancer chemotherapy 21-22 (2015) 41–59.

[2] K.S. Kaye, J.M. Pogue, Infections Caused by Resistant Gram-Negative Bacteria: Epidemiology and Management, Pharmacotherapy 35(10) (2015) 949–62.

[3] J.S. Talwalkar, T.S. Murray, The Approach to Pseudomonas aeruginosa in Cystic Fibrosis, Clin Chest Med 37(1) (2016) 69–81.

[4] L. Martinez-Solano, M.D. Macia, A. Fajardo, A. Oliver, J.L. Martinez, Chronic *Pseudomonas aeruginosa* infection in chronic obstructive pulmonary disease, Clinical Infectious Diseases 47(12) (2008) 1526–33.

[5] J. Eklof, R. Sorensen, T.S. Ingebrigtsen, P. Sivapalan, I. Achir, J.B. Boel, J. Bangsborg, C. Ostergaard, R.B. Dessau, U.S. Jensen, A. Browatzki, T.S. Lapperre, J. Janner, U.M. Weinreich, K. Armbruster, T. Wilcke, N. Seersholm, J.U.S. Jensen, Pseudomonas aeruginosa and risk of death and exacerbations in patients with chronic obstructive pulmonary disease: an observational cohort study of 22 053 patients, Clinical microbiology and infection: the official publication of the European Society of Clinical Microbiology and Infectious Diseases (2019).

[6] M.F. Moradali, S. Ghods, B.H. Rehm, Pseudomonas aeruginosa Lifestyle: A Paradigm for Adaptation, Survival, and Persistence, Frontiers in cellular and infection microbiology 7 (2017) 39.

[7] L. Wiehlmann, G. Wagner, N. Cramer, B. Siebert, P. Gudowius, G. Morales, T. Kohler, C. van Delden, C. Weinel, P. Slickers, B. Tummler, Population structure of Pseudomonas aeruginosa, Proceedings of the National Academy of Sciences of the United States of America 104(19) (2007) 8101–6.

[8] G. Morales, L. Wiehlmann, P. Gudowius, C. van Delden, B. Tummler, J.L. Martinez, F. Rojo, Structure of Pseudomonas aeruginosa populations analyzed by single nucleotide polymorphism and pulsed-field gel electrophoresis genotyping, Journal of bacteriology 186(13) (2004) 4228–37.

[9] A. Alonso, F. Rojo, J.L. Martinez, Environmental and clinical isolates of *Pseudomonas aeruginosa* show pathogenic and biodegradative properties irrespective of their origin, Environmental microbiology 1(5) (1999) 421–30.

[10] J.L. Martinez, Bacterial pathogens: from natural ecosystems to human hosts, Environ Microbiol 15(2) (2013) 325–33.

[11] B. Valot, C. Guyeux, J.Y. Rolland, K. Mazouzi, X. Bertrand, D. Hocquet, What It Takes to Be a Pseudomonas aeruginosa? The Core Genome of the Opportunistic Pathogen Updated, PloS one 10(5) (2015) e0126468.

[12] S. Mahajan-Miklos, L.G. Rahme, F.M. Ausubel, Elucidating the molecular mechanisms of bacterial virulence using non-mammalian hosts, Molecular microbiology 37(5) (2000) 981–8.

[13] S.P. Diggle, M. Whiteley, Microbe Profile: Pseudomonas aeruginosa: opportunistic pathogen and lab rat, Microbiology (Reading, England) (2019).

[14] M.W. Azam, A.U. Khan, Updates on the pathogenicity status of Pseudomonas aeruginosa, Drug discovery today 24(1) (2019) 350–359.

[15] J. Lee, L. Zhang, The hierarchy quorum sensing network in Pseudomonas aeruginosa, Protein Cell 6(1) (2015) 26–41.

[16] P. Williams, M. Camara, Quorum sensing and environmental adaptation in Pseudomonas aeruginosa: a tale of regulatory networks and multifunctional signal molecules, Curr Opin Microbiol 12(2) (2009) 182–91.

[17] A. Sandri, A. Ortombina, F. Boschi, E. Cremonini, M. Boaretti, C. Sorio, P. Melotti, G. Bergamini, M. Lleo, Inhibition of Pseudomonas aeruginosa secreted virulence factors reduces lung inflammation in CF mice, Virulence 9(1) (2018) 1008–1018.

[18] J.L. Kadurugamuwa, T.J. Beveridge, Natural release of virulence factors in membrane vesicles by *Pseudomonas aeruginosa* and the effect of aminoglycoside antibiotics on their release, The Journal of antimicrobial chemotherapy 40(5) (1997) 615–21.

[19] J.L. Kadurugamuwa, T.J. Beveridge, Virulence factors are released from Pseudomonas aeruginosa in association with membrane vesicles during normal growth and exposure to gentamicin: a novel mechanism of enzyme secretion, Journal of bacteriology 177(14) (1995) 3998–4008.

[20] J. Botelho, F. Grosso, L. Peixe, Antibiotic resistance in Pseudomonas aeruginosa - Mechanisms, epidemiology and evolution, Drug resistance updates: reviews and commentaries in antimicrobial and anticancer chemotherapy 44 (2019) 100640.

[21] Z. Pang, R. Raudonis, B.R. Glick, T.J. Lin, Z. Cheng, Antibiotic resistance in Pseudomonas aeruginosa: mechanisms and alternative therapeutic strategies, Biotechnol Adv 37(1) (2019) 177–192.

[22] A. Fajardo, N. Martinez-Martin, M. Mercadillo, J.C. Galan, B. Ghysels, S. Matthijs, P. Cornelis, L. Wiehlmann, B. Tummler, F. Baquero, J.L. Martinez, The neglected intrinsic resistome of bacterial pathogens, PloS one 3(2) (2008) e1619.

[23] T. Miyoshi-Akiyama, T. Tada, N. Ohmagari, N. Viet Hung, P. Tharavichitkul, B.M. Pokhrel, M. Gniadkowski, M. Shimojima, T. Kirikae, Emergence and Spread of Epidemic Multidrug-Resistant Pseudomonas aeruginosa, Genome biology and evolution 9(12) (2017) 3238–3245.

[24] J.A. Reales-Calderon, F. Corona, L. Monteoliva, C. Gil, J.L. Martinez, Quantitative proteomics unravels that the post-transcriptional regulator Crc modulates the generation of vesicles and secreted virulence determinants of Pseudomonas aeruginosa, Journal of proteomics 127(Pt B) (2015) 352–64.

[25] J.F. Linares, R. Moreno, A. Fajardo, L. Martinez-Solano, R. Escalante, F. Rojo, J.L. Martinez, The global regulator Crc modulates metabolism, susceptibility to antibiotics and virulence in Pseudomonas aeruginosa, Environ Microbiol 12(12) (2010) 3196–212.

[26] E. Sonnleitner, A. Wulf, S. Campagne, X.Y. Pei, M.T. Wolfinger, G. Forlani, K. Prindl, L. Abdou, A. Resch, F.H. Allain, B.F. Luisi, H. Urlaub, U. Blasi, Interplay between the catabolite repression control protein Crc, Hfq and RNA in Hfq-dependent translational regulation in Pseudomonas aeruginosa, Nucleic acids research (2017).

[27] E. Sonnleitner, K. Prindl, U. Blasi, The Pseudomonas aeruginosa CrcZ RNA interferes with Hfq-mediated riboregulation, PloS one 12(7) (2017) e0180887.

[28] D.R. Gifford, V. Furio, A. Papkou, T. Vogwill, A. Oliver, R.C. MacLean, Identifying and exploiting genes that potentiate the evolution of antibiotic resistance, Nat Ecol Evol 2(6) (2018) 1033–1039.

[29] C. Lopez-Causape, L.M. Sommer, G. Cabot, R. Rubio, A.A. Ocampo-Sosa, H.K. Johansen, J. Figuerola, R. Canton, T.J. Kidd, S. Molin, A. Oliver, Evolution of the Pseudomonas aeruginosa mutational resistome in an international Cystic Fibrosis clone, Scientific reports 7(1) (2017) 5555.

[30] G. Cabot, L. Zamorano, B. Moya, C. Juan, A. Navas, J. Blazquez, A. Oliver, Evolution of Pseudomonas aeruginosa Antimicrobial Resistance and Fitness under Low and High Mutation Rates, Antimicrob Agents Chemother 60(3) (2016) 1767–78.

[31] F. Sanz-Garcia, C. Alvarez-Ortega, J. Olivares-Pacheco, P. Blanco, J.L. Martinez, S. Hernando-Amado, Analysis of the Pseudomonas aeruginosa Aminoglycoside Differential Resistomes Allows Defining Genes Simultaneously Involved in Intrinsic Antibiotic Resistance and Virulence, Antimicrob Agents Chemother 63(5) (2019).

[32] S. Hernando-Amado, F. Sanz-Garcia, J.L. Martinez, Antibiotic Resistance Evolution Is Contingent on the Quorum-Sensing Response in Pseudomonas aeruginosa, Molecular biology and evolution 36(10) (2019) 2238–2251.

[33] F. Sanz-Garcia, S. Hernando-Amado, J.L. Martinez, Mutation-Driven Evolution of Pseudomonas aeruginosa in the Presence of either Ceftazidime or Ceftazidime-Avibactam, Antimicrob Agents Chemother 62(10) (2018).

[34] F. Sanz-Garcia, S. Hernando-Amado, J.L. Martinez, Mutational Evolution of Pseudomonas aeruginosa Resistance to Ribosome-Targeting Antibiotics, Frontiers in genetics 9 (2018) 451.

[35] G.L. Winsor, E.J. Griffiths, R. Lo, B.K. Dhillon, J.A. Shay, F.S. Brinkman, Enhanced annotations and features for comparing thousands of Pseudomonas genomes in the Pseudomonas genome database, Nucleic acids research 44(D1) (2016) D646–53.

[36] W. Huang, L.K. Brewer, J.W. Jones, A.T. Nguyen, A. Marcu, D.S. Wishart, A.G. Oglesby-Sherrouse, M.A. Kane, A. Wilks, PAMDB: a comprehensive Pseudomonas aeruginosa metabolome database, Nucleic acids research 46(D1) (2018) D575–d580.

[37] C. Gaviard, T. Jouenne, J. Hardouin, Proteomics of Pseudomonas aeruginosa: the increasing role of post-translational modifications, Expert review of proteomics 15(9) (2018) 757–772.

[38] M.O. Hesselager, M.C. Codrea, Z. Sun, E.W. Deutsch, T.B. Bennike, A. Stensballe, L. Bundgaard, R.L. Moritz, E. Bendixen, The Pig PeptideAtlas: A resource for systems biology in animal production and biomedicine, Proteomics 16(4) (2016) 634–44.

[39] J.A. Vizcaino, A. Csordas, N. Del-Toro, J.A. Dianes, J. Griss, I. Lavidas, G. Mayer, Y. Perez-Riverol, F. Reisinger, T. Ternent, Q.W. Xu, R. Wang, H. Hermjakob, 2016 update of the PRIDE database and its related tools, Nucleic acids research 44(22) (2016) 11033.

[40] D. Dacheux, J. Goure, J. Chabert, Y. Usson, I. Attree, Pore-forming activity of type III system-secreted proteins leads to oncosis of Pseudomonas aeruginosa-infected macrophages, Molecular microbiology 40(1) (2001) 76–85.

[41] F. Corona, J.A. Reales-Calderon, C. Gil, J.L. Martinez, The development of a new parameter for tracking post-transcriptional regulation allows the detailed map of the Pseudomonas aeruginosa Crc regulon, Scientific reports 8(1) (2018) 16793.

[42] J.A. Vizcaino, R.G. Cote, A. Csordas, J.A. Dianes, A. Fabregat, J.M. Foster, J. Griss, E. Alpi, M. Birim, J. Contell, G. O’Kelly, A. Schoenegger, D. Ovelleiro, Y. Perez-Riverol, F. Reisinger, D. Rios, R. Wang, H. Hermjakob, The PRoteomics IDEntifications (PRIDE) database and associated tools: status in 2013, Nucleic acids research 41(Database issue) (2013) D1063–9.

[43] F.A. Herbst, M.T. Sondergaard, H. Kjeldal, A. Stensballe, P.H. Nielsen, M.S. Dueholm, Major proteomic changes associated with amyloid-induced biofilm formation in Pseudomonas aeruginosa PAO1, Journal of proteome research 14(1) (2015) 72–81.

[44] A. Penesyan, S.S. Kumar, K. Kamath, A.M. Shathili, V. Venkatakrishnan, C. Krisp, N.H. Packer, M.P. Molloy, I.T. Paulsen, Genetically and Phenotypically Distinct Pseudomonas aeruginosa Cystic Fibrosis Isolates Share a Core Proteomic Signature, PloS one 10(10) (2015) e0138527.

[45] M.G. Casabona, Y. Vandenbrouck, I. Attree, Y. Coute, Proteomic characterization of Pseudomonas aeruginosa PAO1 inner membrane, Proteomics 13(16) (2013) 2419–23.

[46] M. Robert-Genthon, M.G. Casabona, D. Neves, Y. Coute, F. Ciceron, S. Elsen, A. Dessen, I. Attree, Unique features of a Pseudomonas aeruginosa alpha2-macroglobulin homolog, mBio 4(4) (2013).

[47] L. Martens, M. Chambers, M. Sturm, D. Kessner, F. Levander, J. Shofstahl, W.H. Tang, A. Rompp, S. Neumann, A.D. Pizarro, L. Montecchi-Palazzi, N. Tasman, M. Coleman, F. Reisinger, P. Souda, H. Hermjakob, P.A. Binz, E.W. Deutsch, mzML--a community standard for mass spectrometry data, Molecular & cellular proteomics: MCP 10(1) (2011) R110.000133.

[48] J.K. Eng, T.A. Jahan, M.R. Hoopmann, Comet: an open-source MS/MS sequence database search tool, Proteomics 13(1) (2013) 22–4.

[49] J.K. Eng, A.L. McCormack, J.R. Yates, An approach to correlate tandem mass spectral data of peptides with amino acid sequences in a protein database, Journal of the American Society for Mass Spectrometry 5(11) (1994) 976–89.

[50] R. Craig, R.C. Beavis, TANDEM: matching proteins with tandem mass spectra, Bioinformatics (Oxford, England) 20(9) (2004) 1466–7.

[51] B. MacLean, J.K. Eng, R.C. Beavis, M. McIntosh, General framework for developing and evaluating database scoring algorithms using the TANDEM search engine, Bioinformatics (Oxford, England) 22(22) (2006) 2830–2.

[52] L.Y. Geer, S.P. Markey, J.A. Kowalak, L. Wagner, M. Xu, D.M. Maynard, X. Yang, W. Shi, S.H. Bryant, Open mass spectrometry search algorithm, Journal of proteome research 3(5) (2004) 958–64.

[53] E.W. Deutsch, L. Mendoza, D. Shteynberg, J. Slagel, Z. Sun, R.L. Moritz, Trans-Proteomic Pipeline, a standardized data processing pipeline for large-scale reproducible proteomics informatics, Proteomics. Clinical applications 9(7-8) (2015) 745–54.

[54] E.W. Deutsch, L. Mendoza, D. Shteynberg, T. Farrah, H. Lam, N. Tasman, Z. Sun, E. Nilsson, B. Pratt, B. Prazen, J.K. Eng, D.B. Martin, A.I. Nesvizhskii, R. Aebersold, A guided tour of the Trans-Proteomic Pipeline, Proteomics 10(6) (2010) 1150–9.

[55] A. Keller, A.I. Nesvizhskii, E. Kolker, R. Aebersold, Empirical statistical model to estimate the accuracy of peptide identifications made by MS/MS and database search, Analytical chemistry 74(20) (2002) 5383–92.

[56] D. Shteynberg, E.W. Deutsch, H. Lam, J.K. Eng, Z. Sun, N. Tasman, L. Mendoza, R.L. Moritz, R. Aebersold, A.I. Nesvizhskii, iProphet: multi-level integrative analysis of shotgun proteomic data improves peptide and protein identification rates and error estimates, Molecular & cellular proteomics: MCP 10(12) (2011) M111.007690.

[57] A.I. Nesvizhskii, A. Keller, E. Kolker, R. Aebersold, A statistical model for identifying proteins by tandem mass spectrometry, Analytical chemistry 75(17) (2003) 4646–58.

[58] L. Reiter, M. Claassen, S.P. Schrimpf, M. Jovanovic, A. Schmidt, J.M. Buhmann, M.O. Hengartner, R. Aebersold, Protein identification false discovery rates for very large proteomics data sets generated by tandem mass spectrometry, Molecular & cellular proteomics: MCP 8(11) (2009) 2405–17.

[59] M. Kanehisa, Y. Sato, KEGG Mapper for inferring cellular functions from protein sequences, Protein science: a publication of the Protein Society (2019).

[60] M. Kanehisa, M. Furumichi, M. Tanabe, Y. Sato, K. Morishima, KEGG: new perspectives on genomes, pathways, diseases and drugs, Nucleic acids research 45(D1) (2017) D353–d361.

[61] C.K. Stover, X.Q. Pham, A.L. Erwin, S.D. Mizoguchi, P. Warrener, M.J. Hickey, F.S.L. Brinkman, W.O. Hufnagle, D.J. Kowalik, M. Lagrou, R.L. Garber, L. Goltry, E. Tolentino, S. Westbrock-Wadman, Y. Yuan, L.L. Brody, S.N. Coulter, K.R. Folger, A. Kas, K. Larbig, R. Lim, K. Smith, D. Spencer, G.K.-S. Wong, Z. Wu, I.T. Paulsen, J. Reizer, M.H. Saier, R.E.W. Hancock, S. Lory, M.V. Olson, Complete genome sequence of *Pseudomonas aeruginosa* PAO1, an opportunistic pathogen, Nature 406(6799) (2000) 959–964.

[62] M. Schobert, D. Jahn, Anaerobic physiology of Pseudomonas aeruginosa in the cystic fibrosis lung, International journal of medical microbiology: IJMM 300(8) (2010) 549–56.

[63] J. Walker, G. Moore, Pseudomonas aeruginosa in hospital water systems: biofilms, guidelines, and practicalities, The Journal of hospital infection 89(4) (2015) 324–7.

[64] J.F. Linares, I. Gustafsson, F. Baquero, J.L. Martinez, Antibiotics as intermicrobial signaling agents instead of weapons, Proceedings of the National Academy of Sciences of the United States of America 103(51) (2006) 19484–9.

[65] Y. Zhang, Z. Lin, P. Hao, K. Hou, Y. Sui, K. Zhang, Y. He, H. Li, H. Yang, S. Liu, Y. Ren, Improvement of Peptide Separation for Exploring the Missing Proteins Localized on Membranes, Journal of proteome research 17(12) (2018) 4152–4159.

[66] T. Beaudoin, L. Zhang, A.J. Hinz, C.J. Parr, T.F. Mah, The Biofilm-Specific Antibiotic Resistance Gene ndvB Is Important for Expression of Ethanol Oxidation Genes in Pseudomonas aeruginosa Biofilms, Journal of bacteriology 194(12) (2012) 3128–36.

[67] M. Zhu, J. Zhao, H. Kang, W. Kong, Y. Zhao, M. Wu, H. Liang, Corrigendum: Modulation of Type III Secretion System in Pseudomonas aeruginosa: Involvement of the PA4857 Gene Product, Frontiers in microbiology 7 (2016).

[68] M. Zhu, J. Zhao, H. Kang, W. Kong, Y. Zhao, M. Wu, H. Liang, Modulation of Type III Secretion System in Pseudomonas aeruginosa: Involvement of the PA4857 Gene Product, Frontiers in microbiology 7 (2016).

[69] E.E. Smith, D.G. Buckley, Z. Wu, C. Saenphimmachak, L.R. Hoffman, D.A. D’Argenio, S.I. Miller, B.W. Ramsey, D.P. Speert, S.M. Moskowitz, J.L. Burns, R. Kaul, M.V. Olson, Genetic adaptation by Pseudomonas aeruginosa to the airways of cystic fibrosis patients, Proceedings of the National Academy of Sciences of the United States of America 103(22) (2006) 8487–92.

[70] K.M. Colvin, Y. Irie, C.S. Tart, R. Urbano, J.C. Whitney, C. Ryder, P.L. Howell, D.J. Wozniak, M.R. Parsek, The Pel and Psl polysaccharides provide Pseudomonas aeruginosa structural redundancy within the biofilm matrix, Environ Microbiol 14(8) (2012) 1913–28.

[71] L.R. Hoffman, D.A. D’Argenio, M.J. MacCoss, Z. Zhang, R.A. Jones, S.I. Miller, Aminoglycoside antibiotics induce bacterial biofilm formation, Nature 436(7054) (2005) 1171–5.

[72] N. Pakharukova, M. Tuittila, S. Paavilainen, H. Malmi, O. Parilova, S. Teneberg, S.D. Knight, A.V. Zavialov, Structural basis for Acinetobacter baumannii biofilm formation, Proceedings of the National Academy of Sciences of the United States of America 115(21) (2018) 5558–5563.

[73] B.R. Borlee, A.D. Goldman, K. Murakami, R. Samudrala, D.J. Wozniak, M.R. Parsek, Pseudomonas aeruginosa uses a cyclic-di-GMP-regulated adhesin to reinforce the biofilm extracellular matrix, Molecular microbiology 75(4) (2010) 827–42.

[74] A. Balyimez, J.A. Colmer-Hamood, M. San Francisco, A.N. Hamood, Characterization of the Pseudomonas aeruginosa metalloendopeptidase, Mep72, a member of the Vfr regulon, BMC microbiology 13 (2013) 269.

[75] I.J. Passmore, K. Nishikawa, K.S. Lilley, S.D. Bowden, J.C. Chung, M. Welch, Mep72, a metzincin protease that is preferentially secreted by biofilms of Pseudomonas aeruginosa, Journal of bacteriology 197(4) (2015) 762–73.

[76] J.C. Chung, O. Rzhepishevska, M. Ramstedt, M. Welch, Type III secretion system expression in oxygen-limited Pseudomonas aeruginosa cultures is stimulated by isocitrate lyase activity, Open biology 3(1) (2013) 120131.

[77] G.H. Kaihami, L.C.D. Breda, J.R.F. de Almeida, T. de Oliveira Pereira, G.G. Nicastro, A.L. Boechat, S.R. de Almeida, R.L. Baldini, The Atypical Response Regulator AtvR Is a New Player in Pseudomonas aeruginosa Response to Hypoxia and Virulence, Infection and immunity 85(8) (2017).

[78] G.I. Lucchesi, T.A. Lisa, C.H. Casale, C.E. Domenech, Carnitine resembles choline in the induction of cholinesterase, acid phosphatase, and phospholipase C and in its action as an osmoprotectant in Pseudomonas aeruginosa, Current microbiology 30(1) (1995) 55–60.

[79] J.A. Meadows, M.J. Wargo, Characterization of Pseudomonas aeruginosa growth on O-acylcarnitines and identification of a short-chain acylcarnitine hydrolase, Applied and environmental microbiology 79(11) (2013) 3355–63.

[80] K. Koeppen, R. Barnaby, A.A. Jackson, S.A. Gerber, D.A. Hogan, B.A. Stanton, Tobramycin reduces key virulence determinants in the proteome of Pseudomonas aeruginosa outer membrane vesicles, PloS one 14(1) (2019) e0211290.

[81] P. Tielen, H. Kuhn, F. Rosenau, K.E. Jaeger, H.C. Flemming, J. Wingender, Interaction between extracellular lipase LipA and the polysaccharide alginate of Pseudomonas aeruginosa, BMC microbiology 13 (2013) 159.

[82] Y. Jin, H. Yang, M. Qiao, S. Jin, MexT regulates the type III secretion system through MexS and PtrC in Pseudomonas aeruginosa, Journal of bacteriology 193(2) (2011) 399–410.

[83] T. Kohler, S.F. Epp, L.K. Curty, J.C. Pechere, Characterization of MexT, the regulator of the MexE-MexF-OprN multidrug efflux system of Pseudomonas aeruginosa, Journal of bacteriology 181(20) (1999) 6300–5.

[84] S.K. Arora, B.W. Ritchings, E.C. Almira, S. Lory, R. Ramphal, A transcriptional activator, FleQ, regulates mucin adhesion and flagellar gene expression in Pseudomonas aeruginosa in a cascade manner, Journal of bacteriology 179(17) (1997) 5574–81.

[85] J.D. Mougous, M.E. Cuff, S. Raunser, A. Shen, M. Zhou, C.A. Gifford, A.L. Goodman, G. Joachimiak, C.L. Ordonez, S. Lory, T. Walz, A. Joachimiak, J.J. Mekalanos, A virulence locus of Pseudomonas aeruginosa encodes a protein secretion apparatus, Science (New York, N.Y.) 312(5779) (2006) 1526–30.

[86] A. Hachani, N.S. Lossi, A. Hamilton, C. Jones, S. Bleves, D. Albesa-Jove, A. Filloux, Type VI secretion system in Pseudomonas aeruginosa: secretion and multimerization of VgrG proteins, The Journal of biological chemistry 286(14) (2011) 12317–27.

[87] F. Corona, P. Blanco, M. Alcalde-Rico, S. Hernando-Amado, F. Lira, A. Bernardini, M.B. Sanchez, J.L. Martinez, The analysis of the antibiotic resistome offers new opportunities for therapeutic intervention, Future medicinal chemistry 8(10) (2016) 1133–51.

[88] A. Daddaoua, C. Molina-Santiago, J. de la Torre, T. Krell, J.L. Ramos, GtrS and GltR form a two-component system: the central role of 2-ketogluconate in the expression of exotoxin A and glucose catabolic enzymes in Pseudomonas aeruginosa, Nucleic acids research 42(12) (2014) 7654–63.

[89] K.F. Kong, S.R. Jayawardena, S.D. Indulkar, A. Del Puerto, C.L. Koh, N. Hoiby, K. Mathee, Pseudomonas aeruginosa AmpR is a global transcriptional factor that regulates expression of AmpC and PoxB beta-lactamases, proteases, quorum sensing, and other virulence factors, Antimicrob Agents Chemother 49(11) (2005) 4567–75.

[90] G. Dieppois, V. Ducret, O. Caille, K. Perron, The transcriptional regulator CzcR modulates antibiotic resistance and quorum sensing in Pseudomonas aeruginosa, PloS one 7(5) (2012) e38148.

[91] K.N. Raymond, E.A. Dertz, S.S. Kim, Enterobactin: an archetype for microbial iron transport, Proceedings of the National Academy of Sciences of the United States of America 100(7) (2003) 3584–8.

[92] S. Chevalier, E. Bouffartigues, A. Bazire, A. Tahrioui, R. Duchesne, D. Tortuel, O. Maillot, T. Clamens, N. Orange, M.G.J. Feuilloley, O. Lesouhaitier, A. Dufour, P. Cornelis, Extracytoplasmic function sigma factors in Pseudomonas aeruginosa, Biochimica et biophysica acta, Gene regulatory mechanisms 1862(7) (2019) 706–721.

[93] M. Hannauer, A. Braud, F. Hoegy, P. Ronot, A. Boos, I.J. Schalk, The PvdRT-OpmQ efflux pump controls the metal selectivity of the iron uptake pathway mediated by the siderophore pyoverdine in Pseudomonas aeruginosa, Environ Microbiol 14(7) (2012) 1696–708.

[94] S. Elias, E. Degtyar, E. Banin, FvbA is required for vibriobactin utilization in Pseudomonas aeruginosa, Microbiology (Reading) 157(Pt 7) (2011) 2172–2180.

[95] M.A. Llamas, M. Sparrius, R. Kloet, C.R. Jimenez, C. Vandenbroucke-Grauls, W. Bitter, The heterologous siderophores ferrioxamine B and ferrichrome activate signaling pathways in Pseudomonas aeruginosa, Journal of bacteriology 188(5) (2006) 1882–91.

[96] K.C. Bastiaansen, P. van Ulsen, M. Wijtmans, W. Bitter, M.A. Llamas, Self-cleavage of the Pseudomonas aeruginosa Cell-surface Signaling Anti-sigma Factor FoxR Occurs through an N-O Acyl Rearrangement, The Journal of biological chemistry 290(19) (2015) 12237–46.

[97] R. Wegele, R. Tasler, Y. Zeng, M. Rivera, N. Frankenberg-Dinkel, The heme oxygenase(s)-phytochrome system of Pseudomonas aeruginosa, The Journal of biological chemistry 279(44) (2004) 45791–802.

[98] K.A. McFarland, E.L. Dolben, M. LeRoux, T.K. Kambara, K.M. Ramsey, R.L. Kirkpatrick, J.D. Mougous, D.A. Hogan, S.L. Dove, A self-lysis pathway that enhances the virulence of a pathogenic bacterium, Proceedings of the National Academy of Sciences of the United States of America 112(27) (2015) 8433–8.

[99] J. Malmstrom, M. Beck, A. Schmidt, V. Lange, E.W. Deutsch, R. Aebersold, Proteome-wide cellular protein concentrations of the human pathogen Leptospira interrogans, Nature 460(7256) (2009) 762–5.

[100] P.T. Van, A.K. Schmid, N.L. King, A. Kaur, M. Pan, K. Whitehead, T. Koide, M.T. Facciotti, Y.A. Goo, E.W. Deutsch, D.J. Reiss, P. Mallick, N.S. Baliga, Halobacterium salinarum NRC-1 PeptideAtlas: toward strategies for targeted proteomics and improved proteome coverage, Journal of proteome research 7(9) (2008) 3755–64.

[101] V. Lange, J.A. Malmstrom, J. Didion, N.L. King, B.P. Johansson, J. Schafer, J. Rameseder, C.H. Wong, E.W. Deutsch, M.Y. Brusniak, P. Buhlmann, L. Bjorck, B. Domon, R. Aebersold, Targeted quantitative analysis of Streptococcus pyogenes virulence factors by multiple reaction monitoring, Molecular & cellular proteomics: MCP 7(8) (2008) 1489–500.

