## Supplemental Tables S1 and S2 for "A wide-ranging *Pseudomonas aeruginosa* PeptideAtlas build: a useful proteomic resource for a versatile pathogen"

**TABLE S1.** Database search parameters

| | | pao1_fcp001_proteome_4plex/ pao1_vescl_secretome_4plex/ Pseudomonas_Cystic_Fibrosis | pao1_anaerobiosis/ pao1_biofilm/ pao1_growth_20c/ pao1_growth_TCN/ pao1_inner_memb/ pao1_outer_memb/ pao1_secretome_T3SS/ pao1_stat_phase/ pao1_T3SS_induction | pao1_itraq_4plex/ pao1_itraq_6plex/ pao1_itraq_8plex | PAO1_Fap | PAO1_Inner_Membrane_Proteome/ PAO1_MagD-WT/ PAO1_delta-MagD |
|---|---|---|---|---|---|---|
| Instrument | | Triple TOF 5600 System (AB SCIEX, Concord, ON) | LTQ-Orbitrap | AB SCIEX TripleTOF 5600 | Q-Exactive | LTQ-Orbitrap Velos |
| Precursor Monoisotopic Mass Tolerance | comet | 50ppm | 20ppm | 50ppm | 20ppm | 20ppm |
| | X! tandem | 50ppm | 20ppm | 50ppm | 20ppm | 20ppm |
| | OMSSA | 0.1 | 0.1 | 0.1 | 0.1 | 0.1 |
| Fragment Monoisotopic Mass Tolerance | comet | 0.1 | 0.4 | 0.1 | 0.03 | 0.4 |
| | X! tandem | 50ppm | 0.4 | 50ppm | 20ppm | 0.4 |
| | OMSSA | 0.1 | 0.4 | 0.1 | 0.1 | 0.4 |
| Number of tryptic termini | | 1 | 1 | 1 | 1 | 1 |
| Number of allowed missed cleavages | | 2 | 2 | 2 | 2 | 2 |
| Modifications | | Cysteine carboxyamidomethylation was set as fixed modification, oxidation of methionine, and transformation of N-terminal glutamine and N-terminal glutamic acid residue in the pyroglutamic acid form were included as variable modifications. | | | | |
| Extra Static Modification | | Static iTRAQ4plex on Lys and peptide N-termini | | iTRAQ4plex, iTRAQ6plex, or iTRAQ8plex on Lys and peptide N-termini | | |

**TABLE S2.** Proteins identified in each sample.Proteins uniquely identified in each sample are highlighted.

| Sample 15 | Sample 8 | Sample 9 | Sample 10 | Sample 1 | Sample 16 | Sample 4 | Sample 7 | Sample 11 | Sample 5 | Sample 3 | Sample 6 | Sample 14 | Sample 13 | Sample 17 | Sample 2 | Sample 12 | Sample 18 | Sample 19 |
|---|---|---|---|---|---|---|---|---|---|---|---|---|---|---|---|---|---|---|
| PA0001 | PA0001 | PA0001 | PA0001 | PA0001 | PA0001 | PA0001 | N/A | PA0001 | PA0001 | PA0001 | PA0001 | PA0001 | PA0001 | PA0001 | N/A | N/A | PA0001 | N/A |
| PA0002 | PA0002 | PA0002 | PA0002 | PA0002 | PA0002 | PA0002 | N/A | N/A | PA0002 | PA0002 | PA0002 | PA0002 | PA0002 | PA0002 | PA0002 | N/A | PA0002 | PA0002 |
| PA0003 | N/A | N/A | PA0003 | PA0003 | PA0003 | N/A | PA0003 | PA0003 | PA0003 | PA0003 | PA0003 | PA0003 | PA0003 | N/A | N/A | N/A | N/A | N/A |
| PA0004 | PA0004 | PA0004 | PA0004 | PA0004 | PA0004 | PA0004 | PA0004 | PA0004 | PA0004 | PA0004 | PA0004 | PA0004 | PA0004 | PA0004 | PA0004 | PA0004 | PA0004 | N/A |
| PA0005 | PA0005 | PA0005 | PA0005 | PA0005 | PA0005 | PA0005 | N/A | N/A | PA0005 | N/A | PA0005 | PA0005 | PA0005 | PA0005 | PA0005 | N/A | PA0005 | N/A |
| PA0006 | N/A | PA0006 | PA0006 | PA0006 | PA0006 | N/A | N/A | N/A | PA0006 | PA0006 | PA0006 | PA0006 | PA0006 | N/A | PA0006 | N/A | N/A | N/A |
| PA0007 | N/A | N/A | N/A | N/A | PA0007 | N/A | N/A | N/A | N/A | N/A | N/A | N/A | N/A | N/A | N/A | N/A | N/A | N/A |
| PA0008 | PA0008 | PA0008 | PA0008 | PA0008 | PA0008 | PA0008 | PA0008 | PA0008 | PA0008 | PA0008 | PA0008 | PA0008 | PA0008 | PA0008 | PA0008 | N/A | N/A | N/A |
| PA0009 | PA0009 | PA0009 | PA0009 | PA0009 | PA0009 | PA0009 | PA0009 | PA0009 | PA0009 | PA0009 | PA0009 | PA0009 | PA0009 | PA0009 | PA0009 | N/A | N/A | N/A |
| PA0010 | N/A | N/A | N/A | PA0010 | N/A | N/A | N/A | N/A | PA0010 | N/A | N/A | N/A | N/A | N/A | N/A | N/A | N/A | N/A |
| PA0011 | PA0011 | PA0011 | N/A | PA0011 | N/A | PA0011 | PA0011 | PA0011 | PA0011 | PA0011 | PA0011 | PA0011 | PA0011 | PA0011 | N/A | N/A | PA0011 | N/A |
| PA0012 | N/A | N/A | N/A | PA0012 | PA0012 | N/A | N/A | N/A | N/A | N/A | PA0012 | N/A | N/A | N/A | N/A | N/A | N/A | N/A |
| N/A | N/A | N/A | N/A | PA0015 | N/A | N/A | N/A | N/A | N/A | N/A | PA0015 | N/A | N/A | N/A | N/A | N/A | N/A | N/A |
| PA0016 | PA0016 | PA0016 | PA0016 | PA0016 | N/A | N/A | N/A | PA0016 | PA0016 | PA0016 | PA0016 | PA0016 | PA0016 | PA0016 | N/A | N/A | PA0016 | N/A |
| PA0017 | PA0017 | PA0017 | PA0017 | PA0017 | PA0017 | N/A | N/A | PA0017 | PA0017 | N/A | PA0017 | PA0017 | N/A | PA0017 | N/A | N/A | N/A | N/A |
| PA0018 | PA0018 | PA0018 | PA0018 | PA0018 | PA0018 | PA0018 | N/A | N/A | PA0018 | PA0018 | PA0018 | PA0018 | PA0018 | N/A | PA0018 | N/A | N/A | N/A |
| PA0019 | PA0019 | PA0019 | PA0019 | PA0019 | PA0019 | PA0019 | PA0019 | N/A | PA0019 | PA0019 | PA0019 | PA0019 | PA0019 | N/A | PA0019 | N/A | PA0019 | N/A |
| PA0020 | PA0020 | PA0020 | PA0020 | PA0020 | PA0020 | PA0020 | PA0020 | PA0020 | PA0020 | PA0020 | PA0020 | PA0020 | PA0020 | PA0020 | PA0020 | N/A | N/A | N/A |
| PA0022 | PA0022 | PA0022 | N/A | PA0022 | PA0022 | PA0022 | PA0022 | N/A | PA0022 | PA0022 | PA0022 | N/A | PA0022 | PA0022 | N/A | N/A | N/A | N/A |
| PA0023 | N/A | PA0023 | PA0023 | PA0023 | PA0023 | PA0023 | N/A | PA0023 | PA0023 | PA0023 | PA0023 | PA0023 | PA0023 | PA0023 | N/A | N/A | N/A | N/A |
| PA0024 | PA0024 | PA0024 | PA0024 | PA0024 | PA0024 | PA0024 | PA0024 | N/A | PA0024 | PA0024 | PA0024 | PA0024 | PA0024 | N/A | PA0024 | N/A | N/A | N/A |
| PA0025 | PA0025 | PA0025 | N/A | PA0025 | PA0025 | N/A | PA0025 | N/A | PA0025 | N/A | PA0025 | PA0025 | PA0025 | N/A | N/A | N/A | N/A | N/A |
| PA0026 | PA0026 | N/A | N/A | N/A | N/A | N/A | N/A | N/A | N/A | N/A | N/A | N/A | N/A | N/A | PA0026 | PA0026 | N/A | N/A |
| PA0027 | N/A | N/A | N/A | N/A | N/A | N/A | N/A | N/A | N/A | N/A | N/A | N/A | N/A | N/A | N/A | N/A | N/A | N/A |
| PA0032 | N/A | N/A | N/A | N/A | N/A | N/A | N/A | N/A | N/A | N/A | N/A | N/A | N/A | N/A | N/A | N/A | N/A | N/A |
| PA0033 | N/A | N/A | N/A | N/A | N/A | N/A | N/A | N/A | N/A | N/A | N/A | N/A | N/A | N/A | N/A | N/A | N/A | N/A |
| PA0035 | PA0035 | PA0035 | PA0035 | PA0035 | PA0035 | PA0035 | N/A | N/A | PA0035 | PA0035 | PA0035 | PA0035 | PA0035 | N/A | PA0035 | N/A | N/A | N/A |
| PA0036 | PA0036 | PA0036 | PA0036 | PA0036 | PA0036 | PA0036 | N/A | N/A | PA0036 | PA0036 | PA0036 | PA0036 | PA0036 | N/A | N/A | N/A | N/A | N/A |
| PA0037 | N/A | N/A | N/A | PA0037 | PA0037 | N/A | N/A | N/A | N/A | N/A | N/A | N/A | N/A | N/A | N/A | N/A | N/A | N/A |
| PA0038 | N/A | N/A | N/A | PA0038 | PA0038 | PA0038 | PA0038 | N/A | PA0038 | PA0038 | PA0038 | PA0038 | PA0038 | PA0038 | N/A | PA0038 | N/A | N/A |
| PA0039 | N/A | N/A | N/A | N/A | PA0039 | N/A | N/A | N/A | PA0039 | N/A | PA0039 | PA0039 | PA0039 | N/A | N/A | N/A | N/A | N/A |
| PA0040 | N/A | N/A | N/A | PA0040 | N/A | N/A | N/A | PA0040 | PA0040 | PA0040 | PA0040 | PA0040 | PA0040 | N/A | PA0040 | N/A | N/A | N/A |
| PA0041 | N/A | N/A | N/A | PA0041 | PA0041 | N/A | N/A | PA0041 | N/A | PA0041 | PA0041 | N/A | N/A | PA0041 | PA0041 | PA0041 | PA0041 | PA0041 |
| PA0042 | N/A | N/A | N/A | N/A | N/A | N/A | PA0042 | N/A | PA0042 | N/A | PA0042 | PA0042 | N/A | N/A | N/A | N/A | N/A | N/A |
| N/A | N/A | N/A | N/A | N/A | N/A | N/A | N/A | N/A | N/A | N/A | N/A | N/A | N/A | PA0043 | N/A | N/A | N/A | N/A |
| N/A | N/A | N/A | N/A | PA0044 | N/A | N/A | N/A | N/A | N/A | PA0044 | N/A | N/A | N/A | N/A | N/A | N/A | N/A | N/A |
| PA0045 | PA0045 | PA0045 | PA0045 | PA0045 | N/A | PA0045 | PA0045 | PA0045 | PA0045 | PA0045 | PA0045 | PA0045 | N/A | PA0045 | PA0045 | N/A | PA0045 | N/A |
| N/A | PA0046 | PA0046 | PA0046 | PA0046 | N/A | N/A | PA0046 | N/A | PA0046 | PA0046 | PA0046 | N/A | N/A | N/A | PA0046 | N/A | N/A | N/A |
| PA0047 | PA0047 | PA0047 | N/A | PA0047 | N/A | N/A | PA0047 | N/A | PA0047 | PA0047 | N/A | N/A | N/A | N/A | PA0047 | N/A | N/A | N/A |
| PA0048 | N/A | N/A | N/A | N/A | N/A | N/A | N/A | N/A | N/A | N/A | N/A | PA0048 | N/A | N/A | N/A | N/A | N/A | N/A |
| PA0049 | N/A | N/A | N/A | PA0049 | N/A | N/A | N/A | N/A | N/A | N/A | N/A | N/A | N/A | N/A | PA0049 | N/A | N/A | N/A |
| PA0051 | N/A | N/A | N/A | N/A | N/A | N/A | N/A | N/A | PA0051 | PA0051 | PA0051 | PA0051 | PA0051 | N/A | N/A | N/A | N/A | N/A |
| PA0052 | N/A | N/A | N/A | N/A | N/A | N/A | N/A | N/A | N/A | N/A | N/A | PA0052 | PA0052 | N/A | N/A | N/A | N/A | N/A |
| PA0054 | N/A | N/A | N/A | N/A | N/A | N/A | N/A | N/A | N/A | N/A | N/A | N/A | N/A | N/A | N/A | N/A | N/A | N/A |
| PA0055 | PA0055 | PA0055 | PA0055 | PA0055 | N/A | PA0055 | PA0055 | N/A | PA0055 | PA0055 | PA0055 | PA0055 | PA0055 | N/A | PA0055 | N/A | N/A | N/A |
| PA0056 | N/A | N/A | N/A | N/A | PA0056 | N/A | N/A | N/A | N/A | N/A | N/A | N/A | N/A | N/A | N/A | N/A | N/A | N/A |
| PA0057 | N/A | N/A | N/A | N/A | PA0057 | N/A | N/A | N/A | N/A | N/A | N/A | N/A | N/A | N/A | N/A | N/A | N/A | N/A |
| N/A | N/A | N/A | N/A | N/A | PA0058 | N/A | N/A | N/A | N/A | N/A | N/A | N/A | N/A | N/A | N/A | N/A | N/A | N/A |
| PA0059 | N/A | N/A | N/A | N/A | PA0059 | PA0059 | N/A | N/A | PA0059 | PA0059 | PA0059 | PA0059 | PA0059 | N/A | N/A | N/A | N/A | N/A |
| N/A | N/A | N/A | N/A | N/A | N/A | N/A | N/A | N/A | N/A | N/A | PA0060 | N/A | PA0060 | N/A | N/A | N/A | N/A | N/A |
| PA0061 | N/A | N/A | N/A | N/A | N/A | N/A | N/A | N/A | N/A | PA0061 | N/A | PA0061 | N/A | N/A | N/A | N/A | N/A | N/A |
| PA0062 | N/A | N/A | N/A | N/A | N/A | N/A | N/A | N/A | N/A | N/A | N/A | N/A | N/A | N/A | N/A | N/A | N/A | N/A |
| PA0063 | N/A | N/A | N/A | PA0063 | N/A | N/A | N/A | N/A | N/A | N/A | N/A | N/A | N/A | N/A | N/A | N/A | N/A | N/A |
| PA0064 | PA0064 | PA0064 | PA0064 | PA0064 | PA0064 | N/A | N/A | N/A | N/A | N/A | PA0064 | PA0064 | PA0064 | PA0064 | PA0064 | N/A | PA0064 | PA0064 |
| PA0065 | N/A | PA0065 | PA0065 | PA0065 | PA0065 | N/A | PA0065 | PA0065 | PA0065 | N/A | PA0065 | PA0065 | PA0065 | N/A | N/A | N/A | N/A | N/A |
| PA0066 | N/A | N/A | N/A | N/A | PA0066 | PA0066 | N/A | N/A | PA0066 | PA0066 | PA0066 | PA0066 | PA0066 | N/A | N/A | N/A | PA0066 | N/A |
| PA0067 | PA0067 | PA0067 | PA0067 | PA0067 | PA0067 | PA0067 | N/A | N/A | PA0067 | PA0067 | PA0067 | PA0067 | PA0067 | N/A | PA0067 | PA0067 | N/A | N/A |
| PA0068 | N/A | N/A | N/A | N/A | N/A | N/A | N/A | N/A | PA0068 | PA0068 | PA0068 | PA0068 | PA0068 | N/A | N/A | N/A | N/A | N/A |
| N/A | N/A | N/A | N/A | N/A | PA0069 | N/A | N/A | N/A | N/A | N/A | N/A | N/A | N/A | N/A | N/A | N/A | N/A | N/A |
| PA0070 | PA0070 | PA0070 | PA0070 | PA0070 | PA0070 | PA0070 | PA0070 | PA0070 | PA0070 | PA0070 | PA0070 | PA0070 | PA0070 | PA0070 | PA0070 | PA0070 | PA0070 | PA0070 |
| PA0071 | PA0071 | PA0071 | N/A | PA0071 | N/A | N/A | N/A | PA0071 | N/A | PA0071 | PA0071 | N/A | N/A | PA0071 | N/A | N/A | N/A | N/A |
| PA0072 | N/A | N/A | N/A | N/A | N/A | N/A | N/A | N/A | PA0072 | N/A | N/A | N/A | PA0072 | PA0072 | N/A | N/A | N/A | N/A |
| PA0073 | N/A | N/A | N/A | PA0073 | N/A | N/A | PA0073 | N/A | PA0073 | N/A | PA0073 | N/A | PA0073 | PA0073 | N/A | N/A | PA0073 | N/A |
| PA0074 | PA0074 | PA0074 | PA0074 | PA0074 | PA0074 | N/A | PA0074 | PA0074 | PA0074 | PA0074 | PA0074 | PA0074 | PA0074 | PA0074 | N/A | N/A | PA0074 | N/A |
| PA0075 | PA0075 | N/A | PA0075 | PA0075 | PA0075 | N/A | PA0075 | N/A | PA0075 | PA0075 | PA0075 | PA0075 | N/A | N/A | N/A | N/A | N/A | N/A |
| PA0076 | PA0076 | PA0076 | N/A | PA0076 | N/A | N/A | N/A | N/A | N/A | N/A | PA0076 | PA0076 | N/A | N/A | PA0076 | N/A | N/A | N/A |
| PA0077 | PA0077 | PA0077 | PA0077 | PA0077 | N/A | N/A | PA0077 | PA0077 | PA0077 | PA0077 | PA0077 | PA0077 | PA0077 | PA0077 | PA0077 | N/A | PA0077 | PA0077 |
| PA0078 | PA0078 | PA0078 | PA0078 | PA0078 | N/A | N/A | PA0078 | PA0078 | PA0078 | N/A | PA0078 | PA0078 | PA0078 | PA0078 | PA0078 | N/A | PA0078 | PA0078 |
| PA0079 | PA0079 | PA0079 | PA0079 | PA0079 | N/A | PA0079 | PA0079 | PA0079 | PA0079 | PA0079 | PA0079 | PA0079 | PA0079 | PA0079 | PA0079 | N/A | PA0079 | PA0079 |
| PA0080 | PA0080 | PA0080 | PA0080 | PA0080 | N/A | N/A | PA0080 | N/A | PA0080 | PA0080 | PA0080 | PA0080 | PA0080 | PA0080 | PA0080 | N/A | N/A | N/A |
| PA0081 | N/A | N/A | PA0081 | PA0081 | N/A | N/A | PA0081 | PA0081 | N/A | N/A | PA0081 | N/A | N/A | N/A | N/A | N/A | PA0081 | N/A |
| PA0082 | PA0082 | PA0082 | N/A | PA0082 | PA0082 | N/A | PA0082 | N/A | PA0082 | PA0082 | PA0082 | N/A | PA0082 | N/A | N/A | N/A | N/A | N/A |
| PA0083 | PA0083 | PA0083 | PA0083 | PA0083 | PA0083 | PA0083 | PA0083 | PA0083 | PA0083 | PA0083 | PA0083 | PA0083 | PA0083 | N/A | PA0083 | N/A | PA0083 | PA0083 |
| PA0084 | PA0084 | PA0084 | PA0084 | PA0084 | PA0084 | PA0084 | PA0084 | PA0084 | PA0084 | PA0084 | PA0084 | PA0084 | PA0084 | PA0084 | PA0084 | PA0084 | PA0084 | PA0084 |
| PA0085 | PA0085 | PA0085 | PA0085 | PA0085 | PA0085 | PA0085 | PA0085 | PA0085 | PA0085 | PA0085 | PA0085 | PA0085 | PA0085 | PA0085 | PA0085 | N/A | PA0085 | PA0085 |
| PA0086 | PA0086 | PA0086 | PA0086 | PA0086 | PA0086 | N/A | PA0086 | N/A | PA0086 | PA0086 | PA0086 | PA0086 | PA0086 | N/A | PA0086 | N/A | N/A | N/A |
| N/A | N/A | N/A | N/A | N/A | N/A | N/A | PA0087 | N/A | N/A | N/A | N/A | N/A | N/A | N/A | N/A | N/A | N/A | N/A |
| PA0088 | N/A | N/A | N/A | PA0088 | N/A | N/A | N/A | PA0088 | N/A | N/A | N/A | N/A | PA0088 | N/A | N/A | N/A | PA0088 | PA0088 |
| PA0089 | N/A | N/A | N/A | N/A | PA0089 | N/A | N/A | N/A | N/A | N/A | N/A | N/A | N/A | N/A | N/A | N/A | N/A | PA0089 |
| PA0090 | PA0090 | PA0090 | PA0090 | PA0090 | PA0090 | PA0090 | N/A | PA0090 | PA0090 | PA0090 | PA0090 | PA0090 | PA0090 | N/A | N/A | N/A | PA0090 | PA0090 |
| PA0091 | N/A | PA0091 | N/A | PA0091 | N/A | PA0091 | N/A | N/A | PA0091 | PA0091 | PA0091 | PA0091 | PA0091 | PA0091 | N/A | N/A | N/A | N/A |
| PA0092 | PA0092 | PA0092 | N/A | PA0092 | N/A | PA0092 | N/A | N/A | PA0092 | N/A | PA0092 | PA0092 | PA0092 | N/A | N/A | N/A | N/A | N/A |
| PA0093 | N/A | N/A | N/A | PA0093 | PA0093 | N/A | N/A | N/A | N/A | N/A | PA0093 | PA0093 | N/A | N/A | N/A | N/A | N/A | N/A |
| PA0094 | PA0094 | PA0094 | PA0094 | PA0094 | PA0094 | N/A | PA0094 | N/A | PA0094 | PA0094 | PA0094 | PA0094 | PA0094 | N/A | PA0094 | N/A | PA0094 | PA0094 |
| PA0095 | N/A | PA0095 | N/A | PA0095 | N/A | PA0095 | N/A | N/A | PA0095 | PA0095 | PA0095 | PA0095 | PA0095 | PA0095 | N/A | N/A | N/A | N/A |
| PA0097 | N/A | N/A | N/A | N/A | N/A | N/A | N/A | N/A | PA0097 | N/A | N/A | N/A | N/A | N/A | N/A | N/A | N/A | N/A |
| N/A | N/A | N/A | N/A | N/A | PA0099 | N/A | N/A | N/A | N/A | N/A | N/A | N/A | N/A | N/A | N/A | N/A | N/A | N/A |
| PA0100 | PA0100 | N/A | N/A | PA0100 | N/A | N/A | N/A | N/A | N/A | PA0100 | PA0100 | PA0100 | PA0100 | N/A | N/A | N/A | N/A | N/A |
| N/A | N/A | N/A | N/A | N/A | PA0101 | N/A | N/A | N/A | N/A | N/A | N/A | N/A | N/A | N/A | N/A | N/A | N/A | N/A |
| PA0102 | PA0102 | PA0102 | PA0102 | PA0102 | PA0102 | PA0102 | PA0102 | N/A | PA0102 | PA0102 | PA0102 | PA0102 | PA0102 | N/A | PA0102 | N/A | PA0102 | PA0102 |
| PA0105 | N/A | N/A | N/A | N/A | N/A | N/A | N/A | N/A | N/A | N/A | N/A | N/A | N/A | N/A | N/A | N/A | N/A | N/A |
| PA0106 | N/A | N/A | N/A | N/A | N/A | PA0106 | N/A | N/A | N/A | N/A | N/A | N/A | PA0106 | N/A | N/A | N/A | N/A | N/A |
| PA0107 | N/A | N/A | N/A | N/A | N/A | N/A | N/A | N/A | N/A | N/A | N/A | N/A | N/A | PA0107 | N/A | N/A | N/A | N/A |
| N/A | N/A | N/A | N/A | N/A | N/A | PA0108 | N/A | N/A | N/A | N/A | N/A | N/A | N/A | N/A | N/A | N/A | N/A | N/A |
| PA0110 | N/A | N/A | N/A | N/A | N/A | N/A | N/A | N/A | N/A | N/A | N/A | N/A | N/A | N/A | N/A | N/A | N/A | N/A |
| PA0111 | N/A | N/A | N/A | N/A | N/A | N/A | N/A | N/A | N/A | N/A | N/A | N/A | N/A | N/A | N/A | N/A | N/A | N/A |
| N/A | N/A | N/A | PA0112 | N/A | N/A | N/A | N/A | N/A | N/A | N/A | N/A | N/A | N/A | N/A | N/A | N/A | N/A | N/A |
| PA0114 | N/A | PA0114 | PA0114 | PA0114 | PA0114 | PA0114 | PA0114 | N/A | PA0114 | PA0114 | PA0114 | PA0114 | PA0114 | PA0114 | N/A | N/A | PA0114 | N/A |
| PA0115 | N/A | N/A | N/A | PA0115 | PA0115 | N/A | N/A | N/A | PA0115 | PA0115 | PA0115 | PA0115 | PA0115 | N/A | N/A | N/A | N/A | N/A |
| PA0116 | N/A | N/A | N/A | N/A | N/A | N/A | N/A | N/A | N/A | N/A | N/A | PA0116 | PA0116 | N/A | N/A | N/A | N/A | N/A |
| N/A | N/A | N/A | N/A | N/A | PA0117 | N/A | N/A | N/A | N/A | N/A | N/A | N/A | N/A | N/A | N/A | N/A | N/A | N/A |
| PA0118 | N/A | PA0118 | PA0118 | PA0118 | PA0118 | N/A | PA0118 | N/A | PA0118 | N/A | PA0118 | PA0118 | PA0118 | N/A | N/A | N/A | PA0118 | N/A |
| PA0120 | N/A | N/A | N/A | N/A | N/A | N/A | PA0120 | N/A | N/A | PA0120 | N/A | PA0120 | N/A | N/A | N/A | N/A | N/A | N/A |
| N/A | N/A | N/A | N/A | N/A | PA0121 | N/A | N/A | N/A | N/A | N/A | N/A | N/A | N/A | N/A | N/A | N/A | N/A | N/A |
| PA0122 | N/A | N/A | N/A | N/A | PA0122 | PA0122 | N/A | N/A | PA0122 | PA0122 | PA0122 | PA0122 | PA0122 | N/A | N/A | PA0122 | N/A | N/A |
| PA0123 | N/A | N/A | N/A | PA0123 | N/A | N/A | N/A | N/A | N/A | N/A | PA0123 | N/A | N/A | N/A | N/A | N/A | N/A | N/A |

| | | | | | | | | | | | | | | | | | | |
|---|---|---|---|---|---|---|---|---|---|---|---|---|---|---|---|---|---|---|
| PA0124 | PA0124 | N/A | N/A | PA0124 | N/A | N/A | N/A | N/A | PA0124 | N/A | PA0124 | PA0124 | PA0124 | N/A | N/A | N/A | N/A | N/A |
| PA0125 | N/A | N/A | N/A | PA0125 | N/A | N/A | N/A | N/A | PA0125 | N/A | PA0125 | N/A | N/A | N/A | N/A | N/A | N/A | N/A |
| PA0126 | PA0126 | PA0126 | PA0126 | PA0126 | N/A | N/A | PA0126 | N/A | PA0126 | PA0126 | PA0126 | PA0126 | PA0126 | PA0126 | PA0126 | PA0126 | N/A | PA0126 |
| PA0127 | PA0127 | N/A | N/A | PA0127 | PA0127 | PA0127 | N/A | N/A | N/A | PA0127 | PA0127 | PA0127 | PA0127 | PA0127 | N/A | N/A | N/A | N/A |
| PA0128 | PA0128 | PA0128 | PA0128 | PA0128 | PA0128 | PA0128 | PA0128 | N/A | PA0128 | PA0128 | PA0128 | PA0128 | PA0128 | N/A | N/A | N/A | N/A | N/A |
| N/A | N/A | N/A | N/A | PA0129 | PA0129 | N/A | N/A | N/A | PA0129 | N/A | PA0129 | PA0129 | N/A | PA0129 | N/A | N/A | N/A | N/A |
| PA0130 | PA0130 | PA0130 | PA0130 | PA0130 | PA0130 | PA0130 | PA0130 | PA0130 | PA0130 | PA0130 | PA0130 | PA0130 | PA0130 | PA0130 | PA0130 | PA0130 | N/A | N/A |
| N/A | N/A | N/A | N/A | N/A | PA0131 | N/A | N/A | N/A | N/A | N/A | N/A | N/A | N/A | N/A | N/A | N/A | N/A | N/A |
| PA0132 | PA0132 | PA0132 | PA0132 | PA0132 | N/A | PA0132 | PA0132 | N/A | PA0132 | PA0132 | PA0132 | PA0132 | PA0132 | PA0132 | PA0132 | N/A | N/A | N/A |
| N/A | N/A | N/A | N/A | PA0133 | N/A | N/A | N/A | N/A | N/A | N/A | PA0133 | N/A | N/A | N/A | N/A | N/A | N/A | N/A |
| PA0134 | N/A | N/A | N/A | PA0134 | N/A | N/A | N/A | N/A | N/A | N/A | N/A | N/A | PA0134 | N/A | N/A | N/A | N/A | N/A |
| N/A | N/A | N/A | N/A | N/A | PA0138 | N/A | N/A | N/A | N/A | N/A | N/A | N/A | N/A | N/A | N/A | N/A | N/A | N/A |
| PA0139 | PA0139 | PA0139 | PA0139 | PA0139 | PA0139 | PA0139 | PA0139 | PA0139 | PA0139 | PA0139 | PA0139 | PA0139 | PA0139 | PA0139 | PA0139 | PA0139 | PA0139 | PA0139 |
| PA0140 | PA0140 | PA0140 | PA0140 | PA0140 | PA0140 | PA0140 | PA0140 | PA0140 | PA0140 | PA0140 | PA0140 | PA0140 | PA0140 | N/A | PA0140 | PA0140 | N/A | N/A |
| PA0141 | PA0141 | N/A | N/A | PA0141 | PA0141 | PA0141 | PA0141 | PA0141 | N/A | PA0141 | PA0141 | PA0141 | PA0141 | PA0141 | N/A | N/A | PA0141 | PA0141 |
| PA0142 | N/A | N/A | N/A | PA0142 | PA0142 | N/A | N/A | N/A | N/A | N/A | N/A | PA0142 | N/A | N/A | N/A | N/A | N/A | N/A |
| PA0143 | PA0143 | PA0143 | PA0143 | PA0143 | PA0143 | PA0143 | PA0143 | N/A | PA0143 | PA0143 | N/A | PA0143 | PA0143 | PA0143 | PA0143 | PA0143 | N/A | N/A |
| PA0144 | N/A | N/A | N/A | N/A | N/A | N/A | N/A | N/A | PA0144 | N/A | N/A | N/A | PA0144 | N/A | N/A | N/A | N/A | N/A |
| N/A | N/A | N/A | N/A | N/A | PA0147 | N/A | N/A | N/A | N/A | N/A | N/A | N/A | N/A | N/A | N/A | N/A | N/A | N/A |
| PA0148 | PA0148 | PA0148 | PA0148 | PA0148 | PA0148 | N/A | PA0148 | N/A | PA0148 | N/A | PA0148 | PA0148 | PA0148 | N/A | N/A | PA0148 | N/A | N/A |
| PA0150 | N/A | N/A | N/A | PA0150 | N/A | N/A | N/A | N/A | N/A | N/A | N/A | N/A | N/A | N/A | N/A | N/A | N/A | N/A |
| PA0151 | N/A | N/A | N/A | PA0151 | N/A | N/A | N/A | N/A | N/A | N/A | N/A | N/A | N/A | N/A | N/A | N/A | N/A | N/A |
| PA0152 | N/A | N/A | N/A | PA0152 | N/A | N/A | N/A | N/A | N/A | N/A | N/A | N/A | N/A | N/A | N/A | N/A | N/A | N/A |
| N/A | N/A | N/A | N/A | N/A | N/A | PA0153 | N/A | N/A | N/A | N/A | N/A | N/A | N/A | N/A | N/A | N/A | N/A | N/A |
| PA0155 | PA0155 | PA0155 | N/A | PA0155 | N/A | PA0155 | N/A | N/A | PA0155 | PA0155 | PA0155 | PA0155 | PA0155 | N/A | N/A | N/A | N/A | N/A |
| PA0156 | N/A | N/A | N/A | PA0156 | N/A | N/A | PA0156 | PA0156 | N/A | N/A | N/A | N/A | PA0156 | PA0156 | PA0156 | N/A | PA0156 | N/A |
| PA0157 | N/A | N/A | N/A | PA0157 | N/A | PA0157 | PA0157 | N/A | N/A | PA0157 | N/A | N/A | PA0157 | PA0157 | N/A | N/A | PA0157 | N/A |
| PA0158 | N/A | N/A | N/A | PA0158 | N/A | PA0158 | N/A | N/A | PA0158 | PA0158 | PA0158 | PA0158 | PA0158 | PA0158 | N/A | N/A | PA0158 | N/A |
| PA0159 | N/A | N/A | N/A | PA0159 | N/A | N/A | PA0159 | N/A | PA0159 | PA0159 | PA0159 | N/A | PA0159 | N/A | N/A | N/A | N/A | N/A |
| N/A | N/A | N/A | N/A | N/A | PA0161 | N/A | N/A | N/A | N/A | N/A | N/A | N/A | N/A | N/A | N/A | N/A | N/A | N/A |
| PA0162 | PA0162 | PA0162 | PA0162 | PA0162 | PA0162 | PA0162 | PA0162 | PA0162 | PA0162 | PA0162 | PA0162 | PA0162 | PA0162 | N/A | PA0162 | PA0162 | N/A | N/A |
| PA0163 | N/A | N/A | N/A | PA0163 | N/A | N/A | N/A | N/A | N/A | N/A | N/A | N/A | N/A | N/A | N/A | N/A | N/A | N/A |
| N/A | N/A | N/A | N/A | N/A | PA0164 | N/A | N/A | N/A | N/A | N/A | N/A | N/A | N/A | N/A | N/A | N/A | N/A | N/A |
| PA0165 | PA0165 | PA0165 | N/A | PA0165 | N/A | PA0165 | PA0165 | PA0165 | PA0165 | PA0165 | PA0165 | PA0165 | PA0165 | N/A | PA0165 | PA0165 | N/A | N/A |
| PA0167 | N/A | PA0167 | N/A | PA0167 | N/A | N/A | N/A | N/A | PA0167 | PA0167 | PA0167 | PA0167 | N/A | PA0167 | N/A | N/A | N/A | N/A |
| N/A | N/A | N/A | N/A | PA0168 | N/A | N/A | N/A | N/A | N/A | N/A | N/A | N/A | N/A | N/A | N/A | N/A | N/A | N/A |
| PA0169 | N/A | N/A | N/A | PA0169 | PA0169 | N/A | N/A | N/A | N/A | N/A | N/A | PA0169 | N/A | N/A | N/A | N/A | PA0169 | N/A |
| PA0170 | PA0170 | PA0170 | PA0170 | PA0170 | PA0170 | N/A | N/A | N/A | PA0170 | PA0170 | PA0170 | PA0170 | N/A | N/A | PA0170 | N/A | N/A | N/A |
| PA0171 | N/A | PA0171 | PA0171 | PA0171 | N/A | N/A | N/A | N/A | PA0171 | PA0171 | PA0171 | PA0171 | N/A | N/A | N/A | N/A | PA0171 | N/A |
| PA0172 | N/A | N/A | N/A | N/A | N/A | N/A | N/A | N/A | N/A | N/A | N/A | N/A | N/A | PA0172 | N/A | N/A | PA0172 | N/A |
| PA0173 | N/A | N/A | N/A | N/A | N/A | N/A | N/A | N/A | N/A | N/A | N/A | PA0173 | PA0173 | N/A | N/A | N/A | N/A | N/A |
| PA0174 | N/A | N/A | N/A | N/A | N/A | N/A | N/A | N/A | N/A | N/A | N/A | PA0174 | N/A | N/A | N/A | N/A | N/A | N/A |
| N/A | N/A | N/A | N/A | N/A | PA0175 | N/A | N/A | N/A | N/A | N/A | N/A | PA0175 | N/A | PA0175 | N/A | N/A | N/A | N/A |
| PA0176 | N/A | N/A | N/A | N/A | PA0176 | PA0176 | N/A | N/A | N/A | PA0176 | PA0176 | PA0176 | PA0176 | PA0176 | N/A | N/A | PA0176 | PA0176 |
| PA0177 | N/A | N/A | N/A | N/A | N/A | N/A | N/A | N/A | N/A | N/A | PA0177 | PA0177 | PA0177 | N/A | N/A | N/A | N/A | N/A |
| PA0178 | N/A | N/A | N/A | PA0178 | N/A | N/A | N/A | N/A | N/A | N/A | N/A | PA0178 | PA0178 | N/A | N/A | N/A | N/A | N/A |
| PA0179 | N/A | N/A | N/A | N/A | N/A | N/A | N/A | N/A | PA0179 | N/A | PA0179 | PA0179 | PA0179 | N/A | N/A | N/A | PA0179 | N/A |
| PA0180 | PA0180 | PA0180 | PA0180 | PA0180 | N/A | PA0180 | PA0180 | PA0180 | PA0180 | PA0180 | PA0180 | PA0180 | PA0180 | PA0180 | PA0180 | N/A | PA0180 | PA0180 |
| PA0181 | N/A | N/A | N/A | N/A | N/A | N/A | N/A | N/A | PA0181 | N/A | PA0181 | PA0181 | PA0181 | N/A | N/A | N/A | N/A | N/A |
| PA0182 | N/A | N/A | N/A | PA0182 | N/A | PA0182 | N/A | N/A | N/A | N/A | PA0182 | PA0182 | N/A | N/A | N/A | N/A | N/A | N/A |
| PA0183 | N/A | N/A | N/A | N/A | N/A | N/A | N/A | N/A | N/A | N/A | N/A | N/A | N/A | N/A | N/A | N/A | N/A | N/A |
| N/A | N/A | N/A | N/A | N/A | N/A | N/A | N/A | N/A | N/A | N/A | PA0184 | N/A | N/A | N/A | N/A | N/A | N/A | N/A |
| PA0195 | N/A | N/A | N/A | PA0195 | PA0195 | PA0195 | N/A | N/A | N/A | PA0195 | PA0195 | PA0195 | PA0195 | N/A | PA0195 | N/A | N/A | N/A |
| PA0196 | N/A | N/A | N/A | N/A | N/A | N/A | N/A | N/A | N/A | N/A | PA0196 | PA0196 | PA0196 | PA0196 | N/A | N/A | N/A | N/A |
| PA0198 | N/A | N/A | N/A | N/A | N/A | N/A | N/A | N/A | N/A | N/A | N/A | N/A | N/A | N/A | N/A | N/A | N/A | N/A |
| PA0207 | N/A | N/A | N/A | N/A | N/A | N/A | N/A | N/A | N/A | N/A | N/A | N/A | N/A | N/A | N/A | N/A | N/A | N/A |
| PA0208 | N/A | N/A | N/A | N/A | N/A | N/A | N/A | N/A | N/A | N/A | N/A | N/A | N/A | N/A | N/A | N/A | N/A | N/A |
| N/A | N/A | N/A | N/A | N/A | N/A | N/A | N/A | N/A | N/A | N/A | N/A | N/A | N/A | PA0209 | N/A | N/A | N/A | N/A |
| N/A | N/A | N/A | N/A | N/A | N/A | N/A | N/A | N/A | N/A | N/A | N/A | N/A | PA0210 | N/A | N/A | N/A | N/A | N/A |
| PA0211 | N/A | N/A | N/A | N/A | N/A | N/A | N/A | N/A | N/A | N/A | N/A | N/A | PA0211 | N/A | N/A | N/A | N/A | N/A |
| PA0212 | N/A | N/A | N/A | N/A | N/A | N/A | N/A | N/A | N/A | N/A | N/A | N/A | N/A | N/A | N/A | N/A | N/A | N/A |
| PA0214 | N/A | N/A | N/A | N/A | N/A | N/A | N/A | N/A | N/A | N/A | N/A | N/A | N/A | N/A | N/A | N/A | N/A | N/A |
| PA0217 | N/A | N/A | N/A | N/A | N/A | N/A | N/A | N/A | N/A | N/A | N/A | N/A | N/A | N/A | N/A | N/A | N/A | N/A |
| N/A | N/A | N/A | N/A | PA0222 | N/A | N/A | N/A | N/A | N/A | N/A | PA0222 | N/A | PA0222 | N/A | PA0222 | N/A | N/A | N/A |
| N/A | N/A | N/A | N/A | PA0223 | N/A | N/A | N/A | N/A | N/A | N/A | N/A | N/A | N/A | N/A | N/A | N/A | N/A | N/A |
| N/A | N/A | N/A | N/A | N/A | PA0224 | N/A | N/A | N/A | N/A | N/A | N/A | N/A | N/A | N/A | N/A | N/A | N/A | N/A |
| PA0225 | N/A | N/A | N/A | N/A | PA0225 | N/A | N/A | N/A | N/A | N/A | N/A | N/A | N/A | N/A | N/A | N/A | N/A | N/A |
| N/A | N/A | N/A | N/A | N/A | N/A | PA0227 | N/A | N/A | N/A | N/A | N/A | N/A | N/A | N/A | N/A | N/A | N/A | N/A |
| PA0229 | N/A | N/A | N/A | N/A | N/A | N/A | N/A | N/A | N/A | N/A | N/A | N/A | N/A | N/A | N/A | N/A | N/A | N/A |
| PA0230 | N/A | N/A | N/A | PA0230 | PA0230 | PA0230 | N/A | N/A | N/A | PA0230 | N/A | PA0230 | PA0230 | N/A | N/A | N/A | N/A | N/A |
| PA0231 | N/A | N/A | N/A | PA0231 | PA0231 | N/A | N/A | N/A | PA0231 | PA0231 | PA0231 | PA0231 | PA0231 | N/A | N/A | N/A | N/A | N/A |
| PA0232 | N/A | N/A | N/A | N/A | N/A | N/A | N/A | N/A | N/A | N/A | N/A | N/A | PA0232 | N/A | N/A | N/A | N/A | N/A |
| PA0233 | N/A | N/A | N/A | N/A | N/A | N/A | N/A | N/A | N/A | N/A | N/A | N/A | N/A | N/A | N/A | N/A | N/A | N/A |
| PA0236 | PA0236 | N/A | N/A | PA0236 | N/A | PA0236 | PA0236 | PA0236 | PA0236 | PA0236 | PA0236 | PA0236 | PA0236 | PA0236 | N/A | N/A | PA0236 | N/A |
| PA0243 | N/A | N/A | N/A | PA0243 | N/A | N/A | N/A | N/A | PA0243 | N/A | PA0243 | PA0243 | PA0243 | N/A | N/A | N/A | N/A | N/A |
| N/A | N/A | N/A | N/A | N/A | N/A | N/A | N/A | N/A | N/A | N/A | N/A | N/A | N/A | PA0246 | N/A | N/A | N/A | N/A |
| PA0248 | N/A | N/A | N/A | N/A | N/A | N/A | N/A | N/A | N/A | N/A | N/A | N/A | N/A | N/A | N/A | N/A | N/A | N/A |
| N/A | N/A | N/A | N/A | N/A | PA0249 | N/A | N/A | N/A | N/A | N/A | N/A | N/A | N/A | N/A | N/A | N/A | N/A | N/A |
| PA0250 | N/A | N/A | N/A | PA0250 | PA0250 | N/A | N/A | N/A | PA0250 | PA0250 | PA0250 | PA0250 | PA0250 | N/A | N/A | N/A | N/A | N/A |
| N/A | N/A | N/A | N/A | PA0253 | N/A | N/A | PA0253 | N/A | PA0253 | PA0253 | PA0253 | PA0253 | PA0253 | N/A | N/A | N/A | N/A | N/A |
| PA0255 | N/A | N/A | N/A | N/A | N/A | N/A | N/A | N/A | N/A | PA0255 | N/A | N/A | N/A | N/A | N/A | N/A | N/A | N/A |
| PA0259 | N/A | N/A | N/A | PA0259 | N/A | N/A | N/A | PA0259 | PA0259 | N/A | PA0259 | PA0259 | PA0259 | N/A | N/A | N/A | N/A | N/A |
| N/A | N/A | N/A | N/A | N/A | PA0260 | N/A | N/A | N/A | N/A | N/A | N/A | N/A | N/A | N/A | N/A | N/A | N/A | N/A |
| PA0261 | N/A | N/A | N/A | PA0261 | N/A | N/A | N/A | N/A | PA0261 | PA0261 | PA0261 | N/A | PA0261 | N/A | PA0261 | N/A | PA0261 | N/A |
| N/A | N/A | N/A | N/A | PA0262 | N/A | N/A | N/A | N/A | N/A | N/A | N/A | N/A | N/A | N/A | N/A | N/A | PA0262 | N/A |
| PA0263 | N/A | N/A | N/A | PA0263 | N/A | PA0263 | N/A | N/A | PA0263 | PA0263 | N/A | PA0263 | PA0263 | N/A | N/A | N/A | N/A | N/A |
| N/A | N/A | N/A | N/A | N/A | PA0264 | N/A | N/A | N/A | N/A | N/A | N/A | N/A | N/A | N/A | N/A | N/A | N/A | N/A |
| PA0265 | PA0265 | PA0265 | PA0265 | PA0265 | PA0265 | PA0265 | PA0265 | N/A | PA0265 | PA0265 | PA0265 | PA0265 | PA0265 | PA0265 | PA0265 | PA0265 | N/A | N/A |
| PA0266 | PA0266 | PA0266 | PA0266 | PA0266 | PA0266 | PA0266 | PA0266 | N/A | PA0266 | PA0266 | PA0266 | PA0266 | PA0266 | PA0266 | PA0266 | PA0266 | N/A | N/A |
| PA0267 | N/A | N/A | N/A | PA0267 | N/A | N/A | N/A | PA0267 | N/A | N/A | N/A | N/A | N/A | N/A | N/A | N/A | N/A | N/A |
| PA0268 | N/A | N/A | N/A | N/A | PA0268 | N/A | N/A | N/A | N/A | N/A | N/A | N/A | N/A | N/A | N/A | N/A | N/A | N/A |
| PA0269 | N/A | N/A | N/A | PA0269 | PA0269 | PA0269 | N/A | N/A | PA0269 | PA0269 | PA0269 | PA0269 | PA0269 | N/A | N/A | N/A | N/A | N/A |
| PA0270 | PA0270 | N/A | N/A | N/A | PA0270 | N/A | N/A | N/A | N/A | N/A | PA0270 | N/A | PA0270 | N/A | N/A | N/A | N/A | N/A |
| N/A | N/A | N/A | N/A | N/A | N/A | N/A | N/A | N/A | N/A | N/A | N/A | N/A | PA0271 | N/A | N/A | N/A | N/A | N/A |
| N/A | N/A | N/A | N/A | N/A | PA0276 | N/A | N/A | N/A | N/A | PA0276 | N/A | N/A | N/A | N/A | N/A | N/A | N/A | N/A |
| PA0277 | PA0277 | PA0277 | PA0277 | PA0277 | PA0277 | N/A | PA0277 | PA0277 | PA0277 | PA0277 | PA0277 | PA0277 | PA0277 | PA0277 | PA0277 | N/A | PA0277 | PA0277 |
| PA0279 | N/A | N/A | N/A | N/A | N/A | N/A | N/A | N/A | N/A | N/A | N/A | N/A | N/A | N/A | N/A | N/A | N/A | N/A |
| PA0280 | PA0280 | PA0280 | PA0280 | PA0280 | N/A | PA0280 | PA0280 | PA0280 | PA0280 | PA0280 | PA0280 | PA0280 | PA0280 | PA0280 | N/A | N/A | PA0280 | N/A |
| PA0281 | N/A | N/A | N/A | PA0281 | N/A | N/A | N/A | N/A | PA0281 | N/A | PA0281 | PA0281 | PA0281 | PA0281 | N/A | N/A | PA0281 | N/A |
| PA0282 | N/A | N/A | N/A | PA0282 | PA0282 | PA0282 | N/A | N/A | N/A | N/A | PA0282 | PA0282 | N/A | PA0282 | N/A | N/A | PA0282 | N/A |
| PA0283 | N/A | N/A | N/A | N/A | PA0283 | PA0283 | N/A | N/A | PA0283 | PA0283 | PA0283 | PA0283 | PA0283 | N/A | N/A | PA0283 | N/A | N/A |
| PA0284 | N/A | N/A | PA0284 | PA0284 | PA0284 | N/A | N/A | N/A | PA0284 | PA0284 | PA0284 | PA0284 | N/A | N/A | N/A | N/A | N/A | N/A |
| PA0285 | PA0285 | PA0285 | PA0285 | PA0285 | N/A | N/A | N/A | PA0285 | PA0285 | N/A | PA0285 | PA0285 | N/A | PA0285 | PA0285 | N/A | N/A | N/A |
| N/A | N/A | N/A | N/A | N/A | N/A | N/A | N/A | N/A | N/A | N/A | N/A | N/A | N/A | PA0286 | N/A | N/A | N/A | N/A |
| PA0290 | N/A | N/A | N/A | N/A | PA0290 | N/A | N/A | N/A | N/A | N/A | N/A | N/A | N/A | N/A | N/A | N/A | N/A | N/A |
| PA0291 | PA0291 | PA0291 | PA0291 | PA0291 | PA0291 | PA0291 | PA0291 | PA0291 | PA0291 | PA0291 | PA0291 | PA0291 | PA0291 | PA0291 | PA0291 | PA0291 | PA0291 | N/A |
| PA0292 | PA0292 | PA0292 | PA0292 | PA0292 | PA0292 | PA0292 | N/A | N/A | PA0292 | PA0292 | PA0292 | PA0292 | PA0292 | PA0292 | N/A | PA0292 | N/A | N/A |
| PA0293 | PA0293 | N/A | N/A | PA0293 | PA0293 | N/A | PA0293 | N/A | PA0293 | N/A | PA0293 | PA0293 | PA0293 | PA0293 | PA0293 | N/A | N/A | N/A |
| PA0294 | N/A | N/A | N/A | PA0294 | PA0294 | N/A | N/A | N/A | PA0294 | PA0294 | N/A | PA0294 | PA0294 | N/A | N/A | N/A | N/A | N/A |

| | | | | | | | | | | | | | | | | | | |
|---|---|---|---|---|---|---|---|---|---|---|---|---|---|---|---|---|---|---|
| PA0295 | PA0295 | N/A | N/A | PA0295 | PA0295 | N/A | PA0295 | N/A | N/A | PA0295 | PA0295 | PA0295 | PA0295 | PA0295 | N/A | N/A | N/A | N/A |
| PA0296 | PA0296 | PA0296 | PA0296 | PA0296 | N/A | PA0296 | PA0296 | PA0296 | PA0296 | PA0296 | PA0296 | PA0296 | PA0296 | N/A | PA0296 | N/A | N/A | N/A |
| PA0297 | N/A | N/A | N/A | PA0297 | PA0297 | N/A | N/A | N/A | PA0297 | PA0297 | PA0297 | PA0297 | PA0297 | N/A | N/A | N/A | N/A | N/A |
| PA0298 | PA0298 | PA0298 | PA0298 | PA0298 | PA0298 | PA0298 | PA0298 | PA0298 | PA0298 | PA0298 | PA0298 | PA0298 | PA0298 | PA0298 | N/A | N/A | PA0298 | N/A |
| PA0299 | PA0299 | PA0299 | PA0299 | PA0299 | PA0299 | PA0299 | N/A | PA0299 | PA0299 | PA0299 | PA0299 | PA0299 | PA0299 | PA0299 | PA0299 | N/A | N/A | N/A |
| PA0300 | PA0300 | PA0300 | PA0300 | PA0300 | PA0300 | PA0300 | PA0300 | PA0300 | PA0300 | PA0300 | PA0300 | PA0300 | PA0300 | PA0300 | PA0300 | PA0300 | N/A | N/A |
| PA0301 | N/A | PA0301 | N/A | PA0301 | PA0301 | PA0301 | PA0301 | PA0301 | PA0301 | PA0301 | PA0301 | PA0301 | PA0301 | PA0301 | PA0301 | PA0301 | N/A | N/A |
| PA0302 | PA0302 | PA0302 | PA0302 | PA0302 | N/A | PA0302 | N/A | PA0302 | PA0302 | PA0302 | PA0302 | PA0302 | PA0302 | PA0302 | PA0302 | N/A | PA0302 | N/A |
| PA0303 | PA0303 | N/A | PA0303 | PA0303 | N/A | PA0303 | N/A | N/A | PA0303 | PA0303 | PA0303 | PA0303 | PA0303 | PA0303 | N/A | N/A | PA0303 | N/A |
| N/A | N/A | N/A | N/A | PA0304 | N/A | N/A | N/A | N/A | N/A | N/A | N/A | N/A | N/A | PA0304 | N/A | N/A | PA0304 | N/A |
| PA0305 | N/A | N/A | N/A | PA0305 | N/A | N/A | PA0305 | N/A | PA0305 | N/A | N/A | N/A | N/A | PA0305 | N/A | N/A | PA0305 | N/A |
| PA0306.1 | N/A | PA0306.1 | PA0306.1 | PA0306.1 | N/A | PA0306.1 | PA0306.1 | N/A | PA0306.1 | PA0306.1 | PA0306.1 | PA0306.1 | PA0306.1 | PA0306.1 | N/A | N/A | PA0306.1 | PA0306.1 |
| PA0307 | N/A | N/A | N/A | N/A | N/A | N/A | N/A | N/A | N/A | PA0307 | N/A | PA0307 | N/A | PA0307 | N/A | N/A | PA0307 | N/A |
| PA0308 | N/A | N/A | N/A | PA0308 | PA0308 | N/A | PA0308 | N/A | N/A | PA0308 | PA0308 | PA0308 | N/A | PA0308 | N/A | N/A | N/A | N/A |
| PA0309 | PA0309 | N/A | PA0309 | PA0309 | PA0309 | N/A | PA0309 | N/A | PA0309 | PA0309 | PA0309 | PA0309 | PA0309 | N/A | PA0309 | N/A | N/A | N/A |
| PA0310 | N/A | N/A | N/A | PA0310 | N/A | N/A | N/A | N/A | PA0310 | N/A | N/A | N/A | N/A | N/A | N/A | N/A | N/A | N/A |
| PA0312 | N/A | N/A | N/A | N/A | N/A | N/A | N/A | N/A | N/A | PA0312 | PA0312 | PA0312 | PA0312 | N/A | N/A | N/A | N/A | N/A |
| PA0313 | N/A | N/A | N/A | PA0313 | PA0313 | N/A | N/A | N/A | PA0313 | PA0313 | PA0313 | PA0313 | N/A | PA0313 | N/A | N/A | N/A | N/A |
| PA0314 | N/A | PA0314 | N/A | PA0314 | PA0314 | PA0314 | PA0314 | N/A | PA0314 | PA0314 | PA0314 | PA0314 | PA0314 | N/A | PA0314 | PA0314 | N/A | N/A |
| PA0315 | PA0315 | PA0315 | PA0315 | PA0315 | PA0315 | PA0315 | N/A | N/A | PA0315 | PA0315 | PA0315 | PA0315 | PA0315 | N/A | PA0315 | N/A | N/A | N/A |
| PA0316 | PA0316 | PA0316 | PA0316 | PA0316 | PA0316 | PA0316 | PA0316 | PA0316 | PA0316 | PA0316 | PA0316 | PA0316 | PA0316 | PA0316 | PA0316 | N/A | N/A | N/A |
| PA0317 | PA0317 | PA0317 | PA0317 | PA0317 | PA0317 | PA0317 | N/A | N/A | PA0317 | PA0317 | PA0317 | PA0317 | PA0317 | N/A | PA0317 | PA0317 | N/A | N/A |
| PA0318 | PA0318 | PA0318 | PA0318 | PA0318 | PA0318 | PA0318 | PA0318 | PA0318 | PA0318 | PA0318 | PA0318 | PA0318 | PA0318 | N/A | PA0318 | N/A | N/A | N/A |
| PA0319 | N/A | N/A | N/A | PA0319 | N/A | N/A | PA0319 | N/A | PA0319 | N/A | PA0319 | N/A | N/A | PA0319 | N/A | N/A | PA0319 | N/A |
| N/A | N/A | N/A | N/A | N/A | PA0322 | N/A | N/A | N/A | N/A | N/A | N/A | N/A | N/A | N/A | N/A | N/A | N/A | N/A |
| PA0323 | N/A | N/A | N/A | N/A | PA0323 | N/A | N/A | N/A | N/A | N/A | N/A | N/A | N/A | N/A | N/A | N/A | N/A | N/A |
| PA0327 | N/A | N/A | N/A | N/A | N/A | N/A | N/A | N/A | N/A | N/A | N/A | N/A | N/A | PA0327 | N/A | N/A | N/A | N/A |
| PA0328 | PA0328 | N/A | N/A | PA0328 | PA0328 | PA0328 | PA0328 | PA0328 | PA0328 | PA0328 | PA0328 | PA0328 | PA0328 | PA0328 | N/A | N/A | N/A | N/A |
| PA0329 | PA0329 | PA0329 | PA0329 | PA0329 | PA0329 | PA0329 | PA0329 | N/A | PA0329 | PA0329 | PA0329 | PA0329 | PA0329 | PA0329 | PA0329 | N/A | N/A | N/A |
| PA0330 | PA0330 | PA0330 | PA0330 | PA0330 | PA0330 | PA0330 | PA0330 | N/A | PA0330 | PA0330 | PA0330 | PA0330 | PA0330 | N/A | PA0330 | N/A | N/A | N/A |
| PA0331 | PA0331 | PA0331 | PA0331 | PA0331 | PA0331 | PA0331 | N/A | N/A | PA0331 | PA0331 | PA0331 | PA0331 | PA0331 | PA0331 | N/A | PA0331 | N/A | N/A |
| PA0332 | N/A | N/A | N/A | N/A | N/A | PA0332 | N/A | N/A | PA0332 | PA0332 | PA0332 | PA0332 | PA0332 | PA0332 | N/A | N/A | PA0332 | N/A |
| PA0333 | N/A | N/A | N/A | PA0333 | N/A | N/A | N/A | N/A | N/A | N/A | N/A | N/A | N/A | PA0333 | N/A | N/A | N/A | N/A |
| PA0334 | N/A | N/A | N/A | N/A | PA0334 | N/A | N/A | N/A | N/A | N/A | N/A | N/A | N/A | N/A | N/A | N/A | N/A | N/A |
| PA0335 | PA0335 | PA0335 | PA0335 | PA0335 | PA0335 | N/A | PA0335 | N/A | PA0335 | PA0335 | PA0335 | PA0335 | PA0335 | N/A | PA0335 | N/A | N/A | N/A |
| PA0336 | N/A | N/A | N/A | PA0336 | N/A | PA0336 | PA0336 | N/A | PA0336 | PA0336 | PA0336 | PA0336 | N/A | PA0336 | N/A | N/A | PA0336 | N/A |
| PA0337 | PA0337 | PA0337 | PA0337 | PA0337 | PA0337 | PA0337 | N/A | PA0337 | PA0337 | PA0337 | PA0337 | PA0337 | PA0337 | N/A | N/A | N/A | N/A | N/A |
| PA0339 | N/A | N/A | N/A | N/A | N/A | N/A | N/A | N/A | PA0339 | N/A | PA0339 | PA0339 | N/A | N/A | N/A | N/A | N/A | N/A |
| PA0341 | PA0341 | PA0341 | PA0341 | PA0341 | PA0341 | PA0341 | N/A | PA0341 | PA0341 | PA0341 | PA0341 | PA0341 | PA0341 | PA0341 | N/A | N/A | PA0341 | N/A |
| PA0342 | PA0342 | PA0342 | PA0342 | PA0342 | PA0342 | PA0342 | N/A | N/A | PA0342 | PA0342 | PA0342 | PA0342 | N/A | N/A | N/A | N/A | N/A | N/A |
| PA0344 | N/A | N/A | N/A | N/A | N/A | N/A | N/A | N/A | N/A | N/A | N/A | PA0344 | PA0344 | PA0344 | N/A | N/A | N/A | N/A |
| PA0345 | N/A | N/A | N/A | PA0345 | N/A | PA0345 | N/A | N/A | N/A | N/A | N/A | N/A | N/A | PA0345 | N/A | N/A | PA0345 | N/A |
| N/A | N/A | N/A | N/A | N/A | PA0349 | N/A | N/A | N/A | N/A | N/A | N/A | N/A | N/A | N/A | N/A | N/A | N/A | N/A |
| PA0350 | PA0350 | PA0350 | PA0350 | PA0350 | PA0350 | N/A | PA0350 | N/A | PA0350 | PA0350 | PA0350 | PA0350 | PA0350 | N/A | PA0350 | N/A | N/A | N/A |
| N/A | N/A | N/A | N/A | PA0351 | N/A | N/A | N/A | N/A | N/A | N/A | N/A | N/A | N/A | N/A | N/A | N/A | N/A | N/A |
| PA0352 | N/A | N/A | N/A | N/A | PA0352 | N/A | N/A | N/A | N/A | PA0352 | N/A | PA0352 | N/A | PA0352 | N/A | N/A | PA0352 | N/A |
| PA0353 | PA0353 | PA0353 | PA0353 | PA0353 | PA0353 | PA0353 | PA0353 | N/A | PA0353 | PA0353 | PA0353 | PA0353 | PA0353 | PA0353 | PA0353 | PA0353 | N/A | N/A |
| PA0354 | PA0354 | PA0354 | PA0354 | PA0354 | N/A | PA0354 | PA0354 | PA0354 | PA0354 | PA0354 | PA0354 | PA0354 | PA0354 | PA0354 | PA0354 | N/A | N/A | N/A |
| PA0355 | N/A | N/A | N/A | N/A | PA0355 | PA0355 | N/A | N/A | PA0355 | PA0355 | PA0355 | PA0355 | PA0355 | N/A | N/A | N/A | N/A | N/A |
| PA0356 | PA0356 | PA0356 | N/A | PA0356 | N/A | N/A | PA0356 | N/A | PA0356 | PA0356 | PA0356 | N/A | N/A | N/A | N/A | N/A | N/A | N/A |
| PA0357 | PA0357 | N/A | N/A | PA0357 | N/A | PA0357 | N/A | PA0357 | PA0357 | PA0357 | PA0357 | PA0357 | PA0357 | PA0357 | N/A | N/A | PA0357 | N/A |
| N/A | N/A | N/A | N/A | PA0358 | N/A | N/A | N/A | N/A | PA0358 | N/A | N/A | N/A | N/A | N/A | N/A | N/A | N/A | N/A |
| PA0359 | PA0359 | PA0359 | N/A | PA0359 | N/A | PA0359 | N/A | N/A | PA0359 | PA0359 | PA0359 | PA0359 | PA0359 | N/A | N/A | N/A | N/A | N/A |
| N/A | N/A | N/A | N/A | N/A | PA0360 | N/A | N/A | N/A | N/A | N/A | N/A | N/A | N/A | N/A | N/A | N/A | N/A | N/A |
| PA0361 | N/A | N/A | N/A | PA0361 | PA0361 | N/A | PA0361 | PA0361 | N/A | PA0361 | N/A | N/A | PA0361 | N/A | N/A | N/A | N/A | N/A |
| PA0363 | PA0363 | PA0363 | PA0363 | PA0363 | PA0363 | PA0363 | PA0363 | N/A | PA0363 | PA0363 | PA0363 | PA0363 | PA0363 | PA0363 | N/A | N/A | PA0363 | N/A |
| PA0364 | N/A | N/A | N/A | N/A | N/A | N/A | N/A | N/A | N/A | N/A | N/A | N/A | N/A | PA0364 | N/A | N/A | PA0364 | N/A |
| PA0365 | N/A | N/A | N/A | PA0365 | PA0365 | N/A | N/A | N/A | N/A | N/A | N/A | PA0365 | N/A | N/A | N/A | N/A | N/A | N/A |
| PA0366 | N/A | N/A | N/A | PA0366 | PA0366 | N/A | N/A | N/A | PA0366 | PA0366 | N/A | PA0366 | PA0366 | PA0366 | N/A | N/A | PA0366 | N/A |
| PA0367 | N/A | N/A | N/A | N/A | N/A | N/A | N/A | N/A | N/A | N/A | N/A | PA0367 | N/A | N/A | N/A | N/A | N/A | N/A |
| PA0369 | N/A | N/A | N/A | PA0369 | N/A | N/A | N/A | N/A | N/A | N/A | N/A | N/A | N/A | N/A | N/A | N/A | N/A | N/A |
| PA0370 | PA0370 | PA0370 | PA0370 | PA0370 | N/A | N/A | PA0370 | PA0370 | PA0370 | PA0370 | PA0370 | PA0370 | PA0370 | PA0370 | N/A | N/A | N/A | N/A |
| PA0371 | PA0371 | PA0371 | PA0371 | PA0371 | PA0371 | N/A | PA0371 | PA0371 | PA0371 | PA0371 | PA0371 | PA0371 | PA0371 | PA0371 | PA0371 | PA0371 | PA0371 | N/A |
| PA0372 | PA0372 | PA0372 | PA0372 | PA0372 | PA0372 | PA0372 | PA0372 | PA0372 | PA0372 | PA0372 | PA0372 | PA0372 | PA0372 | PA0372 | PA0372 | N/A | PA0372 | PA0372 |
| PA0373 | PA0373 | PA0373 | PA0373 | PA0373 | PA0373 | PA0373 | PA0373 | PA0373 | PA0373 | PA0373 | PA0373 | PA0373 | PA0373 | PA0373 | PA0373 | N/A | PA0373 | PA0373 |
| PA0374 | PA0374 | PA0374 | PA0374 | PA0374 | N/A | N/A | PA0374 | N/A | PA0374 | PA0374 | PA0374 | PA0374 | PA0374 | PA0374 | PA0374 | N/A | PA0374 | PA0374 |
| PA0375 | PA0375 | PA0375 | PA0375 | PA0375 | N/A | PA0375 | N/A | PA0375 | PA0375 | PA0375 | PA0375 | PA0375 | PA0375 | PA0375 | PA0375 | N/A | PA0375 | N/A |
| N/A | N/A | PA0376 | N/A | N/A | N/A | N/A | N/A | N/A | N/A | N/A | N/A | N/A | N/A | N/A | N/A | N/A | N/A | N/A |
| PA0377 | N/A | N/A | N/A | N/A | N/A | N/A | N/A | N/A | N/A | N/A | N/A | N/A | N/A | PA0377 | N/A | N/A | N/A | N/A |
| PA0378 | N/A | N/A | N/A | N/A | N/A | N/A | N/A | N/A | N/A | N/A | N/A | N/A | N/A | PA0378 | PA0378 | N/A | N/A | N/A |
| N/A | N/A | N/A | N/A | N/A | PA0380 | N/A | N/A | N/A | N/A | N/A | PA0380 | N/A | PA0380 | N/A | PA0380 | N/A | N/A | N/A |
| PA0381 | PA0381 | PA0381 | PA0381 | PA0381 | PA0381 | PA0381 | PA0381 | PA0381 | PA0381 | PA0381 | PA0381 | PA0381 | PA0381 | N/A | PA0381 | N/A | PA0381 | PA0381 |
| PA0382 | PA0382 | PA0382 | PA0382 | PA0382 | PA0382 | PA0382 | PA0382 | N/A | PA0382 | PA0382 | PA0382 | PA0382 | PA0382 | PA0382 | PA0382 | N/A | N/A | N/A |
| PA0384 | N/A | N/A | N/A | N/A | N/A | N/A | N/A | N/A | N/A | N/A | PA0384 | PA0384 | PA0384 | N/A | N/A | N/A | N/A | N/A |
| PA0385 | N/A | N/A | N/A | N/A | N/A | PA0385 | N/A | N/A | PA0385 | N/A | PA0385 | N/A | N/A | N/A | N/A | N/A | N/A | N/A |
| N/A | PA0386 | PA0386 | N/A | PA0386 | PA0386 | N/A | PA0386 | PA0386 | N/A | N/A | N/A | N/A | N/A | N/A | PA0386 | N/A | N/A | N/A |
| PA0387 | PA0387 | PA0387 | PA0387 | PA0387 | PA0387 | N/A | PA0387 | N/A | PA0387 | PA0387 | PA0387 | PA0387 | PA0387 | N/A | PA0387 | N/A | N/A | N/A |
| PA0388 | PA0388 | PA0388 | PA0388 | PA0388 | PA0388 | PA0388 | PA0388 | N/A | PA0388 | PA0388 | PA0388 | PA0388 | PA0388 | PA0388 | PA0388 | N/A | PA0388 | N/A |
| PA0389 | PA0389 | PA0389 | PA0389 | PA0389 | N/A | N/A | PA0389 | PA0389 | PA0389 | PA0389 | PA0389 | PA0389 | PA0389 | PA0389 | N/A | N/A | N/A | N/A |
| PA0390 | PA0390 | PA0390 | N/A | PA0390 | N/A | N/A | PA0390 | PA0390 | PA0390 | PA0390 | PA0390 | PA0390 | PA0390 | PA0390 | N/A | N/A | PA0390 | N/A |
| PA0391 | N/A | N/A | N/A | PA0391 | N/A | N/A | N/A | PA0391 | PA0391 | N/A | N/A | N/A | N/A | N/A | N/A | N/A | N/A | N/A |
| PA0392 | PA0392 | PA0392 | N/A | PA0392 | PA0392 | PA0392 | PA0392 | PA0392 | PA0392 | PA0392 | PA0392 | PA0392 | PA0392 | PA0392 | N/A | N/A | PA0392 | N/A |
| PA0393 | PA0393 | PA0393 | PA0393 | PA0393 | PA0393 | N/A | PA0393 | PA0393 | PA0393 | PA0393 | PA0393 | PA0393 | PA0393 | N/A | PA0393 | N/A | N/A | N/A |
| PA0394 | PA0394 | PA0394 | N/A | PA0394 | N/A | PA0394 | PA0394 | N/A | PA0394 | PA0394 | PA0394 | PA0394 | PA0394 | N/A | PA0394 | N/A | N/A | N/A |
| PA0395 | PA0395 | PA0395 | PA0395 | PA0395 | PA0395 | PA0395 | PA0395 | PA0395 | PA0395 | PA0395 | PA0395 | PA0395 | PA0395 | PA0395 | N/A | N/A | N/A | N/A |
| PA0396 | PA0396 | PA0396 | PA0396 | PA0396 | N/A | PA0396 | N/A | PA0396 | PA0396 | PA0396 | PA0396 | PA0396 | PA0396 | PA0396 | N/A | N/A | PA0396 | PA0396 |
| PA0397 | N/A | N/A | N/A | N/A | N/A | N/A | N/A | N/A | N/A | N/A | N/A | N/A | N/A | PA0397 | N/A | N/A | N/A | N/A |
| N/A | N/A | N/A | N/A | N/A | PA0398 | N/A | N/A | N/A | N/A | N/A | N/A | N/A | N/A | N/A | N/A | N/A | N/A | N/A |
| PA0399 | PA0399 | PA0399 | PA0399 | PA0399 | PA0399 | PA0399 | PA0399 | PA0399 | PA0399 | PA0399 | PA0399 | PA0399 | PA0399 | N/A | PA0399 | N/A | PA0399 | N/A |
| PA0400 | PA0400 | PA0400 | PA0400 | PA0400 | PA0400 | PA0400 | PA0400 | PA0400 | PA0400 | PA0400 | PA0400 | PA0400 | PA0400 | PA0400 | PA0400 | PA0400 | N/A | N/A |
| PA0401 | PA0401 | PA0401 | PA0401 | PA0401 | PA0401 | PA0401 | PA0401 | PA0401 | PA0401 | PA0401 | PA0401 | PA0401 | PA0401 | PA0401 | PA0401 | N/A | N/A | N/A |
| PA0402 | PA0402 | PA0402 | PA0402 | PA0402 | PA0402 | PA0402 | N/A | N/A | PA0402 | PA0402 | PA0402 | PA0402 | PA0402 | PA0402 | PA0402 | N/A | PA0402 | N/A |
| PA0403 | PA0403 | PA0403 | PA0403 | PA0403 | PA0403 | PA0403 | N/A | N/A | PA0403 | PA0403 | PA0403 | PA0403 | PA0403 | N/A | PA0403 | N/A | N/A | N/A |
| PA0404 | PA0404 | N/A | N/A | N/A | N/A | N/A | N/A | N/A | PA0404 | N/A | N/A | N/A | N/A | N/A | N/A | N/A | N/A | N/A |
| PA0405 | N/A | PA0405 | N/A | PA0405 | N/A | N/A | N/A | N/A | PA0405 | N/A | PA0405 | N/A | N/A | N/A | N/A | N/A | N/A | N/A |
| PA0406 | N/A | N/A | N/A | PA0406 | PA0406 | N/A | N/A | PA0406 | PA0406 | PA0406 | PA0406 | PA0406 | N/A | PA0406 | PA0406 | N/A | PA0406 | N/A |
| PA0407 | PA0407 | PA0407 | PA0407 | PA0407 | PA0407 | PA0407 | N/A | N/A | PA0407 | PA0407 | PA0407 | PA0407 | PA0407 | N/A | PA0407 | N/A | N/A | N/A |
| PA0408 | PA0408 | PA0408 | PA0408 | PA0408 | PA0408 | PA0408 | PA0408 | N/A | PA0408 | PA0408 | PA0408 | PA0408 | PA0408 | PA0408 | N/A | N/A | PA0408 | PA0408 |
| PA0409 | PA0409 | PA0409 | PA0409 | PA0409 | PA0409 | PA0409 | N/A | N/A | PA0409 | PA0409 | PA0409 | PA0409 | PA0409 | N/A | PA0409 | N/A | N/A | N/A |
| PA0410 | PA0410 | PA0410 | PA0410 | PA0410 | PA0410 | N/A | PA0410 | N/A | PA0410 | PA0410 | PA0410 | PA0410 | N/A | N/A | N/A | N/A | N/A | N/A |
| PA0411 | PA0411 | PA0411 | PA0411 | PA0411 | PA0411 | PA0411 | PA0411 | PA0411 | PA0411 | PA0411 | PA0411 | PA0411 | PA0411 | PA0411 | PA0411 | N/A | PA0411 | PA0411 |
| N/A | N/A | N/A | N/A | N/A | PA0412 | N/A | N/A | N/A | N/A | N/A | N/A | N/A | N/A | N/A | N/A | N/A | N/A | N/A |
| PA0413 | PA0413 | PA0413 | PA0413 | PA0413 | PA0413 | PA0413 | PA0413 | PA0413 | PA0413 | PA0413 | PA0413 | PA0413 | PA0413 | PA0413 | N/A | N/A | PA0413 | N/A |
| N/A | N/A | N/A | N/A | PA0414 | N/A | N/A | N/A | N/A | N/A | PA0414 | N/A | N/A | N/A | N/A | N/A | N/A | N/A | N/A |
| PA0415 | PA0415 | PA0415 | N/A | PA0415 | N/A | PA0415 | PA0415 | PA0415 | PA0415 | PA0415 | PA0415 | PA0415 | PA0415 | N/A | N/A | N/A | N/A | N/A |
| PA0418 | N/A | N/A | N/A | PA0418 | PA0418 | N/A | N/A | N/A | N/A | N/A | PA0418 | N/A | N/A | N/A | N/A | N/A | N/A | N/A |
| PA0419 | PA0419 | PA0419 | PA0419 | PA0419 | PA0419 | N/A | PA0419 | N/A | PA0419 | PA0419 | PA0419 | PA0419 | PA0419 | PA0419 | PA0419 | N/A | N/A | N/A |
| PA0420 | PA0420 | PA0420 | PA0420 | PA0420 | PA0420 | PA0420 | N/A | N/A | PA0420 | PA0420 | PA0420 | PA0420 | PA0420 | N/A | N/A | N/A | N/A | N/A |
| PA0421 | PA0421 | PA0421 | PA0421 | PA0421 | PA0421 | N/A | N/A | PA0421 | PA0421 | PA0421 | PA0421 | PA0421 | PA0421 | PA0421 | PA0421 | N/A | PA0421 | PA0421 |

| | | | | | | | | | | | | | | | | | | |
|---|---|---|---|---|---|---|---|---|---|---|---|---|---|---|---|---|---|---|
| N/A | N/A | N/A | N/A | N/A | PA0422 | PA0422 | N/A | N/A | N/A | PA0422 | N/A | N/A | N/A | N/A | N/A | N/A | N/A | N/A |
| PA0423 | PA0423 | PA0423 | PA0423 | PA0423 | PA0423 | PA0423 | PA0423 | N/A | PA0423 | PA0423 | PA0423 | PA0423 | PA0423 | PA0423 | PA0423 | PA0423 | PA0423 | N/A |
| PA0424 | N/A | PA0424 | N/A | PA0424 | PA0424 | PA0424 | N/A | N/A | PA0424 | N/A | PA0424 | PA0424 | PA0424 | N/A | N/A | N/A | N/A | N/A |
| PA0425 | PA0425 | PA0425 | PA0425 | PA0425 | PA0425 | PA0425 | PA0425 | PA0425 | PA0425 | PA0425 | PA0425 | PA0425 | PA0425 | PA0425 | PA0425 | PA0425 | PA0425 | PA0425 |
| PA0426 | PA0426 | PA0426 | PA0426 | PA0426 | N/A | PA0426 | N/A | PA0426 | PA0426 | PA0426 | PA0426 | PA0426 | PA0426 | PA0426 | PA0426 | N/A | PA0426 | PA0426 |
| PA0427 | PA0427 | PA0427 | PA0427 | PA0427 | PA0427 | PA0427 | PA0427 | PA0427 | PA0427 | PA0427 | PA0427 | PA0427 | PA0427 | PA0427 | PA0427 | PA0427 | N/A | N/A |
| PA0428 | PA0428 | PA0428 | PA0428 | PA0428 | N/A | PA0428 | PA0428 | PA0428 | PA0428 | PA0428 | PA0428 | PA0428 | PA0428 | PA0428 | PA0428 | N/A | PA0428 | PA0428 |
| PA0429 | PA0429 | PA0429 | PA0429 | PA0429 | PA0429 | N/A | PA0429 | PA0429 | PA0429 | PA0429 | PA0429 | PA0429 | PA0429 | PA0429 | N/A | N/A | PA0429 | N/A |
| PA0430 | PA0430 | PA0430 | PA0430 | PA0430 | PA0430 | PA0430 | PA0430 | PA0430 | PA0430 | PA0430 | PA0430 | PA0430 | PA0430 | N/A | PA0430 | N/A | PA0430 | N/A |
| PA0431 | N/A | N/A | N/A | N/A | PA0431 | N/A | N/A | N/A | PA0431 | PA0431 | PA0431 | PA0431 | PA0431 | N/A | N/A | N/A | N/A | N/A |
| PA0432 | PA0432 | PA0432 | PA0432 | PA0432 | PA0432 | PA0432 | PA0432 | PA0432 | PA0432 | PA0432 | PA0432 | PA0432 | PA0432 | PA0432 | PA0432 | PA0432 | N/A | N/A |
| PA0434 | N/A | N/A | N/A | N/A | N/A | PA0434 | N/A | N/A | N/A | N/A | N/A | N/A | N/A | N/A | N/A | N/A | N/A | N/A |
| PA0435 | N/A | N/A | N/A | N/A | N/A | N/A | N/A | N/A | N/A | N/A | N/A | N/A | N/A | N/A | N/A | N/A | N/A | N/A |
| PA0436 | N/A | N/A | N/A | PA0436 | PA0436 | N/A | N/A | N/A | PA0436 | PA0436 | PA0436 | PA0436 | PA0436 | N/A | N/A | N/A | N/A | N/A |
| PA0437 | PA0437 | PA0437 | PA0437 | PA0437 | PA0437 | N/A | N/A | N/A | PA0437 | PA0437 | PA0437 | PA0437 | PA0437 | N/A | N/A | N/A | N/A | N/A |
| PA0438 | N/A | N/A | N/A | N/A | N/A | N/A | N/A | N/A | N/A | N/A | N/A | N/A | N/A | PA0438 | N/A | N/A | N/A | N/A |
| N/A | N/A | N/A | N/A | N/A | PA0443 | N/A | N/A | N/A | N/A | N/A | N/A | N/A | N/A | N/A | N/A | N/A | N/A | N/A |
| N/A | N/A | N/A | N/A | N/A | PA0444 | N/A | N/A | N/A | N/A | N/A | N/A | N/A | N/A | N/A | N/A | N/A | N/A | N/A |
| N/A | N/A | N/A | N/A | N/A | PA0445 | N/A | N/A | N/A | N/A | N/A | N/A | N/A | N/A | N/A | N/A | N/A | N/A | N/A |
| PA0446 | N/A | N/A | N/A | PA0446 | PA0446 | PA0446 | N/A | N/A | PA0446 | PA0446 | PA0446 | PA0446 | PA0446 | N/A | N/A | N/A | N/A | N/A |
| PA0447 | PA0447 | PA0447 | N/A | PA0447 | PA0447 | PA0447 | N/A | PA0447 | PA0447 | PA0447 | PA0447 | PA0447 | PA0447 | PA0447 | PA0447 | N/A | N/A | N/A |
| N/A | N/A | N/A | N/A | N/A | PA0448 | N/A | N/A | N/A | N/A | N/A | N/A | N/A | N/A | N/A | N/A | N/A | N/A | N/A |
| PA0449 | PA0449 | PA0449 | PA0449 | PA0449 | N/A | PA0449 | N/A | N/A | PA0449 | PA0449 | PA0449 | PA0449 | PA0449 | N/A | N/A | N/A | N/A | N/A |
| N/A | N/A | N/A | N/A | PA0451 | N/A | N/A | N/A | N/A | N/A | N/A | N/A | N/A | N/A | N/A | N/A | N/A | N/A | N/A |
| PA0452 | N/A | N/A | N/A | N/A | N/A | PA0452 | N/A | N/A | N/A | N/A | N/A | N/A | N/A | N/A | N/A | N/A | N/A | N/A |
| PA0454 | N/A | N/A | N/A | PA0454 | PA0454 | N/A | N/A | N/A | N/A | N/A | N/A | N/A | N/A | PA0454 | N/A | N/A | N/A | N/A |
| PA0455 | PA0455 | N/A | PA0455 | PA0455 | PA0455 | N/A | N/A | PA0455 | PA0455 | N/A | PA0455 | PA0455 | PA0455 | PA0455 | N/A | N/A | N/A | N/A |
| PA0456 | PA0456 | PA0456 | PA0456 | PA0456 | PA0456 | PA0456 | PA0456 | N/A | PA0456 | PA0456 | PA0456 | PA0456 | PA0456 | PA0456 | PA0456 | PA0456 | N/A | N/A |
| PA0457 | N/A | N/A | N/A | N/A | N/A | N/A | N/A | N/A | PA0457 | N/A | N/A | N/A | N/A | N/A | N/A | N/A | N/A | N/A |
| PA0458 | N/A | N/A | N/A | N/A | PA0458 | N/A | N/A | N/A | N/A | PA0458 | N/A | PA0458 | N/A | PA0458 | N/A | N/A | N/A | N/A |
| PA0459 | PA0459 | PA0459 | PA0459 | PA0459 | PA0459 | PA0459 | N/A | N/A | PA0459 | PA0459 | PA0459 | PA0459 | PA0459 | PA0459 | PA0459 | N/A | PA0459 | PA0459 |
| PA0460 | PA0460 | PA0460 | PA0460 | PA0460 | N/A | PA0460 | N/A | N/A | PA0460 | PA0460 | PA0460 | PA0460 | PA0460 | N/A | PA0460 | N/A | N/A | PA0460 |
| PA0461 | N/A | N/A | N/A | PA0461 | PA0461 | PA0461 | PA0461 | PA0461 | PA0461 | PA0461 | PA0461 | PA0461 | PA0461 | PA0461 | N/A | N/A | PA0461 | N/A |
| PA0462 | N/A | N/A | PA0462 | PA0462 | PA0462 | PA0462 | N/A | N/A | PA0462 | PA0462 | PA0462 | PA0462 | PA0462 | N/A | PA0462 | N/A | N/A | N/A |
| PA0463 | N/A | N/A | N/A | PA0463 | PA0463 | PA0463 | N/A | N/A | N/A | N/A | PA0463 | PA0463 | N/A | N/A | N/A | N/A | N/A | N/A |
| PA0464 | N/A | N/A | N/A | N/A | N/A | N/A | N/A | N/A | N/A | N/A | N/A | N/A | PA0464 | PA0464 | N/A | N/A | PA0464 | N/A |
| PA0467 | PA0467 | N/A | N/A | PA0467 | PA0467 | N/A | N/A | N/A | PA0467 | PA0467 | PA0467 | PA0467 | PA0467 | N/A | N/A | N/A | N/A | N/A |
| PA0468 | N/A | N/A | N/A | PA0468 | PA0468 | N/A | N/A | N/A | N/A | PA0468 | PA0468 | N/A | PA0468 | N/A | PA0468 | N/A | N/A | N/A |
| PA0469 | PA0469 | PA0469 | PA0469 | PA0469 | PA0469 | N/A | PA0469 | N/A | PA0469 | PA0469 | PA0469 | PA0469 | PA0469 | N/A | N/A | N/A | N/A | N/A |
| PA0470 | N/A | N/A | N/A | PA0470 | N/A | PA0470 | N/A | PA0470 | N/A | N/A | N/A | N/A | N/A | N/A | PA0470 | N/A | N/A | N/A |
| PA0471 | N/A | N/A | N/A | N/A | N/A | N/A | N/A | N/A | N/A | N/A | N/A | N/A | N/A | N/A | N/A | N/A | N/A | N/A |
| N/A | N/A | N/A | N/A | N/A | PA0472 | N/A | N/A | N/A | N/A | N/A | N/A | N/A | N/A | N/A | N/A | N/A | N/A | N/A |
| PA0473 | PA0473 | PA0473 | N/A | PA0473 | PA0473 | PA0473 | PA0473 | PA0473 | PA0473 | PA0473 | PA0473 | PA0473 | PA0473 | N/A | N/A | N/A | N/A | N/A |
| PA0475 | N/A | N/A | N/A | PA0475 | N/A | N/A | N/A | N/A | N/A | N/A | N/A | PA0475 | N/A | N/A | N/A | N/A | N/A | N/A |
| N/A | N/A | N/A | N/A | N/A | PA0477 | N/A | N/A | N/A | N/A | N/A | N/A | N/A | N/A | N/A | N/A | N/A | N/A | N/A |
| PA0478 | N/A | N/A | N/A | N/A | PA0478 | N/A | N/A | N/A | N/A | PA0478 | PA0478 | PA0478 | N/A | N/A | N/A | N/A | N/A | N/A |
| PA0479 | N/A | N/A | N/A | N/A | N/A | N/A | N/A | N/A | N/A | N/A | N/A | N/A | N/A | N/A | N/A | N/A | N/A | N/A |
| PA0480 | N/A | N/A | N/A | N/A | N/A | N/A | N/A | N/A | N/A | N/A | N/A | PA0480 | PA0480 | N/A | N/A | N/A | N/A | N/A |
| N/A | N/A | N/A | N/A | N/A | PA0481 | N/A | N/A | N/A | N/A | N/A | N/A | N/A | N/A | N/A | N/A | N/A | N/A | N/A |
| PA0482 | PA0482 | PA0482 | PA0482 | PA0482 | PA0482 | PA0482 | N/A | PA0482 | PA0482 | PA0482 | PA0482 | PA0482 | PA0482 | PA0482 | PA0482 | PA0482 | N/A | N/A |
| PA0483 | N/A | N/A | N/A | N/A | N/A | N/A | N/A | N/A | N/A | N/A | PA0483 | PA0483 | PA0483 | N/A | N/A | N/A | N/A | N/A |
| PA0484 | N/A | N/A | N/A | N/A | N/A | N/A | N/A | N/A | N/A | PA0484 | PA0484 | PA0484 | PA0484 | N/A | N/A | N/A | N/A | N/A |
| PA0485 | N/A | N/A | N/A | N/A | N/A | N/A | N/A | N/A | N/A | N/A | N/A | N/A | N/A | PA0485 | N/A | N/A | N/A | N/A |
| PA0486 | PA0486 | N/A | N/A | PA0486 | PA0486 | PA0486 | PA0486 | PA0486 | PA0486 | PA0486 | PA0486 | PA0486 | PA0486 | N/A | N/A | N/A | PA0486 | N/A |
| PA0487 | N/A | N/A | N/A | PA0487 | N/A | PA0487 | PA0487 | N/A | PA0487 | PA0487 | PA0487 | PA0487 | PA0487 | PA0487 | N/A | N/A | N/A | N/A |
| N/A | N/A | N/A | N/A | N/A | N/A | N/A | N/A | N/A | PA0492 | N/A | N/A | N/A | N/A | N/A | N/A | N/A | N/A | N/A |
| N/A | N/A | N/A | N/A | N/A | N/A | N/A | N/A | N/A | PA0493 | N/A | PA0493 | N/A | N/A | N/A | N/A | N/A | N/A | N/A |
| N/A | N/A | N/A | N/A | N/A | PA0494 | N/A | N/A | PA0494 | N/A | N/A | PA0494 | N/A | N/A | N/A | N/A | N/A | PA0494 | N/A |
| PA0495 | PA0495 | N/A | N/A | PA0495 | N/A | N/A | N/A | PA0495 | PA0495 | PA0495 | PA0495 | N/A | N/A | N/A | N/A | N/A | N/A | N/A |
| N/A | N/A | N/A | N/A | PA0496 | PA0496 | N/A | N/A | N/A | PA0496 | N/A | PA0496 | N/A | N/A | N/A | N/A | N/A | N/A | N/A |
| N/A | N/A | N/A | N/A | N/A | PA0499 | N/A | N/A | N/A | N/A | N/A | N/A | N/A | N/A | N/A | N/A | N/A | N/A | N/A |
| PA0500 | PA0500 | PA0500 | PA0500 | PA0500 | PA0500 | N/A | PA0500 | N/A | PA0500 | PA0500 | PA0500 | PA0500 | N/A | N/A | PA0500 | N/A | N/A | N/A |
| PA0501 | N/A | N/A | N/A | PA0501 | PA0501 | N/A | N/A | PA0501 | PA0501 | N/A | N/A | N/A | PA0501 | N/A | PA0501 | N/A | N/A | N/A |
| PA0502 | N/A | N/A | N/A | PA0502 | PA0502 | N/A | N/A | N/A | N/A | PA0502 | N/A | PA0502 | PA0502 | N/A | N/A | N/A | N/A | N/A |
| PA0503 | N/A | N/A | N/A | PA0503 | N/A | N/A | N/A | PA0503 | PA0503 | N/A | PA0503 | N/A | N/A | N/A | N/A | N/A | N/A | N/A |
| PA0504 | PA0504 | N/A | PA0504 | PA0504 | PA0504 | N/A | N/A | N/A | PA0504 | N/A | PA0504 | N/A | N/A | N/A | N/A | N/A | N/A | N/A |
| N/A | N/A | N/A | N/A | N/A | PA0505 | N/A | N/A | N/A | N/A | N/A | N/A | N/A | N/A | N/A | PA0505 | N/A | N/A | N/A |
| PA0506 | PA0506 | PA0506 | PA0506 | PA0506 | PA0506 | PA0506 | N/A | PA0506 | PA0506 | PA0506 | PA0506 | PA0506 | PA0506 | PA0506 | PA0506 | N/A | N/A | N/A |
| PA0507 | PA0507 | PA0507 | PA0507 | PA0507 | PA0507 | PA0507 | N/A | N/A | PA0507 | PA0507 | PA0507 | PA0507 | PA0507 | PA0507 | N/A | N/A | N/A | N/A |
| PA0508 | PA0508 | PA0508 | PA0508 | PA0508 | PA0508 | PA0508 | N/A | PA0508 | PA0508 | PA0508 | PA0508 | PA0508 | PA0508 | PA0508 | N/A | N/A | N/A | N/A |
| PA0509 | N/A | N/A | N/A | N/A | PA0509 | N/A | N/A | N/A | N/A | PA0509 | N/A | N/A | N/A | N/A | N/A | N/A | N/A | N/A |
| PA0510 | N/A | N/A | N/A | PA0510 | N/A | N/A | N/A | N/A | N/A | N/A | PA0510 | N/A | N/A | N/A | N/A | N/A | N/A | N/A |
| PA0511 | N/A | N/A | N/A | N/A | N/A | N/A | N/A | N/A | N/A | N/A | N/A | N/A | N/A | PA0511 | N/A | N/A | PA0511 | N/A |
| PA0512 | N/A | N/A | N/A | N/A | N/A | N/A | N/A | N/A | N/A | PA0512 | N/A | N/A | N/A | N/A | N/A | N/A | N/A | N/A |
| PA0513 | N/A | N/A | N/A | N/A | N/A | N/A | N/A | N/A | N/A | N/A | N/A | N/A | N/A | N/A | N/A | N/A | N/A | N/A |
| N/A | N/A | N/A | N/A | N/A | PA0514 | N/A | N/A | N/A | N/A | N/A | PA0514 | N/A | N/A | N/A | N/A | N/A | N/A | N/A |
| PA0515 | N/A | N/A | N/A | N/A | PA0515 | N/A | N/A | N/A | N/A | N/A | N/A | N/A | N/A | N/A | N/A | N/A | N/A | N/A |
| PA0516 | N/A | N/A | N/A | N/A | N/A | N/A | N/A | N/A | N/A | PA0516 | N/A | N/A | PA0516 | PA0516 | N/A | N/A | PA0516 | N/A |
| PA0517 | N/A | N/A | N/A | N/A | PA0517 | N/A | N/A | N/A | N/A | N/A | N/A | N/A | N/A | N/A | N/A | N/A | N/A | N/A |
| PA0518 | N/A | N/A | N/A | PA0518 | PA0518 | PA0518 | N/A | N/A | N/A | PA0518 | PA0518 | N/A | N/A | N/A | N/A | N/A | N/A | PA0518 |
| PA0519 | N/A | N/A | N/A | PA0519 | PA0519 | PA0519 | N/A | PA0519 | PA0519 | PA0519 | PA0519 | PA0519 | PA0519 | PA0519 | N/A | N/A | PA0519 | PA0519 |
| N/A | N/A | N/A | N/A | N/A | PA0520 | N/A | N/A | N/A | N/A | N/A | N/A | N/A | N/A | PA0520 | N/A | N/A | PA0520 | N/A |
| PA0523 | N/A | N/A | N/A | N/A | N/A | PA0523 | N/A | N/A | N/A | PA0523 | N/A | N/A | N/A | PA0523 | N/A | N/A | PA0523 | PA0523 |
| N/A | PA0524 | N/A | N/A | N/A | N/A | N/A | N/A | N/A | N/A | PA0524 | N/A | N/A | N/A | PA0524 | N/A | N/A | PA0524 | N/A |
| N/A | N/A | N/A | N/A | N/A | N/A | N/A | N/A | N/A | N/A | N/A | N/A | N/A | N/A | N/A | N/A | N/A | PA0525 | N/A |
| N/A | N/A | N/A | N/A | N/A | PA0526 | N/A | N/A | N/A | N/A | N/A | N/A | N/A | N/A | N/A | N/A | N/A | N/A | N/A |
| PA0527 | N/A | N/A | N/A | PA0527 | PA0527 | PA0527 | PA0527 | N/A | PA0527 | PA0527 | PA0527 | PA0527 | PA0527 | N/A | N/A | N/A | PA0527 | N/A |
| PA0528 | N/A | N/A | N/A | N/A | N/A | N/A | N/A | N/A | N/A | N/A | N/A | N/A | N/A | N/A | N/A | N/A | N/A | N/A |
| PA0533 | N/A | N/A | N/A | PA0533 | N/A | N/A | N/A | N/A | N/A | N/A | N/A | N/A | N/A | N/A | N/A | N/A | N/A | N/A |
| PA0534 | N/A | N/A | N/A | PA0534 | N/A | N/A | N/A | N/A | PA0534 | N/A | PA0534 | PA0534 | PA0534 | PA0534 | N/A | N/A | N/A | N/A |
| PA0535 | N/A | N/A | N/A | PA0535 | N/A | N/A | N/A | N/A | PA0535 | N/A | N/A | N/A | PA0535 | N/A | N/A | N/A | N/A | N/A |
| PA0536 | N/A | N/A | N/A | PA0536 | PA0536 | PA0536 | PA0536 | N/A | N/A | PA0536 | N/A | PA0536 | N/A | PA0536 | N/A | N/A | PA0536 | N/A |
| PA0537 | PA0537 | PA0537 | PA0537 | PA0537 | PA0537 | PA0537 | PA0537 | N/A | PA0537 | PA0537 | PA0537 | PA0537 | PA0537 | PA0537 | PA0537 | N/A | PA0537 | PA0537 |
| PA0538 | N/A | N/A | N/A | N/A | N/A | N/A | N/A | N/A | PA0538 | PA0538 | PA0538 | N/A | N/A | PA0538 | N/A | N/A | N/A | N/A |
| PA0540 | N/A | N/A | N/A | N/A | N/A | N/A | N/A | N/A | N/A | N/A | N/A | PA0540 | PA0540 | PA0540 | N/A | N/A | N/A | N/A |
| PA0541 | N/A | PA0541 | PA0541 | PA0541 | PA0541 | PA0541 | PA0541 | N/A | PA0541 | PA0541 | PA0541 | PA0541 | PA0541 | PA0541 | PA0541 | N/A | PA0541 | N/A |
| PA0542 | N/A | PA0542 | PA0542 | PA0542 | PA0542 | N/A | PA0542 | N/A | PA0542 | PA0542 | PA0542 | PA0542 | PA0542 | PA0542 | N/A | N/A | PA0542 | PA0542 |
| PA0543 | N/A | N/A | N/A | N/A | N/A | N/A | N/A | N/A | N/A | N/A | N/A | N/A | N/A | N/A | N/A | N/A | N/A | N/A |
| PA0544 | N/A | N/A | N/A | N/A | N/A | N/A | N/A | N/A | N/A | N/A | N/A | N/A | N/A | N/A | N/A | N/A | N/A | N/A |
| PA0545 | N/A | N/A | N/A | N/A | PA0545 | N/A | N/A | N/A | N/A | PA0545 | N/A | N/A | N/A | PA0545 | N/A | N/A | PA0545 | N/A |
| PA0546 | PA0546 | PA0546 | PA0546 | PA0546 | PA0546 | PA0546 | PA0546 | PA0546 | PA0546 | PA0546 | PA0546 | PA0546 | PA0546 | PA0546 | PA0546 | PA0546 | PA0546 | N/A |
| PA0547 | N/A | N/A | N/A | PA0547 | PA0547 | N/A | N/A | N/A | N/A | N/A | N/A | N/A | N/A | N/A | N/A | N/A | PA0547 | N/A |
| PA0548 | PA0548 | PA0548 | PA0548 | PA0548 | PA0548 | PA0548 | N/A | N/A | PA0548 | PA0548 | PA0548 | PA0548 | PA0548 | PA0548 | PA0548 | N/A | N/A | N/A |
| N/A | N/A | N/A | N/A | PA0549 | PA0549 | N/A | N/A | N/A | N/A | N/A | N/A | N/A | N/A | N/A | N/A | N/A | N/A | N/A |
| PA0550 | N/A | N/A | N/A | N/A | PA0550 | N/A | N/A | N/A | N/A | N/A | PA0550 | PA0550 | PA0550 | N/A | N/A | N/A | N/A | N/A |
| PA0551 | PA0551 | PA0551 | N/A | PA0551 | PA0551 | N/A | N/A | N/A | PA0551 | PA0551 | PA0551 | PA0551 | PA0551 | N/A | N/A | N/A | N/A | N/A |
| PA0552 | PA0552 | PA0552 | PA0552 | PA0552 | PA0552 | PA0552 | PA0552 | PA0552 | PA0552 | PA0552 | PA0552 | PA0552 | PA0552 | N/A | PA0552 | N/A | PA0552 | PA0552 |
| N/A | N/A | N/A | N/A | N/A | N/A | N/A | N/A | N/A | N/A | PA0553 | N/A | N/A | N/A | N/A | N/A | N/A | N/A | N/A |
| PA0554 | N/A | N/A | N/A | PA0554 | PA0554 | N/A | N/A | N/A | N/A | N/A | N/A | N/A | PA0554 | N/A | N/A | N/A | N/A | N/A |

| | | | | | | | | | | | | | | | | | | |
|---|---|---|---|---|---|---|---|---|---|---|---|---|---|---|---|---|---|---|
| PA0555 | PA0555 | PA0555 | PA0555 | PA0555 | PA0555 | PA0555 | PA0555 | PA0555 | PA0555 | PA0555 | PA0555 | PA0555 | PA0555 | PA0555 | PA0555 | PA0555 | PA0555 | N/A |
| PA0557 | N/A | N/A | N/A | N/A | PA0557 | N/A | N/A | N/A | N/A | PA0557 | N/A | N/A | N/A | N/A | N/A | N/A | N/A | N/A |
| PA0558 | N/A | N/A | N/A | N/A | PA0558 | PA0558 | N/A | N/A | PA0558 | PA0558 | PA0558 | PA0558 | PA0558 | N/A | N/A | N/A | N/A | N/A |
| PA0559 | N/A | N/A | N/A | PA0559 | N/A | N/A | PA0559 | N/A | PA0559 | N/A | PA0559 | N/A | PA0559 | N/A | N/A | N/A | N/A | N/A |
| N/A | N/A | N/A | N/A | N/A | PA0561 | N/A | N/A | N/A | N/A | N/A | N/A | N/A | N/A | PA0561 | N/A | N/A | N/A | N/A |
| PA0562 | PA0562 | PA0562 | PA0562 | PA0562 | PA0562 | N/A | N/A | N/A | PA0562 | N/A | PA0562 | N/A | N/A | N/A | N/A | N/A | N/A | N/A |
| PA0563 | N/A | PA0563 | N/A | PA0563 | N/A | PA0563 | PA0563 | PA0563 | PA0563 | PA0563 | PA0563 | PA0563 | PA0563 | PA0563 | N/A | PA0563 | PA0563 | N/A |
| PA0564 | N/A | N/A | N/A | PA0564 | N/A | N/A | N/A | N/A | N/A | N/A | N/A | N/A | N/A | N/A | N/A | N/A | N/A | N/A |
| PA0565 | N/A | N/A | N/A | N/A | N/A | PA0565 | N/A | N/A | N/A | PA0565 | PA0565 | PA0565 | PA0565 | N/A | N/A | N/A | N/A | N/A |
| N/A | N/A | N/A | N/A | N/A | PA0567 | N/A | N/A | N/A | N/A | N/A | N/A | N/A | N/A | N/A | N/A | N/A | N/A | N/A |
| PA0568 | N/A | N/A | N/A | PA0568 | N/A | N/A | N/A | N/A | PA0568 | PA0568 | PA0568 | PA0568 | PA0568 | N/A | N/A | N/A | N/A | N/A |
| N/A | N/A | N/A | N/A | PA0569 | N/A | N/A | N/A | N/A | PA0569 | N/A | N/A | N/A | N/A | N/A | N/A | N/A | N/A | N/A |
| PA0570 | N/A | N/A | N/A | N/A | N/A | N/A | N/A | N/A | PA0570 | N/A | PA0570 | PA0570 | N/A | N/A | N/A | N/A | N/A | N/A |
| PA0571 | N/A | N/A | N/A | PA0571 | N/A | N/A | PA0571 | N/A | PA0571 | N/A | PA0571 | N/A | N/A | N/A | N/A | N/A | N/A | N/A |
| PA0572 | N/A | N/A | N/A | PA0572 | N/A | N/A | N/A | N/A | N/A | N/A | PA0572 | PA0572 | PA0572 | N/A | PA0572 | PA0572 | N/A | N/A |
| N/A | N/A | N/A | N/A | PA0574 | N/A | N/A | N/A | N/A | N/A | N/A | N/A | N/A | N/A | N/A | N/A | N/A | N/A | N/A |
| PA0575 | N/A | N/A | N/A | N/A | PA0575 | N/A | N/A | N/A | N/A | N/A | N/A | N/A | N/A | N/A | N/A | N/A | N/A | N/A |
| PA0576 | PA0576 | PA0576 | PA0576 | PA0576 | PA0576 | PA0576 | PA0576 | PA0576 | PA0576 | PA0576 | PA0576 | PA0576 | PA0576 | PA0576 | PA0576 | N/A | PA0576 | N/A |
| PA0577 | PA0577 | PA0577 | N/A | PA0577 | N/A | N/A | N/A | PA0577 | PA0577 | PA0577 | PA0577 | PA0577 | N/A | PA0577 | N/A | N/A | N/A | N/A |
| PA0578 | PA0578 | PA0578 | N/A | PA0578 | PA0578 | N/A | N/A | PA0578 | PA0578 | PA0578 | N/A | N/A | PA0578 | N/A | PA0578 | N/A | N/A | N/A |
| PA0579 | PA0579 | PA0579 | PA0579 | PA0579 | PA0579 | PA0579 | PA0579 | PA0579 | PA0579 | PA0579 | PA0579 | PA0579 | PA0579 | PA0579 | PA0579 | N/A | PA0579 | PA0579 |
| PA0580 | PA0580 | PA0580 | PA0580 | PA0580 | N/A | N/A | PA0580 | PA0580 | PA0580 | N/A | PA0580 | PA0580 | PA0580 | N/A | N/A | N/A | PA0580 | N/A |
| PA0581 | N/A | N/A | N/A | N/A | N/A | PA0581 | N/A | N/A | PA0581 | PA0581 | PA0581 | PA0581 | PA0581 | PA0581 | N/A | N/A | N/A | N/A |
| PA0582 | N/A | N/A | N/A | PA0582 | N/A | N/A | N/A | N/A | PA0582 | N/A | PA0582 | PA0582 | PA0582 | N/A | N/A | N/A | N/A | N/A |
| PA0583 | PA0583 | N/A | PA0583 | PA0583 | PA0583 | N/A | PA0583 | N/A | PA0583 | PA0583 | PA0583 | PA0583 | N/A | N/A | N/A | N/A | N/A | N/A |
| PA0584 | N/A | N/A | N/A | PA0584 | PA0584 | N/A | N/A | N/A | N/A | N/A | N/A | N/A | PA0584 | N/A | N/A | N/A | N/A | N/A |
| PA0585 | N/A | N/A | N/A | N/A | PA0585 | N/A | N/A | N/A | N/A | PA0585 | N/A | PA0585 | N/A | N/A | N/A | N/A | N/A | N/A |
| PA0586 | N/A | N/A | N/A | N/A | PA0586 | PA0586 | N/A | N/A | PA0586 | PA0586 | PA0586 | PA0586 | PA0586 | PA0586 | N/A | N/A | N/A | N/A |
| PA0587 | N/A | N/A | N/A | N/A | PA0587 | PA0587 | N/A | N/A | N/A | PA0587 | N/A | PA0587 | PA0587 | PA0587 | N/A | N/A | N/A | N/A |
| PA0588 | N/A | N/A | PA0588 | PA0588 | PA0588 | PA0588 | N/A | N/A | PA0588 | PA0588 | PA0588 | PA0588 | PA0588 | PA0588 | N/A | N/A | N/A | N/A |
| PA0589 | N/A | PA0589 | N/A | N/A | PA0589 | N/A | N/A | N/A | PA0589 | N/A | PA0589 | PA0589 | PA0589 | N/A | N/A | N/A | N/A | N/A |
| PA0590 | N/A | N/A | PA0590 | PA0590 | PA0590 | N/A | PA0590 | N/A | PA0590 | N/A | PA0590 | PA0590 | PA0590 | PA0590 | N/A | N/A | N/A | N/A |
| PA0591 | PA0591 | PA0591 | PA0591 | PA0591 | PA0591 | N/A | PA0591 | N/A | PA0591 | PA0591 | PA0591 | PA0591 | PA0591 | N/A | PA0591 | N/A | N/A | N/A |
| PA0592 | PA0592 | PA0592 | PA0592 | PA0592 | PA0592 | PA0592 | PA0592 | PA0592 | PA0592 | PA0592 | PA0592 | PA0592 | PA0592 | PA0592 | PA0592 | N/A | N/A | N/A |
| PA0593 | PA0593 | PA0593 | PA0593 | PA0593 | PA0593 | N/A | N/A | N/A | PA0593 | PA0593 | PA0593 | PA0593 | PA0593 | N/A | N/A | N/A | N/A | N/A |
| PA0594 | PA0594 | PA0594 | PA0594 | PA0594 | PA0594 | PA0594 | PA0594 | N/A | PA0594 | PA0594 | PA0594 | PA0594 | PA0594 | PA0594 | PA0594 | N/A | N/A | N/A |
| PA0595 | PA0595 | PA0595 | PA0595 | PA0595 | PA0595 | PA0595 | N/A | PA0595 | PA0595 | PA0595 | PA0595 | PA0595 | PA0595 | PA0595 | PA0595 | N/A | PA0595 | N/A |
| PA0596 | PA0596 | PA0596 | PA0596 | PA0596 | PA0596 | N/A | PA0596 | PA0596 | PA0596 | PA0596 | PA0596 | PA0596 | PA0596 | N/A | PA0596 | N/A | N/A | N/A |
| PA0597 | PA0597 | N/A | N/A | PA0597 | PA0597 | N/A | PA0597 | N/A | PA0597 | PA0597 | N/A | N/A | PA0597 | N/A | PA0597 | N/A | N/A | N/A |
| PA0598 | N/A | N/A | N/A | N/A | PA0598 | N/A | N/A | N/A | PA0598 | N/A | N/A | PA0598 | PA0598 | PA0598 | N/A | N/A | N/A | N/A |
| N/A | N/A | N/A | N/A | PA0599 | PA0599 | N/A | N/A | N/A | N/A | PA0599 | N/A | PA0599 | PA0599 | N/A | N/A | N/A | N/A | N/A |
| PA0600 | N/A | N/A | N/A | N/A | PA0600 | N/A | N/A | N/A | N/A | N/A | N/A | N/A | N/A | PA0600 | N/A | N/A | N/A | N/A |
| PA0601 | N/A | N/A | N/A | PA0601 | PA0601 | N/A | N/A | N/A | N/A | PA0601 | PA0601 | N/A | PA0601 | N/A | N/A | N/A | N/A | N/A |
| PA0602 | N/A | N/A | N/A | PA0602 | PA0602 | N/A | PA0602 | N/A | PA0602 | N/A | PA0602 | PA0602 | PA0602 | PA0602 | N/A | N/A | N/A | N/A |
| PA0603 | PA0603 | PA0603 | PA0603 | PA0603 | PA0603 | PA0603 | PA0603 | PA0603 | PA0603 | PA0603 | PA0603 | PA0603 | PA0603 | PA0603 | PA0603 | N/A | PA0603 | PA0603 |
| PA0604 | N/A | N/A | N/A | PA0604 | PA0604 | PA0604 | PA0604 | N/A | PA0604 | PA0604 | PA0604 | PA0604 | PA0604 | N/A | PA0604 | PA0604 | N/A | N/A |
| PA0605 | PA0605 | PA0605 | PA0605 | PA0605 | N/A | PA0605 | N/A | PA0605 | PA0605 | PA0605 | N/A | PA0605 | N/A | PA0605 | PA0605 | N/A | PA0605 | N/A |
| PA0606 | PA0606 | N/A | N/A | PA0606 | PA0606 | PA0606 | N/A | N/A | PA0606 | PA0606 | N/A | N/A | N/A | PA0606 | N/A | N/A | N/A | N/A |
| PA0607 | PA0607 | PA0607 | PA0607 | PA0607 | PA0607 | PA0607 | PA0607 | N/A | PA0607 | PA0607 | PA0607 | PA0607 | PA0607 | N/A | PA0607 | PA0607 | N/A | N/A |
| PA0608 | N/A | N/A | N/A | PA0608 | PA0608 | N/A | PA0608 | N/A | N/A | N/A | N/A | N/A | N/A | N/A | N/A | N/A | N/A | N/A |
| PA0609 | PA0609 | PA0609 | PA0609 | PA0609 | PA0609 | PA0609 | N/A | PA0609 | PA0609 | PA0609 | PA0609 | PA0609 | PA0609 | N/A | N/A | N/A | N/A | N/A |
| PA0611 | N/A | N/A | PA0611 | PA0611 | N/A | N/A | N/A | PA0611 | PA0611 | PA0611 | PA0611 | PA0611 | PA0611 | N/A | PA0611 | N/A | N/A | N/A |
| N/A | N/A | N/A | N/A | N/A | N/A | N/A | N/A | N/A | N/A | N/A | N/A | N/A | N/A | N/A | PA0612 | N/A | N/A | N/A |
| PA0613 | N/A | N/A | N/A | N/A | N/A | N/A | N/A | N/A | N/A | N/A | N/A | N/A | N/A | N/A | PA0613 | N/A | N/A | N/A |
| PA0615 | N/A | N/A | N/A | N/A | N/A | N/A | N/A | N/A | N/A | N/A | N/A | N/A | N/A | N/A | PA0615 | PA0615 | N/A | N/A |
| PA0616 | N/A | N/A | N/A | N/A | N/A | N/A | N/A | N/A | N/A | N/A | N/A | N/A | N/A | N/A | PA0616 | PA0616 | N/A | N/A |
| PA0617 | N/A | N/A | N/A | N/A | N/A | N/A | N/A | N/A | N/A | N/A | N/A | N/A | N/A | N/A | PA0617 | N/A | N/A | N/A |
| PA0618 | N/A | N/A | N/A | PA0618 | N/A | N/A | N/A | N/A | N/A | N/A | N/A | N/A | N/A | N/A | PA0618 | PA0618 | N/A | N/A |
| N/A | N/A | N/A | N/A | N/A | PA0619 | N/A | N/A | N/A | N/A | N/A | N/A | N/A | N/A | N/A | PA0619 | PA0619 | N/A | N/A |
| PA0620 | N/A | N/A | N/A | PA0620 | PA0620 | N/A | N/A | PA0620 | N/A | N/A | N/A | N/A | N/A | PA0620 | PA0620 | PA0620 | N/A | N/A |
| PA0621 | N/A | N/A | N/A | N/A | PA0621 | N/A | N/A | N/A | N/A | N/A | N/A | N/A | N/A | N/A | PA0621 | N/A | N/A | N/A |
| PA0622 | PA0622 | PA0622 | PA0622 | PA0622 | PA0622 | PA0622 | N/A | PA0622 | PA0622 | PA0622 | N/A | PA0622 | PA0622 | N/A | PA0622 | PA0622 | PA0622 | N/A |
| PA0623 | N/A | PA0623 | PA0623 | PA0623 | N/A | PA0623 | PA0623 | N/A | PA0623 | PA0623 | PA0623 | PA0623 | N/A | N/A | PA0623 | PA0623 | PA0623 | N/A |
| N/A | N/A | N/A | N/A | PA0624 | N/A | N/A | N/A | N/A | N/A | N/A | N/A | N/A | N/A | N/A | PA0624 | N/A | N/A | N/A |
| PA0625 | N/A | N/A | N/A | N/A | N/A | N/A | N/A | PA0625 | N/A | N/A | N/A | N/A | N/A | PA0625 | PA0625 | PA0625 | N/A | N/A |
| N/A | N/A | N/A | N/A | N/A | N/A | N/A | N/A | N/A | N/A | N/A | N/A | N/A | N/A | N/A | PA0626 | PA0626 | PA0626 | N/A |
| N/A | N/A | N/A | N/A | N/A | N/A | N/A | N/A | N/A | N/A | N/A | N/A | N/A | N/A | N/A | PA0627 | N/A | N/A | N/A |
| PA0628 | N/A | N/A | N/A | N/A | N/A | N/A | N/A | N/A | N/A | N/A | N/A | N/A | N/A | N/A | PA0628 | PA0628 | PA0628 | PA0628 |
| PA0629 | N/A | N/A | N/A | N/A | N/A | N/A | N/A | N/A | N/A | N/A | N/A | N/A | N/A | N/A | PA0629 | PA0629 | PA0629 | N/A |
| PA0630 | N/A | N/A | N/A | N/A | N/A | N/A | N/A | N/A | N/A | N/A | N/A | N/A | N/A | N/A | PA0630 | N/A | N/A | N/A |
| N/A | N/A | N/A | N/A | N/A | N/A | N/A | N/A | N/A | N/A | N/A | N/A | N/A | N/A | N/A | PA0631 | N/A | N/A | N/A |
| PA0633 | N/A | N/A | N/A | PA0633 | N/A | N/A | PA0633 | N/A | PA0633 | N/A | N/A | N/A | N/A | N/A | PA0633 | PA0633 | N/A | N/A |
| PA0634 | N/A | N/A | N/A | PA0634 | N/A | N/A | N/A | N/A | N/A | N/A | PA0634 | N/A | N/A | N/A | PA0634 | N/A | PA0634 | N/A |
| PA0635 | N/A | N/A | N/A | N/A | N/A | N/A | N/A | N/A | N/A | N/A | N/A | N/A | N/A | N/A | N/A | N/A | N/A | N/A |
| PA0636 | N/A | N/A | N/A | N/A | N/A | N/A | N/A | N/A | N/A | N/A | N/A | N/A | N/A | N/A | PA0636 | N/A | N/A | N/A |
| PA0637 | N/A | N/A | N/A | N/A | N/A | N/A | N/A | N/A | N/A | N/A | N/A | N/A | N/A | N/A | PA0637 | N/A | N/A | N/A |
| PA0638 | N/A | N/A | N/A | N/A | N/A | N/A | N/A | N/A | N/A | N/A | N/A | N/A | N/A | N/A | PA0638 | PA0638 | N/A | N/A |
| N/A | N/A | N/A | N/A | N/A | N/A | N/A | N/A | N/A | N/A | N/A | N/A | N/A | N/A | N/A | PA0639 | N/A | N/A | N/A |
| PA0640 | N/A | N/A | N/A | N/A | N/A | N/A | N/A | N/A | N/A | N/A | N/A | N/A | N/A | N/A | PA0640 | N/A | N/A | N/A |
| PA0641 | N/A | N/A | N/A | PA0641 | N/A | N/A | N/A | N/A | N/A | N/A | N/A | N/A | N/A | N/A | PA0641 | PA0641 | PA0641 | N/A |
| N/A | N/A | N/A | N/A | N/A | N/A | N/A | N/A | N/A | N/A | N/A | N/A | N/A | N/A | N/A | PA0642 | N/A | N/A | N/A |
| N/A | N/A | N/A | N/A | N/A | N/A | N/A | N/A | N/A | N/A | N/A | N/A | N/A | N/A | N/A | PA0643 | N/A | N/A | N/A |
| N/A | N/A | N/A | N/A | N/A | N/A | N/A | N/A | N/A | N/A | N/A | N/A | N/A | N/A | N/A | PA0644 | N/A | N/A | N/A |
| N/A | N/A | N/A | N/A | N/A | N/A | N/A | N/A | N/A | N/A | N/A | N/A | N/A | N/A | N/A | PA0646 | N/A | N/A | N/A |
| N/A | N/A | N/A | N/A | N/A | N/A | N/A | N/A | N/A | N/A | N/A | N/A | N/A | N/A | N/A | PA0647 | N/A | N/A | N/A |
| N/A | N/A | N/A | N/A | N/A | PA0648 | N/A | N/A | N/A | N/A | N/A | N/A | N/A | N/A | N/A | N/A | N/A | N/A | N/A |
| PA0649 | PA0649 | PA0649 | PA0649 | PA0649 | PA0649 | PA0649 | PA0649 | N/A | PA0649 | PA0649 | PA0649 | PA0649 | PA0649 | N/A | PA0649 | N/A | N/A | N/A |
| PA0650 | PA0650 | PA0650 | PA0650 | PA0650 | PA0650 | N/A | N/A | PA0650 | PA0650 | PA0650 | PA0650 | PA0650 | PA0650 | N/A | PA0650 | N/A | N/A | N/A |
| PA0651 | PA0651 | PA0651 | PA0651 | PA0651 | PA0651 | PA0651 | N/A | N/A | PA0651 | PA0651 | PA0651 | PA0651 | PA0651 | N/A | PA0651 | N/A | N/A | N/A |
| PA0652 | PA0652 | PA0652 | PA0652 | PA0652 | PA0652 | PA0652 | PA0652 | PA0652 | PA0652 | PA0652 | PA0652 | PA0652 | PA0652 | PA0652 | PA0652 | N/A | PA0652 | N/A |
| PA0653 | N/A | PA0653 | N/A | PA0653 | PA0653 | PA0653 | N/A | N/A | PA0653 | PA0653 | PA0653 | PA0653 | PA0653 | N/A | N/A | N/A | N/A | N/A |
| PA0654 | PA0654 | PA0654 | PA0654 | PA0654 | PA0654 | PA0654 | PA0654 | N/A | PA0654 | PA0654 | PA0654 | PA0654 | PA0654 | N/A | PA0654 | PA0654 | N/A | N/A |
| PA0655 | PA0655 | PA0655 | PA0655 | PA0655 | N/A | N/A | PA0655 | N/A | PA0655 | PA0655 | PA0655 | PA0655 | PA0655 | PA0655 | N/A | N/A | PA0655 | N/A |
| PA0656 | N/A | N/A | N/A | N/A | N/A | PA0656 | N/A | N/A | N/A | PA0656 | PA0656 | PA0656 | PA0656 | N/A | N/A | N/A | N/A | N/A |
| PA0657 | N/A | N/A | N/A | PA0657 | N/A | N/A | N/A | PA0657 | N/A | PA0657 | N/A | N/A | PA0657 | N/A | N/A | N/A | N/A | N/A |
| PA0658 | N/A | N/A | N/A | PA0658 | N/A | PA0658 | N/A | N/A | PA0658 | PA0658 | PA0658 | PA0658 | PA0658 | PA0658 | N/A | N/A | N/A | N/A |
| PA0659 | PA0659 | PA0659 | PA0659 | PA0659 | PA0659 | PA0659 | N/A | N/A | PA0659 | PA0659 | PA0659 | PA0659 | PA0659 | PA0659 | PA0659 | N/A | PA0659 | N/A |
| PA0660 | N/A | PA0660 | N/A | PA0660 | PA0660 | PA0660 | N/A | N/A | PA0660 | PA0660 | PA0660 | PA0660 | PA0660 | N/A | PA0660 | N/A | N/A | N/A |
| PA0661 | N/A | N/A | N/A | N/A | PA0661 | PA0661 | N/A | N/A | N/A | N/A | N/A | N/A | N/A | PA0661 | N/A | N/A | PA0661 | N/A |
| PA0662 | PA0662 | PA0662 | PA0662 | PA0662 | PA0662 | PA0662 | PA0662 | PA0662 | PA0662 | PA0662 | PA0662 | PA0662 | PA0662 | PA0662 | N/A | PA0662 | N/A | N/A |
| PA0663 | N/A | N/A | N/A | PA0663 | N/A | N/A | N/A | N/A | N/A | N/A | PA0663 | N/A | N/A | PA0663 | PA0663 | N/A | PA0663 | PA0663 |
| PA0664 | PA0664 | PA0664 | PA0664 | PA0664 | N/A | N/A | PA0664 | N/A | PA0664 | PA0664 | PA0664 | PA0664 | PA0664 | N/A | N/A | N/A | PA0664 | N/A |
| PA0665 | PA0665 | PA0665 | PA0665 | PA0665 | N/A | PA0665 | PA0665 | N/A | PA0665 | PA0665 | PA0665 | PA0665 | PA0665 | N/A | PA0665 | N/A | N/A | N/A |
| PA0666 | PA0666 | PA0666 | PA0666 | PA0666 | N/A | N/A | N/A | N/A | N/A | N/A | N/A | N/A | N/A | N/A | N/A | N/A | N/A | N/A |
| N/A | N/A | N/A | N/A | N/A | PA0667 | N/A | N/A | N/A | PA0667 | N/A | N/A | N/A | N/A | PA0667 | N/A | N/A | PA0667 | N/A |
| PA0668 | PA0668 | PA0668 | PA0668 | PA0668 | PA0668 | PA0668 | PA0668 | PA0668 | PA0668 | PA0668 | PA0668 | PA0668 | PA0668 | N/A | PA0668 | PA0668 | N/A | N/A |
| PA0670 | N/A | N/A | N/A | N/A | N/A | N/A | N/A | N/A | N/A | N/A | N/A | N/A | N/A | N/A | N/A | N/A | N/A | N/A |
| PA0672 | N/A | N/A | N/A | PA0672 | N/A | PA0672 | N/A | N/A | PA0672 | N/A | N/A | N/A | PA0672 | N/A | PA0672 | N/A | N/A | N/A |
| PA0686 | PA0686 | N/A | PA0686 | PA0686 | N/A | PA0686 | N/A | N/A | PA0686 | PA0686 | PA0686 | PA0686 | PA0686 | PA0686 | N/A | N/A | N/A | N/A |

| | | | | | | | | | | | | | | | | | | |
|---|---|---|---|---|---|---|---|---|---|---|---|---|---|---|---|---|---|---|
| N/A | N/A | N/A | N/A | N/A | N/A | N/A | N/A | N/A | N/A | PA0688 | N/A | N/A | N/A | N/A | N/A | N/A | N/A | N/A |
| N/A | N/A | N/A | N/A | N/A | N/A | N/A | N/A | N/A | N/A | N/A | N/A | N/A | N/A | PA0703 | N/A | N/A | N/A | N/A |
| PA0704 | N/A | N/A | N/A | N/A | N/A | N/A | N/A | N/A | N/A | N/A | N/A | N/A | PA0704 | N/A | N/A | N/A | N/A | N/A |
| PA0705 | N/A | PA0705 | PA0705 | PA0705 | N/A | N/A | N/A | N/A | N/A | N/A | PA0705 | N/A | N/A | PA0705 | N/A | N/A | PA0705 | N/A |
| N/A | N/A | N/A | N/A | N/A | N/A | N/A | N/A | N/A | PA0706 | N/A | N/A | N/A | N/A | N/A | N/A | N/A | N/A | N/A |
| N/A | PA0708 | N/A | N/A | PA0708 | N/A | N/A | N/A | N/A | N/A | N/A | N/A | N/A | N/A | N/A | N/A | N/A | N/A | N/A |
| PA0711 | N/A | N/A | N/A | N/A | N/A | N/A | N/A | N/A | N/A | N/A | N/A | N/A | N/A | N/A | N/A | N/A | N/A | N/A |
| N/A | N/A | N/A | N/A | N/A | PA0712 | N/A | N/A | N/A | N/A | N/A | N/A | N/A | N/A | N/A | N/A | N/A | N/A | N/A |
| PA0713 | N/A | N/A | N/A | N/A | PA0713 | N/A | N/A | N/A | N/A | PA0713 | N/A | N/A | N/A | N/A | N/A | N/A | N/A | N/A |
| PA0714 | N/A | N/A | N/A | N/A | N/A | N/A | N/A | N/A | N/A | N/A | N/A | N/A | N/A | N/A | N/A | N/A | N/A | N/A |
| PA0715 | N/A | N/A | N/A | PA0715 | N/A | PA0715 | N/A | PA0715 | N/A | N/A | PA0715 | PA0715 | PA0715 | PA0715 | N/A | N/A | N/A | N/A |
| PA0716 | N/A | N/A | N/A | PA0716 | N/A | N/A | N/A | PA0716 | N/A | N/A | N/A | N/A | N/A | N/A | N/A | N/A | N/A | N/A |
| N/A | N/A | N/A | N/A | N/A | N/A | N/A | PA0720 | N/A | PA0720 | PA0720 | PA0720 | N/A | N/A | N/A | N/A | N/A | N/A | N/A |
| PA0729 | N/A | N/A | N/A | PA0729 | N/A | N/A | N/A | N/A | PA0729 | N/A | PA0729 | PA0729 | PA0729 | N/A | N/A | N/A | N/A | N/A |
| N/A | N/A | N/A | N/A | N/A | PA0731 | N/A | N/A | N/A | N/A | N/A | N/A | N/A | N/A | N/A | N/A | N/A | N/A | N/A |
| PA0732 | N/A | N/A | N/A | N/A | PA0732 | N/A | N/A | N/A | N/A | N/A | N/A | N/A | N/A | N/A | N/A | N/A | N/A | N/A |
| PA0735 | N/A | N/A | N/A | PA0735 | N/A | N/A | PA0735 | N/A | PA0735 | N/A | PA0735 | N/A | N/A | PA0735 | PA0735 | N/A | PA0735 | N/A |
| PA0736 | N/A | N/A | N/A | PA0736 | N/A | N/A | N/A | N/A | N/A | N/A | N/A | N/A | N/A | PA0736 | N/A | N/A | N/A | N/A |
| PA0737 | N/A | N/A | N/A | N/A | N/A | N/A | N/A | N/A | N/A | N/A | N/A | N/A | N/A | N/A | N/A | N/A | N/A | N/A |
| N/A | N/A | N/A | N/A | N/A | N/A | N/A | N/A | N/A | N/A | N/A | N/A | N/A | PA0738 | N/A | N/A | N/A | N/A | N/A |
| PA0739 | N/A | N/A | N/A | N/A | N/A | N/A | N/A | N/A | N/A | N/A | N/A | N/A | N/A | N/A | N/A | N/A | N/A | N/A |
| N/A | N/A | N/A | N/A | N/A | PA0740 | N/A | N/A | N/A | N/A | N/A | N/A | N/A | N/A | N/A | N/A | N/A | N/A | N/A |
| PA0741 | N/A | N/A | N/A | PA0741 | PA0741 | N/A | N/A | N/A | PA0741 | PA0741 | N/A | PA0741 | PA0741 | N/A | N/A | N/A | N/A | N/A |
| PA0743 | N/A | N/A | N/A | N/A | PA0743 | PA0743 | N/A | N/A | N/A | N/A | N/A | N/A | PA0743 | N/A | N/A | N/A | N/A | N/A |
| PA0744 | N/A | N/A | N/A | PA0744 | PA0744 | PA0744 | N/A | N/A | PA0744 | PA0744 | PA0744 | PA0744 | PA0744 | N/A | N/A | N/A | N/A | N/A |
| PA0745 | PA0745 | PA0745 | PA0745 | PA0745 | PA0745 | PA0745 | N/A | N/A | PA0745 | PA0745 | PA0745 | PA0745 | PA0745 | PA0745 | N/A | PA0745 | PA0745 | PA0745 |
| PA0746 | N/A | N/A | N/A | PA0746 | PA0746 | N/A | N/A | N/A | N/A | PA0746 | PA0746 | PA0746 | PA0746 | N/A | N/A | N/A | N/A | N/A |
| PA0747 | N/A | N/A | N/A | N/A | PA0747 | PA0747 | N/A | N/A | PA0747 | PA0747 | PA0747 | PA0747 | PA0747 | PA0747 | N/A | PA0747 | N/A | N/A |
| N/A | N/A | N/A | N/A | N/A | PA0749 | N/A | N/A | N/A | N/A | N/A | N/A | N/A | N/A | N/A | N/A | N/A | N/A | N/A |
| PA0750 | PA0750 | PA0750 | PA0750 | PA0750 | PA0750 | PA0750 | PA0750 | N/A | PA0750 | PA0750 | PA0750 | PA0750 | PA0750 | PA0750 | N/A | N/A | N/A | N/A |
| N/A | N/A | N/A | N/A | PA0751 | N/A | N/A | N/A | N/A | N/A | N/A | N/A | N/A | N/A | N/A | N/A | N/A | N/A | N/A |
| PA0753 | N/A | N/A | N/A | N/A | PA0753 | N/A | N/A | N/A | N/A | N/A | N/A | N/A | N/A | N/A | N/A | N/A | N/A | N/A |
| PA0754 | N/A | N/A | N/A | PA0754 | PA0754 | PA0754 | N/A | N/A | N/A | N/A | PA0754 | PA0754 | PA0754 | N/A | PA0754 | N/A | N/A | N/A |
| PA0755 | PA0755 | PA0755 | PA0755 | PA0755 | N/A | PA0755 | N/A | N/A | N/A | PA0755 | N/A | N/A | N/A | N/A | PA0755 | N/A | N/A | N/A |
| PA0756 | PA0756 | PA0756 | N/A | PA0756 | N/A | N/A | N/A | N/A | PA0756 | N/A | PA0756 | PA0756 | PA0756 | N/A | N/A | N/A | N/A | N/A |
| PA0757 | N/A | N/A | N/A | N/A | PA0757 | N/A | N/A | N/A | N/A | N/A | N/A | N/A | N/A | N/A | N/A | N/A | N/A | N/A |
| PA0758 | PA0758 | PA0758 | PA0758 | PA0758 | PA0758 | PA0758 | PA0758 | N/A | PA0758 | PA0758 | PA0758 | PA0758 | PA0758 | N/A | PA0758 | N/A | N/A | N/A |
| PA0759 | PA0759 | PA0759 | PA0759 | PA0759 | PA0759 | PA0759 | PA0759 | N/A | PA0759 | PA0759 | PA0759 | PA0759 | PA0759 | N/A | PA0759 | N/A | N/A | N/A |
| PA0760 | N/A | PA0760 | PA0760 | N/A | N/A | N/A | N/A | N/A | N/A | N/A | PA0760 | PA0760 | PA0760 | N/A | N/A | N/A | N/A | N/A |
| PA0761 | PA0761 | N/A | PA0761 | PA0761 | PA0761 | N/A | N/A | PA0761 | PA0761 | PA0761 | PA0761 | PA0761 | PA0761 | N/A | N/A | N/A | N/A | N/A |
| PA0762 | PA0762 | PA0762 | PA0762 | PA0762 | PA0762 | PA0762 | PA0762 | N/A | PA0762 | PA0762 | PA0762 | PA0762 | PA0762 | PA0762 | PA0762 | N/A | PA0762 | N/A |
| PA0763 | PA0763 | PA0763 | PA0763 | PA0763 | PA0763 | PA0763 | PA0763 | N/A | PA0763 | PA0763 | PA0763 | PA0763 | PA0763 | PA0763 | PA0763 | N/A | PA0763 | PA0763 |
| PA0764 | PA0764 | PA0764 | PA0764 | PA0764 | PA0764 | PA0764 | PA0764 | N/A | PA0764 | PA0764 | PA0764 | PA0764 | PA0764 | PA0764 | PA0764 | N/A | N/A | N/A |
| PA0766 | PA0766 | PA0766 | PA0766 | PA0766 | PA0766 | PA0766 | PA0766 | PA0766 | PA0766 | PA0766 | PA0766 | PA0766 | PA0766 | PA0766 | PA0766 | PA0766 | PA0766 | PA0766 |
| PA0767 | PA0767 | PA0767 | PA0767 | PA0767 | PA0767 | PA0767 | N/A | PA0767 | PA0767 | PA0767 | PA0767 | PA0767 | PA0767 | PA0767 | PA0767 | N/A | PA0767 | N/A |
| PA0768 | PA0768 | PA0768 | PA0768 | PA0768 | N/A | PA0768 | PA0768 | N/A | PA0768 | PA0768 | PA0768 | PA0768 | PA0768 | PA0768 | PA0768 | N/A | PA0768 | N/A |
| PA0769 | N/A | N/A | N/A | PA0769 | PA0769 | PA0769 | PA0769 | N/A | PA0769 | PA0769 | PA0769 | PA0769 | PA0769 | PA0769 | PA0769 | N/A | PA0769 | N/A |
| PA0770 | PA0770 | PA0770 | N/A | PA0770 | PA0770 | N/A | N/A | N/A | PA0770 | PA0770 | PA0770 | PA0770 | PA0770 | PA0770 | N/A | N/A | N/A | N/A |
| PA0771 | PA0771 | PA0771 | PA0771 | PA0771 | PA0771 | N/A | N/A | PA0771 | PA0771 | PA0771 | PA0771 | PA0771 | PA0771 | PA0771 | N/A | N/A | PA0771 | PA0771 |
| PA0772 | N/A | N/A | N/A | PA0772 | PA0772 | N/A | PA0772 | N/A | PA0772 | N/A | N/A | N/A | N/A | N/A | N/A | N/A | N/A | N/A |
| PA0773 | PA0773 | N/A | N/A | PA0773 | PA0773 | PA0773 | N/A | N/A | PA0773 | PA0773 | PA0773 | PA0773 | PA0773 | PA0773 | N/A | N/A | N/A | N/A |
| PA0774 | N/A | N/A | N/A | PA0774 | N/A | N/A | N/A | PA0774 | N/A | N/A | N/A | N/A | N/A | N/A | N/A | N/A | N/A | N/A |
| PA0775 | PA0775 | PA0775 | PA0775 | PA0775 | PA0775 | N/A | N/A | N/A | PA0775 | PA0775 | PA0775 | PA0775 | PA0775 | N/A | PA0775 | N/A | N/A | N/A |
| PA0776 | N/A | N/A | N/A | N/A | N/A | N/A | N/A | N/A | N/A | N/A | N/A | N/A | N/A | N/A | N/A | N/A | N/A | N/A |
| PA0778 | N/A | N/A | PA0778 | PA0778 | PA0778 | PA0778 | PA0778 | N/A | PA0778 | PA0778 | PA0778 | PA0778 | N/A | PA0778 | PA0778 | N/A | N/A | N/A |
| PA0779 | PA0779 | PA0779 | PA0779 | PA0779 | PA0779 | PA0779 | PA0779 | PA0779 | PA0779 | PA0779 | PA0779 | PA0779 | PA0779 | PA0779 | PA0779 | PA0779 | PA0779 | PA0779 |
| PA0781 | N/A | N/A | N/A | N/A | PA0781 | N/A | N/A | PA0781 | N/A | PA0781 | N/A | N/A | N/A | N/A | N/A | N/A | N/A | N/A |
| PA0782 | PA0782 | PA0782 | PA0782 | PA0782 | PA0782 | PA0782 | PA0782 | PA0782 | PA0782 | PA0782 | PA0782 | PA0782 | PA0782 | PA0782 | PA0782 | PA0782 | PA0782 | N/A |
| PA0783 | PA0783 | PA0783 | PA0783 | PA0783 | N/A | PA0783 | N/A | PA0783 | PA0783 | PA0783 | PA0783 | PA0783 | PA0783 | PA0783 | PA0783 | N/A | PA0783 | PA0783 |
| PA0784 | N/A | N/A | N/A | PA0784 | N/A | N/A | N/A | N/A | N/A | N/A | PA0784 | N/A | PA0784 | N/A | N/A | N/A | N/A | N/A |
| PA0785 | N/A | N/A | N/A | PA0785 | N/A | N/A | N/A | N/A | N/A | N/A | N/A | N/A | N/A | N/A | N/A | N/A | N/A | N/A |
| PA0787 | N/A | N/A | N/A | N/A | N/A | N/A | N/A | N/A | N/A | N/A | N/A | N/A | N/A | N/A | N/A | N/A | N/A | N/A |
| PA0788 | N/A | N/A | N/A | N/A | N/A | N/A | N/A | N/A | N/A | N/A | N/A | N/A | N/A | N/A | N/A | N/A | N/A | N/A |
| PA0789 | PA0789 | PA0789 | N/A | PA0789 | N/A | PA0789 | N/A | PA0789 | PA0789 | PA0789 | PA0789 | PA0789 | N/A | PA0789 | PA0789 | N/A | PA0789 | N/A |
| N/A | N/A | N/A | N/A | N/A | PA0791 | N/A | N/A | N/A | N/A | N/A | N/A | N/A | N/A | N/A | N/A | N/A | N/A | N/A |
| PA0792 | N/A | N/A | N/A | N/A | PA0792 | PA0792 | N/A | N/A | N/A | PA0792 | N/A | PA0792 | PA0792 | N/A | N/A | N/A | N/A | N/A |
| PA0793 | N/A | N/A | N/A | PA0793 | N/A | PA0793 | N/A | N/A | N/A | N/A | PA0793 | PA0793 | PA0793 | N/A | N/A | N/A | N/A | N/A |
| PA0794 | PA0794 | PA0794 | PA0794 | PA0794 | N/A | PA0794 | N/A | N/A | PA0794 | PA0794 | PA0794 | PA0794 | PA0794 | PA0794 | PA0794 | N/A | N/A | N/A |
| PA0795 | PA0795 | PA0795 | PA0795 | PA0795 | PA0795 | PA0795 | N/A | N/A | PA0795 | PA0795 | PA0795 | PA0795 | PA0795 | N/A | PA0795 | PA0795 | N/A | N/A |
| PA0796 | PA0796 | PA0796 | PA0796 | PA0796 | PA0796 | PA0796 | PA0796 | PA0796 | PA0796 | PA0796 | PA0796 | PA0796 | PA0796 | PA0796 | N/A | N/A | N/A | N/A |
| PA0797 | N/A | N/A | N/A | N/A | PA0797 | N/A | N/A | N/A | N/A | N/A | PA0797 | PA0797 | PA0797 | N/A | N/A | N/A | N/A | N/A |
| PA0798 | N/A | N/A | N/A | N/A | N/A | N/A | N/A | N/A | N/A | PA0798 | N/A | PA0798 | PA0798 | N/A | N/A | N/A | N/A | N/A |
| PA0799 | N/A | N/A | N/A | PA0799 | N/A | N/A | N/A | PA0799 | PA0799 | PA0799 | PA0799 | N/A | N/A | N/A | N/A | N/A | N/A | N/A |
| PA0801 | N/A | N/A | N/A | PA0801 | N/A | N/A | N/A | N/A | N/A | N/A | N/A | N/A | N/A | N/A | N/A | N/A | N/A | N/A |
| PA0803 | N/A | N/A | N/A | N/A | PA0803 | N/A | N/A | N/A | N/A | N/A | N/A | PA0803 | PA0803 | N/A | N/A | N/A | N/A | N/A |
| PA0804 | N/A | N/A | N/A | PA0804 | PA0804 | N/A | N/A | N/A | N/A | N/A | PA0804 | PA0804 | PA0804 | N/A | N/A | N/A | N/A | N/A |
| N/A | N/A | N/A | N/A | N/A | N/A | N/A | N/A | N/A | N/A | PA0805 | N/A | N/A | N/A | N/A | N/A | N/A | N/A | N/A |
| N/A | N/A | N/A | N/A | N/A | PA0806 | N/A | N/A | N/A | N/A | N/A | N/A | N/A | N/A | N/A | N/A | N/A | N/A | N/A |
| PA0807 | N/A | N/A | N/A | N/A | PA0807 | PA0807 | PA0807 | N/A | PA0807 | PA0807 | N/A | PA0807 | PA0807 | N/A | PA0807 | PA0807 | N/A | N/A |
| PA0808 | N/A | N/A | N/A | N/A | PA0808 | N/A | N/A | N/A | N/A | PA0808 | N/A | N/A | N/A | N/A | N/A | N/A | N/A | N/A |
| PA0810 | N/A | N/A | N/A | N/A | N/A | N/A | N/A | N/A | N/A | N/A | N/A | N/A | PA0810 | N/A | N/A | N/A | N/A | N/A |
| PA0814 | N/A | N/A | N/A | PA0814 | N/A | N/A | N/A | N/A | N/A | N/A | N/A | N/A | N/A | N/A | N/A | N/A | N/A | N/A |
| PA0816 | N/A | N/A | N/A | N/A | N/A | PA0816 | N/A | N/A | N/A | N/A | N/A | N/A | N/A | PA0816 | N/A | N/A | N/A | N/A |
| PA0817 | N/A | N/A | N/A | PA0817 | PA0817 | N/A | N/A | N/A | N/A | N/A | N/A | N/A | N/A | N/A | N/A | N/A | N/A | N/A |
| N/A | N/A | N/A | N/A | PA0820 | N/A | N/A | N/A | N/A | N/A | N/A | PA0820 | PA0820 | N/A | N/A | N/A | N/A | N/A | N/A |
| PA0825 | N/A | N/A | N/A | N/A | N/A | N/A | N/A | N/A | N/A | N/A | N/A | N/A | N/A | N/A | N/A | N/A | N/A | N/A |
| PA0827 | N/A | N/A | N/A | PA0827 | N/A | N/A | N/A | N/A | N/A | N/A | N/A | N/A | N/A | N/A | N/A | N/A | N/A | N/A |
| N/A | N/A | N/A | N/A | PA0830 | N/A | N/A | N/A | N/A | N/A | N/A | N/A | N/A | N/A | N/A | N/A | N/A | N/A | N/A |
| N/A | N/A | N/A | N/A | N/A | PA0831 | N/A | N/A | N/A | N/A | N/A | N/A | N/A | N/A | N/A | N/A | N/A | N/A | N/A |
| PA0832 | PA0832 | PA0832 | N/A | PA0832 | PA0832 | PA0832 | N/A | N/A | PA0832 | PA0832 | PA0832 | PA0832 | PA0832 | N/A | N/A | N/A | N/A | N/A |
| PA0833 | PA0833 | PA0833 | PA0833 | PA0833 | PA0833 | PA0833 | PA0833 | PA0833 | PA0833 | PA0833 | PA0833 | PA0833 | PA0833 | PA0833 | PA0833 | PA0833 | PA0833 | PA0833 |
| PA0834 | N/A | N/A | PA0834 | PA0834 | PA0834 | PA0834 | N/A | PA0834 | PA0834 | PA0834 | PA0834 | PA0834 | PA0834 | PA0834 | PA0834 | N/A | N/A | N/A |
| PA0835 | N/A | N/A | N/A | PA0835 | PA0835 | PA0835 | N/A | PA0835 | PA0835 | PA0835 | PA0835 | PA0835 | PA0835 | PA0835 | N/A | N/A | N/A | N/A |
| PA0836 | PA0836 | N/A | PA0836 | PA0836 | PA0836 | PA0836 | N/A | N/A | PA0836 | PA0836 | PA0836 | PA0836 | PA0836 | N/A | N/A | N/A | N/A | N/A |
| PA0837 | PA0837 | PA0837 | PA0837 | PA0837 | PA0837 | PA0837 | PA0837 | N/A | PA0837 | PA0837 | PA0837 | PA0837 | PA0837 | N/A | PA0837 | N/A | N/A | N/A |
| PA0838 | PA0838 | PA0838 | N/A | PA0838 | PA0838 | PA0838 | N/A | N/A | PA0838 | PA0838 | PA0838 | PA0838 | PA0838 | N/A | PA0838 | N/A | N/A | N/A |
| N/A | N/A | N/A | N/A | N/A | PA0839 | N/A | N/A | N/A | N/A | N/A | N/A | N/A | N/A | N/A | N/A | N/A | N/A | N/A |
| PA0840 | PA0840 | N/A | PA0840 | PA0840 | PA0840 | N/A | N/A | N/A | PA0840 | PA0840 | PA0840 | PA0840 | PA0840 | N/A | PA0840 | N/A | N/A | N/A |
| PA0841 | N/A | N/A | N/A | PA0841 | N/A | N/A | N/A | N/A | N/A | N/A | N/A | N/A | N/A | PA0841 | N/A | N/A | N/A | N/A |
| N/A | N/A | N/A | N/A | N/A | N/A | N/A | N/A | N/A | N/A | N/A | N/A | N/A | N/A | N/A | N/A | PA0844 | N/A | N/A |
| PA0846 | N/A | N/A | N/A | PA0846 | N/A | PA0846 | N/A | N/A | N/A | PA0846 | PA0846 | PA0846 | PA0846 | PA0846 | N/A | N/A | N/A | N/A |
| PA0847 | N/A | N/A | N/A | PA0847 | N/A | N/A | N/A | N/A | N/A | N/A | N/A | N/A | N/A | PA0847 | N/A | N/A | N/A | N/A |
| PA0848 | PA0848 | PA0848 | PA0848 | PA0848 | PA0848 | N/A | N/A | N/A | PA0848 | PA0848 | PA0848 | PA0848 | PA0848 | N/A | PA0848 | N/A | N/A | N/A |
| PA0849 | N/A | N/A | PA0849 | PA0849 | N/A | PA0849 | N/A | N/A | PA0849 | PA0849 | PA0849 | N/A | N/A | N/A | PA0849 | N/A | N/A | N/A |
| PA0850 | N/A | N/A | N/A | N/A | PA0850 | PA0850 | N/A | N/A | N/A | PA0850 | N/A | N/A | N/A | N/A | N/A | N/A | N/A | N/A |
| PA0851 | N/A | N/A | N/A | PA0851 | PA0851 | N/A | N/A | N/A | N/A | N/A | N/A | N/A | PA0851 | N/A | N/A | N/A | N/A | N/A |
| PA0852 | N/A | N/A | N/A | PA0852 | PA0852 | PA0852 | PA0852 | PA0852 | PA0852 | PA0852 | PA0852 | PA0852 | PA0852 | N/A | PA0852 | PA0852 | N/A | PA0852 |
| PA0853 | N/A | N/A | N/A | N/A | PA0853 | PA0853 | PA0853 | N/A | PA0853 | PA0853 | PA0853 | PA0853 | PA0853 | N/A | N/A | N/A | N/A | N/A |
| PA0854 | PA0854 | PA0854 | PA0854 | PA0854 | PA0854 | PA0854 | N/A | N/A | PA0854 | PA0854 | PA0854 | PA0854 | PA0854 | PA0854 | PA0854 | PA0854 | N/A | N/A |

| | | | | | | | | | | | | | | | | | | |
|---|---|---|---|---|---|---|---|---|---|---|---|---|---|---|---|---|---|---|
| N/A | N/A | N/A | N/A | N/A | PA0855 | N/A | N/A | N/A | N/A | N/A | N/A | N/A | N/A | N/A | N/A | N/A | N/A | N/A |
| PA0856 | PA0856 | PA0856 | PA0856 | PA0856 | PA0856 | PA0856 | PA0856 | N/A | PA0856 | PA0856 | PA0856 | PA0856 | PA0856 | PA0856 | PA0856 | N/A | PA0856 | PA0856 |
| PA0857 | PA0857 | PA0857 | PA0857 | PA0857 | PA0857 | PA0857 | N/A | N/A | PA0857 | PA0857 | PA0857 | PA0857 | PA0857 | N/A | PA0857 | N/A | N/A | N/A |
| PA0858 | PA0858 | PA0858 | PA0858 | PA0858 | PA0858 | N/A | PA0858 | PA0858 | PA0858 | PA0858 | PA0858 | PA0858 | PA0858 | N/A | N/A | N/A | N/A | N/A |
| PA0859 | PA0859 | N/A | PA0859 | PA0859 | N/A | N/A | PA0859 | N/A | PA0859 | PA0859 | PA0859 | PA0859 | PA0859 | N/A | N/A | N/A | N/A | N/A |
| PA0860 | N/A | N/A | N/A | PA0860 | N/A | PA0860 | N/A | N/A | N/A | N/A | N/A | N/A | N/A | PA0860 | N/A | N/A | N/A | N/A |
| PA0861 | N/A | N/A | N/A | PA0861 | N/A | N/A | N/A | N/A | N/A | N/A | N/A | N/A | PA0861 | PA0861 | N/A | N/A | N/A | N/A |
| PA0862 | N/A | N/A | N/A | N/A | PA0862 | N/A | N/A | N/A | N/A | N/A | PA0862 | PA0862 | PA0862 | N/A | N/A | N/A | N/A | N/A |
| PA0863 | N/A | N/A | N/A | PA0863 | PA0863 | N/A | N/A | N/A | N/A | N/A | N/A | N/A | N/A | N/A | N/A | N/A | N/A | N/A |
| PA0864 | N/A | N/A | N/A | N/A | PA0864 | N/A | N/A | N/A | N/A | N/A | N/A | N/A | N/A | N/A | N/A | N/A | N/A | N/A |
| PA0865 | PA0865 | PA0865 | PA0865 | PA0865 | PA0865 | PA0865 | PA0865 | PA0865 | PA0865 | PA0865 | PA0865 | PA0865 | PA0865 | PA0865 | PA0865 | PA0865 | PA0865 | N/A |
| PA0866 | N/A | N/A | N/A | PA0866 | PA0866 | N/A | N/A | N/A | N/A | N/A | N/A | N/A | N/A | N/A | N/A | N/A | N/A | N/A |
| PA0867 | PA0867 | PA0867 | PA0867 | PA0867 | PA0867 | PA0867 | PA0867 | N/A | PA0867 | PA0867 | PA0867 | PA0867 | PA0867 | PA0867 | PA0867 | N/A | PA0867 | PA0867 |
| PA0869 | N/A | PA0869 | N/A | PA0869 | PA0869 | N/A | N/A | PA0869 | PA0869 | N/A | PA0869 | N/A | N/A | N/A | N/A | N/A | PA0869 | PA0869 |
| PA0870 | PA0870 | PA0870 | PA0870 | PA0870 | PA0870 | PA0870 | PA0870 | N/A | PA0870 | PA0870 | PA0870 | PA0870 | PA0870 | N/A | PA0870 | PA0870 | N/A | N/A |
| PA0871 | PA0871 | PA0871 | PA0871 | PA0871 | PA0871 | PA0871 | PA0871 | N/A | PA0871 | PA0871 | PA0871 | PA0871 | PA0871 | N/A | PA0871 | PA0871 | N/A | N/A |
| PA0872 | N/A | PA0872 | PA0872 | PA0872 | PA0872 | PA0872 | PA0872 | PA0872 | PA0872 | PA0872 | PA0872 | PA0872 | PA0872 | N/A | PA0872 | PA0872 | N/A | N/A |
| PA0873 | N/A | N/A | N/A | PA0873 | N/A | N/A | N/A | PA0873 | PA0873 | N/A | N/A | N/A | N/A | N/A | N/A | N/A | N/A | N/A |
| N/A | N/A | N/A | N/A | N/A | PA0875 | N/A | N/A | N/A | N/A | N/A | N/A | N/A | N/A | N/A | N/A | N/A | N/A | N/A |
| N/A | N/A | N/A | N/A | N/A | PA0877 | N/A | N/A | N/A | PA0877 | N/A | PA0877 | N/A | N/A | N/A | N/A | N/A | N/A | N/A |
| PA0876 | N/A | N/A | N/A | PA0876 | N/A | PA0876 | N/A | N/A | PA0876 | PA0876 | PA0876 | PA0876 | N/A | N/A | N/A | N/A | PA0876 | N/A |
| N/A | N/A | N/A | N/A | N/A | PA0878 | N/A | N/A | N/A | N/A | N/A | N/A | N/A | N/A | N/A | N/A | N/A | N/A | N/A |
| N/A | N/A | N/A | N/A | N/A | PA0879 | N/A | N/A | N/A | N/A | N/A | N/A | N/A | N/A | N/A | N/A | N/A | N/A | N/A |
| N/A | N/A | N/A | N/A | N/A | PA0880 | PA0880 | N/A | N/A | N/A | N/A | PA0880 | N/A | N/A | N/A | N/A | N/A | N/A | N/A |
| N/A | N/A | N/A | N/A | N/A | PA0883 | N/A | N/A | N/A | N/A | N/A | N/A | N/A | N/A | N/A | N/A | N/A | N/A | N/A |
| PA0884 | N/A | N/A | N/A | PA0884 | PA0884 | PA0884 | N/A | N/A | PA0884 | PA0884 | PA0884 | PA0884 | PA0884 | N/A | PA0884 | N/A | N/A | N/A |
| N/A | N/A | N/A | N/A | N/A | PA0886 | N/A | N/A | N/A | N/A | N/A | N/A | N/A | N/A | N/A | N/A | N/A | N/A | N/A |
| PA0887 | PA0887 | PA0887 | PA0887 | PA0887 | PA0887 | PA0887 | PA0887 | PA0887 | PA0887 | PA0887 | PA0887 | PA0887 | PA0887 | N/A | PA0887 | N/A | N/A | N/A |
| PA0888 | PA0888 | PA0888 | PA0888 | PA0888 | PA0888 | PA0888 | PA0888 | N/A | PA0888 | PA0888 | PA0888 | PA0888 | PA0888 | PA0888 | PA0888 | PA0888 | PA0888 | N/A |
| PA0889 | N/A | PA0889 | N/A | PA0889 | N/A | N/A | N/A | N/A | PA0889 | N/A | PA0889 | PA0889 | N/A | PA0889 | N/A | N/A | PA0889 | N/A |
| PA0890 | N/A | PA0890 | N/A | PA0890 | PA0890 | N/A | N/A | N/A | PA0890 | N/A | N/A | N/A | N/A | PA0890 | N/A | N/A | PA0890 | N/A |
| PA0891 | PA0891 | PA0891 | PA0891 | PA0891 | PA0891 | PA0891 | PA0891 | N/A | PA0891 | PA0891 | PA0891 | PA0891 | PA0891 | PA0891 | N/A | N/A | N/A | N/A |
| PA0892 | PA0892 | PA0892 | PA0892 | PA0892 | N/A | PA0892 | PA0892 | N/A | PA0892 | PA0892 | PA0892 | PA0892 | PA0892 | PA0892 | PA0892 | N/A | PA0892 | PA0892 |
| PA0893 | PA0893 | N/A | PA0893 | PA0893 | N/A | N/A | PA0893 | PA0893 | PA0893 | PA0893 | PA0893 | PA0893 | PA0893 | PA0893 | N/A | N/A | N/A | N/A |
| N/A | N/A | N/A | N/A | N/A | PA0894 | N/A | N/A | N/A | N/A | N/A | N/A | N/A | N/A | N/A | N/A | N/A | N/A | N/A |
| PA0895 | PA0895 | PA0895 | PA0895 | PA0895 | PA0895 | PA0895 | PA0895 | PA0895 | PA0895 | PA0895 | PA0895 | PA0895 | PA0895 | PA0895 | PA0895 | PA0895 | N/A | N/A |
| PA0896 | PA0896 | PA0896 | PA0896 | PA0896 | PA0896 | PA0896 | N/A | PA0896 | PA0896 | PA0896 | PA0896 | PA0896 | PA0896 | N/A | PA0896 | N/A | N/A | N/A |
| PA0897 | PA0897 | PA0897 | PA0897 | PA0897 | PA0897 | N/A | PA0897 | PA0897 | PA0897 | PA0897 | PA0897 | PA0897 | PA0897 | N/A | PA0897 | N/A | N/A | N/A |
| PA0898 | PA0898 | PA0898 | PA0898 | PA0898 | PA0898 | PA0898 | N/A | PA0898 | PA0898 | PA0898 | PA0898 | PA0898 | PA0898 | N/A | PA0898 | PA0898 | N/A | N/A |
| PA0899 | PA0899 | PA0899 | PA0899 | PA0899 | PA0899 | PA0899 | PA0899 | PA0899 | PA0899 | PA0899 | PA0899 | PA0899 | PA0899 | PA0899 | N/A | PA0899 | N/A | N/A |
| PA0900 | PA0900 | N/A | PA0900 | PA0900 | PA0900 | PA0900 | N/A | N/A | PA0900 | PA0900 | PA0900 | PA0900 | PA0900 | N/A | N/A | N/A | N/A | N/A |
| PA0901 | PA0901 | PA0901 | PA0901 | PA0901 | N/A | PA0901 | N/A | N/A | PA0901 | PA0901 | PA0901 | PA0901 | PA0901 | N/A | N/A | N/A | N/A | N/A |
| PA0902 | PA0902 | PA0902 | N/A | PA0902 | N/A | PA0902 | N/A | N/A | PA0902 | PA0902 | PA0902 | PA0902 | PA0902 | N/A | N/A | N/A | PA0902 | N/A |
| PA0903 | PA0903 | PA0903 | PA0903 | PA0903 | PA0903 | PA0903 | PA0903 | PA0903 | PA0903 | PA0903 | PA0903 | PA0903 | PA0903 | PA0903 | PA0903 | N/A | N/A | N/A |
| PA0904 | PA0904 | PA0904 | PA0904 | PA0904 | PA0904 | PA0904 | PA0904 | PA0904 | PA0904 | PA0904 | PA0904 | PA0904 | PA0904 | N/A | PA0904 | N/A | N/A | N/A |
| PA0905 | N/A | PA0905 | N/A | PA0905 | PA0905 | N/A | N/A | N/A | N/A | N/A | N/A | N/A | PA0905 | N/A | N/A | N/A | N/A | N/A |
| PA0907 | N/A | N/A | N/A | N/A | N/A | N/A | N/A | N/A | N/A | N/A | N/A | N/A | N/A | N/A | N/A | N/A | N/A | N/A |
| PA0910 | N/A | N/A | N/A | N/A | N/A | N/A | N/A | N/A | N/A | N/A | N/A | N/A | N/A | N/A | PA0910 | PA0910 | N/A | N/A |
| N/A | N/A | N/A | N/A | N/A | N/A | N/A | N/A | N/A | N/A | N/A | N/A | N/A | N/A | N/A | PA0911 | N/A | N/A | N/A |
| PA0913 | PA0913 | PA0913 | PA0913 | PA0913 | N/A | PA0913 | N/A | PA0913 | PA0913 | PA0913 | PA0913 | PA0913 | PA0913 | PA0913 | N/A | N/A | PA0913 | N/A |
| PA0915 | N/A | N/A | N/A | PA0915 | PA0915 | N/A | PA0915 | N/A | PA0915 | PA0915 | PA0915 | PA0915 | PA0915 | N/A | N/A | N/A | N/A | N/A |
| PA0916 | PA0916 | PA0916 | PA0916 | PA0916 | PA0916 | N/A | N/A | PA0916 | PA0916 | PA0916 | PA0916 | PA0916 | PA0916 | N/A | PA0916 | N/A | N/A | N/A |
| PA0917 | N/A | N/A | N/A | PA0917 | N/A | PA0917 | N/A | PA0917 | PA0917 | N/A | N/A | PA0917 | PA0917 | PA0917 | N/A | N/A | PA0917 | N/A |
| PA0918 | N/A | N/A | N/A | N/A | N/A | PA0918 | N/A | N/A | PA0918 | PA0918 | PA0918 | PA0918 | PA0918 | PA0918 | N/A | N/A | PA0918 | N/A |
| PA0919 | PA0919 | PA0919 | PA0919 | PA0919 | N/A | N/A | N/A | PA0919 | N/A | N/A | N/A | N/A | PA0919 | PA0919 | PA0919 | N/A | PA0919 | N/A |
| PA0920 | N/A | N/A | N/A | PA0920 | N/A | PA0920 | N/A | PA0920 | N/A | PA0920 | PA0920 | N/A | N/A | PA0920 | N/A | N/A | N/A | N/A |
| N/A | N/A | N/A | N/A | N/A | N/A | N/A | N/A | N/A | N/A | N/A | PA0922 | N/A | N/A | N/A | PA0922 | N/A | N/A | N/A |
| PA0925 | PA0925 | PA0925 | PA0925 | N/A | PA0925 | N/A | PA0925 | N/A | PA0925 | PA0925 | N/A | PA0925 | PA0925 | PA0925 | N/A | N/A | N/A | N/A |
| PA0926 | PA0926 | PA0926 | N/A | PA0926 | PA0926 | N/A | PA0926 | N/A | PA0926 | PA0926 | PA0926 | PA0926 | PA0926 | PA0926 | N/A | N/A | N/A | N/A |
| PA0927 | PA0927 | PA0927 | N/A | PA0927 | N/A | N/A | N/A | PA0927 | N/A | PA0927 | N/A | N/A | PA0927 | N/A | N/A | N/A | N/A | N/A |
| PA0928 | N/A | N/A | N/A | PA0928 | N/A | N/A | N/A | N/A | PA0928 | PA0928 | PA0928 | PA0928 | PA0928 | PA0928 | N/A | N/A | N/A | N/A |
| PA0929 | N/A | N/A | N/A | PA0929 | N/A | N/A | N/A | N/A | N/A | N/A | N/A | N/A | N/A | N/A | N/A | N/A | N/A | N/A |
| PA0930 | N/A | N/A | N/A | PA0930 | N/A | N/A | N/A | N/A | N/A | N/A | N/A | N/A | N/A | N/A | N/A | N/A | N/A | N/A |
| PA0931 | N/A | N/A | N/A | PA0931 | PA0931 | N/A | N/A | N/A | N/A | N/A | N/A | N/A | N/A | N/A | PA0931 | N/A | N/A | N/A |
| PA0932 | PA0932 | PA0932 | PA0932 | PA0932 | PA0932 | PA0932 | N/A | N/A | PA0932 | PA0932 | PA0932 | PA0932 | PA0932 | N/A | PA0932 | N/A | N/A | N/A |
| PA0933 | N/A | N/A | N/A | PA0933 | PA0933 | N/A | N/A | PA0933 | N/A | N/A | PA0933 | PA0933 | PA0933 | PA0933 | N/A | N/A | N/A | N/A |
| PA0934 | PA0934 | PA0934 | PA0934 | PA0934 | PA0934 | PA0934 | N/A | PA0934 | PA0934 | PA0934 | PA0934 | PA0934 | PA0934 | N/A | N/A | N/A | PA0934 | N/A |
| PA0935 | PA0935 | PA0935 | N/A | PA0935 | PA0935 | N/A | N/A | N/A | PA0935 | PA0935 | PA0935 | PA0935 | PA0935 | N/A | PA0935 | N/A | N/A | N/A |
| PA0936 | PA0936 | PA0936 | N/A | PA0936 | PA0936 | PA0936 | PA0936 | PA0936 | PA0936 | PA0936 | PA0936 | PA0936 | PA0936 | PA0936 | N/A | N/A | PA0936 | PA0936 |
| PA0937 | PA0937 | PA0937 | PA0937 | PA0937 | PA0937 | PA0937 | PA0937 | PA0937 | PA0937 | PA0937 | PA0937 | PA0937 | PA0937 | PA0937 | PA0937 | N/A | PA0937 | PA0937 |
| PA0938 | PA0938 | PA0938 | PA0938 | PA0938 | N/A | N/A | PA0938 | PA0938 | PA0938 | N/A | PA0938 | PA0938 | PA0938 | PA0938 | PA0938 | N/A | PA0938 | N/A |
| N/A | N/A | N/A | N/A | N/A | PA0940 | N/A | N/A | N/A | N/A | N/A | N/A | N/A | N/A | N/A | N/A | N/A | N/A | N/A |
| PA0941 | PA0941 | PA0941 | PA0941 | PA0941 | PA0941 | N/A | N/A | N/A | N/A | N/A | PA0941 | N/A | PA0941 | N/A | N/A | N/A | N/A | N/A |
| PA0942 | N/A | N/A | N/A | PA0942 | PA0942 | PA0942 | PA0942 | N/A | PA0942 | PA0942 | PA0942 | PA0942 | PA0942 | N/A | N/A | N/A | N/A | N/A |
| PA0943 | N/A | PA0943 | N/A | PA0943 | PA0943 | PA0943 | N/A | N/A | PA0943 | PA0943 | PA0943 | PA0943 | PA0943 | N/A | PA0943 | PA0943 | N/A | N/A |
| PA0944 | PA0944 | PA0944 | PA0944 | PA0944 | PA0944 | N/A | N/A | PA0944 | PA0944 | N/A | PA0944 | N/A | N/A | N/A | N/A | N/A | N/A | N/A |
| PA0945 | PA0945 | PA0945 | PA0945 | PA0945 | PA0945 | PA0945 | PA0945 | PA0945 | PA0945 | PA0945 | PA0945 | PA0945 | PA0945 | N/A | PA0945 | N/A | N/A | N/A |
| PA0946 | PA0946 | PA0946 | PA0946 | PA0946 | PA0946 | N/A | PA0946 | N/A | PA0946 | PA0946 | PA0946 | PA0946 | PA0946 | PA0946 | PA0946 | N/A | PA0946 | PA0946 |
| PA0947 | PA0947 | PA0947 | PA0947 | PA0947 | PA0947 | N/A | PA0947 | PA0947 | PA0947 | PA0947 | PA0947 | PA0947 | PA0947 | N/A | N/A | N/A | N/A | N/A |
| PA0948 | N/A | N/A | N/A | PA0948 | N/A | N/A | N/A | N/A | PA0948 | N/A | N/A | N/A | N/A | PA0948 | N/A | N/A | PA0948 | N/A |
| PA0949 | N/A | N/A | N/A | PA0949 | PA0949 | N/A | N/A | N/A | PA0949 | PA0949 | PA0949 | PA0949 | PA0949 | N/A | N/A | N/A | N/A | N/A |
| PA0950 | PA0950 | PA0950 | PA0950 | PA0950 | PA0950 | PA0950 | N/A | N/A | PA0950 | PA0950 | PA0950 | PA0950 | PA0950 | N/A | PA0950 | N/A | N/A | N/A |
| PA0951 | N/A | N/A | N/A | PA0951 | N/A | N/A | N/A | N/A | N/A | N/A | N/A | N/A | N/A | PA0951 | PA0951 | N/A | N/A | N/A |
| N/A | N/A | N/A | N/A | PA0952 | N/A | N/A | N/A | N/A | PA0952 | PA0952 | N/A | N/A | N/A | N/A | N/A | N/A | N/A | N/A |
| PA0953 | N/A | N/A | N/A | PA0953 | PA0953 | PA0953 | PA0953 | N/A | PA0953 | PA0953 | PA0953 | PA0953 | PA0953 | PA0953 | N/A | N/A | PA0953 | N/A |
| PA0954 | N/A | N/A | N/A | PA0954 | N/A | N/A | N/A | N/A | N/A | N/A | PA0954 | PA0954 | PA0954 | N/A | N/A | N/A | N/A | N/A |
| PA0955 | PA0955 | PA0955 | PA0955 | PA0955 | PA0955 | N/A | PA0955 | PA0955 | PA0955 | PA0955 | PA0955 | PA0955 | PA0955 | PA0955 | PA0955 | N/A | N/A | N/A |
| PA0956 | PA0956 | PA0956 | PA0956 | PA0956 | PA0956 | PA0956 | PA0956 | PA0956 | PA0956 | PA0956 | PA0956 | PA0956 | PA0956 | PA0956 | PA0956 | N/A | N/A | N/A |
| N/A | N/A | N/A | N/A | N/A | PA0957 | N/A | N/A | N/A | N/A | N/A | N/A | N/A | N/A | N/A | N/A | N/A | N/A | N/A |
| PA0958 | PA0958 | PA0958 | PA0958 | PA0958 | PA0958 | PA0958 | PA0958 | PA0958 | PA0958 | PA0958 | PA0958 | PA0958 | PA0958 | PA0958 | PA0958 | PA0958 | PA0958 | PA0958 |
| PA0959 | PA0959 | PA0959 | PA0959 | PA0959 | N/A | PA0959 | PA0959 | N/A | PA0959 | PA0959 | PA0959 | PA0959 | PA0959 | N/A | PA0959 | PA0959 | N/A | N/A |
| N/A | N/A | N/A | N/A | PA0960 | N/A | N/A | N/A | N/A | N/A | N/A | N/A | PA0960 | N/A | N/A | N/A | N/A | N/A | N/A |
| PA0961 | PA0961 | PA0961 | PA0961 | PA0961 | PA0961 | PA0961 | N/A | PA0961 | PA0961 | PA0961 | PA0961 | PA0961 | PA0961 | PA0961 | PA0961 | N/A | PA0961 | N/A |
| PA0962 | PA0962 | PA0962 | PA0962 | PA0962 | PA0962 | PA0962 | PA0962 | N/A | PA0962 | PA0962 | PA0962 | PA0962 | PA0962 | PA0962 | PA0962 | PA0962 | PA0962 | PA0962 |
| PA0963 | PA0963 | PA0963 | PA0963 | PA0963 | PA0963 | PA0963 | PA0963 | PA0963 | PA0963 | PA0963 | PA0963 | PA0963 | PA0963 | PA0963 | PA0963 | N/A | PA0963 | N/A |
| PA0964 | PA0964 | PA0964 | PA0964 | PA0964 | PA0964 | PA0964 | PA0964 | N/A | PA0964 | PA0964 | PA0964 | PA0964 | PA0964 | N/A | PA0964 | N/A | N/A | N/A |
| PA0965 | N/A | N/A | N/A | PA0965 | N/A | N/A | PA0965 | N/A | PA0965 | N/A | PA0965 | PA0965 | PA0965 | PA0965 | N/A | N/A | N/A | N/A |
| PA0966 | PA0966 | PA0966 | N/A | PA0966 | N/A | N/A | PA0966 | N/A | PA0966 | PA0966 | PA0966 | PA0966 | PA0966 | PA0966 | N/A | N/A | N/A | N/A |
| PA0967 | N/A | PA0967 | PA0967 | PA0967 | N/A | N/A | PA0967 | PA0967 | PA0967 | PA0967 | PA0967 | N/A | PA0967 | N/A | N/A | N/A | N/A | N/A |
| PA0968 | PA0968 | PA0968 | PA0968 | PA0968 | PA0968 | N/A | N/A | N/A | PA0968 | PA0968 | PA0968 | PA0968 | PA0968 | PA0968 | PA0968 | N/A | N/A | PA0968 |
| PA0969 | PA0969 | PA0969 | PA0969 | PA0969 | PA0969 | PA0969 | PA0969 | PA0969 | PA0969 | PA0969 | PA0969 | PA0969 | PA0969 | PA0969 | PA0969 | N/A | PA0969 | PA0969 |
| PA0970 | PA0970 | PA0970 | PA0970 | PA0970 | N/A | PA0970 | PA0970 | PA0970 | PA0970 | PA0970 | PA0970 | PA0970 | PA0970 | PA0970 | PA0970 | N/A | PA0970 | N/A |
| PA0971 | PA0971 | N/A | N/A | PA0971 | PA0971 | PA0971 | N/A | PA0971 | PA0971 | PA0971 | PA0971 | PA0971 | N/A | PA0971 | PA0971 | N/A | PA0971 | N/A |
| PA0972 | PA0972 | PA0972 | PA0972 | PA0972 | PA0972 | PA0972 | PA0972 | PA0972 | PA0972 | PA0972 | PA0972 | PA0972 | PA0972 | PA0972 | PA0972 | PA0972 | PA0972 | PA0972 |
| PA0973 | PA0973 | PA0973 | PA0973 | PA0973 | PA0973 | PA0973 | PA0973 | PA0973 | PA0973 | PA0973 | PA0973 | PA0973 | PA0973 | PA0973 | PA0973 | PA0973 | PA0973 | PA0973 |
| PA0974 | PA0974 | PA0974 | PA0974 | PA0974 | PA0974 | PA0974 | PA0974 | N/A | PA0974 | PA0974 | PA0974 | PA0974 | PA0974 | PA0974 | PA0974 | N/A | PA0974 | PA0974 |
| PA0975 | N/A | N/A | N/A | PA0975 | N/A | N/A | N/A | N/A | PA0975 | N/A | N/A | N/A | N/A | N/A | N/A | N/A | N/A | N/A |
| PA0976 | N/A | PA0976 | PA0976 | PA0976 | N/A | PA0976 | N/A | N/A | PA0976 | N/A | PA0976 | N/A | PA0976 | N/A | N/A | N/A | N/A | N/A |
| PA0980 | N/A | N/A | N/A | N/A | PA0980 | N/A | N/A | N/A | N/A | N/A | N/A | N/A | N/A | N/A | N/A | N/A | N/A | N/A |

| | | | | | | | | | | | | | | | | | | |
|---|---|---|---|---|---|---|---|---|---|---|---|---|---|---|---|---|---|---|
| PA0981 | N/A | N/A | N/A | N/A | N/A | N/A | N/A | N/A | PA0981 | N/A | N/A | N/A | N/A | N/A | N/A | N/A | N/A | N/A |
| PA0984 | N/A | N/A | N/A | N/A | N/A | N/A | N/A | N/A | N/A | N/A | N/A | N/A | N/A | PA0984 | N/A | N/A | N/A | N/A |
| N/A | N/A | N/A | N/A | N/A | N/A | N/A | N/A | N/A | N/A | N/A | N/A | N/A | N/A | PA0985 | PA0985 | PA0985 | N/A | N/A |
| PA0988 | N/A | N/A | N/A | N/A | N/A | N/A | N/A | N/A | N/A | N/A | PA0988 | N/A | PA0988 | N/A | N/A | N/A | N/A | N/A |
| PA0989 | N/A | N/A | N/A | PA0989 | N/A | N/A | N/A | N/A | N/A | N/A | N/A | N/A | N/A | N/A | PA0989 | N/A | N/A | PA0989 |
| PA0991 | N/A | N/A | N/A | PA0991 | N/A | N/A | N/A | N/A | PA0991 | N/A | PA0991 | PA0991 | PA0991 | N/A | N/A | N/A | N/A | N/A |
| PA0995 | N/A | N/A | N/A | N/A | PA0995 | N/A | N/A | N/A | N/A | N/A | PA0995 | N/A | N/A | N/A | N/A | N/A | N/A | N/A |
| PA0996 | N/A | N/A | N/A | PA0996 | PA0996 | N/A | N/A | N/A | PA0996 | N/A | PA0996 | PA0996 | PA0996 | PA0996 | N/A | N/A | N/A | N/A |
| PA0997 | N/A | N/A | N/A | PA0997 | PA0997 | PA0997 | N/A | N/A | PA0997 | PA0997 | PA0997 | PA0997 | PA0997 | N/A | N/A | N/A | N/A | N/A |
| PA0998 | N/A | N/A | N/A | PA0998 | PA0998 | PA0998 | N/A | N/A | PA0998 | PA0998 | PA0998 | PA0998 | PA0998 | N/A | N/A | PA0998 | N/A | N/A |
| PA0999 | N/A | N/A | N/A | PA0999 | PA0999 | PA0999 | N/A | N/A | PA0999 | PA0999 | PA0999 | PA0999 | PA0999 | N/A | N/A | N/A | N/A | N/A |
| PA1000 | N/A | N/A | N/A | N/A | PA1000 | N/A | N/A | N/A | PA1000 | N/A | PA1000 | PA1000 | PA1000 | N/A | N/A | N/A | N/A | N/A |
| PA1001 | N/A | N/A | N/A | N/A | N/A | N/A | N/A | N/A | PA1001 | N/A | PA1001 | PA1001 | N/A | N/A | N/A | N/A | N/A | N/A |
| PA1002 | N/A | N/A | N/A | N/A | PA1002 | N/A | N/A | N/A | PA1002 | N/A | PA1002 | PA1002 | PA1002 | N/A | N/A | N/A | N/A | N/A |
| PA1003 | N/A | N/A | N/A | PA1003 | PA1003 | N/A | N/A | PA1003 | PA1003 | N/A | PA1003 | PA1003 | PA1003 | PA1003 | N/A | N/A | PA1003 | N/A |
| PA1004 | PA1004 | PA1004 | PA1004 | PA1004 | PA1004 | PA1004 | PA1004 | PA1004 | PA1004 | PA1004 | PA1004 | PA1004 | PA1004 | N/A | PA1004 | N/A | N/A | N/A |
| PA1005 | N/A | N/A | N/A | PA1005 | PA1005 | N/A | N/A | N/A | N/A | N/A | PA1005 | PA1005 | PA1005 | PA1005 | PA1005 | N/A | PA1005 | PA1005 |
| PA1006 | N/A | PA1006 | N/A | N/A | N/A | N/A | N/A | N/A | PA1006 | N/A | PA1006 | N/A | N/A | N/A | N/A | N/A | N/A | N/A |
| N/A | N/A | N/A | N/A | N/A | PA1007 | N/A | N/A | N/A | N/A | N/A | N/A | N/A | N/A | PA1007 | N/A | N/A | N/A | N/A |
| PA1008 | PA1008 | PA1008 | PA1008 | PA1008 | PA1008 | PA1008 | PA1008 | N/A | PA1008 | PA1008 | PA1008 | PA1008 | PA1008 | PA1008 | PA1008 | N/A | N/A | N/A |
| PA1009 | PA1009 | PA1009 | PA1009 | PA1009 | PA1009 | PA1009 | N/A | N/A | PA1009 | PA1009 | PA1009 | PA1009 | PA1009 | N/A | N/A | N/A | N/A | N/A |
| PA1010 | PA1010 | PA1010 | PA1010 | PA1010 | PA1010 | PA1010 | PA1010 | N/A | PA1010 | PA1010 | PA1010 | PA1010 | PA1010 | PA1010 | PA1010 | PA1010 | N/A | N/A |
| PA1011 | PA1011 | PA1011 | PA1011 | PA1011 | PA1011 | PA1011 | PA1011 | PA1011 | PA1011 | PA1011 | PA1011 | PA1011 | PA1011 | PA1011 | PA1011 | N/A | PA1011 | PA1011 |
| N/A | N/A | N/A | N/A | N/A | PA1012 | N/A | N/A | N/A | N/A | N/A | N/A | N/A | N/A | N/A | N/A | N/A | N/A | N/A |
| PA1013 | PA1013 | PA1013 | PA1013 | PA1013 | PA1013 | PA1013 | PA1013 | PA1013 | PA1013 | PA1013 | PA1013 | PA1013 | PA1013 | N/A | PA1013 | PA1013 | N/A | N/A |
| PA1014 | PA1014 | PA1014 | N/A | PA1014 | N/A | N/A | PA1014 | PA1014 | PA1014 | N/A | PA1014 | N/A | N/A | PA1014 | PA1014 | N/A | PA1014 | N/A |
| PA1015 | N/A | N/A | PA1015 | PA1015 | N/A | N/A | PA1015 | N/A | PA1015 | N/A | PA1015 | PA1015 | N/A | N/A | N/A | N/A | N/A | N/A |
| PA1023 | N/A | N/A | N/A | PA1023 | N/A | N/A | N/A | N/A | N/A | PA1023 | PA1023 | PA1023 | PA1023 | N/A | N/A | N/A | N/A | N/A |
| N/A | N/A | N/A | N/A | PA1024 | N/A | N/A | N/A | N/A | N/A | N/A | N/A | N/A | N/A | N/A | N/A | N/A | N/A | N/A |
| N/A | N/A | N/A | N/A | N/A | N/A | N/A | N/A | PA1025 | PA1025 | PA1025 | N/A | N/A | N/A | N/A | N/A | N/A | N/A | N/A |
| PA1026 | N/A | N/A | N/A | PA1026 | N/A | N/A | N/A | N/A | N/A | N/A | N/A | PA1026 | N/A | N/A | N/A | N/A | N/A | N/A |
| PA1027 | N/A | N/A | N/A | PA1027 | N/A | N/A | N/A | N/A | N/A | PA1027 | N/A | PA1027 | PA1027 | N/A | N/A | N/A | N/A | N/A |
| N/A | N/A | N/A | N/A | N/A | N/A | N/A | N/A | N/A | N/A | N/A | N/A | PA1030 | N/A | N/A | N/A | N/A | N/A | N/A |
| PA1031 | N/A | N/A | N/A | PA1031 | N/A | N/A | N/A | PA1031 | N/A | N/A | N/A | N/A | N/A | PA1031 | N/A | N/A | N/A | N/A |
| PA1032 | N/A | PA1032 | N/A | PA1032 | PA1032 | PA1032 | N/A | PA1032 | N/A | PA1032 | N/A | PA1032 | PA1032 | PA1032 | PA1032 | N/A | N/A | N/A |
| PA1033 | N/A | N/A | N/A | N/A | PA1033 | PA1033 | N/A | N/A | PA1033 | PA1033 | PA1033 | PA1033 | PA1033 | N/A | N/A | N/A | N/A | N/A |
| PA1035 | N/A | N/A | N/A | PA1035 | N/A | PA1035 | PA1035 | N/A | PA1035 | PA1035 | PA1035 | PA1035 | PA1035 | PA1035 | N/A | N/A | N/A | N/A |
| PA1036 | N/A | PA1036 | N/A | PA1036 | N/A | PA1036 | N/A | N/A | N/A | PA1036 | N/A | PA1036 | PA1036 | N/A | N/A | N/A | N/A | N/A |
| PA1039 | N/A | N/A | N/A | PA1039 | N/A | N/A | PA1039 | N/A | PA1039 | PA1039 | PA1039 | PA1039 | PA1039 | N/A | N/A | N/A | N/A | N/A |
| PA1040 | N/A | N/A | N/A | PA1040 | PA1040 | N/A | N/A | N/A | PA1040 | PA1040 | PA1040 | PA1040 | PA1040 | N/A | N/A | N/A | N/A | N/A |
| PA1041 | N/A | N/A | N/A | N/A | PA1041 | PA1041 | N/A | N/A | PA1041 | PA1041 | PA1041 | PA1041 | PA1041 | PA1041 | N/A | PA1041 | PA1041 | N/A |
| N/A | N/A | N/A | N/A | N/A | PA1042 | N/A | N/A | N/A | N/A | N/A | N/A | N/A | N/A | N/A | N/A | N/A | N/A | N/A |
| PA1043 | PA1043 | N/A | N/A | PA1043 | PA1043 | N/A | PA1043 | N/A | N/A | N/A | PA1043 | PA1043 | N/A | PA1043 | N/A | N/A | N/A | N/A |
| PA1044 | N/A | N/A | N/A | N/A | N/A | N/A | N/A | N/A | N/A | N/A | N/A | PA1044 | N/A | PA1044 | N/A | N/A | N/A | N/A |
| PA1045 | PA1045 | PA1045 | N/A | PA1045 | N/A | N/A | N/A | PA1045 | PA1045 | PA1045 | PA1045 | PA1045 | N/A | N/A | PA1045 | N/A | PA1045 | N/A |
| PA1046 | N/A | N/A | N/A | N/A | PA1046 | N/A | N/A | N/A | N/A | N/A | N/A | N/A | N/A | N/A | N/A | N/A | N/A | N/A |
| PA1047 | PA1047 | N/A | N/A | PA1047 | PA1047 | N/A | N/A | N/A | N/A | PA1047 | N/A | PA1047 | PA1047 | N/A | PA1047 | N/A | N/A | N/A |
| PA1048 | PA1048 | N/A | N/A | PA1048 | PA1048 | N/A | N/A | N/A | N/A | N/A | N/A | PA1048 | PA1048 | PA1048 | PA1048 | N/A | N/A | N/A |
| PA1049 | N/A | PA1049 | N/A | PA1049 | PA1049 | PA1049 | PA1049 | N/A | PA1049 | PA1049 | PA1049 | PA1049 | PA1049 | N/A | PA1049 | N/A | PA1049 | N/A |
| PA1050 | N/A | N/A | N/A | N/A | N/A | N/A | N/A | N/A | N/A | PA1050 | N/A | N/A | N/A | N/A | N/A | N/A | N/A | N/A |
| PA1051 | N/A | N/A | N/A | PA1051 | N/A | N/A | N/A | PA1051 | PA1051 | N/A | N/A | PA1051 | N/A | PA1051 | N/A | N/A | N/A | N/A |
| PA1052 | N/A | N/A | N/A | PA1052 | PA1052 | N/A | N/A | N/A | N/A | PA1052 | N/A | N/A | N/A | N/A | N/A | N/A | N/A | N/A |
| PA1053 | PA1053 | PA1053 | PA1053 | PA1053 | PA1053 | PA1053 | PA1053 | PA1053 | PA1053 | PA1053 | PA1053 | PA1053 | PA1053 | PA1053 | PA1053 | PA1053 | PA1053 | PA1053 |
| PA1054 | N/A | N/A | N/A | N/A | N/A | N/A | N/A | N/A | PA1054 | N/A | N/A | N/A | N/A | PA1054 | N/A | N/A | PA1054 | N/A |
| PA1055 | N/A | N/A | N/A | N/A | N/A | N/A | N/A | N/A | N/A | N/A | N/A | N/A | N/A | N/A | N/A | N/A | N/A | N/A |
| PA1056 | N/A | N/A | N/A | PA1056 | N/A | N/A | N/A | N/A | N/A | N/A | N/A | N/A | N/A | PA1056 | N/A | N/A | PA1056 | N/A |
| PA1057 | N/A | PA1057 | N/A | N/A | N/A | PA1057 | N/A | N/A | PA1057 | N/A | N/A | N/A | N/A | N/A | N/A | N/A | N/A | N/A |
| PA1059 | N/A | N/A | N/A | N/A | N/A | N/A | N/A | N/A | N/A | PA1059 | PA1059 | PA1059 | N/A | PA1059 | N/A | N/A | N/A | N/A |
| PA1060 | N/A | N/A | N/A | N/A | PA1060 | N/A | N/A | N/A | N/A | N/A | N/A | N/A | N/A | N/A | N/A | N/A | N/A | N/A |
| PA1061 | N/A | N/A | N/A | PA1061 | N/A | PA1061 | N/A | N/A | PA1061 | PA1061 | PA1061 | PA1061 | PA1061 | N/A | N/A | N/A | N/A | N/A |
| PA1062 | N/A | N/A | N/A | PA1062 | N/A | N/A | N/A | N/A | PA1062 | N/A | PA1062 | PA1062 | N/A | N/A | N/A | N/A | N/A | N/A |
| PA1064 | PA1064 | PA1064 | PA1064 | PA1064 | N/A | PA1064 | PA1064 | N/A | PA1064 | PA1064 | PA1064 | PA1064 | PA1064 | PA1064 | PA1064 | N/A | PA1064 | N/A |
| PA1065 | N/A | N/A | N/A | N/A | PA1065 | N/A | N/A | N/A | N/A | PA1065 | PA1065 | PA1065 | N/A | N/A | N/A | N/A | N/A | N/A |
| PA1066 | N/A | N/A | N/A | N/A | PA1066 | PA1066 | N/A | N/A | PA1066 | PA1066 | PA1066 | PA1066 | PA1066 | PA1066 | N/A | N/A | PA1066 | N/A |
| PA1067 | N/A | N/A | N/A | N/A | PA1067 | N/A | N/A | N/A | N/A | N/A | N/A | N/A | N/A | N/A | N/A | N/A | N/A | N/A |
| PA1068 | PA1068 | PA1068 | PA1068 | PA1068 | PA1068 | N/A | PA1068 | PA1068 | PA1068 | N/A | PA1068 | PA1068 | PA1068 | PA1068 | N/A | N/A | PA1068 | PA1068 |
| PA1069 | PA1069 | PA1069 | PA1069 | PA1069 | PA1069 | PA1069 | PA1069 | PA1069 | PA1069 | PA1069 | PA1069 | PA1069 | PA1069 | PA1069 | PA1069 | PA1069 | PA1069 | N/A |
| PA1070 | PA1070 | PA1070 | N/A | PA1070 | N/A | N/A | PA1070 | N/A | PA1070 | PA1070 | PA1070 | PA1070 | PA1070 | PA1070 | PA1070 | N/A | PA1070 | PA1070 |
| PA1071 | PA1071 | PA1071 | PA1071 | PA1071 | N/A | PA1071 | N/A | N/A | PA1071 | N/A | PA1071 | PA1071 | PA1071 | PA1071 | N/A | N/A | PA1071 | PA1071 |
| PA1072 | N/A | N/A | N/A | N/A | N/A | PA1072 | N/A | N/A | N/A | PA1072 | PA1072 | PA1072 | PA1072 | PA1072 | N/A | N/A | PA1072 | N/A |
| N/A | N/A | N/A | N/A | N/A | PA1073 | N/A | N/A | N/A | N/A | N/A | N/A | N/A | N/A | PA1073 | N/A | N/A | N/A | N/A |
| PA1074 | PA1074 | PA1074 | PA1074 | PA1074 | PA1074 | PA1074 | PA1074 | N/A | PA1074 | PA1074 | PA1074 | PA1074 | PA1074 | PA1074 | PA1074 | PA1074 | PA1074 | N/A |
| PA1075 | N/A | N/A | PA1075 | PA1075 | PA1075 | PA1075 | N/A | N/A | N/A | N/A | PA1075 | N/A | N/A | N/A | N/A | N/A | N/A | N/A |
| PA1076 | PA1076 | PA1076 | PA1076 | PA1076 | PA1076 | PA1076 | N/A | N/A | PA1076 | PA1076 | PA1076 | PA1076 | PA1076 | PA1076 | N/A | N/A | N/A | N/A |
| PA1077 | N/A | N/A | N/A | N/A | N/A | N/A | N/A | N/A | N/A | N/A | N/A | N/A | N/A | N/A | N/A | N/A | N/A | N/A |
| PA1079 | N/A | N/A | N/A | PA1079 | PA1079 | N/A | N/A | N/A | PA1079 | N/A | N/A | N/A | N/A | N/A | PA1079 | PA1079 | N/A | N/A |
| PA1080 | PA1080 | PA1080 | PA1080 | PA1080 | PA1080 | PA1080 | PA1080 | PA1080 | PA1080 | PA1080 | PA1080 | PA1080 | PA1080 | N/A | PA1080 | PA1080 | N/A | N/A |
| N/A | N/A | N/A | N/A | N/A | N/A | N/A | N/A | N/A | N/A | N/A | N/A | N/A | N/A | N/A | PA1081 | N/A | N/A | N/A |
| PA1082 | N/A | N/A | N/A | N/A | N/A | N/A | PA1082 | N/A | N/A | N/A | N/A | N/A | N/A | N/A | PA1082 | PA1082 | N/A | N/A |
| N/A | PA1083 | PA1083 | PA1083 | PA1083 | N/A | PA1083 | PA1083 | N/A | PA1083 | PA1083 | PA1083 | N/A | PA1083 | PA1083 | PA1083 | N/A | N/A | N/A |
| PA1084 | N/A | PA1084 | PA1084 | PA1084 | N/A | PA1084 | PA1084 | PA1084 | PA1084 | PA1084 | PA1084 | N/A | N/A | N/A | N/A | N/A | N/A | N/A |
| N/A | N/A | N/A | N/A | N/A | PA1085 | N/A | N/A | N/A | N/A | N/A | N/A | N/A | N/A | N/A | N/A | N/A | N/A | N/A |
| PA1086 | N/A | N/A | N/A | PA1086 | PA1086 | N/A | PA1086 | PA1086 | PA1086 | PA1086 | N/A | N/A | PA1086 | N/A | PA1086 | PA1086 | N/A | N/A |
| PA1087 | N/A | N/A | N/A | PA1087 | N/A | N/A | N/A | N/A | N/A | N/A | N/A | N/A | N/A | N/A | PA1087 | PA1087 | N/A | N/A |
| PA1088 | N/A | N/A | N/A | N/A | N/A | N/A | N/A | N/A | N/A | N/A | N/A | N/A | N/A | N/A | N/A | N/A | N/A | N/A |
| PA1089 | N/A | N/A | N/A | PA1089 | N/A | N/A | N/A | N/A | N/A | N/A | N/A | N/A | N/A | N/A | N/A | N/A | N/A | N/A |
| PA1090 | N/A | N/A | N/A | PA1090 | N/A | N/A | N/A | N/A | PA1090 | PA1090 | PA1090 | PA1090 | PA1090 | N/A | N/A | N/A | N/A | N/A |
| PA1091 | N/A | PA1091 | PA1091 | PA1091 | PA1091 | N/A | N/A | N/A | PA1091 | PA1091 | PA1091 | PA1091 | PA1091 | PA1091 | N/A | N/A | N/A | N/A |
| PA1092 | PA1092 | PA1092 | PA1092 | PA1092 | PA1092 | PA1092 | PA1092 | PA1092 | PA1092 | PA1092 | PA1092 | PA1092 | PA1092 | PA1092 | PA1092 | PA1092 | PA1092 | PA1092 |
| PA1093 | N/A | N/A | N/A | PA1093 | PA1093 | N/A | N/A | N/A | PA1093 | N/A | PA1093 | PA1093 | PA1093 | N/A | PA1093 | N/A | N/A | N/A |
| PA1094 | PA1094 | PA1094 | N/A | PA1094 | PA1094 | PA1094 | PA1094 | PA1094 | PA1094 | PA1094 | PA1094 | PA1094 | PA1094 | PA1094 | PA1094 | PA1094 | PA1094 | PA1094 |
| PA1095 | N/A | PA1095 | PA1095 | PA1095 | N/A | PA1095 | PA1095 | N/A | PA1095 | N/A | PA1095 | PA1095 | PA1095 | PA1095 | PA1095 | N/A | N/A | N/A |
| PA1096 | PA1096 | N/A | PA1096 | PA1096 | PA1096 | N/A | N/A | N/A | N/A | N/A | PA1096 | PA1096 | PA1096 | N/A | N/A | N/A | N/A | N/A |
| PA1097 | PA1097 | PA1097 | PA1097 | PA1097 | PA1097 | PA1097 | PA1097 | PA1097 | PA1097 | PA1097 | PA1097 | PA1097 | PA1097 | PA1097 | PA1097 | N/A | PA1097 | N/A |
| N/A | N/A | N/A | N/A | N/A | N/A | N/A | N/A | PA1098 | N/A | N/A | N/A | N/A | N/A | N/A | N/A | N/A | N/A | N/A |
| N/A | PA1099 | PA1099 | PA1099 | PA1099 | N/A | N/A | N/A | PA1099 | PA1099 | N/A | N/A | N/A | N/A | N/A | N/A | N/A | N/A | N/A |
| PA1101 | PA1101 | PA1101 | N/A | PA1101 | N/A | N/A | PA1101 | PA1101 | PA1101 | PA1101 | PA1101 | PA1101 | N/A | PA1101 | PA1101 | N/A | PA1101 | N/A |
| PA1102 | PA1102 | PA1102 | PA1102 | PA1102 | N/A | N/A | PA1102 | PA1102 | PA1102 | PA1102 | N/A | PA1102 | PA1102 | PA1102 | N/A | N/A | N/A | N/A |
| PA1103 | N/A | N/A | N/A | PA1103 | N/A | N/A | N/A | N/A | N/A | PA1103 | N/A | N/A | N/A | N/A | N/A | N/A | N/A | N/A |
| PA1104 | N/A | N/A | N/A | PA1104 | N/A | N/A | N/A | N/A | N/A | N/A | N/A | N/A | N/A | N/A | N/A | N/A | N/A | N/A |
| N/A | N/A | N/A | N/A | PA1110 | N/A | N/A | N/A | N/A | PA1110 | N/A | N/A | N/A | N/A | PA1110 | N/A | N/A | N/A | N/A |
| N/A | N/A | N/A | N/A | N/A | PA1111 | N/A | N/A | N/A | N/A | N/A | N/A | N/A | N/A | N/A | N/A | N/A | N/A | N/A |
| PA1112 | N/A | N/A | N/A | N/A | PA1112 | N/A | N/A | N/A | N/A | N/A | N/A | N/A | PA1112 | N/A | N/A | N/A | N/A | N/A |
| PA1113 | N/A | N/A | PA1113 | N/A | N/A | N/A | N/A | N/A | N/A | PA1113 | N/A | N/A | N/A | PA1113 | N/A | N/A | N/A | N/A |
| N/A | N/A | N/A | N/A | N/A | N/A | N/A | N/A | N/A | N/A | N/A | N/A | N/A | N/A | PA1115 | N/A | N/A | N/A | N/A |
| PA1116 | N/A | PA1116 | N/A | PA1116 | N/A | N/A | N/A | N/A | PA1116 | N/A | PA1116 | PA1116 | N/A | N/A | N/A | N/A | N/A | N/A |
| PA1117 | N/A | N/A | N/A | PA1117 | N/A | N/A | N/A | N/A | PA1117 | N/A | N/A | N/A | PA1117 | N/A | N/A | N/A | N/A | N/A |
| PA1118 | N/A | N/A | N/A | N/A | N/A | N/A | N/A | N/A | N/A | N/A | N/A | N/A | N/A | N/A | N/A | N/A | N/A | N/A |
| PA1119 | PA1119 | N/A | PA1119 | PA1119 | N/A | PA1119 | PA1119 | N/A | PA1119 | PA1119 | PA1119 | PA1119 | PA1119 | PA1119 | PA1119 | N/A | N/A | N/A |

| | | | | | | | | | | | | | | | | | | |
|---|---|---|---|---|---|---|---|---|---|---|---|---|---|---|---|---|---|---|
| PA1120 | N/A | N/A | N/A | PA1120 | N/A | N/A | N/A | N/A | N/A | N/A | PA1120 | PA1120 | N/A | PA1120 | N/A | N/A | N/A | N/A |
| PA1121 | N/A | N/A | N/A | N/A | PA1121 | N/A | N/A | N/A | N/A | PA1121 | N/A | PA1121 | PA1121 | PA1121 | N/A | N/A | N/A | N/A |
| PA1122 | N/A | PA1122 | N/A | PA1122 | PA1122 | PA1122 | PA1122 | N/A | PA1122 | PA1122 | PA1122 | PA1122 | PA1122 | N/A | N/A | N/A | N/A | N/A |
| PA1123 | N/A | N/A | N/A | N/A | PA1123 | PA1123 | N/A | N/A | PA1123 | PA1123 | PA1123 | PA1123 | PA1123 | N/A | N/A | N/A | N/A | N/A |
| PA1124 | N/A | N/A | N/A | PA1124 | PA1124 | PA1124 | N/A | N/A | PA1124 | PA1124 | PA1124 | PA1124 | PA1124 | N/A | N/A | N/A | N/A | N/A |
| PA1125 | N/A | N/A | N/A | N/A | N/A | N/A | N/A | N/A | N/A | N/A | N/A | N/A | N/A | N/A | N/A | N/A | N/A | N/A |
| PA1126 | N/A | N/A | N/A | PA1126 | PA1126 | N/A | PA1126 | N/A | PA1126 | PA1126 | PA1126 | N/A | N/A | PA1126 | N/A | N/A | PA1126 | N/A |
| PA1127 | PA1127 | PA1127 | PA1127 | PA1127 | PA1127 | PA1127 | PA1127 | PA1127 | PA1127 | PA1127 | PA1127 | PA1127 | PA1127 | N/A | PA1127 | PA1127 | N/A | N/A |
| PA1128 | N/A | N/A | N/A | N/A | N/A | N/A | N/A | N/A | N/A | N/A | PA1128 | N/A | N/A | N/A | N/A | N/A | N/A | N/A |
| PA1129 | N/A | N/A | N/A | N/A | N/A | N/A | N/A | N/A | N/A | N/A | PA1129 | N/A | N/A | N/A | N/A | N/A | N/A | N/A |
| PA1130 | N/A | N/A | N/A | N/A | N/A | N/A | N/A | N/A | N/A | N/A | N/A | N/A | PA1130 | N/A | N/A | N/A | N/A | N/A |
| N/A | N/A | N/A | N/A | N/A | PA1134 | N/A | N/A | N/A | N/A | N/A | N/A | N/A | N/A | N/A | N/A | N/A | N/A | N/A |
| PA1135 | N/A | N/A | N/A | PA1135 | PA1135 | PA1135 | N/A | N/A | PA1135 | PA1135 | PA1135 | PA1135 | PA1135 | N/A | N/A | N/A | N/A | N/A |
| N/A | N/A | N/A | N/A | PA1137 | N/A | N/A | N/A | N/A | N/A | PA1137 | N/A | N/A | N/A | N/A | N/A | N/A | N/A | N/A |
| N/A | N/A | N/A | N/A | N/A | PA1138 | N/A | N/A | N/A | N/A | N/A | N/A | N/A | N/A | N/A | N/A | N/A | N/A | N/A |
| PA1139 | N/A | N/A | N/A | PA1139 | N/A | N/A | N/A | N/A | N/A | PA1139 | N/A | N/A | N/A | N/A | N/A | N/A | N/A | N/A |
| PA1140 | N/A | N/A | N/A | PA1140 | PA1140 | N/A | N/A | N/A | PA1140 | PA1140 | PA1140 | PA1140 | PA1140 | PA1140 | N/A | N/A | N/A | N/A |
| PA1141 | N/A | N/A | N/A | PA1141 | N/A | N/A | N/A | N/A | N/A | N/A | N/A | N/A | N/A | N/A | N/A | N/A | N/A | N/A |
| PA1142 | N/A | N/A | N/A | PA1142 | N/A | N/A | N/A | N/A | N/A | N/A | N/A | N/A | N/A | N/A | N/A | N/A | N/A | N/A |
| N/A | N/A | N/A | N/A | N/A | N/A | N/A | N/A | N/A | N/A | N/A | N/A | N/A | N/A | N/A | PA1148 | PA1148 | N/A | N/A |
| PA1150 | N/A | N/A | N/A | PA1150 | N/A | N/A | N/A | N/A | N/A | N/A | N/A | N/A | N/A | PA1150 | PA1150 | N/A | PA1150 | N/A |
| PA1151 | PA1151 | PA1151 | PA1151 | PA1151 | N/A | PA1151 | N/A | N/A | PA1151 | PA1151 | PA1151 | PA1151 | PA1151 | N/A | PA1151 | N/A | N/A | N/A |
| N/A | N/A | N/A | N/A | N/A | PA1154 | N/A | N/A | N/A | N/A | N/A | N/A | N/A | N/A | N/A | N/A | N/A | N/A | N/A |
| PA1155 | PA1155 | PA1155 | PA1155 | PA1155 | PA1155 | PA1155 | PA1155 | PA1155 | PA1155 | PA1155 | PA1155 | PA1155 | PA1155 | PA1155 | PA1155 | N/A | PA1155 | N/A |
| PA1156 | PA1156 | PA1156 | PA1156 | PA1156 | PA1156 | PA1156 | PA1156 | PA1156 | PA1156 | PA1156 | PA1156 | PA1156 | PA1156 | PA1156 | PA1156 | PA1156 | PA1156 | PA1156 |
| PA1157 | N/A | N/A | N/A | PA1157 | N/A | PA1157 | N/A | N/A | PA1157 | PA1157 | PA1157 | PA1157 | PA1157 | N/A | PA1157 | N/A | N/A | N/A |
| PA1158 | N/A | N/A | N/A | N/A | PA1158 | N/A | N/A | N/A | N/A | N/A | N/A | N/A | N/A | PA1158 | N/A | N/A | N/A | N/A |
| PA1159 | PA1159 | PA1159 | PA1159 | PA1159 | N/A | PA1159 | PA1159 | N/A | PA1159 | PA1159 | PA1159 | PA1159 | PA1159 | N/A | PA1159 | N/A | N/A | N/A |
| PA1160 | PA1160 | N/A | PA1160 | PA1160 | PA1160 | N/A | PA1160 | N/A | PA1160 | PA1160 | PA1160 | PA1160 | N/A | N/A | PA1160 | N/A | N/A | N/A |
| PA1161 | PA1161 | N/A | N/A | PA1161 | PA1161 | N/A | PA1161 | N/A | PA1161 | N/A | PA1161 | N/A | N/A | N/A | N/A | N/A | N/A | N/A |
| PA1162 | PA1162 | PA1162 | PA1162 | PA1162 | PA1162 | N/A | PA1162 | N/A | PA1162 | PA1162 | PA1162 | PA1162 | PA1162 | N/A | PA1162 | N/A | N/A | N/A |
| PA1163 | N/A | N/A | N/A | N/A | N/A | N/A | N/A | PA1163 | N/A | N/A | N/A | N/A | N/A | PA1163 | N/A | N/A | PA1163 | N/A |
| PA1164 | PA1164 | PA1164 | N/A | PA1164 | N/A | N/A | N/A | N/A | N/A | PA1164 | PA1164 | N/A | N/A | PA1164 | N/A | N/A | N/A | N/A |
| PA1165 | N/A | N/A | N/A | PA1165 | PA1165 | N/A | N/A | N/A | PA1165 | N/A | N/A | N/A | N/A | N/A | N/A | N/A | N/A | N/A |
| PA1166 | N/A | N/A | N/A | N/A | PA1166 | PA1166 | N/A | N/A | PA1166 | PA1166 | PA1166 | PA1166 | PA1166 | N/A | N/A | PA1166 | N/A | N/A |
| PA1170 | N/A | N/A | N/A | N/A | PA1170 | N/A | N/A | N/A | PA1170 | PA1170 | N/A | N/A | N/A | PA1170 | N/A | N/A | N/A | N/A |
| PA1171 | PA1171 | PA1171 | PA1171 | PA1171 | N/A | N/A | PA1171 | PA1171 | PA1171 | PA1171 | PA1171 | PA1171 | PA1171 | PA1171 | PA1171 | PA1171 | PA1171 | PA1171 |
| PA1172 | N/A | N/A | N/A | N/A | N/A | N/A | N/A | N/A | N/A | N/A | N/A | N/A | N/A | PA1172 | N/A | N/A | N/A | N/A |
| PA1173 | N/A | N/A | N/A | N/A | PA1173 | N/A | N/A | N/A | N/A | PA1173 | PA1173 | PA1173 | PA1173 | PA1173 | N/A | N/A | N/A | N/A |
| PA1174 | N/A | N/A | N/A | N/A | PA1174 | PA1174 | N/A | N/A | PA1174 | PA1174 | PA1174 | PA1174 | PA1174 | PA1174 | N/A | N/A | N/A | N/A |
| PA1175 | N/A | N/A | N/A | N/A | N/A | N/A | N/A | N/A | N/A | N/A | N/A | PA1175 | N/A | N/A | N/A | N/A | N/A | N/A |
| N/A | N/A | N/A | N/A | N/A | PA1177 | N/A | N/A | N/A | N/A | N/A | N/A | N/A | N/A | N/A | N/A | N/A | N/A | N/A |
| PA1178 | PA1178 | PA1178 | PA1178 | PA1178 | PA1178 | PA1178 | PA1178 | PA1178 | PA1178 | PA1178 | PA1178 | PA1178 | PA1178 | PA1178 | PA1178 | PA1178 | PA1178 | PA1178 |
| PA1179 | PA1179 | PA1179 | PA1179 | PA1179 | PA1179 | PA1179 | PA1179 | N/A | PA1179 | PA1179 | PA1179 | PA1179 | PA1179 | N/A | PA1179 | N/A | PA1179 | N/A |
| PA1180 | PA1180 | PA1180 | PA1180 | PA1180 | N/A | PA1180 | N/A | PA1180 | N/A | N/A | N/A | N/A | PA1180 | PA1180 | N/A | N/A | PA1180 | N/A |
| PA1181 | N/A | N/A | N/A | PA1181 | N/A | N/A | N/A | PA1181 | N/A | N/A | N/A | PA1181 | PA1181 | N/A | N/A | N/A | N/A | N/A |
| PA1183 | PA1183 | PA1183 | PA1183 | PA1183 | N/A | PA1183 | N/A | PA1183 | PA1183 | PA1183 | N/A | PA1183 | N/A | PA1183 | N/A | N/A | PA1183 | N/A |
| PA1184 | N/A | N/A | N/A | N/A | N/A | N/A | N/A | N/A | N/A | N/A | N/A | N/A | N/A | N/A | N/A | N/A | N/A | N/A |
| PA1189 | N/A | N/A | N/A | N/A | N/A | N/A | N/A | N/A | N/A | N/A | N/A | N/A | N/A | N/A | N/A | N/A | N/A | N/A |
| PA1191 | N/A | N/A | N/A | N/A | N/A | N/A | N/A | N/A | N/A | N/A | N/A | N/A | N/A | N/A | N/A | N/A | N/A | N/A |
| PA1192 | PA1192 | PA1192 | PA1192 | PA1192 | N/A | N/A | PA1192 | PA1192 | PA1192 | PA1192 | PA1192 | PA1192 | PA1192 | PA1192 | N/A | N/A | N/A | N/A |
| PA1193 | N/A | PA1193 | N/A | PA1193 | N/A | N/A | N/A | N/A | N/A | PA1193 | PA1193 | PA1193 | PA1193 | N/A | N/A | N/A | N/A | N/A |
| N/A | N/A | N/A | N/A | N/A | N/A | N/A | N/A | N/A | N/A | PA1195 | N/A | N/A | N/A | N/A | N/A | N/A | N/A | N/A |
| N/A | N/A | N/A | N/A | PA1196 | N/A | N/A | N/A | N/A | N/A | N/A | N/A | N/A | N/A | N/A | N/A | N/A | N/A | N/A |
| PA1197 | N/A | N/A | N/A | PA1197 | N/A | N/A | N/A | N/A | N/A | N/A | N/A | N/A | N/A | N/A | N/A | N/A | N/A | N/A |
| PA1198 | N/A | N/A | N/A | N/A | N/A | N/A | N/A | N/A | PA1198 | PA1198 | N/A | PA1198 | N/A | N/A | N/A | N/A | N/A | N/A |
| N/A | N/A | N/A | N/A | N/A | PA1199 | N/A | N/A | N/A | N/A | N/A | N/A | N/A | N/A | N/A | N/A | N/A | N/A | N/A |
| PA1200 | N/A | N/A | N/A | PA1200 | N/A | PA1200 | N/A | PA1200 | N/A | PA1200 | PA1200 | PA1200 | PA1200 | N/A | N/A | N/A | N/A | N/A |
| PA1201 | N/A | N/A | N/A | PA1201 | PA1201 | N/A | N/A | N/A | N/A | N/A | N/A | N/A | N/A | N/A | N/A | N/A | N/A | N/A |
| PA1202 | N/A | N/A | N/A | PA1202 | PA1202 | PA1202 | PA1202 | N/A | PA1202 | PA1202 | PA1202 | PA1202 | PA1202 | PA1202 | N/A | PA1202 | N/A | N/A |
| PA1203 | PA1203 | N/A | N/A | PA1203 | PA1203 | PA1203 | N/A | N/A | PA1203 | PA1203 | PA1203 | PA1203 | PA1203 | N/A | N/A | N/A | N/A | N/A |
| PA1204 | PA1204 | N/A | N/A | N/A | PA1204 | PA1204 | N/A | N/A | PA1204 | PA1204 | PA1204 | PA1204 | PA1204 | N/A | N/A | N/A | N/A | N/A |
| PA1205 | N/A | N/A | N/A | N/A | PA1205 | N/A | N/A | N/A | N/A | PA1205 | N/A | N/A | N/A | N/A | N/A | N/A | N/A | N/A |
| PA1206 | PA1206 | PA1206 | PA1206 | PA1206 | PA1206 | N/A | PA1206 | N/A | PA1206 | PA1206 | PA1206 | PA1206 | PA1206 | N/A | N/A | N/A | N/A | N/A |
| PA1207 | PA1207 | PA1207 | N/A | PA1207 | N/A | N/A | N/A | PA1207 | PA1207 | PA1207 | PA1207 | PA1207 | N/A | PA1207 | PA1207 | N/A | PA1207 | N/A |
| PA1209 | N/A | N/A | N/A | PA1209 | N/A | N/A | N/A | N/A | N/A | N/A | N/A | N/A | N/A | N/A | N/A | N/A | N/A | N/A |
| N/A | N/A | N/A | N/A | N/A | N/A | N/A | N/A | N/A | N/A | PA1210 | N/A | N/A | N/A | N/A | N/A | N/A | N/A | N/A |
| PA1212 | N/A | N/A | N/A | N/A | N/A | N/A | N/A | N/A | N/A | N/A | N/A | N/A | N/A | N/A | N/A | N/A | N/A | N/A |
| PA1213 | N/A | N/A | N/A | N/A | N/A | N/A | N/A | N/A | N/A | N/A | N/A | N/A | N/A | N/A | N/A | N/A | N/A | N/A |
| PA1214 | N/A | N/A | N/A | N/A | N/A | N/A | N/A | N/A | N/A | N/A | N/A | N/A | N/A | N/A | N/A | N/A | N/A | N/A |
| PA1215 | N/A | N/A | N/A | N/A | PA1215 | N/A | N/A | N/A | N/A | N/A | N/A | N/A | PA1215 | N/A | N/A | N/A | N/A | N/A |
| PA1216 | N/A | N/A | N/A | N/A | PA1216 | N/A | N/A | N/A | PA1216 | PA1216 | PA1216 | PA1216 | PA1216 | N/A | N/A | N/A | N/A | N/A |
| PA1217 | N/A | N/A | N/A | N/A | N/A | N/A | N/A | N/A | N/A | N/A | N/A | N/A | PA1217 | N/A | N/A | N/A | N/A | N/A |
| PA1218 | N/A | N/A | N/A | N/A | N/A | N/A | N/A | N/A | N/A | N/A | PA1218 | N/A | PA1218 | N/A | N/A | N/A | N/A | N/A |
| PA1219 | N/A | N/A | N/A | N/A | N/A | N/A | N/A | N/A | N/A | N/A | N/A | N/A | N/A | N/A | N/A | N/A | N/A | N/A |
| PA1220 | N/A | N/A | N/A | N/A | N/A | N/A | N/A | N/A | N/A | N/A | N/A | N/A | N/A | N/A | N/A | N/A | N/A | N/A |
| PA1222 | PA1222 | PA1222 | N/A | PA1222 | N/A | PA1222 | PA1222 | PA1222 | PA1222 | PA1222 | PA1222 | PA1222 | PA1222 | PA1222 | PA1222 | N/A | PA1222 | N/A |
| PA1223 | N/A | N/A | N/A | PA1223 | N/A | N/A | N/A | PA1223 | N/A | N/A | PA1223 | N/A | N/A | N/A | N/A | N/A | N/A | N/A |
| N/A | N/A | N/A | N/A | N/A | PA1224 | N/A | N/A | N/A | N/A | N/A | N/A | N/A | N/A | N/A | N/A | N/A | N/A | N/A |
| PA1225 | PA1225 | PA1225 | PA1225 | PA1225 | PA1225 | N/A | N/A | N/A | PA1225 | N/A | N/A | PA1225 | N/A | N/A | N/A | N/A | N/A | N/A |
| PA1226 | N/A | N/A | N/A | PA1226 | N/A | N/A | N/A | N/A | PA1226 | N/A | PA1226 | N/A | N/A | N/A | N/A | N/A | N/A | N/A |
| N/A | N/A | N/A | N/A | PA1227 | N/A | N/A | N/A | N/A | N/A | N/A | N/A | N/A | N/A | N/A | N/A | N/A | N/A | N/A |
| PA1228 | PA1228 | PA1228 | PA1228 | PA1228 | N/A | N/A | PA1228 | PA1228 | PA1228 | N/A | PA1228 | PA1228 | N/A | PA1228 | N/A | N/A | PA1228 | PA1228 |
| N/A | N/A | N/A | N/A | N/A | PA1233 | N/A | N/A | N/A | N/A | N/A | N/A | N/A | N/A | N/A | N/A | N/A | N/A | N/A |
| PA1234 | N/A | N/A | N/A | PA1234 | PA1234 | N/A | N/A | N/A | N/A | N/A | N/A | PA1234 | PA1234 | N/A | N/A | N/A | N/A | N/A |
| PA1240 | N/A | N/A | N/A | N/A | N/A | N/A | N/A | N/A | N/A | N/A | N/A | N/A | N/A | N/A | N/A | N/A | N/A | N/A |
| PA1241 | N/A | N/A | N/A | N/A | N/A | N/A | N/A | N/A | PA1241 | N/A | N/A | N/A | N/A | N/A | N/A | N/A | N/A | N/A |
| PA1243 | N/A | N/A | N/A | PA1243 | PA1243 | N/A | N/A | N/A | N/A | N/A | PA1243 | PA1243 | PA1243 | N/A | N/A | N/A | N/A | N/A |
| PA1244 | PA1244 | PA1244 | PA1244 | PA1244 | PA1244 | PA1244 | N/A | N/A | PA1244 | PA1244 | PA1244 | PA1244 | PA1244 | PA1244 | N/A | N/A | N/A | PA1244 |
| PA1245 | N/A | N/A | N/A | N/A | N/A | N/A | N/A | N/A | N/A | N/A | N/A | N/A | N/A | PA1245 | N/A | N/A | N/A | N/A |
| PA1246 | N/A | N/A | N/A | N/A | N/A | N/A | N/A | N/A | PA1246 | N/A | PA1246 | PA1246 | PA1246 | PA1246 | N/A | N/A | N/A | N/A |
| PA1247 | N/A | N/A | N/A | N/A | N/A | N/A | N/A | N/A | N/A | N/A | N/A | N/A | N/A | PA1247 | N/A | N/A | N/A | N/A |
| PA1248 | N/A | N/A | N/A | N/A | N/A | N/A | N/A | N/A | N/A | N/A | N/A | N/A | PA1248 | N/A | N/A | N/A | N/A | N/A |
| N/A | N/A | N/A | N/A | N/A | PA1249 | N/A | N/A | N/A | N/A | N/A | N/A | N/A | N/A | N/A | N/A | PA1249 | N/A | N/A |
| PA1250 | N/A | N/A | N/A | PA1250 | PA1250 | N/A | N/A | N/A | N/A | N/A | PA1250 | PA1250 | N/A | N/A | PA1250 | N/A | N/A | N/A |
| PA1251 | PA1251 | PA1251 | N/A | PA1251 | N/A | N/A | N/A | N/A | N/A | N/A | N/A | N/A | N/A | PA1251 | N/A | N/A | N/A | N/A |
| PA1258 | PA1258 | N/A | N/A | PA1258 | N/A | N/A | N/A | N/A | PA1258 | N/A | N/A | N/A | N/A | PA1258 | PA1258 | N/A | PA1258 | PA1258 |
| PA1263 | N/A | N/A | N/A | PA1263 | N/A | N/A | N/A | N/A | N/A | N/A | N/A | N/A | N/A | N/A | N/A | N/A | N/A | N/A |
| PA1264 | N/A | N/A | N/A | N/A | N/A | N/A | N/A | N/A | N/A | N/A | N/A | N/A | N/A | N/A | N/A | N/A | N/A | N/A |
| PA1269 | N/A | N/A | N/A | PA1269 | N/A | N/A | N/A | N/A | N/A | N/A | N/A | N/A | N/A | N/A | N/A | N/A | N/A | N/A |
| N/A | N/A | N/A | N/A | N/A | PA1270 | N/A | N/A | N/A | N/A | N/A | N/A | N/A | N/A | N/A | N/A | N/A | N/A | N/A |
| PA1271 | PA1271 | N/A | N/A | PA1271 | PA1271 | PA1271 | N/A | PA1271 | PA1271 | PA1271 | N/A | N/A | PA1271 | N/A | PA1271 | N/A | N/A | N/A |
| PA1272 | N/A | PA1272 | N/A | PA1272 | PA1272 | N/A | PA1272 | N/A | PA1272 | N/A | PA1272 | PA1272 | PA1272 | PA1272 | N/A | N/A | N/A | N/A |
| N/A | N/A | N/A | N/A | PA1273 | N/A | N/A | N/A | N/A | N/A | N/A | N/A | N/A | N/A | N/A | N/A | N/A | N/A | N/A |
| PA1274 | N/A | N/A | N/A | PA1274 | N/A | N/A | N/A | N/A | PA1274 | N/A | N/A | N/A | N/A | N/A | N/A | N/A | N/A | N/A |
| PA1275 | N/A | N/A | N/A | PA1275 | PA1275 | N/A | N/A | N/A | N/A | N/A | N/A | N/A | N/A | PA1275 | N/A | N/A | N/A | N/A |
| PA1276 | N/A | N/A | PA1276 | PA1276 | PA1276 | N/A | N/A | N/A | PA1276 | N/A | N/A | N/A | N/A | N/A | N/A | N/A | N/A | N/A |
| PA1277 | N/A | N/A | N/A | PA1277 | PA1277 | N/A | N/A | PA1277 | N/A | N/A | N/A | N/A | N/A | N/A | N/A | N/A | N/A | N/A |
| PA1278 | N/A | PA1278 | N/A | PA1278 | N/A | N/A | N/A | N/A | PA1278 | PA1278 | PA1278 | PA1278 | PA1278 | N/A | N/A | N/A | N/A | N/A |

| | | | | | | | | | | | | | | | | | | |
|---|---|---|---|---|---|---|---|---|---|---|---|---|---|---|---|---|---|---|
| PA1279 | N/A | PA1279 | N/A | PA1279 | N/A | N/A | N/A | PA1279 | PA1279 | N/A | PA1279 | PA1279 | N/A | N/A | PA1279 | N/A | N/A | N/A |
| PA1280 | N/A | N/A | N/A | PA1280 | N/A | N/A | N/A | N/A | N/A | N/A | N/A | N/A | N/A | N/A | N/A | N/A | N/A | N/A |
| PA1281 | N/A | N/A | N/A | N/A | N/A | N/A | N/A | N/A | PA1281 | N/A | N/A | N/A | N/A | N/A | N/A | N/A | N/A | N/A |
| PA1283 | N/A | N/A | N/A | PA1283 | N/A | PA1283 | N/A | N/A | PA1283 | PA1283 | PA1283 | PA1283 | PA1283 | PA1283 | N/A | N/A | N/A | N/A |
| PA1284 | PA1284 | PA1284 | PA1284 | PA1284 | PA1284 | PA1284 | N/A | N/A | PA1284 | PA1284 | PA1284 | PA1284 | PA1284 | PA1284 | N/A | N/A | N/A | N/A |
| PA1285 | N/A | N/A | N/A | PA1285 | N/A | PA1285 | N/A | N/A | PA1285 | PA1285 | PA1285 | PA1285 | PA1285 | N/A | N/A | N/A | N/A | N/A |
| N/A | N/A | N/A | N/A | N/A | PA1286 | N/A | N/A | N/A | N/A | N/A | N/A | N/A | N/A | N/A | N/A | N/A | N/A | N/A |
| PA1287 | N/A | N/A | N/A | PA1287 | PA1287 | N/A | N/A | N/A | PA1287 | PA1287 | PA1287 | PA1287 | PA1287 | N/A | PA1287 | N/A | N/A | N/A |
| PA1288 | PA1288 | PA1288 | PA1288 | PA1288 | PA1288 | PA1288 | PA1288 | PA1288 | PA1288 | PA1288 | PA1288 | PA1288 | PA1288 | PA1288 | PA1288 | PA1288 | PA1288 | N/A |
| N/A | N/A | N/A | N/A | N/A | PA1291 | N/A | N/A | N/A | N/A | N/A | N/A | N/A | N/A | N/A | N/A | N/A | N/A | N/A |
| PA1292 | N/A | PA1292 | N/A | PA1292 | PA1292 | PA1292 | PA1292 | N/A | PA1292 | PA1292 | PA1292 | PA1292 | PA1292 | N/A | N/A | N/A | N/A | N/A |
| PA1293 | PA1293 | PA1293 | PA1293 | PA1293 | PA1293 | N/A | PA1293 | N/A | PA1293 | PA1293 | N/A | PA1293 | PA1293 | PA1293 | PA1293 | N/A | PA1293 | N/A |
| PA1294 | PA1294 | N/A | PA1294 | PA1294 | PA1294 | N/A | PA1294 | PA1294 | PA1294 | PA1294 | PA1294 | PA1294 | PA1294 | N/A | PA1294 | N/A | N/A | N/A |
| PA1295 | PA1295 | PA1295 | N/A | PA1295 | PA1295 | N/A | N/A | N/A | PA1295 | N/A | N/A | N/A | N/A | N/A | PA1295 | N/A | N/A | N/A |
| PA1296 | N/A | N/A | N/A | PA1296 | PA1296 | N/A | N/A | N/A | N/A | N/A | N/A | N/A | N/A | N/A | N/A | N/A | N/A | N/A |
| N/A | N/A | N/A | N/A | PA1297 | N/A | N/A | N/A | N/A | N/A | N/A | N/A | N/A | N/A | N/A | N/A | N/A | N/A | N/A |
| PA1298 | PA1298 | N/A | N/A | PA1298 | N/A | N/A | N/A | N/A | PA1298 | N/A | PA1298 | PA1298 | PA1298 | PA1298 | N/A | N/A | N/A | N/A |
| N/A | N/A | N/A | N/A | N/A | N/A | N/A | N/A | N/A | N/A | N/A | PA1299 | N/A | N/A | N/A | N/A | N/A | N/A | N/A |
| PA1301 | N/A | N/A | N/A | PA1301 | N/A | N/A | N/A | N/A | N/A | N/A | N/A | N/A | PA1301 | N/A | N/A | N/A | N/A | N/A |
| PA1303 | N/A | N/A | N/A | PA1303 | PA1303 | N/A | N/A | N/A | N/A | N/A | N/A | N/A | N/A | PA1303 | N/A | N/A | N/A | N/A |
| PA1304 | N/A | PA1304 | PA1304 | PA1304 | PA1304 | PA1304 | N/A | N/A | PA1304 | PA1304 | PA1304 | PA1304 | PA1304 | N/A | N/A | N/A | N/A | N/A |
| PA1305 | N/A | PA1305 | N/A | N/A | N/A | PA1305 | N/A | N/A | N/A | N/A | N/A | PA1305 | N/A | PA1305 | N/A | N/A | PA1305 | N/A |
| PA1306 | N/A | N/A | N/A | N/A | PA1306 | N/A | N/A | N/A | N/A | N/A | N/A | N/A | N/A | PA1306 | N/A | N/A | N/A | N/A |
| PA1307 | PA1307 | N/A | PA1307 | PA1307 | PA1307 | PA1307 | PA1307 | N/A | PA1307 | PA1307 | PA1307 | PA1307 | PA1307 | N/A | N/A | N/A | N/A | N/A |
| PA1308 | N/A | N/A | N/A | N/A | N/A | PA1308 | PA1308 | N/A | N/A | PA1308 | N/A | PA1308 | PA1308 | PA1308 | N/A | N/A | N/A | N/A |
| PA1309 | N/A | N/A | N/A | N/A | N/A | N/A | N/A | N/A | N/A | N/A | N/A | N/A | N/A | N/A | N/A | N/A | N/A | N/A |
| PA1310 | N/A | N/A | N/A | N/A | N/A | N/A | N/A | N/A | N/A | N/A | N/A | N/A | N/A | N/A | N/A | N/A | N/A | N/A |
| PA1311 | N/A | N/A | N/A | PA1311 | N/A | N/A | N/A | N/A | N/A | N/A | N/A | N/A | N/A | N/A | N/A | N/A | N/A | N/A |
| PA1315 | N/A | N/A | N/A | PA1315 | N/A | N/A | PA1315 | N/A | PA1315 | N/A | PA1315 | PA1315 | N/A | N/A | N/A | N/A | N/A | N/A |
| PA1316 | N/A | N/A | N/A | N/A | N/A | N/A | N/A | N/A | N/A | N/A | N/A | N/A | N/A | PA1316 | N/A | N/A | N/A | N/A |
| PA1317 | PA1317 | PA1317 | N/A | PA1317 | N/A | N/A | PA1317 | PA1317 | PA1317 | N/A | PA1317 | N/A | N/A | PA1317 | N/A | N/A | PA1317 | N/A |
| PA1318 | N/A | PA1318 | N/A | N/A | N/A | N/A | N/A | PA1318 | PA1318 | N/A | PA1318 | PA1318 | N/A | PA1318 | N/A | N/A | PA1318 | N/A |
| PA1323 | N/A | N/A | N/A | N/A | PA1323 | PA1323 | N/A | N/A | N/A | N/A | N/A | PA1323 | PA1323 | PA1323 | N/A | N/A | PA1323 | N/A |
| PA1324 | PA1324 | PA1324 | N/A | PA1324 | PA1324 | PA1324 | PA1324 | N/A | PA1324 | PA1324 | PA1324 | PA1324 | PA1324 | PA1324 | PA1324 | N/A | N/A | N/A |
| N/A | N/A | N/A | N/A | N/A | N/A | N/A | N/A | N/A | N/A | PA1325 | N/A | N/A | N/A | N/A | N/A | N/A | N/A | N/A |
| PA1326 | PA1326 | PA1326 | PA1326 | PA1326 | N/A | N/A | N/A | N/A | PA1326 | PA1326 | PA1326 | PA1326 | N/A | N/A | N/A | PA1326 | N/A | N/A |
| PA1327 | N/A | N/A | N/A | N/A | N/A | PA1327 | N/A | N/A | N/A | N/A | N/A | PA1327 | PA1327 | N/A | N/A | N/A | N/A | N/A |
| N/A | N/A | N/A | N/A | PA1328 | N/A | N/A | N/A | N/A | N/A | N/A | N/A | N/A | N/A | N/A | N/A | N/A | N/A | N/A |
| N/A | N/A | N/A | N/A | N/A | PA1329 | N/A | N/A | N/A | N/A | N/A | N/A | N/A | N/A | N/A | N/A | N/A | N/A | N/A |
| PA1330 | N/A | N/A | N/A | PA1330 | PA1330 | N/A | N/A | N/A | N/A | PA1330 | N/A | N/A | N/A | N/A | PA1330 | N/A | N/A | N/A |
| PA1331 | N/A | N/A | N/A | PA1331 | N/A | N/A | N/A | N/A | N/A | N/A | N/A | PA1331 | N/A | PA1331 | N/A | N/A | N/A | N/A |
| PA1333 | N/A | N/A | N/A | N/A | N/A | N/A | N/A | N/A | N/A | N/A | PA1333 | N/A | N/A | N/A | N/A | N/A | N/A | N/A |
| PA1334 | N/A | N/A | N/A | N/A | N/A | N/A | N/A | N/A | N/A | N/A | N/A | N/A | N/A | N/A | N/A | N/A | N/A | N/A |
| PA1335 | N/A | N/A | N/A | PA1335 | N/A | N/A | N/A | N/A | PA1335 | PA1335 | PA1335 | PA1335 | N/A | N/A | N/A | N/A | N/A | N/A |
| PA1336 | N/A | N/A | N/A | N/A | PA1336 | N/A | N/A | N/A | N/A | N/A | N/A | N/A | N/A | PA1336 | N/A | N/A | N/A | N/A |
| PA1337 | PA1337 | PA1337 | PA1337 | PA1337 | PA1337 | PA1337 | PA1337 | N/A | PA1337 | PA1337 | PA1337 | PA1337 | PA1337 | PA1337 | PA1337 | PA1337 | N/A | PA1337 |
| PA1338 | PA1338 | PA1338 | PA1338 | PA1338 | PA1338 | PA1338 | PA1338 | N/A | PA1338 | PA1338 | PA1338 | PA1338 | PA1338 | N/A | PA1338 | PA1338 | N/A | N/A |
| PA1339 | PA1339 | PA1339 | PA1339 | PA1339 | N/A | PA1339 | PA1339 | N/A | PA1339 | PA1339 | PA1339 | PA1339 | PA1339 | PA1339 | N/A | N/A | PA1339 | PA1339 |
| PA1340 | N/A | PA1340 | N/A | N/A | N/A | PA1340 | N/A | N/A | PA1340 | PA1340 | PA1340 | PA1340 | PA1340 | PA1340 | N/A | N/A | PA1340 | N/A |
| PA1341 | PA1341 | PA1341 | PA1341 | PA1341 | PA1341 | PA1341 | N/A | PA1341 | PA1341 | PA1341 | PA1341 | PA1341 | PA1341 | PA1341 | N/A | N/A | PA1341 | PA1341 |
| PA1342 | PA1342 | PA1342 | PA1342 | PA1342 | PA1342 | PA1342 | PA1342 | N/A | PA1342 | PA1342 | PA1342 | PA1342 | PA1342 | PA1342 | PA1342 | PA1342 | PA1342 | PA1342 |
| PA1343 | N/A | N/A | N/A | PA1343 | PA1343 | PA1343 | N/A | N/A | PA1343 | N/A | PA1343 | N/A | PA1343 | N/A | N/A | N/A | N/A | N/A |
| PA1344 | PA1344 | PA1344 | PA1344 | PA1344 | PA1344 | PA1344 | N/A | N/A | PA1344 | PA1344 | PA1344 | PA1344 | PA1344 | N/A | N/A | N/A | N/A | N/A |
| PA1345 | N/A | N/A | N/A | N/A | N/A | N/A | N/A | N/A | N/A | N/A | N/A | N/A | PA1345 | N/A | N/A | N/A | N/A | N/A |
| PA1349 | N/A | N/A | N/A | N/A | N/A | N/A | N/A | N/A | N/A | N/A | PA1349 | N/A | PA1349 | N/A | N/A | N/A | N/A | N/A |
| PA1353 | N/A | N/A | N/A | N/A | N/A | PA1353 | N/A | N/A | N/A | PA1353 | PA1353 | PA1353 | PA1353 | N/A | N/A | N/A | N/A | N/A |
| PA1354 | N/A | N/A | N/A | N/A | N/A | PA1354 | N/A | N/A | N/A | N/A | N/A | PA1354 | PA1354 | N/A | N/A | N/A | N/A | N/A |
| PA1357 | PA1357 | PA1357 | PA1357 | PA1357 | N/A | N/A | N/A | N/A | N/A | N/A | PA1357 | PA1357 | PA1357 | N/A | N/A | N/A | N/A | N/A |
| PA1358 | N/A | N/A | N/A | N/A | N/A | PA1358 | N/A | N/A | N/A | N/A | PA1358 | PA1358 | PA1358 | N/A | N/A | N/A | N/A | N/A |
| PA1360 | N/A | N/A | N/A | N/A | N/A | N/A | N/A | N/A | N/A | N/A | N/A | N/A | N/A | N/A | N/A | N/A | N/A | N/A |
| PA1361 | N/A | PA1361 | N/A | PA1361 | N/A | N/A | N/A | N/A | N/A | N/A | N/A | PA1361 | N/A | PA1361 | N/A | N/A | N/A | N/A |
| PA1364 | N/A | N/A | N/A | PA1364 | N/A | N/A | PA1364 | N/A | PA1364 | N/A | N/A | N/A | N/A | N/A | N/A | N/A | PA1364 | N/A |
| PA1365 | N/A | N/A | N/A | PA1365 | N/A | N/A | N/A | PA1365 | N/A | N/A | N/A | N/A | N/A | N/A | PA1365 | N/A | N/A | N/A |
| PA1367 | N/A | N/A | N/A | PA1367 | N/A | N/A | N/A | N/A | PA1367 | N/A | PA1367 | N/A | N/A | N/A | N/A | N/A | N/A | N/A |
| N/A | N/A | N/A | N/A | N/A | N/A | N/A | N/A | N/A | PA1369 | N/A | PA1369 | N/A | N/A | N/A | N/A | N/A | N/A | N/A |
| N/A | N/A | N/A | N/A | N/A | N/A | N/A | N/A | PA1370 | N/A | N/A | N/A | N/A | N/A | N/A | N/A | N/A | N/A | N/A |
| N/A | N/A | N/A | N/A | N/A | PA1371 | N/A | N/A | N/A | N/A | N/A | N/A | N/A | N/A | N/A | N/A | N/A | N/A | N/A |
| PA1372 | PA1372 | PA1372 | PA1372 | PA1372 | PA1372 | PA1372 | N/A | PA1372 | PA1372 | PA1372 | PA1372 | PA1372 | PA1372 | PA1372 | N/A | N/A | N/A | N/A |
| PA1373 | PA1373 | PA1373 | PA1373 | PA1373 | PA1373 | N/A | N/A | N/A | N/A | N/A | N/A | N/A | PA1373 | N/A | N/A | N/A | N/A | N/A |
| PA1375 | PA1375 | PA1375 | PA1375 | PA1375 | PA1375 | N/A | N/A | N/A | N/A | PA1375 | N/A | N/A | PA1375 | N/A | N/A | N/A | N/A | N/A |
| PA1376 | N/A | N/A | N/A | PA1376 | N/A | PA1376 | N/A | PA1376 | N/A | PA1376 | N/A | PA1376 | PA1376 | N/A | N/A | N/A | N/A | N/A |
| PA1377 | N/A | N/A | N/A | PA1377 | N/A | PA1377 | N/A | N/A | PA1377 | N/A | PA1377 | PA1377 | PA1377 | N/A | N/A | N/A | N/A | N/A |
| PA1379 | N/A | N/A | N/A | N/A | N/A | N/A | N/A | N/A | N/A | N/A | N/A | N/A | N/A | N/A | N/A | N/A | N/A | N/A |
| PA1380 | N/A | N/A | N/A | N/A | N/A | N/A | N/A | N/A | N/A | PA1380 | N/A | PA1380 | PA1380 | N/A | N/A | N/A | N/A | N/A |
| PA1383 | N/A | N/A | N/A | PA1383 | N/A | N/A | N/A | N/A | N/A | N/A | N/A | N/A | PA1383 | N/A | N/A | N/A | N/A | N/A |
| PA1395 | N/A | N/A | N/A | PA1395 | N/A | N/A | N/A | N/A | N/A | N/A | N/A | N/A | N/A | N/A | N/A | N/A | N/A | N/A |
| PA1396 | N/A | N/A | N/A | N/A | N/A | N/A | N/A | N/A | N/A | N/A | N/A | N/A | N/A | PA1396 | N/A | N/A | PA1396 | N/A |
| PA1397 | N/A | N/A | N/A | PA1397 | N/A | N/A | N/A | N/A | PA1397 | N/A | N/A | N/A | N/A | N/A | N/A | N/A | N/A | N/A |
| PA1398 | N/A | N/A | N/A | N/A | N/A | PA1398 | N/A | N/A | PA1398 | PA1398 | PA1398 | N/A | N/A | PA1398 | N/A | N/A | PA1398 | N/A |
| PA1401 | N/A | N/A | N/A | N/A | N/A | N/A | N/A | N/A | N/A | N/A | PA1401 | PA1401 | PA1401 | N/A | N/A | N/A | N/A | N/A |
| PA1402 | N/A | N/A | N/A | PA1402 | N/A | N/A | N/A | N/A | PA1402 | PA1402 | N/A | PA1402 | N/A | N/A | N/A | N/A | N/A | N/A |
| PA1403 | N/A | N/A | N/A | N/A | N/A | N/A | N/A | N/A | N/A | N/A | N/A | PA1403 | PA1403 | N/A | N/A | N/A | N/A | N/A |
| PA1404 | N/A | N/A | N/A | N/A | N/A | PA1404 | N/A | N/A | N/A | N/A | PA1404 | N/A | PA1404 | N/A | N/A | N/A | N/A | N/A |
| PA1407 | N/A | N/A | PA1407 | PA1407 | N/A | N/A | N/A | N/A | N/A | N/A | N/A | N/A | N/A | N/A | N/A | N/A | N/A | N/A |
| PA1408 | N/A | N/A | N/A | N/A | N/A | N/A | N/A | N/A | N/A | N/A | N/A | PA1408 | PA1408 | PA1408 | N/A | N/A | N/A | N/A |
| PA1410 | N/A | PA1410 | N/A | N/A | N/A | N/A | N/A | N/A | N/A | N/A | N/A | N/A | N/A | N/A | N/A | N/A | N/A | N/A |
| PA1413 | N/A | N/A | N/A | N/A | N/A | N/A | N/A | N/A | N/A | N/A | PA1413 | N/A | N/A | N/A | N/A | N/A | N/A | N/A |
| PA1417 | N/A | PA1417 | N/A | PA1417 | N/A | N/A | N/A | PA1417 | PA1417 | N/A | N/A | PA1417 | N/A | N/A | N/A | N/A | N/A | N/A |
| N/A | N/A | N/A | N/A | N/A | N/A | N/A | N/A | N/A | N/A | N/A | N/A | N/A | N/A | PA1418 | N/A | N/A | N/A | N/A |
| N/A | N/A | N/A | N/A | N/A | PA1420 | N/A | N/A | N/A | N/A | N/A | N/A | N/A | N/A | N/A | N/A | N/A | N/A | N/A |
| PA1421 | PA1421 | PA1421 | PA1421 | PA1421 | PA1421 | PA1421 | PA1421 | N/A | PA1421 | PA1421 | PA1421 | PA1421 | PA1421 | PA1421 | PA1421 | N/A | N/A | N/A |
| PA1422 | N/A | N/A | N/A | PA1422 | N/A | N/A | N/A | N/A | PA1422 | N/A | PA1422 | N/A | PA1422 | PA1422 | N/A | N/A | N/A | N/A |
| PA1423 | PA1423 | PA1423 | PA1423 | PA1423 | N/A | PA1423 | PA1423 | PA1423 | PA1423 | PA1423 | PA1423 | PA1423 | PA1423 | PA1423 | PA1423 | N/A | PA1423 | PA1423 |
| PA1425 | N/A | N/A | N/A | N/A | N/A | N/A | N/A | N/A | N/A | N/A | N/A | N/A | N/A | N/A | N/A | N/A | N/A | N/A |
| PA1428 | N/A | N/A | N/A | PA1428 | N/A | N/A | N/A | N/A | N/A | N/A | N/A | N/A | N/A | N/A | N/A | N/A | N/A | N/A |
| PA1429 | N/A | N/A | N/A | PA1429 | PA1429 | N/A | N/A | N/A | N/A | PA1429 | PA1429 | PA1429 | N/A | PA1429 | N/A | N/A | PA1429 | N/A |
| PA1430 | PA1430 | PA1430 | N/A | PA1430 | N/A | PA1430 | N/A | N/A | PA1430 | PA1430 | PA1430 | PA1430 | PA1430 | PA1430 | N/A | N/A | PA1430 | N/A |
| PA1431 | N/A | N/A | N/A | PA1431 | N/A | N/A | N/A | N/A | PA1431 | N/A | PA1431 | PA1431 | PA1431 | N/A | N/A | N/A | PA1431 | N/A |
| PA1432 | PA1432 | PA1432 | PA1432 | PA1432 | N/A | PA1432 | N/A | PA1432 | PA1432 | PA1432 | N/A | N/A | PA1432 | PA1432 | N/A | N/A | PA1432 | N/A |
| PA1433 | N/A | N/A | N/A | PA1433 | N/A | N/A | N/A | N/A | N/A | PA1433 | N/A | N/A | PA1433 | PA1433 | N/A | N/A | PA1433 | N/A |
| PA1434 | N/A | N/A | N/A | N/A | N/A | N/A | N/A | N/A | N/A | N/A | N/A | N/A | N/A | N/A | N/A | N/A | N/A | N/A |
| PA1437 | N/A | N/A | N/A | N/A | N/A | N/A | N/A | N/A | N/A | N/A | N/A | N/A | N/A | N/A | N/A | N/A | N/A | N/A |
| PA1438 | N/A | N/A | N/A | N/A | N/A | N/A | N/A | N/A | N/A | N/A | N/A | N/A | N/A | PA1438 | N/A | N/A | N/A | N/A |
| N/A | N/A | N/A | N/A | N/A | PA1439 | N/A | N/A | N/A | N/A | N/A | N/A | N/A | N/A | N/A | N/A | N/A | N/A | N/A |
| PA1440 | PA1440 | PA1440 | N/A | PA1440 | PA1440 | PA1440 | PA1440 | N/A | PA1440 | PA1440 | PA1440 | PA1440 | PA1440 | N/A | PA1440 | N/A | N/A | N/A |
| N/A | PA1441 | PA1441 | N/A | PA1441 | N/A | N/A | N/A | N/A | N/A | N/A | N/A | N/A | N/A | N/A | PA1441 | N/A | N/A | N/A |
| PA1442 | PA1442 | N/A | N/A | PA1442 | PA1442 | PA1442 | PA1442 | N/A | PA1442 | PA1442 | PA1442 | PA1442 | PA1442 | PA1442 | N/A | N/A | N/A | N/A |
| PA1443 | PA1443 | PA1443 | N/A | PA1443 | PA1443 | N/A | PA1443 | PA1443 | PA1443 | PA1443 | PA1443 | PA1443 | PA1443 | N/A | N/A | N/A | N/A | N/A |
| PA1444 | PA1444 | PA1444 | PA1444 | PA1444 | PA1444 | N/A | N/A | N/A | PA1444 | PA1444 | PA1444 | PA1444 | PA1444 | N/A | PA1444 | N/A | N/A | N/A |

| | | | | | | | | | | | | | | | | | | |
|---|---|---|---|---|---|---|---|---|---|---|---|---|---|---|---|---|---|---|
| PA1445 | N/A | N/A | N/A | N/A | N/A | PA1445 | N/A | N/A | N/A | N/A | N/A | N/A | N/A | PA1445 | N/A | N/A | N/A | N/A |
| PA1446 | N/A | N/A | N/A | PA1446 | N/A | PA1446 | N/A | N/A | PA1446 | N/A | PA1446 | PA1446 | PA1446 | PA1446 | N/A | N/A | N/A | N/A |
| PA1449 | N/A | N/A | N/A | N/A | N/A | N/A | N/A | N/A | N/A | N/A | N/A | N/A | N/A | N/A | N/A | N/A | N/A | N/A |
| PA1450 | N/A | N/A | N/A | N/A | N/A | N/A | N/A | N/A | N/A | N/A | N/A | PA1450 | N/A | PA1450 | N/A | N/A | PA1450 | N/A |
| PA1451 | N/A | N/A | N/A | N/A | N/A | N/A | N/A | N/A | N/A | N/A | N/A | N/A | N/A | PA1451 | N/A | N/A | PA1451 | N/A |
| PA1452 | N/A | N/A | N/A | PA1452 | N/A | N/A | PA1452 | PA1452 | PA1452 | N/A | PA1452 | N/A | N/A | PA1452 | N/A | N/A | PA1452 | N/A |
| PA1453 | N/A | N/A | N/A | PA1453 | PA1453 | N/A | N/A | PA1453 | N/A | N/A | N/A | N/A | N/A | N/A | N/A | N/A | PA1453 | N/A |
| PA1454 | PA1454 | PA1454 | PA1454 | PA1454 | PA1454 | PA1454 | N/A | PA1454 | PA1454 | PA1454 | PA1454 | PA1454 | PA1454 | PA1454 | N/A | N/A | PA1454 | PA1454 |
| PA1455 | PA1455 | PA1455 | PA1455 | PA1455 | PA1455 | PA1455 | PA1455 | N/A | PA1455 | PA1455 | PA1455 | PA1455 | PA1455 | PA1455 | PA1455 | N/A | N/A | N/A |
| PA1456 | PA1456 | PA1456 | PA1456 | PA1456 | N/A | PA1456 | PA1456 | N/A | PA1456 | PA1456 | PA1456 | PA1456 | PA1456 | N/A | PA1456 | N/A | N/A | N/A |
| PA1457 | PA1457 | PA1457 | PA1457 | PA1457 | N/A | PA1457 | PA1457 | N/A | PA1457 | PA1457 | PA1457 | PA1457 | PA1457 | N/A | PA1457 | N/A | N/A | N/A |
| PA1458 | PA1458 | PA1458 | PA1458 | PA1458 | PA1458 | PA1458 | PA1458 | PA1458 | PA1458 | PA1458 | PA1458 | PA1458 | PA1458 | PA1458 | PA1458 | N/A | PA1458 | N/A |
| PA1459 | PA1459 | PA1459 | PA1459 | PA1459 | PA1459 | PA1459 | PA1459 | PA1459 | PA1459 | PA1459 | PA1459 | PA1459 | PA1459 | PA1459 | PA1459 | N/A | PA1459 | PA1459 |
| PA1460 | N/A | N/A | N/A | PA1460 | N/A | N/A | N/A | N/A | PA1460 | N/A | N/A | PA1460 | N/A | PA1460 | N/A | N/A | N/A | N/A |
| PA1461 | PA1461 | N/A | N/A | PA1461 | PA1461 | N/A | N/A | N/A | PA1461 | N/A | N/A | N/A | N/A | PA1461 | N/A | N/A | N/A | N/A |
| PA1462 | PA1462 | N/A | N/A | PA1462 | N/A | N/A | PA1462 | N/A | PA1462 | PA1462 | PA1462 | PA1462 | PA1462 | PA1462 | N/A | N/A | PA1462 | N/A |
| PA1463 | N/A | PA1463 | N/A | PA1463 | PA1463 | N/A | N/A | N/A | N/A | N/A | N/A | N/A | N/A | N/A | N/A | N/A | N/A | N/A |
| PA1464 | PA1464 | PA1464 | PA1464 | PA1464 | PA1464 | PA1464 | N/A | N/A | PA1464 | PA1464 | PA1464 | PA1464 | PA1464 | PA1464 | PA1464 | PA1464 | N/A | N/A |
| PA1465 | N/A | N/A | N/A | N/A | N/A | PA1465 | PA1465 | N/A | PA1465 | PA1465 | PA1465 | PA1465 | PA1465 | PA1465 | N/A | N/A | PA1465 | N/A |
| PA1466 | N/A | N/A | N/A | PA1466 | N/A | N/A | N/A | N/A | N/A | N/A | N/A | N/A | N/A | N/A | PA1466 | N/A | N/A | N/A |
| N/A | N/A | N/A | N/A | N/A | PA1469 | N/A | N/A | N/A | N/A | N/A | N/A | N/A | N/A | N/A | N/A | N/A | N/A | N/A |
| PA1470 | N/A | N/A | N/A | PA1470 | PA1470 | N/A | N/A | N/A | N/A | PA1470 | PA1470 | PA1470 | PA1470 | N/A | N/A | N/A | N/A | N/A |
| PA1472 | N/A | N/A | N/A | PA1472 | N/A | N/A | N/A | N/A | N/A | N/A | N/A | N/A | N/A | N/A | N/A | N/A | N/A | N/A |
| N/A | N/A | N/A | N/A | PA1473 | N/A | N/A | N/A | N/A | PA1473 | N/A | PA1473 | N/A | PA1473 | N/A | N/A | N/A | N/A | N/A |
| PA1474 | N/A | N/A | N/A | PA1474 | N/A | N/A | PA1474 | N/A | N/A | PA1474 | N/A | PA1474 | PA1474 | N/A | N/A | N/A | N/A | N/A |
| PA1475 | N/A | N/A | N/A | PA1475 | N/A | N/A | PA1475 | PA1475 | PA1475 | N/A | PA1475 | PA1475 | PA1475 | PA1475 | N/A | N/A | PA1475 | N/A |
| PA1476 | N/A | N/A | N/A | PA1476 | N/A | N/A | N/A | N/A | PA1476 | PA1476 | N/A | N/A | PA1476 | PA1476 | N/A | N/A | N/A | N/A |
| N/A | N/A | N/A | N/A | N/A | N/A | PA1477 | N/A | N/A | N/A | N/A | N/A | N/A | N/A | PA1477 | N/A | N/A | N/A | N/A |
| PA1478 | N/A | N/A | N/A | N/A | PA1478 | N/A | N/A | N/A | N/A | N/A | N/A | N/A | N/A | N/A | N/A | N/A | N/A | N/A |
| PA1479 | N/A | PA1479 | PA1479 | PA1479 | PA1479 | PA1479 | PA1479 | PA1479 | PA1479 | PA1479 | PA1479 | PA1479 | PA1479 | PA1479 | PA1479 | N/A | PA1479 | N/A |
| PA1480 | N/A | N/A | N/A | PA1480 | N/A | N/A | N/A | N/A | N/A | PA1480 | PA1480 | PA1480 | N/A | PA1480 | N/A | N/A | N/A | N/A |
| PA1481 | N/A | PA1481 | N/A | PA1481 | PA1481 | PA1481 | PA1481 | N/A | PA1481 | PA1481 | PA1481 | PA1481 | PA1481 | PA1481 | PA1481 | N/A | PA1481 | N/A |
| PA1482 | PA1482 | N/A | N/A | PA1482 | PA1482 | PA1482 | PA1482 | N/A | PA1482 | PA1482 | PA1482 | PA1482 | PA1482 | PA1482 | N/A | N/A | PA1482 | PA1482 |
| PA1483 | PA1483 | PA1483 | PA1483 | PA1483 | PA1483 | PA1483 | PA1483 | PA1483 | PA1483 | PA1483 | PA1483 | PA1483 | PA1483 | PA1483 | PA1483 | N/A | PA1483 | PA1483 |
| PA1486 | N/A | N/A | N/A | PA1486 | N/A | N/A | N/A | N/A | N/A | N/A | N/A | N/A | N/A | N/A | N/A | N/A | N/A | N/A |
| N/A | N/A | N/A | N/A | N/A | N/A | N/A | N/A | N/A | N/A | N/A | N/A | PA1487 | PA1487 | N/A | N/A | N/A | N/A | N/A |
| PA1488 | N/A | N/A | N/A | N/A | N/A | N/A | N/A | N/A | N/A | N/A | N/A | N/A | N/A | N/A | N/A | N/A | N/A | N/A |
| N/A | N/A | N/A | N/A | PA1489 | N/A | N/A | N/A | N/A | N/A | N/A | N/A | N/A | N/A | N/A | N/A | N/A | N/A | N/A |
| PA1490 | N/A | N/A | N/A | PA1490 | N/A | N/A | N/A | N/A | N/A | N/A | PA1490 | PA1490 | N/A | N/A | N/A | N/A | N/A | N/A |
| N/A | N/A | N/A | N/A | N/A | PA1492 | N/A | N/A | N/A | N/A | N/A | N/A | N/A | PA1492 | N/A | N/A | N/A | N/A | N/A |
| PA1493 | PA1493 | PA1493 | PA1493 | PA1493 | PA1493 | PA1493 | PA1493 | N/A | PA1493 | PA1493 | PA1493 | PA1493 | PA1493 | PA1493 | PA1493 | PA1493 | PA1493 | N/A |
| PA1494 | N/A | N/A | N/A | PA1494 | PA1494 | PA1494 | N/A | PA1494 | PA1494 | PA1494 | PA1494 | PA1494 | PA1494 | PA1494 | N/A | N/A | PA1494 | PA1494 |
| PA1495 | N/A | N/A | N/A | N/A | N/A | N/A | N/A | N/A | N/A | N/A | N/A | N/A | N/A | N/A | N/A | N/A | N/A | N/A |
| PA1496 | N/A | N/A | N/A | N/A | N/A | N/A | PA1496 | N/A | N/A | N/A | N/A | N/A | N/A | N/A | N/A | N/A | N/A | N/A |
| PA1501 | N/A | N/A | N/A | N/A | N/A | N/A | N/A | N/A | N/A | N/A | N/A | PA1501 | N/A | N/A | N/A | N/A | N/A | N/A |
| PA1504 | PA1504 | PA1504 | N/A | PA1504 | N/A | PA1504 | PA1504 | N/A | PA1504 | PA1504 | PA1504 | PA1504 | PA1504 | PA1504 | N/A | N/A | PA1504 | N/A |
| N/A | N/A | N/A | N/A | PA1505 | N/A | N/A | N/A | N/A | N/A | N/A | N/A | N/A | N/A | N/A | N/A | N/A | N/A | N/A |
| PA1509 | N/A | N/A | N/A | PA1509 | N/A | N/A | N/A | N/A | PA1509 | PA1509 | N/A | N/A | N/A | PA1509 | N/A | N/A | N/A | N/A |
| N/A | N/A | N/A | N/A | PA1511 | N/A | N/A | N/A | N/A | N/A | N/A | N/A | N/A | N/A | N/A | N/A | N/A | PA1511 | N/A |
| PA1512 | N/A | N/A | N/A | PA1512 | N/A | PA1512 | N/A | N/A | PA1512 | PA1512 | N/A | PA1512 | PA1512 | N/A | N/A | N/A | N/A | N/A |
| PA1513 | N/A | PA1513 | N/A | PA1513 | N/A | PA1513 | N/A | N/A | N/A | N/A | N/A | N/A | N/A | PA1513 | PA1513 | N/A | N/A | N/A |
| PA1514 | N/A | N/A | N/A | PA1514 | N/A | PA1514 | N/A | N/A | PA1514 | N/A | PA1514 | PA1514 | PA1514 | N/A | PA1514 | N/A | N/A | N/A |
| N/A | N/A | N/A | N/A | N/A | PA1515 | N/A | N/A | N/A | N/A | N/A | N/A | N/A | N/A | N/A | N/A | N/A | N/A | N/A |
| PA1516 | PA1516 | N/A | N/A | PA1516 | PA1516 | N/A | N/A | N/A | N/A | PA1516 | PA1516 | N/A | PA1516 | N/A | PA1516 | N/A | N/A | N/A |
| PA1517 | N/A | N/A | N/A | PA1517 | PA1517 | PA1517 | N/A | N/A | N/A | N/A | PA1517 | PA1517 | PA1517 | N/A | N/A | N/A | N/A | N/A |
| PA1518 | PA1518 | PA1518 | N/A | PA1518 | PA1518 | PA1518 | N/A | N/A | PA1518 | PA1518 | PA1518 | PA1518 | PA1518 | N/A | PA1518 | N/A | N/A | N/A |
| N/A | N/A | N/A | N/A | N/A | PA1519 | N/A | N/A | N/A | N/A | N/A | N/A | N/A | N/A | N/A | N/A | N/A | N/A | N/A |
| PA1520 | PA1520 | PA1520 | PA1520 | PA1520 | N/A | PA1520 | PA1520 | PA1520 | PA1520 | PA1520 | PA1520 | PA1520 | PA1520 | PA1520 | PA1520 | N/A | PA1520 | PA1520 |
| PA1521 | PA1521 | N/A | N/A | PA1521 | N/A | PA1521 | N/A | N/A | N/A | N/A | N/A | PA1521 | PA1521 | N/A | N/A | N/A | N/A | N/A |
| PA1523 | N/A | N/A | N/A | PA1523 | N/A | PA1523 | N/A | N/A | N/A | PA1523 | N/A | PA1523 | PA1523 | N/A | N/A | N/A | N/A | N/A |
| N/A | N/A | N/A | N/A | PA1524 | N/A | N/A | N/A | N/A | N/A | N/A | N/A | PA1524 | PA1524 | N/A | N/A | N/A | N/A | N/A |
| N/A | N/A | N/A | N/A | N/A | PA1525 | N/A | N/A | N/A | N/A | N/A | N/A | N/A | N/A | N/A | N/A | N/A | N/A | N/A |
| PA1526 | PA1526 | N/A | N/A | PA1526 | PA1526 | PA1526 | PA1526 | N/A | PA1526 | PA1526 | PA1526 | PA1526 | PA1526 | N/A | N/A | N/A | PA1526 | N/A |
| PA1527 | PA1527 | N/A | PA1527 | PA1527 | PA1527 | N/A | N/A | PA1527 | PA1527 | N/A | PA1527 | PA1527 | PA1527 | N/A | N/A | N/A | PA1527 | N/A |
| PA1528 | PA1528 | PA1528 | PA1528 | PA1528 | PA1528 | PA1528 | PA1528 | PA1528 | PA1528 | PA1528 | PA1528 | PA1528 | PA1528 | PA1528 | PA1528 | PA1528 | PA1528 | PA1528 |
| PA1529 | PA1529 | PA1529 | PA1529 | PA1529 | PA1529 | PA1529 | N/A | PA1529 | PA1529 | PA1529 | PA1529 | PA1529 | PA1529 | PA1529 | PA1529 | N/A | N/A | N/A |
| PA1530 | PA1530 | PA1530 | N/A | PA1530 | N/A | PA1530 | PA1530 | N/A | PA1530 | PA1530 | PA1530 | PA1530 | PA1530 | PA1530 | N/A | N/A | PA1530 | N/A |
| PA1532 | PA1532 | PA1532 | PA1532 | PA1532 | N/A | N/A | N/A | PA1532 | PA1532 | PA1532 | PA1532 | PA1532 | PA1532 | N/A | N/A | N/A | PA1532 | N/A |
| PA1533 | PA1533 | PA1533 | PA1533 | PA1533 | N/A | PA1533 | PA1533 | N/A | PA1533 | PA1533 | PA1533 | PA1533 | PA1533 | PA1533 | PA1533 | N/A | N/A | N/A |
| PA1534 | PA1534 | PA1534 | N/A | PA1534 | N/A | N/A | N/A | N/A | PA1534 | PA1534 | PA1534 | PA1534 | PA1534 | N/A | N/A | N/A | N/A | N/A |
| PA1539 | N/A | N/A | N/A | PA1539 | N/A | N/A | N/A | N/A | N/A | N/A | PA1539 | N/A | N/A | N/A | N/A | N/A | N/A | N/A |
| PA1543 | PA1543 | PA1543 | PA1543 | PA1543 | PA1543 | PA1543 | N/A | N/A | PA1543 | PA1543 | PA1543 | PA1543 | PA1543 | N/A | N/A | N/A | N/A | N/A |
| PA1544 | PA1544 | PA1544 | PA1544 | PA1544 | PA1544 | PA1544 | PA1544 | PA1544 | PA1544 | PA1544 | PA1544 | PA1544 | PA1544 | N/A | PA1544 | N/A | N/A | N/A |
| N/A | N/A | N/A | N/A | PA1545 | PA1545 | N/A | N/A | N/A | N/A | N/A | N/A | N/A | N/A | N/A | N/A | N/A | N/A | N/A |
| PA1546 | N/A | N/A | N/A | PA1546 | PA1546 | PA1546 | N/A | PA1546 | N/A | PA1546 | PA1546 | PA1546 | PA1546 | N/A | N/A | N/A | PA1546 | N/A |
| PA1547 | N/A | N/A | N/A | N/A | N/A | N/A | N/A | N/A | N/A | N/A | N/A | N/A | N/A | PA1547 | N/A | N/A | N/A | N/A |
| PA1548 | N/A | N/A | N/A | N/A | N/A | PA1548 | N/A | N/A | PA1548 | PA1548 | PA1548 | PA1548 | PA1548 | PA1548 | N/A | N/A | N/A | N/A |
| PA1549 | N/A | N/A | N/A | PA1549 | PA1549 | N/A | N/A | N/A | PA1549 | PA1549 | N/A | PA1549 | N/A | PA1549 | N/A | N/A | N/A | N/A |
| PA1550 | N/A | N/A | N/A | PA1550 | PA1550 | PA1550 | PA1550 | N/A | PA1550 | PA1550 | PA1550 | PA1550 | PA1550 | PA1550 | PA1550 | N/A | PA1550 | PA1550 |
| PA1551 | N/A | N/A | N/A | PA1551 | PA1551 | PA1551 | N/A | N/A | N/A | PA1551 | PA1551 | PA1551 | PA1551 | PA1551 | N/A | N/A | PA1551 | N/A |
| PA1552 | PA1552 | PA1552 | PA1552 | PA1552 | PA1552 | PA1552 | PA1552 | PA1552 | PA1552 | PA1552 | PA1552 | PA1552 | PA1552 | PA1552 | PA1552 | N/A | PA1552 | N/A |
| PA1552.1 | N/A | N/A | N/A | PA1552.1 | PA1552.1 | PA1552.1 | N/A | N/A | PA1552.1 | PA1552.1 | PA1552.1 | PA1552.1 | PA1552.1 | PA1552.1 | N/A | N/A | PA1552.1 | N/A |
| PA1553 | PA1553 | PA1553 | PA1553 | PA1553 | PA1553 | PA1553 | PA1553 | PA1553 | PA1553 | PA1553 | PA1553 | PA1553 | PA1553 | PA1553 | PA1553 | PA1553 | PA1553 | PA1553 |
| PA1554 | N/A | PA1554 | N/A | PA1554 | PA1554 | PA1554 | N/A | PA1554 | PA1554 | PA1554 | PA1554 | PA1554 | PA1554 | PA1554 | PA1554 | N/A | PA1554 | PA1554 |
| PA1555 | N/A | PA1555 | N/A | PA1555 | PA1555 | PA1555 | PA1555 | PA1555 | PA1555 | PA1555 | PA1555 | PA1555 | PA1555 | PA1555 | N/A | PA1555 | PA1555 | PA1555 |
| PA1555.1 | N/A | N/A | N/A | PA1555.1 | PA1555.1 | PA1555.1 | N/A | N/A | PA1555.1 | PA1555.1 | PA1555.1 | PA1555.1 | PA1555.1 | PA1555.1 | N/A | N/A | PA1555.1 | N/A |
| PA1556 | N/A | PA1556 | PA1556 | PA1556 | PA1556 | PA1556 | PA1556 | PA1556 | PA1556 | PA1556 | PA1556 | PA1556 | PA1556 | PA1556 | PA1556 | N/A | PA1556 | PA1556 |
| PA1557 | N/A | PA1557 | N/A | PA1557 | N/A | PA1557 | N/A | PA1557 | PA1557 | PA1557 | PA1557 | PA1557 | PA1557 | PA1557 | PA1557 | N/A | PA1557 | PA1557 |
| PA1558 | N/A | N/A | N/A | PA1558 | N/A | N/A | N/A | N/A | PA1558 | PA1558 | PA1558 | PA1558 | PA1558 | N/A | N/A | N/A | N/A | N/A |
| N/A | N/A | N/A | N/A | N/A | PA1560 | N/A | N/A | N/A | N/A | N/A | N/A | N/A | N/A | N/A | N/A | N/A | N/A | N/A |
| PA1561 | PA1561 | PA1561 | PA1561 | PA1561 | PA1561 | PA1561 | PA1561 | PA1561 | PA1561 | PA1561 | PA1561 | PA1561 | PA1561 | PA1561 | PA1561 | N/A | PA1561 | PA1561 |
| PA1562 | PA1562 | PA1562 | PA1562 | PA1562 | PA1562 | PA1562 | N/A | PA1562 | PA1562 | PA1562 | PA1562 | PA1562 | PA1562 | PA1562 | PA1562 | PA1562 | N/A | N/A |
| PA1563 | N/A | N/A | PA1563 | PA1563 | PA1563 | N/A | N/A | PA1563 | PA1563 | N/A | N/A | N/A | PA1563 | PA1563 | N/A | N/A | N/A | N/A |
| PA1564 | N/A | PA1564 | N/A | PA1564 | N/A | N/A | N/A | N/A | N/A | N/A | N/A | N/A | N/A | N/A | N/A | N/A | N/A | N/A |
| PA1566 | N/A | N/A | N/A | N/A | N/A | N/A | N/A | PA1566 | N/A | N/A | PA1566 | N/A | N/A | N/A | N/A | N/A | N/A | N/A |
| PA1570 | N/A | N/A | N/A | PA1570 | N/A | N/A | N/A | N/A | N/A | N/A | PA1570 | N/A | N/A | N/A | N/A | N/A | N/A | N/A |
| N/A | N/A | N/A | N/A | PA1571 | N/A | N/A | N/A | N/A | N/A | N/A | PA1571 | PA1571 | N/A | N/A | N/A | N/A | N/A | N/A |
| PA1572 | N/A | N/A | N/A | PA1572 | N/A | N/A | N/A | N/A | N/A | N/A | N/A | N/A | N/A | N/A | N/A | N/A | N/A | N/A |
| N/A | N/A | N/A | N/A | N/A | PA1573 | N/A | N/A | N/A | N/A | N/A | N/A | N/A | N/A | N/A | N/A | N/A | N/A | N/A |
| PA1574 | N/A | PA1574 | N/A | PA1574 | PA1574 | PA1574 | N/A | N/A | PA1574 | PA1574 | PA1574 | PA1574 | PA1574 | N/A | N/A | N/A | N/A | N/A |
| PA1575 | N/A | N/A | N/A | N/A | PA1575 | N/A | N/A | N/A | N/A | N/A | N/A | N/A | N/A | N/A | N/A | N/A | N/A | N/A |
| PA1576 | N/A | N/A | N/A | PA1576 | PA1576 | PA1576 | N/A | N/A | N/A | PA1576 | PA1576 | PA1576 | PA1576 | N/A | N/A | N/A | N/A | N/A |
| PA1577 | N/A | N/A | N/A | N/A | N/A | N/A | N/A | N/A | N/A | N/A | PA1577 | N/A | N/A | N/A | N/A | N/A | N/A | N/A |
| PA1578 | N/A | N/A | N/A | N/A | PA1578 | N/A | N/A | N/A | N/A | N/A | N/A | N/A | N/A | PA1578 | N/A | N/A | N/A | N/A |
| PA1579 | PA1579 | PA1579 | PA1579 | PA1579 | PA1579 | PA1579 | PA1579 | N/A | PA1579 | PA1579 | PA1579 | PA1579 | PA1579 | N/A | PA1579 | PA1579 | N/A | N/A |
| PA1580 | PA1580 | PA1580 | PA1580 | PA1580 | PA1580 | PA1580 | PA1580 | PA1580 | PA1580 | PA1580 | PA1580 | PA1580 | PA1580 | PA1580 | PA1580 | PA1580 | N/A | N/A |
| PA1581 | PA1581 | PA1581 | PA1581 | PA1581 | N/A | PA1581 | PA1581 | PA1581 | PA1581 | PA1581 | PA1581 | PA1581 | PA1581 | PA1581 | PA1581 | PA1581 | PA1581 | N/A |
| PA1582 | PA1582 | PA1582 | PA1582 | PA1582 | PA1582 | PA1582 | N/A | PA1582 | PA1582 | PA1582 | PA1582 | PA1582 | PA1582 | PA1582 | PA1582 | N/A | PA1582 | PA1582 |
| PA1583 | PA1583 | PA1583 | PA1583 | PA1583 | PA1583 | PA1583 | PA1583 | PA1583 | PA1583 | PA1583 | PA1583 | PA1583 | PA1583 | PA1583 | PA1583 | PA1583 | PA1583 | PA1583 |

| | | | | | | | | | | | | | | | | | | |
|---|---|---|---|---|---|---|---|---|---|---|---|---|---|---|---|---|---|---|
| PA1584 | PA1584 | PA1584 | PA1584 | PA1584 | PA1584 | PA1584 | PA1584 | PA1584 | PA1584 | PA1584 | PA1584 | PA1584 | PA1584 | PA1584 | PA1584 | PA1584 | PA1584 | PA1584 |
| PA1585 | PA1585 | PA1585 | PA1585 | PA1585 | PA1585 | PA1585 | PA1585 | PA1585 | PA1585 | PA1585 | PA1585 | PA1585 | PA1585 | PA1585 | PA1585 | PA1585 | N/A | N/A |
| PA1586 | PA1586 | PA1586 | PA1586 | PA1586 | PA1586 | PA1586 | PA1586 | PA1586 | PA1586 | PA1586 | PA1586 | PA1586 | PA1586 | PA1586 | PA1586 | PA1586 | PA1586 | PA1586 |
| PA1587 | PA1587 | PA1587 | PA1587 | PA1587 | PA1587 | PA1587 | PA1587 | PA1587 | PA1587 | PA1587 | PA1587 | PA1587 | PA1587 | PA1587 | PA1587 | PA1587 | PA1587 | PA1587 |
| PA1588 | PA1588 | PA1588 | PA1588 | PA1588 | PA1588 | PA1588 | PA1588 | PA1588 | PA1588 | PA1588 | PA1588 | PA1588 | PA1588 | PA1588 | PA1588 | PA1588 | PA1588 | PA1588 |
| PA1589 | PA1589 | PA1589 | PA1589 | PA1589 | PA1589 | PA1589 | PA1589 | PA1589 | PA1589 | PA1589 | PA1589 | PA1589 | PA1589 | PA1589 | PA1589 | PA1589 | PA1589 | PA1589 |
| PA1590 | N/A | N/A | N/A | PA1590 | N/A | N/A | N/A | N/A | N/A | N/A | N/A | PA1590 | N/A | PA1590 | N/A | N/A | PA1590 | N/A |
| PA1591 | N/A | N/A | N/A | PA1591 | N/A | N/A | N/A | N/A | N/A | N/A | N/A | N/A | N/A | PA1591 | N/A | N/A | N/A | N/A |
| PA1592 | N/A | N/A | N/A | N/A | N/A | PA1592 | N/A | N/A | PA1592 | PA1592 | N/A | PA1592 | PA1592 | N/A | N/A | N/A | PA1592 | N/A |
| PA1593 | N/A | N/A | N/A | PA1593 | N/A | N/A | N/A | N/A | PA1593 | PA1593 | N/A | N/A | PA1593 | N/A | N/A | N/A | N/A | N/A |
| PA1594 | N/A | N/A | N/A | N/A | N/A | N/A | N/A | N/A | PA1594 | PA1594 | PA1594 | PA1594 | PA1594 | N/A | N/A | N/A | N/A | N/A |
| PA1595 | N/A | N/A | N/A | N/A | PA1595 | N/A | N/A | N/A | N/A | N/A | N/A | N/A | N/A | N/A | N/A | N/A | N/A | N/A |
| PA1596 | PA1596 | PA1596 | PA1596 | PA1596 | PA1596 | PA1596 | PA1596 | PA1596 | PA1596 | PA1596 | PA1596 | PA1596 | PA1596 | PA1596 | PA1596 | PA1596 | PA1596 | PA1596 |
| PA1597 | PA1597 | PA1597 | PA1597 | PA1597 | PA1597 | PA1597 | N/A | N/A | PA1597 | PA1597 | PA1597 | PA1597 | PA1597 | N/A | N/A | N/A | N/A | N/A |
| N/A | PA1600 | N/A | N/A | PA1600 | PA1600 | N/A | N/A | N/A | N/A | N/A | N/A | N/A | N/A | PA1600 | N/A | N/A | PA1600 | N/A |
| PA1601 | N/A | N/A | PA1601 | PA1601 | PA1601 | N/A | N/A | PA1601 | PA1601 | PA1601 | PA1601 | PA1601 | PA1601 | PA1601 | N/A | N/A | PA1601 | N/A |
| PA1602 | PA1602 | PA1602 | N/A | PA1602 | N/A | N/A | N/A | N/A | PA1602 | PA1602 | PA1602 | PA1602 | PA1602 | N/A | N/A | N/A | N/A | N/A |
| PA1603 | N/A | N/A | N/A | N/A | PA1603 | N/A | N/A | N/A | N/A | N/A | N/A | N/A | N/A | N/A | N/A | N/A | N/A | N/A |
| PA1604 | N/A | N/A | N/A | N/A | PA1604 | N/A | N/A | N/A | N/A | N/A | N/A | N/A | N/A | PA1604 | N/A | N/A | N/A | N/A |
| PA1605 | N/A | N/A | N/A | N/A | PA1605 | N/A | N/A | N/A | N/A | N/A | N/A | N/A | N/A | N/A | N/A | N/A | N/A | N/A |
| PA1606 | N/A | N/A | N/A | N/A | N/A | N/A | N/A | N/A | N/A | N/A | N/A | N/A | N/A | N/A | N/A | N/A | N/A | N/A |
| PA1607 | N/A | N/A | N/A | PA1607 | N/A | N/A | N/A | N/A | PA1607 | PA1607 | N/A | PA1607 | PA1607 | N/A | N/A | N/A | N/A | N/A |
| PA1608 | PA1608 | PA1608 | PA1608 | PA1608 | PA1608 | PA1608 | PA1608 | PA1608 | PA1608 | PA1608 | PA1608 | PA1608 | PA1608 | PA1608 | PA1608 | N/A | PA1608 | PA1608 |
| PA1609 | PA1609 | PA1609 | PA1609 | PA1609 | PA1609 | PA1609 | PA1609 | PA1609 | PA1609 | PA1609 | PA1609 | PA1609 | PA1609 | PA1609 | PA1609 | N/A | PA1609 | N/A |
| PA1610 | PA1610 | PA1610 | PA1610 | PA1610 | PA1610 | PA1610 | PA1610 | N/A | PA1610 | PA1610 | PA1610 | PA1610 | PA1610 | PA1610 | PA1610 | N/A | PA1610 | PA1610 |
| PA1611 | N/A | N/A | N/A | PA1611 | N/A | N/A | N/A | N/A | N/A | N/A | N/A | N/A | N/A | PA1611 | N/A | N/A | N/A | N/A |
| PA1612 | N/A | N/A | N/A | PA1612 | N/A | N/A | N/A | N/A | N/A | N/A | N/A | N/A | N/A | PA1612 | N/A | N/A | N/A | N/A |
| PA1613 | N/A | N/A | N/A | N/A | PA1613 | N/A | N/A | N/A | N/A | N/A | N/A | N/A | N/A | N/A | N/A | N/A | N/A | N/A |
| PA1614 | PA1614 | PA1614 | PA1614 | PA1614 | PA1614 | PA1614 | PA1614 | PA1614 | PA1614 | PA1614 | PA1614 | PA1614 | PA1614 | N/A | PA1614 | N/A | N/A | N/A |
| PA1615 | N/A | N/A | N/A | N/A | PA1615 | PA1615 | PA1615 | N/A | PA1615 | PA1615 | PA1615 | PA1615 | PA1615 | PA1615 | PA1615 | N/A | N/A | N/A |
| PA1616 | N/A | PA1616 | N/A | PA1616 | PA1616 | N/A | PA1616 | N/A | PA1616 | PA1616 | PA1616 | PA1616 | PA1616 | N/A | PA1616 | N/A | N/A | N/A |
| PA1617 | N/A | N/A | N/A | N/A | N/A | PA1617 | N/A | N/A | N/A | N/A | N/A | PA1617 | PA1617 | N/A | N/A | N/A | N/A | N/A |
| PA1618 | PA1618 | N/A | N/A | PA1618 | N/A | PA1618 | N/A | N/A | PA1618 | PA1618 | PA1618 | PA1618 | PA1618 | N/A | N/A | N/A | N/A | N/A |
| PA1622 | N/A | N/A | N/A | N/A | PA1622 | N/A | N/A | N/A | N/A | N/A | N/A | N/A | N/A | N/A | N/A | N/A | N/A | N/A |
| PA1623 | PA1623 | N/A | N/A | PA1623 | PA1623 | PA1623 | N/A | N/A | PA1623 | PA1623 | PA1623 | PA1623 | PA1623 | N/A | PA1623 | N/A | PA1623 | PA1623 |
| PA1624 | N/A | N/A | N/A | PA1624 | PA1624 | N/A | N/A | N/A | PA1624 | PA1624 | PA1624 | PA1624 | PA1624 | PA1624 | N/A | N/A | PA1624 | N/A |
| PA1625 | N/A | N/A | N/A | N/A | N/A | N/A | N/A | N/A | N/A | N/A | N/A | N/A | N/A | N/A | N/A | N/A | N/A | N/A |
| N/A | N/A | N/A | N/A | N/A | N/A | N/A | N/A | N/A | N/A | N/A | N/A | PA1626 | N/A | N/A | N/A | N/A | N/A | N/A |
| PA1627 | N/A | N/A | N/A | N/A | N/A | N/A | N/A | N/A | PA1627 | N/A | N/A | N/A | N/A | N/A | N/A | N/A | N/A | N/A |
| PA1628 | N/A | N/A | N/A | N/A | N/A | N/A | N/A | N/A | N/A | N/A | N/A | N/A | PA1628 | N/A | N/A | N/A | N/A | N/A |
| N/A | N/A | N/A | N/A | PA1630 | N/A | N/A | N/A | N/A | N/A | N/A | N/A | N/A | N/A | N/A | N/A | N/A | N/A | N/A |
| N/A | N/A | N/A | N/A | N/A | N/A | N/A | N/A | N/A | N/A | N/A | PA1631 | N/A | N/A | N/A | N/A | N/A | N/A | N/A |
| N/A | N/A | N/A | N/A | PA1633 | N/A | N/A | N/A | N/A | N/A | PA1633 | N/A | N/A | N/A | PA1633 | N/A | N/A | N/A | N/A |
| PA1634 | N/A | N/A | N/A | PA1634 | N/A | PA1634 | N/A | N/A | PA1634 | PA1634 | N/A | N/A | N/A | PA1634 | PA1634 | N/A | PA1634 | N/A |
| PA1635 | N/A | N/A | N/A | PA1635 | N/A | N/A | N/A | N/A | N/A | PA1635 | N/A | N/A | N/A | N/A | N/A | N/A | N/A | N/A |
| PA1636 | N/A | N/A | N/A | PA1636 | N/A | N/A | N/A | N/A | N/A | PA1636 | N/A | N/A | N/A | PA1636 | N/A | N/A | N/A | N/A |
| PA1637 | N/A | N/A | N/A | N/A | N/A | N/A | N/A | N/A | N/A | PA1637 | N/A | N/A | N/A | N/A | N/A | N/A | N/A | N/A |
| PA1638 | N/A | N/A | N/A | PA1638 | N/A | N/A | N/A | N/A | N/A | N/A | PA1638 | N/A | N/A | N/A | N/A | N/A | N/A | N/A |
| PA1639 | N/A | N/A | N/A | PA1639 | N/A | N/A | N/A | N/A | PA1639 | N/A | PA1639 | N/A | PA1639 | N/A | PA1639 | N/A | N/A | N/A |
| PA1640 | N/A | N/A | N/A | PA1640 | N/A | N/A | N/A | N/A | N/A | PA1640 | PA1640 | N/A | N/A | N/A | N/A | N/A | N/A | N/A |
| PA1641 | N/A | N/A | N/A | N/A | PA1641 | PA1641 | N/A | N/A | N/A | PA1641 | PA1641 | PA1641 | PA1641 | PA1641 | N/A | N/A | N/A | N/A |
| PA1642 | PA1642 | PA1642 | PA1642 | PA1642 | PA1642 | PA1642 | PA1642 | PA1642 | PA1642 | PA1642 | PA1642 | PA1642 | PA1642 | N/A | PA1642 | N/A | PA1642 | N/A |
| PA1643 | N/A | N/A | N/A | PA1643 | PA1643 | N/A | N/A | PA1643 | PA1643 | PA1643 | N/A | N/A | N/A | N/A | N/A | N/A | N/A | N/A |
| PA1644 | PA1644 | N/A | PA1644 | PA1644 | PA1644 | N/A | N/A | N/A | PA1644 | N/A | PA1644 | PA1644 | N/A | N/A | N/A | N/A | N/A | N/A |
| PA1645 | N/A | N/A | N/A | PA1645 | PA1645 | N/A | N/A | N/A | N/A | N/A | PA1645 | N/A | PA1645 | N/A | PA1645 | N/A | N/A | N/A |
| PA1646 | PA1646 | PA1646 | PA1646 | PA1646 | N/A | PA1646 | PA1646 | PA1646 | PA1646 | PA1646 | PA1646 | PA1646 | PA1646 | PA1646 | PA1646 | N/A | PA1646 | PA1646 |
| PA1647 | N/A | N/A | N/A | N/A | PA1647 | N/A | N/A | N/A | N/A | PA1647 | N/A | N/A | N/A | PA1647 | N/A | N/A | N/A | N/A |
| PA1648 | N/A | N/A | N/A | PA1648 | PA1648 | PA1648 | N/A | N/A | N/A | PA1648 | PA1648 | PA1648 | PA1648 | N/A | N/A | N/A | N/A | N/A |
| PA1649 | N/A | N/A | N/A | N/A | N/A | N/A | N/A | N/A | N/A | N/A | N/A | N/A | N/A | N/A | N/A | N/A | N/A | N/A |
| PA1651 | N/A | N/A | N/A | PA1651 | N/A | N/A | N/A | N/A | N/A | N/A | N/A | PA1651 | N/A | PA1651 | N/A | N/A | N/A | N/A |
| PA1652 | N/A | N/A | N/A | N/A | N/A | PA1652 | N/A | N/A | PA1652 | PA1652 | PA1652 | PA1652 | PA1652 | PA1652 | N/A | N/A | N/A | N/A |
| PA1653 | N/A | N/A | N/A | N/A | PA1653 | N/A | N/A | N/A | PA1653 | N/A | N/A | N/A | N/A | N/A | N/A | N/A | N/A | N/A |
| PA1654 | PA1654 | PA1654 | PA1654 | PA1654 | PA1654 | N/A | PA1654 | PA1654 | PA1654 | PA1654 | PA1654 | PA1654 | PA1654 | PA1654 | N/A | N/A | N/A | N/A |
| PA1655 | N/A | N/A | N/A | PA1655 | PA1655 | N/A | N/A | N/A | PA1655 | PA1655 | PA1655 | PA1655 | PA1655 | N/A | N/A | N/A | N/A | N/A |
| N/A | N/A | N/A | N/A | N/A | PA1656 | N/A | N/A | N/A | PA1656 | N/A | N/A | N/A | N/A | N/A | N/A | N/A | PA1656 | N/A |
| N/A | N/A | N/A | N/A | PA1657 | PA1657 | N/A | N/A | N/A | PA1657 | PA1657 | PA1657 | PA1657 | PA1657 | N/A | N/A | N/A | PA1657 | N/A |
| PA1658 | N/A | N/A | N/A | PA1658 | PA1658 | PA1658 | PA1658 | PA1658 | PA1658 | PA1658 | PA1658 | PA1658 | PA1658 | N/A | N/A | N/A | PA1658 | N/A |
| N/A | N/A | N/A | N/A | N/A | N/A | N/A | N/A | N/A | PA1660 | N/A | N/A | N/A | N/A | N/A | N/A | N/A | PA1660 | N/A |
| N/A | N/A | N/A | N/A | N/A | PA1661 | N/A | N/A | N/A | N/A | N/A | N/A | N/A | N/A | N/A | N/A | N/A | PA1661 | N/A |
| PA1662 | N/A | N/A | N/A | PA1662 | PA1662 | N/A | N/A | PA1662 | PA1662 | N/A | PA1662 | PA1662 | PA1662 | N/A | N/A | N/A | PA1662 | N/A |
| PA1663 | N/A | PA1663 | PA1663 | N/A | N/A | N/A | N/A | PA1663 | PA1663 | PA1663 | N/A | PA1663 | N/A | N/A | N/A | N/A | PA1663 | N/A |
| N/A | N/A | N/A | N/A | N/A | N/A | N/A | N/A | N/A | N/A | N/A | PA1664 | N/A | N/A | N/A | N/A | N/A | N/A | N/A |
| N/A | N/A | N/A | N/A | N/A | N/A | N/A | N/A | N/A | N/A | N/A | PA1665 | N/A | N/A | PA1665 | N/A | N/A | PA1665 | N/A |
| PA1666 | N/A | N/A | N/A | N/A | N/A | N/A | N/A | N/A | PA1666 | N/A | PA1666 | N/A | N/A | PA1666 | N/A | N/A | PA1666 | N/A |
| N/A | N/A | N/A | N/A | N/A | N/A | N/A | N/A | N/A | N/A | N/A | PA1667 | N/A | PA1667 | N/A | N/A | N/A | PA1667 | N/A |
| N/A | N/A | N/A | N/A | N/A | N/A | N/A | N/A | N/A | PA1668 | N/A | PA1668 | N/A | N/A | PA1668 | N/A | N/A | PA1668 | N/A |
| N/A | N/A | N/A | N/A | PA1669 | N/A | N/A | N/A | PA1669 | PA1669 | N/A | PA1669 | N/A | N/A | PA1669 | N/A | N/A | PA1669 | N/A |
| N/A | N/A | N/A | N/A | N/A | PA1672 | N/A | N/A | N/A | N/A | N/A | PA1672 | N/A | N/A | N/A | N/A | N/A | N/A | N/A |
| PA1673 | N/A | N/A | N/A | PA1673 | PA1673 | PA1673 | N/A | N/A | PA1673 | PA1673 | PA1673 | PA1673 | PA1673 | PA1673 | N/A | N/A | N/A | N/A |
| PA1674 | PA1674 | PA1674 | PA1674 | PA1674 | N/A | PA1674 | N/A | N/A | PA1674 | PA1674 | PA1674 | PA1674 | PA1674 | PA1674 | PA1674 | N/A | N/A | N/A |
| PA1675 | N/A | N/A | N/A | PA1675 | N/A | N/A | PA1675 | N/A | PA1675 | PA1675 | PA1675 | PA1675 | PA1675 | PA1675 | N/A | N/A | N/A | N/A |
| PA1676 | N/A | N/A | N/A | N/A | PA1676 | PA1676 | N/A | N/A | N/A | PA1676 | N/A | N/A | N/A | PA1676 | N/A | N/A | N/A | N/A |
| PA1677 | PA1677 | PA1677 | PA1677 | PA1677 | N/A | PA1677 | PA1677 | N/A | PA1677 | PA1677 | PA1677 | PA1677 | PA1677 | N/A | PA1677 | N/A | N/A | N/A |
| PA1678 | PA1678 | PA1678 | PA1678 | PA1678 | N/A | N/A | N/A | PA1678 | PA1678 | PA1678 | PA1678 | PA1678 | PA1678 | PA1678 | N/A | N/A | N/A | N/A |
| PA1679 | N/A | N/A | N/A | PA1679 | N/A | N/A | N/A | N/A | N/A | N/A | N/A | N/A | N/A | PA1679 | N/A | N/A | N/A | N/A |
| PA1680 | N/A | N/A | N/A | N/A | PA1680 | N/A | N/A | N/A | N/A | N/A | N/A | N/A | N/A | N/A | N/A | N/A | N/A | PA1680 |
| PA1681 | PA1681 | PA1681 | PA1681 | PA1681 | PA1681 | PA1681 | PA1681 | N/A | PA1681 | PA1681 | PA1681 | PA1681 | PA1681 | PA1681 | PA1681 | N/A | N/A | N/A |
| N/A | N/A | N/A | N/A | PA1682 | N/A | N/A | N/A | N/A | N/A | N/A | N/A | N/A | N/A | N/A | N/A | N/A | N/A | N/A |
| PA1683 | N/A | N/A | N/A | PA1683 | PA1683 | PA1683 | N/A | N/A | PA1683 | PA1683 | PA1683 | PA1683 | PA1683 | N/A | N/A | N/A | N/A | N/A |
| PA1684 | PA1684 | PA1684 | PA1684 | PA1684 | N/A | N/A | N/A | N/A | PA1684 | PA1684 | PA1684 | PA1684 | PA1684 | N/A | PA1684 | N/A | N/A | N/A |
| PA1685 | PA1685 | PA1685 | PA1685 | PA1685 | PA1685 | PA1685 | N/A | N/A | PA1685 | PA1685 | PA1685 | PA1685 | PA1685 | N/A | PA1685 | N/A | N/A | N/A |
| PA1686 | N/A | N/A | N/A | PA1686 | N/A | N/A | N/A | N/A | PA1686 | N/A | PA1686 | PA1686 | PA1686 | N/A | N/A | N/A | N/A | N/A |
| PA1687 | PA1687 | PA1687 | PA1687 | PA1687 | PA1687 | N/A | PA1687 | PA1687 | PA1687 | PA1687 | PA1687 | PA1687 | PA1687 | N/A | PA1687 | N/A | N/A | N/A |
| PA1688 | N/A | N/A | N/A | PA1688 | N/A | N/A | N/A | N/A | N/A | N/A | N/A | N/A | N/A | N/A | N/A | N/A | N/A | N/A |
| PA1689 | PA1689 | PA1689 | PA1689 | PA1689 | N/A | PA1689 | PA1689 | PA1689 | PA1689 | PA1689 | PA1689 | PA1689 | PA1689 | PA1689 | PA1689 | N/A | PA1689 | N/A |
| N/A | N/A | N/A | N/A | N/A | N/A | N/A | N/A | N/A | N/A | PA1690 | N/A | N/A | N/A | N/A | N/A | N/A | N/A | N/A |
| N/A | N/A | N/A | N/A | N/A | N/A | N/A | PA1691 | N/A | N/A | N/A | N/A | N/A | N/A | N/A | N/A | N/A | N/A | N/A |
| N/A | N/A | N/A | N/A | PA1695 | N/A | N/A | N/A | N/A | N/A | PA1695 | N/A | N/A | N/A | N/A | N/A | N/A | N/A | N/A |
| N/A | N/A | N/A | N/A | N/A | N/A | N/A | N/A | N/A | N/A | PA1698 | N/A | N/A | N/A | N/A | N/A | N/A | N/A | N/A |
| N/A | PA1706 | PA1706 | N/A | PA1706 | N/A | N/A | N/A | N/A | N/A | PA1706 | N/A | N/A | N/A | N/A | PA1706 | PA1706 | N/A | N/A |
| N/A | PA1707 | N/A | PA1707 | PA1707 | N/A | N/A | N/A | N/A | N/A | PA1707 | PA1707 | N/A | N/A | N/A | N/A | N/A | N/A | N/A |
| N/A | N/A | N/A | N/A | PA1708 | N/A | N/A | N/A | N/A | N/A | PA1708 | N/A | N/A | N/A | N/A | N/A | N/A | N/A | N/A |
| N/A | PA1709 | N/A | N/A | PA1709 | N/A | N/A | N/A | N/A | N/A | PA1709 | PA1709 | N/A | N/A | N/A | N/A | N/A | PA1709 | PA1709 |
| N/A | N/A | N/A | N/A | N/A | N/A | N/A | N/A | N/A | PA1710 | PA1710 | PA1710 | N/A | N/A | N/A | N/A | N/A | N/A | N/A |
| N/A | N/A | N/A | N/A | N/A | N/A | N/A | N/A | N/A | N/A | PA1712 | N/A | N/A | N/A | N/A | N/A | N/A | N/A | N/A |
| PA1713 | N/A | N/A | N/A | N/A | N/A | N/A | N/A | N/A | N/A | PA1713 | PA1713 | N/A | N/A | N/A | N/A | N/A | N/A | N/A |
| N/A | PA1714 | PA1714 | PA1714 | PA1714 | N/A | N/A | N/A | N/A | N/A | PA1714 | PA1714 | PA1714 | N/A | N/A | N/A | N/A | N/A | N/A |
| N/A | PA1715 | N/A | N/A | PA1715 | N/A | N/A | N/A | N/A | N/A | PA1715 | PA1715 | N/A | N/A | N/A | N/A | N/A | N/A | N/A |
| N/A | N/A | N/A | N/A | N/A | N/A | N/A | N/A | N/A | N/A | PA1716 | N/A | N/A | N/A | N/A | N/A | N/A | N/A | N/A |

| | | | | | | | | | | | | | | | | | | |
|---|---|---|---|---|---|---|---|---|---|---|---|---|---|---|---|---|---|---|
| N/A | N/A | N/A | N/A | PA1717 | N/A | N/A | N/A | N/A | N/A | N/A | N/A | N/A | N/A | N/A | N/A | N/A | N/A | N/A |
| N/A | PA1718 | N/A | PA1718 | PA1718 | N/A | N/A | N/A | N/A | N/A | N/A | N/A | N/A | N/A | N/A | N/A | N/A | N/A | N/A |
| N/A | N/A | PA1719 | N/A | N/A | N/A | N/A | N/A | N/A | N/A | N/A | N/A | N/A | N/A | N/A | N/A | N/A | N/A | N/A |
| N/A | N/A | N/A | N/A | PA1720 | N/A | N/A | N/A | N/A | N/A | N/A | PA1720 | N/A | N/A | N/A | N/A | N/A | N/A | N/A |
| N/A | N/A | N/A | PA1723 | PA1723 | N/A | N/A | N/A | N/A | N/A | PA1723 | N/A | N/A | N/A | N/A | PA1723 | N/A | N/A | N/A |
| PA1726 | N/A | N/A | N/A | PA1726 | N/A | N/A | N/A | N/A | PA1726 | N/A | N/A | N/A | N/A | N/A | N/A | N/A | N/A | N/A |
| PA1728 | N/A | N/A | N/A | N/A | PA1728 | N/A | N/A | N/A | N/A | N/A | N/A | PA1728 | N/A | N/A | N/A | N/A | N/A | N/A |
| N/A | N/A | N/A | N/A | N/A | PA1729 | N/A | N/A | N/A | N/A | N/A | N/A | N/A | N/A | N/A | N/A | N/A | N/A | N/A |
| PA1730 | N/A | N/A | N/A | N/A | N/A | N/A | N/A | N/A | N/A | N/A | N/A | N/A | PA1730 | N/A | N/A | N/A | N/A | N/A |
| N/A | N/A | N/A | N/A | N/A | N/A | N/A | N/A | N/A | N/A | N/A | N/A | N/A | PA1731 | N/A | N/A | N/A | N/A | N/A |
| N/A | N/A | N/A | N/A | N/A | N/A | N/A | N/A | N/A | N/A | N/A | N/A | N/A | N/A | N/A | N/A | N/A | PA1732 | N/A |
| PA1733 | N/A | N/A | N/A | N/A | N/A | N/A | N/A | N/A | N/A | PA1733 | N/A | PA1733 | PA1733 | N/A | N/A | N/A | N/A | N/A |
| N/A | N/A | N/A | N/A | PA1734 | N/A | N/A | N/A | N/A | N/A | N/A | N/A | N/A | N/A | N/A | N/A | N/A | N/A | N/A |
| PA1736 | N/A | N/A | N/A | PA1736 | N/A | N/A | N/A | N/A | N/A | N/A | N/A | N/A | N/A | N/A | N/A | N/A | N/A | N/A |
| PA1737 | N/A | N/A | N/A | N/A | N/A | PA1737 | N/A | N/A | N/A | PA1737 | PA1737 | PA1737 | PA1737 | N/A | N/A | N/A | N/A | N/A |
| PA1738 | N/A | N/A | N/A | PA1738 | N/A | N/A | N/A | N/A | N/A | N/A | N/A | N/A | N/A | N/A | N/A | N/A | N/A | N/A |
| PA1741 | N/A | N/A | N/A | N/A | PA1741 | N/A | N/A | N/A | PA1741 | N/A | N/A | N/A | N/A | N/A | N/A | N/A | N/A | N/A |
| PA1742 | PA1742 | PA1742 | PA1742 | PA1742 | PA1742 | PA1742 | PA1742 | PA1742 | PA1742 | PA1742 | PA1742 | PA1742 | PA1742 | N/A | PA1742 | N/A | N/A | N/A |
| PA1745 | N/A | N/A | N/A | PA1745 | PA1745 | PA1745 | N/A | N/A | N/A | PA1745 | PA1745 | PA1745 | PA1745 | N/A | N/A | N/A | N/A | N/A |
| PA1746 | PA1746 | N/A | N/A | PA1746 | PA1746 | PA1746 | PA1746 | N/A | PA1746 | PA1746 | PA1746 | PA1746 | PA1746 | PA1746 | N/A | N/A | N/A | N/A |
| N/A | N/A | N/A | N/A | N/A | PA1747 | N/A | N/A | N/A | N/A | N/A | N/A | N/A | N/A | N/A | N/A | N/A | N/A | N/A |
| PA1748 | PA1748 | PA1748 | PA1748 | PA1748 | PA1748 | PA1748 | PA1748 | N/A | PA1748 | PA1748 | PA1748 | PA1748 | PA1748 | N/A | PA1748 | PA1748 | N/A | N/A |
| PA1749 | PA1749 | PA1749 | PA1749 | PA1749 | PA1749 | PA1749 | N/A | N/A | PA1749 | PA1749 | PA1749 | PA1749 | PA1749 | N/A | PA1749 | N/A | N/A | N/A |
| PA1750 | PA1750 | PA1750 | PA1750 | PA1750 | PA1750 | PA1750 | PA1750 | PA1750 | PA1750 | PA1750 | PA1750 | PA1750 | PA1750 | PA1750 | PA1750 | PA1750 | N/A | N/A |
| PA1752 | N/A | N/A | N/A | PA1752 | N/A | N/A | N/A | N/A | N/A | PA1752 | PA1752 | PA1752 | PA1752 | N/A | N/A | N/A | N/A | N/A |
| PA1753 | N/A | N/A | N/A | PA1753 | PA1753 | N/A | N/A | N/A | N/A | PA1753 | N/A | PA1753 | N/A | N/A | N/A | N/A | N/A | N/A |
| PA1754 | PA1754 | PA1754 | PA1754 | PA1754 | PA1754 | PA1754 | N/A | PA1754 | PA1754 | PA1754 | PA1754 | PA1754 | PA1754 | PA1754 | N/A | PA1754 | N/A | N/A |
| PA1755 | N/A | N/A | N/A | N/A | PA1755 | N/A | N/A | N/A | N/A | N/A | N/A | N/A | N/A | N/A | N/A | N/A | N/A | N/A |
| PA1756 | PA1756 | PA1756 | N/A | PA1756 | N/A | N/A | N/A | N/A | PA1756 | PA1756 | PA1756 | PA1756 | PA1756 | N/A | PA1756 | N/A | N/A | N/A |
| PA1757 | PA1757 | PA1757 | PA1757 | PA1757 | PA1757 | PA1757 | N/A | N/A | PA1757 | PA1757 | PA1757 | PA1757 | N/A | N/A | N/A | N/A | N/A | N/A |
| PA1758 | PA1758 | PA1758 | N/A | PA1758 | PA1758 | N/A | N/A | PA1758 | N/A | N/A | N/A | N/A | PA1758 | N/A | N/A | N/A | N/A | N/A |
| PA1759 | N/A | N/A | N/A | PA1759 | N/A | N/A | N/A | N/A | N/A | N/A | N/A | N/A | N/A | N/A | N/A | N/A | N/A | N/A |
| PA1760 | N/A | PA1760 | N/A | PA1760 | N/A | N/A | N/A | N/A | N/A | N/A | PA1760 | PA1760 | PA1760 | N/A | N/A | N/A | N/A | N/A |
| PA1766 | PA1766 | PA1766 | PA1766 | PA1766 | PA1766 | PA1766 | PA1766 | PA1766 | PA1766 | PA1766 | PA1766 | PA1766 | PA1766 | PA1766 | PA1766 | N/A | PA1766 | N/A |
| PA1767 | PA1767 | PA1767 | PA1767 | PA1767 | PA1767 | PA1767 | N/A | PA1767 | PA1767 | PA1767 | PA1767 | PA1767 | PA1767 | PA1767 | PA1767 | N/A | PA1767 | N/A |
| PA1768 | PA1768 | PA1768 | PA1768 | PA1768 | PA1768 | PA1768 | PA1768 | N/A | PA1768 | PA1768 | PA1768 | PA1768 | PA1768 | PA1768 | PA1768 | N/A | PA1768 | PA1768 |
| PA1769 | N/A | PA1769 | N/A | PA1769 | N/A | PA1769 | N/A | N/A | PA1769 | PA1769 | PA1769 | PA1769 | PA1769 | N/A | N/A | N/A | N/A | N/A |
| PA1770 | PA1770 | PA1770 | PA1770 | PA1770 | PA1770 | PA1770 | PA1770 | PA1770 | PA1770 | PA1770 | PA1770 | PA1770 | PA1770 | PA1770 | PA1770 | PA1770 | N/A | N/A |
| PA1771 | N/A | N/A | N/A | PA1771 | PA1771 | N/A | PA1771 | PA1771 | N/A | N/A | N/A | N/A | N/A | N/A | N/A | N/A | PA1771 | N/A |
| PA1772 | PA1772 | PA1772 | PA1772 | PA1772 | PA1772 | PA1772 | N/A | N/A | PA1772 | PA1772 | PA1772 | PA1772 | PA1772 | N/A | PA1772 | N/A | PA1772 | PA1772 |
| PA1773 | N/A | N/A | N/A | N/A | N/A | N/A | N/A | N/A | PA1773 | N/A | PA1773 | PA1773 | PA1773 | PA1773 | N/A | N/A | N/A | N/A |
| PA1774 | N/A | N/A | N/A | N/A | N/A | PA1774 | N/A | N/A | N/A | N/A | N/A | N/A | N/A | N/A | N/A | N/A | N/A | N/A |
| PA1775 | PA1775 | PA1775 | PA1775 | PA1775 | PA1775 | PA1775 | N/A | PA1775 | PA1775 | PA1775 | PA1775 | PA1775 | PA1775 | PA1775 | PA1775 | N/A | PA1775 | N/A |
| N/A | N/A | N/A | N/A | PA1776 | PA1776 | N/A | N/A | N/A | N/A | N/A | N/A | N/A | N/A | N/A | N/A | N/A | N/A | N/A |
| PA1777 | PA1777 | PA1777 | PA1777 | PA1777 | PA1777 | PA1777 | PA1777 | PA1777 | PA1777 | PA1777 | PA1777 | PA1777 | PA1777 | PA1777 | PA1777 | PA1777 | PA1777 | PA1777 |
| PA1783 | N/A | N/A | N/A | N/A | N/A | N/A | N/A | N/A | N/A | N/A | N/A | N/A | N/A | N/A | N/A | N/A | N/A | N/A |
| PA1784 | N/A | N/A | N/A | N/A | N/A | N/A | N/A | N/A | N/A | N/A | N/A | N/A | PA1784 | N/A | PA1784 | N/A | N/A | N/A |
| N/A | N/A | N/A | N/A | N/A | PA1786 | N/A | N/A | N/A | N/A | N/A | N/A | N/A | N/A | N/A | N/A | N/A | N/A | N/A |
| PA1787 | PA1787 | PA1787 | PA1787 | PA1787 | PA1787 | PA1787 | PA1787 | PA1787 | PA1787 | PA1787 | PA1787 | PA1787 | PA1787 | PA1787 | PA1787 | PA1787 | PA1787 | PA1787 |
| PA1788 | N/A | N/A | N/A | PA1788 | PA1788 | N/A | N/A | N/A | PA1788 | PA1788 | PA1788 | PA1788 | N/A | N/A | N/A | N/A | PA1788 | N/A |
| PA1789 | PA1789 | N/A | N/A | PA1789 | PA1789 | PA1789 | N/A | N/A | PA1789 | PA1789 | PA1789 | PA1789 | PA1789 | PA1789 | N/A | N/A | N/A | N/A |
| PA1790 | N/A | N/A | N/A | PA1790 | PA1790 | N/A | N/A | N/A | PA1790 | PA1790 | PA1790 | N/A | N/A | N/A | N/A | N/A | N/A | N/A |
| PA1791 | PA1791 | PA1791 | N/A | PA1791 | N/A | N/A | PA1791 | PA1791 | PA1791 | N/A | PA1791 | N/A | PA1791 | PA1791 | PA1791 | N/A | PA1791 | PA1791 |
| PA1792 | N/A | N/A | N/A | PA1792 | PA1792 | PA1792 | N/A | N/A | PA1792 | N/A | PA1792 | PA1792 | PA1792 | PA1792 | N/A | N/A | PA1792 | N/A |
| PA1793 | PA1793 | PA1793 | PA1793 | PA1793 | PA1793 | PA1793 | PA1793 | N/A | PA1793 | PA1793 | PA1793 | PA1793 | PA1793 | PA1793 | PA1793 | PA1793 | N/A | N/A |
| PA1794 | PA1794 | PA1794 | PA1794 | PA1794 | PA1794 | PA1794 | PA1794 | PA1794 | PA1794 | PA1794 | PA1794 | PA1794 | PA1794 | N/A | PA1794 | N/A | N/A | N/A |
| PA1795 | PA1795 | PA1795 | PA1795 | PA1795 | PA1795 | PA1795 | PA1795 | PA1795 | PA1795 | PA1795 | PA1795 | PA1795 | PA1795 | N/A | PA1795 | PA1795 | N/A | N/A |
| PA1796 | PA1796 | PA1796 | PA1796 | PA1796 | PA1796 | PA1796 | PA1796 | N/A | PA1796 | PA1796 | PA1796 | PA1796 | PA1796 | N/A | PA1796 | N/A | PA1796 | PA1796 |
| PA1798 | N/A | N/A | N/A | PA1798 | PA1798 | N/A | N/A | N/A | N/A | N/A | N/A | N/A | N/A | PA1798 | N/A | N/A | N/A | N/A |
| PA1799 | PA1799 | PA1799 | N/A | PA1799 | PA1799 | N/A | N/A | N/A | PA1799 | N/A | PA1799 | N/A | N/A | N/A | N/A | N/A | N/A | N/A |
| PA1800 | PA1800 | PA1800 | PA1800 | PA1800 | PA1800 | PA1800 | PA1800 | PA1800 | PA1800 | PA1800 | PA1800 | PA1800 | PA1800 | PA1800 | PA1800 | PA1800 | PA1800 | PA1800 |
| PA1801 | PA1801 | PA1801 | PA1801 | PA1801 | PA1801 | PA1801 | PA1801 | N/A | PA1801 | PA1801 | PA1801 | PA1801 | PA1801 | PA1801 | PA1801 | N/A | N/A | N/A |
| PA1802 | PA1802 | PA1802 | PA1802 | PA1802 | PA1802 | PA1802 | PA1802 | PA1802 | PA1802 | PA1802 | PA1802 | PA1802 | PA1802 | PA1802 | PA1802 | N/A | PA1802 | N/A |
| PA1803 | PA1803 | PA1803 | PA1803 | PA1803 | PA1803 | PA1803 | PA1803 | PA1803 | PA1803 | PA1803 | PA1803 | PA1803 | PA1803 | PA1803 | PA1803 | PA1803 | N/A | PA1803 |
| PA1804 | PA1804 | PA1804 | PA1804 | PA1804 | PA1804 | PA1804 | PA1804 | PA1804 | PA1804 | PA1804 | PA1804 | PA1804 | PA1804 | PA1804 | PA1804 | PA1804 | PA1804 | PA1804 |
| PA1805 | PA1805 | PA1805 | PA1805 | PA1805 | PA1805 | PA1805 | PA1805 | PA1805 | PA1805 | PA1805 | PA1805 | PA1805 | PA1805 | PA1805 | PA1805 | PA1805 | PA1805 | PA1805 |
| PA1806 | PA1806 | PA1806 | PA1806 | PA1806 | N/A | PA1806 | N/A | N/A | PA1806 | PA1806 | PA1806 | PA1806 | PA1806 | N/A | PA1806 | N/A | N/A | N/A |
| PA1807 | PA1807 | N/A | PA1807 | PA1807 | N/A | PA1807 | N/A | PA1807 | PA1807 | PA1807 | PA1807 | PA1807 | PA1807 | PA1807 | N/A | N/A | PA1807 | N/A |
| PA1808 | N/A | N/A | N/A | PA1808 | N/A | N/A | N/A | N/A | N/A | N/A | N/A | N/A | N/A | PA1808 | N/A | N/A | PA1808 | N/A |
| PA1809 | N/A | N/A | N/A | PA1809 | PA1809 | N/A | N/A | N/A | N/A | PA1809 | N/A | PA1809 | N/A | PA1809 | N/A | N/A | PA1809 | N/A |
| PA1810 | PA1810 | PA1810 | PA1810 | PA1810 | PA1810 | N/A | N/A | N/A | PA1810 | PA1810 | PA1810 | PA1810 | PA1810 | PA1810 | PA1810 | N/A | N/A | N/A |
| PA1811 | N/A | N/A | N/A | N/A | N/A | N/A | N/A | N/A | N/A | N/A | N/A | N/A | N/A | N/A | N/A | N/A | N/A | N/A |
| PA1812 | N/A | N/A | N/A | PA1812 | PA1812 | N/A | N/A | PA1812 | PA1812 | PA1812 | N/A | N/A | N/A | PA1812 | N/A | N/A | N/A | N/A |
| PA1813 | N/A | PA1813 | N/A | PA1813 | PA1813 | PA1813 | N/A | N/A | PA1813 | PA1813 | PA1813 | PA1813 | PA1813 | N/A | N/A | N/A | N/A | N/A |
| PA1814 | N/A | N/A | N/A | N/A | PA1814 | N/A | N/A | N/A | N/A | PA1814 | PA1814 | PA1814 | PA1814 | PA1814 | N/A | N/A | N/A | N/A |
| PA1815 | N/A | N/A | N/A | PA1815 | N/A | PA1815 | PA1815 | N/A | PA1815 | PA1815 | PA1815 | N/A | PA1815 | N/A | N/A | N/A | N/A | N/A |
| PA1816 | PA1816 | PA1816 | PA1816 | PA1816 | N/A | PA1816 | PA1816 | N/A | PA1816 | PA1816 | PA1816 | PA1816 | PA1816 | N/A | PA1816 | PA1816 | N/A | N/A |
| PA1817 | N/A | N/A | N/A | PA1817 | PA1817 | PA1817 | N/A | N/A | PA1817 | PA1817 | PA1817 | N/A | PA1817 | N/A | N/A | N/A | N/A | N/A |
| PA1818 | PA1818 | PA1818 | PA1818 | PA1818 | PA1818 | PA1818 | N/A | PA1818 | PA1818 | PA1818 | PA1818 | PA1818 | PA1818 | PA1818 | PA1818 | N/A | N/A | N/A |
| PA1819 | N/A | N/A | N/A | N/A | N/A | PA1819 | N/A | N/A | N/A | PA1819 | N/A | N/A | PA1819 | PA1819 | N/A | N/A | PA1819 | N/A |
| PA1820 | N/A | N/A | N/A | N/A | PA1820 | PA1820 | N/A | N/A | N/A | PA1820 | N/A | PA1820 | PA1820 | PA1820 | N/A | N/A | PA1820 | N/A |
| PA1821 | PA1821 | PA1821 | PA1821 | PA1821 | PA1821 | PA1821 | PA1821 | N/A | PA1821 | PA1821 | PA1821 | PA1821 | PA1821 | PA1821 | N/A | N/A | PA1821 | N/A |
| PA1822 | PA1822 | PA1822 | PA1822 | PA1822 | PA1822 | N/A | PA1822 | PA1822 | PA1822 | PA1822 | PA1822 | PA1822 | PA1822 | PA1822 | PA1822 | N/A | N/A | N/A |
| PA1823 | N/A | N/A | N/A | PA1823 | N/A | N/A | N/A | N/A | PA1823 | PA1823 | PA1823 | PA1823 | PA1823 | N/A | N/A | N/A | N/A | N/A |
| PA1824 | N/A | N/A | N/A | N/A | N/A | N/A | N/A | N/A | N/A | N/A | N/A | N/A | N/A | N/A | N/A | N/A | N/A | N/A |
| PA1825 | N/A | PA1825 | PA1825 | PA1825 | N/A | PA1825 | N/A | N/A | PA1825 | PA1825 | PA1825 | PA1825 | PA1825 | PA1825 | N/A | N/A | PA1825 | N/A |
| N/A | N/A | N/A | N/A | N/A | PA1827 | N/A | N/A | N/A | N/A | N/A | N/A | N/A | N/A | N/A | N/A | N/A | N/A | N/A |
| PA1828 | PA1828 | PA1828 | PA1828 | PA1828 | PA1828 | PA1828 | N/A | N/A | PA1828 | PA1828 | PA1828 | PA1828 | PA1828 | N/A | N/A | N/A | N/A | N/A |
| PA1829 | N/A | N/A | N/A | PA1829 | PA1829 | N/A | N/A | N/A | N/A | PA1829 | PA1829 | PA1829 | PA1829 | N/A | N/A | N/A | N/A | N/A |
| PA1830 | PA1830 | PA1830 | PA1830 | PA1830 | PA1830 | PA1830 | N/A | N/A | N/A | N/A | PA1830 | N/A | PA1830 | N/A | PA1830 | N/A | N/A | N/A |
| PA1831 | PA1831 | N/A | N/A | PA1831 | N/A | N/A | N/A | N/A | N/A | PA1831 | PA1831 | N/A | N/A | N/A | N/A | N/A | N/A | N/A |
| PA1832 | N/A | PA1832 | PA1832 | PA1832 | PA1832 | PA1832 | PA1832 | N/A | PA1832 | PA1832 | PA1832 | PA1832 | PA1832 | PA1832 | N/A | N/A | PA1832 | N/A |
| PA1833 | PA1833 | PA1833 | PA1833 | PA1833 | PA1833 | PA1833 | PA1833 | PA1833 | PA1833 | PA1833 | PA1833 | PA1833 | PA1833 | PA1833 | N/A | N/A | PA1833 | PA1833 |
| N/A | N/A | N/A | N/A | N/A | N/A | N/A | N/A | N/A | N/A | N/A | N/A | PA1835 | PA1835 | N/A | N/A | N/A | N/A | N/A |
| PA1836 | N/A | N/A | N/A | N/A | PA1836 | N/A | N/A | N/A | N/A | N/A | N/A | N/A | N/A | N/A | N/A | N/A | N/A | N/A |
| PA1837 | PA1837 | PA1837 | PA1837 | PA1837 | PA1837 | N/A | PA1837 | N/A | PA1837 | PA1837 | PA1837 | PA1837 | PA1837 | N/A | PA1837 | N/A | N/A | N/A |
| PA1838 | PA1838 | PA1838 | PA1838 | PA1838 | PA1838 | PA1838 | N/A | PA1838 | PA1838 | PA1838 | PA1838 | PA1838 | PA1838 | N/A | PA1838 | N/A | N/A | N/A |
| PA1840 | N/A | PA1840 | N/A | PA1840 | N/A | N/A | PA1840 | N/A | PA1840 | PA1840 | PA1840 | PA1840 | N/A | PA1840 | N/A | N/A | N/A | N/A |
| PA1841 | N/A | N/A | N/A | PA1841 | PA1841 | N/A | PA1841 | N/A | N/A | N/A | PA1841 | N/A | N/A | N/A | N/A | N/A | N/A | N/A |
| PA1842 | N/A | PA1842 | N/A | PA1842 | PA1842 | N/A | PA1842 | N/A | N/A | N/A | PA1842 | PA1842 | PA1842 | PA1842 | N/A | N/A | N/A | N/A |
| PA1843 | PA1843 | PA1843 | PA1843 | PA1843 | N/A | PA1843 | N/A | PA1843 | PA1843 | PA1843 | PA1843 | PA1843 | PA1843 | N/A | PA1843 | N/A | N/A | N/A |
| PA1844 | N/A | N/A | N/A | N/A | N/A | N/A | N/A | N/A | N/A | N/A | N/A | N/A | N/A | N/A | N/A | N/A | PA1844 | N/A |
| PA1845 | N/A | N/A | N/A | N/A | PA1845 | N/A | N/A | N/A | N/A | N/A | N/A | N/A | N/A | N/A | PA1845 | N/A | N/A | N/A |
| PA1846 | N/A | N/A | N/A | PA1846 | N/A | N/A | N/A | N/A | PA1846 | N/A | PA1846 | N/A | N/A | N/A | N/A | N/A | N/A | N/A |
| PA1847 | PA1847 | PA1847 | PA1847 | PA1847 | PA1847 | PA1847 | PA1847 | N/A | PA1847 | PA1847 | PA1847 | PA1847 | PA1847 | PA1847 | PA1847 | N/A | PA1847 | PA1847 |
| N/A | N/A | N/A | N/A | N/A | PA1851 | N/A | N/A | N/A | N/A | N/A | N/A | N/A | N/A | N/A | N/A | N/A | N/A | N/A |
| PA1852 | N/A | PA1852 | N/A | PA1852 | PA1852 | N/A | N/A | N/A | PA1852 | N/A | PA1852 | PA1852 | PA1852 | N/A | N/A | N/A | N/A | N/A |
| PA1853 | N/A | N/A | N/A | N/A | N/A | N/A | N/A | N/A | N/A | N/A | N/A | N/A | N/A | N/A | N/A | N/A | N/A | N/A |
| PA1856 | N/A | N/A | N/A | N/A | N/A | PA1856 | N/A | N/A | N/A | PA1856 | N/A | N/A | N/A | PA1856 | N/A | N/A | N/A | N/A |

| | | | | | | | | | | | | | | | | | | |
|---|---|---|---|---|---|---|---|---|---|---|---|---|---|---|---|---|---|---|
| PA1857 | N/A | N/A | PA1857 | PA1857 | N/A | N/A | N/A | PA1857 | N/A | N/A | PA1857 | PA1857 | PA1857 | PA1857 | N/A | N/A | PA1857 | N/A |
| PA1859 | N/A | N/A | N/A | PA1859 | N/A | N/A | N/A | N/A | N/A | N/A | PA1859 | N/A | N/A | N/A | N/A | N/A | N/A | N/A |
| PA1860 | N/A | N/A | N/A | N/A | N/A | PA1860 | N/A | N/A | N/A | PA1860 | N/A | PA1860 | PA1860 | N/A | N/A | N/A | N/A | N/A |
| PA1861 | N/A | N/A | N/A | N/A | N/A | N/A | N/A | N/A | N/A | PA1861 | N/A | N/A | N/A | N/A | N/A | N/A | N/A | N/A |
| N/A | N/A | N/A | N/A | N/A | PA1862 | N/A | N/A | N/A | N/A | N/A | N/A | N/A | N/A | N/A | N/A | N/A | N/A | N/A |
| PA1863 | N/A | PA1863 | N/A | PA1863 | PA1863 | PA1863 | N/A | N/A | PA1863 | PA1863 | PA1863 | PA1863 | PA1863 | N/A | PA1863 | N/A | N/A | N/A |
| PA1865 | N/A | N/A | N/A | N/A | N/A | N/A | N/A | N/A | N/A | N/A | N/A | N/A | N/A | N/A | N/A | N/A | N/A | N/A |
| PA1866 | N/A | N/A | N/A | PA1866 | N/A | N/A | N/A | N/A | N/A | N/A | N/A | N/A | N/A | N/A | N/A | N/A | N/A | N/A |
| PA1868 | N/A | N/A | N/A | PA1868 | N/A | PA1868 | N/A | PA1868 | PA1868 | PA1868 | N/A | PA1868 | PA1868 | PA1868 | N/A | N/A | N/A | N/A |
| PA1869 | N/A | N/A | N/A | N/A | N/A | N/A | N/A | N/A | N/A | N/A | N/A | N/A | N/A | N/A | N/A | N/A | N/A | N/A |
| N/A | N/A | N/A | N/A | N/A | N/A | N/A | N/A | N/A | N/A | PA1870 | N/A | N/A | N/A | N/A | N/A | N/A | N/A | N/A |
| PA1871 | N/A | N/A | N/A | N/A | N/A | PA1871 | PA1871 | N/A | PA1871 | PA1871 | PA1871 | PA1871 | PA1871 | N/A | PA1871 | PA1871 | N/A | N/A |
| PA1872 | N/A | N/A | N/A | PA1872 | N/A | N/A | N/A | N/A | N/A | N/A | N/A | N/A | N/A | PA1872 | N/A | N/A | PA1872 | N/A |
| PA1874 | N/A | N/A | N/A | N/A | N/A | PA1874 | N/A | N/A | N/A | N/A | N/A | PA1874 | PA1874 | N/A | N/A | N/A | N/A | N/A |
| PA1875 | N/A | N/A | N/A | N/A | N/A | PA1875 | N/A | N/A | N/A | N/A | N/A | PA1875 | PA1875 | N/A | N/A | N/A | N/A | N/A |
| PA1876 | N/A | N/A | N/A | N/A | N/A | N/A | N/A | N/A | N/A | N/A | PA1876 | PA1876 | PA1876 | N/A | N/A | N/A | N/A | N/A |
| PA1877 | N/A | N/A | N/A | N/A | N/A | N/A | N/A | N/A | N/A | N/A | N/A | N/A | PA1877 | N/A | N/A | N/A | N/A | N/A |
| N/A | N/A | N/A | N/A | N/A | PA1879 | N/A | N/A | N/A | N/A | N/A | N/A | N/A | N/A | N/A | N/A | N/A | N/A | N/A |
| PA1880 | N/A | N/A | N/A | PA1880 | PA1880 | PA1880 | N/A | N/A | PA1880 | PA1880 | PA1880 | PA1880 | PA1880 | N/A | N/A | N/A | N/A | N/A |
| PA1881 | N/A | N/A | N/A | N/A | N/A | PA1881 | N/A | N/A | N/A | PA1881 | PA1881 | PA1881 | PA1881 | N/A | N/A | N/A | N/A | N/A |
| PA1884 | N/A | N/A | N/A | N/A | N/A | N/A | N/A | N/A | N/A | N/A | N/A | N/A | N/A | N/A | N/A | N/A | N/A | N/A |
| PA1885 | N/A | N/A | N/A | N/A | N/A | N/A | N/A | N/A | N/A | N/A | N/A | N/A | N/A | N/A | N/A | N/A | N/A | N/A |
| PA1886 | N/A | N/A | N/A | N/A | N/A | N/A | N/A | PA1886 | N/A | N/A | N/A | N/A | N/A | N/A | N/A | N/A | N/A | N/A |
| PA1887 | N/A | N/A | N/A | N/A | N/A | N/A | N/A | N/A | N/A | N/A | N/A | N/A | N/A | N/A | N/A | N/A | N/A | N/A |
| PA1888 | N/A | N/A | N/A | N/A | N/A | N/A | N/A | N/A | N/A | N/A | N/A | N/A | N/A | N/A | N/A | N/A | N/A | N/A |
| PA1889 | N/A | N/A | N/A | PA1889 | PA1889 | PA1889 | N/A | N/A | PA1889 | PA1889 | PA1889 | PA1889 | PA1889 | PA1889 | PA1889 | N/A | PA1889 | N/A |
| PA1890 | N/A | N/A | N/A | PA1890 | PA1890 | N/A | N/A | N/A | PA1890 | PA1890 | PA1890 | PA1890 | PA1890 | N/A | N/A | N/A | N/A | N/A |
| PA1893 | N/A | N/A | N/A | N/A | N/A | N/A | N/A | N/A | N/A | N/A | N/A | N/A | N/A | N/A | N/A | N/A | N/A | N/A |
| PA1894 | N/A | N/A | N/A | N/A | N/A | N/A | N/A | N/A | PA1894 | N/A | PA1894 | PA1894 | N/A | N/A | N/A | N/A | N/A | N/A |
| N/A | N/A | N/A | N/A | N/A | N/A | N/A | N/A | N/A | N/A | N/A | N/A | PA1895 | N/A | N/A | N/A | N/A | N/A | N/A |
| PA1896 | N/A | N/A | N/A | N/A | N/A | N/A | N/A | N/A | N/A | N/A | N/A | N/A | N/A | N/A | N/A | N/A | N/A | N/A |
| PA1897 | N/A | N/A | N/A | N/A | N/A | N/A | N/A | N/A | N/A | N/A | N/A | N/A | N/A | N/A | N/A | N/A | N/A | N/A |
| N/A | N/A | N/A | N/A | N/A | PA1898 | N/A | N/A | N/A | N/A | N/A | N/A | N/A | N/A | N/A | N/A | N/A | N/A | N/A |
| PA1899 | N/A | N/A | N/A | N/A | N/A | N/A | N/A | N/A | N/A | N/A | PA1899 | PA1899 | PA1899 | N/A | N/A | N/A | N/A | N/A |
| PA1900 | N/A | N/A | N/A | N/A | N/A | PA1900 | N/A | N/A | PA1900 | PA1900 | PA1900 | PA1900 | PA1900 | N/A | N/A | PA1900 | N/A | N/A |
| PA1901 | N/A | N/A | N/A | N/A | N/A | N/A | N/A | N/A | PA1901 | N/A | PA1901 | PA1901 | PA1901 | N/A | N/A | N/A | N/A | N/A |
| PA1902 | N/A | N/A | N/A | N/A | N/A | PA1902 | N/A | N/A | PA1902 | PA1902 | PA1902 | PA1902 | PA1902 | N/A | N/A | N/A | N/A | N/A |
| PA1903 | N/A | N/A | N/A | N/A | N/A | PA1903 | N/A | N/A | N/A | PA1903 | PA1903 | PA1903 | PA1903 | N/A | N/A | N/A | N/A | N/A |
| PA1904 | PA1904 | PA1904 | N/A | PA1904 | N/A | PA1904 | N/A | N/A | PA1904 | PA1904 | PA1904 | PA1904 | PA1904 | N/A | N/A | PA1904 | N/A | N/A |
| PA1905 | N/A | N/A | N/A | N/A | N/A | PA1905 | N/A | N/A | PA1905 | PA1905 | PA1905 | PA1905 | PA1905 | N/A | N/A | N/A | N/A | N/A |
| PA1910 | N/A | N/A | N/A | N/A | N/A | N/A | N/A | N/A | N/A | N/A | N/A | N/A | N/A | N/A | N/A | N/A | N/A | N/A |
| PA1913 | N/A | N/A | N/A | PA1913 | PA1913 | N/A | N/A | N/A | PA1913 | N/A | PA1913 | PA1913 | PA1913 | N/A | N/A | N/A | N/A | N/A |
| PA1914 | N/A | N/A | N/A | N/A | N/A | N/A | N/A | N/A | N/A | N/A | N/A | N/A | PA1914 | N/A | N/A | N/A | PA1914 | N/A |
| N/A | N/A | N/A | N/A | N/A | N/A | N/A | N/A | N/A | N/A | N/A | PA1924 | N/A | N/A | N/A | N/A | N/A | N/A | N/A |
| PA1926 | PA1926 | PA1926 | PA1926 | PA1926 | PA1926 | N/A | N/A | PA1926 | PA1926 | N/A | PA1926 | PA1926 | PA1926 | N/A | PA1926 | N/A | N/A | N/A |
| PA1927 | N/A | N/A | N/A | N/A | PA1927 | N/A | N/A | N/A | N/A | N/A | N/A | N/A | N/A | N/A | N/A | N/A | N/A | N/A |
| PA1928 | N/A | N/A | N/A | N/A | N/A | N/A | N/A | N/A | N/A | N/A | N/A | N/A | N/A | N/A | N/A | N/A | N/A | N/A |
| N/A | N/A | N/A | N/A | N/A | N/A | N/A | N/A | N/A | N/A | N/A | N/A | PA1929 | PA1929 | N/A | N/A | N/A | N/A | N/A |
| PA1930 | PA1930 | PA1930 | PA1930 | PA1930 | N/A | PA1930 | PA1930 | PA1930 | PA1930 | PA1930 | PA1930 | PA1930 | PA1930 | PA1930 | PA1930 | N/A | PA1930 | PA1930 |
| N/A | N/A | N/A | N/A | N/A | PA1933 | N/A | N/A | N/A | N/A | N/A | N/A | PA1933 | PA1933 | N/A | N/A | N/A | N/A | N/A |
| PA1934 | N/A | N/A | N/A | N/A | PA1934 | PA1934 | N/A | N/A | PA1934 | PA1934 | PA1934 | PA1934 | N/A | N/A | N/A | N/A | N/A | N/A |
| PA1939 | N/A | N/A | N/A | PA1939 | N/A | N/A | N/A | N/A | PA1939 | N/A | PA1939 | PA1939 | N/A | N/A | N/A | N/A | PA1939 | N/A |
| PA1940 | N/A | N/A | N/A | PA1940 | N/A | N/A | PA1940 | N/A | N/A | N/A | N/A | N/A | N/A | PA1940 | N/A | N/A | PA1940 | N/A |
| PA1941 | N/A | N/A | N/A | PA1941 | N/A | N/A | N/A | N/A | N/A | N/A | N/A | N/A | PA1941 | PA1941 | N/A | N/A | PA1941 | N/A |
| PA1943 | N/A | N/A | N/A | N/A | PA1943 | N/A | N/A | N/A | N/A | PA1943 | N/A | N/A | N/A | N/A | N/A | N/A | N/A | N/A |
| PA1944 | N/A | N/A | N/A | N/A | PA1944 | PA1944 | N/A | N/A | N/A | PA1944 | N/A | PA1944 | PA1944 | N/A | N/A | N/A | N/A | N/A |
| N/A | N/A | N/A | N/A | N/A | PA1945 | N/A | N/A | N/A | N/A | N/A | N/A | N/A | N/A | N/A | N/A | N/A | N/A | N/A |
| PA1946 | N/A | N/A | N/A | N/A | PA1946 | PA1946 | N/A | N/A | N/A | N/A | PA1946 | PA1946 | PA1946 | N/A | N/A | N/A | N/A | N/A |
| PA1947 | N/A | PA1947 | N/A | N/A | N/A | PA1947 | N/A | N/A | N/A | PA1947 | N/A | N/A | PA1947 | N/A | N/A | N/A | N/A | N/A |
| PA1948 | N/A | N/A | N/A | N/A | N/A | PA1948 | N/A | N/A | N/A | N/A | N/A | PA1948 | PA1948 | PA1948 | N/A | N/A | N/A | N/A |
| PA1949 | N/A | N/A | N/A | PA1949 | PA1949 | N/A | N/A | N/A | N/A | N/A | PA1949 | PA1949 | PA1949 | N/A | N/A | N/A | N/A | N/A |
| PA1950 | N/A | N/A | N/A | PA1950 | PA1950 | N/A | N/A | N/A | N/A | PA1950 | PA1950 | PA1950 | PA1950 | N/A | N/A | N/A | N/A | N/A |
| PA1951 | N/A | N/A | N/A | N/A | N/A | N/A | N/A | N/A | N/A | N/A | N/A | N/A | PA1951 | N/A | N/A | N/A | N/A | N/A |
| PA1952 | N/A | N/A | N/A | N/A | N/A | N/A | N/A | N/A | N/A | N/A | N/A | N/A | N/A | N/A | N/A | N/A | N/A | N/A |
| PA1953 | N/A | N/A | N/A | N/A | N/A | N/A | N/A | N/A | N/A | N/A | N/A | N/A | N/A | N/A | N/A | N/A | N/A | N/A |
| PA1954 | N/A | N/A | N/A | N/A | N/A | N/A | N/A | N/A | N/A | N/A | N/A | N/A | N/A | N/A | N/A | N/A | N/A | N/A |
| PA1955 | N/A | N/A | N/A | N/A | N/A | N/A | N/A | N/A | N/A | N/A | N/A | N/A | N/A | N/A | N/A | N/A | N/A | N/A |
| PA1957 | N/A | N/A | N/A | N/A | N/A | N/A | N/A | N/A | N/A | N/A | N/A | N/A | N/A | N/A | N/A | N/A | N/A | N/A |
| PA1959 | N/A | N/A | N/A | N/A | N/A | N/A | N/A | N/A | PA1959 | N/A | PA1959 | PA1959 | PA1959 | PA1959 | N/A | N/A | N/A | N/A |
| PA1960 | N/A | N/A | N/A | N/A | N/A | N/A | N/A | N/A | N/A | N/A | N/A | N/A | N/A | PA1960 | N/A | N/A | N/A | N/A |
| N/A | N/A | N/A | N/A | PA1961 | N/A | N/A | N/A | PA1961 | N/A | N/A | N/A | N/A | N/A | N/A | N/A | N/A | N/A | N/A |
| PA1964 | PA1964 | PA1964 | PA1964 | PA1964 | N/A | N/A | PA1964 | PA1964 | PA1964 | PA1964 | PA1964 | PA1964 | PA1964 | N/A | N/A | N/A | N/A | N/A |
| PA1965 | N/A | N/A | N/A | PA1965 | PA1965 | N/A | PA1965 | N/A | PA1965 | PA1965 | PA1965 | PA1965 | PA1965 | PA1965 | N/A | N/A | PA1965 | N/A |
| PA1966 | N/A | N/A | N/A | N/A | PA1966 | N/A | N/A | N/A | PA1966 | N/A | PA1966 | PA1966 | N/A | N/A | N/A | N/A | N/A | N/A |
| N/A | N/A | N/A | N/A | N/A | PA1968 | N/A | N/A | N/A | N/A | N/A | N/A | N/A | N/A | N/A | N/A | N/A | N/A | N/A |
| PA1969 | PA1969 | PA1969 | PA1969 | PA1969 | PA1969 | PA1969 | PA1969 | N/A | PA1969 | PA1969 | PA1969 | PA1969 | PA1969 | PA1969 | PA1969 | N/A | PA1969 | N/A |
| PA1970 | N/A | N/A | N/A | N/A | N/A | N/A | N/A | N/A | N/A | N/A | N/A | N/A | N/A | N/A | N/A | N/A | N/A | N/A |
| PA1973 | N/A | N/A | N/A | N/A | N/A | N/A | N/A | N/A | N/A | N/A | N/A | N/A | N/A | N/A | N/A | N/A | N/A | N/A |
| N/A | N/A | N/A | N/A | N/A | N/A | PA1974 | N/A | N/A | N/A | N/A | N/A | N/A | N/A | N/A | N/A | N/A | N/A | N/A |
| N/A | N/A | N/A | N/A | N/A | N/A | PA1975 | N/A | N/A | N/A | N/A | N/A | N/A | N/A | N/A | N/A | N/A | N/A | N/A |
| N/A | N/A | N/A | N/A | N/A | N/A | PA1976 | N/A | N/A | N/A | N/A | N/A | N/A | N/A | N/A | N/A | N/A | N/A | N/A |
| N/A | N/A | N/A | N/A | N/A | N/A | PA1978 | N/A | N/A | N/A | N/A | N/A | N/A | N/A | N/A | N/A | N/A | N/A | N/A |
| N/A | N/A | N/A | N/A | N/A | N/A | PA1980 | N/A | N/A | N/A | N/A | N/A | N/A | N/A | N/A | N/A | N/A | N/A | N/A |
| PA1983 | N/A | N/A | N/A | N/A | N/A | PA1983 | N/A | N/A | N/A | N/A | N/A | N/A | N/A | N/A | N/A | N/A | N/A | N/A |
| PA1984 | PA1984 | PA1984 | PA1984 | PA1984 | N/A | PA1984 | PA1984 | PA1984 | PA1984 | PA1984 | PA1984 | PA1984 | PA1984 | PA1984 | PA1984 | N/A | N/A | N/A |
| N/A | N/A | N/A | N/A | N/A | PA1985 | N/A | N/A | N/A | N/A | N/A | N/A | N/A | N/A | N/A | N/A | N/A | N/A | N/A |
| PA1986 | N/A | N/A | N/A | N/A | N/A | PA1986 | N/A | N/A | N/A | N/A | N/A | N/A | N/A | N/A | N/A | N/A | N/A | N/A |
| N/A | N/A | N/A | N/A | N/A | N/A | PA1987 | N/A | N/A | N/A | N/A | N/A | N/A | N/A | N/A | N/A | N/A | N/A | N/A |
| PA1988 | N/A | N/A | N/A | PA1988 | N/A | PA1988 | N/A | N/A | N/A | N/A | PA1988 | N/A | PA1988 | N/A | N/A | N/A | N/A | N/A |
| PA1989 | N/A | N/A | N/A | N/A | N/A | PA1989 | N/A | N/A | N/A | N/A | N/A | N/A | N/A | N/A | N/A | N/A | N/A | N/A |
| N/A | N/A | N/A | N/A | PA1991 | N/A | PA1991 | N/A | N/A | N/A | N/A | N/A | N/A | N/A | N/A | N/A | N/A | N/A | N/A |
| PA1992 | N/A | N/A | N/A | N/A | N/A | PA1992 | N/A | N/A | N/A | N/A | N/A | N/A | N/A | PA1992 | N/A | N/A | N/A | N/A |
| PA1994 | N/A | N/A | N/A | PA1994 | N/A | N/A | N/A | N/A | N/A | N/A | N/A | PA1994 | PA1994 | N/A | N/A | N/A | N/A | N/A |
| PA1995 | N/A | N/A | N/A | N/A | N/A | N/A | N/A | N/A | PA1995 | N/A | PA1995 | PA1995 | PA1995 | N/A | N/A | N/A | N/A | N/A |
| PA1996 | N/A | N/A | N/A | PA1996 | N/A | N/A | N/A | N/A | N/A | N/A | PA1996 | PA1996 | N/A | N/A | N/A | N/A | N/A | N/A |
| PA1997 | N/A | N/A | N/A | N/A | N/A | PA1997 | N/A | N/A | N/A | N/A | PA1997 | PA1997 | PA1997 | N/A | N/A | N/A | N/A | N/A |
| PA1998 | N/A | N/A | N/A | PA1998 | PA1998 | N/A | N/A | N/A | N/A | N/A | N/A | N/A | N/A | N/A | N/A | N/A | N/A | N/A |
| PA1999 | N/A | N/A | N/A | PA1999 | PA1999 | PA1999 | N/A | N/A | PA1999 | PA1999 | PA1999 | PA1999 | PA1999 | N/A | PA1999 | N/A | N/A | N/A |
| PA2000 | N/A | N/A | N/A | PA2000 | PA2000 | PA2000 | N/A | N/A | PA2000 | PA2000 | PA2000 | PA2000 | PA2000 | N/A | PA2000 | N/A | N/A | N/A |
| PA2001 | PA2001 | PA2001 | PA2001 | PA2001 | PA2001 | PA2001 | N/A | N/A | PA2001 | PA2001 | PA2001 | PA2001 | PA2001 | PA2001 | PA2001 | PA2001 | N/A | N/A |
| PA2002 | N/A | N/A | N/A | PA2002 | PA2002 | PA2002 | N/A | N/A | N/A | N/A | N/A | PA2002 | N/A | PA2002 | PA2002 | N/A | N/A | N/A |
| PA2003 | N/A | N/A | N/A | N/A | PA2003 | PA2003 | N/A | N/A | N/A | PA2003 | PA2003 | PA2003 | PA2003 | N/A | N/A | N/A | N/A | N/A |
| PA2004 | N/A | N/A | N/A | N/A | N/A | PA2004 | N/A | N/A | N/A | N/A | N/A | N/A | PA2004 | PA2004 | N/A | N/A | N/A | N/A |
| PA2006 | N/A | N/A | N/A | N/A | PA2006 | N/A | N/A | N/A | N/A | N/A | N/A | N/A | N/A | N/A | N/A | N/A | N/A | N/A |
| PA2007 | N/A | N/A | N/A | PA2007 | PA2007 | PA2007 | PA2007 | PA2007 | PA2007 | PA2007 | PA2007 | PA2007 | PA2007 | N/A | PA2007 | N/A | N/A | N/A |
| PA2008 | PA2008 | PA2008 | PA2008 | PA2008 | PA2008 | PA2008 | PA2008 | PA2008 | PA2008 | PA2008 | PA2008 | PA2008 | PA2008 | N/A | PA2008 | PA2008 | N/A | N/A |
| PA2009 | PA2009 | PA2009 | PA2009 | PA2009 | PA2009 | PA2009 | PA2009 | PA2009 | PA2009 | PA2009 | PA2009 | PA2009 | PA2009 | PA2009 | PA2009 | PA2009 | N/A | N/A |
| PA2010 | N/A | N/A | N/A | PA2010 | N/A | PA2010 | N/A | N/A | N/A | N/A | PA2010 | PA2010 | PA2010 | N/A | N/A | N/A | N/A | N/A |
| PA2011 | N/A | N/A | PA2011 | PA2011 | PA2011 | PA2011 | N/A | N/A | N/A | PA2011 | PA2011 | PA2011 | PA2011 | N/A | N/A | N/A | N/A | N/A |

| | | | | | | | | | | | | | | | | | | |
|---|---|---|---|---|---|---|---|---|---|---|---|---|---|---|---|---|---|---|
| PA2012 | N/A | N/A | N/A | PA2012 | PA2012 | PA2012 | N/A | N/A | PA2012 | PA2012 | PA2012 | PA2012 | PA2012 | PA2012 | N/A | N/A | N/A | N/A |
| PA2013 | N/A | N/A | N/A | PA2013 | PA2013 | PA2013 | N/A | N/A | PA2013 | PA2013 | PA2013 | PA2013 | PA2013 | PA2013 | N/A | PA2013 | N/A | N/A |
| PA2014 | N/A | N/A | N/A | PA2014 | PA2014 | PA2014 | N/A | N/A | PA2014 | PA2014 | PA2014 | PA2014 | PA2014 | PA2014 | PA2014 | N/A | N/A | N/A |
| PA2015 | N/A | N/A | N/A | PA2015 | PA2015 | PA2015 | N/A | PA2015 | PA2015 | PA2015 | PA2015 | PA2015 | PA2015 | PA2015 | N/A | N/A | N/A | N/A |
| PA2016 | N/A | N/A | N/A | PA2016 | PA2016 | PA2016 | N/A | N/A | PA2016 | PA2016 | PA2016 | PA2016 | PA2016 | PA2016 | N/A | N/A | PA2016 | PA2016 |
| PA2018 | N/A | N/A | N/A | PA2018 | PA2018 | N/A | N/A | N/A | PA2018 | N/A | PA2018 | N/A | N/A | PA2018 | N/A | N/A | N/A | N/A |
| PA2019 | N/A | N/A | N/A | PA2019 | PA2019 | N/A | PA2019 | N/A | PA2019 | N/A | PA2019 | N/A | N/A | N/A | N/A | N/A | N/A | N/A |
| PA2020 | N/A | N/A | N/A | PA2020 | N/A | PA2020 | PA2020 | N/A | PA2020 | N/A | PA2020 | PA2020 | PA2020 | PA2020 | N/A | N/A | N/A | N/A |
| PA2021 | N/A | N/A | N/A | N/A | N/A | PA2021 | N/A | N/A | PA2021 | PA2021 | N/A | N/A | PA2021 | N/A | N/A | N/A | N/A | N/A |
| N/A | N/A | N/A | N/A | N/A | PA2022 | N/A | N/A | N/A | N/A | N/A | N/A | N/A | N/A | N/A | N/A | N/A | N/A | N/A |
| PA2023 | PA2023 | PA2023 | PA2023 | PA2023 | PA2023 | PA2023 | PA2023 | PA2023 | PA2023 | PA2023 | PA2023 | PA2023 | PA2023 | N/A | PA2023 | PA2023 | N/A | N/A |
| PA2024 | N/A | N/A | N/A | N/A | PA2024 | PA2024 | N/A | N/A | N/A | N/A | N/A | N/A | N/A | N/A | N/A | N/A | N/A | N/A |
| PA2025 | PA2025 | PA2025 | PA2025 | PA2025 | PA2025 | PA2025 | N/A | N/A | PA2025 | PA2025 | PA2025 | PA2025 | PA2025 | N/A | PA2025 | N/A | N/A | N/A |
| N/A | N/A | N/A | N/A | N/A | N/A | N/A | N/A | N/A | N/A | N/A | N/A | N/A | N/A | PA2026 | N/A | N/A | N/A | N/A |
| PA2028 | N/A | N/A | N/A | PA2028 | N/A | N/A | N/A | N/A | PA2028 | PA2028 | PA2028 | PA2028 | PA2028 | N/A | N/A | N/A | N/A | N/A |
| N/A | N/A | N/A | N/A | N/A | N/A | N/A | N/A | N/A | PA2029 | N/A | N/A | PA2029 | PA2029 | N/A | N/A | N/A | N/A | N/A |
| PA2031 | N/A | N/A | N/A | N/A | N/A | PA2031 | N/A | N/A | N/A | N/A | PA2031 | PA2031 | PA2031 | N/A | N/A | N/A | N/A | N/A |
| N/A | N/A | N/A | N/A | PA2032 | N/A | N/A | N/A | PA2032 | N/A | N/A | N/A | N/A | PA2032 | N/A | N/A | N/A | N/A | N/A |
| N/A | N/A | N/A | N/A | N/A | N/A | N/A | N/A | N/A | N/A | N/A | N/A | N/A | PA2033 | N/A | N/A | N/A | N/A | N/A |
| PA2034 | N/A | N/A | N/A | PA2034 | N/A | N/A | N/A | N/A | N/A | N/A | N/A | N/A | PA2034 | N/A | N/A | N/A | N/A | N/A |
| PA2035 | N/A | N/A | N/A | N/A | N/A | N/A | N/A | N/A | N/A | N/A | N/A | N/A | N/A | N/A | N/A | N/A | N/A | N/A |
| N/A | N/A | N/A | N/A | N/A | PA2039 | N/A | N/A | N/A | N/A | N/A | N/A | N/A | N/A | PA2039 | N/A | N/A | N/A | N/A |
| PA2040 | PA2040 | PA2040 | PA2040 | PA2040 | N/A | PA2040 | PA2040 | PA2040 | PA2040 | PA2040 | PA2040 | PA2040 | PA2040 | N/A | PA2040 | N/A | N/A | N/A |
| PA2042 | N/A | N/A | N/A | PA2042 | N/A | PA2042 | N/A | PA2042 | PA2042 | PA2042 | PA2042 | PA2042 | PA2042 | PA2042 | N/A | N/A | PA2042 | N/A |
| N/A | N/A | N/A | N/A | N/A | PA2043 | N/A | N/A | N/A | N/A | N/A | N/A | N/A | N/A | N/A | N/A | N/A | N/A | N/A |
| PA2044 | PA2044 | PA2044 | PA2044 | PA2044 | PA2044 | N/A | N/A | PA2044 | PA2044 | PA2044 | PA2044 | PA2044 | PA2044 | N/A | PA2044 | N/A | N/A | N/A |
| PA2047 | N/A | N/A | N/A | N/A | N/A | N/A | N/A | N/A | N/A | PA2047 | N/A | N/A | N/A | N/A | N/A | N/A | N/A | N/A |
| PA2049 | N/A | N/A | N/A | PA2049 | N/A | N/A | N/A | N/A | N/A | N/A | N/A | N/A | N/A | PA2049 | N/A | N/A | N/A | N/A |
| PA2051 | N/A | N/A | N/A | N/A | N/A | N/A | N/A | N/A | N/A | N/A | N/A | N/A | N/A | N/A | N/A | N/A | N/A | N/A |
| PA2052 | N/A | N/A | N/A | N/A | N/A | PA2052 | PA2052 | N/A | PA2052 | PA2052 | N/A | PA2052 | PA2052 | PA2052 | N/A | N/A | PA2052 | N/A |
| PA2054 | N/A | N/A | N/A | PA2054 | N/A | N/A | N/A | N/A | N/A | N/A | N/A | PA2054 | N/A | N/A | N/A | N/A | N/A | N/A |
| N/A | PA2055 | N/A | N/A | N/A | N/A | N/A | N/A | N/A | N/A | N/A | N/A | N/A | N/A | N/A | N/A | N/A | N/A | N/A |
| PA2056 | N/A | N/A | N/A | N/A | N/A | N/A | N/A | N/A | N/A | N/A | N/A | N/A | N/A | N/A | N/A | N/A | N/A | N/A |
| PA2061 | N/A | N/A | N/A | N/A | N/A | N/A | N/A | N/A | N/A | N/A | N/A | N/A | N/A | N/A | N/A | N/A | N/A | N/A |
| PA2062 | PA2062 | PA2062 | PA2062 | PA2062 | N/A | N/A | N/A | N/A | N/A | N/A | N/A | N/A | N/A | N/A | PA2062 | N/A | N/A | N/A |
| PA2063 | N/A | N/A | N/A | PA2063 | N/A | N/A | N/A | PA2063 | N/A | N/A | PA2063 | N/A | N/A | PA2063 | N/A | N/A | N/A | N/A |
| PA2066 | N/A | N/A | N/A | N/A | N/A | N/A | N/A | N/A | N/A | N/A | PA2066 | PA2066 | PA2066 | N/A | N/A | N/A | N/A | N/A |
| PA2067 | N/A | N/A | N/A | N/A | N/A | N/A | N/A | N/A | N/A | N/A | N/A | PA2067 | PA2067 | N/A | N/A | N/A | N/A | N/A |
| N/A | N/A | N/A | N/A | N/A | PA2068 | N/A | N/A | N/A | N/A | N/A | N/A | N/A | N/A | N/A | N/A | N/A | N/A | N/A |
| PA2069 | N/A | N/A | N/A | N/A | PA2069 | PA2069 | N/A | N/A | PA2069 | PA2069 | PA2069 | PA2069 | PA2069 | N/A | N/A | PA2069 | N/A | N/A |
| N/A | N/A | N/A | N/A | N/A | PA2070 | N/A | N/A | N/A | N/A | N/A | N/A | N/A | N/A | N/A | N/A | N/A | N/A | N/A |
| PA2071 | PA2071 | PA2071 | PA2071 | PA2071 | PA2071 | PA2071 | PA2071 | PA2071 | PA2071 | PA2071 | PA2071 | PA2071 | PA2071 | PA2071 | PA2071 | PA2071 | PA2071 | N/A |
| PA2072 | N/A | N/A | N/A | N/A | N/A | N/A | N/A | N/A | N/A | N/A | N/A | N/A | PA2072 | PA2072 | N/A | N/A | N/A | N/A |
| N/A | N/A | N/A | N/A | PA2076 | N/A | N/A | N/A | N/A | N/A | N/A | N/A | N/A | N/A | N/A | N/A | N/A | N/A | N/A |
| N/A | N/A | N/A | N/A | N/A | PA2079 | N/A | N/A | N/A | N/A | N/A | N/A | N/A | N/A | N/A | N/A | N/A | N/A | N/A |
| PA2080 | PA2080 | PA2080 | PA2080 | PA2080 | PA2080 | PA2080 | N/A | N/A | PA2080 | PA2080 | PA2080 | PA2080 | PA2080 | N/A | N/A | N/A | N/A | N/A |
| PA2081 | PA2081 | N/A | N/A | PA2081 | PA2081 | PA2081 | N/A | N/A | PA2081 | PA2081 | PA2081 | PA2081 | PA2081 | N/A | N/A | N/A | N/A | N/A |
| PA2082 | N/A | N/A | N/A | PA2082 | N/A | N/A | N/A | N/A | PA2082 | PA2082 | PA2082 | PA2082 | PA2082 | N/A | N/A | N/A | N/A | N/A |
| PA2100 | N/A | N/A | N/A | PA2100 | N/A | N/A | N/A | PA2100 | PA2100 | N/A | N/A | N/A | PA2100 | N/A | N/A | N/A | N/A | N/A |
| N/A | N/A | N/A | N/A | N/A | N/A | N/A | N/A | N/A | N/A | PA2107 | N/A | N/A | N/A | N/A | N/A | N/A | N/A | N/A |
| N/A | N/A | N/A | N/A | PA2109 | N/A | N/A | N/A | N/A | N/A | N/A | N/A | N/A | N/A | N/A | PA2109 | N/A | N/A | N/A |
| PA2110 | N/A | N/A | N/A | PA2110 | N/A | N/A | N/A | N/A | N/A | N/A | PA2110 | PA2110 | PA2110 | PA2110 | PA2110 | N/A | N/A | N/A |
| PA2111 | N/A | N/A | N/A | PA2111 | PA2111 | PA2111 | N/A | N/A | PA2111 | PA2111 | PA2111 | PA2111 | PA2111 | N/A | PA2111 | N/A | N/A | N/A |
| PA2112 | N/A | N/A | N/A | PA2112 | PA2112 | N/A | N/A | N/A | PA2112 | PA2112 | PA2112 | PA2112 | PA2112 | N/A | PA2112 | N/A | PA2112 | N/A |
| PA2113 | N/A | N/A | N/A | PA2113 | N/A | PA2113 | N/A | N/A | N/A | PA2113 | PA2113 | PA2113 | PA2113 | N/A | PA2113 | PA2113 | N/A | N/A |
| PA2114 | N/A | N/A | N/A | PA2114 | N/A | PA2114 | N/A | N/A | N/A | PA2114 | PA2114 | PA2114 | PA2114 | PA2114 | N/A | N/A | PA2114 | N/A |
| PA2115 | N/A | N/A | N/A | PA2115 | PA2115 | N/A | N/A | N/A | N/A | N/A | PA2115 | N/A | N/A | N/A | N/A | N/A | N/A | N/A |
| PA2116 | N/A | N/A | N/A | PA2116 | PA2116 | PA2116 | N/A | N/A | PA2116 | PA2116 | PA2116 | PA2116 | PA2116 | PA2116 | N/A | N/A | N/A | N/A |
| PA2117 | N/A | N/A | N/A | N/A | PA2117 | N/A | N/A | N/A | N/A | N/A | N/A | N/A | N/A | N/A | N/A | N/A | N/A | N/A |
| N/A | N/A | N/A | N/A | N/A | PA2118 | N/A | N/A | N/A | N/A | N/A | N/A | N/A | N/A | N/A | N/A | N/A | N/A | N/A |
| PA2119 | N/A | N/A | N/A | PA2119 | PA2119 | PA2119 | N/A | N/A | N/A | PA2119 | PA2119 | PA2119 | PA2119 | PA2119 | N/A | N/A | N/A | N/A |
| PA2120 | N/A | N/A | N/A | N/A | N/A | N/A | N/A | N/A | N/A | N/A | N/A | N/A | N/A | N/A | N/A | N/A | N/A | N/A |
| PA2121 | N/A | N/A | N/A | N/A | N/A | N/A | N/A | N/A | N/A | N/A | N/A | N/A | N/A | N/A | N/A | N/A | N/A | N/A |
| PA2122 | N/A | N/A | N/A | N/A | N/A | N/A | N/A | N/A | N/A | N/A | N/A | N/A | N/A | N/A | N/A | N/A | N/A | N/A |
| PA2126 | N/A | N/A | N/A | PA2126 | N/A | N/A | N/A | N/A | N/A | N/A | N/A | N/A | N/A | N/A | N/A | N/A | PA2126 | N/A |
| PA2127 | N/A | N/A | N/A | N/A | N/A | N/A | N/A | N/A | N/A | N/A | N/A | N/A | N/A | N/A | N/A | N/A | N/A | N/A |
| N/A | N/A | N/A | N/A | N/A | PA2133 | N/A | N/A | N/A | N/A | N/A | N/A | N/A | N/A | N/A | N/A | N/A | N/A | N/A |
| N/A | N/A | N/A | N/A | N/A | PA2134 | N/A | N/A | N/A | N/A | PA2134 | N/A | N/A | N/A | N/A | PA2134 | N/A | N/A | N/A |
| N/A | N/A | N/A | N/A | N/A | N/A | PA2142 | N/A | N/A | N/A | N/A | N/A | N/A | N/A | N/A | N/A | N/A | N/A | N/A |
| PA2144 | N/A | N/A | N/A | N/A | N/A | PA2144 | N/A | N/A | N/A | N/A | N/A | N/A | PA2144 | N/A | N/A | N/A | N/A | N/A |
| N/A | N/A | N/A | N/A | N/A | PA2146 | N/A | N/A | N/A | N/A | N/A | N/A | N/A | N/A | N/A | N/A | N/A | N/A | N/A |
| PA2147 | N/A | N/A | N/A | N/A | PA2147 | PA2147 | N/A | N/A | N/A | PA2147 | N/A | N/A | PA2147 | N/A | N/A | N/A | N/A | N/A |
| N/A | N/A | N/A | N/A | N/A | N/A | N/A | N/A | N/A | N/A | N/A | N/A | N/A | PA2151 | N/A | N/A | N/A | N/A | N/A |
| PA2152 | N/A | N/A | N/A | N/A | N/A | N/A | N/A | N/A | N/A | N/A | N/A | N/A | PA2152 | N/A | N/A | N/A | N/A | N/A |
| PA2153 | N/A | N/A | N/A | N/A | N/A | N/A | N/A | N/A | N/A | N/A | N/A | N/A | N/A | N/A | N/A | N/A | N/A | N/A |
| N/A | N/A | N/A | N/A | N/A | N/A | PA2159 | N/A | N/A | N/A | N/A | N/A | N/A | N/A | N/A | N/A | N/A | N/A | N/A |
| PA2160 | N/A | N/A | N/A | N/A | N/A | N/A | N/A | N/A | N/A | N/A | N/A | PA2160 | PA2160 | N/A | N/A | N/A | N/A | N/A |
| PA2161 | N/A | N/A | N/A | N/A | N/A | N/A | N/A | N/A | N/A | N/A | N/A | PA2161 | PA2161 | N/A | N/A | N/A | N/A | N/A |
| PA2162 | N/A | N/A | N/A | N/A | N/A | N/A | N/A | N/A | N/A | N/A | N/A | N/A | PA2162 | N/A | N/A | N/A | N/A | N/A |
| N/A | N/A | N/A | N/A | N/A | N/A | N/A | N/A | N/A | N/A | N/A | N/A | N/A | PA2164 | N/A | N/A | N/A | N/A | N/A |
| PA2166 | N/A | N/A | N/A | N/A | N/A | PA2166 | N/A | N/A | N/A | N/A | N/A | PA2166 | PA2166 | N/A | N/A | N/A | N/A | N/A |
| PA2169 | N/A | N/A | N/A | N/A | N/A | N/A | N/A | N/A | N/A | PA2169 | N/A | N/A | PA2169 | N/A | N/A | N/A | N/A | N/A |
| N/A | N/A | N/A | N/A | N/A | PA2170 | N/A | N/A | N/A | N/A | N/A | N/A | N/A | N/A | N/A | N/A | N/A | N/A | N/A |
| PA2171 | N/A | N/A | N/A | N/A | PA2171 | PA2171 | N/A | N/A | N/A | PA2171 | N/A | PA2171 | PA2171 | N/A | N/A | N/A | N/A | N/A |
| N/A | N/A | N/A | N/A | N/A | N/A | PA2172 | N/A | N/A | N/A | N/A | N/A | N/A | N/A | N/A | N/A | N/A | N/A | N/A |
| PA2173 | N/A | N/A | N/A | N/A | N/A | N/A | N/A | N/A | N/A | PA2173 | N/A | N/A | N/A | N/A | N/A | N/A | N/A | N/A |
| N/A | N/A | N/A | N/A | N/A | N/A | N/A | N/A | N/A | N/A | N/A | N/A | N/A | PA2174 | N/A | N/A | N/A | N/A | N/A |
| PA2175 | N/A | N/A | N/A | N/A | N/A | N/A | N/A | N/A | N/A | N/A | N/A | N/A | N/A | N/A | N/A | N/A | N/A | N/A |
| PA2176 | N/A | N/A | N/A | N/A | N/A | PA2176 | N/A | N/A | N/A | N/A | N/A | N/A | PA2176 | N/A | N/A | N/A | N/A | N/A |
| PA2177 | N/A | N/A | N/A | N/A | N/A | N/A | N/A | N/A | N/A | N/A | N/A | N/A | PA2177 | N/A | N/A | N/A | N/A | N/A |
| N/A | N/A | N/A | N/A | N/A | N/A | PA2178 | N/A | N/A | N/A | N/A | N/A | N/A | N/A | N/A | N/A | N/A | N/A | N/A |
| N/A | N/A | N/A | N/A | N/A | PA2183 | N/A | N/A | N/A | N/A | N/A | N/A | N/A | N/A | N/A | N/A | N/A | N/A | N/A |
| PA2184 | N/A | N/A | N/A | N/A | N/A | PA2184 | N/A | N/A | PA2184 | PA2184 | PA2184 | PA2184 | PA2184 | N/A | N/A | N/A | N/A | N/A |
| PA2190 | N/A | N/A | N/A | N/A | N/A | N/A | N/A | N/A | N/A | PA2190 | N/A | PA2190 | PA2190 | N/A | N/A | N/A | N/A | N/A |
| PA2193 | N/A | N/A | N/A | N/A | PA2193 | N/A | N/A | N/A | N/A | N/A | PA2193 | PA2193 | N/A | N/A | N/A | N/A | N/A | N/A |
| PA2194 | N/A | N/A | N/A | PA2194 | PA2194 | PA2194 | N/A | PA2194 | N/A | PA2194 | PA2194 | PA2194 | PA2194 | PA2194 | N/A | N/A | PA2194 | N/A |
| PA2195 | N/A | N/A | N/A | PA2195 | PA2195 | N/A | N/A | PA2195 | PA2195 | PA2195 | PA2195 | PA2195 | PA2195 | PA2195 | N/A | N/A | PA2195 | N/A |
| PA2196 | N/A | PA2196 | N/A | PA2196 | PA2196 | PA2196 | N/A | N/A | PA2196 | PA2196 | PA2196 | PA2196 | N/A | N/A | N/A | N/A | N/A | N/A |
| PA2197 | PA2197 | PA2197 | PA2197 | PA2197 | PA2197 | PA2197 | PA2197 | PA2197 | PA2197 | PA2197 | PA2197 | PA2197 | PA2197 | N/A | N/A | N/A | N/A | N/A |
| PA2198 | N/A | PA2198 | N/A | PA2198 | PA2198 | PA2198 | N/A | N/A | PA2198 | PA2198 | PA2198 | PA2198 | PA2198 | N/A | N/A | N/A | N/A | N/A |
| PA2199 | PA2199 | PA2199 | PA2199 | PA2199 | PA2199 | PA2199 | N/A | N/A | PA2199 | PA2199 | PA2199 | PA2199 | PA2199 | N/A | N/A | N/A | N/A | N/A |
| PA2200 | N/A | N/A | N/A | N/A | N/A | N/A | N/A | N/A | N/A | N/A | N/A | N/A | N/A | N/A | N/A | N/A | N/A | N/A |
| PA2201 | N/A | N/A | N/A | PA2201 | N/A | N/A | N/A | N/A | PA2201 | N/A | PA2201 | PA2201 | N/A | N/A | N/A | N/A | N/A | N/A |
| PA2203 | N/A | N/A | N/A | N/A | PA2203 | N/A | N/A | N/A | N/A | N/A | N/A | N/A | N/A | PA2203 | N/A | N/A | PA2203 | N/A |
| PA2204 | PA2204 | PA2204 | PA2204 | PA2204 | PA2204 | PA2204 | PA2204 | N/A | PA2204 | PA2204 | PA2204 | PA2204 | PA2204 | PA2204 | PA2204 | PA2204 | PA2204 | PA2204 |
| PA2205 | N/A | N/A | N/A | N/A | N/A | N/A | N/A | N/A | N/A | N/A | N/A | N/A | N/A | N/A | N/A | N/A | N/A | N/A |
| N/A | N/A | N/A | N/A | PA2206 | N/A | N/A | N/A | N/A | N/A | N/A | N/A | N/A | N/A | N/A | N/A | N/A | N/A | N/A |
| PA2218 | N/A | N/A | N/A | N/A | N/A | N/A | N/A | N/A | N/A | N/A | N/A | N/A | N/A | N/A | N/A | N/A | N/A | N/A |
| PA2220 | N/A | N/A | N/A | PA2220 | N/A | N/A | N/A | N/A | N/A | N/A | N/A | N/A | N/A | N/A | N/A | N/A | N/A | N/A |

| | | | | | | | | | | | | | | | | | | |
|---|---|---|---|---|---|---|---|---|---|---|---|---|---|---|---|---|---|---|
| PA2222 | N/A | N/A | N/A | PA2222 | PA2222 | PA2222 | N/A | N/A | PA2222 | PA2222 | PA2222 | PA2222 | PA2222 | N/A | N/A | N/A | N/A | N/A |
| PA2223 | N/A | N/A | N/A | N/A | PA2223 | N/A | N/A | N/A | N/A | N/A | PA2223 | PA2223 | PA2223 | N/A | N/A | N/A | N/A | N/A |
| PA2224 | N/A | N/A | N/A | N/A | N/A | N/A | N/A | N/A | N/A | N/A | N/A | N/A | N/A | N/A | N/A | N/A | N/A | N/A |
| PA2225 | N/A | N/A | N/A | N/A | N/A | N/A | N/A | N/A | N/A | N/A | N/A | N/A | N/A | N/A | N/A | N/A | N/A | N/A |
| N/A | N/A | N/A | N/A | N/A | N/A | N/A | N/A | N/A | N/A | N/A | PA2226 | N/A | N/A | N/A | N/A | N/A | N/A | N/A |
| N/A | N/A | N/A | N/A | N/A | PA2228 | N/A | N/A | N/A | N/A | N/A | N/A | N/A | N/A | N/A | N/A | N/A | N/A | N/A |
| PA2229 | N/A | N/A | N/A | PA2229 | N/A | N/A | N/A | N/A | PA2229 | PA2229 | PA2229 | PA2229 | N/A | N/A | N/A | N/A | N/A | N/A |
| N/A | N/A | N/A | N/A | N/A | N/A | N/A | N/A | N/A | N/A | N/A | N/A | N/A | N/A | PA2230 | N/A | N/A | N/A | N/A |
| PA2231 | N/A | N/A | N/A | PA2231 | PA2231 | PA2231 | N/A | N/A | N/A | N/A | PA2231 | PA2231 | PA2231 | PA2231 | N/A | N/A | PA2231 | N/A |
| PA2232 | PA2232 | N/A | PA2232 | PA2232 | N/A | PA2232 | N/A | PA2232 | PA2232 | PA2232 | PA2232 | PA2232 | PA2232 | PA2232 | PA2232 | N/A | PA2232 | PA2232 |
| PA2233 | N/A | N/A | N/A | PA2233 | N/A | N/A | N/A | N/A | N/A | N/A | PA2233 | N/A | N/A | PA2233 | N/A | N/A | PA2233 | N/A |
| PA2234 | PA2234 | PA2234 | PA2234 | PA2234 | PA2234 | PA2234 | PA2234 | PA2234 | PA2234 | PA2234 | PA2234 | PA2234 | PA2234 | PA2234 | PA2234 | N/A | PA2234 | N/A |
| PA2235 | PA2235 | PA2235 | PA2235 | PA2235 | N/A | PA2235 | N/A | PA2235 | PA2235 | PA2235 | PA2235 | PA2235 | PA2235 | PA2235 | PA2235 | N/A | PA2235 | PA2235 |
| PA2236 | N/A | N/A | N/A | PA2236 | N/A | N/A | PA2236 | N/A | N/A | N/A | N/A | N/A | PA2236 | PA2236 | N/A | N/A | PA2236 | N/A |
| PA2237 | N/A | N/A | N/A | PA2237 | N/A | N/A | N/A | N/A | N/A | N/A | N/A | N/A | PA2237 | PA2237 | N/A | N/A | N/A | N/A |
| PA2238 | N/A | N/A | N/A | PA2238 | N/A | N/A | N/A | N/A | N/A | N/A | N/A | N/A | PA2238 | PA2238 | N/A | N/A | N/A | N/A |
| PA2239 | N/A | N/A | N/A | PA2239 | N/A | N/A | N/A | N/A | N/A | N/A | N/A | N/A | N/A | PA2239 | N/A | N/A | N/A | N/A |
| PA2240 | N/A | N/A | N/A | PA2240 | N/A | N/A | N/A | N/A | N/A | N/A | N/A | PA2240 | N/A | PA2240 | N/A | N/A | N/A | N/A |
| PA2241 | N/A | N/A | N/A | PA2241 | N/A | N/A | N/A | N/A | N/A | N/A | N/A | N/A | N/A | PA2241 | N/A | N/A | N/A | N/A |
| PA2242 | N/A | N/A | N/A | N/A | N/A | PA2242 | N/A | PA2242 | PA2242 | PA2242 | PA2242 | PA2242 | PA2242 | PA2242 | N/A | N/A | PA2242 | N/A |
| PA2246 | N/A | N/A | N/A | PA2246 | PA2246 | PA2246 | N/A | N/A | N/A | N/A | N/A | PA2246 | PA2246 | PA2246 | N/A | N/A | N/A | N/A |
| PA2247 | N/A | N/A | N/A | PA2247 | N/A | PA2247 | N/A | PA2247 | PA2247 | PA2247 | PA2247 | PA2247 | PA2247 | PA2247 | PA2247 | PA2247 | PA2247 | N/A |
| PA2248 | N/A | N/A | N/A | PA2248 | N/A | PA2248 | PA2248 | PA2248 | PA2248 | PA2248 | PA2248 | PA2248 | PA2248 | PA2248 | PA2248 | PA2248 | PA2248 | N/A |
| PA2249 | PA2249 | N/A | N/A | PA2249 | N/A | PA2249 | N/A | PA2249 | PA2249 | PA2249 | PA2249 | PA2249 | PA2249 | PA2249 | PA2249 | N/A | N/A | N/A |
| PA2250 | PA2250 | N/A | N/A | PA2250 | N/A | PA2250 | N/A | N/A | PA2250 | PA2250 | PA2250 | PA2250 | PA2250 | PA2250 | PA2250 | PA2250 | PA2250 | N/A |
| PA2251 | N/A | N/A | N/A | N/A | N/A | N/A | N/A | N/A | N/A | N/A | N/A | N/A | N/A | N/A | N/A | N/A | N/A | N/A |
| PA2252 | PA2252 | PA2252 | PA2252 | PA2252 | PA2252 | PA2252 | N/A | PA2252 | PA2252 | N/A | PA2252 | PA2252 | PA2252 | PA2252 | PA2252 | N/A | PA2252 | PA2252 |
| PA2253 | PA2253 | PA2253 | PA2253 | PA2253 | N/A | N/A | N/A | N/A | PA2253 | N/A | PA2253 | PA2253 | PA2253 | N/A | N/A | N/A | N/A | N/A |
| PA2259 | N/A | N/A | N/A | N/A | N/A | N/A | N/A | N/A | N/A | N/A | N/A | N/A | N/A | N/A | N/A | N/A | N/A | N/A |
| PA2260 | N/A | N/A | N/A | N/A | N/A | N/A | N/A | N/A | N/A | N/A | N/A | N/A | N/A | N/A | N/A | N/A | N/A | N/A |
| PA2263 | N/A | N/A | N/A | N/A | N/A | N/A | N/A | N/A | N/A | N/A | N/A | N/A | PA2263 | N/A | N/A | N/A | N/A | N/A |
| PA2264 | PA2264 | PA2264 | PA2264 | PA2264 | N/A | N/A | N/A | N/A | PA2264 | PA2264 | N/A | N/A | N/A | PA2264 | PA2264 | N/A | PA2264 | N/A |
| PA2265 | PA2265 | PA2265 | PA2265 | PA2265 | N/A | N/A | N/A | N/A | PA2265 | PA2265 | N/A | PA2265 | N/A | PA2265 | N/A | N/A | PA2265 | N/A |
| PA2266 | PA2266 | PA2266 | PA2266 | PA2266 | N/A | N/A | PA2266 | PA2266 | PA2266 | PA2266 | N/A | N/A | PA2266 | PA2266 | PA2266 | N/A | PA2266 | N/A |
| PA2267 | N/A | N/A | N/A | N/A | N/A | N/A | N/A | N/A | N/A | N/A | N/A | N/A | N/A | N/A | N/A | N/A | N/A | N/A |
| PA2269 | N/A | N/A | N/A | N/A | N/A | N/A | N/A | N/A | N/A | N/A | N/A | N/A | N/A | PA2269 | N/A | N/A | N/A | N/A |
| PA2270 | N/A | N/A | N/A | N/A | N/A | N/A | N/A | N/A | N/A | N/A | PA2270 | PA2270 | PA2270 | N/A | N/A | N/A | N/A | N/A |
| PA2271 | N/A | N/A | N/A | PA2271 | N/A | N/A | N/A | N/A | N/A | N/A | N/A | N/A | N/A | N/A | N/A | N/A | N/A | N/A |
| PA2273 | N/A | N/A | N/A | N/A | N/A | N/A | N/A | N/A | N/A | N/A | PA2273 | N/A | N/A | N/A | N/A | N/A | N/A | N/A |
| PA2279 | N/A | N/A | N/A | N/A | N/A | N/A | N/A | N/A | N/A | N/A | PA2279 | N/A | N/A | N/A | N/A | N/A | N/A | N/A |
| N/A | N/A | N/A | N/A | PA2280 | N/A | N/A | N/A | N/A | N/A | N/A | N/A | N/A | N/A | N/A | N/A | N/A | N/A | N/A |
| PA2281 | N/A | N/A | N/A | PA2281 | N/A | N/A | N/A | N/A | N/A | N/A | N/A | N/A | N/A | N/A | N/A | N/A | N/A | N/A |
| N/A | N/A | N/A | N/A | PA2282 | N/A | N/A | N/A | N/A | N/A | PA2282 | N/A | N/A | N/A | N/A | PA2282 | N/A | N/A | N/A |
| PA2285 | PA2285 | PA2285 | PA2285 | PA2285 | N/A | N/A | PA2285 | N/A | PA2285 | PA2285 | N/A | N/A | N/A | N/A | PA2285 | N/A | N/A | N/A |
| PA2289 | N/A | N/A | N/A | PA2289 | PA2289 | N/A | N/A | PA2289 | PA2289 | N/A | N/A | N/A | N/A | N/A | N/A | N/A | N/A | N/A |
| PA2290 | PA2290 | PA2290 | PA2290 | PA2290 | PA2290 | N/A | PA2290 | PA2290 | PA2290 | PA2290 | PA2290 | PA2290 | PA2290 | PA2290 | PA2290 | N/A | PA2290 | PA2290 |
| PA2291 | PA2291 | PA2291 | PA2291 | PA2291 | N/A | PA2291 | PA2291 | PA2291 | PA2291 | PA2291 | PA2291 | PA2291 | PA2291 | PA2291 | PA2291 | PA2291 | N/A | N/A |
| PA2299 | N/A | N/A | N/A | N/A | PA2299 | N/A | N/A | N/A | N/A | N/A | N/A | N/A | N/A | N/A | N/A | N/A | N/A | N/A |
| PA2300 | N/A | N/A | N/A | N/A | N/A | PA2300 | N/A | N/A | PA2300 | PA2300 | PA2300 | PA2300 | PA2300 | N/A | N/A | PA2300 | N/A | N/A |
| PA2301 | N/A | N/A | N/A | PA2301 | PA2301 | N/A | N/A | N/A | PA2301 | PA2301 | PA2301 | PA2301 | PA2301 | N/A | N/A | N/A | N/A | N/A |
| PA2302 | N/A | N/A | N/A | PA2302 | N/A | PA2302 | N/A | N/A | PA2302 | PA2302 | PA2302 | PA2302 | PA2302 | N/A | N/A | N/A | N/A | N/A |
| PA2303 | N/A | N/A | N/A | N/A | PA2303 | N/A | N/A | N/A | N/A | N/A | N/A | PA2303 | PA2303 | N/A | N/A | N/A | N/A | N/A |
| PA2304 | N/A | N/A | N/A | PA2304 | N/A | N/A | N/A | N/A | N/A | N/A | N/A | PA2304 | PA2304 | N/A | N/A | N/A | N/A | N/A |
| PA2305 | N/A | N/A | N/A | PA2305 | PA2305 | N/A | N/A | N/A | PA2305 | PA2305 | PA2305 | PA2305 | PA2305 | N/A | N/A | N/A | N/A | N/A |
| PA2306 | N/A | N/A | N/A | N/A | N/A | PA2306 | N/A | PA2306 | PA2306 | PA2306 | PA2306 | PA2306 | PA2306 | PA2306 | N/A | N/A | N/A | N/A |
| PA2309 | N/A | N/A | N/A | N/A | N/A | N/A | N/A | N/A | N/A | N/A | N/A | N/A | N/A | N/A | N/A | N/A | N/A | N/A |
| PA2316 | N/A | N/A | N/A | N/A | N/A | N/A | N/A | N/A | N/A | N/A | N/A | N/A | N/A | N/A | N/A | N/A | N/A | N/A |
| PA2317 | N/A | N/A | N/A | N/A | N/A | N/A | N/A | N/A | N/A | N/A | N/A | N/A | N/A | N/A | N/A | N/A | N/A | N/A |
| PA2318 | N/A | N/A | N/A | PA2318 | N/A | N/A | N/A | N/A | PA2318 | PA2318 | PA2318 | PA2318 | PA2318 | N/A | N/A | N/A | N/A | N/A |
| PA2320 | N/A | N/A | N/A | PA2320 | N/A | N/A | PA2320 | N/A | PA2320 | N/A | PA2320 | PA2320 | N/A | N/A | N/A | N/A | N/A | N/A |
| PA2321 | N/A | N/A | N/A | PA2321 | N/A | N/A | N/A | N/A | PA2321 | PA2321 | PA2321 | PA2321 | PA2321 | N/A | N/A | N/A | N/A | N/A |
| PA2322 | N/A | N/A | N/A | N/A | PA2322 | N/A | N/A | N/A | PA2322 | N/A | N/A | PA2322 | N/A | PA2322 | N/A | N/A | PA2322 | N/A |
| PA2323 | PA2323 | PA2323 | PA2323 | PA2323 | PA2323 | PA2323 | N/A | PA2323 | PA2323 | PA2323 | PA2323 | PA2323 | PA2323 | PA2323 | N/A | N/A | N/A | N/A |
| PA2327 | N/A | N/A | N/A | N/A | PA2327 | N/A | N/A | N/A | PA2327 | N/A | N/A | PA2327 | PA2327 | PA2327 | N/A | N/A | N/A | N/A |
| PA2328 | N/A | N/A | N/A | N/A | N/A | N/A | N/A | N/A | N/A | N/A | N/A | PA2328 | PA2328 | N/A | N/A | N/A | N/A | N/A |
| PA2329 | N/A | N/A | N/A | PA2329 | PA2329 | N/A | N/A | N/A | PA2329 | PA2329 | PA2329 | PA2329 | PA2329 | PA2329 | N/A | N/A | PA2329 | N/A |
| PA2330 | N/A | N/A | N/A | PA2330 | N/A | PA2330 | N/A | N/A | PA2330 | PA2330 | PA2330 | PA2330 | PA2330 | N/A | N/A | N/A | N/A | N/A |
| PA2331 | N/A | N/A | N/A | N/A | N/A | PA2331 | N/A | N/A | PA2331 | PA2331 | PA2331 | PA2331 | PA2331 | N/A | N/A | PA2331 | N/A | N/A |
| PA2332 | N/A | N/A | N/A | PA2332 | N/A | N/A | N/A | N/A | N/A | N/A | PA2332 | PA2332 | PA2332 | N/A | N/A | N/A | N/A | N/A |
| PA2344 | PA2344 | N/A | N/A | PA2344 | N/A | N/A | N/A | N/A | N/A | N/A | N/A | N/A | N/A | N/A | N/A | N/A | N/A | N/A |
| PA2345 | N/A | N/A | N/A | PA2345 | N/A | PA2345 | N/A | N/A | PA2345 | PA2345 | N/A | PA2345 | PA2345 | PA2345 | N/A | N/A | PA2345 | N/A |
| N/A | N/A | PA2349 | PA2349 | N/A | N/A | N/A | N/A | N/A | N/A | N/A | N/A | N/A | N/A | N/A | N/A | N/A | N/A | N/A |
| N/A | N/A | N/A | N/A | N/A | N/A | N/A | N/A | N/A | N/A | N/A | N/A | N/A | N/A | PA2350 | N/A | N/A | N/A | N/A |
| N/A | N/A | N/A | N/A | N/A | PA2351 | N/A | N/A | N/A | N/A | N/A | N/A | N/A | N/A | N/A | N/A | N/A | N/A | N/A |
| PA2352 | N/A | N/A | PA2352 | PA2352 | N/A | N/A | N/A | N/A | PA2352 | N/A | N/A | N/A | N/A | N/A | PA2352 | N/A | N/A | N/A |
| N/A | N/A | N/A | N/A | PA2353 | N/A | N/A | N/A | N/A | N/A | N/A | N/A | N/A | N/A | N/A | N/A | N/A | N/A | N/A |
| PA2360 | N/A | N/A | N/A | N/A | N/A | N/A | N/A | N/A | N/A | N/A | N/A | N/A | N/A | N/A | N/A | N/A | N/A | N/A |
| PA2361 | N/A | N/A | N/A | N/A | N/A | N/A | N/A | N/A | N/A | N/A | N/A | N/A | N/A | N/A | N/A | N/A | N/A | N/A |
| PA2363 | N/A | N/A | N/A | N/A | N/A | N/A | N/A | N/A | N/A | N/A | N/A | PA2363 | PA2363 | N/A | N/A | N/A | N/A | N/A |
| PA2364 | N/A | N/A | N/A | N/A | N/A | N/A | N/A | N/A | N/A | N/A | N/A | N/A | N/A | PA2364 | N/A | N/A | N/A | N/A |
| PA2365 | N/A | N/A | N/A | N/A | N/A | N/A | N/A | N/A | N/A | N/A | PA2365 | PA2365 | PA2365 | N/A | N/A | N/A | N/A | N/A |
| PA2366 | N/A | N/A | N/A | N/A | N/A | N/A | N/A | N/A | N/A | N/A | PA2366 | PA2366 | PA2366 | N/A | N/A | N/A | PA2366 | N/A |
| PA2367 | N/A | N/A | N/A | N/A | N/A | N/A | N/A | N/A | N/A | N/A | PA2367 | PA2367 | PA2367 | N/A | N/A | N/A | N/A | N/A |
| PA2369 | N/A | N/A | N/A | N/A | N/A | N/A | N/A | N/A | N/A | N/A | N/A | N/A | N/A | N/A | N/A | N/A | N/A | N/A |
| PA2371 | N/A | N/A | N/A | PA2371 | N/A | N/A | N/A | N/A | N/A | N/A | N/A | PA2371 | PA2371 | N/A | N/A | N/A | N/A | N/A |
| PA2373 | N/A | N/A | N/A | N/A | N/A | N/A | N/A | N/A | N/A | N/A | N/A | PA2373 | PA2373 | N/A | N/A | N/A | N/A | N/A |
| N/A | N/A | N/A | N/A | N/A | PA2377 | N/A | N/A | N/A | N/A | N/A | N/A | N/A | N/A | N/A | N/A | N/A | N/A | N/A |
| PA2378 | PA2378 | PA2378 | PA2378 | PA2378 | PA2378 | PA2378 | N/A | N/A | PA2378 | PA2378 | PA2378 | PA2378 | PA2378 | N/A | PA2378 | N/A | N/A | N/A |
| PA2379 | N/A | PA2379 | N/A | N/A | N/A | PA2379 | N/A | N/A | PA2379 | PA2379 | N/A | PA2379 | PA2379 | N/A | N/A | N/A | N/A | N/A |
| N/A | N/A | N/A | N/A | N/A | N/A | N/A | N/A | N/A | N/A | N/A | PA2380 | N/A | N/A | N/A | N/A | N/A | N/A | N/A |
| N/A | N/A | N/A | N/A | N/A | N/A | N/A | N/A | N/A | N/A | N/A | N/A | PA2381 | PA2381 | N/A | N/A | N/A | N/A | N/A |
| N/A | N/A | N/A | N/A | N/A | N/A | N/A | N/A | N/A | N/A | PA2382 | N/A | N/A | N/A | N/A | N/A | N/A | N/A | N/A |
| PA2384 | N/A | N/A | N/A | N/A | N/A | N/A | N/A | N/A | PA2384 | N/A | N/A | N/A | PA2384 | N/A | N/A | N/A | N/A | N/A |
| PA2385 | N/A | N/A | N/A | PA2385 | PA2385 | N/A | N/A | N/A | N/A | N/A | N/A | N/A | N/A | N/A | N/A | N/A | N/A | N/A |
| PA2386 | N/A | N/A | N/A | PA2386 | PA2386 | N/A | N/A | N/A | PA2386 | N/A | N/A | N/A | PA2386 | N/A | N/A | PA2386 | N/A | N/A |
| PA2388 | N/A | N/A | N/A | PA2388 | N/A | N/A | PA2388 | N/A | N/A | N/A | N/A | N/A | N/A | PA2388 | N/A | N/A | PA2388 | N/A |
| PA2389 | N/A | N/A | N/A | N/A | N/A | N/A | N/A | N/A | N/A | N/A | N/A | N/A | N/A | N/A | N/A | N/A | N/A | N/A |
| PA2390 | N/A | N/A | N/A | N/A | N/A | N/A | N/A | N/A | N/A | N/A | N/A | N/A | N/A | N/A | N/A | N/A | N/A | N/A |
| PA2391 | N/A | N/A | N/A | PA2391 | N/A | N/A | N/A | N/A | N/A | N/A | N/A | N/A | N/A | N/A | N/A | N/A | N/A | N/A |
| PA2392 | N/A | N/A | N/A | N/A | N/A | N/A | N/A | N/A | N/A | N/A | N/A | N/A | PA2392 | N/A | N/A | N/A | N/A | N/A |
| PA2393 | N/A | N/A | N/A | PA2393 | N/A | PA2393 | N/A | N/A | N/A | N/A | N/A | N/A | PA2393 | N/A | N/A | N/A | N/A | N/A |
| PA2394 | N/A | N/A | N/A | PA2394 | N/A | PA2394 | N/A | N/A | N/A | N/A | N/A | N/A | PA2394 | N/A | N/A | N/A | N/A | N/A |
| PA2395 | N/A | N/A | N/A | PA2395 | N/A | N/A | N/A | N/A | N/A | N/A | N/A | N/A | PA2395 | N/A | N/A | N/A | N/A | N/A |
| PA2396 | N/A | N/A | N/A | PA2396 | N/A | N/A | N/A | N/A | N/A | N/A | N/A | N/A | PA2396 | N/A | N/A | N/A | N/A | N/A |
| PA2397 | N/A | N/A | N/A | N/A | N/A | PA2397 | N/A | N/A | N/A | N/A | N/A | N/A | PA2397 | N/A | N/A | N/A | N/A | N/A |
| PA2398 | N/A | N/A | N/A | PA2398 | N/A | PA2398 | N/A | PA2398 | PA2398 | PA2398 | PA2398 | PA2398 | PA2398 | N/A | N/A | N/A | N/A | N/A |
| PA2399 | PA2399 | N/A | N/A | PA2399 | N/A | N/A | N/A | N/A | PA2399 | N/A | N/A | PA2399 | PA2399 | N/A | N/A | N/A | N/A | N/A |
| PA2400 | N/A | N/A | N/A | PA2400 | N/A | N/A | N/A | N/A | PA2400 | N/A | PA2400 | N/A | PA2400 | N/A | N/A | N/A | N/A | N/A |
| PA2402 | N/A | N/A | N/A | PA2402 | N/A | N/A | N/A | N/A | N/A | N/A | PA2402 | N/A | PA2402 | N/A | N/A | N/A | N/A | N/A |

| | | | | | | | | | | | | | | | | | | |
|---|---|---|---|---|---|---|---|---|---|---|---|---|---|---|---|---|---|---|
| PA2403 | N/A | N/A | N/A | N/A | N/A | N/A | N/A | N/A | N/A | N/A | N/A | N/A | N/A | N/A | N/A | N/A | N/A | N/A |
| PA2404 | N/A | N/A | N/A | PA2404 | N/A | N/A | N/A | N/A | PA2404 | N/A | PA2404 | N/A | N/A | N/A | N/A | N/A | N/A | N/A |
| PA2406 | N/A | N/A | N/A | N/A | PA2406 | N/A | N/A | N/A | PA2406 | N/A | N/A | N/A | N/A | PA2406 | N/A | N/A | N/A | N/A |
| PA2407 | N/A | N/A | N/A | PA2407 | PA2407 | PA2407 | N/A | N/A | PA2407 | N/A | PA2407 | PA2407 | PA2407 | N/A | N/A | N/A | N/A | N/A |
| PA2408 | N/A | N/A | N/A | N/A | N/A | N/A | N/A | N/A | N/A | N/A | N/A | PA2408 | PA2408 | N/A | N/A | N/A | N/A | N/A |
| PA2409 | N/A | N/A | N/A | N/A | PA2409 | N/A | N/A | N/A | N/A | N/A | N/A | N/A | N/A | N/A | N/A | N/A | N/A | N/A |
| PA2410 | N/A | N/A | N/A | PA2410 | PA2410 | N/A | N/A | N/A | PA2410 | N/A | PA2410 | N/A | N/A | N/A | N/A | N/A | N/A | N/A |
| PA2411 | N/A | N/A | N/A | PA2411 | N/A | N/A | N/A | N/A | N/A | N/A | N/A | N/A | PA2411 | N/A | N/A | N/A | N/A | N/A |
| PA2413 | N/A | N/A | N/A | PA2413 | N/A | N/A | N/A | N/A | PA2413 | N/A | N/A | N/A | PA2413 | N/A | N/A | N/A | N/A | N/A |
| PA2414 | N/A | N/A | N/A | N/A | N/A | N/A | N/A | N/A | N/A | N/A | N/A | N/A | PA2414 | PA2414 | N/A | N/A | N/A | N/A |
| PA2416 | N/A | N/A | N/A | N/A | N/A | N/A | N/A | N/A | N/A | N/A | N/A | N/A | N/A | N/A | N/A | N/A | N/A | N/A |
| PA2417 | N/A | N/A | N/A | N/A | N/A | N/A | N/A | PA2417 | N/A | PA2417 | N/A | N/A | PA2417 | N/A | N/A | N/A | N/A | N/A |
| PA2419 | N/A | N/A | N/A | N/A | N/A | PA2419 | N/A | N/A | PA2419 | PA2419 | N/A | PA2419 | PA2419 | N/A | N/A | N/A | N/A | N/A |
| PA2424 | N/A | N/A | N/A | PA2424 | N/A | N/A | N/A | N/A | N/A | N/A | N/A | PA2424 | PA2424 | N/A | N/A | N/A | N/A | N/A |
| N/A | N/A | N/A | N/A | N/A | N/A | N/A | N/A | N/A | N/A | N/A | N/A | N/A | PA2426 | N/A | N/A | N/A | N/A | N/A |
| PA2430 | N/A | N/A | N/A | N/A | N/A | N/A | N/A | N/A | N/A | N/A | N/A | N/A | N/A | PA2430 | N/A | N/A | PA2430 | N/A |
| N/A | N/A | N/A | N/A | N/A | PA2431 | N/A | N/A | N/A | N/A | N/A | N/A | N/A | N/A | N/A | N/A | N/A | N/A | N/A |
| N/A | N/A | N/A | N/A | N/A | PA2432 | N/A | N/A | N/A | N/A | N/A | N/A | N/A | N/A | N/A | N/A | N/A | N/A | N/A |
| N/A | N/A | N/A | N/A | N/A | PA2433 | PA2433 | N/A | N/A | PA2433 | N/A | PA2433 | PA2433 | PA2433 | N/A | N/A | N/A | N/A | N/A |
| PA2435 | N/A | N/A | N/A | N/A | N/A | N/A | N/A | N/A | PA2435 | N/A | N/A | N/A | N/A | PA2435 | N/A | N/A | N/A | N/A |
| PA2436 | N/A | N/A | N/A | PA2436 | N/A | N/A | N/A | N/A | PA2436 | N/A | PA2436 | PA2436 | PA2436 | N/A | N/A | N/A | N/A | PA2436 |
| PA2438 | N/A | N/A | N/A | N/A | N/A | N/A | N/A | N/A | N/A | N/A | N/A | N/A | N/A | N/A | N/A | N/A | N/A | N/A |
| PA2439 | N/A | N/A | N/A | N/A | N/A | N/A | N/A | N/A | N/A | N/A | PA2439 | PA2439 | N/A | N/A | N/A | N/A | N/A | N/A |
| N/A | N/A | N/A | N/A | N/A | PA2441 | N/A | N/A | N/A | N/A | N/A | N/A | N/A | N/A | N/A | N/A | N/A | N/A | N/A |
| PA2442 | N/A | PA2442 | PA2442 | PA2442 | PA2442 | PA2442 | PA2442 | N/A | PA2442 | PA2442 | PA2442 | PA2442 | PA2442 | N/A | PA2442 | PA2442 | N/A | N/A |
| PA2443 | N/A | PA2443 | N/A | PA2443 | PA2443 | PA2443 | N/A | N/A | PA2443 | PA2443 | PA2443 | PA2443 | PA2443 | N/A | PA2443 | N/A | N/A | N/A |
| PA2444 | PA2444 | PA2444 | PA2444 | PA2444 | PA2444 | PA2444 | PA2444 | PA2444 | PA2444 | PA2444 | PA2444 | PA2444 | PA2444 | N/A | PA2444 | N/A | N/A | N/A |
| PA2445 | PA2445 | PA2445 | PA2445 | PA2445 | PA2445 | PA2445 | N/A | N/A | PA2445 | PA2445 | PA2445 | PA2445 | PA2445 | PA2445 | PA2445 | N/A | N/A | N/A |
| PA2446 | N/A | N/A | N/A | PA2446 | PA2446 | PA2446 | N/A | N/A | PA2446 | PA2446 | PA2446 | PA2446 | PA2446 | N/A | PA2446 | N/A | N/A | N/A |
| N/A | N/A | N/A | N/A | N/A | PA2447 | N/A | N/A | N/A | N/A | N/A | N/A | N/A | N/A | N/A | N/A | N/A | N/A | N/A |
| PA2448 | N/A | N/A | N/A | N/A | PA2448 | N/A | N/A | N/A | N/A | N/A | N/A | N/A | PA2448 | N/A | N/A | N/A | N/A | N/A |
| PA2449 | N/A | N/A | N/A | PA2449 | N/A | N/A | N/A | PA2449 | PA2449 | PA2449 | N/A | PA2449 | N/A | N/A | N/A | N/A | N/A | N/A |
| N/A | N/A | N/A | N/A | PA2450 | N/A | N/A | PA2450 | N/A | N/A | N/A | N/A | N/A | N/A | N/A | PA2450 | N/A | N/A | N/A |
| N/A | N/A | N/A | N/A | N/A | PA2452 | N/A | N/A | N/A | N/A | N/A | N/A | N/A | N/A | N/A | N/A | PA2452 | N/A | N/A |
| PA2453 | PA2453 | PA2453 | PA2453 | PA2453 | PA2453 | PA2453 | PA2453 | N/A | PA2453 | PA2453 | PA2453 | PA2453 | PA2453 | PA2453 | PA2453 | PA2453 | N/A | N/A |
| PA2455 | N/A | N/A | N/A | N/A | N/A | N/A | N/A | N/A | PA2455 | N/A | PA2455 | N/A | N/A | N/A | N/A | N/A | N/A | N/A |
| N/A | N/A | N/A | N/A | N/A | N/A | N/A | N/A | N/A | PA2459 | N/A | PA2459 | PA2459 | PA2459 | N/A | N/A | N/A | N/A | N/A |
| PA2461 | N/A | N/A | N/A | PA2461 | PA2461 | N/A | N/A | N/A | N/A | N/A | N/A | N/A | N/A | N/A | N/A | N/A | N/A | N/A |
| PA2462 | PA2462 | N/A | N/A | PA2462 | N/A | N/A | N/A | PA2462 | PA2462 | PA2462 | PA2462 | N/A | N/A | PA2462 | PA2462 | PA2462 | PA2462 | PA2462 |
| PA2463 | N/A | N/A | N/A | PA2463 | PA2463 | N/A | N/A | PA2463 | PA2463 | PA2463 | PA2463 | PA2463 | PA2463 | N/A | PA2463 | N/A | N/A | N/A |
| PA2464 | PA2464 | PA2464 | N/A | PA2464 | PA2464 | N/A | N/A | N/A | PA2464 | PA2464 | PA2464 | PA2464 | PA2464 | N/A | PA2464 | N/A | N/A | N/A |
| PA2466 | N/A | N/A | N/A | N/A | N/A | N/A | N/A | N/A | N/A | N/A | N/A | N/A | PA2466 | N/A | N/A | N/A | N/A | N/A |
| PA2467 | N/A | N/A | N/A | N/A | N/A | N/A | N/A | N/A | N/A | N/A | N/A | N/A | N/A | N/A | N/A | N/A | N/A | N/A |
| PA2469 | N/A | N/A | N/A | N/A | N/A | N/A | N/A | N/A | N/A | N/A | N/A | N/A | N/A | N/A | N/A | N/A | N/A | N/A |
| N/A | N/A | N/A | N/A | N/A | N/A | PA2475 | N/A | N/A | N/A | N/A | N/A | N/A | PA2475 | N/A | N/A | N/A | N/A | N/A |
| PA2476 | N/A | N/A | N/A | PA2476 | N/A | N/A | N/A | N/A | PA2476 | PA2476 | PA2476 | PA2476 | PA2476 | N/A | N/A | N/A | N/A | N/A |
| PA2477 | N/A | N/A | N/A | N/A | PA2477 | N/A | N/A | N/A | N/A | N/A | N/A | N/A | N/A | N/A | N/A | N/A | N/A | N/A |
| PA2478 | N/A | N/A | N/A | N/A | N/A | N/A | N/A | N/A | N/A | N/A | N/A | N/A | N/A | N/A | N/A | N/A | N/A | N/A |
| PA2479 | N/A | N/A | N/A | N/A | N/A | N/A | N/A | N/A | N/A | N/A | N/A | N/A | N/A | N/A | N/A | N/A | N/A | N/A |
| PA2480 | N/A | N/A | N/A | N/A | N/A | N/A | N/A | N/A | N/A | PA2480 | N/A | N/A | N/A | PA2480 | N/A | N/A | N/A | N/A |
| PA2481 | N/A | PA2481 | N/A | PA2481 | N/A | N/A | N/A | N/A | N/A | N/A | PA2481 | N/A | N/A | N/A | N/A | N/A | N/A | N/A |
| PA2482 | PA2482 | N/A | N/A | PA2482 | PA2482 | N/A | N/A | N/A | PA2482 | N/A | PA2482 | PA2482 | N/A | N/A | PA2482 | N/A | N/A | N/A |
| PA2483 | PA2483 | PA2483 | PA2483 | PA2483 | PA2483 | PA2483 | PA2483 | PA2483 | PA2483 | PA2483 | PA2483 | PA2483 | PA2483 | N/A | N/A | N/A | N/A | N/A |
| N/A | N/A | N/A | N/A | N/A | PA2490 | N/A | N/A | N/A | N/A | PA2490 | N/A | N/A | N/A | N/A | N/A | N/A | N/A | N/A |
| PA2491 | PA2491 | PA2491 | PA2491 | PA2491 | PA2491 | PA2491 | N/A | N/A | PA2491 | PA2491 | PA2491 | PA2491 | PA2491 | N/A | N/A | N/A | N/A | N/A |
| PA2492 | N/A | N/A | N/A | N/A | N/A | N/A | N/A | N/A | N/A | N/A | N/A | N/A | N/A | N/A | N/A | N/A | N/A | N/A |
| PA2493 | N/A | N/A | N/A | N/A | N/A | N/A | N/A | N/A | N/A | N/A | N/A | N/A | N/A | PA2493 | N/A | N/A | PA2493 | PA2493 |
| PA2494 | N/A | PA2494 | PA2494 | PA2494 | N/A | PA2494 | N/A | N/A | PA2494 | N/A | PA2494 | PA2494 | PA2494 | PA2494 | N/A | N/A | PA2494 | PA2494 |
| PA2495 | N/A | N/A | N/A | N/A | N/A | N/A | N/A | N/A | N/A | N/A | N/A | N/A | N/A | PA2495 | N/A | N/A | N/A | N/A |
| N/A | N/A | N/A | N/A | PA2497 | N/A | N/A | N/A | N/A | N/A | N/A | N/A | N/A | N/A | N/A | N/A | N/A | N/A | N/A |
| PA2499 | N/A | N/A | N/A | N/A | N/A | N/A | N/A | N/A | N/A | N/A | N/A | N/A | N/A | N/A | N/A | N/A | N/A | N/A |
| N/A | N/A | N/A | N/A | N/A | N/A | PA2501 | N/A | N/A | N/A | N/A | PA2501 | N/A | N/A | N/A | N/A | N/A | N/A | N/A |
| N/A | N/A | N/A | N/A | N/A | N/A | PA3109 | N/A | N/A | N/A | N/A | N/A | N/A | N/A | N/A | N/A | N/A | N/A | N/A |
| N/A | N/A | N/A | N/A | N/A | N/A | PA3236 | N/A | N/A | N/A | PA3236 | N/A | PA3236 | PA3236 | N/A | N/A | N/A | N/A | N/A |
| PA2502 | PA2502 | N/A | N/A | PA2502 | N/A | N/A | N/A | PA2502 | PA2502 | N/A | N/A | PA2502 | PA2502 | N/A | N/A | N/A | N/A | N/A |
| PA2503 | N/A | N/A | N/A | PA2503 | PA2503 | N/A | PA2503 | PA2503 | N/A | N/A | N/A | N/A | PA2503 | N/A | N/A | N/A | N/A | N/A |
| PA2504 | N/A | N/A | N/A | PA2504 | PA2504 | PA2504 | N/A | N/A | PA2504 | PA2504 | PA2504 | PA2504 | PA2504 | N/A | N/A | N/A | N/A | N/A |
| PA2512 | N/A | N/A | N/A | N/A | N/A | N/A | N/A | N/A | N/A | N/A | N/A | N/A | N/A | N/A | N/A | N/A | N/A | N/A |
| PA2513 | N/A | N/A | N/A | N/A | PA2513 | PA2513 | N/A | N/A | N/A | N/A | N/A | PA2513 | PA2513 | N/A | N/A | N/A | N/A | N/A |
| PA2514 | N/A | N/A | N/A | N/A | PA2514 | N/A | N/A | N/A | N/A | N/A | N/A | N/A | N/A | N/A | N/A | N/A | N/A | N/A |
| PA2523 | N/A | N/A | N/A | N/A | N/A | N/A | N/A | N/A | N/A | N/A | N/A | N/A | N/A | N/A | N/A | N/A | N/A | N/A |
| PA2525 | N/A | PA2525 | N/A | PA2525 | N/A | PA2525 | N/A | PA2525 | N/A | PA2525 | N/A | N/A | N/A | N/A | PA2525 | N/A | N/A | N/A |
| PA2526 | N/A | N/A | N/A | N/A | N/A | N/A | N/A | N/A | PA2526 | N/A | N/A | N/A | N/A | PA2526 | N/A | N/A | N/A | N/A |
| PA2527 | N/A | N/A | N/A | PA2527 | N/A | N/A | N/A | N/A | N/A | PA2527 | N/A | N/A | N/A | PA2527 | N/A | N/A | PA2527 | N/A |
| PA2528 | N/A | N/A | N/A | PA2528 | N/A | PA2528 | N/A | PA2528 | PA2528 | PA2528 | N/A | PA2528 | PA2528 | PA2528 | N/A | N/A | PA2528 | PA2528 |
| PA2529 | PA2529 | PA2529 | N/A | PA2529 | PA2529 | N/A | N/A | N/A | N/A | N/A | PA2529 | PA2529 | PA2529 | N/A | N/A | N/A | N/A | N/A |
| PA2530 | PA2530 | N/A | PA2530 | PA2530 | PA2530 | PA2530 | N/A | N/A | N/A | PA2530 | PA2530 | PA2530 | PA2530 | N/A | N/A | N/A | N/A | N/A |
| PA2531 | N/A | N/A | N/A | N/A | PA2531 | N/A | N/A | N/A | N/A | N/A | N/A | N/A | N/A | N/A | N/A | N/A | N/A | N/A |
| PA2532 | PA2532 | PA2532 | PA2532 | PA2532 | PA2532 | PA2532 | PA2532 | N/A | PA2532 | PA2532 | PA2532 | PA2532 | PA2532 | N/A | PA2532 | N/A | PA2532 | N/A |
| PA2533 | N/A | N/A | N/A | PA2533 | N/A | N/A | N/A | N/A | N/A | N/A | PA2533 | PA2533 | N/A | PA2533 | N/A | N/A | PA2533 | N/A |
| PA2535 | PA2535 | PA2535 | N/A | PA2535 | N/A | N/A | N/A | N/A | N/A | N/A | N/A | N/A | N/A | N/A | N/A | PA2535 | N/A | N/A |
| PA2536 | N/A | N/A | N/A | PA2536 | N/A | N/A | N/A | N/A | N/A | PA2536 | PA2536 | PA2536 | PA2536 | PA2536 | N/A | N/A | PA2536 | N/A |
| PA2537 | PA2537 | PA2537 | PA2537 | PA2537 | N/A | N/A | PA2537 | N/A | PA2537 | PA2537 | PA2537 | PA2537 | PA2537 | PA2537 | PA2537 | N/A | PA2537 | PA2537 |
| PA2538 | N/A | N/A | N/A | N/A | N/A | N/A | N/A | N/A | N/A | N/A | PA2538 | N/A | N/A | PA2538 | N/A | N/A | PA2538 | N/A |
| PA2539 | N/A | N/A | N/A | PA2539 | N/A | N/A | N/A | N/A | N/A | N/A | N/A | N/A | N/A | PA2539 | N/A | N/A | PA2539 | N/A |
| PA2540 | PA2540 | PA2540 | PA2540 | PA2540 | N/A | N/A | N/A | PA2540 | PA2540 | PA2540 | PA2540 | PA2540 | PA2540 | PA2540 | N/A | N/A | PA2540 | PA2540 |
| PA2541 | N/A | N/A | N/A | PA2541 | N/A | N/A | N/A | N/A | PA2541 | PA2541 | PA2541 | N/A | N/A | PA2541 | N/A | N/A | PA2541 | PA2541 |
| PA2542 | N/A | N/A | N/A | N/A | N/A | N/A | N/A | N/A | N/A | N/A | N/A | N/A | N/A | N/A | N/A | N/A | N/A | N/A |
| PA2543 | N/A | N/A | N/A | PA2543 | N/A | N/A | N/A | PA2543 | N/A | PA2543 | N/A | PA2543 | PA2543 | N/A | PA2543 | N/A | N/A | N/A |
| PA2544 | N/A | N/A | N/A | N/A | PA2544 | N/A | N/A | N/A | N/A | N/A | N/A | PA2544 | PA2544 | PA2544 | N/A | N/A | N/A | N/A |
| PA2545 | PA2545 | PA2545 | PA2545 | PA2545 | PA2545 | PA2545 | PA2545 | PA2545 | PA2545 | PA2545 | PA2545 | PA2545 | PA2545 | N/A | PA2545 | N/A | N/A | N/A |
| PA2546 | N/A | N/A | N/A | N/A | PA2546 | N/A | N/A | N/A | N/A | N/A | N/A | PA2546 | PA2546 | N/A | N/A | N/A | N/A | N/A |
| N/A | N/A | N/A | N/A | N/A | N/A | N/A | N/A | N/A | N/A | PA2549 | N/A | N/A | N/A | N/A | N/A | N/A | N/A | N/A |
| N/A | N/A | N/A | N/A | PA2550 | N/A | N/A | N/A | N/A | N/A | PA2550 | N/A | N/A | N/A | N/A | N/A | N/A | N/A | N/A |
| PA2551 | PA2551 | N/A | PA2551 | PA2551 | PA2551 | PA2551 | N/A | PA2551 | N/A | N/A | PA2551 | N/A | N/A | N/A | N/A | N/A | N/A | N/A |
| PA2552 | N/A | N/A | N/A | PA2552 | PA2552 | PA2552 | N/A | N/A | PA2552 | PA2552 | PA2552 | PA2552 | PA2552 | PA2552 | N/A | N/A | N/A | N/A |
| PA2553 | N/A | PA2553 | N/A | PA2553 | PA2553 | PA2553 | N/A | N/A | PA2553 | PA2553 | PA2553 | PA2553 | PA2553 | PA2553 | N/A | PA2553 | N/A | N/A |
| PA2554 | N/A | N/A | N/A | PA2554 | PA2554 | PA2554 | N/A | N/A | PA2554 | PA2554 | PA2554 | PA2554 | PA2554 | PA2554 | N/A | N/A | PA2554 | PA2554 |
| PA2555 | N/A | N/A | N/A | PA2555 | N/A | PA2555 | N/A | N/A | PA2555 | PA2555 | PA2555 | PA2555 | PA2555 | N/A | N/A | N/A | N/A | N/A |
| PA2557 | N/A | N/A | N/A | PA2557 | N/A | PA2557 | N/A | N/A | PA2557 | PA2557 | PA2557 | PA2557 | PA2557 | N/A | N/A | N/A | N/A | N/A |
| PA2558 | N/A | N/A | N/A | N/A | N/A | N/A | N/A | N/A | PA2558 | N/A | N/A | N/A | N/A | PA2558 | N/A | N/A | N/A | N/A |
| PA2559 | N/A | PA2559 | N/A | PA2559 | N/A | PA2559 | N/A | N/A | PA2559 | PA2559 | PA2559 | PA2559 | PA2559 | N/A | PA2559 | N/A | N/A | N/A |
| PA2559.1 | N/A | N/A | N/A | N/A | N/A | N/A | N/A | N/A | N/A | N/A | N/A | N/A | N/A | N/A | N/A | N/A | N/A | N/A |
| PA2561 | PA2561 | PA2561 | PA2561 | PA2561 | PA2561 | PA2561 | PA2561 | PA2561 | PA2561 | PA2561 | PA2561 | PA2561 | PA2561 | PA2561 | PA2561 | N/A | PA2561 | PA2561 |
| PA2562 | N/A | N/A | N/A | N/A | PA2562 | PA2562 | N/A | N/A | N/A | PA2562 | N/A | N/A | PA2562 | N/A | N/A | N/A | N/A | N/A |
| PA2564 | N/A | N/A | N/A | N/A | N/A | N/A | N/A | N/A | N/A | N/A | N/A | N/A | PA2564 | N/A | N/A | N/A | N/A | N/A |
| PA2565 | N/A | N/A | N/A | N/A | N/A | N/A | N/A | N/A | N/A | N/A | N/A | PA2565 | PA2565 | N/A | N/A | N/A | N/A | N/A |
| PA2566 | N/A | N/A | N/A | N/A | N/A | N/A | N/A | N/A | N/A | N/A | N/A | N/A | PA2566 | N/A | N/A | N/A | N/A | N/A |
| PA2567 | N/A | N/A | N/A | PA2567 | N/A | N/A | N/A | N/A | N/A | PA2567 | N/A | N/A | N/A | N/A | N/A | N/A | N/A | N/A |

| | | | | | | | | | | | | | | | | | | |
|---|---|---|---|---|---|---|---|---|---|---|---|---|---|---|---|---|---|---|
| PA2568 | N/A | N/A | N/A | N/A | N/A | N/A | N/A | N/A | N/A | N/A | N/A | N/A | N/A | N/A | N/A | N/A | N/A | N/A |
| PA2569 | N/A | N/A | N/A | N/A | PA2569 | N/A | N/A | N/A | N/A | N/A | N/A | N/A | N/A | N/A | N/A | N/A | N/A | N/A |
| PA2570 | N/A | N/A | N/A | N/A | N/A | PA2570 | N/A | N/A | PA2570 | N/A | PA2570 | PA2570 | PA2570 | N/A | N/A | N/A | N/A | PA2570 |
| PA2571 | N/A | N/A | N/A | N/A | PA2571 | N/A | N/A | N/A | N/A | N/A | N/A | N/A | N/A | N/A | N/A | N/A | N/A | N/A |
| PA2572 | N/A | N/A | N/A | N/A | PA2572 | N/A | N/A | N/A | N/A | N/A | N/A | PA2572 | PA2572 | N/A | N/A | N/A | N/A | N/A |
| PA2573 | PA2573 | PA2573 | PA2573 | PA2573 | N/A | PA2573 | PA2573 | PA2573 | PA2573 | PA2573 | PA2573 | PA2573 | PA2573 | PA2573 | PA2573 | N/A | PA2573 | PA2573 |
| N/A | N/A | N/A | N/A | N/A | PA2574 | N/A | N/A | N/A | N/A | N/A | N/A | N/A | N/A | N/A | N/A | N/A | N/A | N/A |
| PA2575 | PA2575 | PA2575 | PA2575 | PA2575 | PA2575 | PA2575 | N/A | N/A | PA2575 | PA2575 | PA2575 | PA2575 | PA2575 | N/A | PA2575 | N/A | N/A | N/A |
| PA2577 | N/A | N/A | N/A | PA2577 | N/A | N/A | N/A | N/A | PA2577 | PA2577 | PA2577 | PA2577 | PA2577 | N/A | N/A | N/A | N/A | PA2577 |
| N/A | N/A | N/A | N/A | N/A | PA2578 | N/A | N/A | N/A | N/A | N/A | N/A | N/A | N/A | N/A | N/A | N/A | N/A | N/A |
| PA2579 | N/A | PA2579 | PA2579 | PA2579 | PA2579 | PA2579 | PA2579 | N/A | PA2579 | PA2579 | PA2579 | PA2579 | PA2579 | N/A | N/A | PA2579 | N/A | N/A |
| PA2580 | N/A | N/A | N/A | N/A | N/A | N/A | N/A | N/A | N/A | N/A | N/A | N/A | N/A | N/A | N/A | N/A | N/A | N/A |
| PA2581 | N/A | N/A | PA2581 | PA2581 | PA2581 | N/A | PA2581 | PA2581 | PA2581 | PA2581 | PA2581 | PA2581 | PA2581 | PA2581 | N/A | N/A | PA2581 | N/A |
| PA2582 | PA2582 | PA2582 | PA2582 | PA2582 | PA2582 | PA2582 | PA2582 | N/A | PA2582 | PA2582 | PA2582 | PA2582 | PA2582 | PA2582 | PA2582 | N/A | N/A | N/A |
| PA2583 | N/A | N/A | N/A | PA2583 | N/A | N/A | N/A | N/A | PA2583 | N/A | PA2583 | PA2583 | N/A | PA2583 | N/A | N/A | N/A | N/A |
| PA2584 | PA2584 | N/A | PA2584 | N/A | N/A | N/A | N/A | N/A | PA2584 | PA2584 | PA2584 | N/A | N/A | PA2584 | N/A | N/A | PA2584 | N/A |
| PA2585 | N/A | N/A | N/A | PA2585 | N/A | N/A | N/A | PA2585 | PA2585 | N/A | N/A | PA2585 | PA2585 | N/A | N/A | N/A | N/A | N/A |
| PA2586 | N/A | N/A | N/A | PA2586 | PA2586 | N/A | N/A | N/A | PA2586 | PA2586 | PA2586 | PA2586 | PA2586 | N/A | N/A | N/A | N/A | N/A |
| PA2587 | N/A | PA2587 | PA2587 | PA2587 | PA2587 | PA2587 | PA2587 | PA2587 | PA2587 | PA2587 | PA2587 | PA2587 | PA2587 | PA2587 | N/A | N/A | PA2587 | N/A |
| PA2588 | N/A | N/A | N/A | N/A | N/A | N/A | N/A | PA2588 | PA2588 | N/A | N/A | PA2588 | PA2588 | N/A | N/A | N/A | N/A | N/A |
| N/A | PA2590 | N/A | N/A | N/A | N/A | N/A | N/A | N/A | N/A | N/A | N/A | N/A | N/A | N/A | N/A | N/A | N/A | N/A |
| N/A | N/A | N/A | N/A | N/A | PA2591 | N/A | N/A | N/A | N/A | N/A | N/A | N/A | N/A | N/A | N/A | N/A | N/A | N/A |
| PA2592 | N/A | N/A | N/A | PA2592 | PA2592 | N/A | PA2592 | PA2592 | PA2592 | PA2592 | PA2592 | PA2592 | PA2592 | N/A | N/A | N/A | N/A | N/A |
| N/A | N/A | N/A | N/A | N/A | N/A | N/A | N/A | N/A | PA2593 | N/A | N/A | N/A | N/A | N/A | N/A | N/A | N/A | N/A |
| PA2594 | N/A | N/A | N/A | PA2594 | N/A | N/A | N/A | N/A | N/A | N/A | N/A | N/A | N/A | N/A | N/A | N/A | N/A | N/A |
| N/A | N/A | N/A | N/A | PA2599 | N/A | N/A | N/A | N/A | N/A | N/A | N/A | N/A | N/A | N/A | N/A | N/A | N/A | N/A |
| PA2601 | N/A | N/A | N/A | PA2601 | N/A | PA2601 | PA2601 | N/A | PA2601 | PA2601 | PA2601 | PA2601 | PA2601 | N/A | N/A | N/A | N/A | N/A |
| PA2603 | N/A | N/A | N/A | PA2603 | N/A | N/A | N/A | N/A | N/A | N/A | N/A | N/A | N/A | N/A | N/A | N/A | N/A | N/A |
| PA2604 | N/A | N/A | N/A | N/A | N/A | N/A | N/A | N/A | N/A | N/A | N/A | PA2604 | N/A | PA2604 | N/A | N/A | PA2604 | N/A |
| PA2605 | PA2605 | N/A | N/A | PA2605 | N/A | PA2605 | N/A | N/A | PA2605 | N/A | PA2605 | PA2605 | PA2605 | N/A | N/A | N/A | N/A | N/A |
| PA2606 | N/A | N/A | PA2606 | N/A | N/A | N/A | N/A | N/A | N/A | N/A | PA2606 | N/A | N/A | N/A | N/A | N/A | N/A | N/A |
| PA2607 | N/A | N/A | N/A | PA2607 | N/A | N/A | N/A | N/A | N/A | N/A | PA2607 | N/A | PA2607 | N/A | N/A | N/A | N/A | N/A |
| N/A | N/A | N/A | N/A | PA2608 | PA2608 | N/A | N/A | N/A | N/A | N/A | N/A | PA2608 | N/A | N/A | N/A | N/A | N/A | N/A |
| PA2609 | N/A | N/A | PA2609 | PA2609 | PA2609 | PA2609 | N/A | N/A | PA2609 | N/A | PA2609 | N/A | PA2609 | N/A | N/A | N/A | N/A | N/A |
| PA2610 | N/A | N/A | N/A | PA2610 | PA2610 | N/A | N/A | N/A | N/A | N/A | N/A | N/A | N/A | N/A | N/A | N/A | N/A | N/A |
| PA2611 | N/A | PA2611 | PA2611 | PA2611 | PA2611 | N/A | N/A | PA2611 | PA2611 | PA2611 | PA2611 | PA2611 | PA2611 | N/A | N/A | N/A | PA2611 | N/A |
| PA2612 | PA2612 | PA2612 | PA2612 | PA2612 | PA2612 | PA2612 | PA2612 | PA2612 | PA2612 | PA2612 | PA2612 | PA2612 | PA2612 | N/A | PA2612 | N/A | N/A | N/A |
| PA2613 | PA2613 | PA2613 | PA2613 | PA2613 | PA2613 | PA2613 | PA2613 | PA2613 | PA2613 | PA2613 | PA2613 | PA2613 | PA2613 | PA2613 | N/A | N/A | N/A | N/A |
| PA2614 | PA2614 | PA2614 | PA2614 | PA2614 | PA2614 | PA2614 | PA2614 | N/A | PA2614 | PA2614 | PA2614 | PA2614 | PA2614 | PA2614 | PA2614 | PA2614 | PA2614 | N/A |
| PA2615 | PA2615 | PA2615 | PA2615 | PA2615 | PA2615 | PA2615 | N/A | PA2615 | PA2615 | PA2615 | PA2615 | PA2615 | PA2615 | PA2615 | N/A | N/A | PA2615 | N/A |
| PA2616 | PA2616 | PA2616 | PA2616 | PA2616 | PA2616 | PA2616 | N/A | N/A | PA2616 | PA2616 | PA2616 | PA2616 | PA2616 | N/A | PA2616 | PA2616 | N/A | N/A |
| PA2617 | N/A | N/A | N/A | PA2617 | N/A | N/A | N/A | N/A | N/A | PA2617 | N/A | N/A | N/A | N/A | N/A | N/A | N/A | N/A |
| PA2618 | N/A | N/A | N/A | N/A | PA2618 | N/A | N/A | N/A | N/A | N/A | N/A | N/A | N/A | N/A | N/A | N/A | N/A | N/A |
| PA2619 | PA2619 | PA2619 | PA2619 | PA2619 | PA2619 | PA2619 | PA2619 | N/A | PA2619 | PA2619 | PA2619 | PA2619 | PA2619 | PA2619 | PA2619 | N/A | PA2619 | PA2619 |
| PA2620 | PA2620 | PA2620 | PA2620 | PA2620 | PA2620 | PA2620 | N/A | PA2620 | PA2620 | PA2620 | PA2620 | PA2620 | PA2620 | PA2620 | N/A | N/A | N/A | N/A |
| PA2621 | PA2621 | N/A | N/A | N/A | PA2621 | N/A | N/A | N/A | N/A | N/A | PA2621 | N/A | PA2621 | N/A | N/A | N/A | N/A | N/A |
| PA2622 | PA2622 | PA2622 | N/A | PA2622 | PA2622 | PA2622 | N/A | N/A | PA2622 | PA2622 | PA2622 | PA2622 | PA2622 | N/A | PA2622 | N/A | PA2622 | N/A |
| PA2623 | PA2623 | PA2623 | PA2623 | PA2623 | PA2623 | PA2623 | PA2623 | PA2623 | PA2623 | PA2623 | PA2623 | PA2623 | PA2623 | PA2623 | PA2623 | PA2623 | PA2623 | PA2623 |
| PA2624 | PA2624 | PA2624 | PA2624 | PA2624 | PA2624 | PA2624 | PA2624 | PA2624 | PA2624 | PA2624 | PA2624 | PA2624 | PA2624 | PA2624 | PA2624 | PA2624 | N/A | N/A |
| PA2625 | N/A | N/A | N/A | PA2625 | PA2625 | N/A | N/A | N/A | PA2625 | N/A | PA2625 | N/A | N/A | N/A | N/A | N/A | N/A | N/A |
| PA2626 | PA2626 | PA2626 | PA2626 | PA2626 | PA2626 | PA2626 | N/A | PA2626 | PA2626 | PA2626 | PA2626 | PA2626 | PA2626 | N/A | PA2626 | N/A | N/A | N/A |
| PA2627 | N/A | PA2627 | PA2627 | PA2627 | N/A | PA2627 | PA2627 | N/A | PA2627 | PA2627 | PA2627 | PA2627 | PA2627 | PA2627 | N/A | N/A | PA2627 | N/A |
| N/A | N/A | N/A | N/A | N/A | PA2628 | N/A | N/A | N/A | N/A | N/A | N/A | N/A | N/A | N/A | N/A | N/A | N/A | N/A |
| PA2629 | PA2629 | PA2629 | PA2629 | PA2629 | PA2629 | PA2629 | PA2629 | N/A | PA2629 | PA2629 | PA2629 | PA2629 | PA2629 | PA2629 | PA2629 | N/A | N/A | N/A |
| PA2630 | PA2630 | PA2630 | PA2630 | PA2630 | PA2630 | PA2630 | PA2630 | PA2630 | PA2630 | PA2630 | PA2630 | PA2630 | PA2630 | PA2630 | PA2630 | N/A | N/A | N/A |
| PA2631 | PA2631 | PA2631 | PA2631 | PA2631 | PA2631 | N/A | N/A | N/A | PA2631 | N/A | PA2631 | PA2631 | N/A | N/A | PA2631 | N/A | N/A | N/A |
| PA2632 | N/A | N/A | N/A | PA2632 | PA2632 | N/A | N/A | N/A | N/A | PA2632 | N/A | N/A | N/A | N/A | N/A | N/A | N/A | N/A |
| PA2633 | N/A | N/A | N/A | PA2633 | N/A | N/A | N/A | N/A | N/A | N/A | PA2633 | N/A | N/A | N/A | N/A | N/A | PA2633 | N/A |
| PA2634 | PA2634 | PA2634 | PA2634 | PA2634 | PA2634 | PA2634 | PA2634 | PA2634 | PA2634 | PA2634 | PA2634 | PA2634 | PA2634 | PA2634 | PA2634 | PA2634 | N/A | N/A |
| PA2637 | PA2637 | PA2637 | N/A | PA2637 | PA2637 | PA2637 | N/A | PA2637 | PA2637 | PA2637 | PA2637 | PA2637 | PA2637 | PA2637 | N/A | N/A | PA2637 | N/A |
| PA2638 | PA2638 | PA2638 | PA2638 | PA2638 | PA2638 | PA2638 | PA2638 | PA2638 | PA2638 | PA2638 | PA2638 | PA2638 | PA2638 | PA2638 | PA2638 | N/A | PA2638 | PA2638 |
| PA2639 | PA2639 | PA2639 | PA2639 | PA2639 | PA2639 | PA2639 | PA2639 | PA2639 | PA2639 | PA2639 | PA2639 | PA2639 | PA2639 | PA2639 | PA2639 | N/A | PA2639 | PA2639 |
| PA2640 | PA2640 | PA2640 | PA2640 | PA2640 | PA2640 | PA2640 | PA2640 | N/A | PA2640 | PA2640 | PA2640 | PA2640 | PA2640 | PA2640 | N/A | N/A | PA2640 | PA2640 |
| PA2641 | PA2641 | PA2641 | PA2641 | PA2641 | PA2641 | PA2641 | PA2641 | PA2641 | PA2641 | PA2641 | PA2641 | PA2641 | PA2641 | PA2641 | PA2641 | PA2641 | PA2641 | PA2641 |
| PA2642 | PA2642 | PA2642 | PA2642 | PA2642 | PA2642 | PA2642 | PA2642 | PA2642 | PA2642 | PA2642 | PA2642 | PA2642 | PA2642 | PA2642 | PA2642 | PA2642 | PA2642 | PA2642 |
| PA2643 | PA2643 | PA2643 | N/A | PA2643 | N/A | PA2643 | N/A | PA2643 | N/A | PA2643 | PA2643 | PA2643 | PA2643 | PA2643 | N/A | N/A | PA2643 | N/A |
| PA2644 | PA2644 | PA2644 | PA2644 | PA2644 | PA2644 | PA2644 | PA2644 | PA2644 | PA2644 | PA2644 | PA2644 | PA2644 | PA2644 | PA2644 | PA2644 | N/A | PA2644 | PA2644 |
| PA2645 | PA2645 | N/A | N/A | N/A | N/A | PA2645 | N/A | N/A | PA2645 | PA2645 | PA2645 | PA2645 | PA2645 | PA2645 | N/A | N/A | PA2645 | N/A |
| PA2646 | N/A | N/A | N/A | N/A | N/A | N/A | N/A | N/A | N/A | N/A | N/A | N/A | N/A | PA2646 | N/A | N/A | PA2646 | N/A |
| PA2647 | PA2647 | PA2647 | PA2647 | PA2647 | N/A | PA2647 | N/A | PA2647 | PA2647 | PA2647 | PA2647 | PA2647 | PA2647 | PA2647 | N/A | N/A | PA2647 | N/A |
| PA2648 | N/A | N/A | N/A | PA2648 | N/A | PA2648 | N/A | PA2648 | PA2648 | PA2648 | PA2648 | PA2648 | PA2648 | PA2648 | N/A | N/A | PA2648 | N/A |
| PA2649 | PA2649 | PA2649 | N/A | PA2649 | N/A | PA2649 | N/A | N/A | PA2649 | PA2649 | PA2649 | PA2649 | PA2649 | PA2649 | N/A | N/A | PA2649 | N/A |
| PA2650 | N/A | N/A | N/A | PA2650 | N/A | N/A | PA2650 | N/A | N/A | N/A | PA2650 | PA2650 | PA2650 | N/A | N/A | N/A | N/A | N/A |
| PA2651 | N/A | N/A | N/A | N/A | N/A | PA2651 | N/A | N/A | N/A | PA2651 | PA2651 | PA2651 | PA2651 | PA2651 | N/A | N/A | N/A | N/A |
| PA2652 | PA2652 | PA2652 | PA2652 | PA2652 | N/A | PA2652 | PA2652 | PA2652 | PA2652 | PA2652 | PA2652 | PA2652 | PA2652 | PA2652 | PA2652 | N/A | PA2652 | PA2652 |
| N/A | PA2653 | PA2653 | N/A | N/A | N/A | N/A | N/A | N/A | N/A | N/A | N/A | N/A | N/A | PA2653 | N/A | N/A | N/A | N/A |
| PA2654 | PA2654 | PA2654 | PA2654 | PA2654 | N/A | PA2654 | PA2654 | PA2654 | PA2654 | PA2654 | PA2654 | PA2654 | PA2654 | PA2654 | PA2654 | N/A | PA2654 | PA2654 |
| PA2656 | N/A | N/A | N/A | PA2656 | N/A | N/A | N/A | N/A | N/A | N/A | N/A | N/A | N/A | PA2656 | N/A | N/A | N/A | N/A |
| PA2657 | N/A | N/A | N/A | PA2657 | N/A | N/A | N/A | N/A | PA2657 | PA2657 | PA2657 | PA2657 | PA2657 | N/A | N/A | N/A | N/A | N/A |
| N/A | N/A | N/A | N/A | N/A | PA2658 | N/A | N/A | N/A | N/A | N/A | N/A | N/A | N/A | N/A | N/A | N/A | N/A | N/A |
| PA2659 | N/A | N/A | N/A | N/A | N/A | N/A | N/A | N/A | PA2659 | PA2659 | PA2659 | PA2659 | PA2659 | N/A | N/A | N/A | N/A | N/A |
| PA2660 | N/A | PA2660 | N/A | PA2660 | N/A | N/A | N/A | PA2660 | N/A | N/A | N/A | N/A | N/A | PA2660 | N/A | N/A | PA2660 | N/A |
| PA2662 | N/A | N/A | N/A | N/A | N/A | N/A | N/A | N/A | N/A | N/A | N/A | N/A | N/A | PA2662 | N/A | N/A | PA2662 | N/A |
| N/A | N/A | N/A | N/A | N/A | PA2663 | N/A | N/A | N/A | N/A | N/A | N/A | N/A | N/A | N/A | N/A | N/A | N/A | N/A |
| PA2664 | N/A | N/A | N/A | PA2664 | PA2664 | N/A | N/A | N/A | N/A | N/A | N/A | N/A | PA2664 | N/A | N/A | N/A | N/A | N/A |
| PA2665 | N/A | N/A | N/A | PA2665 | PA2665 | N/A | N/A | PA2665 | N/A | N/A | N/A | PA2665 | PA2665 | N/A | N/A | N/A | N/A | N/A |
| N/A | N/A | N/A | N/A | N/A | N/A | N/A | N/A | N/A | PA2666 | PA2666 | PA2666 | PA2666 | N/A | N/A | N/A | N/A | N/A | N/A |
| PA2667 | PA2667 | PA2667 | PA2667 | PA2667 | PA2667 | PA2667 | PA2667 | PA2667 | PA2667 | PA2667 | PA2667 | PA2667 | PA2667 | PA2667 | PA2667 | N/A | PA2667 | PA2667 |
| N/A | N/A | N/A | N/A | N/A | N/A | N/A | N/A | PA2677 | N/A | N/A | N/A | N/A | N/A | N/A | N/A | N/A | N/A | N/A |
| N/A | N/A | N/A | N/A | N/A | PA2678 | N/A | N/A | N/A | N/A | N/A | N/A | N/A | N/A | N/A | N/A | N/A | N/A | N/A |
| PA2679 | N/A | N/A | N/A | PA2679 | PA2679 | PA2679 | N/A | N/A | PA2679 | PA2679 | PA2679 | PA2679 | PA2679 | N/A | N/A | N/A | N/A | N/A |
| PA2680 | N/A | N/A | N/A | N/A | N/A | N/A | N/A | N/A | N/A | N/A | PA2680 | N/A | PA2680 | N/A | N/A | N/A | N/A | N/A |
| PA2683 | N/A | N/A | N/A | PA2683 | N/A | N/A | N/A | N/A | N/A | N/A | N/A | N/A | N/A | N/A | N/A | N/A | PA2683 | N/A |
| PA2684 | N/A | N/A | N/A | PA2684 | N/A | N/A | N/A | N/A | PA2684 | PA2684 | PA2684 | PA2684 | PA2684 | N/A | PA2684 | N/A | N/A | PA2684 |
| PA2685 | N/A | PA2685 | N/A | PA2685 | N/A | PA2685 | N/A | N/A | PA2685 | PA2685 | PA2685 | PA2685 | PA2685 | PA2685 | N/A | N/A | N/A | N/A |
| N/A | N/A | N/A | N/A | PA2688 | N/A | N/A | N/A | N/A | N/A | N/A | N/A | N/A | N/A | N/A | N/A | N/A | N/A | N/A |
| N/A | N/A | N/A | N/A | PA2691 | N/A | N/A | N/A | N/A | N/A | N/A | N/A | N/A | N/A | N/A | N/A | N/A | N/A | N/A |
| PA2692 | N/A | N/A | N/A | PA2692 | N/A | PA2692 | N/A | N/A | PA2692 | PA2692 | PA2692 | PA2692 | PA2692 | N/A | N/A | N/A | N/A | N/A |
| N/A | N/A | N/A | N/A | N/A | N/A | N/A | N/A | N/A | PA2694 | N/A | N/A | PA2694 | PA2694 | N/A | N/A | N/A | N/A | N/A |
| PA2695 | N/A | N/A | N/A | PA2695 | N/A | N/A | N/A | N/A | N/A | N/A | N/A | N/A | N/A | N/A | N/A | N/A | N/A | N/A |
| N/A | N/A | N/A | N/A | N/A | PA2697 | N/A | N/A | N/A | N/A | N/A | N/A | N/A | N/A | N/A | N/A | N/A | N/A | N/A |
| N/A | N/A | N/A | N/A | N/A | PA2698 | N/A | N/A | N/A | N/A | N/A | N/A | PA2698 | PA2698 | N/A | N/A | N/A | N/A | N/A |
| PA2699 | N/A | N/A | N/A | N/A | PA2699 | N/A | N/A | N/A | PA2699 | N/A | N/A | PA2699 | PA2699 | N/A | N/A | N/A | N/A | N/A |
| PA2701 | N/A | N/A | N/A | N/A | N/A | N/A | N/A | N/A | N/A | N/A | N/A | N/A | N/A | N/A | N/A | N/A | N/A | N/A |
| PA2702 | N/A | N/A | N/A | N/A | N/A | N/A | N/A | N/A | N/A | N/A | N/A | N/A | N/A | N/A | N/A | N/A | N/A | N/A |
| PA2705 | N/A | N/A | N/A | PA2705 | N/A | PA2705 | PA2705 | PA2705 | N/A | N/A | N/A | PA2705 | PA2705 | N/A | N/A | N/A | N/A | N/A |
| PA2706 | N/A | N/A | N/A | N/A | N/A | N/A | N/A | N/A | N/A | N/A | N/A | N/A | N/A | N/A | N/A | N/A | N/A | N/A |
| PA2707 | PA2707 | PA2707 | PA2707 | PA2707 | N/A | PA2707 | PA2707 | PA2707 | PA2707 | PA2707 | PA2707 | PA2707 | PA2707 | PA2707 | N/A | N/A | PA2707 | N/A |
| PA2708 | N/A | N/A | N/A | N/A | PA2708 | N/A | N/A | N/A | N/A | N/A | N/A | N/A | N/A | N/A | N/A | N/A | N/A | N/A |

| | | | | | | | | | | | | | | | | | | |
|---|---|---|---|---|---|---|---|---|---|---|---|---|---|---|---|---|---|---|
| PA2709 | PA2709 | PA2709 | PA2709 | PA2709 | PA2709 | PA2709 | N/A | N/A | PA2709 | PA2709 | PA2709 | PA2709 | PA2709 | N/A | PA2709 | N/A | N/A | N/A |
| N/A | N/A | N/A | N/A | N/A | PA2710 | N/A | N/A | N/A | N/A | N/A | N/A | N/A | N/A | N/A | N/A | N/A | N/A | N/A |
| N/A | N/A | N/A | N/A | N/A | PA2711 | N/A | N/A | N/A | N/A | N/A | N/A | N/A | N/A | N/A | N/A | N/A | N/A | N/A |
| PA2713 | N/A | N/A | N/A | N/A | N/A | N/A | N/A | N/A | N/A | N/A | N/A | N/A | N/A | N/A | N/A | N/A | N/A | N/A |
| PA2716 | N/A | N/A | N/A | N/A | PA2716 | N/A | N/A | N/A | N/A | N/A | N/A | N/A | N/A | N/A | N/A | N/A | N/A | N/A |
| PA2717 | N/A | N/A | N/A | N/A | PA2717 | PA2717 | N/A | N/A | PA2717 | PA2717 | N/A | PA2717 | PA2717 | N/A | N/A | N/A | N/A | N/A |
| PA2718 | N/A | N/A | N/A | N/A | N/A | N/A | N/A | N/A | N/A | N/A | N/A | N/A | N/A | N/A | N/A | N/A | N/A | N/A |
| PA2719 | N/A | N/A | N/A | N/A | N/A | PA2719 | N/A | N/A | N/A | PA2719 | N/A | PA2719 | PA2719 | N/A | N/A | N/A | N/A | N/A |
| PA2720 | N/A | N/A | N/A | PA2720 | PA2720 | N/A | N/A | N/A | PA2720 | PA2720 | PA2720 | PA2720 | PA2720 | N/A | N/A | N/A | N/A | N/A |
| PA2721 | PA2721 | N/A | N/A | N/A | N/A | PA2721 | N/A | N/A | N/A | PA2721 | N/A | PA2721 | PA2721 | N/A | N/A | N/A | N/A | N/A |
| PA2722 | N/A | N/A | N/A | N/A | N/A | N/A | N/A | N/A | N/A | N/A | N/A | N/A | N/A | N/A | N/A | N/A | N/A | N/A |
| PA2724 | N/A | N/A | N/A | N/A | N/A | N/A | N/A | N/A | N/A | N/A | N/A | N/A | N/A | N/A | N/A | N/A | N/A | N/A |
| PA2725 | PA2725 | N/A | PA2725 | PA2725 | N/A | N/A | N/A | PA2725 | PA2725 | PA2725 | PA2725 | PA2725 | PA2725 | N/A | N/A | N/A | PA2725 | N/A |
| PA2727 | N/A | N/A | N/A | PA2727 | N/A | N/A | N/A | PA2727 | PA2727 | N/A | PA2727 | PA2727 | PA2727 | N/A | N/A | N/A | PA2727 | N/A |
| PA2728 | N/A | N/A | N/A | PA2728 | PA2728 | N/A | N/A | N/A | PA2728 | N/A | PA2728 | PA2728 | N/A | PA2728 | N/A | N/A | PA2728 | N/A |
| PA2729 | N/A | N/A | N/A | N/A | PA2729 | N/A | N/A | N/A | PA2729 | N/A | PA2729 | N/A | PA2729 | N/A | N/A | N/A | N/A | N/A |
| PA2730 | N/A | N/A | PA2730 | PA2730 | N/A | N/A | N/A | PA2730 | PA2730 | PA2730 | PA2730 | PA2730 | PA2730 | N/A | N/A | N/A | N/A | N/A |
| PA2731 | N/A | N/A | N/A | PA2731 | PA2731 | N/A | PA2731 | N/A | PA2731 | PA2731 | PA2731 | PA2731 | PA2731 | PA2731 | N/A | N/A | N/A | N/A |
| PA2732 | PA2732 | PA2732 | N/A | PA2732 | N/A | PA2732 | N/A | PA2732 | PA2732 | PA2732 | PA2732 | PA2732 | PA2732 | PA2732 | N/A | N/A | N/A | N/A |
| PA2734 | PA2734 | PA2734 | PA2734 | PA2734 | PA2734 | N/A | N/A | PA2734 | PA2734 | PA2734 | PA2734 | PA2734 | PA2734 | PA2734 | N/A | N/A | N/A | N/A |
| PA2735 | PA2735 | PA2735 | PA2735 | PA2735 | N/A | PA2735 | N/A | PA2735 | PA2735 | PA2735 | PA2735 | PA2735 | PA2735 | PA2735 | PA2735 | N/A | N/A | N/A |
| N/A | N/A | N/A | N/A | N/A | PA2736 | N/A | N/A | N/A | N/A | N/A | N/A | PA2736 | N/A | N/A | N/A | N/A | N/A | N/A |
| PA2737 | N/A | N/A | N/A | PA2737 | PA2737 | PA2737 | N/A | N/A | PA2737 | PA2737 | PA2737 | PA2737 | PA2737 | N/A | N/A | N/A | N/A | N/A |
| PA2738 | N/A | N/A | PA2738 | PA2738 | PA2738 | PA2738 | N/A | N/A | PA2738 | PA2738 | PA2738 | PA2738 | PA2738 | PA2738 | N/A | N/A | PA2738 | PA2738 |
| PA2739 | PA2739 | PA2739 | PA2739 | PA2739 | PA2739 | PA2739 | N/A | PA2739 | PA2739 | PA2739 | PA2739 | PA2739 | PA2739 | PA2739 | PA2739 | N/A | N/A | N/A |
| PA2740 | PA2740 | PA2740 | PA2740 | PA2740 | PA2740 | PA2740 | PA2740 | PA2740 | PA2740 | PA2740 | PA2740 | PA2740 | PA2740 | PA2740 | PA2740 | N/A | N/A | N/A |
| PA2741 | PA2741 | PA2741 | PA2741 | PA2741 | PA2741 | PA2741 | PA2741 | PA2741 | PA2741 | PA2741 | PA2741 | PA2741 | PA2741 | PA2741 | PA2741 | PA2741 | PA2741 | PA2741 |
| N/A | N/A | PA2742 | PA2742 | PA2742 | N/A | N/A | N/A | N/A | N/A | N/A | N/A | N/A | N/A | N/A | N/A | N/A | N/A | N/A |
| PA2743 | PA2743 | PA2743 | PA2743 | PA2743 | PA2743 | PA2743 | PA2743 | PA2743 | PA2743 | PA2743 | PA2743 | PA2743 | PA2743 | PA2743 | PA2743 | N/A | PA2743 | PA2743 |
| PA2744 | PA2744 | PA2744 | PA2744 | PA2744 | PA2744 | PA2744 | N/A | PA2744 | PA2744 | PA2744 | PA2744 | PA2744 | PA2744 | PA2744 | PA2744 | N/A | N/A | N/A |
| PA2745 | PA2745 | N/A | N/A | PA2745 | N/A | N/A | N/A | N/A | N/A | N/A | PA2745 | N/A | N/A | N/A | N/A | N/A | N/A | N/A |
| PA2747 | N/A | N/A | N/A | N/A | N/A | PA2747 | N/A | N/A | N/A | N/A | PA2747 | PA2747 | PA2747 | N/A | N/A | N/A | N/A | N/A |
| PA2750 | N/A | N/A | N/A | PA2750 | N/A | N/A | N/A | N/A | N/A | N/A | N/A | N/A | N/A | PA2750 | N/A | N/A | N/A | N/A |
| PA2751 | N/A | N/A | N/A | N/A | N/A | PA2751 | N/A | N/A | N/A | N/A | N/A | N/A | N/A | PA2751 | N/A | N/A | N/A | N/A |
| N/A | N/A | N/A | N/A | N/A | N/A | N/A | N/A | N/A | N/A | PA2753 | PA2753 | N/A | N/A | N/A | N/A | N/A | N/A | N/A |
| PA2754 | PA2754 | N/A | PA2754 | N/A | PA2754 | PA2754 | N/A | N/A | PA2754 | PA2754 | PA2754 | PA2754 | PA2754 | PA2754 | N/A | N/A | N/A | N/A |
| PA2755 | PA2755 | PA2755 | PA2755 | PA2755 | PA2755 | PA2755 | PA2755 | N/A | PA2755 | PA2755 | PA2755 | PA2755 | PA2755 | N/A | PA2755 | N/A | PA2755 | N/A |
| PA2756 | N/A | N/A | N/A | PA2756 | N/A | PA2756 | PA2756 | N/A | PA2756 | PA2756 | PA2756 | PA2756 | PA2756 | N/A | N/A | N/A | N/A | N/A |
| PA2759 | N/A | N/A | N/A | N/A | PA2759 | N/A | N/A | N/A | N/A | N/A | N/A | N/A | N/A | N/A | N/A | N/A | N/A | N/A |
| PA2760 | PA2760 | PA2760 | PA2760 | PA2760 | PA2760 | PA2760 | PA2760 | PA2760 | PA2760 | PA2760 | PA2760 | PA2760 | PA2760 | PA2760 | PA2760 | PA2760 | PA2760 | N/A |
| N/A | N/A | N/A | N/A | N/A | N/A | N/A | PA2761 | N/A | N/A | PA2761 | N/A | N/A | N/A | PA2761 | N/A | N/A | N/A | N/A |
| PA2762 | N/A | N/A | N/A | N/A | N/A | PA2762 | N/A | N/A | N/A | N/A | PA2762 | PA2762 | PA2762 | N/A | N/A | N/A | N/A | N/A |
| N/A | N/A | N/A | N/A | N/A | PA2763 | N/A | N/A | N/A | N/A | N/A | N/A | N/A | N/A | N/A | N/A | N/A | N/A | N/A |
| PA2764 | N/A | N/A | N/A | PA2764 | N/A | N/A | N/A | N/A | N/A | PA2764 | PA2764 | N/A | PA2764 | N/A | N/A | N/A | N/A | N/A |
| PA2765 | PA2765 | PA2765 | PA2765 | PA2765 | N/A | PA2765 | PA2765 | N/A | PA2765 | PA2765 | PA2765 | PA2765 | PA2765 | N/A | PA2765 | PA2765 | N/A | N/A |
| PA2766 | N/A | N/A | N/A | N/A | N/A | N/A | N/A | N/A | N/A | N/A | N/A | N/A | N/A | N/A | N/A | N/A | N/A | N/A |
| N/A | N/A | N/A | N/A | N/A | PA2769 | N/A | N/A | N/A | PA2769 | N/A | N/A | N/A | N/A | N/A | N/A | N/A | N/A | N/A |
| PA2770 | PA2770 | PA2770 | PA2770 | PA2770 | PA2770 | PA2770 | PA2770 | N/A | PA2770 | PA2770 | PA2770 | PA2770 | PA2770 | N/A | N/A | N/A | N/A | N/A |
| PA2771 | N/A | N/A | N/A | N/A | PA2771 | N/A | N/A | N/A | N/A | PA2771 | N/A | N/A | PA2771 | N/A | N/A | N/A | N/A | N/A |
| PA2774 | N/A | PA2774 | N/A | PA2774 | N/A | N/A | PA2774 | N/A | PA2774 | PA2774 | PA2774 | N/A | N/A | PA2774 | N/A | N/A | PA2774 | PA2774 |
| PA2775 | PA2775 | N/A | N/A | N/A | PA2775 | N/A | PA2775 | N/A | PA2775 | N/A | PA2775 | PA2775 | PA2775 | PA2775 | N/A | N/A | PA2775 | PA2775 |
| PA2776 | PA2776 | PA2776 | PA2776 | PA2776 | PA2776 | PA2776 | PA2776 | N/A | PA2776 | PA2776 | PA2776 | PA2776 | PA2776 | PA2776 | PA2776 | N/A | PA2776 | PA2776 |
| PA2777 | N/A | N/A | N/A | N/A | N/A | N/A | N/A | N/A | N/A | N/A | N/A | N/A | N/A | N/A | N/A | N/A | N/A | N/A |
| PA2778 | N/A | N/A | N/A | N/A | N/A | N/A | N/A | N/A | N/A | N/A | N/A | N/A | N/A | N/A | N/A | N/A | N/A | N/A |
| N/A | N/A | N/A | N/A | N/A | N/A | N/A | N/A | N/A | N/A | N/A | N/A | N/A | N/A | N/A | PA2783 | PA2783 | PA2783 | PA2783 |
| N/A | N/A | N/A | N/A | N/A | PA2786 | N/A | N/A | N/A | N/A | N/A | N/A | N/A | N/A | N/A | N/A | N/A | N/A | N/A |
| PA2787 | N/A | N/A | N/A | N/A | PA2787 | N/A | N/A | N/A | N/A | N/A | N/A | N/A | PA2787 | N/A | N/A | N/A | N/A | N/A |
| PA2788 | PA2788 | PA2788 | PA2788 | PA2788 | N/A | PA2788 | PA2788 | PA2788 | PA2788 | PA2788 | PA2788 | PA2788 | PA2788 | PA2788 | PA2788 | N/A | PA2788 | PA2788 |
| PA2789 | N/A | N/A | N/A | N/A | N/A | N/A | N/A | N/A | PA2789 | N/A | N/A | N/A | N/A | PA2789 | N/A | N/A | PA2789 | N/A |
| PA2790 | N/A | N/A | N/A | N/A | N/A | N/A | N/A | N/A | N/A | N/A | N/A | N/A | N/A | PA2790 | N/A | N/A | PA2790 | N/A |
| N/A | N/A | PA2792 | N/A | N/A | N/A | N/A | N/A | N/A | N/A | N/A | N/A | N/A | N/A | N/A | PA2792 | N/A | N/A | N/A |
| PA2793 | N/A | PA2793 | N/A | PA2793 | N/A | N/A | PA2793 | N/A | N/A | N/A | N/A | N/A | N/A | N/A | PA2793 | N/A | N/A | N/A |
| PA2794 | N/A | N/A | N/A | N/A | PA2794 | N/A | N/A | N/A | N/A | N/A | N/A | N/A | PA2794 | N/A | N/A | N/A | N/A | N/A |
| PA2795 | N/A | N/A | PA2795 | PA2795 | PA2795 | N/A | N/A | N/A | N/A | N/A | N/A | N/A | N/A | N/A | PA2795 | N/A | N/A | N/A |
| PA2796 | PA2796 | PA2796 | PA2796 | PA2796 | PA2796 | PA2796 | N/A | N/A | PA2796 | PA2796 | PA2796 | PA2796 | PA2796 | N/A | N/A | PA2796 | N/A | N/A |
| PA2797 | PA2797 | PA2797 | PA2797 | PA2797 | PA2797 | PA2797 | PA2797 | N/A | PA2797 | PA2797 | PA2797 | PA2797 | PA2797 | N/A | N/A | N/A | N/A | N/A |
| PA2798 | N/A | PA2798 | PA2798 | PA2798 | N/A | N/A | N/A | N/A | PA2798 | PA2798 | PA2798 | PA2798 | PA2798 | N/A | PA2798 | N/A | N/A | N/A |
| PA2799 | N/A | N/A | N/A | N/A | PA2799 | N/A | N/A | N/A | N/A | N/A | PA2799 | PA2799 | N/A | N/A | N/A | N/A | N/A | N/A |
| PA2800 | PA2800 | PA2800 | PA2800 | PA2800 | N/A | PA2800 | PA2800 | PA2800 | PA2800 | PA2800 | PA2800 | PA2800 | PA2800 | N/A | PA2800 | N/A | PA2800 | PA2800 |
| PA2801 | N/A | N/A | PA2801 | N/A | N/A | PA2801 | N/A | N/A | N/A | N/A | PA2801 | PA2801 | PA2801 | N/A | N/A | N/A | N/A | N/A |
| PA2802 | N/A | N/A | N/A | PA2802 | N/A | N/A | N/A | N/A | N/A | N/A | PA2802 | N/A | N/A | N/A | N/A | N/A | N/A | N/A |
| PA2805 | PA2805 | PA2805 | PA2805 | PA2805 | PA2805 | N/A | N/A | N/A | N/A | N/A | PA2805 | N/A | PA2805 | N/A | N/A | N/A | N/A | N/A |
| PA2806 | PA2806 | PA2806 | N/A | PA2806 | PA2806 | N/A | PA2806 | PA2806 | PA2806 | PA2806 | PA2806 | PA2806 | PA2806 | N/A | PA2806 | N/A | N/A | N/A |
| PA2809 | N/A | N/A | N/A | N/A | N/A | N/A | N/A | N/A | N/A | PA2809 | PA2809 | PA2809 | PA2809 | N/A | N/A | N/A | N/A | N/A |
| PA2810 | N/A | N/A | N/A | N/A | N/A | N/A | N/A | N/A | N/A | N/A | N/A | N/A | N/A | N/A | N/A | N/A | N/A | N/A |
| PA2811 | N/A | N/A | N/A | N/A | N/A | PA2811 | N/A | N/A | N/A | N/A | N/A | N/A | N/A | PA2811 | N/A | N/A | N/A | N/A |
| PA2812 | PA2812 | N/A | PA2812 | PA2812 | PA2812 | N/A | PA2812 | PA2812 | PA2812 | PA2812 | PA2812 | PA2812 | PA2812 | PA2812 | N/A | N/A | PA2812 | N/A |
| PA2813 | PA2813 | N/A | N/A | PA2813 | PA2813 | PA2813 | N/A | N/A | PA2813 | PA2813 | PA2813 | PA2813 | PA2813 | N/A | N/A | N/A | N/A | N/A |
| N/A | N/A | N/A | N/A | N/A | PA2814 | N/A | N/A | N/A | N/A | N/A | N/A | N/A | N/A | N/A | N/A | N/A | N/A | N/A |
| PA2815 | N/A | N/A | N/A | PA2815 | PA2815 | PA2815 | N/A | N/A | PA2815 | PA2815 | PA2815 | PA2815 | PA2815 | PA2815 | N/A | PA2815 | PA2815 | N/A |
| PA2816 | N/A | N/A | N/A | N/A | PA2816 | N/A | N/A | N/A | N/A | PA2816 | N/A | PA2816 | N/A | N/A | N/A | N/A | N/A | N/A |
| PA2817 | PA2817 | PA2817 | PA2817 | PA2817 | PA2817 | N/A | PA2817 | N/A | PA2817 | PA2817 | PA2817 | PA2817 | PA2817 | N/A | N/A | N/A | N/A | N/A |
| PA2818 | N/A | N/A | N/A | N/A | N/A | N/A | N/A | N/A | N/A | N/A | N/A | N/A | N/A | N/A | N/A | N/A | N/A | N/A |
| PA2820 | PA2820 | N/A | N/A | PA2820 | PA2820 | PA2820 | N/A | N/A | PA2820 | N/A | PA2820 | N/A | N/A | PA2820 | N/A | N/A | PA2820 | N/A |
| PA2821 | PA2821 | N/A | N/A | PA2821 | PA2821 | PA2821 | PA2821 | N/A | PA2821 | PA2821 | PA2821 | PA2821 | PA2821 | N/A | N/A | N/A | N/A | N/A |
| PA2822 | PA2822 | N/A | N/A | PA2822 | PA2822 | N/A | N/A | N/A | PA2822 | PA2822 | PA2822 | PA2822 | PA2822 | N/A | N/A | N/A | N/A | N/A |
| PA2823 | PA2823 | PA2823 | PA2823 | PA2823 | PA2823 | PA2823 | PA2823 | N/A | PA2823 | PA2823 | PA2823 | PA2823 | PA2823 | PA2823 | N/A | N/A | N/A | N/A |
| PA2824 | N/A | N/A | N/A | N/A | N/A | N/A | N/A | N/A | PA2824 | PA2824 | N/A | PA2824 | PA2824 | PA2824 | N/A | N/A | N/A | N/A |
| PA2825 | N/A | N/A | N/A | PA2825 | PA2825 | N/A | N/A | N/A | N/A | N/A | PA2825 | PA2825 | N/A | N/A | N/A | N/A | N/A | N/A |
| PA2826 | N/A | PA2826 | PA2826 | PA2826 | PA2826 | PA2826 | PA2826 | N/A | PA2826 | PA2826 | PA2826 | PA2826 | PA2826 | PA2826 | N/A | N/A | N/A | N/A |
| PA2827 | N/A | N/A | N/A | N/A | PA2827 | PA2827 | N/A | N/A | PA2827 | PA2827 | PA2827 | PA2827 | PA2827 | N/A | N/A | N/A | N/A | N/A |
| PA2828 | PA2828 | PA2828 | PA2828 | PA2828 | PA2828 | PA2828 | N/A | N/A | PA2828 | PA2828 | PA2828 | PA2828 | N/A | N/A | PA2828 | N/A | N/A | N/A |
| PA2829 | N/A | N/A | N/A | PA2829 | N/A | N/A | N/A | N/A | PA2829 | N/A | PA2829 | PA2829 | N/A | N/A | N/A | N/A | N/A | N/A |
| PA2830 | PA2830 | PA2830 | PA2830 | PA2830 | PA2830 | PA2830 | N/A | N/A | PA2830 | PA2830 | PA2830 | PA2830 | PA2830 | PA2830 | PA2830 | N/A | PA2830 | N/A |
| PA2831 | PA2831 | PA2831 | N/A | PA2831 | PA2831 | N/A | N/A | N/A | PA2831 | PA2831 | PA2831 | PA2831 | PA2831 | N/A | PA2831 | N/A | N/A | N/A |
| PA2832 | N/A | N/A | N/A | N/A | PA2832 | N/A | N/A | N/A | PA2832 | PA2832 | PA2832 | PA2832 | PA2832 | N/A | N/A | N/A | N/A | N/A |
| PA2833 | N/A | N/A | N/A | N/A | N/A | N/A | N/A | N/A | N/A | N/A | N/A | N/A | N/A | N/A | N/A | N/A | N/A | N/A |
| PA2834 | N/A | N/A | N/A | PA2834 | N/A | N/A | N/A | N/A | N/A | N/A | N/A | N/A | N/A | N/A | N/A | N/A | N/A | N/A |
| N/A | N/A | N/A | N/A | N/A | N/A | N/A | N/A | N/A | N/A | N/A | N/A | N/A | N/A | N/A | PA2837 | N/A | N/A | N/A |
| PA2840 | PA2840 | PA2840 | PA2840 | PA2840 | PA2840 | PA2840 | PA2840 | PA2840 | PA2840 | PA2840 | PA2840 | PA2840 | PA2840 | PA2840 | PA2840 | N/A | PA2840 | N/A |
| PA2841 | N/A | N/A | N/A | PA2841 | PA2841 | PA2841 | N/A | N/A | PA2841 | PA2841 | PA2841 | PA2841 | PA2841 | N/A | N/A | N/A | N/A | N/A |
| N/A | N/A | N/A | N/A | PA2842 | N/A | N/A | N/A | N/A | N/A | N/A | N/A | N/A | N/A | N/A | N/A | N/A | N/A | N/A |
| PA2843 | PA2843 | PA2843 | N/A | PA2843 | PA2843 | N/A | N/A | PA2843 | PA2843 | PA2843 | PA2843 | PA2843 | PA2843 | N/A | PA2843 | N/A | N/A | N/A |
| PA2844 | N/A | N/A | N/A | N/A | N/A | N/A | N/A | N/A | N/A | N/A | N/A | N/A | PA2844 | N/A | N/A | N/A | N/A | N/A |
| PA2846 | N/A | N/A | N/A | N/A | N/A | N/A | N/A | N/A | N/A | N/A | N/A | N/A | N/A | N/A | N/A | N/A | N/A | N/A |
| N/A | N/A | N/A | N/A | N/A | PA2848 | N/A | N/A | N/A | N/A | N/A | N/A | N/A | N/A | N/A | N/A | N/A | N/A | N/A |
| PA2849 | PA2849 | N/A | N/A | PA2849 | PA2849 | PA2849 | N/A | N/A | PA2849 | PA2849 | PA2849 | PA2849 | PA2849 | N/A | N/A | N/A | N/A | N/A |
| PA2850 | PA2850 | PA2850 | PA2850 | PA2850 | PA2850 | PA2850 | PA2850 | N/A | PA2850 | PA2850 | PA2850 | PA2850 | PA2850 | N/A | PA2850 | N/A | N/A | N/A |
| PA2851 | PA2851 | PA2851 | PA2851 | PA2851 | PA2851 | PA2851 | PA2851 | N/A | PA2851 | PA2851 | PA2851 | PA2851 | PA2851 | N/A | PA2851 | N/A | N/A | N/A |

| | | | | | | | | | | | | | | | | | | |
|---|---|---|---|---|---|---|---|---|---|---|---|---|---|---|---|---|---|---|
| PA2852 | N/A | N/A | N/A | PA2852 | PA2852 | N/A | N/A | N/A | N/A | N/A | N/A | N/A | PA2852 | N/A | N/A | N/A | N/A | N/A |
| PA2853 | PA2853 | PA2853 | PA2853 | PA2853 | PA2853 | PA2853 | PA2853 | PA2853 | PA2853 | PA2853 | PA2853 | PA2853 | PA2853 | PA2853 | PA2853 | PA2853 | PA2853 | PA2853 |
| PA2854 | PA2854 | PA2854 | PA2854 | PA2854 | PA2854 | N/A | PA2854 | PA2854 | PA2854 | PA2854 | PA2854 | PA2854 | PA2854 | N/A | PA2854 | N/A | N/A | N/A |
| N/A | N/A | N/A | N/A | N/A | PA2855 | N/A | N/A | N/A | N/A | N/A | N/A | N/A | N/A | N/A | N/A | N/A | N/A | N/A |
| PA2856 | N/A | N/A | N/A | PA2856 | PA2856 | N/A | N/A | N/A | N/A | PA2856 | PA2856 | PA2856 | PA2856 | N/A | PA2856 | PA2856 | N/A | N/A |
| PA2857 | N/A | N/A | N/A | PA2857 | N/A | N/A | PA2857 | N/A | PA2857 | PA2857 | PA2857 | PA2857 | PA2857 | PA2857 | N/A | N/A | PA2857 | N/A |
| PA2858 | N/A | N/A | N/A | PA2858 | PA2858 | N/A | N/A | PA2858 | PA2858 | PA2858 | PA2858 | PA2858 | PA2858 | PA2858 | N/A | N/A | PA2858 | N/A |
| PA2859 | PA2859 | N/A | PA2859 | PA2859 | N/A | N/A | N/A | N/A | PA2859 | PA2859 | PA2859 | PA2859 | PA2859 | PA2859 | N/A | N/A | N/A | N/A |
| N/A | N/A | N/A | N/A | N/A | N/A | N/A | N/A | N/A | N/A | N/A | N/A | N/A | N/A | N/A | N/A | PA2862 | N/A | N/A |
| PA2864 | PA2864 | N/A | PA2864 | PA2864 | N/A | PA2864 | N/A | N/A | PA2864 | PA2864 | PA2864 | PA2864 | PA2864 | PA2864 | N/A | N/A | PA2864 | N/A |
| N/A | N/A | PA2866 | N/A | PA2866 | N/A | N/A | PA2866 | N/A | PA2866 | PA2866 | PA2866 | N/A | N/A | N/A | N/A | N/A | N/A | N/A |
| PA2867 | PA2867 | PA2867 | PA2867 | PA2867 | N/A | PA2867 | PA2867 | PA2867 | PA2867 | PA2867 | PA2867 | PA2867 | PA2867 | PA2867 | PA2867 | N/A | PA2867 | PA2867 |
| PA2868 | N/A | N/A | N/A | N/A | N/A | N/A | N/A | N/A | PA2868 | PA2868 | PA2868 | PA2868 | PA2868 | PA2868 | N/A | N/A | N/A | N/A |
| PA2870 | N/A | N/A | N/A | N/A | PA2870 | N/A | N/A | N/A | N/A | N/A | N/A | N/A | N/A | N/A | N/A | N/A | N/A | N/A |
| PA2871 | N/A | PA2871 | N/A | PA2871 | PA2871 | N/A | N/A | N/A | PA2871 | PA2871 | PA2871 | PA2871 | PA2871 | N/A | N/A | N/A | N/A | N/A |
| PA2872 | N/A | N/A | N/A | N/A | N/A | PA2872 | N/A | N/A | N/A | PA2872 | PA2872 | PA2872 | PA2872 | N/A | N/A | N/A | N/A | N/A |
| N/A | N/A | N/A | N/A | PA2873 | N/A | N/A | N/A | N/A | N/A | N/A | N/A | N/A | N/A | N/A | N/A | N/A | N/A | N/A |
| PA2874 | N/A | N/A | N/A | N/A | N/A | N/A | N/A | N/A | N/A | N/A | N/A | N/A | N/A | N/A | N/A | N/A | N/A | N/A |
| PA2875 | PA2875 | PA2875 | N/A | PA2875 | N/A | N/A | PA2875 | PA2875 | PA2875 | PA2875 | PA2875 | PA2875 | PA2875 | N/A | N/A | N/A | N/A | N/A |
| PA2876 | PA2876 | PA2876 | PA2876 | PA2876 | PA2876 | PA2876 | N/A | N/A | PA2876 | PA2876 | PA2876 | PA2876 | PA2876 | N/A | PA2876 | N/A | N/A | N/A |
| PA2877 | N/A | N/A | N/A | PA2877 | N/A | N/A | N/A | N/A | N/A | PA2877 | PA2877 | PA2877 | PA2877 | N/A | N/A | N/A | N/A | N/A |
| PA2883 | N/A | N/A | PA2883 | N/A | N/A | PA2883 | N/A | N/A | PA2883 | PA2883 | PA2883 | PA2883 | PA2883 | PA2883 | N/A | N/A | PA2883 | N/A |
| PA2885 | PA2885 | PA2885 | N/A | PA2885 | N/A | N/A | N/A | N/A | PA2885 | PA2885 | PA2885 | PA2885 | PA2885 | N/A | N/A | N/A | N/A | N/A |
| PA2887 | N/A | N/A | N/A | N/A | N/A | PA2887 | N/A | N/A | N/A | N/A | N/A | N/A | N/A | N/A | N/A | N/A | N/A | N/A |
| N/A | N/A | N/A | N/A | N/A | N/A | N/A | N/A | N/A | N/A | N/A | N/A | PA2888 | PA2888 | N/A | N/A | N/A | N/A | N/A |
| PA2889 | N/A | N/A | N/A | N/A | N/A | PA2889 | N/A | N/A | N/A | N/A | N/A | PA2889 | PA2889 | N/A | N/A | N/A | N/A | N/A |
| PA2890 | N/A | N/A | N/A | N/A | N/A | N/A | N/A | N/A | N/A | N/A | N/A | N/A | N/A | N/A | N/A | N/A | N/A | N/A |
| PA2891 | N/A | PA2891 | N/A | N/A | N/A | PA2891 | N/A | N/A | PA2891 | PA2891 | PA2891 | PA2891 | PA2891 | N/A | N/A | N/A | N/A | PA2891 |
| N/A | N/A | N/A | N/A | N/A | N/A | N/A | N/A | N/A | N/A | N/A | N/A | N/A | PA2893 | N/A | N/A | N/A | N/A | N/A |
| PA2894 | N/A | N/A | N/A | N/A | N/A | N/A | N/A | N/A | N/A | N/A | N/A | N/A | N/A | N/A | N/A | N/A | N/A | N/A |
| PA2895 | N/A | N/A | PA2895 | PA2895 | N/A | N/A | PA2895 | N/A | PA2895 | N/A | N/A | N/A | N/A | PA2895 | N/A | N/A | PA2895 | PA2895 |
| PA2896 | N/A | N/A | N/A | PA2896 | N/A | N/A | N/A | N/A | N/A | N/A | N/A | N/A | N/A | PA2896 | N/A | N/A | PA2896 | N/A |
| PA2897 | N/A | N/A | N/A | PA2897 | N/A | N/A | N/A | PA2897 | PA2897 | N/A | PA2897 | PA2897 | PA2897 | PA2897 | N/A | N/A | N/A | N/A |
| PA2899 | N/A | N/A | N/A | N/A | N/A | N/A | N/A | N/A | N/A | N/A | N/A | N/A | N/A | N/A | N/A | N/A | N/A | N/A |
| PA2900 | PA2900 | PA2900 | PA2900 | PA2900 | N/A | PA2900 | PA2900 | N/A | PA2900 | PA2900 | PA2900 | PA2900 | PA2900 | PA2900 | PA2900 | N/A | N/A | N/A |
| PA2901 | PA2901 | PA2901 | PA2901 | PA2901 | N/A | PA2901 | PA2901 | N/A | PA2901 | PA2901 | PA2901 | PA2901 | PA2901 | PA2901 | PA2901 | N/A | N/A | N/A |
| PA2902 | N/A | N/A | N/A | PA2902 | N/A | N/A | PA2902 | N/A | N/A | N/A | PA2902 | N/A | PA2902 | N/A | N/A | N/A | N/A | N/A |
| PA2903 | PA2903 | N/A | N/A | PA2903 | N/A | N/A | N/A | PA2903 | PA2903 | PA2903 | N/A | N/A | N/A | N/A | N/A | N/A | N/A | N/A |
| PA2904 | PA2904 | N/A | PA2904 | PA2904 | N/A | N/A | PA2904 | N/A | PA2904 | PA2904 | PA2904 | PA2904 | N/A | N/A | N/A | N/A | N/A | N/A |
| PA2905 | N/A | PA2905 | N/A | PA2905 | PA2905 | N/A | N/A | N/A | PA2905 | N/A | PA2905 | PA2905 | N/A | N/A | N/A | N/A | N/A | N/A |
| N/A | N/A | N/A | N/A | PA2906 | N/A | N/A | N/A | PA2906 | N/A | N/A | N/A | N/A | N/A | N/A | N/A | N/A | N/A | N/A |
| PA2907 | N/A | N/A | N/A | PA2907 | N/A | N/A | N/A | PA2907 | N/A | N/A | N/A | N/A | N/A | N/A | N/A | N/A | N/A | N/A |
| PA2911 | N/A | N/A | N/A | PA2911 | N/A | PA2911 | N/A | PA2911 | PA2911 | N/A | N/A | N/A | N/A | N/A | PA2911 | N/A | N/A | N/A |
| PA2912 | N/A | N/A | N/A | N/A | PA2912 | N/A | N/A | N/A | N/A | N/A | N/A | N/A | N/A | N/A | N/A | N/A | N/A | N/A |
| PA2913 | N/A | N/A | N/A | PA2913 | N/A | N/A | N/A | N/A | PA2913 | N/A | N/A | N/A | N/A | N/A | N/A | N/A | N/A | N/A |
| PA2915 | N/A | N/A | N/A | PA2915 | N/A | PA2915 | N/A | N/A | N/A | PA2915 | PA2915 | N/A | N/A | N/A | N/A | N/A | N/A | N/A |
| N/A | N/A | N/A | N/A | N/A | N/A | N/A | N/A | N/A | N/A | PA2916 | N/A | N/A | N/A | PA2916 | N/A | N/A | N/A | N/A |
| PA2918 | N/A | N/A | N/A | PA2918 | N/A | N/A | N/A | N/A | N/A | N/A | N/A | N/A | PA2918 | N/A | N/A | N/A | N/A | N/A |
| PA2920 | PA2920 | PA2920 | PA2920 | PA2920 | N/A | PA2920 | PA2920 | PA2920 | PA2920 | PA2920 | PA2920 | PA2920 | PA2920 | PA2920 | PA2920 | N/A | PA2920 | PA2920 |
| PA2926 | N/A | N/A | N/A | N/A | PA2926 | N/A | PA2926 | N/A | N/A | N/A | N/A | N/A | N/A | N/A | N/A | N/A | N/A | N/A |
| PA2927 | N/A | N/A | N/A | N/A | PA2927 | N/A | N/A | N/A | N/A | N/A | N/A | N/A | PA2927 | N/A | N/A | N/A | N/A | N/A |
| PA2931 | N/A | N/A | N/A | N/A | N/A | N/A | N/A | N/A | N/A | N/A | N/A | N/A | N/A | N/A | N/A | N/A | N/A | N/A |
| PA2932 | N/A | N/A | N/A | PA2932 | N/A | N/A | N/A | N/A | N/A | N/A | N/A | N/A | N/A | N/A | N/A | N/A | N/A | N/A |
| PA2937 | N/A | N/A | N/A | N/A | N/A | N/A | N/A | N/A | N/A | N/A | N/A | N/A | N/A | N/A | N/A | N/A | N/A | N/A |
| N/A | N/A | N/A | N/A | N/A | PA2938 | N/A | N/A | N/A | N/A | N/A | N/A | N/A | N/A | N/A | N/A | N/A | N/A | N/A |
| PA2939 | N/A | N/A | N/A | N/A | PA2939 | PA2939 | PA2939 | N/A | PA2939 | PA2939 | PA2939 | PA2939 | PA2939 | PA2939 | N/A | PA2939 | PA2939 | N/A |
| PA2941 | N/A | N/A | N/A | N/A | N/A | N/A | N/A | N/A | N/A | N/A | N/A | N/A | N/A | N/A | N/A | N/A | N/A | N/A |
| PA2942 | N/A | N/A | N/A | PA2942 | N/A | N/A | N/A | PA2942 | N/A | N/A | N/A | N/A | N/A | N/A | N/A | N/A | N/A | N/A |
| PA2944 | N/A | N/A | N/A | PA2944 | PA2944 | N/A | N/A | N/A | PA2944 | PA2944 | PA2944 | PA2944 | PA2944 | N/A | N/A | N/A | N/A | N/A |
| PA2945 | PA2945 | PA2945 | PA2945 | PA2945 | PA2945 | N/A | PA2945 | PA2945 | PA2945 | PA2945 | PA2945 | PA2945 | PA2945 | N/A | PA2945 | N/A | N/A | N/A |
| PA2946 | N/A | N/A | N/A | N/A | N/A | N/A | N/A | N/A | N/A | N/A | N/A | N/A | N/A | N/A | N/A | N/A | N/A | N/A |
| PA2947 | N/A | N/A | N/A | N/A | N/A | N/A | N/A | N/A | N/A | N/A | N/A | N/A | N/A | N/A | N/A | N/A | N/A | N/A |
| PA2948 | N/A | N/A | N/A | N/A | N/A | N/A | N/A | N/A | PA2948 | N/A | PA2948 | N/A | N/A | N/A | N/A | N/A | N/A | N/A |
| PA2949 | N/A | N/A | N/A | PA2949 | PA2949 | N/A | N/A | N/A | N/A | PA2949 | PA2949 | PA2949 | PA2949 | PA2949 | N/A | N/A | PA2949 | N/A |
| PA2950 | PA2950 | PA2950 | PA2950 | PA2950 | PA2950 | PA2950 | PA2950 | PA2950 | PA2950 | PA2950 | PA2950 | PA2950 | PA2950 | N/A | PA2950 | PA2950 | N/A | N/A |
| PA2951 | PA2951 | PA2951 | PA2951 | PA2951 | PA2951 | PA2951 | PA2951 | PA2951 | PA2951 | PA2951 | PA2951 | PA2951 | PA2951 | PA2951 | PA2951 | N/A | PA2951 | PA2951 |
| PA2952 | PA2952 | PA2952 | PA2952 | PA2952 | PA2952 | PA2952 | PA2952 | PA2952 | PA2952 | PA2952 | PA2952 | PA2952 | PA2952 | PA2952 | PA2952 | N/A | PA2952 | PA2952 |
| PA2953 | PA2953 | PA2953 | PA2953 | PA2953 | PA2953 | PA2953 | PA2953 | PA2953 | PA2953 | PA2953 | PA2953 | PA2953 | PA2953 | PA2953 | PA2953 | N/A | PA2953 | PA2953 |
| PA2954 | N/A | N/A | N/A | PA2954 | N/A | N/A | N/A | N/A | PA2954 | PA2954 | PA2954 | PA2954 | PA2954 | N/A | N/A | N/A | N/A | N/A |
| PA2955 | N/A | N/A | N/A | PA2955 | N/A | PA2955 | N/A | N/A | PA2955 | PA2955 | N/A | N/A | N/A | N/A | N/A | N/A | N/A | N/A |
| PA2956 | N/A | N/A | N/A | N/A | PA2956 | N/A | N/A | N/A | N/A | N/A | N/A | N/A | PA2956 | N/A | N/A | N/A | N/A | N/A |
| PA2957 | PA2957 | PA2957 | PA2957 | PA2957 | PA2957 | PA2957 | N/A | N/A | PA2957 | PA2957 | PA2957 | PA2957 | PA2957 | PA2957 | PA2957 | N/A | N/A | N/A |
| PA2958 | N/A | N/A | N/A | PA2958 | PA2958 | N/A | N/A | N/A | N/A | PA2958 | N/A | N/A | N/A | N/A | N/A | N/A | N/A | N/A |
| PA2959 | PA2959 | PA2959 | PA2959 | PA2959 | PA2959 | PA2959 | N/A | N/A | PA2959 | PA2959 | PA2959 | PA2959 | PA2959 | N/A | N/A | PA2959 | N/A | N/A |
| PA2960 | PA2960 | N/A | N/A | PA2960 | PA2960 | N/A | PA2960 | N/A | PA2960 | PA2960 | PA2960 | PA2960 | PA2960 | N/A | PA2960 | N/A | PA2960 | N/A |
| PA2961 | PA2961 | PA2961 | N/A | PA2961 | PA2961 | N/A | PA2961 | N/A | PA2961 | N/A | PA2961 | PA2961 | N/A | N/A | PA2961 | N/A | N/A | N/A |
| PA2962 | N/A | PA2962 | N/A | PA2962 | PA2962 | N/A | PA2962 | N/A | PA2962 | PA2962 | PA2962 | PA2962 | PA2962 | N/A | N/A | N/A | N/A | N/A |
| PA2963 | N/A | N/A | N/A | PA2963 | N/A | N/A | N/A | PA2963 | PA2963 | N/A | N/A | N/A | N/A | PA2963 | N/A | N/A | PA2963 | N/A |
| PA2964 | N/A | N/A | N/A | PA2964 | PA2964 | N/A | N/A | N/A | N/A | N/A | N/A | N/A | N/A | N/A | N/A | N/A | N/A | N/A |
| PA2965 | PA2965 | PA2965 | PA2965 | PA2965 | PA2965 | PA2965 | PA2965 | N/A | PA2965 | PA2965 | PA2965 | PA2965 | PA2965 | PA2965 | PA2965 | N/A | N/A | N/A |
| PA2966 | PA2966 | PA2966 | PA2966 | PA2966 | PA2966 | PA2966 | PA2966 | PA2966 | PA2966 | PA2966 | PA2966 | PA2966 | PA2966 | PA2966 | N/A | PA2966 | PA2966 | N/A |
| PA2967 | PA2967 | PA2967 | PA2967 | PA2967 | PA2967 | PA2967 | PA2967 | PA2967 | PA2967 | PA2967 | PA2967 | PA2967 | PA2967 | PA2967 | PA2967 | PA2967 | PA2967 | PA2967 |
| PA2968 | PA2968 | PA2968 | PA2968 | PA2968 | PA2968 | PA2968 | PA2968 | N/A | PA2968 | PA2968 | PA2968 | PA2968 | PA2968 | N/A | PA2968 | N/A | N/A | N/A |
| PA2969 | N/A | N/A | N/A | PA2969 | PA2969 | N/A | PA2969 | N/A | PA2969 | N/A | PA2969 | N/A | N/A | PA2969 | N/A | N/A | N/A | N/A |
| PA2970 | PA2970 | PA2970 | PA2970 | PA2970 | PA2970 | PA2970 | PA2970 | PA2970 | PA2970 | PA2970 | PA2970 | PA2970 | PA2970 | PA2970 | N/A | PA2970 | PA2970 | PA2970 |
| PA2971 | N/A | PA2971 | PA2971 | PA2971 | PA2971 | N/A | N/A | N/A | PA2971 | PA2971 | PA2971 | PA2971 | PA2971 | N/A | PA2971 | N/A | N/A | N/A |
| PA2972 | PA2972 | PA2972 | PA2972 | PA2972 | PA2972 | N/A | N/A | N/A | PA2972 | PA2972 | PA2972 | PA2972 | PA2972 | N/A | N/A | N/A | N/A | N/A |
| PA2973 | PA2973 | PA2973 | PA2973 | PA2973 | N/A | PA2973 | PA2973 | PA2973 | PA2973 | PA2973 | PA2973 | PA2973 | PA2973 | PA2973 | PA2973 | N/A | PA2973 | PA2973 |
| N/A | N/A | PA2974 | PA2974 | PA2974 | N/A | N/A | PA2974 | N/A | PA2974 | N/A | N/A | N/A | N/A | N/A | N/A | N/A | N/A | N/A |
| PA2975 | PA2975 | PA2975 | PA2975 | PA2975 | N/A | PA2975 | N/A | PA2975 | PA2975 | PA2975 | PA2975 | PA2975 | PA2975 | PA2975 | PA2975 | N/A | PA2975 | N/A |
| PA2976 | PA2976 | PA2976 | PA2976 | PA2976 | N/A | PA2976 | PA2976 | PA2976 | PA2976 | PA2976 | PA2976 | PA2976 | PA2976 | PA2976 | PA2976 | PA2976 | PA2976 | N/A |
| PA2977 | N/A | N/A | PA2977 | PA2977 | PA2977 | N/A | N/A | N/A | PA2977 | N/A | N/A | N/A | PA2977 | N/A | N/A | N/A | N/A | N/A |
| PA2978 | N/A | PA2978 | N/A | PA2978 | N/A | N/A | N/A | N/A | PA2978 | N/A | PA2978 | PA2978 | PA2978 | N/A | N/A | N/A | PA2978 | N/A |
| PA2979 | PA2979 | PA2979 | PA2979 | PA2979 | N/A | N/A | N/A | N/A | PA2979 | PA2979 | PA2979 | PA2979 | PA2979 | N/A | PA2979 | N/A | N/A | N/A |
| PA2980 | PA2980 | PA2980 | PA2980 | PA2980 | N/A | PA2980 | N/A | N/A | PA2980 | N/A | PA2980 | PA2980 | PA2980 | PA2980 | N/A | N/A | N/A | N/A |
| PA2981 | N/A | N/A | N/A | PA2981 | N/A | N/A | N/A | N/A | N/A | PA2981 | N/A | N/A | N/A | PA2981 | N/A | N/A | N/A | N/A |
| PA2982 | PA2982 | N/A | N/A | PA2982 | N/A | N/A | PA2982 | PA2982 | PA2982 | PA2982 | PA2982 | PA2982 | N/A | PA2982 | PA2982 | N/A | PA2982 | N/A |
| PA2983 | PA2983 | PA2983 | PA2983 | PA2983 | N/A | PA2983 | PA2983 | PA2983 | PA2983 | PA2983 | PA2983 | PA2983 | PA2983 | PA2983 | PA2983 | N/A | PA2983 | N/A |
| PA2986 | N/A | PA2986 | N/A | PA2986 | N/A | N/A | N/A | N/A | N/A | N/A | N/A | PA2986 | N/A | PA2986 | PA2986 | N/A | PA2986 | N/A |
| PA2987 | PA2987 | PA2987 | PA2987 | PA2987 | N/A | N/A | PA2987 | N/A | PA2987 | PA2987 | PA2987 | PA2987 | PA2987 | PA2987 | PA2987 | N/A | N/A | N/A |
| PA2988 | N/A | PA2988 | PA2988 | PA2988 | N/A | PA2988 | N/A | PA2988 | N/A | PA2988 | N/A | PA2988 | PA2988 | PA2988 | N/A | N/A | PA2988 | N/A |
| PA2989 | N/A | N/A | N/A | N/A | N/A | PA2989 | PA2989 | N/A | N/A | PA2989 | PA2989 | N/A | PA2989 | PA2989 | N/A | N/A | N/A | N/A |
| PA2990 | N/A | N/A | N/A | PA2990 | PA2990 | PA2990 | PA2990 | PA2990 | PA2990 | PA2990 | PA2990 | PA2990 | PA2990 | PA2990 | N/A | N/A | N/A | N/A |
| PA2991 | PA2991 | PA2991 | PA2991 | PA2991 | PA2991 | PA2991 | PA2991 | PA2991 | PA2991 | PA2991 | PA2991 | PA2991 | PA2991 | PA2991 | PA2991 | PA2991 | N/A | N/A |
| PA2993 | N/A | N/A | N/A | PA2993 | N/A | N/A | PA2993 | N/A | PA2993 | PA2993 | PA2993 | PA2993 | N/A | PA2993 | N/A | N/A | PA2993 | N/A |
| PA2994 | PA2994 | PA2994 | PA2994 | PA2994 | N/A | PA2994 | PA2994 | PA2994 | PA2994 | PA2994 | PA2994 | PA2994 | PA2994 | PA2994 | N/A | N/A | PA2994 | N/A |
| PA2996 | N/A | N/A | PA2996 | PA2996 | N/A | PA2996 | N/A | N/A | N/A | PA2996 | N/A | PA2996 | PA2996 | PA2996 | N/A | N/A | N/A | N/A |
| PA2997 | PA2997 | PA2997 | PA2997 | PA2997 | N/A | PA2997 | PA2997 | PA2997 | PA2997 | PA2997 | PA2997 | PA2997 | PA2997 | PA2997 | PA2997 | N/A | PA2997 | N/A |
| PA2998 | PA2998 | PA2998 | PA2998 | PA2998 | PA2998 | PA2998 | N/A | N/A | PA2998 | PA2998 | PA2998 | PA2998 | PA2998 | PA2998 | N/A | N/A | PA2998 | N/A |

| | | | | | | | | | | | | | | | | | | |
|---|---|---|---|---|---|---|---|---|---|---|---|---|---|---|---|---|---|---|
| PA2999 | PA2999 | PA2999 | PA2999 | PA2999 | PA2999 | PA2999 | PA2999 | PA2999 | PA2999 | PA2999 | PA2999 | PA2999 | PA2999 | PA2999 | PA2999 | N/A | PA2999 | PA2999 |
| N/A | N/A | N/A | N/A | N/A | PA3000 | N/A | N/A | N/A | N/A | N/A | N/A | N/A | N/A | PA3000 | N/A | N/A | N/A | N/A |
| PA3001 | PA3001 | PA3001 | PA3001 | PA3001 | PA3001 | PA3001 | PA3001 | PA3001 | PA3001 | PA3001 | PA3001 | PA3001 | PA3001 | PA3001 | PA3001 | N/A | PA3001 | PA3001 |
| PA3002 | PA3002 | PA3002 | PA3002 | PA3002 | PA3002 | PA3002 | N/A | PA3002 | PA3002 | PA3002 | PA3002 | PA3002 | PA3002 | PA3002 | PA3002 | N/A | N/A | N/A |
| PA3003 | PA3003 | PA3003 | N/A | PA3003 | PA3003 | N/A | N/A | N/A | PA3003 | PA3003 | PA3003 | PA3003 | PA3003 | N/A | PA3003 | N/A | N/A | N/A |
| PA3004 | PA3004 | PA3004 | PA3004 | PA3004 | PA3004 | PA3004 | N/A | N/A | PA3004 | PA3004 | PA3004 | PA3004 | PA3004 | N/A | N/A | N/A | N/A | N/A |
| PA3005 | PA3005 | PA3005 | PA3005 | PA3005 | PA3005 | PA3005 | PA3005 | PA3005 | PA3005 | PA3005 | PA3005 | PA3005 | PA3005 | N/A | PA3005 | N/A | N/A | N/A |
| PA3006 | PA3006 | PA3006 | PA3006 | PA3006 | PA3006 | PA3006 | PA3006 | N/A | PA3006 | PA3006 | PA3006 | PA3006 | PA3006 | PA3006 | N/A | N/A | N/A | N/A |
| PA3007 | PA3007 | PA3007 | PA3007 | PA3007 | PA3007 | N/A | N/A | N/A | PA3007 | PA3007 | PA3007 | PA3007 | PA3007 | N/A | PA3007 | N/A | N/A | N/A |
| PA3009 | PA3009 | PA3009 | N/A | PA3009 | N/A | N/A | N/A | N/A | PA3009 | N/A | PA3009 | PA3009 | N/A | N/A | N/A | N/A | N/A | N/A |
| PA3010 | N/A | PA3010 | N/A | PA3010 | N/A | PA3010 | N/A | N/A | PA3010 | PA3010 | PA3010 | PA3010 | PA3010 | N/A | N/A | N/A | N/A | N/A |
| PA3011 | PA3011 | PA3011 | PA3011 | PA3011 | PA3011 | PA3011 | N/A | PA3011 | PA3011 | PA3011 | PA3011 | PA3011 | PA3011 | PA3011 | PA3011 | N/A | N/A | N/A |
| PA3012 | N/A | PA3012 | N/A | PA3012 | PA3012 | PA3012 | N/A | N/A | PA3012 | N/A | PA3012 | PA3012 | PA3012 | N/A | N/A | N/A | N/A | N/A |
| PA3013 | PA3013 | PA3013 | PA3013 | PA3013 | PA3013 | PA3013 | PA3013 | N/A | PA3013 | PA3013 | PA3013 | PA3013 | PA3013 | PA3013 | PA3013 | N/A | PA3013 | PA3013 |
| PA3014 | PA3014 | PA3014 | PA3014 | PA3014 | N/A | PA3014 | N/A | PA3014 | PA3014 | PA3014 | PA3014 | PA3014 | PA3014 | PA3014 | PA3014 | N/A | PA3014 | PA3014 |
| PA3015 | N/A | N/A | N/A | N/A | N/A | N/A | N/A | N/A | N/A | N/A | N/A | PA3015 | PA3015 | N/A | N/A | N/A | N/A | N/A |
| PA3017 | N/A | N/A | N/A | N/A | N/A | PA3017 | N/A | N/A | PA3017 | PA3017 | PA3017 | PA3017 | PA3017 | PA3017 | N/A | N/A | N/A | N/A |
| PA3018 | N/A | N/A | N/A | N/A | N/A | N/A | N/A | N/A | N/A | N/A | PA3018 | PA3018 | PA3018 | N/A | N/A | N/A | N/A | N/A |
| PA3019 | PA3019 | PA3019 | PA3019 | PA3019 | PA3019 | N/A | PA3019 | PA3019 | PA3019 | PA3019 | PA3019 | PA3019 | PA3019 | N/A | N/A | N/A | N/A | N/A |
| PA3020 | PA3020 | N/A | PA3020 | PA3020 | PA3020 | N/A | N/A | PA3020 | PA3020 | N/A | PA3020 | PA3020 | PA3020 | PA3020 | PA3020 | N/A | PA3020 | PA3020 |
| PA3021 | PA3021 | PA3021 | PA3021 | PA3021 | N/A | PA3021 | N/A | N/A | PA3021 | PA3021 | PA3021 | PA3021 | PA3021 | N/A | PA3021 | N/A | PA3021 | PA3021 |
| PA3022 | PA3022 | PA3022 | PA3022 | PA3022 | N/A | PA3022 | PA3022 | N/A | PA3022 | N/A | PA3022 | PA3022 | PA3022 | N/A | N/A | N/A | N/A | N/A |
| PA3023 | N/A | N/A | N/A | PA3023 | PA3023 | PA3023 | N/A | N/A | N/A | PA3023 | N/A | PA3023 | PA3023 | N/A | N/A | N/A | N/A | N/A |
| PA3024 | N/A | N/A | N/A | N/A | N/A | N/A | N/A | N/A | N/A | N/A | N/A | N/A | N/A | N/A | N/A | N/A | N/A | N/A |
| PA3025 | N/A | N/A | N/A | N/A | N/A | N/A | N/A | N/A | N/A | N/A | N/A | N/A | N/A | N/A | N/A | N/A | N/A | N/A |
| PA3026 | N/A | N/A | N/A | N/A | N/A | PA3026 | N/A | N/A | N/A | PA3026 | N/A | PA3026 | N/A | PA3026 | N/A | N/A | N/A | N/A |
| PA3027 | N/A | N/A | N/A | PA3027 | PA3027 | N/A | N/A | N/A | N/A | N/A | PA3027 | PA3027 | N/A | N/A | N/A | N/A | N/A | N/A |
| PA3028 | N/A | N/A | N/A | PA3028 | N/A | N/A | N/A | N/A | PA3028 | N/A | PA3028 | N/A | PA3028 | N/A | N/A | N/A | N/A | N/A |
| PA3029 | PA3029 | PA3029 | PA3029 | PA3029 | PA3029 | PA3029 | PA3029 | N/A | PA3029 | PA3029 | PA3029 | PA3029 | PA3029 | PA3029 | PA3029 | PA3029 | N/A | N/A |
| PA3030 | N/A | N/A | N/A | PA3030 | PA3030 | N/A | N/A | N/A | N/A | PA3030 | N/A | N/A | N/A | N/A | N/A | N/A | N/A | N/A |
| PA3031 | N/A | PA3031 | N/A | PA3031 | PA3031 | PA3031 | PA3031 | N/A | PA3031 | PA3031 | PA3031 | PA3031 | PA3031 | PA3031 | PA3031 | PA3031 | PA3031 | PA3031 |
| PA3032 | N/A | N/A | N/A | N/A | N/A | N/A | N/A | N/A | N/A | PA3032 | N/A | N/A | N/A | N/A | N/A | N/A | N/A | N/A |
| PA3034 | N/A | N/A | N/A | N/A | N/A | N/A | N/A | N/A | PA3034 | PA3034 | N/A | PA3034 | N/A | PA3034 | N/A | N/A | N/A | N/A |
| N/A | N/A | N/A | N/A | N/A | N/A | N/A | N/A | N/A | N/A | N/A | N/A | PA3035 | N/A | N/A | N/A | N/A | N/A | N/A |
| N/A | N/A | N/A | N/A | N/A | PA3037 | N/A | N/A | N/A | N/A | N/A | N/A | N/A | N/A | N/A | N/A | N/A | N/A | N/A |
| PA3038 | N/A | PA3038 | N/A | N/A | N/A | PA3038 | N/A | PA3038 | N/A | PA3038 | PA3038 | PA3038 | PA3038 | N/A | N/A | N/A | N/A | N/A |
| N/A | N/A | PA3039 | PA3039 | N/A | PA3039 | N/A | N/A | N/A | N/A | N/A | N/A | N/A | N/A | N/A | N/A | N/A | N/A | N/A |
| PA3040 | PA3040 | PA3040 | PA3040 | PA3040 | N/A | PA3040 | PA3040 | N/A | PA3040 | PA3040 | PA3040 | PA3040 | PA3040 | PA3040 | N/A | PA3040 | PA3040 | PA3040 |
| PA3041 | N/A | N/A | N/A | N/A | N/A | PA3041 | N/A | N/A | PA3041 | PA3041 | PA3041 | PA3041 | PA3041 | PA3041 | N/A | N/A | N/A | N/A |
| PA3042 | N/A | N/A | N/A | N/A | N/A | PA3042 | PA3042 | N/A | PA3042 | PA3042 | PA3042 | PA3042 | PA3042 | PA3042 | N/A | N/A | N/A | N/A |
| PA3043 | N/A | N/A | N/A | N/A | N/A | PA3043 | N/A | N/A | PA3043 | PA3043 | PA3043 | PA3043 | PA3043 | N/A | N/A | N/A | N/A | N/A |
| N/A | N/A | N/A | N/A | N/A | PA3045 | N/A | N/A | N/A | PA3045 | N/A | N/A | N/A | N/A | N/A | N/A | N/A | N/A | N/A |
| PA3046 | PA3046 | PA3046 | PA3046 | PA3046 | PA3046 | PA3046 | PA3046 | N/A | PA3046 | PA3046 | PA3046 | PA3046 | PA3046 | PA3046 | PA3046 | N/A | N/A | N/A |
| PA3047 | PA3047 | PA3047 | PA3047 | PA3047 | PA3047 | N/A | PA3047 | PA3047 | PA3047 | PA3047 | PA3047 | PA3047 | PA3047 | PA3047 | PA3047 | N/A | PA3047 | PA3047 |
| PA3048 | PA3048 | PA3048 | PA3048 | PA3048 | PA3048 | N/A | N/A | PA3048 | PA3048 | N/A | PA3048 | PA3048 | PA3048 | PA3048 | PA3048 | N/A | N/A | N/A |
| PA3049 | N/A | N/A | N/A | N/A | N/A | PA3049 | N/A | N/A | N/A | N/A | N/A | N/A | N/A | PA3049 | N/A | N/A | N/A | N/A |
| PA3050 | PA3050 | PA3050 | PA3050 | PA3050 | N/A | PA3050 | PA3050 | PA3050 | PA3050 | PA3050 | PA3050 | PA3050 | PA3050 | PA3050 | PA3050 | N/A | PA3050 | N/A |
| N/A | N/A | N/A | N/A | N/A | PA3051 | N/A | N/A | N/A | N/A | N/A | N/A | N/A | N/A | N/A | N/A | N/A | N/A | N/A |
| PA3052 | N/A | N/A | N/A | PA3052 | N/A | N/A | N/A | N/A | N/A | N/A | PA3052 | N/A | PA3052 | N/A | N/A | N/A | N/A | N/A |
| PA3053 | N/A | N/A | N/A | PA3053 | PA3053 | PA3053 | N/A | N/A | N/A | N/A | PA3053 | PA3053 | N/A | N/A | N/A | N/A | N/A | N/A |
| PA3054 | N/A | N/A | N/A | N/A | PA3054 | N/A | N/A | N/A | N/A | PA3054 | N/A | N/A | N/A | PA3054 | N/A | N/A | N/A | N/A |
| PA3055 | N/A | N/A | N/A | PA3055 | N/A | N/A | N/A | N/A | N/A | N/A | PA3055 | N/A | N/A | N/A | N/A | N/A | N/A | N/A |
| PA3056 | N/A | PA3056 | N/A | PA3056 | N/A | N/A | N/A | N/A | N/A | N/A | PA3056 | N/A | N/A | PA3056 | N/A | N/A | N/A | N/A |
| PA3061 | N/A | N/A | N/A | N/A | N/A | N/A | N/A | N/A | N/A | N/A | N/A | N/A | N/A | N/A | N/A | N/A | N/A | N/A |
| PA3062 | N/A | N/A | N/A | PA3062 | N/A | N/A | N/A | N/A | N/A | PA3062 | N/A | N/A | N/A | N/A | N/A | N/A | N/A | N/A |
| PA3063 | N/A | PA3063 | N/A | N/A | N/A | N/A | N/A | N/A | N/A | N/A | N/A | N/A | N/A | N/A | N/A | N/A | N/A | N/A |
| PA3066 | N/A | N/A | N/A | N/A | N/A | N/A | N/A | N/A | N/A | N/A | N/A | PA3066 | PA3066 | N/A | N/A | N/A | PA3066 | N/A |
| PA3067 | N/A | N/A | N/A | PA3067 | PA3067 | N/A | N/A | N/A | N/A | N/A | N/A | N/A | N/A | N/A | N/A | N/A | N/A | N/A |
| PA3068 | PA3068 | PA3068 | PA3068 | PA3068 | PA3068 | PA3068 | PA3068 | PA3068 | PA3068 | PA3068 | PA3068 | PA3068 | PA3068 | PA3068 | N/A | PA3068 | PA3068 | PA3068 |
| PA3069 | N/A | N/A | N/A | N/A | N/A | N/A | N/A | N/A | N/A | PA3069 | N/A | N/A | N/A | N/A | N/A | N/A | N/A | N/A |
| PA3070 | PA3070 | N/A | PA3070 | PA3070 | N/A | N/A | PA3070 | PA3070 | PA3070 | PA3070 | PA3070 | PA3070 | PA3070 | PA3070 | PA3070 | N/A | N/A | N/A |
| PA3071 | N/A | N/A | N/A | PA3071 | N/A | N/A | N/A | N/A | N/A | N/A | N/A | N/A | N/A | PA3071 | N/A | N/A | N/A | N/A |
| PA3072 | N/A | N/A | N/A | N/A | N/A | N/A | N/A | N/A | N/A | PA3072 | PA3072 | N/A | N/A | PA3072 | N/A | N/A | N/A | N/A |
| PA3073 | N/A | N/A | N/A | PA3073 | N/A | N/A | PA3073 | N/A | N/A | N/A | PA3073 | N/A | N/A | PA3073 | N/A | N/A | N/A | N/A |
| PA3074 | N/A | N/A | N/A | N/A | N/A | N/A | N/A | N/A | PA3074 | PA3074 | N/A | PA3074 | PA3074 | PA3074 | N/A | N/A | PA3074 | N/A |
| PA3075 | N/A | N/A | N/A | PA3075 | N/A | N/A | N/A | N/A | N/A | PA3075 | N/A | N/A | N/A | PA3075 | N/A | N/A | PA3075 | N/A |
| PA3076 | N/A | N/A | N/A | PA3076 | N/A | N/A | N/A | N/A | N/A | N/A | N/A | N/A | N/A | N/A | N/A | N/A | N/A | N/A |
| PA3077 | N/A | N/A | N/A | N/A | N/A | N/A | N/A | N/A | PA3077 | N/A | PA3077 | PA3077 | PA3077 | N/A | N/A | N/A | N/A | N/A |
| PA3078 | N/A | N/A | N/A | N/A | N/A | N/A | N/A | N/A | N/A | N/A | N/A | N/A | N/A | PA3078 | N/A | N/A | N/A | N/A |
| PA3079 | N/A | N/A | N/A | N/A | N/A | PA3079 | N/A | N/A | N/A | N/A | PA3079 | PA3079 | PA3079 | PA3079 | N/A | N/A | N/A | N/A |
| PA3080 | N/A | N/A | N/A | PA3080 | PA3080 | N/A | N/A | N/A | N/A | N/A | PA3080 | PA3080 | PA3080 | PA3080 | N/A | N/A | PA3080 | N/A |
| PA3081 | PA3081 | PA3081 | PA3081 | PA3081 | PA3081 | PA3081 | N/A | PA3081 | PA3081 | PA3081 | PA3081 | PA3081 | PA3081 | PA3081 | PA3081 | N/A | N/A | N/A |
| PA3082 | N/A | PA3082 | PA3082 | PA3082 | PA3082 | PA3082 | N/A | PA3082 | PA3082 | PA3082 | PA3082 | PA3082 | PA3082 | N/A | PA3082 | N/A | N/A | N/A |
| PA3083 | PA3083 | PA3083 | PA3083 | PA3083 | PA3083 | PA3083 | N/A | PA3083 | PA3083 | PA3083 | PA3083 | PA3083 | PA3083 | PA3083 | PA3083 | N/A | N/A | N/A |
| PA3084 | N/A | N/A | N/A | PA3084 | N/A | N/A | N/A | N/A | PA3084 | N/A | PA3084 | PA3084 | N/A | N/A | N/A | N/A | N/A | N/A |
| N/A | N/A | PA3085 | N/A | PA3085 | N/A | N/A | N/A | N/A | PA3085 | N/A | N/A | PA3085 | N/A | N/A | N/A | N/A | N/A | N/A |
| PA3086 | N/A | N/A | N/A | N/A | PA3086 | N/A | N/A | N/A | N/A | N/A | N/A | N/A | N/A | PA3086 | N/A | N/A | N/A | N/A |
| PA3087 | N/A | N/A | N/A | PA3087 | PA3087 | N/A | N/A | N/A | PA3087 | N/A | N/A | PA3087 | PA3087 | N/A | N/A | N/A | N/A | N/A |
| PA3088 | PA3088 | PA3088 | N/A | PA3088 | N/A | PA3088 | PA3088 | N/A | PA3088 | PA3088 | PA3088 | PA3088 | PA3088 | N/A | N/A | N/A | N/A | N/A |
| PA3089 | N/A | N/A | N/A | N/A | N/A | N/A | N/A | N/A | N/A | N/A | N/A | N/A | N/A | N/A | N/A | N/A | N/A | N/A |
| PA3091 | N/A | N/A | N/A | N/A | PA3091 | N/A | N/A | N/A | N/A | PA3091 | N/A | N/A | N/A | N/A | N/A | N/A | N/A | N/A |
| PA3092 | N/A | N/A | N/A | PA3092 | PA3092 | PA3092 | N/A | PA3092 | PA3092 | PA3092 | PA3092 | PA3092 | PA3092 | N/A | N/A | N/A | N/A | N/A |
| PA3093 | PA3093 | PA3093 | N/A | PA3093 | N/A | PA3093 | PA3093 | N/A | N/A | N/A | PA3093 | PA3093 | N/A | PA3093 | PA3093 | N/A | PA3093 | N/A |
| PA3095 | N/A | N/A | N/A | N/A | N/A | N/A | PA3095 | N/A | PA3095 | PA3095 | PA3095 | PA3095 | PA3095 | PA3095 | N/A | N/A | PA3095 | N/A |
| PA3096 | N/A | N/A | N/A | N/A | N/A | N/A | PA3096 | N/A | N/A | N/A | PA3096 | PA3096 | N/A | PA3096 | N/A | N/A | N/A | N/A |
| PA3097 | N/A | N/A | N/A | N/A | N/A | N/A | N/A | N/A | N/A | N/A | N/A | N/A | N/A | PA3097 | N/A | N/A | N/A | N/A |
| PA3098 | N/A | N/A | N/A | N/A | N/A | N/A | N/A | N/A | N/A | N/A | N/A | N/A | N/A | PA3098 | N/A | N/A | N/A | N/A |
| PA3099 | N/A | N/A | N/A | N/A | N/A | N/A | N/A | N/A | N/A | N/A | N/A | N/A | N/A | PA3099 | N/A | N/A | N/A | N/A |
| PA3100 | N/A | N/A | N/A | N/A | N/A | N/A | N/A | N/A | PA3100 | N/A | PA3100 | PA3100 | N/A | PA3100 | N/A | N/A | N/A | N/A |
| PA3101 | N/A | PA3101 | N/A | PA3101 | N/A | PA3101 | PA3101 | N/A | PA3101 | PA3101 | PA3101 | PA3101 | PA3101 | PA3101 | N/A | N/A | PA3101 | N/A |
| PA3102 | N/A | N/A | N/A | PA3102 | N/A | N/A | N/A | N/A | PA3102 | N/A | PA3102 | PA3102 | PA3102 | PA3102 | N/A | N/A | N/A | N/A |
| PA3103 | N/A | N/A | N/A | PA3103 | N/A | N/A | N/A | PA3103 | PA3103 | PA3103 | N/A | PA3103 | N/A | PA3103 | N/A | N/A | PA3103 | N/A |
| PA3104 | N/A | N/A | N/A | N/A | N/A | N/A | PA3104 | N/A | PA3104 | N/A | PA3104 | PA3104 | N/A | PA3104 | N/A | N/A | N/A | N/A |
| PA3105 | N/A | N/A | PA3105 | PA3105 | N/A | PA3105 | PA3105 | PA3105 | PA3105 | PA3105 | PA3105 | PA3105 | PA3105 | PA3105 | PA3105 | N/A | PA3105 | N/A |
| PA3106 | PA3106 | PA3106 | PA3106 | PA3106 | PA3106 | N/A | N/A | N/A | PA3106 | PA3106 | PA3106 | PA3106 | N/A | N/A | N/A | N/A | N/A | N/A |
| PA3107 | PA3107 | PA3107 | PA3107 | PA3107 | PA3107 | PA3107 | PA3107 | N/A | PA3107 | PA3107 | PA3107 | PA3107 | PA3107 | PA3107 | PA3107 | PA3107 | N/A | N/A |
| PA3108 | PA3108 | PA3108 | PA3108 | PA3108 | PA3108 | PA3108 | N/A | PA3108 | PA3108 | PA3108 | PA3108 | PA3108 | PA3108 | PA3108 | PA3108 | N/A | N/A | PA3108 |
| PA3110 | PA3110 | PA3110 | N/A | PA3110 | PA3110 | N/A | N/A | N/A | N/A | PA3110 | PA3110 | N/A | PA3110 | PA3110 | PA3110 | N/A | N/A | N/A |
| PA3111 | N/A | PA3111 | PA3111 | PA3111 | PA3111 | N/A | N/A | PA3111 | PA3111 | PA3111 | PA3111 | PA3111 | PA3111 | N/A | N/A | N/A | N/A | N/A |
| PA3112 | PA3112 | PA3112 | PA3112 | PA3112 | PA3112 | PA3112 | PA3112 | PA3112 | PA3112 | PA3112 | PA3112 | PA3112 | PA3112 | PA3112 | PA3112 | N/A | PA3112 | PA3112 |
| PA3113 | N/A | PA3113 | N/A | PA3113 | N/A | PA3113 | N/A | N/A | PA3113 | PA3113 | PA3113 | PA3113 | PA3113 | N/A | N/A | N/A | N/A | N/A |
| PA3114 | PA3114 | PA3114 | PA3114 | PA3114 | PA3114 | PA3114 | N/A | N/A | PA3114 | PA3114 | PA3114 | PA3114 | PA3114 | N/A | N/A | N/A | N/A | N/A |
| PA3115 | PA3115 | PA3115 | PA3115 | PA3115 | PA3115 | PA3115 | PA3115 | PA3115 | PA3115 | PA3115 | PA3115 | PA3115 | PA3115 | PA3115 | PA3115 | N/A | PA3115 | PA3115 |
| PA3116 | PA3116 | PA3116 | PA3116 | PA3116 | PA3116 | N/A | PA3116 | PA3116 | PA3116 | PA3116 | PA3116 | PA3116 | PA3116 | N/A | N/A | N/A | N/A | N/A |
| PA3117 | PA3117 | PA3117 | PA3117 | PA3117 | PA3117 | PA3117 | PA3117 | N/A | PA3117 | PA3117 | PA3117 | PA3117 | PA3117 | N/A | PA3117 | PA3117 | N/A | N/A |
| PA3118 | PA3118 | PA3118 | N/A | PA3118 | PA3118 | PA3118 | N/A | PA3118 | PA3118 | PA3118 | PA3118 | PA3118 | PA3118 | N/A | PA3118 | N/A | N/A | N/A |
| PA3119 | N/A | N/A | N/A | N/A | PA3119 | N/A | PA3119 | N/A | N/A | N/A | N/A | N/A | N/A | N/A | N/A | N/A | N/A | N/A |
| PA3120 | PA3120 | PA3120 | N/A | PA3120 | PA3120 | PA3120 | PA3120 | N/A | PA3120 | PA3120 | PA3120 | PA3120 | PA3120 | N/A | N/A | N/A | N/A | N/A |
| PA3121 | PA3121 | PA3121 | N/A | PA3121 | PA3121 | N/A | N/A | N/A | N/A | PA3121 | PA3121 | PA3121 | PA3121 | N/A | N/A | N/A | N/A | N/A |

| | | | | | | | | | | | | | | | | | | |
|---|---|---|---|---|---|---|---|---|---|---|---|---|---|---|---|---|---|---|
| PA3122 | N/A | N/A | N/A | PA3122 | PA3122 | N/A | N/A | N/A | N/A | PA3122 | N/A | N/A | N/A | N/A | N/A | N/A | N/A | N/A |
| PA3123 | N/A | N/A | N/A | PA3123 | PA3123 | PA3123 | N/A | N/A | PA3123 | PA3123 | PA3123 | PA3123 | PA3123 | N/A | N/A | N/A | N/A | N/A |
| PA3124 | N/A | N/A | N/A | N/A | N/A | N/A | N/A | N/A | N/A | N/A | N/A | N/A | N/A | N/A | N/A | N/A | N/A | N/A |
| N/A | N/A | N/A | N/A | N/A | PA3125 | N/A | N/A | N/A | N/A | N/A | N/A | N/A | N/A | N/A | N/A | N/A | N/A | N/A |
| PA3126 | PA3126 | PA3126 | PA3126 | PA3126 | PA3126 | PA3126 | PA3126 | N/A | PA3126 | PA3126 | PA3126 | PA3126 | PA3126 | PA3126 | PA3126 | N/A | PA3126 | PA3126 |
| PA3127 | N/A | N/A | N/A | PA3127 | N/A | N/A | N/A | N/A | PA3127 | PA3127 | PA3127 | N/A | PA3127 | N/A | N/A | N/A | N/A | N/A |
| PA3128 | N/A | N/A | N/A | PA3128 | N/A | PA3128 | N/A | N/A | N/A | N/A | N/A | N/A | PA3128 | N/A | N/A | N/A | N/A | N/A |
| N/A | N/A | N/A | N/A | N/A | PA3129 | N/A | N/A | N/A | N/A | N/A | N/A | N/A | N/A | N/A | N/A | N/A | N/A | N/A |
| PA3130 | PA3130 | N/A | N/A | PA3130 | PA3130 | PA3130 | N/A | N/A | PA3130 | PA3130 | PA3130 | PA3130 | PA3130 | N/A | N/A | N/A | N/A | N/A |
| PA3131 | PA3131 | PA3131 | PA3131 | PA3131 | PA3131 | PA3131 | PA3131 | N/A | PA3131 | PA3131 | PA3131 | PA3131 | PA3131 | N/A | PA3131 | N/A | N/A | N/A |
| PA3133 | N/A | N/A | N/A | PA3133 | PA3133 | N/A | PA3133 | N/A | PA3133 | N/A | PA3133 | PA3133 | PA3133 | N/A | N/A | N/A | N/A | N/A |
| PA3134 | PA3134 | PA3134 | PA3134 | PA3134 | PA3134 | PA3134 | PA3134 | PA3134 | PA3134 | PA3134 | PA3134 | PA3134 | PA3134 | PA3134 | PA3134 | PA3134 | N/A | N/A |
| PA3135 | N/A | N/A | N/A | N/A | N/A | N/A | N/A | N/A | N/A | N/A | PA3135 | N/A | N/A | N/A | N/A | N/A | N/A | N/A |
| PA3138 | PA3138 | PA3138 | PA3138 | PA3138 | PA3138 | PA3138 | N/A | PA3138 | PA3138 | PA3138 | PA3138 | PA3138 | PA3138 | N/A | N/A | N/A | N/A | N/A |
| PA3139 | PA3139 | PA3139 | PA3139 | PA3139 | PA3139 | PA3139 | N/A | PA3139 | PA3139 | PA3139 | PA3139 | PA3139 | PA3139 | PA3139 | PA3139 | N/A | N/A | N/A |
| PA3141 | PA3141 | PA3141 | PA3141 | PA3141 | PA3141 | PA3141 | N/A | PA3141 | PA3141 | PA3141 | PA3141 | PA3141 | PA3141 | PA3141 | PA3141 | N/A | PA3141 | N/A |
| PA3145 | N/A | N/A | N/A | PA3145 | N/A | N/A | N/A | N/A | N/A | N/A | N/A | N/A | N/A | PA3145 | N/A | N/A | N/A | N/A |
| PA3146 | N/A | PA3146 | PA3146 | PA3146 | N/A | N/A | PA3146 | PA3146 | PA3146 | PA3146 | PA3146 | PA3146 | PA3146 | PA3146 | N/A | N/A | PA3146 | N/A |
| PA3147 | PA3147 | PA3147 | N/A | PA3147 | PA3147 | N/A | N/A | PA3147 | PA3147 | PA3147 | PA3147 | PA3147 | PA3147 | PA3147 | PA3147 | N/A | PA3147 | N/A |
| PA3148 | PA3148 | PA3148 | PA3148 | PA3148 | N/A | PA3148 | PA3148 | PA3148 | PA3148 | PA3148 | PA3148 | PA3148 | PA3148 | PA3148 | PA3148 | N/A | PA3148 | N/A |
| PA3149 | PA3149 | PA3149 | N/A | PA3149 | N/A | N/A | N/A | PA3149 | PA3149 | PA3149 | PA3149 | PA3149 | PA3149 | PA3149 | N/A | N/A | PA3149 | N/A |
| PA3150 | PA3150 | PA3150 | PA3150 | PA3150 | PA3150 | PA3150 | PA3150 | PA3150 | PA3150 | PA3150 | PA3150 | PA3150 | PA3150 | PA3150 | PA3150 | N/A | PA3150 | PA3150 |
| PA3151 | PA3151 | PA3151 | PA3151 | PA3151 | N/A | PA3151 | N/A | PA3151 | PA3151 | PA3151 | PA3151 | PA3151 | PA3151 | N/A | N/A | N/A | N/A | N/A |
| PA3152 | PA3152 | PA3152 | PA3152 | PA3152 | N/A | N/A | PA3152 | N/A | PA3152 | PA3152 | PA3152 | PA3152 | PA3152 | PA3152 | PA3152 | N/A | PA3152 | N/A |
| PA3153 | PA3153 | N/A | N/A | N/A | N/A | N/A | N/A | PA3153 | PA3153 | N/A | PA3153 | PA3153 | PA3153 | PA3153 | N/A | N/A | PA3153 | N/A |
| PA3154 | N/A | N/A | N/A | N/A | PA3154 | N/A | N/A | PA3154 | PA3154 | PA3154 | PA3154 | PA3154 | PA3154 | PA3154 | N/A | N/A | PA3154 | N/A |
| PA3155 | PA3155 | PA3155 | PA3155 | PA3155 | N/A | PA3155 | PA3155 | PA3155 | PA3155 | PA3155 | PA3155 | PA3155 | PA3155 | N/A | PA3155 | N/A | PA3155 | N/A |
| PA3156 | PA3156 | PA3156 | PA3156 | PA3156 | N/A | PA3156 | PA3156 | PA3156 | PA3156 | PA3156 | PA3156 | PA3156 | PA3156 | PA3156 | N/A | N/A | PA3156 | N/A |
| N/A | N/A | N/A | N/A | PA3157 | PA3157 | N/A | N/A | N/A | N/A | N/A | N/A | N/A | N/A | N/A | N/A | N/A | N/A | N/A |
| PA3158 | PA3158 | PA3158 | PA3158 | PA3158 | PA3158 | PA3158 | PA3158 | PA3158 | PA3158 | PA3158 | PA3158 | PA3158 | PA3158 | PA3158 | N/A | N/A | PA3158 | N/A |
| PA3159 | PA3159 | PA3159 | PA3159 | PA3159 | N/A | PA3159 | PA3159 | PA3159 | PA3159 | PA3159 | PA3159 | PA3159 | PA3159 | PA3159 | PA3159 | PA3159 | PA3159 | PA3159 |
| PA3160 | PA3160 | PA3160 | PA3160 | PA3160 | N/A | PA3160 | PA3160 | PA3160 | PA3160 | PA3160 | PA3160 | PA3160 | PA3160 | PA3160 | PA3160 | N/A | PA3160 | PA3160 |
| PA3161 | N/A | N/A | N/A | PA3161 | PA3161 | PA3161 | PA3161 | N/A | PA3161 | PA3161 | PA3161 | PA3161 | PA3161 | N/A | N/A | N/A | N/A | N/A |
| PA3162 | PA3162 | PA3162 | PA3162 | PA3162 | PA3162 | PA3162 | PA3162 | PA3162 | PA3162 | PA3162 | PA3162 | PA3162 | PA3162 | PA3162 | PA3162 | PA3162 | PA3162 | PA3162 |
| PA3163 | PA3163 | PA3163 | PA3163 | PA3163 | PA3163 | N/A | PA3163 | N/A | PA3163 | PA3163 | PA3163 | PA3163 | PA3163 | N/A | PA3163 | N/A | N/A | N/A |
| PA3165 | PA3165 | PA3165 | PA3165 | PA3165 | PA3165 | N/A | PA3165 | PA3165 | PA3165 | PA3165 | PA3165 | PA3165 | PA3165 | N/A | N/A | N/A | N/A | N/A |
| PA3166 | PA3166 | PA3166 | PA3166 | PA3166 | PA3166 | PA3166 | PA3166 | PA3166 | PA3166 | PA3166 | PA3166 | PA3166 | PA3166 | PA3166 | PA3166 | N/A | N/A | N/A |
| PA3167 | PA3167 | PA3167 | PA3167 | PA3167 | PA3167 | PA3167 | PA3167 | PA3167 | PA3167 | PA3167 | PA3167 | PA3167 | PA3167 | N/A | N/A | N/A | PA3167 | N/A |
| PA3168 | PA3168 | PA3168 | PA3168 | PA3168 | PA3168 | PA3168 | N/A | PA3168 | PA3168 | PA3168 | PA3168 | PA3168 | PA3168 | PA3168 | PA3168 | PA3168 | N/A | N/A |
| PA3169 | PA3169 | PA3169 | PA3169 | PA3169 | PA3169 | PA3169 | N/A | PA3169 | PA3169 | PA3169 | PA3169 | PA3169 | PA3169 | N/A | PA3169 | N/A | N/A | N/A |
| PA3170 | PA3170 | N/A | PA3170 | PA3170 | PA3170 | PA3170 | N/A | N/A | PA3170 | PA3170 | PA3170 | PA3170 | PA3170 | N/A | N/A | N/A | N/A | N/A |
| PA3171 | PA3171 | PA3171 | PA3171 | PA3171 | PA3171 | PA3171 | PA3171 | N/A | PA3171 | PA3171 | PA3171 | PA3171 | PA3171 | PA3171 | PA3171 | N/A | PA3171 | PA3171 |
| PA3172 | PA3172 | PA3172 | PA3172 | PA3172 | PA3172 | N/A | PA3172 | N/A | PA3172 | PA3172 | PA3172 | PA3172 | PA3172 | N/A | PA3172 | N/A | N/A | N/A |
| PA3173 | PA3173 | PA3173 | PA3173 | PA3173 | PA3173 | PA3173 | PA3173 | PA3173 | PA3173 | PA3173 | PA3173 | PA3173 | PA3173 | PA3173 | PA3173 | N/A | PA3173 | PA3173 |
| PA3177 | N/A | N/A | N/A | N/A | N/A | N/A | N/A | N/A | N/A | N/A | N/A | N/A | N/A | N/A | N/A | N/A | N/A | N/A |
| N/A | N/A | N/A | N/A | N/A | PA3178 | N/A | N/A | N/A | N/A | N/A | N/A | N/A | N/A | N/A | N/A | N/A | N/A | N/A |
| PA3179 | PA3179 | PA3179 | PA3179 | PA3179 | PA3179 | PA3179 | PA3179 | PA3179 | PA3179 | PA3179 | PA3179 | PA3179 | PA3179 | PA3179 | PA3179 | N/A | PA3179 | N/A |
| PA3180 | N/A | N/A | N/A | N/A | PA3180 | N/A | N/A | N/A | N/A | N/A | N/A | PA3180 | N/A | N/A | N/A | N/A | N/A | N/A |
| PA3181 | PA3181 | PA3181 | PA3181 | PA3181 | PA3181 | PA3181 | PA3181 | N/A | PA3181 | PA3181 | PA3181 | PA3181 | PA3181 | N/A | PA3181 | PA3181 | N/A | N/A |
| PA3182 | PA3182 | PA3182 | PA3182 | PA3182 | PA3182 | PA3182 | PA3182 | N/A | PA3182 | PA3182 | PA3182 | PA3182 | PA3182 | N/A | PA3182 | PA3182 | N/A | N/A |
| PA3183 | PA3183 | PA3183 | PA3183 | PA3183 | N/A | PA3183 | N/A | N/A | PA3183 | PA3183 | PA3183 | PA3183 | PA3183 | PA3183 | PA3183 | N/A | N/A | N/A |
| PA3184 | PA3184 | N/A | N/A | PA3184 | N/A | PA3184 | N/A | N/A | N/A | N/A | PA3184 | N/A | PA3184 | PA3184 | N/A | N/A | N/A | N/A |
| PA3185 | N/A | N/A | PA3185 | N/A | PA3185 | N/A | N/A | PA3185 | PA3185 | PA3185 | PA3185 | N/A | PA3185 | PA3185 | N/A | N/A | PA3185 | PA3185 |
| PA3186 | PA3186 | PA3186 | PA3186 | PA3186 | PA3186 | PA3186 | PA3186 | PA3186 | PA3186 | PA3186 | PA3186 | PA3186 | PA3186 | PA3186 | PA3186 | PA3186 | N/A | N/A |
| PA3187 | N/A | N/A | N/A | PA3187 | N/A | PA3187 | N/A | N/A | PA3187 | PA3187 | PA3187 | PA3187 | PA3187 | PA3187 | PA3187 | N/A | PA3187 | PA3187 |
| N/A | N/A | N/A | N/A | PA3188 | N/A | PA3188 | N/A | N/A | PA3188 | PA3188 | PA3188 | PA3188 | PA3188 | PA3188 | N/A | N/A | N/A | N/A |
| PA3189 | N/A | N/A | N/A | PA3189 | PA3189 | PA3189 | N/A | N/A | N/A | PA3189 | PA3189 | PA3189 | PA3189 | PA3189 | PA3189 | N/A | PA3189 | N/A |
| PA3190 | N/A | N/A | N/A | PA3190 | PA3190 | PA3190 | N/A | N/A | PA3190 | PA3190 | PA3190 | PA3190 | PA3190 | N/A | PA3190 | PA3190 | N/A | N/A |
| PA3191 | N/A | N/A | N/A | PA3191 | N/A | N/A | N/A | N/A | N/A | N/A | N/A | N/A | N/A | PA3191 | N/A | N/A | N/A | N/A |
| N/A | N/A | N/A | N/A | N/A | PA3192 | N/A | N/A | N/A | N/A | N/A | PA3192 | N/A | N/A | N/A | N/A | N/A | N/A | N/A |
| PA3193 | PA3193 | PA3193 | PA3193 | PA3193 | PA3193 | PA3193 | N/A | N/A | PA3193 | PA3193 | PA3193 | PA3193 | PA3193 | N/A | PA3193 | N/A | N/A | N/A |
| PA3194 | PA3194 | PA3194 | PA3194 | PA3194 | N/A | PA3194 | N/A | N/A | PA3194 | PA3194 | PA3194 | PA3194 | PA3194 | PA3194 | PA3194 | PA3194 | N/A | N/A |
| PA3195 | PA3195 | PA3195 | PA3195 | PA3195 | PA3195 | PA3195 | N/A | N/A | PA3195 | PA3195 | PA3195 | PA3195 | PA3195 | N/A | PA3195 | N/A | N/A | N/A |
| PA3196 | N/A | PA3196 | PA3196 | PA3196 | N/A | N/A | PA3196 | N/A | PA3196 | N/A | PA3196 | N/A | N/A | N/A | N/A | N/A | N/A | N/A |
| PA3197 | N/A | N/A | N/A | N/A | N/A | N/A | N/A | N/A | N/A | N/A | N/A | N/A | N/A | N/A | N/A | N/A | N/A | N/A |
| PA3198 | N/A | N/A | N/A | N/A | PA3198 | N/A | N/A | N/A | N/A | N/A | N/A | N/A | N/A | N/A | N/A | N/A | N/A | N/A |
| PA3199 | PA3199 | N/A | N/A | PA3199 | PA3199 | N/A | N/A | N/A | PA3199 | PA3199 | PA3199 | PA3199 | PA3199 | N/A | N/A | N/A | N/A | N/A |
| PA3200 | N/A | N/A | N/A | PA3200 | N/A | N/A | N/A | N/A | N/A | N/A | N/A | N/A | N/A | N/A | N/A | N/A | N/A | N/A |
| PA3201 | N/A | N/A | N/A | N/A | PA3201 | N/A | N/A | N/A | N/A | N/A | N/A | N/A | N/A | PA3201 | N/A | N/A | N/A | N/A |
| PA3202 | PA3202 | PA3202 | N/A | N/A | PA3202 | N/A | N/A | N/A | N/A | N/A | PA3202 | PA3202 | N/A | N/A | N/A | N/A | N/A | N/A |
| PA3203 | PA3203 | N/A | N/A | PA3203 | N/A | PA3203 | PA3203 | N/A | PA3203 | N/A | PA3203 | PA3203 | N/A | PA3203 | N/A | N/A | PA3203 | PA3203 |
| PA3204 | N/A | N/A | N/A | N/A | N/A | N/A | N/A | N/A | N/A | N/A | N/A | N/A | N/A | N/A | N/A | N/A | N/A | N/A |
| PA3205 | N/A | PA3205 | N/A | PA3205 | N/A | N/A | N/A | N/A | N/A | PA3205 | N/A | PA3205 | N/A | N/A | N/A | N/A | N/A | N/A |
| PA3206 | N/A | N/A | N/A | N/A | N/A | N/A | N/A | N/A | N/A | N/A | N/A | N/A | N/A | PA3206 | N/A | N/A | N/A | N/A |
| N/A | N/A | N/A | N/A | N/A | PA3207 | N/A | N/A | N/A | N/A | N/A | N/A | N/A | N/A | N/A | N/A | N/A | N/A | N/A |
| PA3208 | PA3208 | PA3208 | PA3208 | PA3208 | PA3208 | PA3208 | N/A | N/A | PA3208 | PA3208 | PA3208 | PA3208 | PA3208 | N/A | PA3208 | N/A | N/A | N/A |
| PA3210 | N/A | N/A | N/A | N/A | N/A | N/A | N/A | PA3210 | N/A | N/A | N/A | N/A | N/A | PA3210 | N/A | N/A | N/A | N/A |
| PA3211 | N/A | N/A | N/A | PA3211 | N/A | N/A | N/A | N/A | PA3211 | PA3211 | PA3211 | PA3211 | PA3211 | PA3211 | N/A | N/A | PA3211 | N/A |
| PA3212 | N/A | N/A | N/A | PA3212 | N/A | N/A | PA3212 | N/A | PA3212 | PA3212 | PA3212 | PA3212 | PA3212 | PA3212 | N/A | N/A | PA3212 | N/A |
| PA3213 | PA3213 | N/A | N/A | PA3213 | N/A | N/A | PA3213 | N/A | N/A | PA3213 | PA3213 | PA3213 | PA3213 | PA3213 | N/A | N/A | PA3213 | N/A |
| PA3214 | N/A | N/A | N/A | PA3214 | N/A | N/A | PA3214 | N/A | PA3214 | PA3214 | PA3214 | PA3214 | PA3214 | PA3214 | PA3214 | N/A | PA3214 | N/A |
| PA3217 | N/A | N/A | N/A | PA3217 | N/A | N/A | N/A | N/A | N/A | PA3217 | N/A | N/A | N/A | PA3217 | N/A | N/A | N/A | N/A |
| N/A | N/A | N/A | N/A | N/A | PA3220 | N/A | N/A | N/A | N/A | N/A | N/A | N/A | N/A | N/A | N/A | N/A | N/A | N/A |
| PA3221 | N/A | N/A | N/A | PA3221 | PA3221 | PA3221 | N/A | N/A | PA3221 | N/A | PA3221 | PA3221 | PA3221 | N/A | N/A | N/A | N/A | N/A |
| PA3222 | N/A | N/A | N/A | N/A | N/A | N/A | N/A | N/A | N/A | N/A | N/A | N/A | N/A | PA3222 | N/A | N/A | N/A | N/A |
| PA3224 | N/A | N/A | N/A | N/A | PA3224 | N/A | N/A | N/A | PA3224 | PA3224 | PA3224 | PA3224 | PA3224 | N/A | PA3224 | N/A | N/A | N/A |
| PA3225 | PA3225 | PA3225 | N/A | PA3225 | N/A | PA3225 | N/A | N/A | PA3225 | PA3225 | PA3225 | PA3225 | PA3225 | PA3225 | N/A | N/A | PA3225 | N/A |
| PA3226 | N/A | N/A | N/A | PA3226 | PA3226 | N/A | N/A | N/A | N/A | N/A | N/A | N/A | N/A | PA3226 | N/A | N/A | PA3226 | N/A |
| PA3227 | PA3227 | PA3227 | PA3227 | PA3227 | PA3227 | N/A | PA3227 | N/A | PA3227 | PA3227 | PA3227 | PA3227 | PA3227 | N/A | PA3227 | PA3227 | N/A | N/A |
| PA3228 | N/A | N/A | N/A | PA3228 | N/A | PA3228 | N/A | N/A | N/A | PA3228 | PA3228 | PA3228 | PA3228 | PA3228 | N/A | N/A | PA3228 | N/A |
| PA3230 | N/A | N/A | N/A | N/A | N/A | N/A | N/A | PA3230 | N/A | N/A | N/A | N/A | N/A | N/A | N/A | N/A | N/A | N/A |
| PA3233 | N/A | N/A | N/A | N/A | N/A | N/A | N/A | N/A | N/A | N/A | N/A | N/A | N/A | N/A | N/A | N/A | PA3233 | N/A |
| PA3234 | N/A | N/A | N/A | N/A | N/A | PA3234 | N/A | N/A | N/A | N/A | PA3234 | PA3234 | PA3234 | PA3234 | N/A | N/A | N/A | N/A |
| PA3235 | N/A | N/A | N/A | N/A | N/A | PA3235 | N/A | N/A | N/A | N/A | PA3235 | PA3235 | N/A | N/A | N/A | N/A | N/A | N/A |
| PA3238 | N/A | N/A | N/A | PA3238 | N/A | PA3238 | N/A | PA3238 | N/A | N/A | N/A | N/A | PA3238 | N/A | PA3238 | N/A | N/A | N/A |
| PA3239 | N/A | N/A | N/A | PA3239 | PA3239 | PA3239 | PA3239 | PA3239 | PA3239 | PA3239 | PA3239 | PA3239 | PA3239 | N/A | N/A | N/A | N/A | N/A |
| PA3240 | N/A | N/A | N/A | PA3240 | PA3240 | N/A | N/A | N/A | N/A | PA3240 | PA3240 | N/A | N/A | N/A | N/A | N/A | N/A | N/A |
| PA3241 | N/A | N/A | N/A | PA3241 | N/A | N/A | N/A | N/A | N/A | N/A | N/A | N/A | N/A | PA3241 | N/A | N/A | N/A | N/A |
| PA3242 | N/A | N/A | N/A | PA3242 | PA3242 | PA3242 | N/A | N/A | PA3242 | PA3242 | PA3242 | PA3242 | N/A | PA3242 | PA3242 | N/A | N/A | N/A |
| PA3243 | PA3243 | PA3243 | PA3243 | PA3243 | PA3243 | PA3243 | PA3243 | PA3243 | PA3243 | PA3243 | PA3243 | PA3243 | PA3243 | PA3243 | PA3243 | N/A | PA3243 | PA3243 |
| PA3244 | PA3244 | PA3244 | PA3244 | PA3244 | PA3244 | PA3244 | PA3244 | PA3244 | PA3244 | PA3244 | PA3244 | PA3244 | PA3244 | PA3244 | PA3244 | N/A | PA3244 | N/A |
| PA3245 | PA3245 | PA3245 | PA3245 | PA3245 | PA3245 | PA3245 | PA3245 | N/A | PA3245 | PA3245 | PA3245 | PA3245 | PA3245 | PA3245 | PA3245 | N/A | PA3245 | N/A |
| PA3246 | PA3246 | N/A | N/A | PA3246 | N/A | N/A | N/A | N/A | PA3246 | N/A | N/A | N/A | N/A | N/A | N/A | N/A | N/A | N/A |
| PA3247 | PA3247 | N/A | N/A | PA3247 | PA3247 | PA3247 | N/A | N/A | PA3247 | PA3247 | PA3247 | PA3247 | PA3247 | PA3247 | N/A | PA3247 | N/A | N/A |
| PA3248 | N/A | N/A | N/A | N/A | N/A | N/A | N/A | N/A | N/A | N/A | N/A | N/A | N/A | N/A | N/A | N/A | N/A | N/A |
| PA3249 | N/A | N/A | N/A | N/A | PA3249 | N/A | N/A | N/A | N/A | N/A | N/A | N/A | PA3249 | N/A | N/A | N/A | N/A | N/A |
| PA3250 | N/A | N/A | N/A | N/A | N/A | PA3250 | N/A | N/A | PA3250 | PA3250 | PA3250 | PA3250 | PA3250 | N/A | N/A | PA3250 | N/A | N/A |
| PA3251 | N/A | N/A | N/A | N/A | N/A | PA3251 | N/A | N/A | N/A | PA3251 | N/A | PA3251 | PA3251 | N/A | N/A | N/A | N/A | N/A |
| PA3254 | N/A | N/A | N/A | N/A | PA3254 | N/A | N/A | N/A | N/A | N/A | N/A | N/A | N/A | N/A | N/A | N/A | N/A | N/A |

| | | | | | | | | | | | | | | | | | | |
|---|---|---|---|---|---|---|---|---|---|---|---|---|---|---|---|---|---|---|
| PA3255 | N/A | PA3255 | N/A | PA3255 | PA3255 | PA3255 | N/A | N/A | PA3255 | PA3255 | PA3255 | PA3255 | PA3255 | N/A | N/A | N/A | N/A | N/A |
| PA3256 | PA3256 | N/A | N/A | PA3256 | PA3256 | PA3256 | PA3256 | PA3256 | PA3256 | PA3256 | PA3256 | PA3256 | PA3256 | N/A | N/A | N/A | PA3256 | PA3256 |
| PA3257 | PA3257 | PA3257 | PA3257 | PA3257 | PA3257 | PA3257 | N/A | PA3257 | PA3257 | PA3257 | PA3257 | PA3257 | PA3257 | PA3257 | PA3257 | N/A | PA3257 | N/A |
| PA3259 | N/A | N/A | N/A | PA3259 | N/A | N/A | N/A | N/A | PA3259 | N/A | PA3259 | PA3259 | PA3259 | PA3259 | N/A | N/A | PA3259 | N/A |
| PA3259.1 | N/A | N/A | N/A | PA3259.1 | N/A | N/A | N/A | N/A | PA3259.1 | PA3259.1 | PA3259.1 | PA3259.1 | PA3259.1 | N/A | N/A | N/A | N/A | N/A |
| PA3260 | N/A | N/A | N/A | PA3260 | N/A | N/A | N/A | N/A | PA3260 | PA3260 | PA3260 | PA3260 | PA3260 | N/A | N/A | N/A | N/A | N/A |
| N/A | N/A | N/A | N/A | N/A | PA3261 | N/A | N/A | N/A | N/A | N/A | N/A | N/A | N/A | N/A | N/A | N/A | N/A | N/A |
| PA3262 | PA3262 | PA3262 | PA3262 | PA3262 | PA3262 | PA3262 | PA3262 | PA3262 | PA3262 | PA3262 | PA3262 | PA3262 | PA3262 | PA3262 | PA3262 | PA3262 | PA3262 | PA3262 |
| PA3263 | PA3263 | PA3263 | PA3263 | PA3263 | N/A | PA3263 | PA3263 | PA3263 | PA3263 | PA3263 | PA3263 | PA3263 | PA3263 | PA3263 | PA3263 | N/A | N/A | N/A |
| N/A | N/A | N/A | N/A | N/A | PA3265 | N/A | N/A | N/A | N/A | N/A | N/A | N/A | N/A | N/A | N/A | N/A | N/A | N/A |
| PA3266 | PA3266 | PA3266 | PA3266 | PA3266 | PA3266 | PA3266 | PA3266 | N/A | PA3266 | PA3266 | PA3266 | PA3266 | PA3266 | N/A | PA3266 | N/A | N/A | N/A |
| PA3267 | N/A | N/A | N/A | PA3267 | N/A | PA3267 | N/A | PA3267 | PA3267 | N/A | N/A | PA3267 | PA3267 | PA3267 | N/A | N/A | PA3267 | N/A |
| PA3268 | N/A | PA3268 | N/A | PA3268 | N/A | N/A | N/A | PA3268 | PA3268 | N/A | PA3268 | PA3268 | N/A | N/A | PA3268 | N/A | N/A | N/A |
| N/A | N/A | N/A | N/A | N/A | PA3269 | N/A | N/A | N/A | N/A | N/A | N/A | N/A | N/A | N/A | N/A | N/A | N/A | N/A |
| PA3270 | PA3270 | PA3270 | PA3270 | PA3270 | PA3270 | PA3270 | PA3270 | N/A | PA3270 | PA3270 | PA3270 | PA3270 | PA3270 | N/A | N/A | N/A | N/A | N/A |
| PA3271 | N/A | N/A | N/A | PA3271 | N/A | N/A | N/A | PA3271 | PA3271 | N/A | PA3271 | PA3271 | PA3271 | PA3271 | N/A | N/A | PA3271 | N/A |
| PA3272 | N/A | N/A | N/A | N/A | N/A | N/A | N/A | N/A | N/A | N/A | N/A | N/A | N/A | N/A | N/A | N/A | N/A | N/A |
| PA3273 | N/A | N/A | N/A | N/A | N/A | N/A | N/A | N/A | N/A | N/A | N/A | N/A | N/A | N/A | N/A | N/A | N/A | N/A |
| PA3276 | N/A | N/A | N/A | PA3276 | N/A | N/A | N/A | N/A | PA3276 | N/A | PA3276 | PA3276 | N/A | N/A | N/A | N/A | N/A | N/A |
| PA3277 | N/A | N/A | N/A | N/A | N/A | N/A | N/A | N/A | PA3277 | N/A | N/A | N/A | PA3277 | N/A | N/A | N/A | N/A | N/A |
| PA3281 | N/A | N/A | N/A | N/A | N/A | PA3281 | N/A | N/A | N/A | PA3281 | N/A | N/A | N/A | N/A | N/A | N/A | N/A | N/A |
| N/A | N/A | N/A | N/A | N/A | N/A | N/A | N/A | N/A | N/A | PA3282 | N/A | N/A | N/A | N/A | N/A | N/A | N/A | N/A |
| N/A | N/A | N/A | N/A | PA3283 | N/A | N/A | N/A | N/A | N/A | PA3283 | N/A | N/A | N/A | N/A | N/A | N/A | N/A | N/A |
| N/A | N/A | N/A | N/A | N/A | N/A | N/A | N/A | N/A | N/A | PA3284 | N/A | N/A | N/A | N/A | N/A | N/A | N/A | N/A |
| PA3285 | N/A | N/A | N/A | PA3285 | PA3285 | N/A | N/A | N/A | N/A | N/A | PA3285 | N/A | N/A | N/A | N/A | N/A | N/A | N/A |
| PA3286 | PA3286 | PA3286 | PA3286 | PA3286 | PA3286 | PA3286 | N/A | PA3286 | PA3286 | PA3286 | PA3286 | PA3286 | PA3286 | N/A | PA3286 | N/A | N/A | N/A |
| PA3287 | N/A | N/A | N/A | PA3287 | N/A | N/A | N/A | N/A | N/A | N/A | PA3287 | N/A | PA3287 | N/A | N/A | N/A | N/A | N/A |
| N/A | N/A | N/A | N/A | N/A | N/A | N/A | N/A | N/A | N/A | PA3288 | PA3288 | PA3288 | PA3288 | N/A | N/A | N/A | N/A | N/A |
| PA3289 | N/A | N/A | N/A | N/A | N/A | N/A | N/A | N/A | N/A | PA3289 | PA3289 | PA3289 | PA3289 | N/A | N/A | N/A | N/A | N/A |
| N/A | N/A | N/A | N/A | N/A | PA3294 | N/A | N/A | N/A | PA3294 | N/A | N/A | N/A | N/A | N/A | N/A | N/A | N/A | N/A |
| PA3295 | PA3295 | PA3295 | PA3295 | PA3295 | PA3295 | PA3295 | PA3295 | N/A | PA3295 | PA3295 | PA3295 | PA3295 | PA3295 | N/A | PA3295 | N/A | N/A | N/A |
| PA3297 | PA3297 | PA3297 | PA3297 | PA3297 | N/A | N/A | N/A | PA3297 | PA3297 | PA3297 | PA3297 | PA3297 | PA3297 | PA3297 | N/A | N/A | PA3297 | N/A |
| N/A | N/A | N/A | N/A | N/A | PA3298 | N/A | N/A | N/A | N/A | N/A | N/A | N/A | N/A | N/A | N/A | N/A | N/A | N/A |
| PA3299 | PA3299 | PA3299 | PA3299 | PA3299 | PA3299 | PA3299 | PA3299 | PA3299 | PA3299 | PA3299 | PA3299 | PA3299 | PA3299 | PA3299 | N/A | N/A | PA3299 | PA3299 |
| PA3300 | N/A | N/A | N/A | PA3300 | N/A | PA3300 | N/A | N/A | N/A | N/A | N/A | PA3300 | PA3300 | PA3300 | N/A | N/A | N/A | N/A |
| PA3301 | N/A | PA3301 | N/A | PA3301 | PA3301 | PA3301 | PA3301 | N/A | PA3301 | PA3301 | N/A | PA3301 | PA3301 | PA3301 | N/A | N/A | PA3301 | N/A |
| PA3302 | PA3302 | PA3302 | PA3302 | PA3302 | PA3302 | PA3302 | PA3302 | N/A | PA3302 | PA3302 | PA3302 | PA3302 | PA3302 | N/A | PA3302 | N/A | N/A | N/A |
| PA3304 | N/A | N/A | N/A | N/A | N/A | N/A | N/A | N/A | N/A | PA3304 | N/A | N/A | N/A | PA3304 | PA3304 | N/A | N/A | N/A |
| PA3305 | N/A | N/A | N/A | N/A | N/A | N/A | N/A | N/A | N/A | N/A | N/A | N/A | N/A | N/A | N/A | N/A | N/A | N/A |
| PA3306 | N/A | N/A | N/A | N/A | N/A | N/A | N/A | N/A | N/A | N/A | N/A | N/A | N/A | N/A | N/A | N/A | N/A | N/A |
| N/A | N/A | N/A | N/A | N/A | PA3307 | N/A | N/A | N/A | N/A | N/A | N/A | N/A | N/A | N/A | N/A | N/A | N/A | N/A |
| PA3308 | PA3308 | PA3308 | PA3308 | PA3308 | PA3308 | PA3308 | N/A | PA3308 | PA3308 | PA3308 | PA3308 | PA3308 | PA3308 | PA3308 | PA3308 | N/A | N/A | N/A |
| PA3309 | PA3309 | PA3309 | PA3309 | PA3309 | PA3309 | PA3309 | PA3309 | PA3309 | PA3309 | PA3309 | PA3309 | PA3309 | PA3309 | PA3309 | N/A | PA3309 | PA3309 | PA3309 |
| PA3310 | N/A | N/A | N/A | PA3310 | N/A | N/A | N/A | N/A | N/A | N/A | N/A | N/A | N/A | PA3310 | N/A | N/A | N/A | N/A |
| PA3311 | N/A | N/A | N/A | N/A | N/A | N/A | N/A | N/A | N/A | N/A | N/A | PA3311 | PA3311 | N/A | N/A | N/A | N/A | N/A |
| PA3312 | PA3312 | N/A | N/A | PA3312 | PA3312 | N/A | N/A | N/A | N/A | N/A | PA3312 | N/A | N/A | N/A | PA3312 | N/A | N/A | N/A |
| PA3313 | PA3313 | PA3313 | PA3313 | PA3313 | PA3313 | PA3313 | PA3313 | N/A | PA3313 | PA3313 | PA3313 | PA3313 | PA3313 | N/A | PA3313 | PA3313 | PA3313 | N/A |
| PA3314 | PA3314 | PA3314 | N/A | PA3314 | N/A | N/A | PA3314 | N/A | N/A | N/A | PA3314 | N/A | N/A | PA3314 | N/A | N/A | PA3314 | N/A |
| PA3315 | N/A | N/A | N/A | N/A | N/A | N/A | N/A | N/A | N/A | N/A | N/A | N/A | N/A | N/A | N/A | N/A | N/A | N/A |
| PA3316 | N/A | N/A | N/A | N/A | N/A | N/A | N/A | N/A | N/A | N/A | N/A | N/A | N/A | N/A | N/A | N/A | N/A | N/A |
| PA3317 | N/A | PA3317 | PA3317 | PA3317 | N/A | N/A | PA3317 | N/A | PA3317 | PA3317 | PA3317 | PA3317 | PA3317 | N/A | N/A | N/A | PA3317 | N/A |
| PA3322 | N/A | N/A | N/A | PA3322 | N/A | PA3322 | PA3322 | N/A | PA3322 | PA3322 | PA3322 | PA3322 | PA3322 | N/A | N/A | N/A | N/A | N/A |
| N/A | N/A | N/A | N/A | N/A | PA3325 | N/A | N/A | N/A | N/A | N/A | N/A | N/A | N/A | N/A | N/A | N/A | N/A | N/A |
| PA3326 | N/A | N/A | N/A | PA3326 | PA3326 | PA3326 | PA3326 | N/A | PA3326 | PA3326 | PA3326 | PA3326 | PA3326 | PA3326 | N/A | PA3326 | PA3326 | N/A |
| PA3327 | N/A | N/A | N/A | N/A | PA3327 | PA3327 | N/A | N/A | PA3327 | PA3327 | PA3327 | PA3327 | PA3327 | N/A | N/A | N/A | PA3327 | N/A |
| PA3328 | N/A | N/A | N/A | N/A | PA3328 | N/A | N/A | N/A | N/A | PA3328 | PA3328 | PA3328 | PA3328 | N/A | N/A | N/A | PA3328 | N/A |
| PA3329 | N/A | N/A | N/A | N/A | N/A | N/A | N/A | N/A | PA3329 | PA3329 | PA3329 | PA3329 | PA3329 | N/A | N/A | N/A | N/A | N/A |
| PA3330 | N/A | N/A | N/A | PA3330 | PA3330 | N/A | N/A | N/A | PA3330 | PA3330 | PA3330 | PA3330 | PA3330 | N/A | N/A | N/A | PA3330 | N/A |
| PA3331 | N/A | N/A | N/A | PA3331 | PA3331 | N/A | N/A | N/A | PA3331 | N/A | PA3331 | PA3331 | PA3331 | N/A | N/A | N/A | N/A | N/A |
| PA3332 | N/A | N/A | N/A | N/A | PA3332 | PA3332 | N/A | N/A | PA3332 | PA3332 | PA3332 | PA3332 | PA3332 | N/A | N/A | N/A | N/A | N/A |
| PA3333 | N/A | N/A | N/A | N/A | PA3333 | N/A | N/A | N/A | PA3333 | N/A | PA3333 | PA3333 | PA3333 | N/A | N/A | N/A | N/A | N/A |
| PA3335 | N/A | N/A | N/A | N/A | N/A | N/A | N/A | N/A | N/A | N/A | PA3335 | PA3335 | N/A | N/A | N/A | N/A | N/A | N/A |
| PA3336 | N/A | N/A | N/A | N/A | PA3336 | N/A | N/A | N/A | N/A | N/A | N/A | N/A | N/A | N/A | N/A | N/A | N/A | N/A |
| PA3337 | N/A | N/A | N/A | PA3337 | PA3337 | N/A | PA3337 | N/A | N/A | PA3337 | PA3337 | PA3337 | PA3337 | PA3337 | N/A | N/A | N/A | N/A |
| N/A | N/A | N/A | N/A | N/A | PA3338 | N/A | N/A | N/A | N/A | N/A | N/A | PA3338 | PA3338 | N/A | N/A | N/A | N/A | N/A |
| PA3339 | N/A | N/A | N/A | PA3339 | N/A | PA3339 | N/A | PA3339 | PA3339 | PA3339 | N/A | N/A | PA3339 | N/A | N/A | N/A | N/A | N/A |
| PA3340 | N/A | N/A | N/A | PA3340 | PA3340 | N/A | N/A | N/A | PA3340 | N/A | PA3340 | PA3340 | PA3340 | PA3340 | PA3340 | N/A | PA3340 | N/A |
| PA3341 | PA3341 | PA3341 | PA3341 | PA3341 | PA3341 | PA3341 | PA3341 | N/A | PA3341 | PA3341 | PA3341 | PA3341 | PA3341 | PA3341 | PA3341 | N/A | PA3341 | N/A |
| PA3342 | N/A | N/A | N/A | N/A | N/A | N/A | N/A | N/A | N/A | N/A | N/A | N/A | N/A | PA3342 | N/A | N/A | N/A | N/A |
| PA3343 | N/A | N/A | N/A | N/A | N/A | N/A | N/A | N/A | N/A | N/A | N/A | PA3343 | N/A | PA3343 | N/A | N/A | N/A | N/A |
| PA3344 | N/A | N/A | N/A | PA3344 | N/A | N/A | N/A | PA3344 | PA3344 | PA3344 | PA3344 | PA3344 | PA3344 | N/A | N/A | N/A | N/A | N/A |
| PA3345 | N/A | N/A | PA3345 | PA3345 | N/A | N/A | N/A | N/A | N/A | N/A | PA3345 | N/A | N/A | N/A | N/A | N/A | N/A | N/A |
| PA3346 | N/A | N/A | N/A | N/A | PA3346 | N/A | N/A | N/A | N/A | N/A | N/A | N/A | N/A | N/A | N/A | N/A | N/A | N/A |
| PA3347 | N/A | N/A | PA3347 | PA3347 | PA3347 | PA3347 | N/A | N/A | PA3347 | PA3347 | PA3347 | PA3347 | PA3347 | N/A | PA3347 | PA3347 | N/A | N/A |
| PA3348 | N/A | PA3348 | N/A | PA3348 | PA3348 | PA3348 | N/A | N/A | PA3348 | PA3348 | PA3348 | PA3348 | PA3348 | PA3348 | N/A | N/A | PA3348 | N/A |
| PA3349 | PA3349 | PA3349 | PA3349 | PA3349 | PA3349 | PA3349 | PA3349 | PA3349 | PA3349 | PA3349 | PA3349 | PA3349 | PA3349 | PA3349 | N/A | N/A | PA3349 | N/A |
| PA3350 | PA3350 | N/A | N/A | PA3350 | N/A | N/A | PA3350 | N/A | PA3350 | N/A | PA3350 | PA3350 | N/A | PA3350 | N/A | N/A | N/A | N/A |
| PA3351 | N/A | PA3351 | N/A | PA3351 | N/A | PA3351 | N/A | N/A | PA3351 | PA3351 | PA3351 | PA3351 | PA3351 | N/A | N/A | N/A | N/A | N/A |
| PA3352 | PA3352 | PA3352 | PA3352 | PA3352 | N/A | N/A | N/A | N/A | PA3352 | PA3352 | PA3352 | PA3352 | PA3352 | N/A | PA3352 | N/A | N/A | N/A |
| PA3353 | N/A | N/A | PA3353 | PA3353 | PA3353 | N/A | N/A | N/A | PA3353 | PA3353 | PA3353 | PA3353 | PA3353 | N/A | N/A | N/A | N/A | N/A |
| PA3354 | N/A | N/A | N/A | PA3354 | PA3354 | N/A | N/A | N/A | N/A | PA3354 | PA3354 | PA3354 | PA3354 | N/A | N/A | N/A | N/A | N/A |
| N/A | N/A | N/A | N/A | N/A | PA3355 | N/A | N/A | N/A | N/A | N/A | N/A | N/A | N/A | N/A | N/A | N/A | N/A | N/A |
| PA3356 | PA3356 | PA3356 | PA3356 | PA3356 | PA3356 | N/A | PA3356 | PA3356 | PA3356 | PA3356 | PA3356 | PA3356 | PA3356 | PA3356 | PA3356 | N/A | N/A | N/A |
| PA3357 | N/A | PA3357 | PA3357 | PA3357 | PA3357 | N/A | N/A | PA3357 | PA3357 | PA3357 | PA3357 | PA3357 | PA3357 | N/A | N/A | N/A | N/A | N/A |
| PA3361 | N/A | N/A | N/A | N/A | N/A | PA3361 | N/A | N/A | PA3361 | PA3361 | PA3361 | PA3361 | PA3361 | N/A | N/A | PA3361 | PA3361 | PA3361 |
| PA3363 | N/A | N/A | N/A | PA3363 | N/A | N/A | N/A | N/A | N/A | N/A | N/A | PA3363 | PA3363 | N/A | N/A | N/A | N/A | N/A |
| PA3364 | N/A | N/A | N/A | PA3364 | N/A | N/A | N/A | N/A | N/A | PA3364 | N/A | N/A | PA3364 | N/A | N/A | N/A | N/A | N/A |
| PA3365 | N/A | N/A | N/A | PA3365 | PA3365 | N/A | N/A | N/A | N/A | N/A | N/A | N/A | N/A | N/A | N/A | N/A | N/A | N/A |
| PA3366 | N/A | N/A | N/A | PA3366 | PA3366 | PA3366 | N/A | N/A | N/A | PA3366 | N/A | PA3366 | PA3366 | PA3366 | N/A | N/A | N/A | N/A |
| N/A | N/A | N/A | N/A | N/A | PA3384 | N/A | N/A | N/A | N/A | N/A | N/A | N/A | N/A | N/A | N/A | N/A | N/A | N/A |
| PA3385 | PA3385 | PA3385 | PA3385 | PA3385 | PA3385 | PA3385 | N/A | N/A | PA3385 | PA3385 | N/A | PA3385 | PA3385 | N/A | N/A | N/A | N/A | N/A |
| PA3391 | N/A | N/A | N/A | N/A | PA3391 | N/A | N/A | N/A | N/A | PA3391 | N/A | N/A | N/A | PA3391 | N/A | N/A | PA3391 | N/A |
| PA3392 | N/A | N/A | N/A | N/A | PA3392 | PA3392 | N/A | N/A | N/A | PA3392 | N/A | PA3392 | N/A | PA3392 | N/A | N/A | N/A | N/A |
| N/A | N/A | N/A | N/A | N/A | N/A | N/A | N/A | N/A | N/A | N/A | N/A | N/A | N/A | PA3393 | N/A | N/A | N/A | N/A |
| PA3394 | N/A | N/A | N/A | N/A | N/A | N/A | N/A | N/A | N/A | N/A | N/A | N/A | PA3994 | PA3394 | N/A | N/A | N/A | N/A |
| N/A | N/A | N/A | N/A | N/A | N/A | N/A | N/A | N/A | N/A | N/A | N/A | N/A | N/A | PA3395 | N/A | N/A | N/A | N/A |
| PA3396 | N/A | N/A | N/A | N/A | PA3396 | N/A | N/A | N/A | N/A | PA3396 | N/A | N/A | N/A | PA3396 | N/A | N/A | PA3396 | N/A |
| PA3397 | PA3397 | PA3397 | PA3397 | PA3397 | PA3397 | PA3397 | PA3397 | PA3397 | PA3397 | PA3397 | PA3397 | PA3397 | PA3397 | N/A | PA3397 | PA3397 | PA3397 | PA3397 |
| PA3398 | N/A | N/A | N/A | PA3398 | N/A | N/A | N/A | N/A | N/A | N/A | N/A | N/A | N/A | N/A | N/A | N/A | N/A | N/A |
| PA3400 | N/A | N/A | N/A | N/A | N/A | N/A | N/A | N/A | N/A | N/A | N/A | N/A | N/A | PA3400 | N/A | N/A | N/A | N/A |
| PA3402 | N/A | N/A | N/A | PA3402 | N/A | N/A | PA3402 | PA3402 | N/A | N/A | PA3402 | N/A | N/A | PA3402 | N/A | N/A | PA3402 | N/A |
| PA3403 | N/A | N/A | N/A | N/A | N/A | N/A | N/A | N/A | N/A | N/A | N/A | N/A | N/A | N/A | N/A | N/A | N/A | N/A |
| PA3409 | N/A | N/A | N/A | N/A | N/A | N/A | N/A | N/A | N/A | N/A | N/A | N/A | N/A | N/A | N/A | N/A | N/A | N/A |
| PA3413 | PA3413 | PA3413 | PA3413 | PA3413 | N/A | N/A | N/A | N/A | PA3413 | PA3413 | PA3413 | PA3413 | PA3413 | N/A | PA3413 | N/A | N/A | N/A |
| PA3414 | N/A | N/A | N/A | PA3414 | N/A | N/A | PA3414 | N/A | PA3414 | PA3414 | PA3414 | PA3414 | PA3414 | N/A | PA3414 | N/A | N/A | N/A |
| PA3415 | N/A | N/A | N/A | N/A | N/A | PA3415 | N/A | N/A | N/A | PA3415 | N/A | PA3415 | PA3415 | N/A | N/A | N/A | N/A | N/A |
| PA3416 | N/A | N/A | N/A | N/A | N/A | PA3416 | N/A | N/A | N/A | PA3416 | N/A | PA3416 | PA3416 | N/A | N/A | N/A | N/A | N/A |
| PA3417 | N/A | N/A | N/A | N/A | PA3417 | PA3417 | N/A | N/A | N/A | PA3417 | N/A | PA3417 | PA3417 | N/A | N/A | N/A | N/A | N/A |
| PA3418 | N/A | N/A | N/A | N/A | PA3418 | PA3418 | N/A | N/A | N/A | PA3418 | N/A | PA3418 | PA3418 | N/A | N/A | N/A | N/A | N/A |

| | | | | | | | | | | | | | | | | | | |
|---|---|---|---|---|---|---|---|---|---|---|---|---|---|---|---|---|---|---|
| PA3419 | N/A | N/A | N/A | N/A | N/A | N/A | N/A | N/A | N/A | N/A | N/A | N/A | N/A | N/A | N/A | N/A | N/A | N/A |
| PA3421 | N/A | N/A | N/A | N/A | N/A | PA3421 | N/A | N/A | PA3421 | PA3421 | N/A | N/A | PA3421 | N/A | N/A | PA3421 | N/A | N/A |
| PA3422 | N/A | N/A | N/A | N/A | N/A | PA3422 | N/A | N/A | N/A | N/A | N/A | N/A | PA3422 | N/A | N/A | N/A | N/A | N/A |
| N/A | N/A | N/A | N/A | N/A | N/A | N/A | N/A | N/A | N/A | PA3423 | N/A | N/A | N/A | N/A | N/A | N/A | N/A | N/A |
| PA3424 | N/A | N/A | N/A | N/A | N/A | N/A | N/A | N/A | N/A | N/A | N/A | N/A | N/A | PA3424 | N/A | N/A | N/A | N/A |
| N/A | N/A | N/A | N/A | N/A | N/A | N/A | N/A | N/A | N/A | N/A | N/A | N/A | PA3425 | N/A | N/A | N/A | N/A | N/A |
| PA3426 | N/A | N/A | N/A | N/A | N/A | N/A | N/A | N/A | N/A | PA3426 | N/A | PA3426 | PA3426 | N/A | N/A | N/A | N/A | N/A |
| PA3427 | N/A | PA3427 | N/A | PA3427 | N/A | PA3427 | N/A | N/A | PA3427 | PA3427 | PA3427 | PA3427 | PA3427 | N/A | N/A | N/A | N/A | N/A |
| PA3429 | N/A | N/A | N/A | N/A | N/A | N/A | N/A | N/A | N/A | PA3429 | N/A | PA3429 | N/A | N/A | N/A | N/A | N/A | N/A |
| PA3430 | N/A | N/A | N/A | N/A | N/A | PA3430 | N/A | N/A | N/A | PA3430 | PA3430 | PA3430 | PA3430 | N/A | N/A | N/A | N/A | N/A |
| PA3433 | N/A | N/A | N/A | PA3433 | N/A | N/A | N/A | N/A | N/A | N/A | N/A | N/A | N/A | N/A | N/A | N/A | N/A | N/A |
| N/A | N/A | N/A | N/A | N/A | PA3434 | N/A | N/A | N/A | N/A | N/A | N/A | N/A | N/A | N/A | N/A | N/A | N/A | N/A |
| PA3435 | PA3435 | PA3435 | PA3435 | PA3435 | PA3435 | N/A | N/A | N/A | PA3435 | PA3435 | PA3435 | PA3435 | PA3435 | N/A | PA3435 | N/A | N/A | N/A |
| PA3437 | N/A | N/A | N/A | PA3437 | N/A | N/A | N/A | N/A | PA3437 | N/A | N/A | N/A | N/A | N/A | N/A | N/A | N/A | N/A |
| PA3438 | PA3438 | PA3438 | PA3438 | PA3438 | PA3438 | PA3438 | N/A | N/A | PA3438 | PA3438 | PA3438 | PA3438 | PA3438 | N/A | N/A | N/A | N/A | N/A |
| PA3439 | PA3439 | PA3439 | PA3439 | PA3439 | N/A | PA3439 | N/A | N/A | PA3439 | PA3439 | PA3439 | PA3439 | PA3439 | N/A | PA3439 | N/A | N/A | N/A |
| PA3440 | PA3440 | PA3440 | PA3440 | PA3440 | PA3440 | PA3440 | N/A | N/A | PA3440 | PA3440 | PA3440 | PA3440 | PA3440 | N/A | PA3440 | N/A | PA3440 | PA3440 |
| PA3440.1 | PA3440.1 | PA3440.1 | PA3440.1 | PA3440.1 | N/A | N/A | N/A | N/A | PA3440.1 | PA3440.1 | PA3440.1 | PA3440.1 | PA3440.1 | N/A | PA3440.1 | N/A | N/A | N/A |
| N/A | N/A | N/A | N/A | N/A | PA3441 | N/A | N/A | N/A | N/A | N/A | N/A | PA3441 | PA3441 | N/A | N/A | N/A | N/A | N/A |
| PA3442 | N/A | N/A | N/A | N/A | N/A | N/A | N/A | N/A | N/A | N/A | PA3442 | N/A | N/A | PA3442 | N/A | N/A | N/A | N/A |
| PA3443 | N/A | N/A | N/A | N/A | PA3443 | N/A | N/A | N/A | N/A | N/A | N/A | N/A | N/A | PA3443 | N/A | N/A | N/A | N/A |
| PA3444 | N/A | N/A | N/A | N/A | PA3444 | N/A | N/A | N/A | N/A | N/A | PA3444 | PA3444 | N/A | PA3444 | N/A | N/A | N/A | N/A |
| PA3445 | N/A | N/A | N/A | N/A | PA3445 | N/A | N/A | N/A | N/A | N/A | N/A | PA3445 | N/A | N/A | N/A | N/A | N/A | N/A |
| PA3446 | N/A | N/A | N/A | N/A | N/A | N/A | N/A | N/A | N/A | N/A | PA3446 | N/A | N/A | N/A | N/A | N/A | PA3446 | N/A |
| PA3448 | N/A | N/A | N/A | N/A | N/A | N/A | N/A | N/A | N/A | N/A | N/A | N/A | N/A | N/A | N/A | N/A | N/A | N/A |
| N/A | N/A | N/A | N/A | N/A | PA3449 | N/A | N/A | N/A | N/A | N/A | N/A | N/A | N/A | N/A | N/A | N/A | N/A | N/A |
| PA3450 | N/A | PA3450 | N/A | PA3450 | PA3450 | N/A | N/A | N/A | N/A | PA3450 | PA3450 | PA3450 | PA3450 | N/A | N/A | N/A | N/A | N/A |
| PA3452 | PA3452 | PA3452 | PA3452 | PA3452 | PA3452 | PA3452 | PA3452 | PA3452 | N/A | PA3452 | PA3452 | PA3452 | PA3452 | PA3452 | PA3452 | N/A | PA3452 | N/A |
| PA3453 | PA3453 | PA3453 | PA3453 | PA3453 | PA3453 | N/A | PA3453 | N/A | PA3453 | PA3453 | PA3453 | PA3453 | PA3453 | PA3453 | PA3453 | N/A | N/A | N/A |
| N/A | N/A | N/A | N/A | N/A | PA3454 | N/A | N/A | N/A | N/A | N/A | N/A | N/A | N/A | N/A | N/A | N/A | N/A | N/A |
| PA3455 | N/A | N/A | N/A | PA3455 | PA3455 | PA3455 | N/A | PA3455 | PA3455 | PA3455 | PA3455 | PA3455 | PA3455 | PA3455 | N/A | N/A | N/A | N/A |
| PA3456 | N/A | N/A | N/A | PA3456 | N/A | N/A | N/A | PA3456 | N/A | PA3456 | N/A | N/A | PA3456 | N/A | N/A | N/A | N/A | N/A |
| PA3457 | N/A | N/A | N/A | N/A | PA3457 | N/A | N/A | N/A | N/A | N/A | N/A | N/A | N/A | PA3457 | N/A | N/A | N/A | N/A |
| PA3458 | N/A | N/A | N/A | PA3458 | PA3458 | PA3458 | N/A | N/A | PA3458 | PA3458 | PA3458 | PA3458 | PA3458 | N/A | N/A | N/A | N/A | N/A |
| PA3459 | N/A | N/A | N/A | N/A | PA3459 | PA3459 | N/A | N/A | N/A | PA3459 | N/A | PA3459 | PA3459 | N/A | N/A | N/A | N/A | N/A |
| PA3460 | N/A | N/A | N/A | N/A | PA3460 | N/A | N/A | N/A | N/A | PA3460 | N/A | N/A | N/A | PA3460 | N/A | N/A | N/A | N/A |
| PA3461 | N/A | N/A | N/A | N/A | PA3461 | PA3461 | N/A | N/A | N/A | PA3461 | N/A | PA3461 | PA3461 | N/A | N/A | N/A | N/A | N/A |
| PA3462 | N/A | N/A | N/A | N/A | N/A | N/A | N/A | N/A | N/A | N/A | N/A | N/A | N/A | N/A | N/A | N/A | N/A | N/A |
| PA3465 | N/A | N/A | N/A | PA3465 | PA3465 | PA3465 | N/A | N/A | N/A | PA3465 | PA3465 | PA3465 | PA3465 | PA3465 | N/A | N/A | PA3465 | N/A |
| PA3466 | PA3466 | PA3466 | PA3466 | PA3466 | N/A | PA3466 | N/A | PA3466 | PA3466 | PA3466 | PA3466 | PA3466 | PA3466 | PA3466 | PA3466 | N/A | PA3466 | N/A |
| PA3468 | PA3468 | PA3468 | PA3468 | PA3468 | N/A | PA3468 | N/A | N/A | N/A | PA3468 | PA3468 | PA3468 | PA3468 | PA3468 | N/A | N/A | PA3468 | N/A |
| PA3469 | N/A | N/A | N/A | PA3469 | N/A | PA3469 | N/A | N/A | PA3469 | PA3469 | PA3469 | PA3469 | PA3469 | N/A | N/A | N/A | N/A | N/A |
| PA3470 | PA3470 | N/A | N/A | PA3470 | PA3470 | N/A | N/A | N/A | N/A | PA3470 | PA3470 | PA3470 | N/A | N/A | N/A | N/A | N/A | N/A |
| PA3471 | PA3471 | PA3471 | PA3471 | PA3471 | PA3471 | PA3471 | N/A | PA3471 | PA3471 | PA3471 | PA3471 | PA3471 | PA3471 | PA3471 | N/A | N/A | N/A | N/A |
| PA3475 | PA3475 | N/A | N/A | PA3475 | N/A | N/A | N/A | N/A | N/A | N/A | PA3475 | N/A | PA3475 | N/A | N/A | N/A | N/A | N/A |
| PA3476 | N/A | N/A | N/A | N/A | PA3476 | N/A | N/A | N/A | PA3476 | N/A | PA3476 | PA3476 | N/A | N/A | N/A | N/A | N/A | N/A |
| PA3477 | N/A | N/A | N/A | PA3477 | PA3477 | PA3477 | PA3477 | N/A | PA3477 | PA3477 | PA3477 | PA3477 | PA3477 | PA3477 | N/A | N/A | PA3477 | N/A |
| PA3478 | N/A | N/A | N/A | N/A | PA3478 | N/A | N/A | N/A | N/A | N/A | N/A | N/A | PA3478 | N/A | N/A | N/A | N/A | N/A |
| PA3479 | N/A | N/A | N/A | N/A | PA3479 | N/A | N/A | N/A | N/A | N/A | PA3479 | PA3479 | PA3479 | N/A | N/A | N/A | N/A | N/A |
| PA3480 | PA3480 | PA3480 | PA3480 | PA3480 | PA3480 | PA3480 | PA3480 | N/A | PA3480 | PA3480 | PA3480 | PA3480 | PA3480 | N/A | PA3480 | N/A | PA3480 | PA3480 |
| PA3481 | PA3481 | PA3481 | PA3481 | PA3481 | PA3481 | PA3481 | N/A | PA3481 | PA3481 | PA3481 | PA3481 | PA3481 | PA3481 | N/A | PA3481 | N/A | N/A | N/A |
| PA3482 | PA3482 | PA3482 | PA3482 | PA3482 | PA3482 | PA3482 | PA3482 | PA3482 | PA3482 | PA3482 | PA3482 | PA3482 | PA3482 | N/A | PA3482 | PA3482 | N/A | N/A |
| PA3483 | PA3483 | PA3483 | N/A | PA3483 | N/A | PA3483 | PA3483 | N/A | PA3483 | PA3483 | PA3483 | PA3483 | PA3483 | PA3483 | PA3483 | N/A | PA3483 | PA3483 |
| PA3484 | N/A | N/A | N/A | PA3484 | N/A | N/A | PA3484 | N/A | N/A | N/A | N/A | N/A | N/A | PA3484 | N/A | N/A | PA3484 | N/A |
| PA3485 | N/A | N/A | N/A | N/A | N/A | N/A | PA3485 | N/A | N/A | PA3485 | N/A | N/A | N/A | PA3485 | N/A | N/A | PA3485 | N/A |
| N/A | N/A | N/A | N/A | N/A | N/A | N/A | N/A | N/A | PA3486 | N/A | N/A | N/A | N/A | N/A | N/A | N/A | N/A | N/A |
| N/A | N/A | N/A | N/A | N/A | N/A | N/A | N/A | N/A | PA3487 | N/A | N/A | N/A | N/A | N/A | N/A | N/A | PA3487 | N/A |
| PA3490 | N/A | N/A | N/A | N/A | N/A | N/A | N/A | N/A | N/A | N/A | N/A | N/A | N/A | PA3490 | N/A | N/A | N/A | N/A |
| PA3491 | N/A | N/A | N/A | PA3491 | N/A | N/A | N/A | N/A | N/A | N/A | N/A | N/A | N/A | PA3491 | N/A | N/A | PA3491 | N/A |
| PA3493 | N/A | N/A | N/A | N/A | N/A | N/A | N/A | N/A | N/A | N/A | N/A | N/A | N/A | N/A | N/A | N/A | N/A | N/A |
| PA3494 | N/A | N/A | N/A | N/A | N/A | N/A | N/A | N/A | N/A | N/A | N/A | N/A | N/A | PA3494 | N/A | N/A | N/A | N/A |
| PA3495 | N/A | PA3495 | PA3495 | PA3495 | PA3495 | N/A | N/A | N/A | PA3495 | PA3495 | PA3495 | PA3495 | PA3495 | N/A | N/A | N/A | N/A | N/A |
| PA3496 | PA3496 | PA3496 | N/A | N/A | N/A | N/A | N/A | N/A | PA3496 | PA3496 | PA3496 | PA3496 | N/A | N/A | N/A | N/A | N/A | N/A |
| N/A | N/A | N/A | N/A | PA3505 | N/A | N/A | N/A | N/A | N/A | N/A | N/A | N/A | N/A | N/A | N/A | N/A | N/A | N/A |
| PA3508 | N/A | N/A | N/A | N/A | N/A | N/A | N/A | N/A | N/A | N/A | PA3508 | N/A | N/A | N/A | N/A | N/A | N/A | N/A |
| N/A | N/A | N/A | N/A | N/A | N/A | N/A | N/A | N/A | N/A | N/A | PA3513 | N/A | N/A | N/A | N/A | N/A | N/A | N/A |
| N/A | N/A | N/A | N/A | N/A | N/A | N/A | N/A | N/A | N/A | N/A | N/A | PA3515 | N/A | N/A | N/A | N/A | N/A | N/A |
| N/A | N/A | N/A | N/A | N/A | N/A | N/A | N/A | N/A | N/A | PA3518 | PA3518 | PA3518 | N/A | N/A | N/A | N/A | N/A | N/A |
| N/A | N/A | N/A | N/A | N/A | N/A | N/A | N/A | N/A | N/A | N/A | N/A | PA3519 | N/A | N/A | N/A | N/A | N/A | N/A |
| PA3520 | N/A | N/A | N/A | N/A | N/A | N/A | N/A | N/A | N/A | N/A | N/A | N/A | N/A | N/A | N/A | N/A | N/A | N/A |
| N/A | N/A | N/A | N/A | N/A | PA3523 | N/A | N/A | N/A | N/A | N/A | N/A | N/A | N/A | N/A | N/A | N/A | N/A | N/A |
| PA3524 | N/A | PA3524 | PA3524 | PA3524 | PA3524 | PA3524 | N/A | N/A | PA3524 | PA3524 | PA3524 | PA3524 | PA3524 | N/A | N/A | N/A | N/A | N/A |
| PA3525 | PA3525 | PA3525 | PA3525 | PA3525 | PA3525 | PA3525 | PA3525 | PA3525 | PA3525 | PA3525 | PA3525 | PA3525 | PA3525 | PA3525 | PA3525 | N/A | N/A | N/A |
| PA3526 | N/A | N/A | N/A | PA3526 | PA3526 | N/A | N/A | N/A | N/A | N/A | N/A | N/A | N/A | N/A | N/A | N/A | N/A | N/A |
| PA3527 | PA3527 | PA3527 | PA3527 | PA3527 | PA3527 | PA3527 | N/A | N/A | PA3527 | PA3527 | PA3527 | PA3527 | PA3527 | N/A | PA3527 | N/A | N/A | N/A |
| PA3528 | PA3528 | PA3528 | PA3528 | PA3528 | PA3528 | PA3528 | PA3528 | N/A | PA3528 | PA3528 | PA3528 | PA3528 | PA3528 | N/A | PA3528 | N/A | N/A | N/A |
| PA3529 | PA3529 | PA3529 | PA3529 | PA3529 | N/A | PA3529 | PA3529 | PA3529 | PA3529 | PA3529 | PA3529 | PA3529 | PA3529 | PA3529 | PA3529 | PA3529 | PA3529 | PA3529 |
| N/A | N/A | N/A | N/A | N/A | PA3530 | N/A | N/A | N/A | N/A | N/A | N/A | N/A | N/A | N/A | N/A | N/A | N/A | N/A |
| PA3531 | PA3531 | PA3531 | PA3531 | PA3531 | PA3531 | PA3531 | PA3531 | PA3531 | PA3531 | PA3531 | PA3531 | PA3531 | PA3531 | PA3531 | PA3531 | PA3531 | N/A | PA3531 |
| N/A | N/A | N/A | N/A | N/A | PA3532 | N/A | N/A | N/A | N/A | N/A | N/A | N/A | N/A | PA3532 | N/A | N/A | N/A | N/A |
| PA3533 | PA3533 | PA3533 | PA3533 | PA3533 | PA3533 | PA3533 | PA3533 | N/A | PA3533 | PA3533 | PA3533 | PA3533 | PA3533 | PA3533 | PA3533 | N/A | PA3533 | PA3533 |
| PA3534 | N/A | N/A | N/A | N/A | N/A | N/A | N/A | N/A | N/A | N/A | N/A | N/A | N/A | N/A | N/A | N/A | N/A | N/A |
| PA3535 | N/A | N/A | N/A | N/A | PA3535 | PA3535 | N/A | PA3535 | N/A | N/A | N/A | PA3535 | PA3535 | N/A | N/A | N/A | N/A | N/A |
| N/A | PA3536 | PA3536 | N/A | N/A | N/A | N/A | N/A | N/A | N/A | N/A | N/A | N/A | N/A | N/A | PA3536 | N/A | N/A | N/A |
| PA3537 | PA3537 | PA3537 | N/A | PA3537 | N/A | PA3537 | N/A | N/A | PA3537 | PA3537 | PA3537 | PA3537 | PA3537 | N/A | N/A | N/A | N/A | N/A |
| PA3538 | PA3538 | PA3538 | PA3538 | PA3538 | PA3538 | N/A | PA3538 | PA3538 | PA3538 | N/A | PA3538 | PA3538 | PA3538 | PA3538 | N/A | N/A | PA3538 | N/A |
| PA3539 | PA3539 | PA3539 | N/A | PA3539 | PA3539 | N/A | PA3539 | N/A | PA3539 | PA3539 | PA3539 | PA3539 | PA3539 | N/A | PA3539 | N/A | N/A | N/A |
| PA3540 | N/A | N/A | N/A | N/A | PA3540 | N/A | N/A | N/A | N/A | N/A | N/A | N/A | N/A | N/A | N/A | N/A | N/A | N/A |
| PA3541 | N/A | N/A | N/A | N/A | N/A | N/A | N/A | N/A | N/A | N/A | N/A | N/A | N/A | N/A | N/A | N/A | N/A | N/A |
| PA3542 | N/A | N/A | N/A | N/A | PA3542 | N/A | N/A | N/A | N/A | N/A | N/A | N/A | N/A | N/A | N/A | N/A | N/A | N/A |
| PA3543 | N/A | N/A | N/A | N/A | PA3543 | N/A | N/A | N/A | N/A | N/A | N/A | N/A | N/A | N/A | N/A | N/A | N/A | N/A |
| PA3544 | N/A | N/A | N/A | N/A | PA3544 | N/A | N/A | N/A | N/A | N/A | N/A | N/A | N/A | N/A | N/A | N/A | N/A | N/A |
| PA3545 | N/A | N/A | N/A | N/A | PA3545 | N/A | N/A | N/A | N/A | N/A | N/A | N/A | N/A | N/A | N/A | N/A | N/A | N/A |
| PA3546 | N/A | N/A | N/A | N/A | PA3546 | N/A | N/A | N/A | N/A | N/A | N/A | N/A | N/A | N/A | N/A | N/A | N/A | N/A |
| N/A | N/A | N/A | N/A | N/A | PA3547 | N/A | N/A | N/A | N/A | N/A | N/A | N/A | N/A | N/A | N/A | N/A | N/A | N/A |
| PA3548 | N/A | N/A | N/A | N/A | N/A | N/A | N/A | N/A | N/A | N/A | N/A | N/A | N/A | N/A | N/A | N/A | N/A | N/A |
| PA3549 | N/A | N/A | N/A | N/A | PA3549 | N/A | N/A | N/A | N/A | N/A | N/A | N/A | N/A | N/A | N/A | N/A | N/A | N/A |
| PA3550 | N/A | N/A | N/A | N/A | N/A | N/A | N/A | N/A | N/A | N/A | N/A | N/A | N/A | N/A | N/A | N/A | N/A | N/A |
| PA3551 | N/A | N/A | N/A | N/A | PA3551 | N/A | N/A | N/A | N/A | N/A | N/A | N/A | N/A | N/A | N/A | N/A | N/A | N/A |
| N/A | PA3552 | PA3552 | N/A | PA3552 | PA3552 | PA3552 | N/A | PA3552 | PA3552 | N/A | N/A | N/A | PA3552 | N/A | N/A | N/A | N/A | N/A |
| N/A | PA3553 | PA3553 | PA3553 | PA3553 | N/A | N/A | N/A | N/A | PA3553 | N/A | N/A | N/A | N/A | PA3553 | PA3553 | N/A | PA3553 | N/A |
| N/A | PA3554 | PA3554 | PA3554 | PA3554 | N/A | N/A | N/A | PA3554 | PA3554 | N/A | PA3554 | N/A | PA3554 | PA3554 | PA3554 | N/A | PA3554 | N/A |
| N/A | N/A | N/A | N/A | PA3555 | N/A | N/A | N/A | N/A | N/A | N/A | N/A | N/A | N/A | PA3555 | N/A | N/A | N/A | N/A |
| N/A | N/A | PA3556 | N/A | PA3556 | N/A | N/A | N/A | N/A | PA3556 | N/A | N/A | N/A | N/A | PA3556 | N/A | N/A | N/A | N/A |
| N/A | N/A | N/A | N/A | N/A | N/A | N/A | N/A | N/A | N/A | N/A | N/A | N/A | N/A | PA3558 | N/A | N/A | N/A | N/A |
| N/A | PA3559 | PA3559 | PA3559 | N/A | N/A | N/A | N/A | PA3559 | PA3559 | N/A | N/A | N/A | N/A | PA3559 | N/A | N/A | N/A | N/A |
| PA3560 | N/A | N/A | N/A | PA3560 | N/A | N/A | N/A | PA3560 | PA3560 | N/A | N/A | PA3560 | PA3560 | PA3560 | N/A | N/A | PA3560 | N/A |
| PA3561 | PA3561 | PA3561 | PA3561 | PA3561 | PA3561 | N/A | PA3561 | PA3561 | PA3561 | N/A | PA3561 | PA3561 | PA3561 | N/A | PA3561 | N/A | PA3561 | N/A |
| PA3562 | PA3562 | PA3562 | PA3562 | PA3562 | N/A | N/A | N/A | PA3562 | PA3562 | N/A | PA3562 | PA3562 | PA3562 | PA3562 | N/A | N/A | PA3562 | N/A |

| | | | | | | | | | | | | | | | | | | |
|---|---|---|---|---|---|---|---|---|---|---|---|---|---|---|---|---|---|---|
| PA3563 | PA3563 | PA3563 | PA3563 | PA3563 | N/A | N/A | N/A | PA3563 | PA3563 | PA3563 | PA3563 | PA3563 | PA3563 | N/A | N/A | N/A | N/A | N/A |
| PA3564 | N/A | N/A | N/A | N/A | N/A | N/A | N/A | N/A | N/A | N/A | N/A | N/A | N/A | N/A | N/A | N/A | N/A | N/A |
| PA3565 | N/A | N/A | N/A | N/A | PA3565 | N/A | N/A | N/A | N/A | N/A | PA3565 | N/A | N/A | N/A | N/A | N/A | N/A | N/A |
| PA3566 | PA3566 | PA3566 | PA3566 | PA3566 | PA3566 | N/A | N/A | N/A | PA3566 | PA3566 | PA3566 | PA3566 | PA3566 | N/A | PA3566 | N/A | N/A | N/A |
| PA3567 | PA3567 | PA3567 | PA3567 | PA3567 | PA3567 | PA3567 | N/A | N/A | PA3567 | PA3567 | PA3567 | PA3567 | PA3567 | N/A | PA3567 | N/A | N/A | N/A |
| PA3568 | N/A | N/A | N/A | N/A | N/A | PA3568 | N/A | N/A | N/A | N/A | PA3568 | PA3568 | PA3568 | N/A | N/A | N/A | N/A | N/A |
| PA3569 | N/A | N/A | N/A | N/A | PA3569 | PA3569 | N/A | N/A | PA3569 | PA3569 | PA3569 | PA3569 | PA3569 | N/A | N/A | N/A | N/A | PA3569 |
| PA3570 | N/A | N/A | N/A | N/A | PA3570 | PA3570 | N/A | N/A | PA3570 | PA3570 | PA3570 | PA3570 | PA3570 | PA3570 | N/A | PA3570 | N/A | N/A |
| PA3574 | PA3574 | N/A | PA3574 | PA3574 | PA3574 | PA3574 | PA3574 | N/A | PA3574 | PA3574 | PA3574 | PA3574 | PA3574 | N/A | N/A | N/A | N/A | N/A |
| PA3574.1 | N/A | PA3574.1 | N/A | PA3574.1 | N/A | PA3574.1 | N/A | N/A | PA3574.1 | PA3574.1 | PA3574.1 | PA3574.1 | PA3574.1 | N/A | N/A | N/A | N/A | N/A |
| PA3575 | PA3575 | N/A | N/A | PA3575 | N/A | PA3575 | N/A | N/A | PA3575 | PA3575 | PA3575 | PA3575 | N/A | PA3575 | N/A | N/A | PA3575 | N/A |
| PA3576 | N/A | N/A | N/A | N/A | N/A | N/A | N/A | N/A | N/A | N/A | N/A | N/A | N/A | N/A | N/A | N/A | N/A | N/A |
| N/A | N/A | N/A | N/A | N/A | PA3578 | N/A | N/A | N/A | N/A | PA3578 | N/A | N/A | PA3578 | N/A | N/A | N/A | N/A | N/A |
| PA3579 | PA3579 | PA3579 | PA3579 | PA3579 | PA3579 | N/A | N/A | PA3579 | PA3579 | PA3579 | N/A | N/A | PA3579 | N/A | N/A | N/A | N/A | N/A |
| PA3580 | N/A | N/A | PA3580 | PA3580 | PA3580 | N/A | PA3580 | N/A | PA3580 | PA3580 | PA3580 | PA3580 | PA3580 | PA3580 | PA3580 | N/A | N/A | N/A |
| PA3581 | PA3581 | N/A | N/A | N/A | PA3581 | PA3581 | N/A | N/A | N/A | PA3581 | N/A | N/A | N/A | PA3581 | N/A | N/A | N/A | N/A |
| PA3582 | PA3582 | PA3582 | PA3582 | PA3582 | PA3582 | PA3582 | N/A | PA3582 | PA3582 | PA3582 | PA3582 | PA3582 | PA3582 | N/A | PA3582 | N/A | PA3582 | N/A |
| PA3583 | PA3583 | PA3583 | N/A | PA3583 | PA3583 | PA3583 | N/A | PA3583 | PA3583 | PA3583 | PA3583 | PA3583 | PA3583 | N/A | PA3583 | N/A | N/A | N/A |
| PA3584 | PA3584 | PA3584 | PA3584 | PA3584 | PA3584 | PA3584 | PA3584 | PA3584 | PA3584 | PA3584 | PA3584 | N/A | N/A | PA3584 | PA3584 | N/A | PA3584 | N/A |
| PA3587 | N/A | N/A | N/A | PA3587 | N/A | N/A | N/A | N/A | N/A | PA3587 | PA3587 | N/A | N/A | N/A | N/A | N/A | N/A | N/A |
| N/A | N/A | N/A | N/A | N/A | N/A | N/A | N/A | N/A | N/A | PA3590 | N/A | N/A | N/A | N/A | N/A | N/A | N/A | N/A |
| PA3593 | N/A | N/A | N/A | N/A | N/A | N/A | N/A | N/A | N/A | N/A | N/A | PA3593 | PA3593 | N/A | N/A | N/A | N/A | N/A |
| PA3598 | N/A | N/A | N/A | N/A | N/A | N/A | N/A | N/A | N/A | N/A | N/A | N/A | N/A | N/A | N/A | N/A | N/A | N/A |
| PA3599 | N/A | N/A | N/A | N/A | N/A | N/A | N/A | N/A | N/A | N/A | N/A | N/A | N/A | N/A | N/A | N/A | N/A | N/A |
| PA3601 | N/A | N/A | N/A | N/A | N/A | N/A | N/A | N/A | N/A | N/A | N/A | N/A | N/A | N/A | N/A | N/A | N/A | N/A |
| PA3602 | N/A | PA3602 | PA3602 | PA3602 | N/A | PA3602 | PA3602 | PA3602 | PA3602 | PA3602 | PA3602 | PA3602 | PA3602 | PA3602 | N/A | N/A | PA3602 | N/A |
| PA3603 | N/A | N/A | N/A | N/A | PA3603 | N/A | N/A | N/A | N/A | N/A | N/A | N/A | N/A | PA3603 | N/A | N/A | N/A | N/A |
| PA3604 | PA3604 | PA3604 | PA3604 | PA3604 | PA3604 | PA3604 | N/A | N/A | PA3604 | PA3604 | PA3604 | PA3604 | PA3604 | N/A | N/A | N/A | N/A | N/A |
| PA3606 | N/A | N/A | N/A | PA3606 | PA3606 | N/A | N/A | N/A | PA3606 | N/A | PA3606 | N/A | N/A | N/A | N/A | N/A | N/A | N/A |
| PA3607 | PA3607 | PA3607 | PA3607 | PA3607 | N/A | N/A | PA3607 | PA3607 | PA3607 | N/A | PA3607 | N/A | PA3607 | PA3607 | PA3607 | N/A | PA3607 | N/A |
| PA3608 | N/A | N/A | N/A | N/A | N/A | N/A | N/A | N/A | N/A | N/A | N/A | N/A | N/A | PA3608 | N/A | N/A | N/A | N/A |
| N/A | N/A | N/A | N/A | N/A | N/A | N/A | N/A | N/A | N/A | N/A | N/A | N/A | N/A | PA3609 | N/A | N/A | N/A | N/A |
| PA3610 | PA3610 | PA3610 | PA3610 | PA3610 | PA3610 | N/A | PA3610 | N/A | PA3610 | PA3610 | PA3610 | PA3610 | PA3610 | N/A | PA3610 | N/A | N/A | N/A |
| PA3611 | N/A | PA3611 | N/A | PA3611 | PA3611 | PA3611 | PA3611 | N/A | PA3611 | PA3611 | PA3611 | PA3611 | PA3611 | PA3611 | PA3611 | N/A | PA3611 | PA3611 |
| N/A | N/A | N/A | N/A | N/A | PA3612 | N/A | N/A | N/A | N/A | N/A | N/A | N/A | N/A | N/A | N/A | N/A | N/A | N/A |
| PA3613 | N/A | N/A | N/A | PA3613 | PA3613 | PA3613 | N/A | PA3613 | PA3613 | PA3613 | PA3613 | PA3613 | PA3613 | PA3613 | N/A | N/A | N/A | N/A |
| PA3614 | N/A | N/A | N/A | PA3614 | N/A | PA3614 | N/A | N/A | PA3614 | PA3614 | PA3614 | PA3614 | PA3614 | PA3614 | N/A | PA3614 | N/A | N/A |
| PA3615 | PA3615 | PA3615 | N/A | PA3615 | PA3615 | PA3615 | N/A | N/A | PA3615 | PA3615 | PA3615 | PA3615 | PA3615 | N/A | N/A | N/A | N/A | N/A |
| PA3616 | N/A | N/A | N/A | N/A | PA3616 | N/A | N/A | N/A | N/A | N/A | N/A | N/A | N/A | N/A | N/A | N/A | N/A | N/A |
| PA3617 | PA3617 | PA3617 | PA3617 | PA3617 | PA3617 | PA3617 | PA3617 | PA3617 | PA3617 | PA3617 | PA3617 | PA3617 | PA3617 | PA3617 | PA3617 | N/A | PA3617 | PA3617 |
| PA3618 | PA3618 | PA3618 | PA3618 | PA3618 | PA3618 | PA3618 | N/A | N/A | PA3618 | PA3618 | PA3618 | PA3618 | PA3618 | N/A | N/A | N/A | PA3618 | PA3618 |
| N/A | N/A | N/A | N/A | N/A | PA3619 | N/A | N/A | N/A | N/A | N/A | N/A | N/A | N/A | N/A | N/A | N/A | N/A | N/A |
| PA3620 | PA3620 | PA3620 | PA3620 | PA3620 | PA3620 | PA3620 | PA3620 | PA3620 | PA3620 | PA3620 | PA3620 | PA3620 | PA3620 | PA3620 | N/A | N/A | N/A | N/A |
| PA3621 | PA3621 | PA3621 | PA3621 | PA3621 | PA3621 | PA3621 | PA3621 | PA3621 | PA3621 | PA3621 | PA3621 | PA3621 | PA3621 | PA3621 | N/A | N/A | N/A | N/A |
| PA3622 | PA3622 | PA3622 | N/A | PA3622 | PA3622 | PA3622 | PA3622 | PA3622 | PA3622 | PA3622 | PA3622 | PA3622 | PA3622 | PA3622 | N/A | N/A | PA3622 | N/A |
| PA3623 | N/A | N/A | N/A | PA3623 | N/A | N/A | N/A | N/A | PA3623 | PA3623 | PA3623 | PA3623 | PA3623 | PA3623 | PA3623 | PA3623 | PA3623 | PA3623 |
| PA3624 | PA3624 | PA3624 | PA3624 | PA3624 | PA3624 | N/A | PA3624 | N/A | PA3624 | PA3624 | PA3624 | PA3624 | PA3624 | N/A | PA3624 | N/A | N/A | N/A |
| PA3625 | PA3625 | PA3625 | PA3625 | PA3625 | PA3625 | N/A | PA3625 | N/A | PA3625 | PA3625 | PA3625 | PA3625 | PA3625 | N/A | PA3625 | N/A | N/A | N/A |
| PA3626 | N/A | N/A | N/A | PA3626 | PA3626 | N/A | N/A | N/A | N/A | N/A | N/A | N/A | PA3626 | N/A | N/A | N/A | N/A | N/A |
| PA3627 | PA3627 | PA3627 | PA3627 | PA3627 | PA3627 | PA3627 | N/A | N/A | PA3627 | PA3627 | PA3627 | PA3627 | PA3627 | N/A | PA3627 | N/A | N/A | N/A |
| PA3628 | PA3628 | PA3628 | N/A | PA3628 | PA3628 | PA3628 | N/A | N/A | N/A | PA3628 | PA3628 | PA3628 | PA3628 | N/A | N/A | N/A | N/A | N/A |
| PA3629 | PA3629 | PA3629 | PA3629 | PA3629 | PA3629 | PA3629 | N/A | N/A | PA3629 | PA3629 | PA3629 | PA3629 | PA3629 | N/A | N/A | N/A | N/A | N/A |
| PA3631 | PA3631 | PA3631 | N/A | PA3631 | PA3631 | N/A | N/A | PA3631 | PA3631 | PA3631 | PA3631 | PA3631 | PA3631 | PA3631 | N/A | N/A | PA3631 | N/A |
| PA3632 | N/A | N/A | N/A | PA3632 | PA3632 | N/A | N/A | N/A | PA3632 | N/A | PA3632 | N/A | N/A | N/A | N/A | N/A | N/A | N/A |
| PA3633 | PA3633 | PA3633 | N/A | PA3633 | N/A | N/A | PA3633 | N/A | PA3633 | PA3633 | PA3633 | PA3633 | N/A | N/A | N/A | N/A | N/A | N/A |
| PA3634 | N/A | N/A | N/A | N/A | PA3634 | N/A | N/A | N/A | PA3634 | N/A | PA3634 | N/A | N/A | PA3634 | N/A | N/A | N/A | N/A |
| PA3635 | PA3635 | PA3635 | PA3635 | PA3635 | PA3635 | PA3635 | PA3635 | PA3635 | PA3635 | PA3635 | PA3635 | PA3635 | PA3635 | PA3635 | PA3635 | PA3635 | PA3635 | N/A |
| PA3636 | PA3636 | PA3636 | PA3636 | PA3636 | PA3636 | PA3636 | PA3636 | PA3636 | PA3636 | PA3636 | PA3636 | PA3636 | PA3636 | PA3636 | PA3636 | N/A | N/A | N/A |
| PA3637 | PA3637 | PA3637 | PA3637 | PA3637 | PA3637 | PA3637 | PA3637 | PA3637 | PA3637 | PA3637 | PA3637 | PA3637 | PA3637 | PA3637 | PA3637 | N/A | PA3637 | PA3637 |
| PA3638 | N/A | N/A | N/A | PA3638 | PA3638 | N/A | N/A | N/A | N/A | N/A | N/A | N/A | N/A | N/A | N/A | N/A | N/A | N/A |
| PA3639 | PA3639 | PA3639 | PA3639 | PA3639 | PA3639 | PA3639 | PA3639 | PA3639 | PA3639 | PA3639 | PA3639 | PA3639 | PA3639 | PA3639 | PA3639 | N/A | PA3639 | PA3639 |
| PA3640 | PA3640 | PA3640 | PA3640 | PA3640 | PA3640 | PA3640 | N/A | PA3640 | PA3640 | PA3640 | PA3640 | PA3640 | PA3640 | PA3640 | PA3640 | N/A | PA3640 | N/A |
| PA3641 | PA3641 | PA3641 | PA3641 | PA3641 | N/A | PA3641 | N/A | PA3641 | PA3641 | PA3641 | PA3641 | PA3641 | PA3641 | PA3641 | N/A | N/A | PA3641 | PA3641 |
| PA3642 | N/A | PA3642 | PA3642 | PA3642 | PA3642 | N/A | N/A | N/A | PA3642 | PA3642 | N/A | PA3642 | N/A | N/A | N/A | N/A | N/A | N/A |
| PA3643 | PA3643 | PA3643 | PA3643 | PA3643 | PA3643 | N/A | PA3643 | PA3643 | PA3643 | PA3643 | N/A | PA3643 | PA3643 | PA3643 | N/A | N/A | PA3643 | N/A |
| PA3644 | PA3644 | PA3644 | PA3644 | PA3644 | PA3644 | PA3644 | PA3644 | N/A | PA3644 | PA3644 | PA3644 | PA3644 | PA3644 | PA3644 | PA3644 | N/A | N/A | N/A |
| PA3645 | PA3645 | PA3645 | PA3645 | PA3645 | PA3645 | PA3645 | PA3645 | PA3645 | PA3645 | PA3645 | PA3645 | PA3645 | PA3645 | PA3645 | PA3645 | N/A | PA3645 | PA3645 |
| PA3646 | PA3646 | PA3646 | PA3646 | PA3646 | PA3646 | PA3646 | N/A | PA3646 | PA3646 | PA3646 | PA3646 | PA3646 | PA3646 | PA3646 | PA3646 | N/A | N/A | N/A |
| PA3647 | PA3647 | PA3647 | PA3647 | PA3647 | PA3647 | PA3647 | PA3647 | PA3647 | PA3647 | PA3647 | PA3647 | PA3647 | PA3647 | PA3647 | PA3647 | N/A | PA3647 | PA3647 |
| PA3648 | PA3648 | PA3648 | PA3648 | PA3648 | N/A | PA3648 | PA3648 | PA3648 | PA3648 | PA3648 | PA3648 | PA3648 | PA3648 | PA3648 | PA3648 | PA3648 | PA3648 | N/A |
| PA3649 | N/A | PA3649 | PA3649 | PA3649 | N/A | N/A | N/A | N/A | PA3649 | N/A | N/A | PA3649 | PA3649 | PA3649 | PA3649 | N/A | N/A | N/A |
| PA3650 | PA3650 | PA3650 | PA3650 | PA3650 | PA3650 | N/A | N/A | PA3650 | PA3650 | PA3650 | PA3650 | PA3650 | PA3650 | N/A | N/A | N/A | N/A | N/A |
| PA3651 | PA3651 | PA3651 | PA3651 | N/A | N/A | PA3651 | N/A | PA3651 | PA3651 | N/A | N/A | PA3651 | N/A | PA3651 | N/A | N/A | PA3651 | N/A |
| PA3652 | PA3652 | PA3652 | PA3652 | PA3652 | PA3652 | N/A | N/A | N/A | PA3652 | N/A | PA3652 | N/A | N/A | PA3652 | N/A | N/A | N/A | N/A |
| PA3653 | PA3653 | PA3653 | PA3653 | PA3653 | PA3653 | PA3653 | PA3653 | N/A | PA3653 | PA3653 | PA3653 | PA3653 | PA3653 | N/A | PA3653 | N/A | PA3653 | PA3653 |
| PA3654 | PA3654 | PA3654 | PA3654 | PA3654 | PA3654 | PA3654 | PA3654 | PA3654 | PA3654 | PA3654 | PA3654 | PA3654 | PA3654 | N/A | PA3654 | N/A | PA3654 | N/A |
| PA3655 | PA3655 | PA3655 | PA3655 | PA3655 | PA3655 | PA3655 | PA3655 | PA3655 | PA3655 | PA3655 | PA3655 | PA3655 | PA3655 | PA3655 | PA3655 | PA3655 | PA3655 | PA3655 |
| PA3656 | PA3656 | PA3656 | PA3656 | PA3656 | PA3656 | PA3656 | PA3656 | PA3656 | PA3656 | PA3656 | PA3656 | PA3656 | PA3656 | PA3656 | PA3656 | PA3656 | PA3656 | PA3656 |
| PA3657 | PA3657 | PA3657 | PA3657 | PA3657 | PA3657 | PA3657 | PA3657 | PA3657 | PA3657 | PA3657 | PA3657 | PA3657 | PA3657 | PA3657 | PA3657 | N/A | N/A | N/A |
| PA3658 | N/A | PA3658 | N/A | PA3658 | PA3658 | N/A | N/A | N/A | PA3658 | N/A | PA3658 | PA3658 | PA3658 | N/A | N/A | N/A | N/A | N/A |
| PA3659 | PA3659 | PA3659 | PA3659 | PA3659 | PA3659 | N/A | N/A | PA3659 | PA3659 | N/A | PA3659 | PA3659 | PA3659 | N/A | PA3659 | N/A | N/A | PA3659 |
| PA3660 | N/A | N/A | N/A | N/A | N/A | N/A | N/A | N/A | N/A | PA3660 | N/A | N/A | N/A | PA3660 | N/A | N/A | N/A | N/A |
| N/A | N/A | N/A | N/A | PA3661 | N/A | N/A | N/A | N/A | N/A | N/A | N/A | N/A | N/A | N/A | N/A | N/A | N/A | N/A |
| N/A | N/A | N/A | N/A | N/A | PA3663 | N/A | N/A | N/A | N/A | N/A | N/A | N/A | N/A | N/A | N/A | N/A | N/A | N/A |
| PA3664 | PA3664 | PA3664 | PA3664 | PA3664 | PA3664 | PA3664 | N/A | N/A | PA3664 | PA3664 | PA3664 | PA3664 | PA3664 | N/A | PA3664 | N/A | N/A | N/A |
| N/A | N/A | N/A | N/A | N/A | PA3665 | N/A | N/A | N/A | N/A | N/A | N/A | N/A | N/A | N/A | N/A | N/A | N/A | N/A |
| PA3666 | PA3666 | PA3666 | PA3666 | PA3666 | PA3666 | PA3666 | PA3666 | N/A | PA3666 | PA3666 | PA3666 | PA3666 | PA3666 | N/A | PA3666 | PA3666 | N/A | N/A |
| PA3667 | N/A | N/A | PA3667 | PA3667 | PA3667 | N/A | N/A | N/A | N/A | N/A | N/A | N/A | PA3667 | N/A | N/A | N/A | N/A | N/A |
| PA3668 | N/A | N/A | N/A | PA3668 | N/A | N/A | N/A | N/A | N/A | N/A | N/A | PA3668 | PA3668 | N/A | N/A | N/A | N/A | N/A |
| PA3669 | N/A | N/A | N/A | PA3669 | N/A | N/A | N/A | N/A | N/A | PA3669 | N/A | PA3669 | N/A | PA3669 | N/A | N/A | PA3669 | N/A |
| PA3670 | N/A | N/A | N/A | PA3670 | N/A | PA3670 | N/A | N/A | N/A | N/A | PA3670 | PA3670 | PA3670 | PA3670 | N/A | N/A | PA3670 | N/A |
| PA3671 | N/A | N/A | N/A | N/A | N/A | N/A | N/A | N/A | N/A | N/A | N/A | N/A | N/A | N/A | N/A | N/A | N/A | N/A |
| PA3672 | N/A | N/A | N/A | PA3672 | N/A | N/A | N/A | N/A | N/A | N/A | N/A | PA3672 | N/A | PA3672 | N/A | N/A | PA3672 | N/A |
| PA3673 | PA3673 | PA3673 | PA3673 | PA3673 | PA3673 | PA3673 | N/A | PA3673 | PA3673 | PA3673 | PA3673 | PA3673 | PA3673 | PA3673 | N/A | N/A | PA3673 | N/A |
| PA3674 | PA3674 | PA3674 | PA3674 | PA3674 | PA3674 | PA3674 | PA3674 | N/A | PA3674 | PA3674 | PA3674 | PA3674 | PA3674 | PA3674 | PA3674 | N/A | PA3674 | N/A |
| PA3675 | PA3675 | PA3675 | PA3675 | PA3675 | PA3675 | N/A | N/A | N/A | PA3675 | PA3675 | N/A | N/A | N/A | PA3675 | PA3675 | N/A | PA3675 | PA3675 |
| PA3676 | N/A | N/A | N/A | PA3676 | N/A | PA3676 | N/A | N/A | N/A | N/A | PA3676 | PA3676 | N/A | PA3676 | N/A | N/A | N/A | N/A |
| PA3677 | N/A | N/A | N/A | PA3677 | PA3677 | PA3677 | N/A | N/A | N/A | N/A | PA3677 | N/A | N/A | N/A | N/A | N/A | N/A | N/A |
| PA3678 | N/A | N/A | N/A | PA3678 | PA3678 | PA3678 | N/A | N/A | PA3678 | PA3678 | PA3678 | PA3678 | PA3678 | N/A | N/A | N/A | N/A | N/A |
| N/A | N/A | N/A | N/A | N/A | PA3679 | N/A | N/A | N/A | N/A | N/A | N/A | N/A | N/A | N/A | N/A | N/A | N/A | N/A |
| PA3680 | PA3680 | PA3680 | N/A | PA3680 | PA3680 | N/A | PA3680 | PA3680 | PA3680 | N/A | PA3680 | PA3680 | PA3680 | PA3680 | N/A | N/A | N/A | N/A |
| PA3681 | N/A | N/A | N/A | N/A | N/A | N/A | N/A | N/A | N/A | N/A | N/A | N/A | N/A | N/A | N/A | N/A | N/A | N/A |
| PA3682 | N/A | N/A | N/A | PA3682 | PA3682 | N/A | N/A | N/A | N/A | N/A | N/A | N/A | PA3682 | N/A | N/A | N/A | N/A | N/A |
| PA3683 | N/A | N/A | N/A | PA3683 | N/A | N/A | PA3683 | PA3683 | N/A | N/A | PA3683 | N/A | N/A | N/A | N/A | N/A | N/A | N/A |
| N/A | N/A | N/A | N/A | N/A | PA3684 | N/A | N/A | N/A | N/A | N/A | N/A | N/A | N/A | N/A | N/A | N/A | N/A | N/A |
| PA3685 | N/A | N/A | N/A | PA3685 | PA3685 | N/A | PA3685 | PA3685 | N/A | PA3685 | PA3685 | PA3685 | PA3685 | N/A | N/A | N/A | N/A | N/A |
| PA3686 | PA3686 | PA3686 | PA3686 | PA3686 | PA3686 | PA3686 | PA3686 | N/A | PA3686 | PA3686 | PA3686 | PA3686 | PA3686 | N/A | PA3686 | N/A | PA3686 | N/A |
| PA3687 | N/A | N/A | N/A | PA3687 | N/A | PA3687 | N/A | N/A | PA3687 | PA3687 | PA3687 | PA3687 | PA3687 | N/A | N/A | N/A | N/A | N/A |
| PA3688 | N/A | N/A | N/A | N/A | N/A | PA3688 | N/A | N/A | N/A | PA3688 | N/A | PA3688 | PA3688 | N/A | N/A | N/A | N/A | N/A |

| | | | | | | | | | | | | | | | | | | |
|---|---|---|---|---|---|---|---|---|---|---|---|---|---|---|---|---|---|---|
| PA3689 | PA3689 | N/A | N/A | N/A | PA3689 | N/A | N/A | N/A | N/A | N/A | PA3689 | N/A | N/A | N/A | N/A | N/A | N/A | N/A |
| PA3690 | PA3690 | N/A | N/A | PA3690 | PA3690 | PA3690 | N/A | PA3690 | PA3690 | PA3690 | PA3690 | PA3690 | PA3690 | PA3690 | PA3690 | N/A | PA3690 | N/A |
| PA3691 | PA3691 | PA3691 | PA3691 | PA3691 | PA3691 | PA3691 | PA3691 | N/A | PA3691 | PA3691 | PA3691 | PA3691 | PA3691 | PA3691 | PA3691 | N/A | N/A | PA3691 |
| PA3692 | PA3692 | PA3692 | PA3692 | PA3692 | PA3692 | PA3692 | PA3692 | PA3692 | PA3692 | PA3692 | PA3692 | PA3692 | PA3692 | PA3692 | PA3692 | PA3692 | PA3692 | PA3692 |
| PA3693 | N/A | N/A | N/A | PA3693 | N/A | N/A | N/A | N/A | N/A | N/A | N/A | PA3693 | N/A | N/A | N/A | N/A | N/A | N/A |
| PA3694 | PA3694 | PA3694 | PA3694 | PA3694 | N/A | N/A | PA3694 | N/A | PA3694 | PA3694 | PA3694 | PA3694 | PA3694 | PA3694 | PA3694 | N/A | PA3694 | PA3694 |
| PA3695 | N/A | N/A | N/A | N/A | N/A | N/A | N/A | N/A | N/A | N/A | N/A | N/A | N/A | PA3695 | N/A | N/A | N/A | N/A |
| PA3696 | N/A | PA3696 | N/A | PA3696 | N/A | PA3696 | N/A | N/A | PA3696 | PA3696 | PA3696 | PA3696 | PA3696 | PA3696 | N/A | N/A | N/A | N/A |
| PA3697 | N/A | N/A | N/A | N/A | PA3697 | N/A | N/A | N/A | N/A | N/A | N/A | N/A | PA3697 | PA3697 | N/A | N/A | N/A | PA3697 |
| PA3698 | N/A | N/A | N/A | N/A | PA3698 | PA3698 | N/A | N/A | PA3698 | PA3698 | PA3698 | N/A | PA3698 | N/A | N/A | N/A | N/A | N/A |
| PA3699 | N/A | N/A | N/A | PA3699 | N/A | N/A | N/A | N/A | PA3699 | PA3699 | PA3699 | PA3699 | PA3699 | N/A | N/A | N/A | N/A | N/A |
| PA3700 | PA3700 | PA3700 | PA3700 | PA3700 | PA3700 | PA3700 | PA3700 | PA3700 | PA3700 | PA3700 | PA3700 | PA3700 | PA3700 | PA3700 | PA3700 | PA3700 | N/A | N/A |
| N/A | N/A | N/A | N/A | N/A | PA3701 | N/A | N/A | N/A | N/A | N/A | N/A | N/A | N/A | N/A | N/A | N/A | N/A | N/A |
| PA3702 | N/A | N/A | N/A | PA3702 | N/A | N/A | N/A | PA3702 | N/A | PA3702 | N/A | N/A | PA3702 | N/A | N/A | N/A | N/A | N/A |
| PA3703 | N/A | N/A | N/A | PA3703 | N/A | N/A | N/A | N/A | N/A | N/A | N/A | N/A | N/A | N/A | N/A | N/A | N/A | N/A |
| PA3704 | N/A | N/A | N/A | PA3704 | N/A | N/A | N/A | N/A | PA3704 | PA3704 | N/A | PA3704 | PA3704 | PA3704 | N/A | N/A | PA3704 | N/A |
| PA3705 | N/A | N/A | N/A | PA3705 | N/A | N/A | N/A | N/A | N/A | N/A | N/A | N/A | N/A | N/A | N/A | N/A | N/A | N/A |
| N/A | N/A | N/A | N/A | PA3706 | N/A | N/A | N/A | N/A | N/A | N/A | N/A | N/A | N/A | N/A | N/A | N/A | N/A | N/A |
| PA3707 | N/A | N/A | N/A | PA3707 | N/A | PA3707 | N/A | N/A | PA3707 | PA3707 | PA3707 | PA3707 | PA3707 | N/A | N/A | N/A | N/A | N/A |
| PA3708 | PA3708 | PA3708 | PA3708 | PA3708 | N/A | PA3708 | N/A | PA3708 | PA3708 | PA3708 | PA3708 | PA3708 | PA3708 | PA3708 | PA3708 | N/A | PA3708 | N/A |
| PA3709 | N/A | N/A | N/A | N/A | N/A | N/A | N/A | N/A | N/A | N/A | N/A | PA3709 | PA3709 | PA3709 | N/A | N/A | N/A | N/A |
| PA3710 | N/A | N/A | N/A | N/A | N/A | N/A | N/A | N/A | N/A | N/A | N/A | PA3710 | PA3710 | N/A | N/A | N/A | N/A | N/A |
| N/A | N/A | N/A | N/A | N/A | PA3711 | N/A | N/A | N/A | N/A | N/A | N/A | N/A | N/A | N/A | N/A | N/A | N/A | N/A |
| PA3712 | PA3712 | N/A | N/A | N/A | PA3712 | PA3712 | PA3712 | N/A | PA3712 | PA3712 | PA3712 | PA3712 | PA3712 | PA3712 | N/A | N/A | PA3712 | N/A |
| PA3713 | PA3713 | PA3713 | PA3713 | PA3713 | PA3713 | PA3713 | N/A | PA3713 | PA3713 | N/A | PA3713 | PA3713 | PA3713 | PA3713 | PA3713 | N/A | N/A | N/A |
| N/A | N/A | N/A | N/A | N/A | PA3714 | N/A | N/A | N/A | N/A | N/A | N/A | N/A | N/A | N/A | N/A | N/A | N/A | N/A |
| N/A | N/A | N/A | N/A | PA3715 | PA3715 | N/A | PA3715 | N/A | N/A | N/A | N/A | N/A | N/A | N/A | PA3715 | N/A | PA3715 | N/A |
| PA3716 | PA3716 | PA3716 | PA3716 | PA3716 | N/A | N/A | PA3716 | PA3716 | PA3716 | PA3716 | PA3716 | PA3716 | PA3716 | PA3716 | PA3716 | PA3716 | PA3716 | PA3716 |
| PA3717 | PA3717 | N/A | N/A | PA3717 | N/A | N/A | N/A | N/A | PA3717 | N/A | PA3717 | PA3717 | PA3717 | N/A | N/A | N/A | N/A | N/A |
| PA3720 | N/A | N/A | N/A | PA3720 | N/A | N/A | N/A | N/A | PA3720 | PA3720 | N/A | PA3720 | N/A | N/A | N/A | N/A | N/A | N/A |
| PA3721 | N/A | N/A | N/A | PA3721 | N/A | N/A | N/A | N/A | PA3721 | N/A | N/A | N/A | N/A | N/A | N/A | N/A | N/A | N/A |
| N/A | N/A | N/A | N/A | N/A | PA3722 | N/A | N/A | N/A | N/A | N/A | N/A | N/A | N/A | N/A | N/A | N/A | N/A | N/A |
| PA3723 | N/A | N/A | N/A | PA3723 | PA3723 | PA3723 | N/A | N/A | PA3723 | PA3723 | PA3723 | PA3723 | PA3723 | N/A | N/A | N/A | N/A | N/A |
| PA3724 | N/A | N/A | N/A | N/A | PA3724 | PA3724 | PA3724 | PA3724 | PA3724 | PA3724 | PA3724 | PA3724 | PA3724 | N/A | PA3724 | PA3724 | N/A | N/A |
| PA3725 | PA3725 | N/A | N/A | PA3725 | PA3725 | N/A | N/A | PA3725 | PA3725 | PA3725 | PA3725 | PA3725 | PA3725 | N/A | N/A | N/A | N/A | N/A |
| PA3726 | PA3726 | PA3726 | PA3726 | PA3726 | PA3726 | N/A | N/A | N/A | PA3726 | PA3726 | PA3726 | PA3726 | PA3726 | N/A | PA3726 | N/A | N/A | N/A |
| PA3727 | PA3727 | N/A | N/A | PA3727 | PA3727 | N/A | N/A | N/A | N/A | N/A | PA3727 | N/A | N/A | N/A | N/A | N/A | N/A | N/A |
| PA3728 | PA3728 | PA3728 | PA3728 | PA3728 | N/A | N/A | N/A | PA3728 | PA3728 | N/A | PA3728 | PA3728 | PA3728 | N/A | N/A | N/A | PA3728 | N/A |
| PA3729 | PA3729 | PA3729 | PA3729 | PA3729 | N/A | PA3729 | N/A | PA3729 | PA3729 | PA3729 | PA3729 | PA3729 | PA3729 | PA3729 | PA3729 | N/A | PA3729 | PA3729 |
| PA3730 | PA3730 | PA3730 | PA3730 | PA3730 | PA3730 | N/A | N/A | N/A | PA3730 | N/A | PA3730 | N/A | N/A | PA3730 | N/A | N/A | PA3730 | N/A |
| PA3731 | PA3731 | PA3731 | PA3731 | PA3731 | PA3731 | PA3731 | PA3731 | PA3731 | PA3731 | PA3731 | PA3731 | PA3731 | PA3731 | PA3731 | N/A | N/A | PA3731 | PA3731 |
| PA3732 | PA3732 | PA3732 | PA3732 | PA3732 | N/A | N/A | PA3732 | N/A | PA3732 | PA3732 | PA3732 | PA3732 | PA3732 | N/A | PA3732 | PA3732 | N/A | N/A |
| PA3733 | N/A | N/A | N/A | PA3733 | PA3733 | N/A | N/A | N/A | PA3733 | N/A | N/A | N/A | PA3733 | N/A | N/A | N/A | N/A | N/A |
| PA3734 | N/A | N/A | N/A | PA3734 | PA3734 | PA3734 | N/A | N/A | N/A | N/A | N/A | N/A | N/A | N/A | N/A | PA3734 | N/A | N/A |
| PA3735 | PA3735 | PA3735 | PA3735 | PA3735 | PA3735 | PA3735 | PA3735 | PA3735 | PA3735 | PA3735 | PA3735 | PA3735 | PA3735 | PA3735 | PA3735 | N/A | N/A | N/A |
| PA3736 | PA3736 | PA3736 | PA3736 | PA3736 | PA3736 | PA3736 | PA3736 | PA3736 | PA3736 | PA3736 | PA3736 | PA3736 | PA3736 | PA3736 | PA3736 | N/A | PA3736 | N/A |
| PA3737 | PA3737 | PA3737 | PA3737 | PA3737 | PA3737 | N/A | PA3737 | N/A | PA3737 | PA3737 | PA3737 | PA3737 | PA3737 | PA3737 | PA3737 | N/A | PA3737 | N/A |
| PA3738 | N/A | N/A | PA3738 | PA3738 | N/A | N/A | N/A | N/A | N/A | N/A | N/A | N/A | N/A | N/A | N/A | N/A | N/A | N/A |
| PA3739 | N/A | N/A | N/A | PA3739 | N/A | N/A | N/A | N/A | N/A | PA3739 | PA3739 | PA3739 | PA3739 | PA3739 | N/A | N/A | PA3739 | N/A |
| PA3740 | N/A | N/A | N/A | N/A | N/A | N/A | N/A | N/A | N/A | N/A | N/A | N/A | N/A | N/A | N/A | N/A | PA3740 | N/A |
| N/A | N/A | N/A | N/A | N/A | PA3741 | N/A | N/A | N/A | PA3741 | PA3741 | N/A | N/A | N/A | N/A | N/A | N/A | N/A | N/A |
| PA3742 | PA3742 | PA3742 | PA3742 | PA3742 | PA3742 | PA3742 | PA3742 | PA3742 | PA3742 | PA3742 | PA3742 | PA3742 | PA3742 | PA3742 | PA3742 | PA3742 | PA3742 | PA3742 |
| PA3743 | PA3743 | PA3743 | PA3743 | PA3743 | N/A | N/A | N/A | N/A | PA3743 | PA3743 | PA3743 | N/A | N/A | N/A | N/A | N/A | N/A | N/A |
| PA3744 | N/A | PA3744 | N/A | N/A | PA3744 | PA3744 | PA3744 | PA3744 | PA3744 | PA3744 | PA3744 | PA3744 | PA3744 | PA3744 | N/A | N/A | N/A | N/A |
| PA3745 | PA3745 | PA3745 | PA3745 | PA3745 | PA3745 | PA3745 | PA3745 | PA3745 | PA3745 | PA3745 | PA3745 | PA3745 | PA3745 | PA3745 | PA3745 | PA3745 | PA3745 | PA3745 |
| PA3746 | PA3746 | PA3746 | PA3746 | PA3746 | N/A | PA3746 | PA3746 | PA3746 | PA3746 | PA3746 | PA3746 | PA3746 | PA3746 | PA3746 | PA3746 | N/A | PA3746 | PA3746 |
| PA3748 | N/A | N/A | N/A | PA3748 | N/A | N/A | N/A | PA3748 | PA3748 | PA3748 | PA3748 | PA3748 | PA3748 | PA3748 | N/A | N/A | N/A | N/A |
| N/A | N/A | N/A | N/A | N/A | PA3750 | N/A | N/A | N/A | N/A | N/A | N/A | N/A | N/A | N/A | N/A | N/A | N/A | N/A |
| PA3751 | PA3751 | PA3751 | PA3751 | PA3751 | PA3751 | PA3751 | N/A | N/A | PA3751 | PA3751 | PA3751 | PA3751 | PA3751 | N/A | PA3751 | N/A | N/A | N/A |
| N/A | N/A | N/A | N/A | N/A | PA3752 | N/A | N/A | N/A | N/A | N/A | N/A | N/A | N/A | N/A | N/A | N/A | N/A | N/A |
| PA3753 | N/A | N/A | N/A | N/A | PA3753 | PA3753 | N/A | N/A | PA3753 | PA3753 | PA3753 | PA3753 | PA3753 | N/A | N/A | N/A | N/A | N/A |
| PA3754 | N/A | N/A | N/A | PA3754 | N/A | N/A | N/A | N/A | PA3754 | N/A | PA3754 | PA3754 | PA3754 | N/A | N/A | N/A | N/A | N/A |
| N/A | N/A | N/A | N/A | N/A | N/A | N/A | N/A | N/A | PA3755 | N/A | PA3755 | PA3755 | PA3755 | N/A | N/A | N/A | N/A | N/A |
| N/A | N/A | N/A | N/A | PA3756 | N/A | N/A | N/A | N/A | PA3756 | N/A | PA3756 | N/A | PA3756 | N/A | N/A | N/A | N/A | N/A |
| N/A | N/A | N/A | N/A | PA3757 | N/A | N/A | N/A | N/A | N/A | N/A | N/A | N/A | N/A | N/A | N/A | N/A | N/A | N/A |
| PA3758 | N/A | N/A | N/A | PA3758 | N/A | N/A | N/A | N/A | N/A | PA3758 | N/A | PA3758 | N/A | N/A | N/A | N/A | N/A | N/A |
| PA3759 | N/A | N/A | N/A | PA3759 | N/A | N/A | N/A | N/A | N/A | N/A | PA3759 | N/A | N/A | N/A | N/A | N/A | PA3759 | N/A |
| PA3760 | N/A | N/A | N/A | PA3760 | N/A | N/A | N/A | PA3760 | PA3760 | N/A | PA3760 | PA3760 | PA3760 | PA3760 | N/A | N/A | N/A | N/A |
| PA3761 | PA3761 | N/A | N/A | PA3761 | N/A | N/A | N/A | N/A | PA3761 | N/A | N/A | N/A | N/A | N/A | PA3761 | N/A | PA3761 | PA3761 |
| PA3762 | N/A | N/A | N/A | N/A | PA3762 | N/A | N/A | N/A | PA3762 | N/A | PA3762 | PA3762 | PA3762 | N/A | N/A | N/A | N/A | N/A |
| PA3763 | PA3763 | PA3763 | PA3763 | PA3763 | PA3763 | PA3763 | PA3763 | PA3763 | PA3763 | PA3763 | PA3763 | PA3763 | PA3763 | PA3763 | PA3763 | N/A | N/A | N/A |
| PA3764 | N/A | N/A | N/A | PA3764 | N/A | N/A | PA3764 | N/A | PA3764 | PA3764 | N/A | PA3764 | PA3764 | PA3764 | N/A | N/A | PA3764 | N/A |
| PA3765 | N/A | N/A | N/A | PA3765 | N/A | PA3765 | N/A | N/A | PA3765 | PA3765 | N/A | N/A | N/A | N/A | N/A | N/A | N/A | N/A |
| PA3766 | N/A | N/A | PA3766 | N/A | N/A | N/A | N/A | PA3766 | N/A | N/A | N/A | N/A | N/A | PA3766 | N/A | N/A | PA3766 | N/A |
| PA3767 | N/A | N/A | N/A | PA3767 | N/A | N/A | N/A | N/A | PA3767 | N/A | PA3767 | PA3767 | N/A | N/A | N/A | N/A | N/A | N/A |
| PA3768 | PA3768 | PA3768 | N/A | PA3768 | PA3768 | N/A | PA3768 | PA3768 | PA3768 | PA3768 | N/A | PA3768 | PA3768 | PA3768 | N/A | N/A | PA3768 | N/A |
| PA3769 | PA3769 | PA3769 | PA3769 | PA3769 | PA3769 | PA3769 | N/A | PA3769 | PA3769 | PA3769 | PA3769 | PA3769 | PA3769 | PA3769 | PA3769 | N/A | N/A | N/A |
| PA3770 | PA3770 | PA3770 | PA3770 | PA3770 | N/A | PA3770 | PA3770 | PA3770 | PA3770 | PA3770 | PA3770 | PA3770 | PA3770 | PA3770 | PA3770 | PA3770 | PA3770 | PA3770 |
| PA3777 | N/A | PA3777 | N/A | PA3777 | N/A | N/A | N/A | N/A | PA3777 | N/A | N/A | N/A | PA3777 | N/A | N/A | N/A | N/A | N/A |
| N/A | N/A | N/A | N/A | N/A | PA3778 | N/A | N/A | N/A | N/A | N/A | N/A | N/A | N/A | PA3778 | N/A | N/A | N/A | N/A |
| PA3779 | PA3779 | N/A | N/A | PA3779 | PA3779 | N/A | N/A | N/A | N/A | N/A | PA3779 | N/A | N/A | N/A | PA3779 | N/A | N/A | N/A |
| N/A | N/A | N/A | N/A | PA3783 | N/A | N/A | N/A | N/A | N/A | N/A | N/A | N/A | N/A | N/A | N/A | N/A | N/A | N/A |
| PA3784 | N/A | N/A | N/A | N/A | PA3784 | N/A | N/A | N/A | N/A | N/A | N/A | N/A | N/A | N/A | N/A | N/A | N/A | N/A |
| PA3785 | N/A | N/A | PA3785 | PA3785 | PA3785 | PA3785 | N/A | N/A | PA3785 | PA3785 | PA3785 | PA3785 | PA3785 | N/A | PA3785 | N/A | N/A | N/A |
| N/A | N/A | N/A | N/A | N/A | PA3786 | N/A | N/A | N/A | N/A | N/A | N/A | N/A | N/A | N/A | N/A | N/A | N/A | N/A |
| PA3787 | PA3787 | N/A | N/A | PA3787 | PA3787 | N/A | N/A | N/A | N/A | N/A | PA3787 | N/A | PA3787 | PA3787 | PA3787 | N/A | PA3787 | PA3787 |
| PA3789 | N/A | N/A | N/A | PA3789 | PA3789 | N/A | N/A | N/A | N/A | N/A | N/A | N/A | N/A | PA3789 | N/A | N/A | PA3789 | N/A |
| PA3790 | N/A | PA3790 | N/A | PA3790 | N/A | PA3790 | N/A | PA3790 | PA3790 | PA3790 | N/A | PA3790 | PA3790 | PA3790 | PA3790 | N/A | N/A | N/A |
| N/A | N/A | N/A | N/A | N/A | PA3791 | N/A | N/A | N/A | N/A | N/A | N/A | N/A | N/A | N/A | N/A | N/A | N/A | N/A |
| PA3792 | PA3792 | PA3792 | PA3792 | PA3792 | PA3792 | PA3792 | N/A | PA3792 | PA3792 | PA3792 | PA3792 | PA3792 | PA3792 | PA3792 | PA3792 | PA3792 | N/A | N/A |
| PA3793 | N/A | PA3793 | N/A | PA3793 | PA3793 | N/A | N/A | N/A | PA3793 | PA3793 | N/A | PA3793 | PA3793 | N/A | PA3793 | N/A | N/A | N/A |
| PA3794 | N/A | N/A | N/A | N/A | PA3794 | PA3794 | N/A | N/A | N/A | PA3794 | PA3794 | N/A | N/A | PA3794 | N/A | N/A | N/A | N/A |
| PA3795 | N/A | N/A | N/A | PA3795 | PA3795 | PA3795 | N/A | N/A | PA3795 | PA3795 | PA3795 | PA3795 | PA3795 | N/A | N/A | N/A | N/A | N/A |
| PA3796 | PA3796 | N/A | PA3796 | N/A | PA3796 | N/A | N/A | N/A | PA3796 | PA3796 | PA3796 | PA3796 | PA3796 | N/A | N/A | N/A | N/A | N/A |
| PA3797 | N/A | N/A | N/A | N/A | PA3797 | PA3797 | N/A | N/A | PA3797 | PA3797 | PA3797 | PA3797 | PA3797 | N/A | N/A | N/A | N/A | N/A |
| PA3798 | PA3798 | PA3798 | PA3798 | PA3798 | PA3798 | N/A | PA3798 | PA3798 | PA3798 | PA3798 | PA3798 | PA3798 | PA3798 | N/A | N/A | N/A | N/A | N/A |
| PA3799 | PA3799 | PA3799 | PA3799 | PA3799 | PA3799 | PA3799 | N/A | PA3799 | PA3799 | PA3799 | PA3799 | PA3799 | PA3799 | PA3799 | PA3799 | N/A | PA3799 | PA3799 |
| PA3800 | PA3800 | PA3800 | PA3800 | PA3800 | PA3800 | PA3800 | PA3800 | PA3800 | PA3800 | PA3800 | PA3800 | PA3800 | PA3800 | PA3800 | PA3800 | N/A | PA3800 | N/A |
| PA3801 | PA3801 | PA3801 | PA3801 | PA3801 | PA3801 | PA3801 | PA3801 | PA3801 | PA3801 | PA3801 | PA3801 | PA3801 | PA3801 | PA3801 | PA3801 | PA3801 | PA3801 | PA3801 |
| PA3802 | PA3802 | PA3802 | PA3802 | PA3802 | PA3802 | PA3802 | PA3802 | PA3802 | PA3802 | PA3802 | PA3802 | PA3802 | PA3802 | PA3802 | PA3802 | PA3802 | N/A | N/A |
| PA3803 | PA3803 | PA3803 | PA3803 | PA3803 | PA3803 | PA3803 | N/A | PA3803 | PA3803 | PA3803 | PA3803 | PA3803 | PA3803 | N/A | PA3803 | N/A | PA3803 | N/A |
| PA3804 | PA3804 | PA3804 | PA3804 | PA3804 | N/A | PA3804 | PA3804 | PA3804 | PA3804 | PA3804 | PA3804 | PA3804 | PA3804 | PA3804 | PA3804 | N/A | PA3804 | N/A |
| PA3805 | PA3805 | PA3805 | PA3805 | PA3805 | PA3805 | PA3805 | PA3805 | PA3805 | PA3805 | PA3805 | PA3805 | PA3805 | PA3805 | PA3805 | PA3805 | N/A | PA3805 | PA3805 |
| PA3806 | PA3806 | PA3806 | PA3806 | PA3806 | PA3806 | PA3806 | PA3806 | PA3806 | PA3806 | PA3806 | PA3806 | PA3806 | PA3806 | PA3806 | N/A | N/A | PA3806 | N/A |
| PA3807 | PA3807 | PA3807 | PA3807 | PA3807 | PA3807 | PA3807 | PA3807 | N/A | PA3807 | PA3807 | PA3807 | PA3807 | PA3807 | PA3807 | PA3807 | PA3807 | PA3807 | N/A |
| PA3808 | PA3808 | PA3808 | PA3808 | PA3808 | PA3808 | N/A | N/A | N/A | PA3808 | N/A | PA3808 | PA3808 | PA3808 | N/A | PA3808 | N/A | N/A | N/A |
| PA3809 | N/A | N/A | N/A | PA3809 | PA3809 | PA3809 | N/A | N/A | PA3809 | PA3809 | PA3809 | PA3809 | PA3809 | N/A | N/A | N/A | N/A | N/A |
| PA3810 | PA3810 | PA3810 | PA3810 | PA3810 | PA3810 | N/A | N/A | PA3810 | PA3810 | PA3810 | PA3810 | PA3810 | PA3810 | N/A | PA3810 | N/A | N/A | N/A |
| PA3811 | N/A | PA3811 | N/A | PA3811 | PA3811 | N/A | N/A | N/A | PA3811 | PA3811 | N/A | PA3811 | PA3811 | N/A | N/A | N/A | N/A | N/A |

| | | | | | | | | | | | | | | | | | | |
|---|---|---|---|---|---|---|---|---|---|---|---|---|---|---|---|---|---|---|
| PA3812 | PA3812 | PA3812 | PA3812 | PA3812 | PA3812 | N/A | N/A | N/A | PA3812 | N/A | PA3812 | PA3812 | PA3812 | N/A | PA3812 | N/A | N/A | N/A |
| PA3813 | PA3813 | PA3813 | PA3813 | PA3813 | PA3813 | PA3813 | PA3813 | N/A | PA3813 | PA3813 | PA3813 | PA3813 | PA3813 | PA3813 | PA3813 | N/A | PA3813 | PA3813 |
| PA3814 | PA3814 | PA3814 | PA3814 | PA3814 | PA3814 | PA3814 | PA3814 | PA3814 | PA3814 | PA3814 | PA3814 | PA3814 | PA3814 | PA3814 | PA3814 | N/A | N/A | N/A |
| PA3815 | PA3815 | PA3815 | PA3815 | PA3815 | N/A | PA3815 | PA3815 | N/A | PA3815 | PA3815 | PA3815 | PA3815 | PA3815 | PA3815 | PA3815 | N/A | N/A | N/A |
| PA3816 | PA3816 | PA3816 | PA3816 | PA3816 | PA3816 | PA3816 | PA3816 | N/A | PA3816 | PA3816 | PA3816 | PA3816 | PA3816 | PA3816 | PA3816 | N/A | PA3816 | N/A |
| PA3817 | PA3817 | PA3817 | PA3817 | PA3817 | PA3817 | N/A | N/A | N/A | PA3817 | PA3817 | PA3817 | PA3817 | PA3817 | N/A | PA3817 | N/A | N/A | N/A |
| PA3818 | PA3818 | PA3818 | PA3818 | PA3818 | PA3818 | PA3818 | PA3818 | PA3818 | PA3818 | PA3818 | PA3818 | PA3818 | PA3818 | PA3818 | PA3818 | PA3818 | N/A | PA3818 |
| PA3819 | PA3819 | PA3819 | PA3819 | PA3819 | PA3819 | PA3819 | PA3819 | N/A | PA3819 | PA3819 | PA3819 | PA3819 | PA3819 | PA3819 | PA3819 | N/A | PA3819 | N/A |
| PA3820 | PA3820 | PA3820 | PA3820 | PA3820 | N/A | PA3820 | N/A | PA3820 | PA3820 | PA3820 | PA3820 | PA3820 | PA3820 | PA3820 | PA3820 | N/A | PA3820 | PA3820 |
| PA3821 | PA3821 | PA3821 | PA3821 | PA3821 | PA3821 | PA3821 | N/A | PA3821 | PA3821 | PA3821 | PA3821 | PA3821 | PA3821 | PA3821 | PA3821 | N/A | PA3821 | PA3821 |
| PA3822 | PA3822 | PA3822 | PA3822 | PA3822 | PA3822 | PA3822 | PA3822 | PA3822 | PA3822 | PA3822 | PA3822 | PA3822 | PA3822 | PA3822 | PA3822 | N/A | PA3822 | PA3822 |
| PA3823 | PA3823 | PA3823 | PA3823 | PA3823 | PA3823 | PA3823 | PA3823 | PA3823 | PA3823 | PA3823 | PA3823 | PA3823 | PA3823 | PA3823 | PA3823 | PA3823 | PA3823 | PA3823 |
| PA3824 | PA3824 | PA3824 | PA3824 | PA3824 | PA3824 | N/A | PA3824 | PA3824 | PA3824 | PA3824 | PA3824 | PA3824 | N/A | N/A | PA3824 | N/A | N/A | N/A |
| PA3825 | N/A | N/A | N/A | N/A | N/A | N/A | N/A | N/A | N/A | N/A | N/A | N/A | N/A | PA3825 | N/A | N/A | N/A | N/A |
| PA3826 | N/A | N/A | N/A | N/A | N/A | PA3826 | N/A | N/A | PA3826 | PA3826 | PA3826 | N/A | N/A | PA3826 | PA3826 | N/A | PA3826 | N/A |
| PA3827 | PA3827 | PA3827 | PA3827 | PA3827 | N/A | PA3827 | N/A | N/A | PA3827 | PA3827 | PA3827 | PA3827 | PA3827 | PA3827 | PA3827 | N/A | PA3827 | N/A |
| PA3828 | PA3828 | PA3828 | PA3828 | PA3828 | N/A | PA3828 | N/A | PA3828 | PA3828 | PA3828 | PA3828 | PA3828 | PA3828 | PA3828 | PA3828 | N/A | PA3828 | N/A |
| N/A | N/A | N/A | N/A | N/A | PA3830 | N/A | N/A | N/A | N/A | N/A | N/A | N/A | N/A | N/A | N/A | N/A | N/A | N/A |
| PA3831 | PA3831 | PA3831 | PA3831 | PA3831 | PA3831 | PA3831 | PA3831 | PA3831 | PA3831 | PA3831 | PA3831 | PA3831 | PA3831 | PA3831 | PA3831 | PA3831 | PA3831 | PA3831 |
| PA3832 | N/A | PA3832 | N/A | PA3832 | N/A | N/A | PA3832 | N/A | PA3832 | N/A | PA3832 | PA3832 | N/A | N/A | N/A | N/A | N/A | N/A |
| PA3833 | PA3833 | N/A | N/A | PA3833 | PA3833 | N/A | N/A | N/A | N/A | PA3833 | N/A | N/A | PA3833 | N/A | N/A | N/A | N/A | N/A |
| PA3834 | PA3834 | PA3834 | PA3834 | PA3834 | PA3834 | PA3834 | N/A | PA3834 | PA3834 | PA3834 | PA3834 | PA3834 | PA3834 | PA3834 | PA3834 | N/A | N/A | N/A |
| N/A | N/A | N/A | N/A | N/A | PA3835 | N/A | N/A | N/A | N/A | N/A | N/A | N/A | N/A | N/A | N/A | N/A | N/A | N/A |
| PA3836 | PA3836 | PA3836 | PA3836 | PA3836 | PA3836 | PA3836 | PA3836 | N/A | PA3836 | PA3836 | PA3836 | PA3836 | PA3836 | N/A | PA3836 | PA3836 | N/A | N/A |
| PA3838 | PA3838 | PA3838 | PA3838 | PA3838 | N/A | PA3838 | PA3838 | N/A | PA3838 | N/A | PA3838 | PA3838 | PA3838 | PA3838 | PA3838 | N/A | PA3838 | N/A |
| PA3839 | N/A | N/A | N/A | N/A | N/A | N/A | N/A | N/A | N/A | N/A | N/A | N/A | N/A | PA3839 | N/A | N/A | N/A | N/A |
| PA3840 | N/A | N/A | N/A | PA3840 | N/A | N/A | N/A | PA3840 | N/A | N/A | N/A | N/A | N/A | PA3840 | N/A | N/A | N/A | N/A |
| N/A | N/A | N/A | N/A | PA3841 | N/A | N/A | PA3841 | N/A | N/A | PA3841 | N/A | N/A | N/A | N/A | N/A | PA3841 | N/A | N/A |
| N/A | PA3842 | PA3842 | PA3842 | PA3842 | N/A | N/A | N/A | N/A | PA3842 | PA3842 | N/A | N/A | N/A | N/A | N/A | N/A | N/A | N/A |
| N/A | N/A | N/A | N/A | N/A | PA3843 | N/A | N/A | N/A | N/A | N/A | N/A | N/A | N/A | N/A | N/A | N/A | N/A | N/A |
| PA3844 | N/A | N/A | N/A | N/A | N/A | N/A | N/A | N/A | N/A | PA3844 | N/A | PA3844 | PA3844 | N/A | N/A | N/A | N/A | N/A |
| N/A | N/A | N/A | N/A | N/A | PA3845 | N/A | N/A | N/A | N/A | N/A | N/A | N/A | N/A | N/A | N/A | N/A | N/A | N/A |
| PA3846 | N/A | N/A | N/A | PA3846 | PA3846 | PA3846 | N/A | N/A | PA3846 | PA3846 | PA3846 | PA3846 | PA3846 | N/A | N/A | N/A | N/A | N/A |
| PA3847 | N/A | N/A | N/A | N/A | N/A | N/A | N/A | N/A | N/A | N/A | N/A | N/A | N/A | PA3847 | N/A | N/A | N/A | N/A |
| PA3848 | PA3848 | PA3848 | PA3848 | PA3848 | PA3848 | PA3848 | N/A | N/A | PA3848 | PA3848 | PA3848 | PA3848 | PA3848 | PA3848 | PA3848 | N/A | PA3848 | N/A |
| PA3849 | PA3849 | PA3849 | PA3849 | PA3849 | PA3849 | PA3849 | PA3849 | PA3849 | PA3849 | PA3849 | PA3849 | PA3849 | PA3849 | PA3849 | N/A | N/A | N/A | N/A |
| N/A | N/A | N/A | N/A | PA3850 | N/A | N/A | N/A | N/A | PA3850 | N/A | N/A | N/A | N/A | N/A | N/A | N/A | N/A | N/A |
| PA3851 | N/A | N/A | N/A | N/A | N/A | N/A | N/A | N/A | N/A | N/A | N/A | N/A | N/A | N/A | N/A | N/A | N/A | N/A |
| PA3852 | N/A | N/A | N/A | PA3852 | N/A | PA3852 | N/A | N/A | PA3852 | PA3852 | PA3852 | PA3852 | PA3852 | N/A | N/A | N/A | N/A | N/A |
| PA3853 | N/A | N/A | N/A | PA3853 | N/A | PA3853 | N/A | N/A | N/A | PA3853 | PA3853 | N/A | N/A | PA3853 | N/A | N/A | N/A | N/A |
| PA3854 | N/A | N/A | N/A | PA3854 | N/A | N/A | N/A | N/A | N/A | N/A | N/A | N/A | N/A | N/A | N/A | N/A | N/A | N/A |
| PA3855 | N/A | N/A | N/A | PA3855 | N/A | PA3855 | N/A | N/A | N/A | N/A | N/A | PA3855 | PA3855 | PA3855 | N/A | N/A | PA3855 | N/A |
| PA3856 | N/A | N/A | N/A | PA3856 | N/A | N/A | N/A | N/A | N/A | N/A | N/A | PA3856 | PA3856 | PA3856 | N/A | N/A | PA3856 | N/A |
| N/A | N/A | N/A | N/A | N/A | PA3857 | N/A | N/A | N/A | N/A | N/A | N/A | N/A | N/A | N/A | N/A | N/A | N/A | N/A |
| PA3858 | N/A | N/A | N/A | N/A | PA3858 | N/A | N/A | N/A | N/A | PA3858 | PA3858 | PA3858 | PA3858 | N/A | N/A | N/A | N/A | N/A |
| PA3859 | PA3859 | PA3859 | PA3859 | PA3859 | PA3859 | PA3859 | N/A | N/A | PA3859 | PA3859 | PA3859 | PA3859 | PA3859 | N/A | PA3859 | PA3859 | N/A | N/A |
| PA3860 | N/A | N/A | N/A | N/A | N/A | N/A | N/A | N/A | N/A | N/A | N/A | N/A | PA3860 | N/A | N/A | N/A | N/A | N/A |
| PA3861 | PA3861 | PA3861 | PA3861 | PA3861 | PA3861 | PA3861 | N/A | PA3861 | PA3861 | PA3861 | PA3861 | PA3861 | PA3861 | PA3861 | PA3861 | N/A | N/A | PA3861 |
| PA3862 | PA3862 | PA3862 | PA3862 | PA3862 | PA3862 | N/A | PA3862 | PA3862 | PA3862 | PA3862 | PA3862 | PA3862 | PA3862 | N/A | N/A | N/A | N/A | N/A |
| N/A | N/A | N/A | N/A | PA3863 | N/A | N/A | N/A | PA3863 | N/A | PA3863 | N/A | N/A | N/A | N/A | N/A | N/A | N/A | N/A |
| PA3864 | N/A | N/A | N/A | PA3864 | PA3864 | PA3864 | N/A | N/A | N/A | N/A | N/A | N/A | N/A | N/A | N/A | N/A | N/A | N/A |
| PA3865 | N/A | N/A | N/A | PA3865 | PA3865 | N/A | N/A | N/A | PA3865 | PA3865 | PA3865 | N/A | PA3865 | N/A | PA3865 | N/A | N/A | N/A |
| PA3865.1 | PA3865.1 | PA3865.1 | PA3865.1 | PA3865.1 | N/A | PA3865.1 | PA3865.1 | N/A | PA3865.1 | PA3865.1 | PA3865.1 | PA3865.1 | PA3865.1 | PA3865.1 | PA3865.1 | PA3865.1 | PA3865.1 | N/A |
| PA3866 | N/A | N/A | N/A | PA3866 | N/A | N/A | N/A | PA3866 | N/A | N/A | N/A | PA3866 | PA3866 | PA3866 | PA3866 | N/A | PA3866 | PA3866 |
| N/A | N/A | N/A | N/A | PA3870 | N/A | N/A | N/A | N/A | N/A | N/A | N/A | N/A | N/A | N/A | N/A | N/A | N/A | N/A |
| N/A | N/A | N/A | N/A | N/A | N/A | N/A | N/A | N/A | N/A | N/A | N/A | N/A | N/A | PA3872 | N/A | N/A | PA3872 | N/A |
| PA3874 | N/A | N/A | N/A | N/A | N/A | PA3874 | N/A | N/A | PA3874 | PA3874 | N/A | PA3874 | PA3874 | PA3874 | N/A | N/A | PA3874 | N/A |
| N/A | N/A | N/A | N/A | N/A | PA3873 | N/A | N/A | N/A | N/A | N/A | N/A | N/A | N/A | N/A | N/A | N/A | N/A | N/A |
| PA3875 | N/A | N/A | N/A | PA3875 | PA3875 | PA3875 | N/A | N/A | N/A | PA3875 | PA3875 | PA3875 | PA3875 | PA3875 | N/A | N/A | PA3875 | PA3875 |
| N/A | N/A | N/A | N/A | N/A | N/A | N/A | N/A | N/A | N/A | N/A | N/A | N/A | N/A | PA3876 | N/A | N/A | PA3876 | N/A |
| PA3877 | N/A | N/A | N/A | N/A | N/A | N/A | N/A | N/A | N/A | PA3877 | N/A | N/A | N/A | PA3877 | N/A | N/A | PA3877 | N/A |
| PA3878 | N/A | N/A | N/A | N/A | N/A | PA3878 | N/A | PA3878 | N/A | PA3878 | N/A | PA3878 | PA3878 | PA3878 | N/A | N/A | PA3878 | N/A |
| PA3879 | N/A | PA3879 | N/A | PA3879 | N/A | N/A | N/A | N/A | N/A | PA3879 | PA3879 | PA3879 | PA3879 | N/A | N/A | N/A | N/A | N/A |
| N/A | N/A | N/A | N/A | N/A | N/A | N/A | N/A | N/A | N/A | PA3880 | PA3880 | PA3880 | N/A | PA3880 | N/A | N/A | N/A | N/A |
| PA3881 | N/A | N/A | N/A | N/A | PA3881 | PA3881 | N/A | N/A | PA3881 | PA3881 | PA3881 | PA3881 | N/A | PA3881 | N/A | N/A | PA3881 | N/A |
| PA3882 | N/A | N/A | N/A | N/A | PA3882 | N/A | N/A | N/A | N/A | PA3882 | PA3882 | PA3882 | PA3882 | N/A | N/A | N/A | N/A | N/A |
| PA3883 | N/A | N/A | N/A | PA3883 | N/A | N/A | N/A | N/A | N/A | N/A | N/A | N/A | N/A | PA3883 | N/A | N/A | PA3883 | N/A |
| N/A | N/A | N/A | N/A | N/A | PA3885 | N/A | N/A | N/A | N/A | N/A | N/A | N/A | N/A | N/A | N/A | N/A | N/A | N/A |
| PA3886 | PA3886 | N/A | PA3886 | PA3886 | PA3886 | N/A | N/A | N/A | N/A | PA3886 | PA3886 | N/A | PA3886 | N/A | N/A | N/A | N/A | N/A |
| PA3887 | PA3887 | PA3887 | PA3887 | PA3887 | N/A | PA3887 | N/A | N/A | N/A | PA3887 | N/A | PA3887 | N/A | PA3887 | N/A | N/A | PA3887 | N/A |
| PA3888 | N/A | N/A | N/A | N/A | PA3888 | N/A | N/A | N/A | N/A | N/A | N/A | N/A | N/A | N/A | N/A | N/A | N/A | N/A |
| PA3889 | N/A | N/A | N/A | N/A | PA3889 | N/A | N/A | N/A | N/A | N/A | N/A | N/A | N/A | N/A | N/A | N/A | N/A | N/A |
| PA3890 | N/A | N/A | N/A | N/A | N/A | N/A | N/A | N/A | N/A | N/A | N/A | N/A | N/A | N/A | N/A | N/A | N/A | N/A |
| PA3891 | N/A | N/A | N/A | N/A | N/A | N/A | N/A | N/A | N/A | N/A | N/A | PA3891 | PA3891 | PA3891 | N/A | N/A | PA3891 | N/A |
| N/A | N/A | N/A | N/A | PA3892 | N/A | N/A | N/A | N/A | N/A | N/A | N/A | N/A | N/A | N/A | PA3892 | N/A | N/A | N/A |
| N/A | N/A | N/A | N/A | PA3893 | N/A | N/A | N/A | N/A | N/A | N/A | N/A | N/A | N/A | N/A | N/A | N/A | N/A | N/A |
| N/A | N/A | N/A | N/A | PA3894 | N/A | N/A | N/A | N/A | N/A | N/A | N/A | N/A | N/A | N/A | PA3894 | N/A | N/A | N/A |
| PA3895 | PA3895 | N/A | N/A | PA3895 | N/A | PA3895 | N/A | PA3895 | PA3895 | PA3895 | PA3895 | PA3895 | PA3895 | N/A | N/A | N/A | PA3895 | N/A |
| PA3896 | PA3896 | N/A | PA3896 | PA3896 | N/A | N/A | N/A | PA3896 | PA3896 | N/A | PA3896 | PA3896 | PA3896 | N/A | N/A | N/A | N/A | N/A |
| PA3900 | N/A | N/A | N/A | PA3900 | N/A | N/A | N/A | N/A | N/A | N/A | N/A | N/A | N/A | N/A | N/A | N/A | N/A | N/A |
| PA3901 | PA3901 | PA3901 | PA3901 | PA3901 | PA3901 | PA3901 | N/A | PA3901 | PA3901 | N/A | PA3901 | N/A | PA3901 | PA3901 | PA3901 | N/A | N/A | N/A |
| PA3902 | N/A | N/A | N/A | N/A | PA3902 | N/A | N/A | N/A | PA3902 | PA3902 | N/A | N/A | PA3902 | N/A | PA3902 | N/A | N/A | N/A |
| PA3903 | PA3903 | PA3903 | PA3903 | PA3903 | PA3903 | PA3903 | PA3903 | PA3903 | PA3903 | PA3903 | PA3903 | PA3903 | PA3903 | PA3903 | PA3903 | N/A | N/A | N/A |
| N/A | N/A | N/A | N/A | PA3905 | N/A | N/A | PA3905 | N/A | PA3905 | N/A | N/A | N/A | N/A | N/A | N/A | N/A | N/A | N/A |
| N/A | N/A | N/A | N/A | N/A | N/A | N/A | N/A | N/A | PA3906 | N/A | N/A | N/A | N/A | N/A | N/A | N/A | N/A | N/A |
| N/A | N/A | N/A | N/A | N/A | PA3907 | N/A | N/A | N/A | N/A | N/A | N/A | N/A | N/A | N/A | N/A | N/A | N/A | N/A |
| PA3908 | N/A | N/A | N/A | PA3908 | PA3908 | PA3908 | PA3908 | N/A | PA3908 | PA3908 | PA3908 | PA3908 | PA3908 | N/A | N/A | PA3908 | N/A | N/A |
| PA3911 | N/A | N/A | N/A | N/A | N/A | N/A | N/A | N/A | N/A | PA3911 | PA3911 | N/A | N/A | PA3911 | N/A | N/A | N/A | N/A |
| N/A | N/A | N/A | N/A | N/A | N/A | N/A | N/A | N/A | N/A | PA3912 | N/A | N/A | N/A | N/A | N/A | N/A | PA3912 | N/A |
| N/A | N/A | N/A | N/A | N/A | N/A | N/A | N/A | N/A | N/A | PA3913 | PA3913 | N/A | N/A | N/A | N/A | N/A | PA3913 | N/A |
| PA3914 | N/A | N/A | N/A | N/A | N/A | N/A | N/A | N/A | N/A | N/A | PA3914 | PA3914 | PA3914 | N/A | N/A | N/A | N/A | N/A |
| PA3915 | N/A | N/A | N/A | PA3915 | N/A | N/A | N/A | N/A | N/A | N/A | N/A | N/A | N/A | N/A | N/A | N/A | N/A | N/A |
| N/A | N/A | N/A | N/A | N/A | N/A | N/A | N/A | N/A | N/A | N/A | N/A | PA3916 | PA3916 | N/A | N/A | N/A | N/A | N/A |
| N/A | PA3917 | PA3917 | PA3917 | PA3917 | N/A | N/A | N/A | N/A | PA3917 | N/A | PA3917 | N/A | PA3917 | N/A | PA3917 | N/A | N/A | N/A |
| PA3918 | PA3918 | N/A | PA3918 | PA3918 | PA3918 | PA3918 | PA3918 | N/A | PA3918 | PA3918 | PA3918 | PA3918 | PA3918 | N/A | N/A | N/A | N/A | N/A |
| PA3919 | N/A | N/A | N/A | PA3919 | PA3919 | N/A | N/A | N/A | N/A | PA3919 | N/A | PA3919 | PA3919 | PA3919 | N/A | N/A | N/A | N/A |
| PA3920 | N/A | N/A | N/A | PA3920 | N/A | PA3920 | N/A | N/A | PA3920 | PA3920 | PA3920 | PA3920 | PA3920 | PA3920 | N/A | N/A | N/A | N/A |
| PA3921 | N/A | N/A | N/A | N/A | PA3921 | PA3921 | N/A | N/A | PA3921 | N/A | PA3921 | PA3921 | PA3921 | N/A | N/A | N/A | N/A | N/A |
| PA3922 | N/A | N/A | N/A | N/A | PA3922 | PA3922 | N/A | N/A | PA3922 | PA3922 | PA3922 | PA3922 | PA3922 | PA3922 | N/A | PA3922 | N/A | N/A |
| PA3923 | N/A | N/A | N/A | N/A | N/A | PA3923 | N/A | N/A | PA3923 | PA3923 | PA3923 | PA3923 | PA3923 | N/A | N/A | N/A | N/A | N/A |
| PA3924 | N/A | N/A | N/A | N/A | PA3924 | PA3924 | N/A | N/A | PA3924 | PA3924 | PA3924 | PA3924 | PA3924 | N/A | N/A | N/A | N/A | N/A |
| PA3925 | PA3925 | PA3925 | PA3925 | PA3925 | PA3925 | PA3925 | N/A | N/A | PA3925 | PA3925 | PA3925 | PA3925 | PA3925 | N/A | PA3925 | N/A | N/A | N/A |
| PA3928 | N/A | N/A | N/A | N/A | N/A | N/A | N/A | N/A | N/A | N/A | N/A | N/A | PA3928 | N/A | N/A | N/A | N/A | N/A |
| PA3929 | N/A | N/A | N/A | N/A | N/A | PA3929 | N/A | N/A | N/A | PA3929 | N/A | N/A | PA3929 | PA3929 | N/A | N/A | N/A | N/A |
| PA3930 | N/A | N/A | N/A | N/A | PA3930 | PA3930 | N/A | N/A | N/A | PA3930 | N/A | PA3930 | PA3930 | PA3930 | N/A | N/A | N/A | N/A |
| PA3931 | N/A | PA3931 | PA3931 | PA3931 | PA3931 | PA3931 | N/A | N/A | PA3931 | PA3931 | PA3931 | PA3931 | PA3931 | N/A | PA3931 | PA3931 | N/A | N/A |
| PA3932 | N/A | N/A | N/A | N/A | N/A | N/A | N/A | N/A | N/A | N/A | PA3932 | PA3932 | N/A | N/A | N/A | N/A | N/A | N/A |
| N/A | N/A | N/A | N/A | N/A | N/A | N/A | N/A | N/A | N/A | N/A | N/A | N/A | N/A | PA3933 | N/A | N/A | N/A | N/A |
| PA3934 | N/A | N/A | N/A | PA3934 | N/A | PA3934 | N/A | PA3934 | PA3934 | PA3934 | PA3934 | PA3934 | PA3934 | PA3934 | PA3934 | N/A | PA3934 | N/A |

| | | | | | | | | | | | | | | | | | | |
|---|---|---|---|---|---|---|---|---|---|---|---|---|---|---|---|---|---|---|
| PA3935 | N/A | N/A | N/A | N/A | N/A | N/A | N/A | N/A | N/A | N/A | PA3935 | N/A | N/A | N/A | N/A | N/A | N/A | N/A |
| PA3936 | N/A | N/A | N/A | N/A | N/A | N/A | N/A | N/A | N/A | N/A | PA3936 | N/A | N/A | N/A | N/A | N/A | N/A | N/A |
| PA3937 | N/A | N/A | N/A | N/A | PA3937 | N/A | N/A | N/A | N/A | N/A | PA3937 | N/A | N/A | PA3937 | N/A | N/A | N/A | N/A |
| PA3938 | N/A | N/A | N/A | N/A | PA3938 | N/A | N/A | N/A | N/A | N/A | N/A | N/A | N/A | N/A | N/A | N/A | N/A | N/A |
| N/A | N/A | N/A | N/A | N/A | PA3939 | N/A | N/A | N/A | N/A | N/A | N/A | N/A | N/A | N/A | N/A | N/A | N/A | N/A |
| PA3940 | PA3940 | PA3940 | PA3940 | PA3940 | PA3940 | PA3940 | PA3940 | PA3940 | PA3940 | PA3940 | PA3940 | PA3940 | PA3940 | PA3940 | PA3940 | N/A | PA3940 | N/A |
| PA3941 | PA3941 | PA3941 | N/A | PA3941 | PA3941 | N/A | PA3941 | N/A | N/A | N/A | PA3941 | PA3941 | N/A | N/A | N/A | N/A | N/A | N/A |
| PA3942 | PA3942 | PA3942 | N/A | PA3942 | PA3942 | PA3942 | N/A | N/A | PA3942 | PA3942 | PA3942 | PA3942 | N/A | N/A | N/A | N/A | N/A | N/A |
| PA3944 | N/A | N/A | N/A | PA3944 | PA3944 | N/A | N/A | N/A | PA3944 | N/A | PA3944 | PA3944 | N/A | N/A | N/A | N/A | N/A | N/A |
| PA3945 | N/A | N/A | N/A | N/A | N/A | N/A | N/A | N/A | N/A | N/A | N/A | N/A | N/A | N/A | N/A | N/A | N/A | N/A |
| N/A | PA3946 | N/A | N/A | N/A | N/A | N/A | N/A | N/A | N/A | N/A | N/A | N/A | N/A | N/A | N/A | N/A | N/A | N/A |
| PA3948 | PA3948 | PA3948 | PA3948 | PA3948 | N/A | PA3948 | PA3948 | PA3948 | PA3948 | PA3948 | PA3948 | PA3948 | PA3948 | PA3948 | N/A | N/A | PA3948 | N/A |
| PA3949 | N/A | N/A | N/A | PA3949 | PA3949 | N/A | N/A | PA3949 | N/A | N/A | PA3949 | N/A | PA3949 | PA3949 | N/A | N/A | N/A | N/A |
| PA3950 | PA3950 | N/A | PA3950 | PA3950 | PA3950 | N/A | N/A | PA3950 | PA3950 | PA3950 | N/A | PA3950 | PA3950 | PA3950 | N/A | N/A | PA3950 | PA3950 |
| PA3951 | N/A | N/A | N/A | PA3951 | N/A | N/A | N/A | N/A | PA3951 | N/A | PA3951 | N/A | N/A | N/A | N/A | N/A | N/A | N/A |
| PA3952 | PA3952 | N/A | N/A | PA3952 | PA3952 | N/A | N/A | N/A | N/A | PA3952 | PA3952 | PA3952 | PA3952 | N/A | N/A | N/A | N/A | N/A |
| PA3955 | N/A | N/A | N/A | N/A | PA3955 | PA3955 | N/A | N/A | PA3955 | N/A | PA3955 | N/A | N/A | PA3955 | N/A | N/A | N/A | N/A |
| PA3956 | N/A | N/A | N/A | N/A | PA3956 | PA3956 | N/A | N/A | PA3956 | PA3956 | PA3956 | PA3956 | PA3956 | N/A | N/A | N/A | N/A | N/A |
| PA3957 | N/A | N/A | N/A | N/A | N/A | N/A | N/A | N/A | N/A | N/A | N/A | N/A | N/A | N/A | N/A | N/A | N/A | N/A |
| PA3958 | N/A | N/A | N/A | PA3958 | N/A | N/A | PA3958 | N/A | N/A | PA3958 | N/A | N/A | N/A | PA3958 | N/A | N/A | PA3958 | N/A |
| PA3961 | N/A | N/A | N/A | PA3961 | N/A | N/A | N/A | PA3961 | PA3961 | PA3961 | PA3961 | PA3961 | PA3961 | N/A | N/A | N/A | N/A | N/A |
| N/A | N/A | N/A | N/A | N/A | N/A | N/A | N/A | N/A | N/A | N/A | PA3962 | N/A | N/A | N/A | N/A | N/A | N/A | N/A |
| PA3963 | N/A | N/A | N/A | N/A | N/A | N/A | N/A | N/A | N/A | N/A | N/A | N/A | N/A | N/A | N/A | N/A | N/A | N/A |
| N/A | N/A | N/A | N/A | N/A | PA3964 | N/A | N/A | N/A | N/A | N/A | N/A | N/A | N/A | N/A | N/A | N/A | N/A | N/A |
| PA3965 | PA3965 | PA3965 | PA3965 | PA3965 | N/A | PA3965 | PA3965 | N/A | PA3965 | PA3965 | PA3965 | PA3965 | PA3965 | N/A | PA3965 | N/A | N/A | N/A |
| N/A | N/A | N/A | N/A | PA3966 | PA3966 | N/A | N/A | N/A | PA3966 | N/A | N/A | N/A | N/A | N/A | N/A | N/A | N/A | N/A |
| PA3967 | PA3967 | PA3967 | PA3967 | PA3967 | PA3967 | N/A | PA3967 | N/A | PA3967 | N/A | PA3967 | PA3967 | PA3967 | N/A | N/A | N/A | N/A | N/A |
| PA3968 | N/A | N/A | N/A | PA3968 | N/A | N/A | N/A | N/A | N/A | N/A | N/A | N/A | N/A | PA3968 | N/A | N/A | N/A | N/A |
| PA3969 | N/A | N/A | N/A | N/A | PA3969 | N/A | N/A | N/A | N/A | N/A | N/A | N/A | N/A | N/A | N/A | N/A | N/A | N/A |
| PA3970 | PA3970 | PA3970 | PA3970 | PA3970 | PA3970 | PA3970 | N/A | PA3970 | PA3970 | PA3970 | PA3970 | PA3970 | PA3970 | PA3970 | N/A | N/A | N/A | N/A |
| N/A | N/A | N/A | N/A | N/A | PA3971 | N/A | N/A | N/A | N/A | N/A | N/A | N/A | N/A | N/A | N/A | N/A | N/A | N/A |
| PA3972 | N/A | PA3972 | PA3972 | PA3972 | PA3972 | PA3972 | N/A | N/A | N/A | PA3972 | PA3972 | PA3972 | PA3972 | PA3972 | N/A | N/A | N/A | N/A |
| PA3974 | N/A | N/A | N/A | PA3974 | PA3974 | N/A | N/A | PA3974 | PA3974 | N/A | PA3974 | PA3974 | PA3974 | PA3974 | N/A | N/A | N/A | N/A |
| PA3975 | PA3975 | PA3975 | PA3975 | PA3975 | PA3975 | PA3975 | PA3975 | N/A | PA3975 | PA3975 | PA3975 | PA3975 | PA3975 | N/A | PA3975 | N/A | PA3975 | N/A |
| PA3976 | PA3976 | PA3976 | PA3976 | PA3976 | PA3976 | N/A | N/A | N/A | PA3976 | PA3976 | PA3976 | PA3976 | PA3976 | N/A | N/A | N/A | N/A | N/A |
| PA3977 | PA3977 | PA3977 | PA3977 | PA3977 | PA3977 | PA3977 | PA3977 | PA3977 | PA3977 | PA3977 | PA3977 | PA3977 | PA3977 | N/A | PA3977 | N/A | N/A | N/A |
| PA3978 | N/A | PA3978 | PA3978 | PA3978 | N/A | N/A | N/A | N/A | PA3978 | PA3978 | PA3978 | PA3978 | PA3978 | PA3978 | N/A | N/A | N/A | N/A |
| PA3979 | PA3979 | PA3979 | PA3979 | PA3979 | PA3979 | N/A | N/A | N/A | PA3979 | N/A | PA3979 | PA3979 | PA3979 | N/A | PA3979 | N/A | N/A | N/A |
| PA3980 | PA3980 | PA3980 | PA3980 | PA3980 | PA3980 | PA3980 | N/A | PA3980 | PA3980 | PA3980 | PA3980 | PA3980 | PA3980 | N/A | N/A | N/A | PA3980 | N/A |
| PA3981 | PA3981 | PA3981 | PA3981 | PA3981 | PA3981 | N/A | N/A | PA3981 | PA3981 | PA3981 | PA3981 | PA3981 | PA3981 | PA3981 | PA3981 | N/A | N/A | N/A |
| PA3982 | N/A | PA3982 | N/A | PA3982 | PA3982 | N/A | PA3982 | N/A | N/A | PA3982 | PA3982 | PA3982 | N/A | N/A | PA3982 | N/A | N/A | N/A |
| PA3983 | PA3983 | PA3983 | PA3983 | PA3983 | PA3983 | N/A | N/A | N/A | PA3983 | PA3983 | PA3983 | PA3983 | PA3983 | PA3983 | PA3983 | N/A | N/A | N/A |
| PA3984 | PA3984 | PA3984 | PA3984 | PA3984 | N/A | PA3984 | N/A | PA3984 | PA3984 | PA3984 | PA3984 | PA3984 | PA3984 | PA3984 | PA3984 | N/A | PA3984 | N/A |
| PA3985 | N/A | N/A | N/A | PA3985 | N/A | PA3985 | N/A | N/A | N/A | N/A | N/A | N/A | N/A | PA3985 | N/A | N/A | PA3985 | N/A |
| PA3986 | N/A | N/A | N/A | N/A | PA3986 | N/A | N/A | N/A | N/A | N/A | N/A | N/A | N/A | N/A | N/A | N/A | N/A | N/A |
| PA3987 | PA3987 | PA3987 | PA3987 | PA3987 | N/A | PA3987 | N/A | PA3987 | PA3987 | PA3987 | PA3987 | PA3987 | PA3987 | PA3987 | PA3987 | PA3987 | N/A | N/A |
| PA3988 | PA3988 | PA3988 | PA3988 | PA3988 | N/A | PA3988 | PA3988 | PA3988 | PA3988 | PA3988 | PA3988 | PA3988 | PA3988 | PA3988 | PA3988 | PA3988 | PA3988 | N/A |
| PA3989 | PA3989 | PA3989 | PA3989 | PA3989 | N/A | N/A | PA3989 | PA3989 | PA3989 | PA3989 | PA3989 | PA3989 | PA3989 | PA3989 | N/A | N/A | PA3989 | N/A |
| N/A | N/A | N/A | N/A | N/A | PA3991 | N/A | N/A | N/A | N/A | N/A | N/A | N/A | N/A | N/A | N/A | N/A | N/A | N/A |
| PA3992 | PA3992 | PA3992 | PA3992 | PA3992 | PA3992 | PA3992 | PA3992 | PA3992 | PA3992 | PA3992 | PA3992 | PA3992 | PA3992 | PA3992 | PA3992 | N/A | PA3992 | N/A |
| N/A | N/A | N/A | N/A | N/A | N/A | N/A | N/A | N/A | N/A | N/A | N/A | N/A | PA3994 | N/A | N/A | N/A | N/A | N/A |
| PA3995 | N/A | N/A | N/A | PA3995 | PA3995 | N/A | N/A | N/A | N/A | N/A | N/A | N/A | N/A | N/A | N/A | N/A | N/A | N/A |
| PA3996 | PA3996 | PA3996 | PA3996 | PA3996 | PA3996 | N/A | PA3996 | PA3996 | PA3996 | PA3996 | PA3996 | PA3996 | PA3996 | PA3996 | N/A | N/A | N/A | N/A |
| PA3997 | N/A | N/A | N/A | PA3997 | PA3997 | N/A | PA3997 | N/A | PA3997 | PA3997 | PA3997 | PA3997 | PA3997 | N/A | N/A | N/A | N/A | N/A |
| PA3998 | PA3998 | PA3998 | PA3998 | PA3998 | PA3998 | N/A | N/A | PA3998 | PA3998 | PA3998 | PA3998 | PA3998 | PA3998 | N/A | PA3998 | N/A | N/A | N/A |
| PA3999 | PA3999 | PA3999 | PA3999 | PA3999 | PA3999 | PA3999 | PA3999 | PA3999 | PA3999 | PA3999 | PA3999 | PA3999 | PA3999 | PA3999 | PA3999 | PA3999 | PA3999 | PA3999 |
| PA4000 | PA4000 | PA4000 | PA4000 | PA4000 | PA4000 | N/A | PA4000 | PA4000 | PA4000 | N/A | N/A | N/A | N/A | PA4000 | PA4000 | PA4000 | N/A | PA4000 |
| PA4001 | PA4001 | N/A | N/A | PA4001 | N/A | N/A | PA4001 | N/A | PA4001 | PA4001 | PA4001 | PA4001 | PA4001 | PA4001 | PA4001 | N/A | N/A | N/A |
| PA4002 | N/A | N/A | N/A | N/A | N/A | N/A | N/A | N/A | N/A | N/A | N/A | N/A | N/A | PA4002 | N/A | N/A | N/A | N/A |
| PA4003 | PA4003 | N/A | N/A | PA4003 | N/A | N/A | N/A | PA4003 | PA4003 | PA4003 | N/A | PA4003 | PA4003 | PA4003 | PA4003 | N/A | PA4003 | N/A |
| PA4004 | PA4004 | PA4004 | N/A | PA4004 | PA4004 | N/A | PA4004 | N/A | PA4004 | PA4004 | PA4004 | PA4004 | N/A | PA4004 | PA4004 | N/A | PA4004 | N/A |
| PA4005 | PA4005 | PA4005 | PA4005 | PA4005 | PA4005 | N/A | PA4005 | PA4005 | PA4005 | PA4005 | PA4005 | PA4005 | PA4005 | PA4005 | PA4005 | N/A | N/A | N/A |
| PA4006 | N/A | N/A | PA4006 | PA4006 | PA4006 | N/A | N/A | N/A | PA4006 | PA4006 | PA4006 | PA4006 | N/A | N/A | PA4006 | N/A | N/A | N/A |
| PA4007 | PA4007 | PA4007 | PA4007 | PA4007 | PA4007 | PA4007 | PA4007 | N/A | PA4007 | PA4007 | PA4007 | PA4007 | PA4007 | PA4007 | PA4007 | N/A | N/A | N/A |
| PA4010 | PA4010 | PA4010 | N/A | PA4010 | N/A | PA4010 | N/A | N/A | PA4010 | N/A | PA4010 | N/A | PA4010 | N/A | N/A | N/A | N/A | N/A |
| PA4011 | PA4011 | N/A | N/A | PA4011 | N/A | PA4011 | N/A | PA4011 | PA4011 | PA4011 | PA4011 | PA4011 | PA4011 | PA4011 | N/A | N/A | PA4011 | N/A |
| PA4012 | N/A | N/A | N/A | PA4012 | N/A | N/A | N/A | N/A | PA4012 | PA4012 | PA4012 | PA4012 | PA4012 | N/A | N/A | N/A | N/A | N/A |
| N/A | N/A | N/A | N/A | N/A | PA4014 | N/A | N/A | N/A | N/A | N/A | N/A | N/A | N/A | N/A | N/A | N/A | N/A | N/A |
| PA4015 | PA4015 | PA4015 | PA4015 | PA4015 | PA4015 | PA4015 | N/A | N/A | PA4015 | PA4015 | PA4015 | PA4015 | PA4015 | N/A | N/A | PA4015 | N/A | N/A |
| PA4016 | N/A | N/A | N/A | PA4016 | PA4016 | N/A | N/A | N/A | N/A | N/A | N/A | N/A | N/A | N/A | N/A | N/A | N/A | N/A |
| PA4017 | PA4017 | PA4017 | PA4017 | PA4017 | PA4017 | PA4017 | PA4017 | N/A | PA4017 | PA4017 | PA4017 | PA4017 | PA4017 | N/A | N/A | N/A | N/A | N/A |
| PA4018 | N/A | N/A | N/A | N/A | N/A | N/A | N/A | N/A | N/A | N/A | PA4018 | N/A | N/A | N/A | N/A | N/A | N/A | N/A |
| PA4019 | N/A | PA4019 | N/A | PA4019 | PA4019 | PA4019 | N/A | N/A | PA4019 | PA4019 | N/A | PA4019 | PA4019 | N/A | N/A | N/A | N/A | N/A |
| PA4020 | PA4020 | PA4020 | PA4020 | PA4020 | PA4020 | N/A | N/A | N/A | PA4020 | PA4020 | PA4020 | PA4020 | PA4020 | N/A | PA4020 | N/A | N/A | N/A |
| N/A | N/A | N/A | N/A | N/A | PA4021 | N/A | N/A | N/A | N/A | N/A | N/A | N/A | N/A | N/A | N/A | N/A | N/A | N/A |
| PA4022 | PA4022 | PA4022 | PA4022 | PA4022 | N/A | PA4022 | PA4022 | PA4022 | PA4022 | PA4022 | PA4022 | PA4022 | PA4022 | PA4022 | PA4022 | N/A | N/A | N/A |
| PA4023 | N/A | N/A | N/A | N/A | N/A | N/A | N/A | N/A | N/A | N/A | N/A | N/A | N/A | N/A | N/A | N/A | N/A | N/A |
| PA4024 | N/A | N/A | N/A | PA4024 | N/A | N/A | N/A | N/A | N/A | N/A | PA4024 | PA4024 | PA4024 | N/A | N/A | N/A | N/A | N/A |
| PA4025 | N/A | PA4025 | N/A | PA4025 | PA4025 | PA4025 | N/A | N/A | N/A | N/A | PA4025 | PA4025 | PA4025 | N/A | N/A | N/A | N/A | N/A |
| PA4026 | PA4026 | PA4026 | N/A | PA4026 | PA4026 | PA4026 | N/A | N/A | PA4026 | PA4026 | PA4026 | PA4026 | PA4026 | PA4026 | N/A | N/A | PA4026 | N/A |
| PA4029 | N/A | N/A | N/A | PA4029 | N/A | N/A | N/A | N/A | N/A | PA4029 | N/A | N/A | N/A | PA4029 | PA4029 | N/A | N/A | N/A |
| N/A | N/A | N/A | N/A | N/A | PA4030 | N/A | N/A | N/A | N/A | N/A | N/A | N/A | N/A | PA4030 | N/A | N/A | N/A | N/A |
| PA4031 | PA4031 | PA4031 | PA4031 | PA4031 | PA4031 | PA4031 | PA4031 | N/A | PA4031 | PA4031 | PA4031 | PA4031 | PA4031 | PA4031 | PA4031 | PA4031 | PA4031 | N/A |
| PA4032 | N/A | N/A | N/A | PA4032 | N/A | N/A | N/A | N/A | N/A | N/A | N/A | N/A | N/A | N/A | N/A | N/A | N/A | N/A |
| N/A | N/A | N/A | N/A | N/A | PA4034 | N/A | N/A | N/A | N/A | N/A | N/A | N/A | N/A | N/A | N/A | N/A | N/A | N/A |
| PA4035 | N/A | PA4035 | PA4035 | PA4035 | PA4035 | PA4035 | N/A | N/A | PA4035 | PA4035 | PA4035 | PA4035 | PA4035 | N/A | PA4035 | N/A | N/A | N/A |
| PA4036 | N/A | N/A | N/A | N/A | N/A | N/A | N/A | N/A | N/A | N/A | N/A | N/A | N/A | PA4036 | N/A | N/A | N/A | N/A |
| PA4041 | N/A | N/A | N/A | N/A | N/A | N/A | N/A | N/A | N/A | N/A | N/A | N/A | PA4041 | N/A | N/A | N/A | N/A | N/A |
| PA4042 | PA4042 | PA4042 | PA4042 | PA4042 | PA4042 | N/A | N/A | N/A | PA4042 | PA4042 | PA4042 | PA4042 | PA4042 | N/A | N/A | N/A | N/A | N/A |
| PA4043 | N/A | PA4043 | PA4043 | PA4043 | N/A | N/A | PA4043 | PA4043 | N/A | N/A | PA4043 | N/A | N/A | N/A | PA4043 | N/A | N/A | N/A |
| PA4044 | PA4044 | PA4044 | PA4044 | PA4044 | N/A | N/A | PA4044 | PA4044 | PA4044 | PA4044 | PA4044 | PA4044 | PA4044 | PA4044 | N/A | N/A | PA4044 | N/A |
| PA4045 | N/A | N/A | N/A | PA4045 | N/A | N/A | N/A | N/A | PA4045 | N/A | N/A | N/A | N/A | N/A | N/A | N/A | N/A | N/A |
| PA4046 | N/A | N/A | N/A | PA4046 | PA4046 | N/A | N/A | N/A | PA4046 | N/A | PA4046 | PA4046 | N/A | PA4046 | PA4046 | N/A | PA4046 | N/A |
| PA4047 | PA4047 | PA4047 | PA4047 | PA4047 | PA4047 | PA4047 | PA4047 | N/A | PA4047 | PA4047 | PA4047 | PA4047 | N/A | N/A | N/A | N/A | N/A | N/A |
| PA4048 | N/A | N/A | N/A | PA4048 | N/A | N/A | N/A | N/A | PA4048 | PA4048 | PA4048 | N/A | N/A | N/A | PA4048 | N/A | N/A | N/A |
| PA4049 | N/A | N/A | N/A | N/A | N/A | N/A | N/A | N/A | N/A | N/A | N/A | N/A | PA4049 | N/A | N/A | N/A | N/A | N/A |
| PA4050 | N/A | N/A | N/A | N/A | N/A | N/A | N/A | N/A | N/A | N/A | PA4050 | N/A | N/A | PA4050 | N/A | N/A | N/A | N/A |
| N/A | N/A | N/A | N/A | PA4051 | PA4051 | N/A | N/A | N/A | N/A | N/A | N/A | N/A | N/A | N/A | N/A | N/A | N/A | N/A |
| PA4052 | PA4052 | PA4052 | PA4052 | PA4052 | PA4052 | PA4052 | PA4052 | PA4052 | PA4052 | PA4052 | PA4052 | PA4052 | PA4052 | N/A | PA4052 | N/A | N/A | N/A |
| PA4053 | PA4053 | PA4053 | PA4053 | PA4053 | PA4053 | PA4053 | PA4053 | PA4053 | PA4053 | PA4053 | PA4053 | PA4053 | PA4053 | PA4053 | PA4053 | PA4053 | N/A | N/A |
| PA4054 | PA4054 | PA4054 | PA4054 | PA4054 | PA4054 | PA4054 | PA4054 | PA4054 | PA4054 | PA4054 | PA4054 | PA4054 | PA4054 | N/A | PA4054 | N/A | N/A | N/A |
| PA4055 | PA4055 | PA4055 | PA4055 | PA4055 | PA4055 | PA4055 | PA4055 | N/A | PA4055 | PA4055 | PA4055 | PA4055 | PA4055 | N/A | PA4055 | N/A | N/A | N/A |
| PA4056 | PA4056 | N/A | N/A | PA4056 | PA4056 | N/A | N/A | N/A | N/A | PA4056 | N/A | PA4056 | PA4056 | N/A | N/A | N/A | N/A | N/A |
| PA4057 | PA4057 | PA4057 | PA4057 | PA4057 | PA4057 | PA4057 | PA4057 | N/A | PA4057 | PA4057 | PA4057 | PA4057 | PA4057 | N/A | N/A | N/A | N/A | N/A |
| PA4058 | PA4058 | N/A | PA4058 | PA4058 | PA4058 | PA4058 | PA4058 | N/A | PA4058 | PA4058 | PA4058 | PA4058 | PA4058 | N/A | PA4058 | N/A | N/A | N/A |
| PA4059 | N/A | PA4059 | N/A | PA4059 | N/A | N/A | N/A | N/A | PA4059 | PA4059 | PA4059 | PA4059 | PA4059 | N/A | N/A | N/A | N/A | N/A |
| PA4060 | N/A | N/A | N/A | PA4060 | PA4060 | N/A | N/A | N/A | PA4060 | N/A | N/A | PA4060 | N/A | N/A | N/A | N/A | N/A | N/A |
| PA4061 | PA4061 | PA4061 | PA4061 | PA4061 | PA4061 | PA4061 | PA4061 | PA4061 | PA4061 | PA4061 | PA4061 | PA4061 | PA4061 | N/A | PA4061 | PA4061 | PA4061 | PA4061 |
| PA4062 | N/A | N/A | N/A | PA4062 | PA4062 | N/A | N/A | N/A | PA4062 | PA4062 | PA4062 | PA4062 | PA4062 | N/A | N/A | N/A | N/A | N/A |

| | | | | | | | | | | | | | | | | | | |
|---|---|---|---|---|---|---|---|---|---|---|---|---|---|---|---|---|---|---|
| PA4063 | N/A | N/A | N/A | PA4063 | PA4063 | N/A | N/A | N/A | N/A | PA4063 | N/A | N/A | N/A | N/A | N/A | N/A | N/A | N/A |
| PA4064 | N/A | N/A | N/A | PA4064 | N/A | N/A | N/A | N/A | N/A | N/A | N/A | N/A | N/A | PA4064 | N/A | N/A | N/A | N/A |
| PA4065 | N/A | N/A | N/A | N/A | N/A | N/A | N/A | N/A | N/A | PA4065 | N/A | N/A | N/A | PA4065 | N/A | N/A | N/A | N/A |
| PA4066 | N/A | N/A | N/A | N/A | PA4066 | N/A | N/A | N/A | N/A | N/A | N/A | N/A | N/A | N/A | N/A | N/A | N/A | N/A |
| PA4067 | PA4067 | PA4067 | PA4067 | PA4067 | PA4067 | PA4067 | PA4067 | PA4067 | PA4067 | PA4067 | PA4067 | PA4067 | PA4067 | PA4067 | PA4067 | PA4067 | PA4067 | PA4067 |
| PA4068 | PA4068 | PA4068 | PA4068 | PA4068 | PA4068 | N/A | N/A | N/A | PA4068 | PA4068 | PA4068 | PA4068 | PA4068 | N/A | PA4068 | PA4068 | N/A | N/A |
| PA4069 | N/A | PA4069 | N/A | PA4069 | PA4069 | N/A | N/A | N/A | PA4069 | PA4069 | PA4069 | N/A | PA4069 | N/A | N/A | N/A | N/A | N/A |
| PA4075 | N/A | N/A | N/A | N/A | N/A | PA4075 | N/A | N/A | N/A | N/A | N/A | N/A | N/A | N/A | N/A | N/A | N/A | N/A |
| PA4077 | N/A | N/A | N/A | N/A | PA4077 | N/A | N/A | N/A | N/A | N/A | PA4077 | N/A | N/A | N/A | N/A | N/A | N/A | N/A |
| PA4078 | N/A | N/A | N/A | N/A | PA4078 | PA4078 | N/A | N/A | PA4078 | PA4078 | PA4078 | PA4078 | PA4078 | N/A | N/A | N/A | N/A | N/A |
| PA4079 | PA4079 | PA4079 | PA4079 | PA4079 | PA4079 | PA4079 | N/A | N/A | PA4079 | PA4079 | PA4079 | PA4079 | PA4079 | N/A | N/A | N/A | N/A | N/A |
| PA4080 | N/A | N/A | N/A | N/A | N/A | N/A | N/A | N/A | PA4080 | N/A | N/A | N/A | N/A | N/A | N/A | N/A | N/A | N/A |
| PA4090 | PA4090 | N/A | N/A | PA4090 | N/A | PA4090 | N/A | N/A | PA4090 | PA4090 | PA4090 | PA4090 | PA4090 | N/A | PA4090 | N/A | N/A | N/A |
| PA4101 | N/A | N/A | N/A | N/A | N/A | N/A | N/A | N/A | N/A | N/A | PA4101 | PA4101 | PA4101 | N/A | N/A | N/A | N/A | N/A |
| PA4102 | N/A | N/A | N/A | N/A | N/A | N/A | N/A | N/A | N/A | N/A | N/A | N/A | PA4102 | PA4102 | N/A | N/A | N/A | N/A |
| N/A | N/A | N/A | N/A | N/A | PA4109 | N/A | N/A | N/A | N/A | N/A | N/A | N/A | N/A | N/A | N/A | N/A | N/A | N/A |
| PA4110 | N/A | N/A | N/A | N/A | PA4110 | N/A | N/A | N/A | N/A | N/A | N/A | N/A | N/A | PA4110 | N/A | N/A | N/A | N/A |
| PA4112 | N/A | N/A | N/A | N/A | N/A | N/A | N/A | N/A | N/A | N/A | N/A | PA4112 | PA4112 | N/A | N/A | N/A | PA4112 | N/A |
| PA4114 | N/A | N/A | N/A | N/A | PA4114 | N/A | N/A | N/A | N/A | N/A | PA4114 | N/A | PA4114 | N/A | N/A | N/A | N/A | N/A |
| PA4115 | PA4115 | PA4115 | N/A | PA4115 | PA4115 | N/A | PA4115 | PA4115 | PA4115 | PA4115 | PA4115 | PA4115 | PA4115 | PA4115 | N/A | N/A | N/A | N/A |
| N/A | N/A | N/A | N/A | N/A | N/A | N/A | N/A | N/A | PA4116 | N/A | N/A | N/A | N/A | N/A | N/A | N/A | N/A | N/A |
| PA4117 | N/A | N/A | N/A | PA4117 | N/A | N/A | N/A | N/A | N/A | N/A | N/A | N/A | N/A | N/A | N/A | N/A | N/A | N/A |
| N/A | N/A | N/A | N/A | PA4118 | PA4118 | N/A | N/A | N/A | PA4118 | N/A | PA4118 | N/A | N/A | PA4118 | N/A | N/A | PA4118 | PA4118 |
| N/A | N/A | N/A | N/A | PA4119 | PA4119 | N/A | N/A | N/A | PA4119 | N/A | PA4119 | N/A | N/A | N/A | N/A | N/A | N/A | N/A |
| N/A | N/A | N/A | N/A | N/A | PA4128 | N/A | N/A | N/A | N/A | N/A | N/A | N/A | N/A | N/A | N/A | N/A | N/A | N/A |
| PA4129 | N/A | N/A | N/A | N/A | PA4129 | PA4129 | N/A | N/A | PA4129 | PA4129 | PA4129 | PA4129 | PA4129 | N/A | N/A | N/A | N/A | N/A |
| PA4130 | N/A | N/A | N/A | N/A | PA4130 | PA4130 | N/A | N/A | PA4130 | PA4130 | PA4130 | PA4130 | PA4130 | N/A | N/A | N/A | N/A | N/A |
| PA4131 | N/A | N/A | N/A | N/A | PA4131 | PA4131 | N/A | N/A | PA4131 | PA4131 | PA4131 | PA4131 | PA4131 | PA4131 | N/A | N/A | PA4131 | PA4131 |
| PA4132 | N/A | N/A | N/A | N/A | N/A | PA4132 | N/A | N/A | PA4132 | N/A | PA4132 | PA4132 | PA4132 | PA4132 | N/A | N/A | N/A | N/A |
| PA4133 | N/A | N/A | N/A | N/A | N/A | PA4133 | N/A | N/A | PA4133 | PA4133 | PA4133 | PA4133 | PA4133 | PA4133 | N/A | N/A | PA4133 | N/A |
| PA4134 | N/A | N/A | N/A | N/A | PA4134 | N/A | N/A | N/A | N/A | N/A | PA4134 | N/A | PA4134 | N/A | N/A | N/A | N/A | N/A |
| PA4135 | PA4135 | PA4135 | PA4135 | PA4135 | PA4135 | PA4135 | PA4135 | N/A | PA4135 | PA4135 | PA4135 | PA4135 | PA4135 | N/A | PA4135 | N/A | N/A | N/A |
| N/A | N/A | N/A | N/A | N/A | PA4137 | N/A | N/A | N/A | N/A | N/A | N/A | N/A | N/A | N/A | N/A | N/A | N/A | N/A |
| PA4138 | PA4138 | PA4138 | PA4138 | PA4138 | PA4138 | N/A | N/A | PA4138 | N/A | PA4138 | N/A | N/A | N/A | N/A | N/A | N/A | N/A | N/A |
| PA4139 | N/A | N/A | N/A | N/A | N/A | N/A | N/A | N/A | N/A | N/A | N/A | N/A | N/A | N/A | N/A | N/A | N/A | N/A |
| N/A | N/A | N/A | N/A | N/A | N/A | N/A | N/A | N/A | N/A | N/A | N/A | N/A | N/A | N/A | N/A | PA4140 | N/A | N/A |
| PA4145 | N/A | N/A | N/A | N/A | N/A | N/A | N/A | N/A | N/A | N/A | N/A | N/A | N/A | N/A | N/A | N/A | N/A | N/A |
| N/A | N/A | N/A | N/A | N/A | PA4149 | N/A | N/A | N/A | N/A | N/A | N/A | N/A | N/A | N/A | N/A | N/A | N/A | N/A |
| N/A | N/A | N/A | N/A | N/A | PA4150 | N/A | N/A | N/A | N/A | N/A | N/A | N/A | N/A | N/A | N/A | N/A | N/A | N/A |
| N/A | N/A | N/A | N/A | N/A | PA4151 | N/A | N/A | N/A | N/A | N/A | N/A | N/A | N/A | N/A | N/A | N/A | N/A | N/A |
| N/A | N/A | N/A | N/A | N/A | PA4152 | N/A | N/A | N/A | N/A | N/A | N/A | N/A | N/A | N/A | N/A | N/A | N/A | N/A |
| PA4154 | PA4154 | N/A | PA4154 | PA4154 | N/A | N/A | PA4154 | N/A | PA4154 | N/A | PA4154 | PA4154 | N/A | PA4154 | N/A | N/A | PA4154 | PA4154 |
| PA4156 | N/A | N/A | N/A | N/A | N/A | N/A | N/A | N/A | N/A | N/A | N/A | N/A | N/A | N/A | N/A | N/A | N/A | N/A |
| PA4157 | N/A | N/A | N/A | PA4157 | N/A | N/A | N/A | N/A | N/A | N/A | N/A | N/A | N/A | N/A | N/A | N/A | N/A | N/A |
| N/A | N/A | N/A | N/A | PA4159 | N/A | N/A | N/A | N/A | N/A | N/A | N/A | N/A | N/A | N/A | N/A | N/A | N/A | N/A |
| PA4162 | N/A | N/A | N/A | N/A | PA4162 | N/A | N/A | N/A | N/A | PA4162 | N/A | PA4162 | N/A | N/A | N/A | N/A | N/A | N/A |
| PA4163 | PA4163 | PA4163 | PA4163 | PA4163 | PA4163 | PA4163 | N/A | PA4163 | PA4163 | PA4163 | PA4163 | PA4163 | PA4163 | N/A | PA4163 | N/A | N/A | N/A |
| PA4164 | N/A | N/A | N/A | N/A | N/A | N/A | N/A | N/A | N/A | N/A | N/A | N/A | N/A | N/A | N/A | N/A | N/A | N/A |
| PA4165 | N/A | N/A | N/A | N/A | N/A | N/A | N/A | N/A | N/A | N/A | N/A | N/A | N/A | N/A | N/A | N/A | N/A | N/A |
| PA4168 | N/A | N/A | N/A | PA4168 | N/A | N/A | N/A | N/A | N/A | N/A | N/A | N/A | N/A | N/A | N/A | N/A | N/A | N/A |
| PA4169 | N/A | N/A | N/A | N/A | N/A | N/A | N/A | N/A | PA4169 | PA4169 | PA4169 | PA4169 | PA4169 | N/A | N/A | N/A | N/A | N/A |
| PA4170 | N/A | N/A | N/A | N/A | N/A | PA4170 | N/A | N/A | N/A | N/A | N/A | N/A | N/A | N/A | N/A | N/A | N/A | N/A |
| N/A | N/A | N/A | N/A | N/A | N/A | PA4171 | N/A | N/A | N/A | N/A | N/A | N/A | PA4171 | N/A | N/A | N/A | N/A | N/A |
| PA4174 | N/A | N/A | N/A | PA4174 | PA4174 | N/A | N/A | N/A | N/A | N/A | N/A | N/A | N/A | N/A | N/A | N/A | N/A | N/A |
| N/A | N/A | N/A | N/A | N/A | PA4175 | PA4175 | N/A | N/A | N/A | N/A | N/A | PA4175 | PA4175 | N/A | PA4175 | PA4175 | N/A | N/A |
| PA4176 | PA4176 | PA4176 | PA4176 | PA4176 | PA4176 | PA4176 | PA4176 | N/A | PA4176 | PA4176 | PA4176 | PA4176 | PA4176 | N/A | PA4176 | N/A | N/A | N/A |
| PA4179 | N/A | N/A | N/A | PA4179 | PA4179 | N/A | N/A | PA4179 | PA4179 | N/A | N/A | N/A | N/A | N/A | N/A | N/A | N/A | N/A |
| PA4180 | N/A | N/A | N/A | PA4180 | PA4180 | PA4180 | N/A | N/A | PA4180 | PA4180 | PA4180 | PA4180 | PA4180 | N/A | N/A | N/A | N/A | N/A |
| N/A | N/A | N/A | N/A | PA4181 | N/A | N/A | N/A | N/A | N/A | N/A | N/A | N/A | N/A | N/A | N/A | N/A | N/A | N/A |
| PA4182 | N/A | N/A | N/A | N/A | N/A | N/A | N/A | N/A | N/A | PA4182 | N/A | N/A | PA4182 | N/A | N/A | N/A | N/A | N/A |
| PA4183 | N/A | N/A | N/A | N/A | N/A | N/A | N/A | N/A | N/A | N/A | PA4183 | PA4183 | PA4183 | N/A | N/A | N/A | N/A | N/A |
| PA4184 | N/A | N/A | N/A | N/A | N/A | N/A | N/A | N/A | N/A | N/A | N/A | N/A | N/A | N/A | N/A | N/A | N/A | N/A |
| PA4185 | N/A | N/A | N/A | N/A | N/A | N/A | N/A | N/A | N/A | N/A | N/A | N/A | N/A | N/A | N/A | N/A | N/A | N/A |
| PA4188 | N/A | N/A | N/A | N/A | N/A | N/A | N/A | N/A | N/A | N/A | N/A | PA4188 | PA4188 | N/A | N/A | N/A | N/A | N/A |
| N/A | N/A | N/A | N/A | N/A | PA4189 | N/A | N/A | N/A | N/A | N/A | N/A | N/A | N/A | N/A | N/A | N/A | N/A | N/A |
| PA4190 | N/A | N/A | N/A | PA4190 | PA4190 | PA4190 | N/A | PA4190 | PA4190 | N/A | PA4190 | PA4190 | PA4190 | N/A | N/A | N/A | N/A | N/A |
| N/A | N/A | N/A | N/A | PA4191 | N/A | N/A | N/A | N/A | N/A | N/A | N/A | N/A | N/A | N/A | N/A | N/A | N/A | N/A |
| N/A | N/A | N/A | N/A | N/A | N/A | PA4192 | PA4192 | N/A | PA4192 | PA4192 | PA4192 | PA4192 | PA4192 | PA4192 | N/A | N/A | PA4192 | PA4192 |
| PA4194 | N/A | N/A | N/A | N/A | PA4194 | N/A | N/A | N/A | N/A | N/A | N/A | N/A | N/A | N/A | N/A | N/A | N/A | N/A |
| PA4195 | N/A | N/A | N/A | N/A | PA4195 | N/A | N/A | N/A | PA4195 | N/A | PA4195 | N/A | N/A | N/A | N/A | N/A | N/A | N/A |
| PA4196 | N/A | N/A | N/A | PA4196 | N/A | N/A | N/A | N/A | N/A | N/A | N/A | N/A | N/A | N/A | N/A | N/A | N/A | N/A |
| PA4197 | N/A | N/A | N/A | N/A | N/A | N/A | N/A | PA4197 | PA4197 | N/A | PA4197 | PA4197 | N/A | N/A | N/A | N/A | N/A | N/A |
| PA4198 | PA4198 | N/A | N/A | PA4198 | PA4198 | N/A | N/A | N/A | N/A | N/A | PA4198 | PA4198 | N/A | N/A | N/A | N/A | N/A | N/A |
| PA4199 | PA4199 | PA4199 | PA4199 | PA4199 | PA4199 | PA4199 | N/A | PA4199 | PA4199 | PA4199 | PA4199 | PA4199 | PA4199 | PA4199 | N/A | N/A | N/A | N/A |
| PA4200 | PA4200 | N/A | PA4200 | PA4200 | PA4200 | N/A | PA4200 | N/A | N/A | PA4200 | PA4200 | PA4200 | PA4200 | N/A | N/A | N/A | N/A | N/A |
| PA4201 | N/A | N/A | PA4201 | PA4201 | PA4201 | N/A | N/A | N/A | N/A | PA4201 | PA4201 | PA4201 | PA4201 | N/A | N/A | N/A | N/A | N/A |
| PA4202 | PA4202 | PA4202 | PA4202 | PA4202 | PA4202 | PA4202 | N/A | N/A | PA4202 | PA4202 | PA4202 | PA4202 | PA4202 | N/A | N/A | N/A | N/A | N/A |
| PA4203 | N/A | N/A | N/A | PA4203 | PA4203 | N/A | N/A | N/A | N/A | N/A | N/A | N/A | N/A | N/A | N/A | N/A | N/A | N/A |
| PA4204 | N/A | PA4204 | N/A | PA4204 | PA4204 | N/A | N/A | N/A | N/A | N/A | N/A | N/A | PA4204 | N/A | N/A | N/A | N/A | N/A |
| PA4205 | N/A | N/A | N/A | N/A | PA4205 | PA4205 | N/A | N/A | PA4205 | PA4205 | PA4205 | PA4205 | PA4205 | N/A | N/A | N/A | N/A | N/A |
| PA4206 | N/A | N/A | N/A | N/A | PA4206 | PA4206 | N/A | N/A | N/A | PA4206 | N/A | PA4206 | PA4206 | N/A | N/A | N/A | PA4206 | N/A |
| PA4207 | N/A | N/A | N/A | PA4207 | N/A | PA4207 | N/A | N/A | N/A | PA4207 | PA4207 | PA4207 | PA4207 | PA4207 | N/A | N/A | PA4207 | N/A |
| PA4208 | N/A | N/A | N/A | N/A | PA4208 | PA4208 | N/A | N/A | N/A | N/A | N/A | PA4208 | PA4208 | N/A | N/A | N/A | N/A | N/A |
| PA4209 | N/A | N/A | N/A | N/A | N/A | PA4209 | N/A | N/A | PA4209 | PA4209 | PA4209 | PA4209 | PA4209 | N/A | N/A | N/A | N/A | N/A |
| PA4210 | N/A | N/A | N/A | N/A | PA4210 | N/A | N/A | N/A | N/A | N/A | PA4210 | PA4210 | PA4210 | N/A | N/A | N/A | N/A | N/A |
| PA4211 | N/A | N/A | N/A | N/A | PA4211 | PA4211 | N/A | N/A | PA4211 | PA4211 | PA4211 | PA4211 | PA4211 | N/A | N/A | PA4211 | N/A | N/A |
| PA4212 | N/A | N/A | N/A | N/A | PA4212 | N/A | N/A | N/A | PA4212 | N/A | PA4212 | PA4212 | PA4212 | N/A | N/A | N/A | N/A | N/A |
| PA4213 | N/A | N/A | N/A | N/A | PA4213 | PA4213 | N/A | N/A | PA4213 | PA4213 | PA4213 | PA4213 | PA4213 | N/A | N/A | N/A | N/A | N/A |
| PA4214 | N/A | N/A | N/A | N/A | PA4214 | PA4214 | N/A | N/A | N/A | PA4214 | PA4214 | PA4214 | PA4214 | N/A | N/A | N/A | N/A | N/A |
| PA4215 | PA4215 | PA4215 | N/A | PA4215 | N/A | PA4215 | N/A | N/A | PA4215 | PA4215 | PA4215 | PA4215 | PA4215 | N/A | N/A | PA4215 | N/A | N/A |
| PA4216 | N/A | N/A | N/A | N/A | PA4216 | PA4216 | N/A | N/A | PA4216 | PA4216 | PA4216 | PA4216 | PA4216 | N/A | N/A | N/A | N/A | N/A |
| PA4217 | N/A | N/A | N/A | N/A | PA4217 | PA4217 | N/A | N/A | PA4217 | PA4217 | PA4217 | PA4217 | PA4217 | N/A | N/A | PA4217 | N/A | N/A |
| PA4218 | PA4218 | N/A | PA4218 | N/A | N/A | PA4218 | N/A | N/A | N/A | N/A | N/A | N/A | PA4218 | PA4218 | N/A | N/A | N/A | N/A |
| PA4219 | N/A | N/A | N/A | N/A | N/A | N/A | N/A | N/A | N/A | N/A | N/A | N/A | N/A | N/A | N/A | N/A | N/A | N/A |
| PA4220 | N/A | N/A | N/A | N/A | N/A | PA4220 | N/A | N/A | N/A | N/A | N/A | N/A | N/A | N/A | N/A | N/A | N/A | N/A |
| PA4221 | PA4221 | PA4221 | PA4221 | PA4221 | N/A | PA4221 | N/A | N/A | N/A | PA4221 | N/A | N/A | PA4221 | N/A | N/A | N/A | N/A | N/A |
| PA4222 | N/A | N/A | N/A | N/A | N/A | PA4222 | N/A | N/A | N/A | N/A | N/A | N/A | N/A | N/A | N/A | N/A | N/A | N/A |
| PA4223 | PA4223 | PA4223 | PA4223 | PA4223 | PA4223 | PA4223 | N/A | N/A | N/A | N/A | N/A | N/A | N/A | PA4223 | N/A | N/A | N/A | N/A |
| PA4224 | PA4224 | PA4224 | PA4224 | N/A | PA4224 | N/A | N/A | N/A | N/A | N/A | N/A | N/A | PA4224 | N/A | N/A | N/A | N/A | N/A |
| PA4225 | PA4225 | PA4225 | PA4225 | N/A | PA4225 | N/A | N/A | N/A | N/A | N/A | N/A | N/A | PA4225 | N/A | N/A | N/A | N/A | N/A |
| PA4226 | PA4226 | PA4226 | PA4226 | PA4226 | PA4226 | N/A | N/A | N/A | PA4226 | N/A | N/A | PA4226 | PA4226 | N/A | N/A | N/A | N/A | N/A |
| PA4227 | N/A | N/A | N/A | N/A | PA4227 | N/A | N/A | N/A | N/A | N/A | N/A | N/A | PA4227 | N/A | N/A | N/A | N/A | N/A |
| PA4228 | PA4228 | PA4228 | PA4228 | N/A | PA4228 | N/A | N/A | N/A | N/A | N/A | N/A | N/A | PA4228 | N/A | N/A | N/A | N/A | N/A |
| PA4229 | N/A | N/A | N/A | N/A | PA4229 | N/A | N/A | N/A | N/A | N/A | N/A | N/A | N/A | N/A | N/A | N/A | N/A | N/A |
| PA4230 | PA4230 | PA4230 | N/A | N/A | PA4230 | N/A | N/A | N/A | N/A | N/A | N/A | N/A | PA4230 | N/A | N/A | N/A | N/A | N/A |
| PA4231 | N/A | PA4231 | PA4231 | N/A | N/A | N/A | N/A | N/A | N/A | N/A | N/A | N/A | PA4231 | N/A | N/A | N/A | N/A | N/A |
| PA4232 | PA4232 | PA4232 | PA4232 | PA4232 | PA4232 | PA4232 | PA4232 | PA4232 | PA4232 | PA4232 | PA4232 | PA4232 | PA4232 | PA4232 | PA4232 | PA4232 | PA4232 | PA4232 |
| PA4233 | PA4233 | PA4233 | N/A | PA4233 | PA4233 | N/A | N/A | N/A | PA4233 | PA4233 | PA4233 | PA4233 | PA4233 | PA4233 | PA4233 | N/A | PA4233 | N/A |
| PA4234 | PA4234 | PA4234 | PA4234 | PA4234 | PA4234 | PA4234 | N/A | PA4234 | PA4234 | PA4234 | PA4234 | PA4234 | PA4234 | PA4234 | N/A | N/A | PA4234 | N/A |

| | | | | | | | | | | | | | | | | | | |
|---|---|---|---|---|---|---|---|---|---|---|---|---|---|---|---|---|---|---|
| PA4235 | PA4235 | PA4235 | PA4235 | PA4235 | PA4235 | PA4235 | PA4235 | N/A | PA4235 | PA4235 | PA4235 | PA4235 | PA4235 | PA4235 | PA4235 | PA4235 | N/A | N/A |
| PA4236 | PA4236 | PA4236 | PA4236 | PA4236 | PA4236 | PA4236 | PA4236 | N/A | PA4236 | PA4236 | PA4236 | PA4236 | PA4236 | PA4236 | PA4236 | PA4236 | PA4236 | PA4236 |
| PA4237 | PA4237 | PA4237 | PA4237 | PA4237 | PA4237 | PA4237 | PA4237 | PA4237 | PA4237 | PA4237 | PA4237 | PA4237 | PA4237 | PA4237 | PA4237 | PA4237 | PA4237 | PA4237 |
| PA4238 | PA4238 | PA4238 | PA4238 | PA4238 | PA4238 | PA4238 | PA4238 | PA4238 | PA4238 | PA4238 | PA4238 | PA4238 | PA4238 | PA4238 | PA4238 | PA4238 | PA4238 | N/A |
| PA4239 | PA4239 | PA4239 | PA4239 | PA4239 | PA4239 | PA4239 | PA4239 | PA4239 | PA4239 | PA4239 | PA4239 | PA4239 | PA4239 | PA4239 | PA4239 | PA4239 | PA4239 | PA4239 |
| PA4240 | PA4240 | PA4240 | PA4240 | PA4240 | PA4240 | PA4240 | PA4240 | PA4240 | PA4240 | PA4240 | PA4240 | PA4240 | PA4240 | PA4240 | PA4240 | PA4240 | PA4240 | PA4240 |
| PA4241 | PA4241 | PA4241 | PA4241 | PA4241 | PA4241 | PA4241 | PA4241 | PA4241 | PA4241 | PA4241 | PA4241 | PA4241 | PA4241 | PA4241 | PA4241 | PA4241 | PA4241 | PA4241 |
| N/A | N/A | N/A | N/A | N/A | PA4242 | PA4242 | N/A | N/A | N/A | N/A | PA4242 | N/A | PA4242 | N/A | N/A | N/A | N/A | N/A |
| PA4243 | PA4243 | PA4243 | PA4243 | PA4243 | PA4243 | PA4243 | N/A | PA4243 | PA4243 | PA4243 | PA4243 | PA4243 | PA4243 | PA4243 | PA4243 | N/A | PA4243 | PA4243 |
| PA4244 | PA4244 | PA4244 | PA4244 | PA4244 | PA4244 | PA4244 | PA4244 | PA4244 | PA4244 | PA4244 | PA4244 | PA4244 | PA4244 | PA4244 | PA4244 | PA4244 | PA4244 | PA4244 |
| PA4245 | PA4245 | PA4245 | PA4245 | PA4245 | PA4245 | PA4245 | PA4245 | PA4245 | PA4245 | PA4245 | PA4245 | PA4245 | PA4245 | PA4245 | PA4245 | PA4245 | PA4245 | PA4245 |
| PA4246 | PA4246 | PA4246 | PA4246 | PA4246 | PA4246 | PA4246 | PA4246 | PA4246 | PA4246 | PA4246 | PA4246 | PA4246 | PA4246 | PA4246 | PA4246 | PA4246 | PA4246 | PA4246 |
| PA4247 | PA4247 | PA4247 | PA4247 | PA4247 | PA4247 | PA4247 | PA4247 | PA4247 | PA4247 | PA4247 | PA4247 | PA4247 | PA4247 | PA4247 | PA4247 | PA4247 | PA4247 | PA4247 |
| PA4248 | PA4248 | PA4248 | PA4248 | PA4248 | PA4248 | PA4248 | PA4248 | PA4248 | PA4248 | PA4248 | PA4248 | PA4248 | PA4248 | PA4248 | PA4248 | PA4248 | PA4248 | PA4248 |
| PA4249 | PA4249 | PA4249 | PA4249 | PA4249 | PA4249 | PA4249 | PA4249 | PA4249 | PA4249 | PA4249 | PA4249 | PA4249 | PA4249 | PA4249 | PA4249 | PA4249 | PA4249 | PA4249 |
| PA4250 | PA4250 | PA4250 | PA4250 | PA4250 | PA4250 | PA4250 | PA4250 | PA4250 | PA4250 | PA4250 | PA4250 | PA4250 | PA4250 | PA4250 | PA4250 | PA4250 | PA4250 | PA4250 |
| PA4251 | PA4251 | PA4251 | PA4251 | PA4251 | PA4251 | PA4251 | PA4251 | PA4251 | PA4251 | PA4251 | PA4251 | PA4251 | PA4251 | PA4251 | PA4251 | PA4251 | PA4251 | PA4251 |
| PA4252 | PA4252 | PA4252 | PA4252 | PA4252 | PA4252 | PA4252 | PA4252 | PA4252 | PA4252 | PA4252 | PA4252 | PA4252 | PA4252 | PA4252 | PA4252 | PA4252 | PA4252 | PA4252 |
| PA4253 | PA4253 | PA4253 | PA4253 | PA4253 | PA4253 | PA4253 | PA4253 | PA4253 | PA4253 | PA4253 | PA4253 | PA4253 | PA4253 | PA4253 | PA4253 | PA4253 | PA4253 | PA4253 |
| PA4254 | PA4254 | PA4254 | PA4254 | PA4254 | PA4254 | PA4254 | PA4254 | PA4254 | PA4254 | PA4254 | PA4254 | PA4254 | PA4254 | PA4254 | PA4254 | PA4254 | PA4254 | PA4254 |
| PA4255 | PA4255 | PA4255 | PA4255 | PA4255 | PA4255 | PA4255 | PA4255 | PA4255 | PA4255 | PA4255 | PA4255 | PA4255 | PA4255 | PA4255 | PA4255 | PA4255 | PA4255 | PA4255 |
| PA4256 | PA4256 | PA4256 | PA4256 | PA4256 | PA4256 | PA4256 | PA4256 | PA4256 | PA4256 | PA4256 | PA4256 | PA4256 | PA4256 | PA4256 | PA4256 | PA4256 | PA4256 | PA4256 |
| PA4257 | PA4257 | PA4257 | PA4257 | PA4257 | PA4257 | PA4257 | PA4257 | PA4257 | PA4257 | PA4257 | PA4257 | PA4257 | PA4257 | PA4257 | PA4257 | PA4257 | PA4257 | PA4257 |
| PA4258 | PA4258 | PA4258 | PA4258 | PA4258 | PA4258 | PA4258 | PA4258 | PA4258 | PA4258 | PA4258 | PA4258 | PA4258 | PA4258 | PA4258 | PA4258 | PA4258 | PA4258 | PA4258 |
| PA4259 | PA4259 | PA4259 | PA4259 | PA4259 | PA4259 | PA4259 | PA4259 | PA4259 | PA4259 | PA4259 | PA4259 | PA4259 | PA4259 | PA4259 | PA4259 | PA4259 | PA4259 | PA4259 |
| PA4260 | PA4260 | PA4260 | PA4260 | PA4260 | PA4260 | PA4260 | PA4260 | PA4260 | PA4260 | PA4260 | PA4260 | PA4260 | PA4260 | PA4260 | PA4260 | PA4260 | PA4260 | PA4260 |
| PA4261 | PA4261 | PA4261 | PA4261 | PA4261 | PA4261 | PA4261 | PA4261 | PA4261 | PA4261 | PA4261 | PA4261 | PA4261 | PA4261 | PA4261 | PA4261 | PA4261 | PA4261 | PA4261 |
| PA4262 | PA4262 | PA4262 | PA4262 | PA4262 | PA4262 | PA4262 | PA4262 | PA4262 | PA4262 | PA4262 | PA4262 | PA4262 | PA4262 | PA4262 | PA4262 | PA4262 | PA4262 | PA4262 |
| PA4263 | PA4263 | PA4263 | PA4263 | PA4263 | PA4263 | PA4263 | PA4263 | PA4263 | PA4263 | PA4263 | PA4263 | PA4263 | PA4263 | PA4263 | PA4263 | PA4263 | PA4263 | PA4263 |
| PA4264 | PA4264 | PA4264 | PA4264 | PA4264 | PA4264 | PA4264 | PA4264 | PA4264 | PA4264 | PA4264 | PA4264 | PA4264 | PA4264 | PA4264 | PA4264 | PA4264 | PA4264 | PA4264 |
| PA4265 | PA4265 | PA4265 | PA4265 | PA4265 | PA4265 | PA4265 | PA4265 | PA4265 | PA4265 | PA4265 | PA4265 | PA4265 | PA4265 | PA4265 | PA4265 | PA4265 | PA4265 | PA4265 |
| PA4266 | PA4266 | PA4266 | PA4266 | PA4266 | PA4266 | PA4266 | PA4266 | PA4266 | PA4266 | PA4266 | PA4266 | PA4266 | PA4266 | PA4266 | PA4266 | PA4266 | PA4266 | PA4266 |
| PA4267 | PA4267 | PA4267 | PA4267 | PA4267 | PA4267 | PA4267 | PA4267 | PA4267 | PA4267 | PA4267 | PA4267 | PA4267 | PA4267 | PA4267 | PA4267 | PA4267 | PA4267 | PA4267 |
| PA4268 | PA4268 | PA4268 | PA4268 | PA4268 | PA4268 | PA4268 | PA4268 | PA4268 | PA4268 | PA4268 | PA4268 | PA4268 | PA4268 | PA4268 | PA4268 | PA4268 | PA4268 | PA4268 |
| PA4269 | PA4269 | PA4269 | PA4269 | PA4269 | PA4269 | PA4269 | PA4269 | PA4269 | PA4269 | PA4269 | PA4269 | PA4269 | PA4269 | PA4269 | PA4269 | PA4269 | PA4269 | N/A |
| PA4270 | PA4270 | PA4270 | PA4270 | PA4270 | PA4270 | PA4270 | PA4270 | PA4270 | PA4270 | PA4270 | PA4270 | PA4270 | PA4270 | PA4270 | PA4270 | PA4270 | PA4270 | PA4270 |
| PA4271 | PA4271 | PA4271 | PA4271 | PA4271 | PA4271 | PA4271 | PA4271 | PA4271 | PA4271 | PA4271 | PA4271 | PA4271 | PA4271 | PA4271 | PA4271 | PA4271 | PA4271 | PA4271 |
| PA4272 | PA4272 | PA4272 | PA4272 | PA4272 | PA4272 | PA4272 | PA4272 | PA4272 | PA4272 | PA4272 | PA4272 | PA4272 | PA4272 | PA4272 | PA4272 | PA4272 | PA4272 | PA4272 |
| PA4273 | PA4273 | PA4273 | PA4273 | PA4273 | PA4273 | PA4273 | PA4273 | PA4273 | PA4273 | PA4273 | PA4273 | PA4273 | PA4273 | PA4273 | PA4273 | PA4273 | PA4273 | PA4273 |
| PA4274 | PA4274 | PA4274 | PA4274 | PA4274 | PA4274 | PA4274 | PA4274 | PA4274 | PA4274 | PA4274 | PA4274 | PA4274 | PA4274 | PA4274 | PA4274 | PA4274 | PA4274 | PA4274 |
| PA4275 | PA4275 | PA4275 | PA4275 | PA4275 | PA4275 | PA4275 | PA4275 | PA4275 | PA4275 | PA4275 | PA4275 | PA4275 | PA4275 | PA4275 | PA4275 | N/A | PA4275 | PA4275 |
| PA4276 | PA4276 | PA4276 | PA4276 | PA4276 | PA4276 | PA4276 | N/A | N/A | PA4276 | PA4276 | PA4276 | PA4276 | PA4276 | PA4276 | N/A | N/A | PA4276 | N/A |
| PA4277 | PA4277 | PA4277 | PA4277 | PA4277 | N/A | PA4277 | PA4277 | PA4277 | PA4277 | PA4277 | PA4277 | PA4277 | PA4277 | PA4277 | PA4277 | PA4277 | PA4277 | PA4277 |
| PA4278 | N/A | N/A | N/A | PA4278 | PA4278 | N/A | PA4278 | N/A | PA4278 | PA4278 | PA4278 | PA4278 | N/A | PA4278 | N/A | N/A | PA4278 | N/A |
| PA4279 | PA4279 | PA4279 | N/A | PA4279 | PA4279 | N/A | N/A | N/A | PA4279 | PA4279 | PA4279 | PA4279 | PA4279 | N/A | N/A | N/A | N/A | N/A |
| PA4280 | PA4280 | N/A | N/A | PA4280 | N/A | N/A | N/A | PA4280 | N/A | N/A | N/A | N/A | N/A | N/A | N/A | N/A | N/A | N/A |
| PA4281 | N/A | N/A | N/A | PA4281 | N/A | N/A | N/A | N/A | PA4281 | N/A | N/A | N/A | PA4281 | N/A | N/A | N/A | N/A | N/A |
| PA4282 | N/A | N/A | N/A | PA4282 | N/A | N/A | N/A | N/A | PA4282 | N/A | PA4282 | N/A | N/A | N/A | N/A | N/A | PA4282 | N/A |
| PA4283 | N/A | N/A | N/A | PA4283 | N/A | N/A | N/A | N/A | N/A | N/A | PA4283 | PA4283 | PA4283 | N/A | N/A | N/A | N/A | N/A |
| PA4284 | N/A | N/A | N/A | PA4284 | PA4284 | N/A | N/A | PA4284 | PA4284 | PA4284 | PA4284 | PA4284 | PA4284 | N/A | N/A | N/A | PA4284 | N/A |
| PA4285 | PA4285 | N/A | N/A | PA4285 | PA4285 | N/A | N/A | PA4285 | PA4285 | N/A | PA4285 | PA4285 | PA4285 | N/A | N/A | N/A | N/A | N/A |
| PA4286 | N/A | N/A | N/A | PA4286 | PA4286 | N/A | N/A | N/A | PA4286 | N/A | PA4286 | N/A | N/A | N/A | N/A | N/A | N/A | N/A |
| PA4289 | N/A | N/A | N/A | N/A | N/A | N/A | N/A | N/A | N/A | N/A | N/A | N/A | N/A | N/A | N/A | N/A | N/A | N/A |
| PA4290 | N/A | N/A | PA4290 | PA4290 | N/A | PA4290 | N/A | PA4290 | PA4290 | N/A | PA4290 | PA4290 | N/A | PA4290 | N/A | N/A | PA4290 | N/A |
| N/A | N/A | N/A | N/A | N/A | N/A | N/A | N/A | N/A | N/A | N/A | PA4291 | N/A | N/A | N/A | N/A | N/A | N/A | N/A |
| PA4292 | PA4292 | PA4292 | PA4292 | PA4292 | N/A | N/A | N/A | PA4292 | PA4292 | PA4292 | N/A | PA4292 | PA4292 | PA4292 | PA4292 | N/A | PA4292 | N/A |
| PA4293 | N/A | N/A | N/A | PA4293 | N/A | N/A | N/A | N/A | N/A | N/A | N/A | PA4293 | PA4293 | N/A | PA4293 | N/A | N/A | N/A |
| PA4294 | N/A | N/A | N/A | N/A | N/A | N/A | N/A | N/A | N/A | N/A | N/A | N/A | N/A | N/A | N/A | N/A | N/A | N/A |
| PA4296 | N/A | N/A | N/A | PA4296 | PA4296 | N/A | N/A | N/A | N/A | PA4296 | PA4296 | PA4296 | PA4296 | N/A | N/A | N/A | N/A | N/A |
| PA4297 | N/A | N/A | N/A | N/A | N/A | N/A | N/A | N/A | N/A | N/A | N/A | N/A | N/A | N/A | N/A | N/A | N/A | N/A |
| N/A | N/A | N/A | N/A | N/A | N/A | N/A | N/A | N/A | N/A | N/A | N/A | N/A | PA4298 | N/A | N/A | N/A | N/A | N/A |
| PA4299 | N/A | N/A | N/A | N/A | N/A | N/A | N/A | N/A | N/A | N/A | N/A | PA4299 | PA4299 | N/A | N/A | N/A | N/A | N/A |
| PA4300 | N/A | N/A | N/A | N/A | N/A | N/A | N/A | N/A | N/A | N/A | N/A | N/A | PA4300 | N/A | N/A | N/A | N/A | N/A |
| PA4301 | N/A | N/A | N/A | N/A | N/A | N/A | N/A | N/A | N/A | N/A | N/A | N/A | N/A | N/A | N/A | N/A | N/A | N/A |
| PA4302 | N/A | N/A | N/A | N/A | N/A | N/A | N/A | N/A | PA4302 | N/A | N/A | PA4302 | PA4302 | N/A | N/A | N/A | N/A | N/A |
| PA4303 | N/A | N/A | N/A | N/A | N/A | N/A | N/A | N/A | N/A | N/A | N/A | PA4303 | PA4303 | N/A | N/A | N/A | N/A | N/A |
| PA4304 | N/A | N/A | N/A | N/A | N/A | PA4304 | N/A | N/A | N/A | N/A | N/A | N/A | N/A | N/A | N/A | N/A | N/A | N/A |
| PA4305 | N/A | N/A | N/A | N/A | N/A | N/A | N/A | N/A | N/A | N/A | N/A | N/A | N/A | N/A | N/A | N/A | N/A | N/A |
| PA4306 | N/A | N/A | N/A | N/A | N/A | N/A | N/A | N/A | N/A | N/A | PA4306 | N/A | PA4306 | N/A | N/A | N/A | N/A | N/A |
| PA4307 | PA4307 | PA4307 | PA4307 | PA4307 | PA4307 | PA4307 | PA4307 | PA4307 | PA4307 | PA4307 | PA4307 | PA4307 | PA4307 | PA4307 | PA4307 | N/A | PA4307 | PA4307 |
| PA4308 | N/A | N/A | N/A | PA4308 | PA4308 | PA4308 | N/A | PA4308 | PA4308 | PA4308 | PA4308 | PA4308 | PA4308 | PA4308 | N/A | N/A | PA4308 | N/A |
| PA4309 | PA4309 | PA4309 | PA4309 | PA4309 | PA4309 | PA4309 | PA4309 | PA4309 | PA4309 | PA4309 | PA4309 | PA4309 | PA4309 | PA4309 | PA4309 | N/A | PA4309 | PA4309 |
| PA4310 | PA4310 | PA4310 | PA4310 | PA4310 | N/A | PA4310 | PA4310 | PA4310 | PA4310 | PA4310 | PA4310 | PA4310 | PA4310 | PA4310 | PA4310 | PA4310 | PA4310 | PA4310 |
| PA4311 | N/A | N/A | N/A | N/A | N/A | N/A | N/A | N/A | N/A | N/A | N/A | PA4311 | N/A | N/A | N/A | N/A | N/A | N/A |
| PA4312 | N/A | N/A | N/A | N/A | N/A | N/A | N/A | N/A | N/A | N/A | N/A | PA4312 | PA4312 | N/A | N/A | N/A | N/A | N/A |
| N/A | N/A | N/A | N/A | N/A | PA4313 | N/A | N/A | N/A | N/A | N/A | N/A | N/A | N/A | PA4313 | N/A | N/A | N/A | N/A |
| PA4314 | PA4314 | PA4314 | PA4314 | PA4314 | PA4314 | PA4314 | N/A | PA4314 | PA4314 | PA4314 | PA4314 | PA4314 | PA4314 | PA4314 | PA4314 | PA4314 | N/A | N/A |
| PA4315 | PA4315 | PA4315 | PA4315 | PA4315 | PA4315 | PA4315 | PA4315 | PA4315 | PA4315 | PA4315 | PA4315 | PA4315 | PA4315 | PA4315 | PA4315 | N/A | PA4315 | N/A |
| PA4316 | PA4316 | N/A | N/A | PA4316 | PA4316 | PA4316 | N/A | N/A | PA4316 | PA4316 | N/A | PA4316 | PA4316 | N/A | N/A | N/A | N/A | N/A |
| PA4317 | PA4317 | PA4317 | PA4317 | PA4317 | N/A | PA4317 | N/A | PA4317 | PA4317 | PA4317 | PA4317 | PA4317 | PA4317 | PA4317 | N/A | N/A | PA4317 | PA4317 |
| PA4318 | N/A | N/A | N/A | PA4318 | N/A | N/A | PA4318 | N/A | PA4318 | N/A | N/A | N/A | N/A | PA4318 | N/A | N/A | PA4318 | N/A |
| PA4319 | N/A | N/A | N/A | N/A | N/A | N/A | N/A | PA4319 | N/A | N/A | PA4319 | PA4319 | N/A | PA4319 | N/A | N/A | PA4319 | N/A |
| PA4320 | PA4320 | N/A | PA4320 | PA4320 | N/A | N/A | N/A | PA4320 | N/A | N/A | PA4320 | PA4320 | N/A | PA4320 | PA4320 | N/A | PA4320 | N/A |
| PA4321 | N/A | N/A | N/A | PA4321 | PA4321 | N/A | N/A | PA4321 | N/A | PA4321 | N/A | N/A | N/A | PA4321 | N/A | N/A | PA4321 | N/A |
| PA4322 | PA4322 | PA4322 | PA4322 | PA4322 | N/A | N/A | PA4322 | PA4322 | PA4322 | PA4322 | PA4322 | PA4322 | PA4322 | PA4322 | PA4322 | N/A | PA4322 | PA4322 |
| PA4323 | PA4323 | PA4323 | N/A | PA4323 | N/A | N/A | N/A | PA4323 | N/A | N/A | N/A | N/A | N/A | PA4323 | PA4323 | N/A | PA4323 | N/A |
| N/A | N/A | N/A | N/A | N/A | N/A | N/A | N/A | N/A | N/A | N/A | PA4324 | N/A | N/A | N/A | N/A | N/A | N/A | N/A |
| PA4326 | N/A | N/A | N/A | N/A | N/A | PA4326 | N/A | N/A | N/A | PA4326 | PA4326 | PA4326 | PA4326 | N/A | N/A | N/A | N/A | N/A |
| PA4327 | N/A | N/A | N/A | N/A | PA4327 | N/A | N/A | N/A | N/A | N/A | N/A | N/A | N/A | N/A | N/A | N/A | N/A | N/A |
| PA4328 | N/A | N/A | N/A | PA4328 | PA4328 | N/A | N/A | N/A | N/A | PA4328 | PA4328 | PA4328 | PA4328 | PA4328 | N/A | N/A | N/A | N/A |
| PA4329 | PA4329 | PA4329 | PA4329 | PA4329 | PA4329 | PA4329 | PA4329 | PA4329 | PA4329 | PA4329 | PA4329 | PA4329 | PA4329 | PA4329 | PA4329 | PA4329 | N/A | N/A |
| N/A | N/A | N/A | N/A | PA4330 | N/A | N/A | N/A | N/A | N/A | N/A | N/A | N/A | N/A | N/A | N/A | N/A | N/A | N/A |
| PA4331 | N/A | N/A | N/A | N/A | N/A | N/A | N/A | N/A | N/A | N/A | PA4331 | N/A | N/A | N/A | N/A | N/A | N/A | N/A |
| PA4332 | N/A | N/A | N/A | N/A | PA4332 | N/A | N/A | N/A | N/A | N/A | N/A | N/A | N/A | PA4332 | N/A | N/A | PA4332 | N/A |
| PA4333 | PA4333 | PA4333 | PA4333 | PA4333 | PA4333 | PA4333 | PA4333 | PA4333 | PA4333 | PA4333 | PA4333 | PA4333 | PA4333 | N/A | PA4333 | N/A | N/A | N/A |
| N/A | N/A | N/A | N/A | N/A | PA4335 | N/A | N/A | N/A | N/A | N/A | PA4335 | PA4335 | PA4335 | N/A | N/A | N/A | N/A | N/A |
| PA4336 | PA4336 | PA4336 | PA4336 | PA4336 | PA4336 | PA4336 | N/A | N/A | PA4336 | PA4336 | PA4336 | PA4336 | PA4336 | N/A | PA4336 | N/A | N/A | N/A |
| PA4338 | N/A | N/A | N/A | N/A | N/A | N/A | N/A | N/A | N/A | N/A | N/A | N/A | N/A | PA4338 | N/A | N/A | N/A | N/A |
| PA4339 | N/A | N/A | N/A | N/A | PA4339 | N/A | N/A | N/A | N/A | N/A | N/A | N/A | N/A | PA4339 | N/A | N/A | N/A | N/A |
| PA4340 | N/A | N/A | N/A | PA4340 | PA4340 | N/A | N/A | N/A | PA4340 | N/A | PA4340 | PA4340 | PA4340 | N/A | PA4340 | N/A | N/A | N/A |
| N/A | PA4342 | N/A | N/A | N/A | N/A | N/A | N/A | N/A | N/A | N/A | N/A | N/A | N/A | N/A | PA4342 | N/A | N/A | N/A |
| N/A | N/A | N/A | N/A | N/A | PA4344 | N/A | N/A | N/A | N/A | N/A | N/A | N/A | N/A | N/A | N/A | N/A | N/A | N/A |
| PA4345 | N/A | N/A | N/A | N/A | PA4345 | N/A | N/A | N/A | N/A | PA4345 | N/A | N/A | PA4345 | N/A | N/A | N/A | N/A | N/A |
| PA4346 | N/A | N/A | N/A | N/A | N/A | N/A | N/A | N/A | N/A | N/A | N/A | N/A | N/A | N/A | N/A | N/A | N/A | N/A |
| PA4347 | N/A | N/A | N/A | N/A | PA4347 | N/A | N/A | N/A | N/A | N/A | N/A | N/A | N/A | N/A | N/A | N/A | N/A | N/A |
| PA4348 | N/A | N/A | N/A | N/A | PA4348 | PA4348 | N/A | N/A | PA4348 | PA4348 | PA4348 | PA4348 | PA4348 | N/A | N/A | N/A | N/A | N/A |
| PA4349 | N/A | N/A | N/A | PA4349 | N/A | N/A | N/A | N/A | N/A | PA4349 | PA4349 | PA4349 | PA4349 | N/A | N/A | N/A | N/A | N/A |
| N/A | N/A | N/A | N/A | N/A | PA4351 | N/A | N/A | N/A | N/A | N/A | N/A | N/A | N/A | N/A | N/A | N/A | N/A | N/A |
| PA4352 | PA4352 | PA4352 | N/A | PA4352 | PA4352 | PA4352 | PA4352 | PA4352 | PA4352 | PA4352 | PA4352 | PA4352 | PA4352 | PA4352 | N/A | PA4352 | PA4352 | PA4352 |

| | | | | | | | | | | | | | | | | | | |
|---|---|---|---|---|---|---|---|---|---|---|---|---|---|---|---|---|---|---|
| PA4353 | N/A | N/A | N/A | N/A | N/A | N/A | N/A | N/A | N/A | N/A | N/A | N/A | N/A | N/A | N/A | N/A | N/A | N/A |
| PA4354 | N/A | N/A | N/A | N/A | N/A | N/A | N/A | N/A | N/A | N/A | PA4354 | N/A | PA4354 | N/A | N/A | N/A | N/A | N/A |
| N/A | N/A | N/A | N/A | N/A | PA4355 | N/A | N/A | N/A | N/A | N/A | N/A | N/A | N/A | N/A | N/A | N/A | N/A | N/A |
| PA4356 | PA4356 | PA4356 | PA4356 | PA4356 | PA4356 | N/A | PA4356 | PA4356 | PA4356 | PA4356 | PA4356 | PA4356 | PA4356 | PA4356 | PA4356 | PA4356 | N/A | N/A |
| PA4357 | N/A | N/A | N/A | PA4357 | N/A | N/A | N/A | N/A | N/A | N/A | PA4357 | PA4357 | PA4357 | N/A | N/A | N/A | N/A | N/A |
| PA4358 | N/A | N/A | N/A | N/A | N/A | N/A | N/A | N/A | N/A | N/A | PA4358 | PA4358 | PA4358 | PA4358 | N/A | N/A | N/A | N/A |
| PA4359 | N/A | N/A | N/A | N/A | N/A | N/A | N/A | N/A | N/A | N/A | PA4359 | PA4359 | PA4359 | PA4359 | N/A | N/A | N/A | N/A |
| PA4360 | PA4360 | PA4360 | PA4360 | PA4360 | N/A | N/A | PA4360 | N/A | PA4360 | PA4360 | PA4360 | PA4360 | N/A | PA4360 | PA4360 | N/A | PA4360 | PA4360 |
| PA4361 | N/A | N/A | N/A | PA4361 | PA4361 | PA4361 | N/A | N/A | N/A | N/A | PA4361 | N/A | PA4361 | N/A | N/A | N/A | N/A | N/A |
| PA4362 | N/A | N/A | N/A | N/A | PA4362 | PA4362 | N/A | N/A | N/A | N/A | N/A | PA4362 | PA4362 | PA4362 | N/A | N/A | PA4362 | N/A |
| PA4363 | N/A | N/A | N/A | PA4363 | N/A | N/A | N/A | N/A | N/A | N/A | N/A | N/A | N/A | N/A | N/A | N/A | N/A | N/A |
| N/A | N/A | N/A | N/A | N/A | PA4365 | N/A | N/A | N/A | N/A | N/A | N/A | N/A | N/A | N/A | N/A | N/A | N/A | N/A |
| PA4366 | PA4366 | PA4366 | PA4366 | PA4366 | PA4366 | PA4366 | PA4366 | N/A | PA4366 | PA4366 | PA4366 | PA4366 | PA4366 | PA4366 | PA4366 | PA4366 | PA4366 | N/A |
| PA4367 | N/A | N/A | N/A | PA4367 | N/A | N/A | N/A | N/A | N/A | N/A | PA4367 | PA4367 | PA4367 | PA4367 | N/A | N/A | N/A | N/A |
| PA4368 | N/A | N/A | N/A | N/A | N/A | N/A | N/A | N/A | N/A | N/A | N/A | N/A | N/A | PA4368 | N/A | N/A | N/A | N/A |
| PA4369 | N/A | PA4369 | N/A | PA4369 | PA4369 | PA4369 | PA4369 | N/A | PA4369 | PA4369 | PA4369 | PA4369 | PA4369 | PA4369 | N/A | N/A | PA4369 | N/A |
| PA4370 | PA4370 | PA4370 | PA4370 | PA4370 | PA4370 | PA4370 | PA4370 | PA4370 | PA4370 | PA4370 | PA4370 | PA4370 | PA4370 | PA4370 | PA4370 | PA4370 | PA4370 | PA4370 |
| PA4371 | N/A | N/A | N/A | PA4371 | PA4371 | N/A | PA4371 | PA4371 | N/A | N/A | N/A | N/A | N/A | PA4371 | PA4371 | N/A | N/A | N/A |
| PA4372 | PA4372 | PA4372 | PA4372 | PA4372 | PA4372 | PA4372 | PA4372 | PA4372 | PA4372 | PA4372 | PA4372 | PA4372 | PA4372 | PA4372 | PA4372 | N/A | PA4372 | N/A |
| PA4373 | N/A | N/A | N/A | PA4373 | PA4373 | N/A | PA4373 | PA4373 | N/A | N/A | N/A | N/A | N/A | PA4373 | N/A | N/A | N/A | N/A |
| PA4374 | PA4374 | N/A | N/A | PA4374 | N/A | PA4374 | N/A | PA4374 | N/A | PA4374 | N/A | N/A | N/A | PA4374 | PA4374 | N/A | PA4374 | N/A |
| PA4375 | N/A | N/A | N/A | N/A | PA4375 | PA4375 | N/A | PA4375 | PA4375 | PA4375 | PA4375 | PA4375 | PA4375 | PA4375 | N/A | N/A | PA4375 | N/A |
| PA4376 | PA4376 | PA4376 | PA4376 | PA4376 | PA4376 | PA4376 | PA4376 | PA4376 | PA4376 | PA4376 | PA4376 | PA4376 | PA4376 | N/A | N/A | N/A | N/A | N/A |
| PA4378 | N/A | N/A | N/A | PA4378 | N/A | N/A | N/A | PA4378 | PA4378 | N/A | N/A | N/A | N/A | N/A | N/A | N/A | N/A | N/A |
| PA4380 | N/A | N/A | N/A | N/A | N/A | N/A | N/A | N/A | N/A | N/A | N/A | N/A | N/A | PA4380 | N/A | N/A | N/A | N/A |
| PA4381 | N/A | N/A | N/A | PA4381 | PA4381 | PA4381 | N/A | N/A | PA4381 | PA4381 | PA4381 | PA4381 | PA4381 | N/A | N/A | N/A | N/A | N/A |
| PA4384 | N/A | N/A | N/A | N/A | PA4384 | N/A | N/A | N/A | N/A | N/A | N/A | N/A | PA4384 | N/A | N/A | N/A | N/A | N/A |
| PA4385 | PA4385 | PA4385 | PA4385 | PA4385 | PA4385 | PA4385 | PA4385 | PA4385 | PA4385 | PA4385 | PA4385 | PA4385 | PA4385 | PA4385 | PA4385 | PA4385 | PA4385 | PA4385 |
| PA4386 | PA4386 | PA4386 | PA4386 | PA4386 | N/A | PA4386 | PA4386 | N/A | PA4386 | PA4386 | PA4386 | PA4386 | PA4386 | PA4386 | PA4386 | N/A | PA4386 | PA4386 |
| PA4387 | N/A | N/A | N/A | PA4387 | PA4387 | PA4387 | N/A | N/A | PA4387 | PA4387 | PA4387 | PA4387 | N/A | PA4387 | PA4387 | N/A | N/A | N/A |
| PA4388 | PA4388 | PA4388 | PA4388 | PA4388 | N/A | PA4388 | N/A | N/A | PA4388 | PA4388 | PA4388 | PA4388 | PA4388 | N/A | N/A | N/A | PA4388 | N/A |
| PA4389 | PA4389 | PA4389 | PA4389 | PA4389 | PA4389 | PA4389 | PA4389 | PA4389 | PA4389 | PA4389 | PA4389 | PA4389 | PA4389 | PA4389 | PA4389 | N/A | PA4389 | PA4389 |
| PA4390 | PA4390 | PA4390 | PA4390 | PA4390 | N/A | PA4390 | PA4390 | PA4390 | PA4390 | PA4390 | PA4390 | PA4390 | N/A | N/A | PA4390 | PA4390 | N/A | N/A |
| PA4392 | N/A | N/A | N/A | N/A | N/A | N/A | N/A | N/A | N/A | N/A | PA4392 | N/A | N/A | N/A | N/A | N/A | N/A | N/A |
| PA4393 | N/A | N/A | N/A | N/A | N/A | PA4393 | N/A | PA4393 | PA4393 | PA4393 | PA4393 | PA4393 | PA4393 | PA4393 | N/A | N/A | N/A | N/A |
| PA4394 | PA4394 | PA4394 | PA4394 | PA4394 | PA4394 | PA4394 | N/A | N/A | PA4394 | PA4394 | PA4394 | PA4394 | PA4394 | PA4394 | PA4394 | N/A | PA4394 | N/A |
| PA4395 | PA4395 | PA4395 | PA4395 | PA4395 | N/A | PA4395 | PA4395 | N/A | PA4395 | PA4395 | PA4395 | PA4395 | PA4395 | PA4395 | PA4395 | PA4395 | PA4395 | PA4395 |
| N/A | N/A | N/A | N/A | PA4396 | N/A | N/A | N/A | PA4396 | N/A | PA4396 | N/A | N/A | N/A | N/A | N/A | N/A | N/A | N/A |
| PA4397 | N/A | N/A | N/A | N/A | N/A | N/A | N/A | N/A | N/A | N/A | N/A | N/A | N/A | N/A | N/A | N/A | N/A | N/A |
| PA4398 | N/A | N/A | N/A | PA4398 | N/A | N/A | N/A | PA4398 | N/A | N/A | N/A | N/A | N/A | PA4398 | N/A | N/A | PA4398 | N/A |
| PA4399 | N/A | N/A | N/A | PA4399 | PA4399 | PA4399 | N/A | N/A | PA4399 | PA4399 | PA4399 | PA4399 | N/A | PA4399 | N/A | N/A | N/A | N/A |
| PA4400 | PA4400 | PA4400 | PA4400 | PA4400 | PA4400 | N/A | PA4400 | PA4400 | PA4400 | PA4400 | PA4400 | PA4400 | PA4400 | N/A | N/A | N/A | N/A | N/A |
| PA4401 | N/A | N/A | N/A | N/A | PA4401 | N/A | N/A | N/A | PA4401 | N/A | PA4401 | PA4401 | PA4401 | N/A | N/A | N/A | N/A | N/A |
| PA4402 | PA4402 | PA4402 | PA4402 | PA4402 | PA4402 | PA4402 | PA4402 | N/A | PA4402 | PA4402 | PA4402 | PA4402 | PA4402 | N/A | PA4402 | N/A | N/A | N/A |
| PA4403 | PA4403 | PA4403 | PA4403 | PA4403 | PA4403 | PA4403 | PA4403 | PA4403 | PA4403 | PA4403 | PA4403 | PA4403 | PA4403 | PA4403 | PA4403 | N/A | PA4403 | N/A |
| N/A | N/A | N/A | N/A | N/A | N/A | N/A | N/A | N/A | N/A | N/A | N/A | N/A | N/A | N/A | N/A | N/A | PA4404 | N/A |
| PA4405 | N/A | N/A | N/A | N/A | PA4405 | N/A | N/A | N/A | PA4405 | PA4405 | PA4405 | PA4405 | PA4405 | N/A | N/A | N/A | N/A | N/A |
| PA4406 | PA4406 | PA4406 | PA4406 | PA4406 | PA4406 | PA4406 | PA4406 | PA4406 | PA4406 | PA4406 | PA4406 | PA4406 | PA4406 | PA4406 | PA4406 | N/A | PA4406 | PA4406 |
| PA4407 | PA4407 | PA4407 | PA4407 | PA4407 | PA4407 | PA4407 | PA4407 | PA4407 | PA4407 | PA4407 | PA4407 | PA4407 | PA4407 | PA4407 | PA4407 | N/A | PA4407 | PA4407 |
| PA4408 | PA4408 | PA4408 | PA4408 | PA4408 | PA4408 | PA4408 | PA4408 | PA4408 | PA4408 | PA4408 | PA4408 | PA4408 | PA4408 | PA4408 | PA4408 | N/A | PA4408 | N/A |
| PA4409 | N/A | N/A | N/A | PA4409 | PA4409 | N/A | PA4409 | N/A | N/A | N/A | N/A | N/A | N/A | PA4409 | N/A | N/A | PA4409 | N/A |
| PA4410 | PA4410 | PA4410 | PA4410 | PA4410 | PA4410 | PA4410 | PA4410 | PA4410 | PA4410 | PA4410 | PA4410 | PA4410 | PA4410 | N/A | PA4410 | N/A | N/A | N/A |
| PA4411 | PA4411 | PA4411 | PA4411 | PA4411 | PA4411 | PA4411 | N/A | PA4411 | PA4411 | PA4411 | PA4411 | PA4411 | PA4411 | N/A | PA4411 | N/A | N/A | N/A |
| PA4412 | PA4412 | PA4412 | PA4412 | PA4412 | PA4412 | PA4412 | PA4412 | PA4412 | PA4412 | PA4412 | PA4412 | PA4412 | PA4412 | PA4412 | PA4412 | N/A | PA4412 | N/A |
| PA4413 | N/A | N/A | N/A | N/A | PA4413 | N/A | N/A | N/A | N/A | N/A | N/A | N/A | PA4413 | PA4413 | N/A | N/A | N/A | N/A |
| PA4414 | PA4414 | PA4414 | PA4414 | PA4414 | PA4414 | N/A | N/A | PA4414 | PA4414 | PA4414 | PA4414 | PA4414 | PA4414 | N/A | PA4414 | N/A | N/A | N/A |
| PA4415 | N/A | N/A | N/A | N/A | PA4415 | N/A | N/A | N/A | PA4415 | PA4415 | PA4415 | PA4415 | PA4415 | PA4415 | N/A | N/A | PA4415 | N/A |
| PA4416 | PA4416 | PA4416 | PA4416 | PA4416 | PA4416 | PA4416 | N/A | PA4416 | PA4416 | PA4416 | PA4416 | PA4416 | PA4416 | N/A | PA4416 | N/A | N/A | N/A |
| PA4417 | PA4417 | PA4417 | PA4417 | PA4417 | PA4417 | PA4417 | N/A | PA4417 | PA4417 | PA4417 | PA4417 | PA4417 | PA4417 | N/A | PA4417 | N/A | N/A | N/A |
| PA4418 | N/A | PA4418 | PA4418 | PA4418 | N/A | N/A | N/A | PA4418 | PA4418 | N/A | PA4418 | PA4418 | N/A | PA4418 | PA4418 | N/A | PA4418 | N/A |
| PA4419 | N/A | N/A | N/A | N/A | PA4419 | N/A | N/A | N/A | PA4419 | N/A | PA4419 | PA4419 | PA4419 | PA4419 | N/A | N/A | N/A | N/A |
| PA4420 | PA4420 | PA4420 | PA4420 | PA4420 | PA4420 | PA4420 | N/A | PA4420 | PA4420 | PA4420 | PA4420 | PA4420 | PA4420 | N/A | PA4420 | N/A | N/A | N/A |
| PA4421 | N/A | N/A | N/A | PA4421 | N/A | PA4421 | N/A | N/A | N/A | PA4421 | N/A | PA4421 | PA4421 | N/A | N/A | N/A | N/A | N/A |
| PA4422 | PA4422 | PA4422 | PA4422 | PA4422 | PA4422 | PA4422 | PA4422 | PA4422 | PA4422 | PA4422 | PA4422 | PA4422 | PA4422 | N/A | N/A | N/A | N/A | N/A |
| PA4423 | PA4423 | PA4423 | PA4423 | PA4423 | N/A | PA4423 | PA4423 | PA4423 | PA4423 | PA4423 | PA4423 | PA4423 | PA4423 | PA4423 | PA4423 | PA4423 | PA4423 | N/A |
| N/A | N/A | N/A | N/A | N/A | PA4424 | N/A | N/A | N/A | PA4424 | N/A | N/A | N/A | N/A | N/A | N/A | N/A | N/A | N/A |
| PA4425 | PA4425 | PA4425 | PA4425 | PA4425 | PA4425 | PA4425 | PA4425 | N/A | PA4425 | PA4425 | PA4425 | PA4425 | PA4425 | N/A | N/A | PA4425 | N/A | N/A |
| PA4426 | PA4426 | PA4426 | PA4426 | PA4426 | N/A | PA4426 | PA4426 | N/A | PA4426 | PA4426 | PA4426 | PA4426 | PA4426 | PA4426 | PA4426 | PA4426 | N/A | N/A |
| PA4427 | PA4427 | PA4427 | N/A | PA4427 | PA4427 | N/A | PA4427 | N/A | PA4427 | PA4427 | PA4427 | PA4427 | PA4427 | N/A | PA4427 | N/A | N/A | N/A |
| PA4428 | PA4428 | PA4428 | PA4428 | PA4428 | PA4428 | PA4428 | PA4428 | N/A | PA4428 | PA4428 | PA4428 | PA4428 | PA4428 | N/A | N/A | N/A | N/A | N/A |
| PA4429 | PA4429 | PA4429 | PA4429 | PA4429 | PA4429 | PA4429 | PA4429 | PA4429 | PA4429 | PA4429 | PA4429 | PA4429 | PA4429 | PA4429 | PA4429 | PA4429 | PA4429 | PA4429 |
| PA4430 | PA4430 | PA4430 | PA4430 | PA4430 | PA4430 | PA4430 | PA4430 | PA4430 | PA4430 | PA4430 | PA4430 | PA4430 | PA4430 | PA4430 | N/A | PA4430 | PA4430 | PA4430 |
| PA4431 | PA4431 | PA4431 | PA4431 | PA4431 | PA4431 | PA4431 | PA4431 | PA4431 | PA4431 | PA4431 | PA4431 | PA4431 | PA4431 | PA4431 | PA4431 | PA4431 | PA4431 | PA4431 |
| PA4432 | PA4432 | PA4432 | PA4432 | PA4432 | PA4432 | PA4432 | PA4432 | PA4432 | PA4432 | PA4432 | PA4432 | PA4432 | PA4432 | PA4432 | PA4432 | PA4432 | PA4432 | PA4432 |
| PA4433 | PA4433 | PA4433 | PA4433 | PA4433 | N/A | PA4433 | PA4433 | PA4433 | PA4433 | PA4433 | PA4433 | PA4433 | PA4433 | PA4433 | PA4433 | PA4433 | PA4433 | PA4433 |
| PA4434 | PA4434 | PA4434 | PA4434 | PA4434 | PA4434 | N/A | N/A | N/A | PA4434 | PA4434 | PA4434 | PA4434 | PA4434 | N/A | N/A | PA4434 | N/A | N/A |
| PA4435 | N/A | N/A | N/A | N/A | N/A | N/A | N/A | N/A | N/A | N/A | N/A | N/A | N/A | N/A | N/A | N/A | N/A | N/A |
| PA4436 | N/A | N/A | N/A | N/A | N/A | N/A | N/A | N/A | N/A | N/A | N/A | N/A | N/A | N/A | N/A | N/A | N/A | N/A |
| N/A | N/A | N/A | N/A | N/A | PA4437 | N/A | N/A | N/A | N/A | N/A | N/A | N/A | N/A | N/A | N/A | N/A | N/A | N/A |
| PA4438 | PA4438 | PA4438 | PA4438 | PA4438 | PA4438 | N/A | PA4438 | PA4438 | PA4438 | PA4438 | PA4438 | PA4438 | PA4438 | PA4438 | PA4438 | N/A | PA4438 | N/A |
| PA4439 | PA4439 | PA4439 | PA4439 | PA4439 | PA4439 | PA4439 | N/A | PA4439 | PA4439 | PA4439 | PA4439 | PA4439 | PA4439 | N/A | PA4439 | N/A | N/A | N/A |
| PA4440 | PA4440 | PA4440 | N/A | PA4440 | N/A | N/A | PA4440 | N/A | PA4440 | PA4440 | PA4440 | PA4440 | PA4440 | N/A | PA4440 | N/A | N/A | N/A |
| PA4441 | PA4441 | PA4441 | PA4441 | PA4441 | PA4441 | PA4441 | PA4441 | N/A | PA4441 | PA4441 | PA4441 | PA4441 | PA4441 | PA4441 | PA4441 | PA4441 | PA4441 | PA4441 |
| PA4442 | PA4442 | PA4442 | PA4442 | PA4442 | PA4442 | PA4442 | N/A | PA4442 | PA4442 | PA4442 | PA4442 | PA4442 | PA4442 | PA4442 | PA4442 | N/A | N/A | N/A |
| PA4443 | PA4443 | PA4443 | PA4443 | PA4443 | PA4443 | PA4443 | N/A | PA4443 | PA4443 | PA4443 | PA4443 | PA4443 | PA4443 | PA4443 | PA4443 | N/A | N/A | N/A |
| PA4444 | N/A | N/A | N/A | PA4444 | PA4444 | N/A | N/A | N/A | N/A | N/A | N/A | N/A | N/A | N/A | N/A | N/A | N/A | N/A |
| PA4445 | PA4445 | PA4445 | N/A | PA4445 | PA4445 | PA4445 | N/A | N/A | PA4445 | PA4445 | PA4445 | PA4445 | PA4445 | N/A | PA4445 | N/A | N/A | N/A |
| PA4446 | PA4446 | PA4446 | PA4446 | PA4446 | PA4446 | PA4446 | PA4446 | PA4446 | PA4446 | PA4446 | PA4446 | PA4446 | PA4446 | PA4446 | PA4446 | N/A | PA4446 | N/A |
| PA4447 | PA4447 | PA4447 | PA4447 | PA4447 | PA4447 | PA4447 | N/A | PA4447 | PA4447 | PA4447 | PA4447 | PA4447 | PA4447 | N/A | PA4447 | N/A | N/A | N/A |
| PA4448 | PA4448 | PA4448 | PA4448 | PA4448 | PA4448 | PA4448 | N/A | N/A | PA4448 | PA4448 | PA4448 | PA4448 | PA4448 | N/A | PA4448 | N/A | N/A | N/A |
| PA4449 | PA4449 | PA4449 | PA4449 | PA4449 | PA4449 | PA4449 | PA4449 | N/A | PA4449 | PA4449 | PA4449 | PA4449 | PA4449 | PA4449 | PA4449 | N/A | PA4449 | N/A |
| PA4450 | PA4450 | PA4450 | PA4450 | PA4450 | PA4450 | PA4450 | PA4450 | PA4450 | PA4450 | PA4450 | PA4450 | PA4450 | PA4450 | N/A | PA4450 | N/A | PA4450 | N/A |
| N/A | N/A | N/A | PA4451 | PA4451 | PA4451 | PA4451 | N/A | N/A | PA4451 | N/A | PA4451 | PA4451 | PA4451 | N/A | N/A | N/A | N/A | N/A |
| PA4452 | PA4452 | N/A | N/A | PA4452 | PA4452 | N/A | PA4452 | N/A | PA4452 | N/A | PA4452 | PA4452 | PA4452 | PA4452 | N/A | N/A | PA4452 | N/A |
| PA4453 | PA4453 | PA4453 | PA4453 | PA4453 | PA4453 | PA4453 | PA4453 | N/A | PA4453 | PA4453 | PA4453 | PA4453 | PA4453 | PA4453 | PA4453 | PA4453 | PA4453 | PA4453 |
| PA4454 | PA4454 | PA4454 | PA4454 | PA4454 | PA4454 | PA4454 | PA4454 | N/A | PA4454 | PA4454 | PA4454 | PA4454 | PA4454 | PA4454 | PA4454 | N/A | PA4454 | N/A |
| PA4455 | PA4455 | PA4455 | PA4455 | PA4455 | N/A | PA4455 | N/A | N/A | PA4455 | PA4455 | PA4455 | PA4455 | PA4455 | PA4455 | PA4455 | N/A | PA4455 | N/A |
| PA4456 | PA4456 | PA4456 | PA4456 | PA4456 | PA4456 | PA4456 | PA4456 | N/A | PA4456 | PA4456 | PA4456 | PA4456 | PA4456 | PA4456 | PA4456 | N/A | PA4456 | PA4456 |
| PA4457 | PA4457 | PA4457 | PA4457 | PA4457 | PA4457 | N/A | PA4457 | PA4457 | PA4457 | PA4457 | PA4457 | PA4457 | PA4457 | N/A | N/A | N/A | PA4457 | N/A |
| PA4458 | PA4458 | PA4458 | PA4458 | PA4458 | PA4458 | PA4458 | N/A | N/A | PA4458 | PA4458 | PA4458 | PA4458 | PA4458 | N/A | PA4458 | N/A | N/A | N/A |
| PA4459 | PA4459 | PA4459 | PA4459 | PA4459 | PA4459 | PA4459 | PA4459 | PA4459 | PA4459 | PA4459 | PA4459 | PA4459 | PA4459 | PA4459 | PA4459 | N/A | PA4459 | N/A |
| PA4460 | PA4460 | PA4460 | PA4460 | PA4460 | PA4460 | PA4460 | PA4460 | N/A | PA4460 | PA4460 | PA4460 | PA4460 | PA4460 | PA4460 | PA4460 | N/A | PA4460 | PA4460 |
| PA4461 | PA4461 | PA4461 | PA4461 | PA4461 | PA4461 | PA4461 | PA4461 | PA4461 | PA4461 | PA4461 | PA4461 | PA4461 | PA4461 | PA4461 | PA4461 | PA4461 | PA4461 | N/A |
| PA4462 | PA4462 | PA4462 | PA4462 | PA4462 | N/A | PA4462 | N/A | PA4462 | PA4462 | PA4462 | PA4462 | PA4462 | PA4462 | PA4462 | N/A | N/A | N/A | N/A |
| PA4463 | PA4463 | PA4463 | PA4463 | PA4463 | PA4463 | PA4463 | N/A | N/A | PA4463 | PA4463 | PA4463 | PA4463 | PA4463 | N/A | N/A | N/A | N/A | N/A |
| PA4464 | PA4464 | PA4464 | PA4464 | PA4464 | PA4464 | PA4464 | PA4464 | N/A | PA4464 | PA4464 | PA4464 | PA4464 | PA4464 | N/A | PA4464 | N/A | N/A | N/A |
| PA4465 | PA4465 | PA4465 | PA4465 | PA4465 | N/A | N/A | PA4465 | N/A | PA4465 | PA4465 | PA4465 | PA4465 | PA4465 | PA4465 | N/A | N/A | N/A | N/A |
| N/A | N/A | PA4466 | N/A | PA4466 | N/A | N/A | N/A | N/A | N/A | N/A | PA4466 | PA4466 | PA4466 | N/A | PA4466 | N/A | N/A | N/A |
| PA4468 | N/A | N/A | N/A | N/A | N/A | PA4468 | N/A | N/A | PA4468 | N/A | N/A | N/A | PA4468 | N/A | N/A | N/A | N/A | N/A |

| | | | | | | | | | | | | | | | | | | |
|---|---|---|---|---|---|---|---|---|---|---|---|---|---|---|---|---|---|---|
| N/A | N/A | N/A | N/A | N/A | N/A | N/A | N/A | N/A | N/A | N/A | N/A | N/A | PA4469 | N/A | N/A | N/A | N/A | N/A |
| PA4470 | PA4470 | PA4470 | N/A | PA4470 | N/A | PA4470 | N/A | N/A | N/A | N/A | N/A | N/A | PA4470 | N/A | N/A | N/A | N/A | N/A |
| N/A | N/A | N/A | N/A | N/A | PA4471 | N/A | N/A | N/A | N/A | N/A | N/A | N/A | N/A | N/A | N/A | N/A | N/A | N/A |
| PA4472 | PA4472 | PA4472 | PA4472 | PA4472 | PA4472 | PA4472 | N/A | N/A | PA4472 | PA4472 | PA4472 | PA4472 | PA4472 | N/A | PA4472 | N/A | N/A | N/A |
| PA4473 | PA4473 | PA4473 | PA4473 | PA4473 | PA4473 | PA4473 | PA4473 | PA4473 | PA4473 | PA4473 | PA4473 | PA4473 | PA4473 | PA4473 | PA4473 | N/A | PA4473 | N/A |
| PA4474 | PA4474 | PA4474 | PA4474 | PA4474 | PA4474 | PA4474 | N/A | N/A | PA4474 | PA4474 | PA4474 | PA4474 | PA4474 | N/A | PA4474 | N/A | N/A | N/A |
| PA4475 | N/A | N/A | N/A | PA4475 | PA4475 | PA4475 | N/A | N/A | N/A | PA4475 | PA4475 | PA4475 | PA4475 | N/A | N/A | N/A | N/A | N/A |
| PA4476 | N/A | N/A | N/A | PA4476 | PA4476 | N/A | N/A | PA4476 | PA4476 | PA4476 | PA4476 | PA4476 | PA4476 | PA4476 | N/A | N/A | PA4476 | N/A |
| PA4477 | N/A | PA4477 | PA4477 | PA4477 | PA4477 | N/A | N/A | PA4477 | PA4477 | PA4477 | PA4477 | PA4477 | PA4477 | N/A | N/A | N/A | N/A | N/A |
| PA4478 | N/A | N/A | N/A | PA4478 | N/A | N/A | N/A | N/A | PA4478 | N/A | PA4478 | PA4478 | PA4478 | N/A | N/A | N/A | N/A | N/A |
| PA4480 | PA4480 | PA4480 | PA4480 | PA4480 | PA4480 | N/A | PA4480 | PA4480 | PA4480 | PA4480 | PA4480 | PA4480 | PA4480 | PA4480 | PA4480 | N/A | PA4480 | N/A |
| PA4481 | PA4481 | PA4481 | PA4481 | PA4481 | PA4481 | PA4481 | PA4481 | PA4481 | PA4481 | PA4481 | PA4481 | PA4481 | PA4481 | PA4481 | PA4481 | N/A | PA4481 | PA4481 |
| PA4482 | PA4482 | PA4482 | PA4482 | PA4482 | PA4482 | PA4482 | N/A | N/A | PA4482 | PA4482 | PA4482 | PA4482 | PA4482 | N/A | N/A | N/A | N/A | N/A |
| PA4483 | PA4483 | PA4483 | PA4483 | PA4483 | PA4483 | PA4483 | PA4483 | PA4483 | PA4483 | PA4483 | PA4483 | PA4483 | PA4483 | PA4483 | PA4483 | PA4483 | PA4483 | N/A |
| PA4484 | PA4484 | PA4484 | PA4484 | PA4484 | PA4484 | PA4484 | N/A | PA4484 | PA4484 | PA4484 | PA4484 | PA4484 | PA4484 | PA4484 | PA4484 | PA4484 | N/A | N/A |
| PA4485 | N/A | N/A | N/A | N/A | N/A | N/A | N/A | N/A | N/A | N/A | N/A | N/A | N/A | N/A | N/A | N/A | N/A | N/A |
| PA4486 | N/A | N/A | N/A | PA4486 | N/A | PA4486 | N/A | N/A | PA4486 | PA4486 | PA4486 | PA4486 | PA4486 | N/A | N/A | N/A | N/A | N/A |
| PA4487 | N/A | PA4487 | N/A | PA4487 | N/A | N/A | PA4487 | N/A | PA4487 | PA4487 | PA4487 | N/A | PA4487 | PA4487 | PA4487 | N/A | PA4487 | PA4487 |
| PA4488 | N/A | N/A | N/A | PA4488 | N/A | N/A | N/A | PA4488 | N/A | N/A | N/A | N/A | N/A | N/A | N/A | N/A | PA4488 | PA4488 |
| PA4489 | PA4489 | PA4489 | PA4489 | PA4489 | PA4489 | PA4489 | PA4489 | PA4489 | PA4489 | PA4489 | PA4489 | PA4489 | PA4489 | PA4489 | PA4489 | N/A | PA4489 | PA4489 |
| PA4490 | N/A | N/A | N/A | N/A | N/A | N/A | N/A | N/A | N/A | N/A | N/A | N/A | N/A | N/A | N/A | N/A | PA4490 | PA4490 |
| PA4491 | PA4491 | PA4491 | PA4491 | PA4491 | N/A | N/A | PA4491 | PA4491 | PA4491 | PA4491 | PA4491 | PA4491 | PA4491 | PA4491 | PA4491 | N/A | PA4491 | PA4491 |
| PA4492 | N/A | N/A | N/A | N/A | PA4492 | N/A | PA4492 | N/A | N/A | N/A | PA4492 | N/A | N/A | PA4492 | N/A | N/A | PA4492 | N/A |
| PA4493 | PA4493 | PA4493 | PA4493 | PA4493 | PA4493 | PA4493 | PA4493 | N/A | PA4493 | PA4493 | PA4493 | PA4493 | PA4493 | PA4493 | N/A | N/A | PA4493 | PA4493 |
| PA4494 | N/A | N/A | N/A | PA4494 | PA4494 | N/A | N/A | N/A | N/A | PA4494 | PA4494 | PA4494 | PA4494 | PA4494 | N/A | N/A | PA4494 | N/A |
| PA4495 | PA4495 | PA4495 | PA4495 | PA4495 | PA4495 | PA4495 | PA4495 | N/A | PA4495 | PA4495 | PA4495 | PA4495 | PA4495 | PA4495 | PA4495 | PA4495 | PA4495 | PA4495 |
| PA4496 | N/A | N/A | PA4496 | PA4496 | N/A | PA4496 | N/A | N/A | PA4496 | PA4496 | PA4496 | PA4496 | PA4496 | N/A | PA4496 | PA4496 | N/A | N/A |
| PA4497 | N/A | N/A | N/A | N/A | PA4497 | PA4497 | N/A | N/A | N/A | PA4497 | PA4497 | PA4497 | PA4497 | N/A | N/A | N/A | N/A | N/A |
| PA4498 | PA4498 | N/A | N/A | PA4498 | PA4498 | N/A | PA4498 | N/A | PA4498 | PA4498 | PA4498 | PA4498 | PA4498 | N/A | PA4498 | N/A | N/A | N/A |
| PA4499 | N/A | N/A | N/A | PA4499 | PA4499 | N/A | N/A | N/A | PA4499 | N/A | N/A | N/A | N/A | N/A | N/A | N/A | N/A | N/A |
| PA4500 | PA4500 | PA4500 | PA4500 | PA4500 | PA4500 | PA4500 | PA4500 | N/A | PA4500 | PA4500 | PA4500 | PA4500 | PA4500 | N/A | PA4500 | PA4500 | N/A | N/A |
| PA4501 | N/A | N/A | N/A | PA4501 | PA4501 | PA4501 | N/A | PA4501 | N/A | N/A | PA4501 | PA4501 | PA4501 | N/A | PA4501 | PA4501 | N/A | N/A |
| PA4502 | N/A | N/A | N/A | PA4502 | PA4502 | N/A | N/A | N/A | PA4502 | PA4502 | PA4502 | PA4502 | PA4502 | N/A | PA4502 | N/A | N/A | N/A |
| PA4503 | PA4503 | N/A | N/A | PA4503 | N/A | N/A | N/A | N/A | N/A | PA4503 | PA4503 | PA4503 | PA4503 | PA4503 | PA4503 | N/A | N/A | N/A |
| PA4504 | N/A | N/A | N/A | PA4504 | N/A | PA4504 | N/A | N/A | PA4504 | PA4504 | PA4504 | PA4504 | PA4504 | PA4504 | N/A | N/A | PA4504 | N/A |
| PA4505 | N/A | N/A | N/A | PA4505 | PA4505 | N/A | PA4505 | N/A | PA4505 | N/A | PA4505 | PA4505 | PA4505 | PA4505 | PA4505 | N/A | N/A | N/A |
| PA4506 | N/A | PA4506 | PA4506 | PA4506 | N/A | N/A | N/A | N/A | PA4506 | PA4506 | PA4506 | PA4506 | PA4506 | PA4506 | PA4506 | N/A | PA4506 | PA4506 |
| PA4508 | N/A | N/A | N/A | PA4508 | N/A | N/A | N/A | N/A | N/A | N/A | N/A | N/A | N/A | N/A | N/A | N/A | N/A | N/A |
| PA4511 | N/A | N/A | N/A | PA4511 | N/A | N/A | N/A | N/A | N/A | PA4511 | PA4511 | PA4511 | PA4511 | N/A | N/A | N/A | N/A | N/A |
| PA4512 | PA4512 | N/A | N/A | PA4512 | N/A | N/A | N/A | PA4512 | PA4512 | PA4512 | PA4512 | PA4512 | PA4512 | PA4512 | N/A | N/A | PA4512 | N/A |
| PA4513 | N/A | N/A | PA4513 | PA4513 | N/A | N/A | N/A | PA4513 | N/A | N/A | N/A | N/A | N/A | PA4513 | N/A | N/A | PA4513 | N/A |
| PA4514 | PA4514 | PA4514 | N/A | PA4514 | N/A | PA4514 | N/A | PA4514 | PA4514 | N/A | N/A | PA4514 | N/A | N/A | PA4514 | N/A | N/A | N/A |
| PA4515 | PA4515 | PA4515 | N/A | PA4515 | N/A | PA4515 | N/A | N/A | PA4515 | N/A | PA4515 | PA4515 | PA4515 | N/A | N/A | N/A | N/A | N/A |
| PA4516 | N/A | N/A | N/A | PA4516 | N/A | N/A | PA4516 | N/A | N/A | N/A | N/A | N/A | N/A | N/A | N/A | N/A | N/A | N/A |
| N/A | N/A | N/A | N/A | PA4517 | N/A | N/A | N/A | N/A | N/A | N/A | N/A | N/A | N/A | N/A | N/A | N/A | N/A | N/A |
| PA4518 | PA4518 | N/A | N/A | PA4518 | PA4518 | N/A | N/A | N/A | PA4518 | N/A | PA4518 | PA4518 | PA4518 | N/A | N/A | N/A | N/A | N/A |
| PA4519 | PA4519 | PA4519 | PA4519 | PA4519 | PA4519 | PA4519 | N/A | PA4519 | PA4519 | PA4519 | PA4519 | PA4519 | PA4519 | N/A | PA4519 | N/A | N/A | N/A |
| PA4520 | PA4520 | PA4520 | PA4520 | PA4520 | N/A | PA4520 | PA4520 | PA4520 | PA4520 | PA4520 | PA4520 | PA4520 | PA4520 | PA4520 | PA4520 | N/A | PA4520 | PA4520 |
| PA4521 | N/A | N/A | N/A | PA4521 | N/A | PA4521 | N/A | N/A | PA4521 | PA4521 | PA4521 | PA4521 | PA4521 | PA4521 | N/A | N/A | PA4521 | N/A |
| PA4522 | N/A | N/A | N/A | N/A | N/A | N/A | N/A | N/A | N/A | PA4522 | N/A | PA4522 | N/A | N/A | N/A | N/A | N/A | N/A |
| PA4523 | N/A | N/A | N/A | N/A | PA4523 | N/A | N/A | PA4523 | PA4523 | PA4523 | PA4523 | PA4523 | PA4523 | N/A | N/A | N/A | N/A | N/A |
| PA4524 | PA4524 | PA4524 | PA4524 | PA4524 | PA4524 | PA4524 | PA4524 | N/A | PA4524 | PA4524 | PA4524 | PA4524 | PA4524 | N/A | PA4524 | PA4524 | N/A | N/A |
| PA4525 | PA4525 | PA4525 | PA4525 | PA4525 | PA4525 | PA4525 | PA4525 | PA4525 | PA4525 | PA4525 | PA4525 | PA4525 | PA4525 | PA4525 | PA4525 | PA4525 | PA4525 | PA4525 |
| PA4526 | PA4526 | N/A | N/A | PA4526 | N/A | PA4526 | N/A | PA4526 | PA4526 | PA4526 | PA4526 | PA4526 | PA4526 | PA4526 | N/A | N/A | N/A | N/A |
| PA4528 | N/A | N/A | N/A | N/A | PA4528 | N/A | N/A | N/A | PA4528 | PA4528 | PA4528 | PA4528 | N/A | PA4528 | N/A | N/A | PA4528 | N/A |
| PA4529 | PA4529 | PA4529 | PA4529 | PA4529 | PA4529 | N/A | PA4529 | N/A | PA4529 | PA4529 | PA4529 | PA4529 | PA4529 | N/A | N/A | N/A | N/A | N/A |
| PA4532 | N/A | N/A | N/A | N/A | PA4532 | N/A | N/A | N/A | N/A | N/A | N/A | N/A | N/A | PA4532 | N/A | N/A | N/A | N/A |
| PA4533 | N/A | N/A | N/A | PA4533 | PA4533 | PA4533 | N/A | N/A | N/A | PA4533 | PA4533 | PA4533 | PA4533 | N/A | PA4533 | N/A | N/A | N/A |
| PA4535 | N/A | N/A | N/A | PA4535 | N/A | N/A | N/A | N/A | N/A | N/A | N/A | N/A | N/A | N/A | N/A | N/A | N/A | N/A |
| PA4536 | N/A | N/A | N/A | PA4536 | N/A | N/A | N/A | N/A | N/A | N/A | N/A | N/A | N/A | N/A | N/A | N/A | N/A | N/A |
| PA4537 | N/A | N/A | N/A | N/A | N/A | N/A | N/A | N/A | N/A | N/A | PA4537 | N/A | PA4537 | N/A | N/A | N/A | N/A | N/A |
| PA4538 | PA4538 | N/A | N/A | PA4538 | N/A | N/A | N/A | PA4538 | PA4538 | N/A | PA4538 | PA4538 | PA4538 | PA4538 | N/A | N/A | PA4538 | N/A |
| PA4539 | N/A | N/A | N/A | N/A | N/A | N/A | N/A | N/A | N/A | N/A | N/A | N/A | N/A | N/A | N/A | N/A | N/A | N/A |
| N/A | N/A | N/A | N/A | N/A | PA4541 | N/A | N/A | N/A | N/A | N/A | N/A | N/A | N/A | N/A | N/A | N/A | N/A | N/A |
| PA4542 | PA4542 | PA4542 | PA4542 | PA4542 | PA4542 | PA4542 | PA4542 | PA4542 | PA4542 | PA4542 | PA4542 | PA4542 | PA4542 | PA4542 | PA4542 | PA4542 | PA4542 | PA4542 |
| PA4543 | N/A | N/A | N/A | PA4543 | PA4543 | PA4543 | N/A | N/A | N/A | N/A | N/A | PA4543 | N/A | N/A | N/A | N/A | N/A | N/A |
| PA4544 | PA4544 | PA4544 | PA4544 | PA4544 | PA4544 | N/A | N/A | PA4544 | PA4544 | PA4544 | PA4544 | PA4544 | PA4544 | N/A | PA4544 | N/A | N/A | N/A |
| PA4545 | PA4545 | PA4545 | PA4545 | PA4545 | N/A | PA4545 | PA4545 | PA4545 | PA4545 | PA4545 | PA4545 | PA4545 | PA4545 | PA4545 | PA4545 | N/A | PA4545 | N/A |
| PA4546 | N/A | N/A | N/A | PA4546 | PA4546 | N/A | N/A | N/A | N/A | N/A | N/A | N/A | N/A | PA4546 | N/A | N/A | N/A | N/A |
| PA4547 | PA4547 | N/A | PA4547 | PA4547 | PA4547 | PA4547 | PA4547 | PA4547 | PA4547 | PA4547 | PA4547 | PA4547 | PA4547 | PA4547 | N/A | N/A | PA4547 | PA4547 |
| PA4548 | PA4548 | PA4548 | PA4548 | PA4548 | PA4548 | N/A | PA4548 | PA4548 | PA4548 | PA4548 | PA4548 | PA4548 | PA4548 | N/A | N/A | N/A | N/A | N/A |
| PA4550 | N/A | N/A | N/A | N/A | N/A | PA4550 | PA4550 | N/A | N/A | PA4550 | PA4550 | PA4550 | PA4550 | PA4550 | N/A | N/A | N/A | N/A |
| PA4551 | N/A | N/A | N/A | N/A | N/A | N/A | PA4551 | N/A | PA4551 | PA4551 | PA4551 | PA4551 | PA4551 | PA4551 | N/A | N/A | PA4551 | N/A |
| PA4552 | N/A | N/A | N/A | PA4552 | N/A | N/A | PA4552 | N/A | PA4552 | PA4552 | PA4552 | N/A | N/A | PA4552 | N/A | N/A | PA4552 | N/A |
| PA4553 | N/A | N/A | N/A | PA4553 | PA4553 | N/A | PA4553 | N/A | PA4553 | PA4553 | PA4553 | PA4553 | PA4553 | PA4553 | N/A | N/A | PA4553 | N/A |
| PA4554 | PA4554 | PA4554 | PA4554 | PA4554 | PA4554 | PA4554 | N/A | PA4554 | PA4554 | PA4554 | PA4554 | PA4554 | PA4554 | PA4554 | PA4554 | N/A | PA4554 | N/A |
| PA4555 | N/A | N/A | N/A | PA4555 | N/A | N/A | N/A | N/A | PA4555 | PA4555 | N/A | PA4555 | N/A | N/A | N/A | N/A | N/A | N/A |
| PA4556 | N/A | N/A | PA4556 | PA4556 | PA4556 | PA4556 | N/A | N/A | PA4556 | PA4556 | PA4556 | PA4556 | PA4556 | PA4556 | N/A | N/A | PA4556 | N/A |
| PA4557 | PA4557 | PA4557 | PA4557 | PA4557 | PA4557 | N/A | N/A | N/A | PA4557 | PA4557 | PA4557 | PA4557 | PA4557 | N/A | N/A | N/A | N/A | N/A |
| PA4558 | PA4558 | PA4558 | PA4558 | PA4558 | PA4558 | PA4558 | PA4558 | N/A | PA4558 | PA4558 | PA4558 | PA4558 | PA4558 | N/A | PA4558 | N/A | N/A | N/A |
| N/A | N/A | N/A | N/A | N/A | PA4559 | N/A | N/A | N/A | N/A | N/A | N/A | N/A | N/A | PA4559 | N/A | N/A | N/A | N/A |
| PA4560 | PA4560 | PA4560 | PA4560 | PA4560 | PA4560 | PA4560 | N/A | PA4560 | PA4560 | PA4560 | PA4560 | PA4560 | PA4560 | PA4560 | PA4560 | PA4560 | N/A | N/A |
| PA4561 | PA4561 | PA4561 | PA4561 | PA4561 | PA4561 | N/A | N/A | N/A | PA4561 | PA4561 | PA4561 | PA4561 | PA4561 | N/A | N/A | N/A | N/A | N/A |
| PA4562 | N/A | N/A | N/A | PA4562 | PA4562 | N/A | N/A | PA4562 | N/A | N/A | PA4562 | N/A | N/A | PA4562 | N/A | N/A | N/A | N/A |
| PA4563 | PA4563 | PA4563 | PA4563 | PA4563 | PA4563 | PA4563 | PA4563 | PA4563 | PA4563 | PA4563 | PA4563 | PA4563 | PA4563 | PA4563 | PA4563 | PA4563 | PA4563 | PA4563 |
| PA4564 | PA4564 | PA4564 | PA4564 | PA4564 | PA4564 | PA4564 | N/A | N/A | PA4564 | PA4564 | PA4564 | PA4564 | PA4564 | PA4564 | PA4564 | N/A | PA4564 | N/A |
| PA4565 | PA4565 | PA4565 | PA4565 | PA4565 | PA4565 | PA4565 | N/A | PA4565 | PA4565 | PA4565 | PA4565 | PA4565 | PA4565 | PA4565 | N/A | N/A | PA4565 | PA4565 |
| PA4566 | PA4566 | PA4566 | PA4566 | PA4566 | PA4566 | PA4566 | PA4566 | PA4566 | PA4566 | PA4566 | PA4566 | PA4566 | PA4566 | PA4566 | PA4566 | N/A | N/A | N/A |
| PA4567 | PA4567 | PA4567 | PA4567 | PA4567 | PA4567 | PA4567 | PA4567 | PA4567 | PA4567 | PA4567 | PA4567 | PA4567 | PA4567 | PA4567 | PA4567 | PA4567 | PA4567 | N/A |
| PA4568 | PA4568 | PA4568 | PA4568 | PA4568 | PA4568 | PA4568 | PA4568 | PA4568 | PA4568 | PA4568 | PA4568 | PA4568 | PA4568 | PA4568 | PA4568 | PA4568 | PA4568 | N/A |
| PA4569 | PA4569 | PA4569 | PA4569 | PA4569 | PA4569 | PA4569 | PA4569 | PA4569 | PA4569 | PA4569 | PA4569 | PA4569 | PA4569 | N/A | PA4569 | N/A | N/A | N/A |
| N/A | N/A | N/A | N/A | N/A | N/A | N/A | N/A | N/A | N/A | N/A | N/A | N/A | PA4570 | N/A | N/A | N/A | N/A | N/A |
| PA4571 | N/A | N/A | N/A | PA4571 | N/A | N/A | PA4571 | N/A | N/A | PA4571 | N/A | N/A | N/A | PA4571 | N/A | N/A | PA4571 | N/A |
| PA4572 | PA4572 | PA4572 | PA4572 | PA4572 | PA4572 | N/A | PA4572 | N/A | PA4572 | PA4572 | PA4572 | PA4572 | PA4572 | N/A | PA4572 | N/A | N/A | N/A |
| PA4573 | N/A | N/A | N/A | N/A | N/A | N/A | N/A | N/A | N/A | N/A | N/A | N/A | N/A | N/A | N/A | N/A | N/A | N/A |
| PA4574 | PA4574 | PA4574 | PA4574 | N/A | N/A | PA4574 | N/A | N/A | N/A | N/A | N/A | N/A | N/A | PA4574 | N/A | N/A | N/A | N/A |
| N/A | N/A | N/A | N/A | N/A | PA4575 | N/A | N/A | N/A | N/A | N/A | N/A | N/A | PA4575 | N/A | N/A | N/A | N/A | N/A |
| PA4576 | PA4576 | PA4576 | PA4576 | PA4576 | PA4576 | PA4576 | N/A | PA4576 | PA4576 | PA4576 | PA4576 | PA4576 | PA4576 | N/A | PA4576 | PA4576 | N/A | N/A |
| PA4577 | N/A | N/A | N/A | PA4577 | PA4577 | PA4577 | N/A | N/A | PA4577 | PA4577 | PA4577 | PA4577 | PA4577 | N/A | N/A | N/A | N/A | N/A |
| N/A | N/A | N/A | N/A | N/A | N/A | PA4578 | PA4578 | N/A | PA4578 | PA4578 | PA4578 | PA4578 | PA4578 | PA4578 | N/A | N/A | PA4578 | N/A |
| PA4579 | N/A | N/A | N/A | PA4579 | N/A | PA4579 | N/A | N/A | N/A | N/A | N/A | N/A | N/A | N/A | N/A | N/A | N/A | N/A |
| PA4580 | N/A | N/A | N/A | N/A | N/A | N/A | N/A | N/A | N/A | N/A | N/A | N/A | N/A | N/A | N/A | N/A | N/A | N/A |
| PA4581 | N/A | N/A | N/A | PA4581 | PA4581 | N/A | N/A | PA4581 | N/A | N/A | N/A | N/A | N/A | N/A | N/A | N/A | N/A | N/A |
| N/A | N/A | N/A | N/A | PA4582 | PA4582 | N/A | N/A | N/A | N/A | N/A | N/A | N/A | N/A | N/A | N/A | N/A | PA4582 | N/A |
| PA4583 | PA4583 | PA4583 | PA4583 | PA4583 | PA4583 | N/A | PA4583 | PA4583 | N/A | PA4583 | N/A | N/A | N/A | N/A | N/A | N/A | PA4583 | N/A |
| N/A | N/A | N/A | N/A | N/A | PA4584 | N/A | N/A | N/A | N/A | N/A | N/A | N/A | N/A | N/A | N/A | N/A | N/A | N/A |
| PA4585 | N/A | N/A | N/A | PA4585 | PA4585 | N/A | N/A | N/A | N/A | N/A | N/A | N/A | N/A | N/A | N/A | N/A | PA4585 | N/A |
| N/A | N/A | N/A | N/A | N/A | PA4586 | N/A | N/A | N/A | N/A | N/A | N/A | N/A | N/A | N/A | N/A | N/A | N/A | N/A |
| PA4587 | N/A | N/A | N/A | PA4587 | PA4587 | PA4587 | N/A | N/A | PA4587 | PA4587 | PA4587 | PA4587 | PA4587 | PA4587 | N/A | N/A | N/A | N/A |

| | | | | | | | | | | | | | | | | | | |
|---|---|---|---|---|---|---|---|---|---|---|---|---|---|---|---|---|---|---|
| N/A | N/A | N/A | N/A | N/A | N/A | PA4588 | N/A | N/A | N/A | PA4588 | N/A | N/A | PA4588 | N/A | N/A | N/A | N/A | N/A |
| N/A | N/A | N/A | N/A | N/A | PA4589 | N/A | N/A | N/A | N/A | N/A | N/A | N/A | N/A | N/A | N/A | N/A | N/A | N/A |
| PA4590 | N/A | N/A | N/A | N/A | PA4590 | PA4590 | N/A | N/A | PA4590 | PA4590 | N/A | PA4590 | PA4590 | N/A | N/A | N/A | N/A | N/A |
| PA4591 | N/A | N/A | N/A | PA4591 | N/A | N/A | N/A | N/A | N/A | N/A | N/A | N/A | N/A | PA4591 | N/A | N/A | N/A | N/A |
| N/A | N/A | N/A | N/A | N/A | PA4594 | N/A | N/A | N/A | N/A | N/A | N/A | N/A | N/A | N/A | N/A | N/A | N/A | N/A |
| PA4595 | PA4595 | PA4595 | PA4595 | PA4595 | PA4595 | PA4595 | PA4595 | PA4595 | PA4595 | PA4595 | PA4595 | PA4595 | PA4595 | PA4595 | PA4595 | N/A | PA4595 | N/A |
| PA4598 | N/A | PA4598 | PA4598 | PA4598 | N/A | PA4598 | N/A | N/A | PA4598 | N/A | PA4598 | PA4598 | PA4598 | PA4598 | N/A | N/A | PA4598 | N/A |
| N/A | N/A | N/A | N/A | PA4599 | N/A | N/A | N/A | N/A | N/A | PA4599 | N/A | N/A | N/A | N/A | N/A | N/A | N/A | N/A |
| PA4600 | PA4600 | PA4600 | PA4600 | PA4600 | PA4600 | N/A | N/A | N/A | PA4600 | PA4600 | PA4600 | PA4600 | PA4600 | N/A | N/A | N/A | N/A | N/A |
| PA4601 | N/A | N/A | N/A | PA4601 | PA4601 | N/A | N/A | N/A | N/A | N/A | N/A | N/A | N/A | PA4601 | N/A | N/A | N/A | N/A |
| PA4602 | PA4602 | PA4602 | PA4602 | PA4602 | N/A | PA4602 | PA4602 | PA4602 | PA4602 | PA4602 | PA4602 | PA4602 | PA4602 | PA4602 | PA4602 | N/A | N/A | PA4602 |
| N/A | N/A | N/A | N/A | N/A | PA4603 | N/A | N/A | N/A | N/A | N/A | N/A | N/A | N/A | N/A | N/A | N/A | N/A | N/A |
| PA4604 | PA4604 | PA4604 | PA4604 | PA4604 | PA4604 | N/A | N/A | PA4604 | PA4604 | PA4604 | PA4604 | PA4604 | PA4604 | N/A | N/A | N/A | N/A | N/A |
| PA4606 | PA4606 | PA4606 | PA4606 | PA4606 | N/A | PA4606 | N/A | PA4606 | PA4606 | PA4606 | PA4606 | PA4606 | PA4606 | PA4606 | N/A | N/A | PA4606 | N/A |
| PA4607 | N/A | N/A | N/A | N/A | N/A | PA4607 | N/A | N/A | PA4607 | PA4607 | PA4607 | PA4607 | PA4607 | PA4607 | N/A | N/A | N/A | N/A |
| PA4608 | N/A | N/A | N/A | N/A | N/A | PA4608 | N/A | N/A | PA4608 | PA4608 | PA4608 | PA4608 | PA4608 | N/A | N/A | N/A | N/A | N/A |
| PA4609 | PA4609 | N/A | PA4609 | PA4609 | PA4609 | N/A | N/A | PA4609 | PA4609 | PA4609 | N/A | PA4609 | PA4609 | N/A | N/A | N/A | N/A | N/A |
| PA4610 | N/A | N/A | N/A | N/A | PA4610 | PA4610 | N/A | N/A | N/A | PA4610 | N/A | N/A | N/A | PA4610 | N/A | N/A | PA4610 | N/A |
| PA4611 | N/A | N/A | N/A | PA4611 | PA4611 | PA4611 | N/A | N/A | PA4611 | PA4611 | PA4611 | PA4611 | PA4611 | N/A | N/A | N/A | N/A | N/A |
| PA4613 | N/A | N/A | N/A | PA4613 | PA4613 | PA4613 | N/A | N/A | N/A | N/A | PA4613 | N/A | PA4613 | N/A | PA4613 | N/A | N/A | N/A |
| PA4614 | PA4614 | PA4614 | PA4614 | PA4614 | PA4614 | PA4614 | PA4614 | N/A | PA4614 | PA4614 | PA4614 | PA4614 | PA4614 | PA4614 | N/A | N/A | PA4614 | N/A |
| PA4615 | PA4615 | PA4615 | PA4615 | PA4615 | PA4615 | PA4615 | N/A | N/A | PA4615 | N/A | PA4615 | PA4615 | PA4615 | PA4615 | PA4615 | N/A | N/A | N/A |
| PA4616 | PA4616 | PA4616 | PA4616 | PA4616 | N/A | N/A | PA4616 | PA4616 | PA4616 | PA4616 | PA4616 | PA4616 | PA4616 | PA4616 | PA4616 | N/A | N/A | N/A |
| PA4617 | PA4617 | N/A | N/A | PA4617 | N/A | N/A | N/A | PA4617 | N/A | PA4617 | N/A | N/A | N/A | PA4617 | N/A | N/A | N/A | N/A |
| PA4618 | N/A | N/A | N/A | PA4618 | N/A | N/A | N/A | N/A | N/A | N/A | N/A | N/A | PA4618 | N/A | N/A | N/A | N/A | N/A |
| PA4619 | PA4619 | PA4619 | PA4619 | PA4619 | N/A | N/A | PA4619 | N/A | PA4619 | PA4619 | PA4619 | PA4619 | N/A | N/A | N/A | N/A | N/A | N/A |
| PA4620 | N/A | N/A | PA4620 | PA4620 | PA4620 | N/A | N/A | N/A | PA4620 | PA4620 | PA4620 | PA4620 | N/A | N/A | PA4620 | N/A | N/A | N/A |
| PA4621 | PA4621 | PA4621 | PA4621 | PA4621 | PA4621 | N/A | N/A | N/A | PA4621 | PA4621 | N/A | PA4621 | PA4621 | N/A | PA4621 | N/A | N/A | N/A |
| PA4622 | N/A | N/A | N/A | N/A | N/A | N/A | N/A | N/A | N/A | N/A | N/A | N/A | N/A | PA4622 | N/A | N/A | N/A | N/A |
| PA4624 | N/A | N/A | N/A | PA4624 | PA4624 | PA4624 | N/A | N/A | N/A | N/A | N/A | PA4624 | PA4624 | N/A | N/A | N/A | N/A | N/A |
| PA4625 | N/A | N/A | N/A | N/A | N/A | PA4625 | N/A | N/A | N/A | N/A | N/A | N/A | N/A | N/A | PA4625 | PA4625 | N/A | N/A |
| PA4626 | PA4626 | N/A | N/A | PA4626 | N/A | PA4626 | N/A | N/A | PA4626 | N/A | PA4626 | PA4626 | PA4626 | N/A | N/A | N/A | N/A | N/A |
| PA4627 | PA4627 | PA4627 | PA4627 | PA4627 | N/A | N/A | PA4627 | N/A | PA4627 | PA4627 | PA4627 | PA4627 | PA4627 | PA4627 | N/A | N/A | N/A | N/A |
| PA4628 | N/A | N/A | PA4628 | PA4628 | N/A | PA4628 | N/A | PA4628 | PA4628 | N/A | PA4628 | PA4628 | PA4628 | PA4628 | N/A | N/A | PA4628 | N/A |
| PA4629 | N/A | N/A | N/A | N/A | N/A | N/A | N/A | N/A | N/A | N/A | N/A | N/A | N/A | N/A | N/A | N/A | N/A | N/A |
| PA4631 | PA4631 | N/A | N/A | PA4631 | PA4631 | N/A | PA4631 | PA4631 | PA4631 | N/A | PA4631 | PA4631 | PA4631 | N/A | N/A | N/A | N/A | N/A |
| PA4632 | PA4632 | PA4632 | PA4632 | PA4632 | PA4632 | PA4632 | PA4632 | N/A | PA4632 | PA4632 | PA4632 | PA4632 | PA4632 | PA4632 | PA4632 | N/A | PA4632 | PA4632 |
| PA4633 | PA4633 | PA4633 | PA4633 | PA4633 | N/A | PA4633 | PA4633 | PA4633 | PA4633 | PA4633 | PA4633 | PA4633 | PA4633 | PA4633 | PA4633 | N/A | PA4633 | PA4633 |
| N/A | N/A | PA4635 | N/A | N/A | PA4635 | N/A | N/A | N/A | N/A | N/A | N/A | N/A | N/A | N/A | N/A | N/A | N/A | N/A |
| PA4636 | PA4636 | PA4636 | PA4636 | PA4636 | PA4636 | N/A | PA4636 | PA4636 | PA4636 | PA4636 | PA4636 | PA4636 | PA4636 | PA4636 | PA4636 | N/A | PA4636 | N/A |
| N/A | N/A | N/A | N/A | N/A | PA4638 | N/A | N/A | N/A | N/A | N/A | N/A | N/A | N/A | N/A | N/A | N/A | N/A | N/A |
| PA4639 | PA4639 | PA4639 | PA4639 | PA4639 | PA4639 | PA4639 | PA4639 | N/A | PA4639 | PA4639 | PA4639 | PA4639 | PA4639 | PA4639 | PA4639 | PA4639 | PA4639 | PA4639 |
| PA4640 | PA4640 | PA4640 | PA4640 | PA4640 | PA4640 | PA4640 | PA4640 | PA4640 | PA4640 | PA4640 | PA4640 | PA4640 | PA4640 | PA4640 | PA4640 | PA4640 | N/A | PA4640 |
| PA4642 | PA4642 | PA4642 | PA4642 | PA4642 | PA4642 | PA4642 | N/A | N/A | PA4642 | PA4642 | PA4642 | PA4642 | PA4642 | N/A | PA4642 | N/A | N/A | N/A |
| PA4643 | N/A | PA4643 | N/A | PA4643 | PA4643 | PA4643 | PA4643 | N/A | PA4643 | PA4643 | PA4643 | PA4643 | PA4643 | PA4643 | N/A | N/A | N/A | N/A |
| PA4644 | PA4644 | PA4644 | PA4644 | PA4644 | PA4644 | N/A | PA4644 | N/A | PA4644 | PA4644 | PA4644 | PA4644 | PA4644 | N/A | PA4644 | N/A | N/A | N/A |
| PA4645 | PA4645 | PA4645 | PA4645 | PA4645 | PA4645 | PA4645 | PA4645 | N/A | PA4645 | PA4645 | PA4645 | PA4645 | PA4645 | N/A | PA4645 | N/A | N/A | N/A |
| PA4646 | PA4646 | PA4646 | PA4646 | PA4646 | PA4646 | PA4646 | PA4646 | N/A | PA4646 | PA4646 | PA4646 | PA4646 | PA4646 | N/A | PA4646 | N/A | PA4646 | N/A |
| PA4647 | N/A | N/A | N/A | PA4647 | PA4647 | PA4647 | N/A | N/A | N/A | N/A | N/A | PA4647 | N/A | PA4647 | N/A | N/A | PA4647 | N/A |
| PA4648 | N/A | N/A | N/A | N/A | PA4648 | PA4648 | N/A | N/A | N/A | N/A | N/A | N/A | PA4648 | N/A | N/A | PA4648 | N/A | N/A |
| PA4650 | N/A | N/A | N/A | N/A | N/A | N/A | N/A | N/A | N/A | N/A | N/A | N/A | N/A | N/A | N/A | N/A | N/A | N/A |
| PA4651 | N/A | N/A | N/A | N/A | N/A | N/A | N/A | N/A | N/A | N/A | PA4651 | PA4651 | PA4651 | N/A | N/A | N/A | N/A | N/A |
| PA4652 | N/A | N/A | N/A | N/A | N/A | N/A | N/A | N/A | N/A | N/A | N/A | N/A | N/A | N/A | N/A | N/A | N/A | N/A |
| PA4655 | N/A | N/A | N/A | PA4655 | PA4655 | PA4655 | PA4655 | PA4655 | PA4655 | PA4655 | PA4655 | PA4655 | PA4655 | PA4655 | N/A | N/A | PA4655 | N/A |
| PA4656 | N/A | N/A | N/A | PA4656 | N/A | PA4656 | PA4656 | N/A | N/A | PA4656 | N/A | PA4656 | PA4656 | PA4656 | N/A | N/A | N/A | N/A |
| PA4657 | PA4657 | N/A | N/A | PA4657 | PA4657 | PA4657 | N/A | N/A | PA4657 | PA4657 | PA4657 | PA4657 | PA4657 | N/A | PA4657 | N/A | N/A | N/A |
| PA4659 | N/A | N/A | N/A | PA4659 | N/A | N/A | N/A | N/A | PA4659 | N/A | PA4659 | PA4659 | PA4659 | N/A | N/A | N/A | N/A | N/A |
| PA4660 | N/A | N/A | N/A | PA4660 | PA4660 | N/A | N/A | N/A | N/A | N/A | N/A | N/A | N/A | N/A | N/A | N/A | N/A | N/A |
| PA4661 | PA4661 | PA4661 | PA4661 | PA4661 | PA4661 | PA4661 | PA4661 | PA4661 | PA4661 | PA4661 | PA4661 | PA4661 | PA4661 | PA4661 | PA4661 | PA4661 | PA4661 | PA4661 |
| PA4662 | PA4662 | PA4662 | N/A | PA4662 | PA4662 | N/A | N/A | N/A | PA4662 | PA4662 | PA4662 | N/A | PA4662 | N/A | N/A | N/A | N/A | N/A |
| PA4663 | N/A | PA4663 | N/A | PA4663 | PA4663 | N/A | N/A | N/A | PA4663 | PA4663 | PA4663 | PA4663 | PA4663 | N/A | N/A | N/A | N/A | N/A |
| PA4664 | N/A | N/A | N/A | PA4664 | PA4664 | N/A | N/A | N/A | N/A | N/A | N/A | N/A | N/A | N/A | N/A | N/A | N/A | N/A |
| PA4665 | PA4665 | PA4665 | PA4665 | PA4665 | PA4665 | N/A | PA4665 | N/A | PA4665 | PA4665 | PA4665 | N/A | PA4665 | N/A | PA4665 | N/A | N/A | N/A |
| PA4666 | PA4666 | PA4666 | PA4666 | PA4666 | N/A | PA4666 | PA4666 | PA4666 | PA4666 | PA4666 | PA4666 | PA4666 | PA4666 | PA4666 | N/A | N/A | PA4666 | N/A |
| PA4667 | PA4667 | PA4667 | PA4667 | PA4667 | N/A | PA4667 | N/A | PA4667 | PA4667 | PA4667 | PA4667 | PA4667 | PA4667 | PA4667 | PA4667 | N/A | N/A | N/A |
| PA4668 | PA4668 | N/A | PA4668 | PA4668 | N/A | PA4668 | PA4668 | N/A | PA4668 | PA4668 | PA4668 | PA4668 | PA4668 | PA4668 | PA4668 | N/A | N/A | N/A |
| PA4669 | N/A | N/A | N/A | PA4669 | PA4669 | N/A | PA4669 | N/A | N/A | PA4669 | PA4669 | PA4669 | N/A | PA4669 | N/A | N/A | N/A | N/A |
| PA4670 | PA4670 | PA4670 | PA4670 | PA4670 | PA4670 | PA4670 | PA4670 | PA4670 | PA4670 | PA4670 | PA4670 | PA4670 | PA4670 | PA4670 | PA4670 | N/A | PA4670 | PA4670 |
| PA4671 | PA4671 | PA4671 | N/A | PA4671 | PA4671 | PA4671 | PA4671 | PA4671 | PA4671 | PA4671 | PA4671 | PA4671 | PA4671 | PA4671 | PA4671 | PA4671 | N/A | PA4671 |
| PA4672 | PA4672 | PA4672 | PA4672 | PA4672 | PA4672 | N/A | PA4672 | N/A | PA4672 | PA4672 | PA4672 | PA4672 | PA4672 | N/A | N/A | N/A | N/A | N/A |
| PA4673 | PA4673 | PA4673 | PA4673 | PA4673 | PA4673 | PA4673 | PA4673 | PA4673 | PA4673 | PA4673 | PA4673 | PA4673 | PA4673 | N/A | PA4673 | N/A | N/A | N/A |
| PA4674 | N/A | N/A | N/A | PA4674 | PA4674 | PA4674 | N/A | N/A | N/A | N/A | PA4674 | N/A | PA4674 | N/A | N/A | N/A | N/A | N/A |
| PA4675 | PA4675 | PA4675 | PA4675 | PA4675 | PA4675 | PA4675 | PA4675 | PA4675 | PA4675 | PA4675 | PA4675 | PA4675 | PA4675 | N/A | PA4675 | PA4675 | N/A | N/A |
| PA4676 | PA4676 | PA4676 | PA4676 | PA4676 | PA4676 | PA4676 | PA4676 | N/A | PA4676 | PA4676 | PA4676 | PA4676 | PA4676 | N/A | PA4676 | PA4676 | N/A | N/A |
| PA4677 | N/A | N/A | N/A | N/A | N/A | N/A | N/A | N/A | N/A | N/A | N/A | N/A | PA4677 | N/A | N/A | N/A | N/A | N/A |
| PA4678 | N/A | N/A | N/A | N/A | N/A | N/A | N/A | N/A | N/A | N/A | N/A | N/A | N/A | N/A | N/A | N/A | N/A | N/A |
| PA4679 | PA4679 | PA4679 | PA4679 | PA4679 | N/A | N/A | PA4679 | N/A | PA4679 | N/A | PA4679 | PA4679 | PA4679 | N/A | N/A | N/A | PA4679 | PA4679 |
| N/A | N/A | N/A | N/A | PA4683 | PA4683 | N/A | N/A | N/A | N/A | N/A | N/A | N/A | N/A | N/A | N/A | N/A | N/A | N/A |
| PA4684 | N/A | N/A | N/A | PA4684 | PA4684 | N/A | PA4684 | N/A | PA4684 | N/A | PA4684 | N/A | N/A | N/A | N/A | N/A | N/A | N/A |
| N/A | PA4685 | N/A | N/A | PA4685 | PA4685 | N/A | PA4685 | PA4685 | PA4685 | N/A | PA4685 | N/A | N/A | PA4685 | N/A | N/A | PA4685 | N/A |
| PA4686 | PA4686 | PA4686 | PA4686 | PA4686 | PA4686 | N/A | N/A | PA4686 | PA4686 | PA4686 | PA4686 | PA4686 | PA4686 | N/A | PA4686 | N/A | N/A | N/A |
| PA4687 | PA4687 | PA4687 | PA4687 | PA4687 | PA4687 | N/A | PA4687 | PA4687 | PA4687 | PA4687 | PA4687 | PA4687 | PA4687 | PA4687 | PA4687 | PA4687 | N/A | N/A |
| PA4688 | N/A | N/A | N/A | PA4688 | N/A | N/A | N/A | N/A | N/A | N/A | N/A | N/A | N/A | N/A | N/A | N/A | N/A | N/A |
| PA4689 | PA4689 | PA4689 | N/A | PA4689 | N/A | PA4689 | N/A | PA4689 | PA4689 | PA4689 | PA4689 | PA4689 | PA4689 | PA4689 | PA4689 | N/A | PA4689 | N/A |
| N/A | N/A | N/A | N/A | N/A | N/A | N/A | N/A | N/A | N/A | PA4691 | N/A | N/A | N/A | N/A | N/A | N/A | N/A | N/A |
| PA4692 | N/A | N/A | N/A | N/A | PA4692 | PA4692 | N/A | N/A | N/A | PA4692 | N/A | N/A | N/A | N/A | N/A | N/A | N/A | N/A |
| PA4693 | PA4693 | PA4693 | N/A | PA4693 | PA4693 | PA4693 | N/A | PA4693 | PA4693 | PA4693 | PA4693 | PA4693 | N/A | PA4693 | PA4693 | N/A | PA4693 | PA4693 |
| PA4694 | PA4694 | PA4694 | PA4694 | PA4694 | PA4694 | PA4694 | PA4694 | PA4694 | PA4694 | PA4694 | PA4694 | PA4694 | PA4694 | PA4694 | PA4694 | PA4694 | PA4694 | N/A |
| PA4695 | PA4695 | PA4695 | PA4695 | PA4695 | PA4695 | PA4695 | PA4695 | PA4695 | PA4695 | PA4695 | PA4695 | PA4695 | PA4695 | PA4695 | PA4695 | N/A | PA4695 | PA4695 |
| PA4696 | PA4696 | PA4696 | PA4696 | PA4696 | PA4696 | PA4696 | PA4696 | PA4696 | PA4696 | PA4696 | PA4696 | PA4696 | PA4696 | PA4696 | PA4696 | N/A | N/A | PA4696 |
| PA4697 | N/A | N/A | N/A | N/A | N/A | N/A | N/A | N/A | N/A | N/A | N/A | N/A | N/A | N/A | N/A | N/A | N/A | N/A |
| PA4698 | N/A | N/A | PA4698 | PA4698 | N/A | N/A | PA4698 | N/A | PA4698 | N/A | PA4698 | PA4698 | PA4698 | N/A | N/A | N/A | N/A | N/A |
| PA4699 | PA4699 | PA4699 | PA4699 | PA4699 | N/A | PA4699 | PA4699 | N/A | PA4699 | PA4699 | PA4699 | PA4699 | PA4699 | PA4699 | PA4699 | PA4699 | PA4699 | PA4699 |
| PA4700 | PA4700 | PA4700 | PA4700 | PA4700 | N/A | N/A | N/A | PA4700 | PA4700 | N/A | PA4700 | PA4700 | PA4700 | PA4700 | PA4700 | N/A | PA4700 | N/A |
| PA4701 | PA4701 | PA4701 | PA4701 | PA4701 | N/A | PA4701 | N/A | PA4701 | PA4701 | PA4701 | PA4701 | PA4701 | PA4701 | N/A | PA4701 | PA4701 | N/A | N/A |
| PA4702 | N/A | N/A | N/A | N/A | N/A | PA4702 | N/A | N/A | N/A | N/A | PA4702 | PA4702 | PA4702 | N/A | N/A | N/A | N/A | N/A |
| PA4703 | N/A | N/A | N/A | N/A | N/A | N/A | N/A | N/A | N/A | N/A | PA4703 | N/A | PA4703 | N/A | N/A | N/A | N/A | N/A |
| PA4704 | PA4704 | N/A | N/A | PA4704 | N/A | N/A | N/A | N/A | PA4704 | PA4704 | PA4704 | PA4704 | PA4704 | N/A | N/A | N/A | N/A | N/A |
| PA4705 | N/A | N/A | N/A | PA4705 | N/A | N/A | N/A | N/A | N/A | N/A | N/A | N/A | N/A | N/A | N/A | N/A | N/A | N/A |
| PA4706 | N/A | N/A | N/A | PA4706 | N/A | N/A | PA4706 | N/A | N/A | N/A | N/A | N/A | N/A | PA4706 | N/A | N/A | N/A | N/A |
| PA4707 | N/A | N/A | N/A | N/A | PA4707 | N/A | N/A | N/A | N/A | N/A | N/A | N/A | N/A | PA4707 | N/A | N/A | N/A | N/A |
| PA4708 | N/A | N/A | N/A | PA4708 | PA4708 | PA4708 | N/A | N/A | PA4708 | PA4708 | PA4708 | PA4708 | PA4708 | N/A | PA4708 | PA4708 | N/A | N/A |
| PA4709 | N/A | N/A | N/A | PA4709 | N/A | N/A | N/A | N/A | N/A | N/A | N/A | N/A | PA4709 | N/A | N/A | N/A | N/A | N/A |
| PA4710 | N/A | N/A | N/A | PA4710 | N/A | PA4710 | N/A | PA4710 | N/A | N/A | N/A | N/A | PA4710 | N/A | N/A | N/A | N/A | N/A |
| N/A | N/A | N/A | N/A | PA4711 | N/A | N/A | N/A | N/A | PA4711 | N/A | N/A | N/A | N/A | N/A | PA4711 | N/A | N/A | N/A |
| PA4712 | N/A | N/A | N/A | N/A | N/A | N/A | N/A | N/A | N/A | N/A | N/A | N/A | N/A | N/A | N/A | N/A | N/A | N/A |
| PA4713 | N/A | N/A | N/A | N/A | N/A | N/A | N/A | N/A | N/A | N/A | N/A | N/A | N/A | PA4713 | N/A | N/A | N/A | N/A |
| N/A | N/A | N/A | N/A | N/A | N/A | N/A | N/A | N/A | PA4714 | N/A | PA4714 | N/A | N/A | N/A | N/A | N/A | N/A | N/A |
| PA4715 | PA4715 | PA4715 | N/A | PA4715 | PA4715 | PA4715 | N/A | PA4715 | PA4715 | PA4715 | PA4715 | PA4715 | PA4715 | PA4715 | N/A | PA4715 | PA4715 | PA4715 |

| | | | | | | | | | | | | | | | | | | |
|---|---|---|---|---|---|---|---|---|---|---|---|---|---|---|---|---|---|---|
| PA4716 | N/A | N/A | N/A | PA4716 | N/A | N/A | N/A | N/A | PA4716 | PA4716 | PA4716 | N/A | PA4716 | N/A | N/A | N/A | N/A | N/A |
| PA4717 | N/A | N/A | N/A | PA4717 | N/A | N/A | PA4717 | N/A | PA4717 | PA4717 | PA4717 | PA4717 | PA4717 | PA4717 | PA4717 | N/A | PA4717 | PA4717 |
| PA4718 | N/A | N/A | N/A | PA4718 | N/A | N/A | N/A | N/A | N/A | PA4718 | N/A | N/A | N/A | N/A | PA4718 | N/A | N/A | N/A |
| PA4719 | N/A | N/A | N/A | N/A | PA4719 | PA4719 | N/A | N/A | N/A | N/A | N/A | PA4719 | N/A | PA4719 | N/A | N/A | N/A | N/A |
| PA4720 | PA4720 | PA4720 | PA4720 | PA4720 | PA4720 | N/A | N/A | PA4720 | PA4720 | N/A | PA4720 | N/A | PA4720 | N/A | N/A | N/A | N/A | N/A |
| N/A | N/A | N/A | N/A | N/A | N/A | N/A | N/A | N/A | PA4721 | N/A | PA4721 | PA4721 | PA4721 | N/A | N/A | N/A | N/A | N/A |
| PA4722 | PA4722 | PA4722 | PA4722 | PA4722 | PA4722 | PA4722 | PA4722 | PA4722 | PA4722 | PA4722 | PA4722 | PA4722 | PA4722 | N/A | N/A | N/A | PA4722 | N/A |
| PA4723 | PA4723 | PA4723 | PA4723 | PA4723 | PA4723 | PA4723 | PA4723 | N/A | PA4723 | PA4723 | PA4723 | PA4723 | PA4723 | N/A | PA4723 | N/A | N/A | N/A |
| PA4724 | N/A | N/A | N/A | PA4724 | N/A | N/A | N/A | N/A | N/A | N/A | N/A | N/A | N/A | N/A | N/A | N/A | N/A | N/A |
| PA4725 | N/A | N/A | N/A | PA4725 | PA4725 | PA4725 | N/A | PA4725 | PA4725 | N/A | PA4725 | PA4725 | PA4725 | PA4725 | N/A | N/A | N/A | N/A |
| PA4726 | PA4726 | PA4726 | PA4726 | PA4726 | PA4726 | PA4726 | N/A | PA4726 | PA4726 | PA4726 | PA4726 | PA4726 | PA4726 | N/A | PA4726 | N/A | N/A | N/A |
| PA4727 | PA4727 | PA4727 | PA4727 | PA4727 | N/A | PA4727 | N/A | PA4727 | PA4727 | PA4727 | PA4727 | PA4727 | PA4727 | PA4727 | PA4727 | N/A | N/A | N/A |
| PA4728 | N/A | N/A | PA4728 | PA4728 | PA4728 | N/A | PA4728 | N/A | PA4728 | N/A | PA4728 | PA4728 | PA4728 | N/A | PA4728 | N/A | N/A | N/A |
| PA4729 | PA4729 | PA4729 | PA4729 | PA4729 | PA4729 | PA4729 | PA4729 | N/A | PA4729 | PA4729 | PA4729 | PA4729 | PA4729 | PA4729 | PA4729 | N/A | N/A | N/A |
| PA4730 | PA4730 | PA4730 | PA4730 | PA4730 | PA4730 | PA4730 | PA4730 | N/A | PA4730 | PA4730 | PA4730 | PA4730 | PA4730 | N/A | PA4730 | N/A | N/A | N/A |
| PA4731 | N/A | N/A | N/A | N/A | PA4731 | N/A | N/A | N/A | PA4731 | N/A | PA4731 | PA4731 | PA4731 | N/A | N/A | N/A | N/A | N/A |
| PA4732 | PA4732 | PA4732 | PA4732 | PA4732 | PA4732 | PA4732 | N/A | N/A | PA4732 | PA4732 | PA4732 | PA4732 | PA4732 | PA4732 | PA4732 | N/A | N/A | N/A |
| PA4733 | PA4733 | N/A | PA4733 | PA4733 | N/A | PA4733 | N/A | PA4733 | PA4733 | PA4733 | PA4733 | PA4733 | PA4733 | N/A | N/A | N/A | N/A | N/A |
| PA4734 | N/A | N/A | N/A | N/A | N/A | N/A | N/A | N/A | N/A | N/A | N/A | N/A | N/A | N/A | N/A | N/A | N/A | N/A |
| PA4735 | PA4735 | PA4735 | PA4735 | PA4735 | N/A | PA4735 | N/A | PA4735 | PA4735 | PA4735 | PA4735 | PA4735 | PA4735 | PA4735 | PA4735 | N/A | PA4735 | N/A |
| PA4736 | N/A | N/A | N/A | N/A | N/A | N/A | N/A | N/A | PA4736 | N/A | PA4736 | PA4736 | PA4736 | N/A | N/A | N/A | N/A | N/A |
| PA4737 | N/A | N/A | N/A | N/A | N/A | N/A | N/A | N/A | PA4737 | N/A | PA4737 | PA4737 | PA4737 | N/A | N/A | N/A | N/A | N/A |
| PA4738 | N/A | N/A | N/A | PA4738 | PA4738 | N/A | N/A | N/A | PA4738 | N/A | N/A | PA4738 | PA4738 | N/A | N/A | N/A | N/A | N/A |
| PA4739 | N/A | PA4739 | PA4739 | PA4739 | PA4739 | PA4739 | N/A | N/A | PA4739 | PA4739 | PA4739 | PA4739 | PA4739 | N/A | PA4739 | N/A | N/A | N/A |
| PA4740 | PA4740 | PA4740 | PA4740 | PA4740 | PA4740 | PA4740 | PA4740 | PA4740 | PA4740 | PA4740 | PA4740 | PA4740 | PA4740 | PA4740 | PA4740 | PA4740 | PA4740 | N/A |
| PA4741 | PA4741 | PA4741 | PA4741 | PA4741 | PA4741 | PA4741 | PA4741 | PA4741 | PA4741 | PA4741 | PA4741 | PA4741 | PA4741 | PA4741 | PA4741 | PA4741 | PA4741 | PA4741 |
| PA4742 | PA4742 | PA4742 | PA4742 | PA4742 | PA4742 | N/A | PA4742 | PA4742 | PA4742 | PA4742 | PA4742 | PA4742 | PA4742 | N/A | PA4742 | N/A | N/A | N/A |
| PA4743 | PA4743 | PA4743 | PA4743 | PA4743 | PA4743 | PA4743 | PA4743 | N/A | PA4743 | PA4743 | PA4743 | PA4743 | PA4743 | PA4743 | PA4743 | N/A | N/A | N/A |
| PA4744 | PA4744 | PA4744 | PA4744 | PA4744 | PA4744 | PA4744 | PA4744 | PA4744 | PA4744 | PA4744 | PA4744 | PA4744 | PA4744 | PA4744 | PA4744 | PA4744 | PA4744 | PA4744 |
| PA4745 | PA4745 | PA4745 | PA4745 | PA4745 | PA4745 | PA4745 | PA4745 | PA4745 | PA4745 | PA4745 | PA4745 | PA4745 | PA4745 | PA4745 | PA4745 | PA4745 | N/A | N/A |
| PA4746 | N/A | PA4746 | N/A | PA4746 | N/A | N/A | PA4746 | N/A | PA4746 | PA4746 | PA4746 | PA4746 | PA4746 | PA4746 | N/A | N/A | N/A | N/A |
| PA4747 | PA4747 | PA4747 | PA4747 | PA4747 | PA4747 | PA4747 | PA4747 | PA4747 | PA4747 | PA4747 | PA4747 | PA4747 | PA4747 | PA4747 | PA4747 | N/A | PA4747 | PA4747 |
| PA4748 | PA4748 | PA4748 | PA4748 | PA4748 | PA4748 | PA4748 | PA4748 | PA4748 | PA4748 | PA4748 | PA4748 | PA4748 | PA4748 | N/A | PA4748 | N/A | PA4748 | N/A |
| PA4749 | PA4749 | PA4749 | PA4749 | PA4749 | PA4749 | PA4749 | PA4749 | PA4749 | PA4749 | PA4749 | PA4749 | PA4749 | PA4749 | PA4749 | PA4749 | N/A | PA4749 | N/A |
| PA4750 | PA4750 | PA4750 | PA4750 | PA4750 | PA4750 | PA4750 | PA4750 | PA4750 | PA4750 | PA4750 | PA4750 | PA4750 | PA4750 | N/A | N/A | N/A | N/A | N/A |
| PA4751 | PA4751 | PA4751 | PA4751 | PA4751 | PA4751 | PA4751 | PA4751 | PA4751 | PA4751 | PA4751 | PA4751 | PA4751 | PA4751 | PA4751 | PA4751 | N/A | PA4751 | N/A |
| PA4752 | N/A | PA4752 | N/A | PA4752 | PA4752 | PA4752 | PA4752 | PA4752 | PA4752 | PA4752 | N/A | N/A | PA4752 | PA4752 | N/A | N/A | N/A | N/A |
| PA4753 | PA4753 | PA4753 | PA4753 | PA4753 | PA4753 | PA4753 | PA4753 | N/A | PA4753 | PA4753 | PA4753 | PA4753 | PA4753 | PA4753 | PA4753 | N/A | PA4753 | N/A |
| N/A | N/A | N/A | N/A | N/A | PA4754 | N/A | N/A | N/A | N/A | N/A | N/A | N/A | N/A | N/A | N/A | N/A | N/A | N/A |
| PA4755 | PA4755 | PA4755 | PA4755 | PA4755 | PA4755 | PA4755 | PA4755 | N/A | PA4755 | PA4755 | PA4755 | PA4755 | PA4755 | N/A | PA4755 | N/A | N/A | N/A |
| PA4756 | PA4756 | PA4756 | PA4756 | PA4756 | PA4756 | PA4756 | PA4756 | PA4756 | PA4756 | PA4756 | PA4756 | PA4756 | PA4756 | PA4756 | PA4756 | N/A | N/A | N/A |
| N/A | N/A | PA4757 | N/A | PA4757 | PA4757 | N/A | N/A | N/A | N/A | N/A | N/A | N/A | N/A | N/A | N/A | N/A | N/A | N/A |
| PA4758 | PA4758 | PA4758 | PA4758 | PA4758 | PA4758 | PA4758 | PA4758 | PA4758 | PA4758 | PA4758 | PA4758 | PA4758 | PA4758 | PA4758 | PA4758 | N/A | N/A | N/A |
| PA4759 | PA4759 | PA4759 | PA4759 | PA4759 | PA4759 | PA4759 | N/A | N/A | PA4759 | PA4759 | PA4759 | PA4759 | PA4759 | PA4759 | PA4759 | PA4759 | N/A | N/A |
| PA4760 | PA4760 | PA4760 | PA4760 | PA4760 | PA4760 | PA4760 | PA4760 | PA4760 | PA4760 | PA4760 | PA4760 | PA4760 | PA4760 | PA4760 | PA4760 | N/A | PA4760 | PA4760 |
| PA4761 | PA4761 | PA4761 | PA4761 | PA4761 | PA4761 | PA4761 | PA4761 | PA4761 | PA4761 | PA4761 | PA4761 | PA4761 | PA4761 | PA4761 | PA4761 | PA4761 | PA4761 | PA4761 |
| PA4762 | PA4762 | PA4762 | PA4762 | PA4762 | PA4762 | PA4762 | PA4762 | N/A | PA4762 | PA4762 | PA4762 | PA4762 | PA4762 | N/A | PA4762 | PA4762 | PA4762 | N/A |
| PA4763 | N/A | PA4763 | N/A | PA4763 | PA4763 | PA4763 | N/A | PA4763 | N/A | PA4763 | PA4763 | N/A | PA4763 | N/A | PA4763 | N/A | N/A | N/A |
| PA4764 | PA4764 | PA4764 | PA4764 | PA4764 | PA4764 | PA4764 | PA4764 | N/A | PA4764 | PA4764 | PA4764 | PA4764 | PA4764 | PA4764 | PA4764 | PA4764 | PA4764 | PA4764 |
| PA4765 | PA4765 | PA4765 | PA4765 | PA4765 | N/A | PA4765 | PA4765 | PA4765 | PA4765 | PA4765 | PA4765 | PA4765 | PA4765 | PA4765 | PA4765 | PA4765 | N/A | N/A |
| N/A | N/A | N/A | N/A | N/A | PA4766 | N/A | N/A | N/A | N/A | N/A | PA4766 | N/A | N/A | N/A | N/A | N/A | N/A | N/A |
| PA4767 | N/A | N/A | PA4767 | PA4767 | PA4767 | PA4767 | PA4767 | N/A | PA4767 | PA4767 | PA4767 | PA4767 | PA4767 | N/A | N/A | N/A | N/A | N/A |
| PA4768 | PA4768 | PA4768 | PA4768 | PA4768 | PA4768 | PA4768 | N/A | N/A | PA4768 | PA4768 | PA4768 | PA4768 | PA4768 | PA4768 | PA4768 | N/A | N/A | N/A |
| PA4769 | PA4769 | PA4769 | PA4769 | PA4769 | N/A | N/A | PA4769 | N/A | PA4769 | PA4769 | PA4769 | PA4769 | PA4769 | PA4769 | PA4769 | N/A | PA4769 | N/A |
| PA4770 | PA4770 | PA4770 | PA4770 | PA4770 | PA4770 | PA4770 | N/A | N/A | PA4770 | PA4770 | N/A | PA4770 | N/A | PA4770 | N/A | N/A | PA4770 | N/A |
| PA4771 | PA4771 | PA4771 | PA4771 | PA4771 | PA4771 | PA4771 | PA4771 | PA4771 | PA4771 | PA4771 | PA4771 | PA4771 | PA4771 | PA4771 | N/A | N/A | PA4771 | N/A |
| PA4772 | PA4772 | PA4772 | PA4772 | PA4772 | N/A | N/A | PA4772 | N/A | PA4772 | PA4772 | PA4772 | PA4772 | N/A | PA4772 | N/A | N/A | PA4772 | N/A |
| N/A | PA4773 | PA4773 | PA4773 | PA4773 | N/A | N/A | PA4773 | N/A | PA4773 | N/A | PA4773 | N/A | N/A | N/A | N/A | N/A | N/A | N/A |
| N/A | PA4774 | PA4774 | PA4774 | PA4774 | N/A | N/A | N/A | N/A | PA4774 | N/A | N/A | N/A | N/A | N/A | N/A | N/A | N/A | N/A |
| N/A | N/A | N/A | N/A | N/A | PA4775 | N/A | N/A | N/A | PA4775 | N/A | N/A | N/A | N/A | PA4775 | N/A | N/A | N/A | N/A |
| PA4776 | PA4776 | PA4776 | PA4776 | PA4776 | PA4776 | N/A | PA4776 | N/A | PA4776 | PA4776 | PA4776 | N/A | PA4776 | N/A | N/A | N/A | N/A | N/A |
| PA4777 | N/A | N/A | N/A | N/A | PA4777 | N/A | N/A | PA4777 | N/A | N/A | N/A | N/A | N/A | PA4777 | N/A | N/A | PA4777 | N/A |
| PA4778 | N/A | N/A | N/A | PA4778 | N/A | PA4778 | PA4778 | N/A | PA4778 | PA4778 | PA4778 | PA4778 | PA4778 | PA4778 | N/A | N/A | PA4778 | N/A |
| PA4780 | N/A | N/A | N/A | PA4780 | N/A | N/A | N/A | N/A | PA4780 | N/A | N/A | N/A | N/A | N/A | N/A | N/A | N/A | N/A |
| PA4781 | N/A | N/A | N/A | N/A | N/A | N/A | N/A | N/A | N/A | N/A | N/A | N/A | PA4781 | N/A | N/A | N/A | N/A | N/A |
| N/A | N/A | N/A | N/A | N/A | N/A | N/A | N/A | N/A | N/A | N/A | PA4782 | N/A | N/A | N/A | N/A | N/A | N/A | N/A |
| N/A | N/A | N/A | N/A | N/A | N/A | N/A | N/A | N/A | N/A | N/A | N/A | N/A | N/A | PA4783 | N/A | N/A | N/A | N/A |
| PA4784 | N/A | N/A | N/A | PA4784 | N/A | N/A | N/A | N/A | N/A | PA4784 | PA4784 | PA4784 | PA4784 | N/A | N/A | N/A | N/A | N/A |
| PA4785 | N/A | N/A | N/A | PA4785 | PA4785 | PA4785 | N/A | N/A | N/A | PA4785 | N/A | PA4785 | PA4785 | PA4785 | N/A | N/A | PA4785 | N/A |
| PA4786 | N/A | N/A | N/A | PA4786 | N/A | PA4786 | PA4786 | N/A | PA4786 | PA4786 | PA4786 | PA4786 | PA4786 | PA4786 | N/A | N/A | PA4786 | N/A |
| PA4787 | N/A | N/A | N/A | PA4787 | N/A | PA4787 | N/A | PA4787 | N/A | N/A | PA4787 | PA4787 | PA4787 | N/A | N/A | N/A | N/A | N/A |
| PA4788 | N/A | N/A | N/A | N/A | N/A | PA4788 | N/A | N/A | N/A | N/A | PA4788 | PA4788 | PA4788 | N/A | N/A | N/A | PA4788 | N/A |
| PA4790 | N/A | N/A | N/A | PA4790 | N/A | N/A | N/A | N/A | PA4790 | N/A | N/A | N/A | N/A | N/A | N/A | N/A | N/A | N/A |
| PA4791 | N/A | N/A | N/A | N/A | N/A | N/A | N/A | N/A | N/A | N/A | N/A | N/A | N/A | N/A | N/A | N/A | N/A | N/A |
| PA4792 | N/A | N/A | N/A | N/A | N/A | N/A | N/A | N/A | N/A | PA4792 | PA4792 | N/A | N/A | N/A | N/A | PA4792 | N/A | N/A |
| PA4793 | N/A | N/A | N/A | PA4793 | PA4793 | PA4793 | PA4793 | N/A | PA4793 | PA4793 | PA4793 | PA4793 | PA4793 | N/A | N/A | N/A | N/A | N/A |
| PA4794 | N/A | N/A | N/A | N/A | N/A | N/A | N/A | N/A | N/A | N/A | N/A | PA4794 | N/A | N/A | N/A | N/A | N/A | N/A |
| PA4795 | N/A | N/A | N/A | N/A | N/A | N/A | N/A | N/A | N/A | PA4795 | PA4795 | PA4795 | PA4795 | N/A | N/A | N/A | N/A | N/A |
| PA4796 | N/A | N/A | N/A | PA4796 | N/A | N/A | N/A | N/A | PA4796 | PA4796 | PA4796 | PA4796 | PA4796 | N/A | N/A | N/A | N/A | N/A |
| PA4798 | N/A | N/A | N/A | N/A | N/A | N/A | N/A | N/A | PA4798 | N/A | N/A | PA4798 | PA4798 | N/A | N/A | N/A | N/A | N/A |
| PA4802 | N/A | N/A | N/A | N/A | N/A | N/A | N/A | N/A | N/A | N/A | PA4802 | N/A | N/A | N/A | N/A | N/A | N/A | N/A |
| PA4803 | N/A | N/A | N/A | N/A | N/A | N/A | N/A | N/A | N/A | N/A | N/A | N/A | N/A | N/A | N/A | N/A | N/A | N/A |
| PA4807 | N/A | N/A | N/A | PA4807 | N/A | N/A | N/A | N/A | N/A | N/A | N/A | N/A | N/A | PA4807 | N/A | N/A | N/A | N/A |
| PA4808 | N/A | N/A | PA4808 | PA4808 | PA4808 | PA4808 | N/A | PA4808 | PA4808 | PA4808 | PA4808 | PA4808 | PA4808 | PA4808 | N/A | N/A | N/A | N/A |
| PA4809 | PA4809 | N/A | N/A | PA4809 | N/A | N/A | N/A | N/A | N/A | N/A | PA4809 | N/A | N/A | N/A | N/A | N/A | N/A | N/A |
| PA4810 | N/A | N/A | N/A | N/A | N/A | N/A | N/A | N/A | N/A | N/A | N/A | N/A | N/A | PA4810 | N/A | N/A | PA4810 | N/A |
| PA4811 | N/A | N/A | N/A | N/A | N/A | N/A | N/A | N/A | N/A | PA4811 | N/A | N/A | N/A | PA4811 | N/A | N/A | PA4811 | N/A |
| PA4812 | PA4812 | N/A | N/A | PA4812 | N/A | PA4812 | N/A | N/A | PA4812 | PA4812 | PA4812 | PA4812 | PA4812 | PA4812 | N/A | N/A | PA4812 | N/A |
| N/A | N/A | N/A | N/A | N/A | N/A | N/A | N/A | PA4817 | N/A | PA4817 | N/A | N/A | N/A | N/A | N/A | N/A | N/A | N/A |
| N/A | N/A | N/A | N/A | N/A | N/A | N/A | N/A | N/A | PA4824 | N/A | PA4824 | N/A | PA4824 | N/A | N/A | N/A | N/A | N/A |
| N/A | N/A | N/A | N/A | N/A | N/A | N/A | N/A | N/A | PA4825 | N/A | N/A | N/A | PA4825 | PA4825 | N/A | N/A | PA4825 | N/A |
| PA4827 | N/A | N/A | N/A | PA4827 | N/A | N/A | N/A | N/A | N/A | PA4827 | PA4827 | PA4827 | PA4827 | N/A | N/A | N/A | N/A | N/A |
| PA4828 | N/A | N/A | N/A | N/A | N/A | PA4828 | N/A | N/A | N/A | N/A | N/A | N/A | N/A | N/A | N/A | N/A | N/A | N/A |
| PA4829 | N/A | N/A | N/A | N/A | N/A | PA4829 | N/A | N/A | N/A | PA4829 | N/A | N/A | N/A | N/A | N/A | N/A | N/A | N/A |
| PA4831 | N/A | N/A | N/A | N/A | N/A | N/A | N/A | N/A | PA4831 | PA4831 | PA4831 | N/A | PA4831 | N/A | N/A | N/A | N/A | N/A |
| N/A | N/A | N/A | N/A | PA4838 | N/A | N/A | N/A | N/A | N/A | N/A | N/A | N/A | N/A | PA4838 | N/A | N/A | N/A | N/A |
| PA4839 | PA4839 | PA4839 | PA4839 | PA4839 | N/A | PA4839 | N/A | PA4839 | PA4839 | PA4839 | PA4839 | PA4839 | PA4839 | N/A | N/A | PA4839 | N/A | N/A |
| PA4840 | PA4840 | PA4840 | PA4840 | PA4840 | PA4840 | N/A | N/A | N/A | PA4840 | PA4840 | PA4840 | PA4840 | PA4840 | PA4840 | N/A | N/A | N/A | N/A |
| PA4841 | PA4841 | PA4841 | PA4841 | PA4841 | PA4841 | N/A | N/A | PA4841 | PA4841 | N/A | PA4841 | PA4841 | N/A | N/A | N/A | N/A | N/A | N/A |
| PA4842 | PA4842 | PA4842 | PA4842 | PA4842 | PA4842 | PA4842 | PA4842 | PA4842 | PA4842 | PA4842 | PA4842 | PA4842 | PA4842 | PA4842 | PA4842 | PA4842 | PA4842 | PA4842 |
| PA4843 | N/A | N/A | N/A | PA4843 | N/A | N/A | N/A | N/A | N/A | N/A | PA4843 | N/A | PA4843 | N/A | N/A | N/A | N/A | N/A |
| PA4844 | PA4844 | PA4844 | PA4844 | PA4844 | N/A | N/A | N/A | N/A | PA4844 | N/A | N/A | PA4844 | N/A | PA4844 | N/A | N/A | PA4844 | N/A |
| PA4845 | N/A | N/A | N/A | PA4845 | PA4845 | PA4845 | N/A | PA4845 | N/A | PA4845 | PA4845 | PA4845 | PA4845 | PA4845 | N/A | N/A | PA4845 | N/A |
| PA4846 | N/A | N/A | N/A | PA4846 | PA4846 | PA4846 | N/A | N/A | PA4846 | PA4846 | PA4846 | PA4846 | PA4846 | PA4846 | N/A | N/A | N/A | N/A |
| PA4847 | PA4847 | PA4847 | PA4847 | PA4847 | PA4847 | PA4847 | PA4847 | PA4847 | PA4847 | PA4847 | PA4847 | PA4847 | PA4847 | PA4847 | PA4847 | PA4847 | PA4847 | N/A |
| PA4848 | PA4848 | PA4848 | PA4848 | PA4848 | PA4848 | PA4848 | PA4848 | PA4848 | PA4848 | PA4848 | PA4848 | PA4848 | PA4848 | PA4848 | PA4848 | N/A | PA4848 | PA4848 |
| N/A | N/A | N/A | N/A | N/A | PA4849 | N/A | N/A | N/A | N/A | N/A | N/A | N/A | N/A | N/A | N/A | N/A | N/A | N/A |
| PA4850 | PA4850 | PA4850 | PA4850 | PA4850 | PA4850 | N/A | PA4850 | PA4850 | PA4850 | PA4850 | PA4850 | PA4850 | PA4850 | N/A | N/A | N/A | N/A | N/A |
| PA4851 | PA4851 | PA4851 | PA4851 | PA4851 | N/A | PA4851 | PA4851 | N/A | PA4851 | PA4851 | PA4851 | N/A | N/A | PA4851 | N/A | N/A | PA4851 | N/A |

| | | | | | | | | | | | | | | | | | | |
|---|---|---|---|---|---|---|---|---|---|---|---|---|---|---|---|---|---|---|
| N/A | N/A | N/A | N/A | N/A | N/A | N/A | N/A | PA4852 | N/A | PA4852 | N/A | N/A | N/A | N/A | N/A | N/A | N/A | N/A |
| PA4853 | PA4853 | PA4853 | PA4853 | PA4853 | PA4853 | N/A | N/A | N/A | PA4853 | N/A | PA4853 | PA4853 | N/A | N/A | N/A | N/A | N/A | N/A |
| PA4854 | PA4854 | PA4854 | PA4854 | PA4854 | PA4854 | PA4854 | N/A | N/A | PA4854 | PA4854 | PA4854 | PA4854 | PA4854 | PA4854 | PA4854 | N/A | N/A | N/A |
| PA4855 | PA4855 | PA4855 | PA4855 | PA4855 | PA4855 | PA4855 | PA4855 | N/A | PA4855 | PA4855 | PA4855 | PA4855 | PA4855 | N/A | PA4855 | PA4855 | N/A | N/A |
| PA4856 | N/A | N/A | N/A | PA4856 | N/A | N/A | N/A | PA4856 | PA4856 | PA4856 | PA4856 | PA4856 | PA4856 | PA4856 | N/A | N/A | N/A | N/A |
| N/A | N/A | N/A | N/A | N/A | N/A | PA4857 | N/A | N/A | N/A | N/A | N/A | N/A | N/A | N/A | N/A | N/A | N/A | N/A |
| N/A | N/A | N/A | N/A | N/A | N/A | N/A | N/A | N/A | N/A | N/A | N/A | N/A | N/A | N/A | N/A | PA4861 | N/A | N/A |
| PA4865 | N/A | N/A | N/A | N/A | N/A | N/A | N/A | N/A | N/A | PA4865 | PA4865 | PA4865 | PA4865 | N/A | N/A | N/A | N/A | N/A |
| PA4866 | N/A | N/A | N/A | PA4866 | N/A | N/A | PA4866 | N/A | N/A | PA4866 | PA4866 | PA4866 | N/A | N/A | N/A | N/A | N/A | N/A |
| PA4867 | N/A | N/A | N/A | N/A | N/A | N/A | N/A | N/A | N/A | PA4867 | PA4867 | PA4867 | N/A | N/A | N/A | N/A | N/A | N/A |
| PA4868 | N/A | N/A | N/A | PA4868 | N/A | N/A | N/A | N/A | PA4868 | PA4868 | PA4868 | N/A | N/A | PA4868 | N/A | N/A | N/A | N/A |
| PA4869 | N/A | N/A | N/A | N/A | N/A | N/A | N/A | N/A | N/A | N/A | N/A | N/A | N/A | N/A | N/A | N/A | N/A | N/A |
| PA4871 | N/A | N/A | N/A | N/A | PA4871 | N/A | N/A | N/A | N/A | N/A | N/A | N/A | N/A | N/A | N/A | N/A | N/A | N/A |
| PA4872 | PA4872 | PA4872 | PA4872 | PA4872 | PA4872 | PA4872 | N/A | N/A | PA4872 | PA4872 | PA4872 | PA4872 | PA4872 | N/A | PA4872 | N/A | N/A | N/A |
| PA4873 | N/A | PA4873 | PA4873 | PA4873 | N/A | N/A | PA4873 | PA4873 | PA4873 | N/A | PA4873 | PA4873 | N/A | PA4873 | N/A | N/A | N/A | N/A |
| N/A | N/A | N/A | N/A | N/A | PA4875 | N/A | N/A | N/A | N/A | N/A | N/A | N/A | N/A | N/A | N/A | N/A | N/A | N/A |
| PA4876 | N/A | N/A | PA4876 | PA4876 | PA4876 | PA4876 | PA4876 | N/A | PA4876 | PA4876 | PA4876 | PA4876 | PA4876 | PA4876 | N/A | PA4876 | PA4876 | N/A |
| PA4877 | N/A | N/A | N/A | N/A | N/A | N/A | N/A | N/A | N/A | PA4877 | N/A | N/A | PA4877 | N/A | N/A | N/A | N/A | N/A |
| PA4878 | N/A | N/A | N/A | PA4878 | N/A | N/A | N/A | N/A | N/A | N/A | N/A | PA4878 | N/A | N/A | N/A | N/A | N/A | N/A |
| PA4879 | N/A | N/A | N/A | PA4879 | PA4879 | N/A | N/A | N/A | N/A | PA4879 | N/A | N/A | PA4879 | PA4879 | N/A | N/A | PA4879 | N/A |
| PA4880 | PA4880 | PA4880 | PA4880 | PA4880 | PA4880 | PA4880 | N/A | N/A | PA4880 | PA4880 | PA4880 | PA4880 | PA4880 | N/A | N/A | N/A | N/A | N/A |
| N/A | N/A | N/A | N/A | N/A | N/A | N/A | N/A | N/A | N/A | PA4884 | N/A | N/A | N/A | N/A | N/A | N/A | N/A | N/A |
| PA4886 | N/A | N/A | N/A | N/A | N/A | N/A | N/A | N/A | N/A | N/A | N/A | N/A | N/A | N/A | N/A | N/A | N/A | N/A |
| PA4887 | N/A | N/A | N/A | PA4887 | N/A | N/A | N/A | N/A | PA4887 | N/A | N/A | PA4887 | N/A | PA4887 | PA4887 | N/A | N/A | N/A |
| N/A | N/A | N/A | N/A | N/A | N/A | N/A | N/A | N/A | PA4888 | N/A | N/A | N/A | N/A | PA4888 | N/A | N/A | N/A | N/A |
| N/A | N/A | PA4889 | N/A | PA4889 | N/A | N/A | PA4889 | PA4889 | PA4889 | N/A | N/A | N/A | N/A | PA4889 | N/A | N/A | N/A | N/A |
| PA4890 | PA4890 | PA4890 | PA4890 | PA4890 | N/A | N/A | N/A | N/A | PA4890 | PA4890 | PA4890 | PA4890 | N/A | N/A | N/A | N/A | N/A | N/A |
| N/A | N/A | N/A | N/A | PA4893 | N/A | N/A | N/A | N/A | N/A | N/A | N/A | N/A | N/A | N/A | N/A | N/A | N/A | N/A |
| PA4894 | N/A | N/A | N/A | N/A | N/A | N/A | N/A | N/A | N/A | N/A | N/A | N/A | N/A | N/A | N/A | N/A | N/A | N/A |
| PA4899 | PA4899 | PA4899 | PA4899 | PA4899 | PA4899 | N/A | N/A | N/A | N/A | PA4899 | N/A | N/A | N/A | N/A | N/A | N/A | N/A | N/A |
| N/A | N/A | N/A | N/A | PA4901 | N/A | N/A | N/A | N/A | N/A | N/A | N/A | N/A | N/A | N/A | N/A | N/A | N/A | N/A |
| PA4902 | N/A | N/A | N/A | PA4902 | N/A | N/A | N/A | N/A | N/A | N/A | PA4902 | N/A | N/A | N/A | N/A | N/A | N/A | N/A |
| PA4906 | N/A | N/A | N/A | PA4906 | PA4906 | N/A | N/A | N/A | N/A | N/A | N/A | N/A | N/A | N/A | N/A | N/A | N/A | N/A |
| PA4907 | PA4907 | PA4907 | PA4907 | PA4907 | PA4907 | PA4907 | PA4907 | N/A | PA4907 | PA4907 | PA4907 | PA4907 | PA4907 | N/A | PA4907 | N/A | N/A | N/A |
| PA4909 | N/A | N/A | N/A | N/A | N/A | N/A | N/A | N/A | N/A | N/A | N/A | PA4909 | N/A | N/A | N/A | N/A | N/A | N/A |
| PA4911 | N/A | N/A | N/A | N/A | N/A | N/A | N/A | N/A | N/A | N/A | N/A | N/A | N/A | PA4911 | N/A | N/A | N/A | N/A |
| N/A | N/A | N/A | N/A | N/A | PA4912 | N/A | N/A | N/A | N/A | N/A | N/A | N/A | N/A | N/A | N/A | N/A | N/A | N/A |
| PA4913 | N/A | N/A | N/A | N/A | PA4913 | PA4913 | N/A | N/A | N/A | N/A | N/A | PA4913 | PA4913 | N/A | N/A | N/A | N/A | N/A |
| PA4914 | N/A | N/A | N/A | N/A | N/A | N/A | N/A | N/A | N/A | N/A | N/A | PA4914 | PA4914 | PA4914 | N/A | N/A | PA4914 | N/A |
| PA4915 | PA4915 | PA4915 | PA4915 | PA4915 | PA4915 | PA4915 | PA4915 | PA4915 | PA4915 | PA4915 | PA4915 | PA4915 | PA4915 | PA4915 | PA4915 | N/A | PA4915 | PA4915 |
| PA4916 | N/A | PA4916 | PA4916 | PA4916 | PA4916 | PA4916 | PA4916 | N/A | PA4916 | PA4916 | PA4916 | PA4916 | PA4916 | N/A | PA4916 | PA4916 | N/A | N/A |
| PA4918 | N/A | PA4918 | N/A | PA4918 | PA4918 | N/A | N/A | N/A | PA4918 | PA4918 | PA4918 | PA4918 | PA4918 | N/A | N/A | N/A | N/A | N/A |
| PA4919 | PA4919 | PA4919 | PA4919 | PA4919 | PA4919 | PA4919 | PA4919 | PA4919 | PA4919 | PA4919 | PA4919 | PA4919 | PA4919 | N/A | N/A | N/A | N/A | N/A |
| PA4920 | PA4920 | PA4920 | PA4920 | PA4920 | PA4920 | PA4920 | PA4920 | PA4920 | PA4920 | PA4920 | PA4920 | PA4920 | PA4920 | N/A | PA4920 | PA4920 | PA4920 | N/A |
| N/A | N/A | N/A | N/A | N/A | PA4921 | N/A | N/A | N/A | N/A | N/A | N/A | N/A | N/A | N/A | N/A | N/A | N/A | N/A |
| PA4922 | PA4922 | PA4922 | PA4922 | PA4922 | PA4922 | PA4922 | N/A | N/A | PA4922 | PA4922 | PA4922 | PA4922 | PA4922 | N/A | PA4922 | PA4922 | N/A | PA4922 |
| PA4923 | PA4923 | PA4923 | PA4923 | PA4923 | PA4923 | N/A | PA4923 | N/A | PA4923 | PA4923 | PA4923 | PA4923 | PA4923 | N/A | PA4923 | N/A | N/A | N/A |
| PA4924 | N/A | N/A | N/A | N/A | N/A | N/A | N/A | N/A | PA4924 | N/A | N/A | N/A | N/A | N/A | PA4924 | PA4924 | PA4924 | PA4924 |
| PA4925 | N/A | N/A | N/A | N/A | N/A | PA4925 | N/A | N/A | N/A | N/A | N/A | PA4925 | PA4925 | N/A | N/A | N/A | N/A | N/A |
| N/A | PA4928 | PA4928 | N/A | PA4928 | N/A | N/A | N/A | PA4928 | PA4928 | N/A | PA4928 | PA4928 | N/A | PA4928 | N/A | N/A | N/A | N/A |
| PA4929 | N/A | N/A | N/A | N/A | PA4929 | N/A | N/A | N/A | N/A | N/A | N/A | N/A | N/A | N/A | N/A | N/A | N/A | N/A |
| PA4930 | PA4930 | PA4930 | PA4930 | PA4930 | PA4930 | N/A | N/A | N/A | PA4930 | PA4930 | PA4930 | PA4930 | PA4930 | N/A | PA4930 | N/A | N/A | N/A |
| PA4931 | PA4931 | PA4931 | PA4931 | PA4931 | PA4931 | N/A | N/A | PA4931 | PA4931 | PA4931 | PA4931 | PA4931 | PA4931 | N/A | N/A | N/A | PA4931 | N/A |
| PA4932 | PA4932 | PA4932 | PA4932 | PA4932 | PA4932 | PA4932 | PA4932 | PA4932 | PA4932 | PA4932 | PA4932 | PA4932 | PA4932 | PA4932 | PA4932 | PA4932 | PA4932 | PA4932 |
| PA4933 | N/A | N/A | N/A | PA4933 | PA4933 | N/A | N/A | PA4933 | PA4933 | PA4933 | PA4933 | PA4933 | PA4933 | PA4933 | N/A | N/A | PA4933 | N/A |
| PA4934 | PA4934 | PA4934 | PA4934 | PA4934 | PA4934 | PA4934 | PA4934 | PA4934 | PA4934 | PA4934 | PA4934 | PA4934 | PA4934 | PA4934 | PA4934 | N/A | PA4934 | PA4934 |
| PA4935 | PA4935 | PA4935 | PA4935 | PA4935 | PA4935 | PA4935 | PA4935 | PA4935 | PA4935 | PA4935 | PA4935 | PA4935 | PA4935 | PA4935 | PA4935 | PA4935 | PA4935 | PA4935 |
| PA4936 | PA4936 | PA4936 | N/A | PA4936 | N/A | PA4936 | PA4936 | PA4936 | PA4936 | PA4936 | PA4936 | PA4936 | PA4936 | PA4936 | N/A | N/A | N/A | N/A |
| PA4937 | PA4937 | PA4937 | PA4937 | PA4937 | PA4937 | PA4937 | N/A | PA4937 | PA4937 | PA4937 | PA4937 | PA4937 | PA4937 | PA4937 | PA4937 | N/A | PA4937 | N/A |
| PA4938 | PA4938 | PA4938 | PA4938 | PA4938 | PA4938 | PA4938 | PA4938 | PA4938 | PA4938 | PA4938 | PA4938 | PA4938 | PA4938 | PA4938 | PA4938 | PA4938 | N/A | N/A |
| PA4939 | PA4939 | PA4939 | PA4939 | PA4939 | PA4939 | PA4939 | PA4939 | N/A | PA4939 | PA4939 | PA4939 | PA4939 | PA4939 | PA4939 | PA4939 | PA4939 | N/A | N/A |
| N/A | N/A | N/A | N/A | N/A | PA4940 | N/A | N/A | N/A | N/A | N/A | N/A | N/A | N/A | N/A | N/A | N/A | N/A | N/A |
| PA4941 | PA4941 | PA4941 | PA4941 | PA4941 | PA4941 | PA4941 | PA4941 | PA4941 | PA4941 | PA4941 | PA4941 | PA4941 | PA4941 | PA4941 | PA4941 | N/A | PA4941 | PA4941 |
| PA4942 | PA4942 | PA4942 | PA4942 | PA4942 | PA4942 | PA4942 | PA4942 | PA4942 | PA4942 | PA4942 | PA4942 | PA4942 | PA4942 | PA4942 | PA4942 | N/A | PA4942 | PA4942 |
| PA4943 | PA4943 | PA4943 | PA4943 | PA4943 | N/A | N/A | PA4943 | PA4943 | PA4943 | N/A | PA4943 | PA4943 | PA4943 | PA4943 | N/A | N/A | N/A | N/A |
| PA4944 | PA4944 | PA4944 | PA4944 | PA4944 | PA4944 | PA4944 | PA4944 | N/A | PA4944 | PA4944 | PA4944 | PA4944 | PA4944 | PA4944 | PA4944 | PA4944 | PA4944 | PA4944 |
| PA4945 | N/A | N/A | N/A | PA4945 | N/A | N/A | N/A | N/A | N/A | N/A | N/A | N/A | N/A | N/A | N/A | N/A | N/A | N/A |
| PA4946 | PA4946 | PA4946 | PA4946 | PA4946 | N/A | N/A | N/A | PA4946 | PA4946 | PA4946 | PA4946 | PA4946 | PA4946 | N/A | N/A | N/A | PA4946 | N/A |
| PA4947 | PA4947 | PA4947 | PA4947 | PA4947 | PA4947 | N/A | N/A | PA4947 | PA4947 | N/A | PA4947 | PA4947 | PA4947 | PA4947 | PA4947 | PA4947 | PA4947 | PA4947 |
| PA4948 | N/A | N/A | PA4948 | PA4948 | PA4948 | N/A | PA4948 | N/A | PA4948 | PA4948 | PA4948 | PA4948 | PA4948 | N/A | N/A | N/A | N/A | N/A |
| PA4949 | PA4949 | N/A | PA4949 | PA4949 | PA4949 | PA4949 | N/A | PA4949 | PA4949 | PA4949 | PA4949 | PA4949 | PA4949 | N/A | PA4949 | N/A | N/A | N/A |
| N/A | N/A | N/A | N/A | PA4950 | N/A | N/A | N/A | N/A | N/A | N/A | N/A | N/A | N/A | N/A | N/A | N/A | N/A | N/A |
| PA4951 | PA4951 | PA4951 | PA4951 | PA4951 | PA4951 | PA4951 | N/A | N/A | PA4951 | PA4951 | PA4951 | PA4951 | PA4951 | N/A | PA4951 | N/A | N/A | N/A |
| PA4952 | N/A | PA4952 | N/A | PA4952 | N/A | N/A | N/A | N/A | PA4952 | PA4952 | N/A | N/A | N/A | PA4952 | N/A | N/A | N/A | N/A |
| PA4953 | PA4953 | N/A | N/A | PA4953 | N/A | N/A | PA4953 | N/A | PA4953 | PA4953 | N/A | PA4953 | PA4953 | PA4953 | PA4953 | N/A | PA4953 | N/A |
| PA4954 | N/A | N/A | N/A | N/A | N/A | PA4954 | N/A | N/A | PA4954 | PA4954 | PA4954 | PA4954 | PA4954 | PA4954 | N/A | N/A | PA4954 | N/A |
| PA4955 | N/A | N/A | N/A | PA4955 | PA4955 | N/A | N/A | N/A | N/A | PA4955 | N/A | PA4955 | PA4955 | N/A | N/A | N/A | N/A | N/A |
| PA4956 | PA4956 | PA4956 | PA4956 | PA4956 | PA4956 | PA4956 | PA4956 | N/A | PA4956 | PA4956 | PA4956 | PA4956 | PA4956 | N/A | PA4956 | N/A | N/A | N/A |
| PA4957 | PA4957 | PA4957 | PA4957 | PA4957 | PA4957 | PA4957 | PA4957 | N/A | PA4957 | PA4957 | PA4957 | PA4957 | PA4957 | PA4957 | PA4957 | N/A | PA4957 | N/A |
| PA4958 | N/A | N/A | PA4958 | PA4958 | PA4958 | N/A | N/A | PA4958 | PA4958 | PA4958 | PA4958 | PA4958 | PA4958 | PA4958 | N/A | N/A | N/A | N/A |
| PA4959 | PA4959 | PA4959 | PA4959 | PA4959 | PA4959 | PA4959 | N/A | PA4959 | PA4959 | PA4959 | PA4959 | PA4959 | PA4959 | N/A | N/A | N/A | N/A | N/A |
| PA4960 | PA4960 | PA4960 | PA4960 | PA4960 | PA4960 | N/A | N/A | PA4960 | PA4960 | PA4960 | PA4960 | PA4960 | PA4960 | N/A | PA4960 | N/A | N/A | N/A |
| PA4961 | PA4961 | PA4961 | PA4961 | PA4961 | N/A | PA4961 | PA4961 | PA4961 | PA4961 | PA4961 | PA4961 | PA4961 | PA4961 | PA4961 | N/A | N/A | PA4961 | N/A |
| PA4962 | N/A | N/A | N/A | N/A | N/A | N/A | N/A | N/A | N/A | N/A | N/A | N/A | N/A | N/A | N/A | N/A | N/A | N/A |
| PA4963 | PA4963 | PA4963 | PA4963 | PA4963 | PA4963 | N/A | PA4963 | N/A | PA4963 | PA4963 | PA4963 | PA4963 | PA4963 | PA4963 | PA4963 | N/A | N/A | N/A |
| PA4964 | PA4964 | PA4964 | PA4964 | PA4964 | N/A | PA4964 | N/A | PA4964 | PA4964 | PA4964 | PA4964 | PA4964 | PA4964 | PA4964 | N/A | N/A | N/A | N/A |
| PA4965 | PA4965 | N/A | N/A | N/A | PA4965 | N/A | N/A | N/A | N/A | N/A | N/A | N/A | N/A | PA4965 | N/A | N/A | PA4965 | N/A |
| PA4966 | PA4966 | PA4966 | N/A | PA4966 | PA4966 | N/A | PA4966 | N/A | PA4966 | PA4966 | PA4966 | PA4966 | PA4966 | PA4966 | N/A | N/A | N/A | N/A |
| PA4967 | PA4967 | PA4967 | PA4967 | PA4967 | PA4967 | PA4967 | N/A | PA4967 | PA4967 | PA4967 | PA4967 | PA4967 | PA4967 | PA4967 | N/A | N/A | PA4967 | N/A |
| PA4968 | PA4968 | PA4968 | N/A | PA4968 | PA4968 | N/A | N/A | PA4968 | PA4968 | PA4968 | PA4968 | N/A | N/A | N/A | N/A | N/A | N/A | N/A |
| PA4969 | N/A | N/A | N/A | PA4969 | PA4969 | N/A | N/A | N/A | PA4969 | PA4969 | PA4969 | PA4969 | PA4969 | N/A | N/A | N/A | N/A | N/A |
| N/A | N/A | N/A | N/A | N/A | PA4970 | N/A | N/A | N/A | N/A | N/A | N/A | N/A | N/A | N/A | N/A | N/A | N/A | N/A |
| PA4971 | PA4971 | PA4971 | PA4971 | PA4971 | PA4971 | PA4971 | PA4971 | N/A | PA4971 | PA4971 | PA4971 | PA4971 | PA4971 | N/A | PA4971 | N/A | N/A | N/A |
| PA4972 | PA4972 | PA4972 | N/A | PA4972 | N/A | PA4972 | PA4972 | N/A | PA4972 | PA4972 | PA4972 | PA4972 | PA4972 | PA4972 | PA4972 | PA4972 | N/A | N/A |
| PA4973 | PA4973 | N/A | N/A | PA4973 | PA4973 | N/A | N/A | N/A | N/A | N/A | N/A | N/A | PA4973 | N/A | N/A | N/A | N/A | N/A |
| PA4974 | PA4974 | PA4974 | PA4974 | PA4974 | N/A | PA4974 | PA4974 | PA4974 | PA4974 | PA4974 | PA4974 | PA4974 | PA4974 | PA4974 | PA4974 | PA4974 | PA4974 | PA4974 |
| PA4975 | N/A | N/A | N/A | PA4975 | PA4975 | N/A | N/A | N/A | N/A | N/A | PA4975 | PA4975 | N/A | N/A | N/A | N/A | N/A | N/A |
| PA4976 | N/A | N/A | N/A | PA4976 | PA4976 | N/A | N/A | N/A | N/A | N/A | PA4976 | N/A | PA4976 | N/A | N/A | N/A | N/A | N/A |
| PA4977 | N/A | N/A | N/A | PA4977 | N/A | N/A | N/A | N/A | N/A | N/A | N/A | N/A | N/A | N/A | N/A | N/A | N/A | N/A |
| PA4978 | N/A | N/A | N/A | PA4978 | N/A | N/A | N/A | N/A | N/A | N/A | N/A | N/A | PA4978 | N/A | N/A | N/A | N/A | N/A |
| N/A | N/A | N/A | N/A | PA4980 | N/A | N/A | N/A | N/A | N/A | N/A | N/A | N/A | N/A | N/A | N/A | N/A | N/A | N/A |
| PA4983 | N/A | N/A | N/A | PA4983 | N/A | N/A | N/A | N/A | N/A | N/A | N/A | N/A | N/A | N/A | N/A | N/A | N/A | N/A |
| N/A | N/A | N/A | N/A | PA4986 | N/A | N/A | N/A | N/A | N/A | N/A | N/A | N/A | N/A | N/A | N/A | N/A | N/A | N/A |
| PA4987 | N/A | N/A | N/A | N/A | N/A | N/A | N/A | N/A | PA4987 | PA4987 | N/A | PA4987 | N/A | N/A | N/A | N/A | N/A | N/A |
| PA4988 | PA4988 | N/A | PA4988 | PA4988 | N/A | N/A | N/A | N/A | N/A | PA4988 | N/A | N/A | PA4988 | PA4988 | N/A | N/A | PA4988 | N/A |
| PA4989 | N/A | N/A | N/A | N/A | N/A | N/A | N/A | N/A | N/A | N/A | N/A | N/A | N/A | N/A | N/A | N/A | N/A | N/A |
| N/A | N/A | N/A | N/A | N/A | PA4990 | N/A | N/A | N/A | N/A | N/A | N/A | N/A | N/A | N/A | N/A | N/A | N/A | N/A |
| PA4991 | PA4991 | PA4991 | PA4991 | PA4991 | PA4991 | N/A | PA4991 | PA4991 | PA4991 | PA4991 | PA4991 | PA4991 | PA4991 | PA4991 | N/A | N/A | N/A | N/A |
| PA4992 | N/A | N/A | N/A | PA4992 | PA4992 | N/A | PA4992 | N/A | PA4992 | PA4992 | PA4992 | PA4992 | PA4992 | N/A | N/A | N/A | N/A | N/A |

| | | | | | | | | | | | | | | | | | | |
|---|---|---|---|---|---|---|---|---|---|---|---|---|---|---|---|---|---|---|
| PA4993 | N/A | N/A | N/A | N/A | N/A | N/A | N/A | N/A | N/A | N/A | N/A | N/A | N/A | N/A | N/A | N/A | N/A | N/A |
| N/A | N/A | N/A | N/A | N/A | PA4995 | N/A | N/A | N/A | N/A | N/A | N/A | N/A | N/A | N/A | N/A | N/A | N/A | N/A |
| PA4996 | PA4996 | PA4996 | PA4996 | PA4996 | N/A | PA4996 | PA4996 | PA4996 | PA4996 | PA4996 | PA4996 | PA4996 | PA4996 | N/A | PA4996 | N/A | PA4996 | N/A |
| PA4997 | PA4997 | PA4997 | PA4997 | PA4997 | PA4997 | PA4997 | PA4997 | PA4997 | PA4997 | PA4997 | PA4997 | PA4997 | PA4997 | PA4997 | PA4997 | N/A | PA4997 | N/A |
| PA4998 | PA4998 | PA4998 | N/A | PA4998 | PA4998 | N/A | PA4998 | PA4998 | PA4998 | PA4998 | PA4998 | PA4998 | PA4998 | PA4998 | N/A | N/A | PA4998 | PA4998 |
| PA4999 | N/A | N/A | N/A | PA4999 | N/A | N/A | N/A | N/A | N/A | N/A | N/A | PA4999 | N/A | PA4999 | N/A | N/A | N/A | N/A |
| PA5000 | N/A | PA5000 | N/A | PA5000 | N/A | N/A | N/A | PA5000 | PA5000 | PA5000 | PA5000 | PA5000 | PA5000 | PA5000 | N/A | N/A | PA5000 | N/A |
| PA5001 | PA5001 | PA5001 | PA5001 | PA5001 | N/A | PA5001 | PA5001 | PA5001 | PA5001 | PA5001 | PA5001 | PA5001 | PA5001 | PA5001 | N/A | N/A | PA5001 | N/A |
| PA5002 | N/A | PA5002 | N/A | PA5002 | N/A | N/A | PA5002 | PA5002 | PA5002 | PA5002 | PA5002 | PA5002 | PA5002 | N/A | PA5002 | N/A | N/A | N/A |
| PA5003 | PA5003 | N/A | PA5003 | PA5003 | N/A | N/A | PA5003 | N/A | PA5003 | PA5003 | PA5003 | PA5003 | N/A | PA5003 | N/A | N/A | PA5003 | N/A |
| PA5004 | PA5004 | PA5004 | PA5004 | PA5004 | N/A | PA5004 | PA5004 | PA5004 | PA5004 | PA5004 | PA5004 | PA5004 | PA5004 | PA5004 | PA5004 | N/A | PA5004 | N/A |
| PA5005 | PA5005 | PA5005 | PA5005 | PA5005 | PA5005 | PA5005 | PA5005 | PA5005 | PA5005 | PA5005 | PA5005 | PA5005 | PA5005 | PA5005 | PA5005 | N/A | PA5005 | N/A |
| PA5006 | PA5006 | PA5006 | N/A | PA5006 | N/A | PA5006 | PA5006 | PA5006 | PA5006 | PA5006 | PA5006 | PA5006 | PA5006 | PA5006 | PA5006 | N/A | PA5006 | N/A |
| PA5007 | PA5007 | PA5007 | PA5007 | PA5007 | N/A | N/A | PA5007 | N/A | PA5007 | N/A | PA5007 | PA5007 | N/A | PA5007 | N/A | N/A | N/A | N/A |
| PA5008 | N/A | PA5008 | N/A | PA5008 | N/A | N/A | PA5008 | N/A | N/A | N/A | N/A | N/A | N/A | PA5008 | PA5008 | N/A | N/A | N/A |
| PA5009 | N/A | N/A | N/A | PA5009 | PA5009 | N/A | PA5009 | N/A | PA5009 | N/A | PA5009 | N/A | N/A | PA5009 | N/A | N/A | N/A | N/A |
| PA5010 | PA5010 | PA5010 | N/A | PA5010 | PA5010 | N/A | PA5010 | N/A | N/A | N/A | PA5010 | N/A | N/A | PA5010 | N/A | N/A | N/A | N/A |
| PA5011 | PA5011 | PA5011 | PA5011 | PA5011 | N/A | N/A | PA5011 | N/A | PA5011 | PA5011 | PA5011 | PA5011 | N/A | PA5011 | N/A | N/A | PA5011 | N/A |
| PA5012 | PA5012 | PA5012 | PA5012 | PA5012 | PA5012 | N/A | PA5012 | N/A | PA5012 | PA5012 | N/A | PA5012 | N/A | N/A | N/A | N/A | PA5012 | N/A |
| PA5013 | PA5013 | PA5013 | PA5013 | PA5013 | PA5013 | PA5013 | N/A | N/A | PA5013 | PA5013 | PA5013 | PA5013 | PA5013 | PA5013 | PA5013 | PA5013 | PA5013 | N/A |
| PA5014 | PA5014 | N/A | N/A | PA5014 | PA5014 | N/A | N/A | PA5014 | PA5014 | PA5014 | PA5014 | PA5014 | PA5014 | N/A | N/A | N/A | N/A | N/A |
| PA5015 | PA5015 | PA5015 | PA5015 | PA5015 | PA5015 | PA5015 | PA5015 | PA5015 | PA5015 | PA5015 | PA5015 | PA5015 | PA5015 | PA5015 | PA5015 | PA5015 | N/A | N/A |
| PA5016 | PA5016 | PA5016 | PA5016 | PA5016 | PA5016 | PA5016 | PA5016 | PA5016 | PA5016 | PA5016 | PA5016 | PA5016 | PA5016 | PA5016 | PA5016 | PA5016 | PA5016 | N/A |
| PA5017 | N/A | N/A | N/A | PA5017 | PA5017 | N/A | N/A | PA5017 | PA5017 | PA5017 | N/A | PA5017 | PA5017 | N/A | N/A | N/A | N/A | N/A |
| PA5018 | PA5018 | PA5018 | PA5018 | PA5018 | PA5018 | PA5018 | N/A | N/A | PA5018 | PA5018 | PA5018 | PA5018 | PA5018 | N/A | N/A | N/A | N/A | N/A |
| PA5019 | PA5019 | PA5019 | PA5019 | PA5019 | N/A | N/A | PA5019 | N/A | PA5019 | PA5019 | PA5019 | PA5019 | PA5019 | N/A | PA5019 | N/A | N/A | N/A |
| PA5020 | PA5020 | PA5020 | PA5020 | PA5020 | N/A | PA5020 | N/A | N/A | PA5020 | PA5020 | PA5020 | PA5020 | PA5020 | PA5020 | N/A | N/A | N/A | N/A |
| PA5021 | N/A | N/A | N/A | PA5021 | N/A | N/A | N/A | N/A | N/A | N/A | N/A | N/A | PA5021 | PA5021 | N/A | N/A | N/A | N/A |
| PA5022 | PA5022 | PA5022 | N/A | PA5022 | N/A | PA5022 | N/A | PA5022 | PA5022 | PA5022 | PA5022 | PA5022 | PA5022 | PA5022 | PA5022 | N/A | PA5022 | N/A |
| N/A | PA5023 | N/A | N/A | PA5023 | N/A | N/A | N/A | PA5023 | PA5023 | N/A | N/A | N/A | PA5023 | N/A | N/A | N/A | N/A | N/A |
| N/A | N/A | N/A | N/A | N/A | PA5024 | N/A | N/A | N/A | N/A | N/A | N/A | N/A | N/A | PA5024 | N/A | N/A | N/A | N/A |
| PA5025 | PA5025 | PA5025 | PA5025 | PA5025 | PA5025 | PA5025 | N/A | N/A | PA5025 | PA5025 | PA5025 | PA5025 | PA5025 | PA5025 | N/A | N/A | N/A | N/A |
| N/A | N/A | N/A | N/A | N/A | PA5026 | N/A | N/A | N/A | N/A | N/A | N/A | N/A | N/A | N/A | N/A | N/A | N/A | N/A |
| PA5027 | N/A | N/A | PA5027 | PA5027 | PA5027 | N/A | PA5027 | PA5027 | N/A | PA5027 | PA5027 | PA5027 | PA5027 | PA5027 | N/A | N/A | PA5027 | PA5027 |
| PA5028 | N/A | N/A | N/A | N/A | N/A | PA5028 | PA5028 | PA5028 | PA5028 | PA5028 | PA5028 | PA5028 | PA5028 | N/A | N/A | N/A | N/A | N/A |
| N/A | N/A | N/A | N/A | N/A | N/A | N/A | N/A | N/A | N/A | N/A | N/A | N/A | N/A | N/A | PA5033 | PA5033 | N/A | N/A |
| PA5034 | N/A | PA5034 | N/A | PA5034 | N/A | N/A | N/A | N/A | PA5034 | PA5034 | PA5034 | PA5034 | PA5034 | N/A | PA5034 | N/A | N/A | N/A |
| PA5035 | PA5035 | N/A | PA5035 | PA5035 | PA5035 | N/A | N/A | PA5035 | PA5035 | PA5035 | PA5035 | PA5035 | PA5035 | PA5035 | N/A | N/A | N/A | N/A |
| PA5036 | PA5036 | PA5036 | N/A | PA5036 | N/A | PA5036 | N/A | N/A | PA5036 | PA5036 | PA5036 | PA5036 | PA5036 | PA5036 | N/A | N/A | N/A | N/A |
| PA5037 | PA5037 | PA5037 | PA5037 | PA5037 | PA5037 | PA5037 | PA5037 | PA5037 | PA5037 | PA5037 | PA5037 | PA5037 | PA5037 | PA5037 | PA5037 | N/A | PA5037 | PA5037 |
| PA5038 | PA5038 | PA5038 | PA5038 | PA5038 | PA5038 | PA5038 | PA5038 | PA5038 | PA5038 | PA5038 | PA5038 | PA5038 | PA5038 | PA5038 | PA5038 | N/A | N/A | N/A |
| PA5039 | N/A | N/A | N/A | PA5039 | PA5039 | N/A | N/A | N/A | PA5039 | PA5039 | PA5039 | PA5039 | PA5039 | N/A | N/A | N/A | N/A | N/A |
| PA5040 | PA5040 | PA5040 | PA5040 | PA5040 | PA5040 | PA5040 | PA5040 | PA5040 | PA5040 | PA5040 | PA5040 | PA5040 | PA5040 | PA5040 | PA5040 | N/A | PA5040 | N/A |
| PA5041 | N/A | PA5041 | N/A | PA5041 | N/A | N/A | PA5041 | N/A | PA5041 | PA5041 | PA5041 | PA5041 | PA5041 | PA5041 | N/A | N/A | PA5041 | N/A |
| PA5042 | PA5042 | PA5042 | PA5042 | PA5042 | N/A | PA5042 | PA5042 | N/A | PA5042 | PA5042 | PA5042 | PA5042 | PA5042 | PA5042 | PA5042 | N/A | PA5042 | N/A |
| PA5043 | PA5043 | PA5043 | PA5043 | PA5043 | N/A | PA5043 | PA5043 | N/A | PA5043 | PA5043 | PA5043 | PA5043 | PA5043 | PA5043 | PA5043 | N/A | PA5043 | N/A |
| PA5044 | PA5044 | PA5044 | PA5044 | PA5044 | N/A | PA5044 | N/A | PA5044 | PA5044 | PA5044 | PA5044 | PA5044 | PA5044 | PA5044 | N/A | N/A | PA5044 | N/A |
| PA5045 | PA5045 | PA5045 | PA5045 | PA5045 | PA5045 | PA5045 | N/A | PA5045 | PA5045 | PA5045 | PA5045 | PA5045 | PA5045 | PA5045 | PA5045 | N/A | PA5045 | N/A |
| PA5046 | PA5046 | PA5046 | PA5046 | PA5046 | PA5046 | PA5046 | PA5046 | PA5046 | PA5046 | PA5046 | PA5046 | PA5046 | PA5046 | PA5046 | PA5046 | PA5046 | N/A | N/A |
| PA5047 | N/A | N/A | N/A | PA5047 | N/A | N/A | N/A | N/A | N/A | N/A | N/A | N/A | N/A | PA5047 | PA5047 | N/A | N/A | N/A |
| N/A | N/A | N/A | N/A | PA5048 | PA5048 | N/A | N/A | N/A | N/A | N/A | N/A | N/A | N/A | N/A | N/A | N/A | N/A | N/A |
| PA5049 | PA5049 | PA5049 | PA5049 | PA5049 | PA5049 | PA5049 | PA5049 | PA5049 | PA5049 | PA5049 | PA5049 | PA5049 | PA5049 | PA5049 | PA5049 | PA5049 | PA5049 | PA5049 |
| PA5050 | N/A | N/A | N/A | PA5050 | N/A | N/A | N/A | PA5050 | N/A | N/A | PA5050 | PA5050 | PA5050 | N/A | N/A | N/A | N/A | N/A |
| PA5051 | PA5051 | PA5051 | PA5051 | PA5051 | PA5051 | PA5051 | PA5051 | PA5051 | PA5051 | PA5051 | PA5051 | PA5051 | PA5051 | PA5051 | PA5051 | PA5051 | N/A | N/A |
| PA5052 | PA5052 | PA5052 | N/A | PA5052 | N/A | PA5052 | PA5052 | PA5052 | PA5052 | PA5052 | PA5052 | PA5052 | PA5052 | PA5052 | PA5052 | PA5052 | PA5052 | N/A |
| PA5053 | PA5053 | PA5053 | PA5053 | PA5053 | PA5053 | PA5053 | N/A | N/A | PA5053 | PA5053 | PA5053 | PA5053 | PA5053 | PA5053 | PA5053 | N/A | N/A | N/A |
| PA5054 | PA5054 | PA5054 | PA5054 | PA5054 | N/A | PA5054 | PA5054 | PA5054 | PA5054 | PA5054 | PA5054 | PA5054 | PA5054 | PA5054 | PA5054 | N/A | PA5054 | N/A |
| PA5055 | N/A | PA5055 | N/A | PA5055 | PA5055 | N/A | N/A | N/A | PA5055 | N/A | PA5055 | PA5055 | PA5055 | N/A | PA5055 | N/A | N/A | N/A |
| PA5056 | N/A | N/A | N/A | N/A | N/A | PA5056 | N/A | N/A | N/A | N/A | PA5056 | PA5056 | PA5056 | N/A | N/A | N/A | PA5056 | N/A |
| PA5058 | N/A | N/A | N/A | N/A | N/A | N/A | N/A | N/A | PA5058 | N/A | PA5058 | PA5058 | PA5058 | N/A | N/A | N/A | N/A | N/A |
| PA5059 | N/A | N/A | N/A | N/A | PA5059 | PA5059 | N/A | N/A | PA5059 | N/A | N/A | PA5059 | PA5059 | N/A | N/A | N/A | N/A | N/A |
| PA5060 | PA5060 | PA5060 | PA5060 | PA5060 | PA5060 | PA5060 | N/A | PA5060 | PA5060 | PA5060 | PA5060 | PA5060 | PA5060 | PA5060 | PA5060 | PA5060 | PA5060 | PA5060 |
| PA5061 | N/A | N/A | N/A | N/A | N/A | PA5061 | N/A | N/A | PA5061 | PA5061 | PA5061 | PA5061 | PA5061 | PA5061 | N/A | N/A | PA5061 | N/A |
| PA5062 | PA5062 | PA5062 | N/A | PA5062 | PA5062 | N/A | N/A | N/A | PA5062 | N/A | PA5062 | PA5062 | PA5062 | PA5062 | PA5062 | N/A | N/A | N/A |
| PA5063 | PA5063 | PA5063 | PA5063 | PA5063 | PA5063 | PA5063 | PA5063 | PA5063 | PA5063 | PA5063 | PA5063 | PA5063 | PA5063 | PA5063 | N/A | N/A | PA5063 | PA5063 |
| PA5064 | PA5064 | PA5064 | PA5064 | PA5064 | PA5064 | PA5064 | PA5064 | N/A | PA5064 | PA5064 | PA5064 | PA5064 | PA5064 | PA5064 | N/A | N/A | N/A | N/A |
| PA5065 | PA5065 | PA5065 | PA5065 | PA5065 | N/A | PA5065 | N/A | PA5065 | PA5065 | PA5065 | PA5065 | PA5065 | PA5065 | PA5065 | PA5065 | N/A | PA5065 | N/A |
| PA5066 | PA5066 | PA5066 | N/A | PA5066 | PA5066 | PA5066 | N/A | N/A | PA5066 | PA5066 | PA5066 | PA5066 | PA5066 | N/A | PA5066 | N/A | N/A | N/A |
| PA5067 | PA5067 | PA5067 | PA5067 | PA5067 | N/A | PA5067 | N/A | N/A | PA5067 | PA5067 | PA5067 | PA5067 | PA5067 | N/A | N/A | N/A | N/A | N/A |
| PA5068 | PA5068 | PA5068 | PA5068 | PA5068 | N/A | PA5068 | PA5068 | PA5068 | PA5068 | PA5068 | PA5068 | PA5068 | PA5068 | PA5068 | N/A | N/A | PA5068 | N/A |
| PA5069 | N/A | N/A | N/A | PA5069 | N/A | PA5069 | PA5069 | N/A | N/A | PA5069 | PA5069 | PA5069 | PA5069 | PA5069 | N/A | N/A | N/A | N/A |
| PA5070 | N/A | PA5070 | N/A | PA5070 | N/A | PA5070 | N/A | N/A | PA5070 | PA5070 | PA5070 | PA5070 | PA5070 | N/A | N/A | N/A | PA5070 | N/A |
| PA5071 | N/A | N/A | N/A | PA5071 | N/A | N/A | N/A | N/A | PA5071 | N/A | PA5071 | N/A | N/A | N/A | N/A | N/A | N/A | N/A |
| PA5072 | PA5072 | PA5072 | PA5072 | PA5072 | N/A | PA5072 | PA5072 | PA5072 | PA5072 | PA5072 | PA5072 | PA5072 | PA5072 | PA5072 | PA5072 | N/A | PA5072 | PA5072 |
| N/A | N/A | N/A | N/A | N/A | N/A | N/A | PA5073 | N/A | N/A | N/A | N/A | N/A | N/A | N/A | N/A | N/A | N/A | N/A |
| PA5074 | N/A | PA5074 | PA5074 | PA5074 | N/A | PA5074 | PA5074 | N/A | PA5074 | PA5074 | PA5074 | PA5074 | PA5074 | PA5074 | N/A | N/A | PA5074 | N/A |
| PA5075 | PA5075 | PA5075 | PA5075 | PA5075 | PA5075 | N/A | N/A | N/A | PA5075 | PA5075 | PA5075 | PA5075 | N/A | PA5075 | N/A | N/A | PA5075 | N/A |
| PA5076 | PA5076 | PA5076 | PA5076 | PA5076 | PA5076 | PA5076 | PA5076 | PA5076 | PA5076 | PA5076 | PA5076 | PA5076 | PA5076 | PA5076 | PA5076 | PA5076 | PA5076 | PA5076 |
| PA5077 | PA5077 | PA5077 | PA5077 | PA5077 | PA5077 | PA5077 | N/A | PA5077 | PA5077 | PA5077 | PA5077 | PA5077 | PA5077 | PA5077 | PA5077 | N/A | PA5077 | PA5077 |
| PA5078 | PA5078 | PA5078 | PA5078 | PA5078 | PA5078 | PA5078 | PA5078 | PA5078 | PA5078 | PA5078 | PA5078 | PA5078 | PA5078 | PA5078 | PA5078 | PA5078 | PA5078 | N/A |
| PA5079 | N/A | PA5079 | PA5079 | PA5079 | PA5079 | PA5079 | N/A | N/A | PA5079 | PA5079 | PA5079 | PA5079 | PA5079 | N/A | N/A | N/A | N/A | N/A |
| PA5080 | PA5080 | PA5080 | PA5080 | PA5080 | PA5080 | PA5080 | N/A | N/A | PA5080 | PA5080 | PA5080 | PA5080 | PA5080 | N/A | N/A | N/A | N/A | N/A |
| PA5081 | N/A | N/A | N/A | N/A | PA5081 | N/A | N/A | N/A | N/A | N/A | N/A | N/A | N/A | N/A | N/A | N/A | N/A | N/A |
| PA5082 | N/A | PA5082 | N/A | PA5082 | PA5082 | PA5082 | PA5082 | N/A | PA5082 | PA5082 | PA5082 | PA5082 | PA5082 | N/A | PA5082 | PA5082 | N/A | N/A |
| PA5083 | N/A | N/A | N/A | N/A | PA5083 | N/A | N/A | N/A | PA5083 | N/A | PA5083 | PA5083 | PA5083 | N/A | N/A | N/A | N/A | N/A |
| PA5084 | N/A | N/A | N/A | N/A | N/A | N/A | N/A | N/A | N/A | N/A | N/A | N/A | PA5084 | N/A | N/A | N/A | N/A | N/A |
| PA5085 | N/A | N/A | N/A | PA5085 | N/A | N/A | N/A | N/A | N/A | N/A | N/A | N/A | N/A | N/A | N/A | N/A | N/A | N/A |
| N/A | N/A | N/A | N/A | N/A | PA5090 | N/A | N/A | N/A | PA5090 | N/A | N/A | N/A | N/A | N/A | N/A | N/A | N/A | N/A |
| PA5091 | N/A | N/A | N/A | PA5091 | PA5091 | PA5091 | N/A | N/A | N/A | PA5091 | PA5091 | PA5091 | PA5091 | N/A | N/A | PA5091 | N/A | N/A |
| PA5092 | N/A | N/A | N/A | PA5092 | PA5092 | PA5092 | N/A | N/A | N/A | PA5092 | N/A | PA5092 | PA5092 | N/A | N/A | N/A | N/A | N/A |
| N/A | N/A | N/A | N/A | N/A | N/A | N/A | N/A | N/A | N/A | N/A | N/A | N/A | PA5093 | N/A | N/A | N/A | N/A | N/A |
| PA5094 | N/A | N/A | N/A | N/A | N/A | PA5094 | N/A | N/A | N/A | N/A | N/A | PA5094 | PA5094 | PA5094 | N/A | N/A | N/A | N/A |
| PA5095 | N/A | N/A | N/A | N/A | N/A | PA5095 | N/A | N/A | N/A | N/A | N/A | PA5095 | PA5095 | PA5095 | N/A | N/A | N/A | N/A |
| PA5096 | N/A | N/A | N/A | N/A | N/A | N/A | N/A | N/A | N/A | N/A | N/A | N/A | PA5096 | N/A | N/A | N/A | N/A | N/A |
| N/A | N/A | N/A | N/A | N/A | PA5097 | N/A | N/A | N/A | N/A | N/A | N/A | N/A | N/A | N/A | N/A | N/A | N/A | N/A |
| PA5098 | N/A | N/A | N/A | N/A | PA5098 | PA5098 | N/A | N/A | PA5098 | PA5098 | N/A | PA5098 | PA5098 | N/A | N/A | N/A | N/A | N/A |
| N/A | N/A | N/A | N/A | N/A | PA5099 | N/A | N/A | N/A | N/A | N/A | N/A | N/A | N/A | N/A | N/A | N/A | N/A | N/A |
| PA5100 | N/A | N/A | N/A | PA5100 | PA5100 | PA5100 | N/A | N/A | PA5100 | PA5100 | PA5100 | PA5100 | PA5100 | PA5100 | N/A | PA5100 | N/A | N/A |
| N/A | N/A | N/A | N/A | N/A | PA5102 | N/A | N/A | N/A | N/A | N/A | N/A | N/A | N/A | N/A | N/A | N/A | N/A | N/A |
| PA5103 | N/A | N/A | N/A | N/A | PA5103 | N/A | N/A | N/A | N/A | N/A | N/A | N/A | N/A | N/A | N/A | N/A | N/A | N/A |
| PA5104 | N/A | N/A | N/A | N/A | PA5104 | N/A | N/A | N/A | N/A | N/A | N/A | PA5104 | PA5104 | N/A | N/A | N/A | N/A | N/A |
| PA5105 | N/A | PA5105 | N/A | PA5105 | PA5105 | N/A | N/A | N/A | N/A | PA5105 | PA5105 | PA5105 | PA5105 | N/A | N/A | N/A | N/A | N/A |
| PA5106 | N/A | N/A | N/A | N/A | N/A | N/A | N/A | N/A | N/A | N/A | N/A | N/A | PA5106 | N/A | N/A | N/A | N/A | N/A |
| PA5107 | N/A | N/A | N/A | PA5107 | N/A | N/A | PA5107 | N/A | PA5107 | PA5107 | PA5107 | PA5107 | PA5107 | PA5107 | N/A | N/A | N/A | N/A |
| PA5108 | PA5108 | PA5108 | PA5108 | PA5108 | PA5108 | PA5108 | PA5108 | N/A | PA5108 | PA5108 | PA5108 | PA5108 | PA5108 | PA5108 | PA5108 | N/A | PA5108 | N/A |
| PA5109 | N/A | N/A | N/A | PA5109 | PA5109 | N/A | N/A | N/A | PA5109 | PA5109 | PA5109 | N/A | PA5109 | N/A | N/A | N/A | N/A | N/A |
| PA5110 | PA5110 | PA5110 | PA5110 | PA5110 | PA5110 | PA5110 | PA5110 | PA5110 | PA5110 | PA5110 | PA5110 | PA5110 | PA5110 | PA5110 | PA5110 | N/A | N/A | N/A |
| PA5111 | PA5111 | PA5111 | PA5111 | PA5111 | PA5111 | PA5111 | N/A | N/A | PA5111 | PA5111 | PA5111 | PA5111 | PA5111 | N/A | PA5111 | N/A | N/A | N/A |
| PA5112 | PA5112 | PA5112 | PA5112 | PA5112 | PA5112 | PA5112 | PA5112 | PA5112 | PA5112 | PA5112 | PA5112 | PA5112 | PA5112 | PA5112 | PA5112 | PA5112 | N/A | N/A |

| | | | | | | | | | | | | | | | | | | |
|---|---|---|---|---|---|---|---|---|---|---|---|---|---|---|---|---|---|---|
| PA5113 | PA5113 | PA5113 | N/A | PA5113 | N/A | PA5113 | PA5113 | N/A | PA5113 | PA5113 | PA5113 | N/A | N/A | PA5113 | PA5113 | N/A | PA5113 | N/A |
| PA5114 | N/A | N/A | PA5114 | N/A | PA5114 | N/A | N/A | N/A | PA5114 | PA5114 | PA5114 | PA5114 | N/A | PA5114 | N/A | N/A | PA5114 | N/A |
| PA5115 | N/A | N/A | N/A | N/A | PA5115 | N/A | N/A | N/A | N/A | PA5115 | PA5115 | PA5115 | PA5115 | N/A | N/A | N/A | N/A | N/A |
| N/A | N/A | N/A | N/A | N/A | PA5116 | N/A | N/A | N/A | N/A | N/A | N/A | N/A | N/A | N/A | N/A | N/A | N/A | N/A |
| PA5117 | PA5117 | PA5117 | PA5117 | PA5117 | PA5117 | PA5117 | PA5117 | PA5117 | PA5117 | PA5117 | PA5117 | PA5117 | PA5117 | PA5117 | PA5117 | N/A | N/A | N/A |
| PA5118 | PA5118 | PA5118 | PA5118 | PA5118 | PA5118 | N/A | PA5118 | N/A | PA5118 | PA5118 | PA5118 | PA5118 | PA5118 | PA5118 | N/A | N/A | N/A | N/A |
| PA5119 | PA5119 | PA5119 | PA5119 | PA5119 | PA5119 | PA5119 | PA5119 | PA5119 | PA5119 | PA5119 | PA5119 | PA5119 | PA5119 | PA5119 | PA5119 | PA5119 | N/A | N/A |
| PA5121 | N/A | N/A | N/A | PA5121 | N/A | N/A | N/A | PA5121 | PA5121 | N/A | PA5121 | PA5121 | PA5121 | PA5121 | N/A | N/A | N/A | N/A |
| PA5122 | N/A | N/A | N/A | PA5122 | PA5122 | N/A | N/A | N/A | PA5122 | PA5122 | PA5122 | N/A | PA5122 | N/A | PA5122 | N/A | N/A | N/A |
| N/A | N/A | N/A | N/A | PA5123 | N/A | N/A | N/A | N/A | N/A | PA5123 | N/A | N/A | N/A | N/A | N/A | N/A | N/A | N/A |
| PA5124 | N/A | N/A | N/A | PA5124 | PA5124 | N/A | N/A | N/A | N/A | N/A | N/A | N/A | N/A | N/A | N/A | N/A | N/A | N/A |
| PA5125 | N/A | N/A | N/A | PA5125 | PA5125 | N/A | N/A | PA5125 | N/A | N/A | N/A | PA5125 | PA5125 | N/A | N/A | N/A | N/A | N/A |
| PA5126 | N/A | N/A | N/A | N/A | N/A | N/A | N/A | N/A | N/A | N/A | N/A | N/A | N/A | PA5126 | N/A | N/A | N/A | N/A |
| PA5127 | N/A | N/A | N/A | PA5127 | PA5127 | N/A | N/A | N/A | PA5127 | PA5127 | PA5127 | PA5127 | PA5127 | N/A | N/A | N/A | N/A | N/A |
| PA5128 | PA5128 | PA5128 | PA5128 | PA5128 | PA5128 | PA5128 | PA5128 | PA5128 | PA5128 | PA5128 | PA5128 | PA5128 | PA5128 | PA5128 | PA5128 | PA5128 | N/A | N/A |
| N/A | N/A | N/A | N/A | N/A | N/A | N/A | N/A | N/A | PA5129 | N/A | PA5129 | PA5129 | PA5129 | N/A | N/A | N/A | N/A | N/A |
| PA5130 | PA5130 | PA5130 | PA5130 | PA5130 | PA5130 | PA5130 | PA5130 | N/A | PA5130 | PA5130 | PA5130 | PA5130 | PA5130 | PA5130 | PA5130 | N/A | PA5130 | N/A |
| PA5131 | PA5131 | PA5131 | PA5131 | PA5131 | PA5131 | PA5131 | PA5131 | N/A | PA5131 | PA5131 | PA5131 | PA5131 | PA5131 | N/A | PA5131 | PA5131 | N/A | N/A |
| N/A | N/A | N/A | N/A | N/A | N/A | N/A | N/A | N/A | N/A | N/A | N/A | N/A | N/A | PA5132 | N/A | N/A | N/A | N/A |
| PA5133 | N/A | N/A | N/A | PA5133 | PA5133 | N/A | N/A | N/A | N/A | N/A | N/A | N/A | N/A | PA5133 | N/A | N/A | PA5133 | PA5133 |
| PA5134 | PA5134 | PA5134 | PA5134 | PA5134 | PA5134 | PA5134 | PA5134 | PA5134 | PA5134 | PA5134 | PA5134 | PA5134 | PA5134 | PA5134 | PA5134 | N/A | PA5134 | PA5134 |
| PA5135 | N/A | PA5135 | N/A | PA5135 | N/A | N/A | PA5135 | PA5135 | N/A | N/A | N/A | PA5135 | N/A | PA5135 | N/A | N/A | PA5135 | PA5135 |
| PA5136 | PA5136 | N/A | PA5136 | PA5136 | PA5136 | N/A | PA5136 | PA5136 | PA5136 | N/A | N/A | PA5136 | N/A | PA5136 | PA5136 | N/A | PA5136 | PA5136 |
| PA5137 | N/A | N/A | N/A | N/A | PA5137 | PA5137 | N/A | N/A | N/A | PA5137 | PA5137 | PA5137 | PA5137 | N/A | N/A | N/A | N/A | N/A |
| PA5138 | PA5138 | PA5138 | PA5138 | PA5138 | PA5138 | PA5138 | PA5138 | N/A | PA5138 | PA5138 | PA5138 | PA5138 | PA5138 | N/A | PA5138 | N/A | N/A | N/A |
| PA5139 | PA5139 | PA5139 | PA5139 | PA5139 | PA5139 | N/A | N/A | N/A | PA5139 | PA5139 | PA5139 | PA5139 | PA5139 | N/A | PA5139 | N/A | N/A | N/A |
| PA5140 | PA5140 | PA5140 | PA5140 | PA5140 | PA5140 | PA5140 | PA5140 | N/A | PA5140 | PA5140 | PA5140 | PA5140 | PA5140 | N/A | PA5140 | N/A | N/A | N/A |
| PA5141 | PA5141 | PA5141 | PA5141 | PA5141 | PA5141 | PA5141 | PA5141 | PA5141 | PA5141 | PA5141 | PA5141 | PA5141 | PA5141 | N/A | PA5141 | N/A | N/A | N/A |
| PA5142 | PA5142 | PA5142 | PA5142 | PA5142 | PA5142 | PA5142 | N/A | N/A | PA5142 | PA5142 | PA5142 | PA5142 | PA5142 | N/A | N/A | N/A | N/A | N/A |
| PA5143 | PA5143 | PA5143 | PA5143 | PA5143 | PA5143 | PA5143 | PA5143 | N/A | PA5143 | PA5143 | PA5143 | PA5143 | PA5143 | PA5143 | PA5143 | N/A | N/A | N/A |
| PA5144 | N/A | N/A | N/A | N/A | N/A | N/A | N/A | N/A | N/A | N/A | N/A | N/A | N/A | N/A | N/A | N/A | N/A | N/A |
| N/A | N/A | N/A | N/A | N/A | PA5145 | N/A | N/A | N/A | N/A | N/A | N/A | N/A | N/A | N/A | N/A | N/A | N/A | N/A |
| PA5146 | PA5146 | PA5146 | PA5146 | PA5146 | PA5146 | PA5146 | PA5146 | PA5146 | PA5146 | PA5146 | PA5146 | PA5146 | PA5146 | PA5146 | PA5146 | N/A | PA5146 | N/A |
| PA5147 | N/A | PA5147 | PA5147 | PA5147 | N/A | N/A | N/A | PA5147 | N/A | N/A | N/A | N/A | N/A | N/A | N/A | N/A | N/A | N/A |
| PA5148 | N/A | PA5148 | PA5148 | PA5148 | N/A | PA5148 | N/A | N/A | PA5148 | PA5148 | PA5148 | PA5148 | PA5148 | N/A | N/A | N/A | N/A | PA5148 |
| PA5149 | N/A | N/A | N/A | N/A | N/A | N/A | N/A | N/A | N/A | N/A | N/A | N/A | PA5149 | N/A | N/A | N/A | N/A | N/A |
| PA5150 | N/A | N/A | N/A | N/A | N/A | N/A | N/A | N/A | N/A | N/A | N/A | N/A | PA5150 | N/A | N/A | N/A | N/A | N/A |
| PA5151 | N/A | N/A | N/A | N/A | N/A | N/A | PA5151 | N/A | N/A | N/A | N/A | N/A | N/A | PA5151 | N/A | N/A | N/A | N/A |
| PA5152 | N/A | N/A | PA5152 | PA5152 | PA5152 | PA5152 | PA5152 | PA5152 | PA5152 | PA5152 | PA5152 | PA5152 | PA5152 | PA5152 | N/A | N/A | PA5152 | N/A |
| PA5153 | PA5153 | PA5153 | N/A | PA5153 | PA5153 | PA5153 | PA5153 | N/A | PA5153 | PA5153 | PA5153 | PA5153 | PA5153 | N/A | PA5153 | PA5153 | N/A | PA5153 |
| PA5154 | N/A | N/A | N/A | PA5154 | N/A | PA5154 | N/A | N/A | PA5154 | N/A | PA5154 | PA5154 | PA5154 | PA5154 | N/A | N/A | PA5154 | N/A |
| PA5155 | N/A | N/A | N/A | PA5155 | N/A | PA5155 | N/A | N/A | N/A | N/A | PA5155 | PA5155 | N/A | PA5155 | N/A | N/A | PA5155 | N/A |
| N/A | N/A | N/A | N/A | PA5156 | N/A | N/A | N/A | PA5156 | N/A | N/A | PA5156 | PA5156 | PA5156 | N/A | N/A | N/A | N/A | N/A |
| PA5158 | N/A | N/A | N/A | PA5158 | N/A | N/A | N/A | N/A | N/A | N/A | N/A | N/A | N/A | N/A | N/A | N/A | N/A | N/A |
| N/A | N/A | N/A | N/A | PA5159 | N/A | N/A | N/A | N/A | N/A | N/A | N/A | N/A | N/A | N/A | N/A | N/A | N/A | N/A |
| N/A | N/A | N/A | N/A | N/A | PA5160 | N/A | N/A | N/A | N/A | N/A | N/A | N/A | N/A | N/A | N/A | N/A | N/A | N/A |
| PA5161 | PA5161 | PA5161 | PA5161 | PA5161 | PA5161 | PA5161 | PA5161 | PA5161 | PA5161 | PA5161 | PA5161 | PA5161 | PA5161 | N/A | PA5161 | PA5161 | N/A | N/A |
| PA5162 | PA5162 | PA5162 | PA5162 | PA5162 | PA5162 | PA5162 | N/A | PA5162 | PA5162 | PA5162 | PA5162 | PA5162 | PA5162 | N/A | N/A | N/A | N/A | N/A |
| PA5163 | PA5163 | PA5163 | PA5163 | PA5163 | PA5163 | PA5163 | PA5163 | N/A | PA5163 | PA5163 | PA5163 | PA5163 | PA5163 | N/A | PA5163 | N/A | N/A | N/A |
| PA5164 | PA5164 | PA5164 | PA5164 | PA5164 | PA5164 | PA5164 | PA5164 | N/A | PA5164 | PA5164 | PA5164 | PA5164 | PA5164 | N/A | PA5164 | PA5164 | N/A | N/A |
| PA5165 | N/A | N/A | N/A | PA5165 | PA5165 | N/A | N/A | N/A | N/A | N/A | N/A | N/A | N/A | PA5165 | N/A | N/A | N/A | N/A |
| PA5166 | PA5166 | N/A | PA5166 | PA5166 | N/A | PA5166 | N/A | PA5166 | PA5166 | PA5166 | N/A | PA5166 | PA5166 | N/A | N/A | N/A | N/A | N/A |
| PA5167 | PA5167 | PA5167 | PA5167 | PA5167 | PA5167 | PA5167 | PA5167 | N/A | PA5167 | PA5167 | PA5167 | PA5167 | PA5167 | N/A | PA5167 | PA5167 | N/A | N/A |
| PA5168 | PA5168 | PA5168 | N/A | PA5168 | N/A | PA5168 | N/A | N/A | PA5168 | N/A | N/A | PA5168 | PA5168 | PA5168 | N/A | N/A | N/A | N/A |
| PA5170 | N/A | N/A | N/A | PA5170 | PA5170 | PA5170 | N/A | N/A | N/A | PA5170 | PA5170 | PA5170 | PA5170 | PA5170 | N/A | N/A | PA5170 | N/A |
| PA5171 | PA5171 | PA5171 | PA5171 | PA5171 | PA5171 | PA5171 | PA5171 | PA5171 | PA5171 | PA5171 | PA5171 | PA5171 | PA5171 | PA5171 | PA5171 | PA5171 | PA5171 | N/A |
| PA5172 | PA5172 | PA5172 | PA5172 | PA5172 | PA5172 | PA5172 | PA5172 | PA5172 | PA5172 | PA5172 | PA5172 | PA5172 | PA5172 | PA5172 | PA5172 | PA5172 | PA5172 | N/A |
| PA5173 | PA5173 | PA5173 | PA5173 | PA5173 | PA5173 | PA5173 | PA5173 | PA5173 | PA5173 | PA5173 | PA5173 | PA5173 | PA5173 | PA5173 | PA5173 | PA5173 | PA5173 | N/A |
| PA5174 | PA5174 | PA5174 | PA5174 | PA5174 | PA5174 | PA5174 | N/A | PA5174 | PA5174 | PA5174 | PA5174 | PA5174 | PA5174 | PA5174 | PA5174 | PA5174 | N/A | N/A |
| PA5175 | PA5175 | N/A | PA5175 | PA5175 | PA5175 | N/A | N/A | N/A | PA5175 | N/A | PA5175 | N/A | N/A | N/A | N/A | N/A | N/A | N/A |
| PA5176 | PA5176 | PA5176 | PA5176 | PA5176 | PA5176 | PA5176 | N/A | N/A | PA5176 | PA5176 | PA5176 | PA5176 | PA5176 | N/A | PA5176 | N/A | N/A | N/A |
| PA5177 | N/A | N/A | N/A | PA5177 | PA5177 | N/A | N/A | N/A | PA5177 | N/A | N/A | N/A | N/A | N/A | N/A | N/A | N/A | N/A |
| PA5178 | PA5178 | PA5178 | PA5178 | PA5178 | PA5178 | PA5178 | PA5178 | PA5178 | PA5178 | PA5178 | PA5178 | PA5178 | PA5178 | PA5178 | PA5178 | PA5178 | PA5178 | PA5178 |
| PA5179 | N/A | N/A | N/A | N/A | N/A | N/A | N/A | N/A | N/A | PA5179 | N/A | N/A | N/A | N/A | N/A | N/A | N/A | N/A |
| PA5181 | N/A | N/A | N/A | N/A | N/A | N/A | N/A | N/A | PA5181 | N/A | N/A | N/A | PA5181 | N/A | N/A | N/A | N/A | N/A |
| PA5183 | N/A | N/A | N/A | PA5183 | PA5183 | N/A | N/A | N/A | N/A | N/A | N/A | N/A | N/A | N/A | PA5183 | N/A | N/A | N/A |
| N/A | N/A | N/A | N/A | PA5183.1 | N/A | N/A | N/A | N/A | PA5183.1 | N/A | PA5183.1 | PA5183.1 | PA5183.1 | N/A | N/A | N/A | PA5183.1 | N/A |
| PA5184 | PA5184 | PA5184 | N/A | PA5184 | PA5184 | PA5184 | N/A | N/A | PA5184 | PA5184 | PA5184 | PA5184 | PA5184 | N/A | PA5184 | N/A | PA5184 | PA5184 |
| PA5186 | N/A | N/A | N/A | N/A | N/A | PA5186 | N/A | N/A | N/A | PA5186 | N/A | N/A | PA5186 | N/A | N/A | N/A | N/A | N/A |
| PA5187 | PA5187 | PA5187 | PA5187 | PA5187 | N/A | PA5187 | N/A | N/A | PA5187 | PA5187 | PA5187 | PA5187 | PA5187 | PA5187 | N/A | N/A | N/A | N/A |
| PA5188 | N/A | N/A | N/A | PA5188 | N/A | PA5188 | N/A | N/A | N/A | N/A | N/A | N/A | PA5188 | N/A | N/A | N/A | N/A | N/A |
| PA5189 | N/A | N/A | N/A | N/A | PA5189 | N/A | N/A | N/A | N/A | N/A | N/A | N/A | N/A | N/A | N/A | N/A | N/A | N/A |
| PA5190 | PA5190 | PA5190 | PA5190 | PA5190 | PA5190 | N/A | PA5190 | N/A | PA5190 | PA5190 | PA5190 | PA5190 | N/A | N/A | N/A | N/A | N/A | N/A |
| N/A | N/A | N/A | N/A | N/A | PA5191 | N/A | N/A | N/A | N/A | N/A | N/A | N/A | N/A | N/A | N/A | N/A | N/A | N/A |
| PA5192 | PA5192 | PA5192 | PA5192 | PA5192 | PA5192 | PA5192 | PA5192 | PA5192 | PA5192 | PA5192 | PA5192 | PA5192 | PA5192 | PA5192 | PA5192 | PA5192 | N/A | N/A |
| PA5193 | PA5193 | PA5193 | PA5193 | PA5193 | PA5193 | N/A | PA5193 | N/A | PA5193 | PA5193 | PA5193 | PA5193 | PA5193 | N/A | PA5193 | N/A | N/A | N/A |
| PA5194 | PA5194 | PA5194 | PA5194 | PA5194 | N/A | PA5194 | PA5194 | PA5194 | PA5194 | PA5194 | PA5194 | PA5194 | PA5194 | PA5194 | N/A | N/A | PA5194 | N/A |
| PA5195 | N/A | N/A | N/A | PA5195 | PA5195 | N/A | N/A | N/A | PA5195 | PA5195 | PA5195 | PA5195 | PA5195 | PA5195 | PA5195 | N/A | N/A | N/A |
| PA5196 | N/A | N/A | N/A | N/A | PA5196 | PA5196 | N/A | N/A | PA5196 | PA5196 | PA5196 | PA5196 | N/A | PA5196 | N/A | N/A | N/A | N/A |
| PA5197 | N/A | N/A | N/A | PA5197 | PA5197 | N/A | N/A | PA5197 | PA5197 | PA5197 | PA5197 | PA5197 | PA5197 | PA5197 | N/A | N/A | N/A | N/A |
| PA5198 | PA5198 | PA5198 | PA5198 | PA5198 | PA5198 | PA5198 | PA5198 | N/A | PA5198 | PA5198 | PA5198 | PA5198 | PA5198 | PA5198 | PA5198 | PA5198 | N/A | N/A |
| PA5199 | N/A | PA5199 | N/A | PA5199 | PA5199 | N/A | N/A | PA5199 | PA5199 | PA5199 | PA5199 | PA5199 | PA5199 | PA5199 | N/A | N/A | PA5199 | N/A |
| PA5200 | PA5200 | PA5200 | PA5200 | PA5200 | PA5200 | PA5200 | PA5200 | N/A | PA5200 | PA5200 | PA5200 | PA5200 | PA5200 | PA5200 | PA5200 | N/A | PA5200 | N/A |
| PA5201 | PA5201 | PA5201 | PA5201 | PA5201 | PA5201 | PA5201 | PA5201 | PA5201 | PA5201 | PA5201 | PA5201 | PA5201 | PA5201 | PA5201 | PA5201 | N/A | N/A | N/A |
| PA5202 | N/A | N/A | PA5202 | PA5202 | PA5202 | N/A | N/A | N/A | N/A | N/A | PA5202 | N/A | N/A | N/A | N/A | N/A | N/A | N/A |
| PA5203 | N/A | PA5203 | PA5203 | PA5203 | N/A | PA5203 | N/A | PA5203 | PA5203 | PA5203 | PA5203 | PA5203 | PA5203 | N/A | PA5203 | N/A | N/A | N/A |
| PA5204 | PA5204 | PA5204 | PA5204 | PA5204 | N/A | N/A | N/A | PA5204 | PA5204 | PA5204 | PA5204 | PA5204 | PA5204 | PA5204 | PA5204 | N/A | N/A | N/A |
| PA5205 | N/A | N/A | N/A | N/A | PA5205 | PA5205 | N/A | N/A | N/A | N/A | N/A | N/A | N/A | PA5205 | N/A | N/A | N/A | N/A |
| PA5206 | N/A | N/A | N/A | PA5206 | PA5206 | N/A | N/A | N/A | N/A | PA5206 | N/A | N/A | N/A | N/A | N/A | N/A | N/A | N/A |
| PA5207 | N/A | N/A | N/A | N/A | N/A | N/A | N/A | N/A | N/A | N/A | N/A | N/A | N/A | N/A | N/A | N/A | N/A | N/A |
| PA5208 | N/A | N/A | N/A | PA5208 | PA5208 | PA5208 | N/A | N/A | PA5208 | PA5208 | PA5208 | PA5208 | PA5208 | PA5208 | N/A | N/A | PA5208 | N/A |
| PA5209 | PA5209 | PA5209 | PA5209 | PA5209 | PA5209 | PA5209 | N/A | PA5209 | PA5209 | PA5209 | PA5209 | PA5209 | PA5209 | PA5209 | PA5209 | N/A | N/A | N/A |
| PA5210 | PA5210 | PA5210 | N/A | PA5210 | PA5210 | N/A | N/A | PA5210 | PA5210 | PA5210 | PA5210 | PA5210 | PA5210 | N/A | N/A | N/A | N/A | N/A |
| PA5211 | N/A | N/A | N/A | N/A | N/A | N/A | N/A | N/A | N/A | N/A | N/A | N/A | N/A | PA5211 | N/A | N/A | PA5211 | N/A |
| N/A | N/A | N/A | N/A | N/A | PA5212 | N/A | N/A | N/A | N/A | N/A | N/A | N/A | N/A | N/A | N/A | N/A | N/A | N/A |
| PA5213 | N/A | N/A | N/A | PA5213 | PA5213 | N/A | N/A | N/A | N/A | PA5213 | N/A | PA5213 | PA5213 | N/A | N/A | N/A | N/A | N/A |
| PA5214 | PA5214 | PA5214 | PA5214 | PA5214 | PA5214 | PA5214 | N/A | N/A | PA5214 | PA5214 | PA5214 | PA5214 | PA5214 | N/A | PA5214 | PA5214 | N/A | N/A |
| PA5215 | PA5215 | PA5215 | PA5215 | PA5215 | PA5215 | PA5215 | PA5215 | PA5215 | PA5215 | PA5215 | PA5215 | PA5215 | PA5215 | N/A | PA5215 | PA5215 | N/A | N/A |
| PA5216 | N/A | N/A | N/A | PA5216 | PA5216 | N/A | N/A | N/A | N/A | N/A | N/A | N/A | N/A | PA5216 | N/A | N/A | N/A | N/A |
| PA5217 | PA5217 | PA5217 | PA5217 | PA5217 | PA5217 | PA5217 | N/A | N/A | PA5217 | PA5217 | PA5217 | PA5217 | PA5217 | PA5217 | PA5217 | PA5217 | N/A | N/A |
| PA5218 | N/A | N/A | N/A | N/A | N/A | N/A | N/A | N/A | N/A | N/A | N/A | N/A | PA5218 | N/A | N/A | N/A | N/A | N/A |
| N/A | N/A | N/A | N/A | N/A | PA5219 | N/A | N/A | N/A | N/A | N/A | N/A | N/A | N/A | N/A | N/A | N/A | N/A | N/A |
| PA5220 | N/A | N/A | N/A | N/A | PA5220 | PA5220 | PA5220 | PA5220 | PA5220 | PA5220 | PA5220 | PA5220 | PA5220 | N/A | N/A | N/A | N/A | N/A |
| PA5221 | PA5221 | PA5221 | PA5221 | PA5221 | N/A | N/A | PA5221 | PA5221 | PA5221 | PA5221 | PA5221 | PA5221 | PA5221 | N/A | N/A | N/A | N/A | N/A |
| N/A | N/A | N/A | N/A | N/A | PA5222 | N/A | N/A | N/A | N/A | N/A | N/A | N/A | N/A | N/A | N/A | N/A | N/A | N/A |
| PA5223 | PA5223 | PA5223 | PA5223 | PA5223 | N/A | N/A | PA5223 | PA5223 | PA5223 | PA5223 | PA5223 | PA5223 | PA5223 | PA5223 | N/A | N/A | N/A | N/A |
| PA5224 | PA5224 | PA5224 | PA5224 | PA5224 | PA5224 | PA5224 | PA5224 | PA5224 | PA5224 | PA5224 | PA5224 | PA5224 | PA5224 | N/A | PA5224 | PA5224 | N/A | N/A |
| PA5225 | PA5225 | PA5225 | N/A | PA5225 | N/A | N/A | PA5225 | PA5225 | PA5225 | PA5225 | PA5225 | PA5225 | PA5225 | N/A | PA5225 | N/A | N/A | N/A |
| N/A | N/A | N/A | N/A | PA5226 | N/A | PA5226 | N/A | N/A | N/A | N/A | PA5226 | N/A | PA5226 | N/A | N/A | N/A | N/A | N/A |

| | | | | | | | | | | | | | | | | | | |
|---|---|---|---|---|---|---|---|---|---|---|---|---|---|---|---|---|---|---|
| PA5227 | PA5227 | PA5227 | PA5227 | PA5227 | N/A | PA5227 | PA5227 | N/A | PA5227 | PA5227 | PA5227 | PA5227 | PA5227 | N/A | N/A | N/A | N/A | N/A |
| PA5228 | N/A | N/A | N/A | N/A | PA5228 | N/A | N/A | N/A | N/A | PA5228 | N/A | N/A | N/A | N/A | N/A | N/A | N/A | N/A |
| PA5229 | PA5229 | PA5229 | PA5229 | PA5229 | PA5229 | N/A | PA5229 | N/A | PA5229 | PA5229 | PA5229 | PA5229 | PA5229 | N/A | PA5229 | N/A | N/A | N/A |
| PA5230 | N/A | N/A | N/A | PA5230 | N/A | N/A | N/A | N/A | N/A | PA5230 | N/A | PA5230 | PA5230 | PA5230 | N/A | N/A | PA5230 | N/A |
| PA5231 | N/A | N/A | N/A | PA5231 | PA5231 | PA5231 | N/A | N/A | PA5231 | PA5231 | PA5231 | PA5231 | PA5231 | PA5231 | N/A | N/A | PA5231 | N/A |
| PA5232 | PA5232 | N/A | PA5232 | PA5232 | PA5232 | PA5232 | PA5232 | PA5232 | PA5232 | PA5232 | PA5232 | PA5232 | PA5232 | PA5232 | PA5232 | N/A | PA5232 | PA5232 |
| PA5233 | PA5233 | N/A | N/A | PA5233 | N/A | N/A | N/A | N/A | PA5233 | PA5233 | PA5233 | PA5233 | PA5233 | PA5233 | N/A | N/A | PA5233 | N/A |
| PA5234 | N/A | N/A | N/A | PA5234 | N/A | N/A | N/A | N/A | PA5234 | PA5234 | PA5234 | PA5234 | PA5234 | N/A | N/A | N/A | N/A | N/A |
| PA5235 | N/A | PA5235 | N/A | PA5235 | N/A | PA5235 | N/A | PA5235 | PA5235 | PA5235 | PA5235 | PA5235 | PA5235 | PA5235 | N/A | N/A | PA5235 | N/A |
| PA5236 | N/A | N/A | N/A | PA5236 | PA5236 | N/A | N/A | N/A | N/A | N/A | PA5236 | N/A | N/A | N/A | N/A | N/A | N/A | N/A |
| PA5237 | PA5237 | PA5237 | N/A | PA5237 | PA5237 | PA5237 | PA5237 | PA5237 | PA5237 | PA5237 | PA5237 | PA5237 | PA5237 | PA5237 | N/A | N/A | N/A | N/A |
| N/A | N/A | N/A | N/A | N/A | PA5238 | N/A | N/A | N/A | N/A | N/A | N/A | N/A | N/A | N/A | N/A | N/A | N/A | N/A |
| PA5239 | PA5239 | PA5239 | PA5239 | PA5239 | PA5239 | PA5239 | PA5239 | PA5239 | PA5239 | PA5239 | PA5239 | PA5239 | PA5239 | PA5239 | PA5239 | N/A | PA5239 | PA5239 |
| PA5240 | PA5240 | PA5240 | PA5240 | PA5240 | PA5240 | PA5240 | PA5240 | N/A | PA5240 | PA5240 | PA5240 | PA5240 | PA5240 | PA5240 | PA5240 | PA5240 | PA5240 | N/A |
| PA5241 | PA5241 | PA5241 | PA5241 | PA5241 | N/A | N/A | N/A | PA5241 | PA5241 | PA5241 | PA5241 | PA5241 | PA5241 | N/A | N/A | N/A | N/A | N/A |
| PA5242 | PA5242 | PA5242 | PA5242 | PA5242 | PA5242 | PA5242 | N/A | PA5242 | PA5242 | PA5242 | PA5242 | PA5242 | PA5242 | PA5242 | N/A | N/A | PA5242 | N/A |
| PA5243 | PA5243 | PA5243 | PA5243 | PA5243 | PA5243 | PA5243 | PA5243 | PA5243 | PA5243 | PA5243 | PA5243 | PA5243 | PA5243 | PA5243 | PA5243 | PA5243 | N/A | N/A |
| N/A | N/A | N/A | N/A | N/A | PA5244 | N/A | N/A | N/A | N/A | N/A | N/A | N/A | N/A | PA5244 | N/A | N/A | N/A | N/A |
| PA5245 | PA5245 | PA5245 | PA5245 | PA5245 | PA5245 | PA5245 | N/A | N/A | PA5245 | PA5245 | PA5245 | PA5245 | PA5245 | PA5245 | PA5245 | N/A | N/A | N/A |
| PA5247 | N/A | N/A | N/A | N/A | N/A | N/A | N/A | N/A | N/A | N/A | N/A | N/A | N/A | PA5247 | N/A | N/A | N/A | N/A |
| PA5248 | N/A | N/A | N/A | PA5248 | N/A | N/A | N/A | N/A | PA5248 | N/A | N/A | PA5248 | N/A | PA5248 | N/A | N/A | PA5248 | N/A |
| N/A | N/A | N/A | N/A | PA5249 | N/A | N/A | N/A | N/A | N/A | N/A | N/A | N/A | N/A | N/A | N/A | N/A | N/A | N/A |
| PA5250 | N/A | N/A | N/A | N/A | N/A | PA5250 | N/A | N/A | PA5250 | N/A | PA5250 | PA5250 | PA5250 | PA5250 | N/A | N/A | N/A | N/A |
| PA5251 | N/A | N/A | N/A | N/A | N/A | PA5251 | N/A | N/A | PA5251 | PA5251 | PA5251 | PA5251 | N/A | PA5251 | N/A | N/A | PA5251 | N/A |
| PA5252 | PA5252 | PA5252 | PA5252 | PA5252 | PA5252 | N/A | N/A | PA5252 | PA5252 | PA5252 | PA5252 | PA5252 | PA5252 | N/A | PA5252 | N/A | PA5252 | N/A |
| PA5253 | PA5253 | PA5253 | PA5253 | PA5253 | PA5253 | PA5253 | PA5253 | PA5253 | PA5253 | PA5253 | PA5253 | PA5253 | PA5253 | PA5253 | PA5253 | PA5253 | PA5253 | PA5253 |
| PA5254 | N/A | N/A | N/A | N/A | N/A | N/A | N/A | N/A | N/A | N/A | N/A | N/A | N/A | N/A | N/A | N/A | N/A | N/A |
| PA5255 | PA5255 | PA5255 | PA5255 | PA5255 | N/A | PA5255 | N/A | N/A | PA5255 | PA5255 | PA5255 | PA5255 | PA5255 | N/A | N/A | N/A | N/A | N/A |
| N/A | N/A | N/A | N/A | N/A | PA5256 | N/A | N/A | N/A | N/A | N/A | N/A | N/A | N/A | N/A | N/A | N/A | N/A | N/A |
| PA5257 | PA5257 | PA5257 | PA5257 | PA5257 | PA5257 | PA5257 | PA5257 | PA5257 | PA5257 | PA5257 | PA5257 | PA5257 | PA5257 | PA5257 | PA5257 | N/A | PA5257 | PA5257 |
| PA5258 | PA5258 | PA5258 | PA5258 | PA5258 | PA5258 | PA5258 | PA5258 | PA5258 | PA5258 | PA5258 | PA5258 | PA5258 | PA5258 | PA5258 | PA5258 | N/A | PA5258 | PA5258 |
| PA5259 | N/A | N/A | N/A | PA5259 | PA5259 | N/A | N/A | N/A | PA5259 | N/A | PA5259 | N/A | N/A | N/A | N/A | N/A | N/A | N/A |
| PA5260 | PA5260 | PA5260 | PA5260 | PA5260 | PA5260 | PA5260 | N/A | N/A | PA5260 | PA5260 | PA5260 | PA5260 | PA5260 | N/A | PA5260 | PA5260 | N/A | N/A |
| PA5261 | PA5261 | PA5261 | PA5261 | PA5261 | PA5261 | PA5261 | N/A | N/A | PA5261 | PA5261 | PA5261 | PA5261 | PA5261 | PA5261 | N/A | N/A | N/A | N/A |
| PA5262 | N/A | N/A | N/A | N/A | PA5262 | N/A | N/A | N/A | N/A | N/A | N/A | N/A | N/A | PA5262 | N/A | N/A | N/A | N/A |
| PA5263 | PA5263 | PA5263 | PA5263 | PA5263 | PA5263 | PA5263 | PA5263 | PA5263 | PA5263 | PA5263 | PA5263 | PA5263 | PA5263 | PA5263 | PA5263 | PA5263 | N/A | N/A |
| N/A | N/A | N/A | N/A | N/A | PA5266 | N/A | N/A | N/A | PA5266 | N/A | N/A | N/A | N/A | N/A | N/A | N/A | N/A | N/A |
| PA5267 | N/A | N/A | N/A | PA5267 | N/A | PA5267 | N/A | N/A | PA5267 | PA5267 | N/A | PA5267 | PA5267 | N/A | N/A | N/A | N/A | N/A |
| PA5268 | N/A | N/A | N/A | N/A | N/A | N/A | N/A | N/A | N/A | N/A | PA5268 | PA5268 | PA5268 | PA5268 | N/A | N/A | PA5268 | N/A |
| PA5269 | N/A | PA5269 | PA5269 | PA5269 | N/A | N/A | N/A | N/A | PA5269 | PA5269 | PA5269 | N/A | N/A | N/A | N/A | N/A | N/A | N/A |
| PA5270 | N/A | N/A | N/A | N/A | PA5270 | N/A | N/A | N/A | N/A | N/A | PA5270 | N/A | N/A | PA5270 | N/A | N/A | N/A | N/A |
| N/A | N/A | N/A | N/A | N/A | N/A | N/A | N/A | N/A | N/A | N/A | N/A | N/A | PA5271 | N/A | N/A | N/A | N/A | N/A |
| PA5272 | N/A | N/A | N/A | PA5272 | N/A | N/A | N/A | PA5272 | N/A | N/A | N/A | PA5272 | PA5272 | N/A | N/A | N/A | N/A | N/A |
| N/A | N/A | N/A | N/A | N/A | PA5273 | N/A | N/A | N/A | N/A | N/A | N/A | N/A | N/A | N/A | N/A | N/A | N/A | N/A |
| PA5274 | PA5274 | PA5274 | PA5274 | PA5274 | N/A | PA5274 | PA5274 | N/A | PA5274 | PA5274 | PA5274 | PA5274 | PA5274 | N/A | PA5274 | PA5274 | N/A | N/A |
| PA5275 | PA5275 | PA5275 | PA5275 | PA5275 | N/A | N/A | N/A | N/A | PA5275 | N/A | PA5275 | PA5275 | PA5275 | N/A | PA5275 | N/A | N/A | N/A |
| PA5276 | N/A | N/A | N/A | N/A | PA5276 | PA5276 | N/A | N/A | PA5276 | PA5276 | PA5276 | PA5276 | PA5276 | N/A | N/A | N/A | N/A | N/A |
| PA5277 | PA5277 | PA5277 | PA5277 | PA5277 | PA5277 | PA5277 | PA5277 | PA5277 | PA5277 | PA5277 | PA5277 | PA5277 | PA5277 | N/A | PA5277 | N/A | N/A | N/A |
| PA5278 | PA5278 | PA5278 | PA5278 | PA5278 | PA5278 | PA5278 | N/A | PA5278 | PA5278 | PA5278 | PA5278 | PA5278 | PA5278 | N/A | PA5278 | N/A | PA5278 | N/A |
| PA5279 | PA5279 | PA5279 | PA5279 | PA5279 | N/A | N/A | PA5279 | PA5279 | PA5279 | PA5279 | PA5279 | PA5279 | PA5279 | PA5279 | N/A | N/A | PA5279 | N/A |
| PA5280 | N/A | N/A | N/A | N/A | PA5280 | N/A | N/A | N/A | N/A | N/A | N/A | N/A | N/A | N/A | N/A | N/A | N/A | N/A |
| PA5281 | N/A | N/A | N/A | PA5281 | PA5281 | N/A | N/A | N/A | PA5281 | N/A | N/A | N/A | N/A | N/A | N/A | N/A | N/A | N/A |
| N/A | N/A | N/A | N/A | N/A | N/A | N/A | N/A | PA5283 | N/A | N/A | N/A | N/A | PA5283 | N/A | N/A | N/A | N/A | N/A |
| N/A | N/A | N/A | N/A | N/A | PA5284 | N/A | N/A | N/A | N/A | N/A | N/A | N/A | N/A | N/A | N/A | N/A | N/A | N/A |
| PA5285 | PA5285 | PA5285 | N/A | PA5285 | PA5285 | PA5285 | N/A | N/A | PA5285 | PA5285 | PA5285 | PA5285 | PA5285 | N/A | N/A | N/A | N/A | N/A |
| PA5286 | PA5286 | PA5286 | PA5286 | N/A | PA5286 | N/A | N/A | N/A | N/A | PA5286 | PA5286 | PA5286 | PA5286 | N/A | N/A | N/A | N/A | N/A |
| PA5287 | N/A | N/A | N/A | N/A | PA5287 | PA5287 | N/A | N/A | N/A | N/A | N/A | N/A | N/A | PA5287 | N/A | N/A | PA5287 | N/A |
| PA5288 | PA5288 | PA5288 | PA5288 | PA5288 | PA5288 | PA5288 | PA5288 | N/A | PA5288 | PA5288 | PA5288 | PA5288 | PA5288 | PA5288 | PA5288 | N/A | PA5288 | PA5288 |
| PA5289 | PA5289 | PA5289 | PA5289 | PA5289 | N/A | PA5289 | N/A | N/A | PA5289 | PA5289 | PA5289 | PA5289 | PA5289 | N/A | N/A | N/A | N/A | N/A |
| N/A | N/A | N/A | N/A | N/A | N/A | N/A | N/A | PA5290 | N/A | N/A | N/A | N/A | N/A | N/A | N/A | N/A | N/A | N/A |
| PA5291 | N/A | PA5291 | PA5291 | PA5291 | N/A | PA5291 | N/A | PA5291 | PA5291 | PA5291 | PA5291 | PA5291 | PA5291 | PA5291 | N/A | N/A | PA5291 | N/A |
| PA5295 | N/A | N/A | N/A | PA5295 | N/A | N/A | N/A | PA5295 | N/A | N/A | N/A | N/A | N/A | N/A | N/A | N/A | N/A | N/A |
| PA5296 | PA5296 | N/A | PA5296 | PA5296 | N/A | N/A | N/A | PA5296 | N/A | N/A | PA5296 | N/A | PA5296 | N/A | N/A | N/A | N/A | N/A |
| N/A | N/A | N/A | N/A | N/A | PA5297 | N/A | N/A | N/A | N/A | PA5297 | N/A | N/A | PA5297 | N/A | N/A | N/A | N/A | N/A |
| PA5298 | PA5298 | PA5298 | PA5298 | PA5298 | PA5298 | PA5298 | N/A | N/A | PA5298 | PA5298 | PA5298 | PA5298 | PA5298 | N/A | PA5298 | N/A | PA5298 | N/A |
| PA5299 | N/A | N/A | N/A | N/A | PA5299 | N/A | N/A | N/A | N/A | N/A | N/A | PA5299 | PA5299 | N/A | N/A | N/A | N/A | N/A |
| PA5300 | PA5300 | PA5300 | PA5300 | PA5300 | N/A | PA5300 | PA5300 | PA5300 | PA5300 | PA5300 | PA5300 | PA5300 | PA5300 | PA5300 | PA5300 | N/A | PA5300 | PA5300 |
| PA5301 | PA5301 | PA5301 | PA5301 | PA5301 | N/A | PA5301 | N/A | N/A | PA5301 | PA5301 | PA5301 | PA5301 | PA5301 | PA5301 | PA5301 | N/A | PA5301 | N/A |
| PA5302 | PA5302 | PA5302 | PA5302 | PA5302 | PA5302 | N/A | N/A | N/A | PA5302 | PA5302 | PA5302 | PA5302 | PA5302 | N/A | PA5302 | N/A | N/A | N/A |
| PA5303 | N/A | N/A | N/A | PA5303 | PA5303 | PA5303 | N/A | N/A | PA5303 | PA5303 | PA5303 | PA5303 | PA5303 | N/A | N/A | N/A | N/A | N/A |
| PA5304 | PA5304 | PA5304 | PA5304 | PA5304 | PA5304 | PA5304 | PA5304 | PA5304 | PA5304 | PA5304 | PA5304 | PA5304 | PA5304 | PA5304 | PA5304 | N/A | PA5304 | PA5304 |
| N/A | N/A | N/A | N/A | PA5305 | N/A | PA5305 | N/A | N/A | PA5305 | PA5305 | PA5305 | N/A | PA5305 | PA5305 | N/A | N/A | N/A | N/A |
| PA5306 | N/A | N/A | N/A | PA5306 | N/A | PA5306 | N/A | N/A | PA5306 | N/A | PA5306 | PA5306 | PA5306 | PA5306 | N/A | N/A | N/A | N/A |
| PA5307 | N/A | N/A | N/A | N/A | PA5307 | N/A | N/A | N/A | N/A | PA5307 | N/A | N/A | N/A | PA5307 | N/A | N/A | PA5307 | N/A |
| PA5308 | PA5308 | PA5308 | N/A | PA5308 | PA5308 | PA5308 | N/A | PA5308 | PA5308 | PA5308 | PA5308 | PA5308 | PA5308 | PA5308 | N/A | N/A | PA5308 | N/A |
| PA5309 | PA5309 | PA5309 | PA5309 | PA5309 | PA5309 | N/A | N/A | PA5309 | N/A | N/A | N/A | N/A | PA5309 | PA5309 | N/A | N/A | PA5309 | N/A |
| PA5310 | N/A | N/A | N/A | PA5310 | N/A | N/A | N/A | N/A | N/A | N/A | N/A | N/A | N/A | N/A | N/A | N/A | N/A | N/A |
| N/A | N/A | N/A | N/A | N/A | PA5311 | N/A | N/A | N/A | N/A | N/A | N/A | N/A | N/A | N/A | N/A | N/A | N/A | N/A |
| PA5312 | PA5312 | PA5312 | PA5312 | PA5312 | PA5312 | PA5312 | PA5312 | N/A | PA5312 | PA5312 | PA5312 | PA5312 | PA5312 | PA5312 | PA5312 | PA5312 | N/A | N/A |
| PA5313 | PA5313 | PA5313 | PA5313 | PA5313 | PA5313 | PA5313 | N/A | N/A | PA5313 | PA5313 | PA5313 | PA5313 | PA5313 | N/A | N/A | N/A | N/A | N/A |
| PA5314 | N/A | N/A | N/A | PA5314 | PA5314 | PA5314 | N/A | N/A | PA5314 | N/A | PA5314 | PA5314 | PA5314 | N/A | N/A | N/A | N/A | N/A |
| PA5315 | PA5315 | PA5315 | N/A | PA5315 | PA5315 | PA5315 | PA5315 | PA5315 | PA5315 | PA5315 | PA5315 | PA5315 | PA5315 | PA5315 | N/A | PA5315 | PA5315 | PA5315 |
| PA5316 | PA5316 | PA5316 | PA5316 | PA5316 | PA5316 | PA5316 | PA5316 | PA5316 | PA5316 | PA5316 | PA5316 | PA5316 | PA5316 | PA5316 | PA5316 | PA5316 | PA5316 | PA5316 |
| PA5317 | PA5317 | PA5317 | N/A | PA5317 | PA5317 | PA5317 | N/A | N/A | PA5317 | PA5317 | PA5317 | PA5317 | PA5317 | PA5317 | PA5317 | N/A | N/A | N/A |
| PA5318 | N/A | N/A | N/A | N/A | N/A | N/A | N/A | N/A | N/A | N/A | N/A | N/A | PA5318 | PA5318 | N/A | N/A | N/A | N/A |
| N/A | N/A | N/A | N/A | N/A | PA5319 | N/A | N/A | N/A | N/A | N/A | N/A | N/A | N/A | N/A | N/A | N/A | N/A | N/A |
| PA5320 | PA5320 | PA5320 | PA5320 | PA5320 | PA5320 | N/A | PA5320 | PA5320 | PA5320 | PA5320 | PA5320 | PA5320 | PA5320 | PA5320 | N/A | N/A | N/A | N/A |
| PA5321 | PA5321 | PA5321 | PA5321 | PA5321 | PA5321 | PA5321 | N/A | N/A | PA5321 | PA5321 | PA5321 | PA5321 | PA5321 | N/A | PA5321 | N/A | N/A | N/A |
| PA5322 | PA5322 | PA5322 | PA5322 | PA5322 | PA5322 | PA5322 | PA5322 | PA5322 | PA5322 | PA5322 | PA5322 | PA5322 | PA5322 | PA5322 | PA5322 | N/A | PA5322 | N/A |
| PA5323 | PA5323 | PA5323 | PA5323 | PA5323 | PA5323 | PA5323 | PA5323 | PA5323 | PA5323 | PA5323 | PA5323 | PA5323 | PA5323 | PA5323 | PA5323 | N/A | PA5323 | PA5323 |
| N/A | N/A | N/A | N/A | N/A | PA5328 | N/A | N/A | N/A | N/A | N/A | N/A | N/A | N/A | N/A | N/A | N/A | N/A | N/A |
| PA5329 | N/A | N/A | N/A | N/A | N/A | PA5329 | N/A | N/A | N/A | PA5329 | N/A | PA5329 | PA5329 | N/A | N/A | N/A | N/A | N/A |
| PA5330 | PA5330 | PA5330 | PA5330 | PA5330 | PA5330 | N/A | PA5330 | N/A | PA5330 | PA5330 | PA5330 | PA5330 | PA5330 | PA5330 | PA5330 | N/A | PA5330 | PA5330 |
| PA5331 | PA5331 | PA5331 | PA5331 | PA5331 | PA5331 | PA5331 | PA5331 | PA5331 | PA5331 | PA5331 | PA5331 | PA5331 | PA5331 | N/A | PA5331 | PA5331 | N/A | N/A |
| PA5332 | PA5332 | PA5332 | PA5332 | PA5332 | PA5332 | PA5332 | PA5332 | PA5332 | PA5332 | PA5332 | PA5332 | PA5332 | PA5332 | PA5332 | PA5332 | PA5332 | N/A | N/A |
| PA5333 | N/A | N/A | N/A | PA5333 | PA5333 | PA5333 | N/A | PA5333 | PA5333 | PA5333 | PA5333 | PA5333 | PA5333 | PA5333 | N/A | N/A | PA5333 | N/A |
| PA5334 | PA5334 | PA5334 | PA5334 | PA5334 | PA5334 | PA5334 | PA5334 | PA5334 | PA5334 | PA5334 | PA5334 | PA5334 | PA5334 | PA5334 | PA5334 | PA5334 | N/A | N/A |
| PA5335 | PA5335 | PA5335 | PA5335 | PA5335 | N/A | PA5335 | PA5335 | N/A | PA5335 | PA5335 | PA5335 | PA5335 | PA5335 | PA5335 | PA5335 | PA5335 | N/A | N/A |
| PA5336 | PA5336 | PA5336 | PA5336 | PA5336 | N/A | N/A | N/A | N/A | PA5336 | N/A | PA5336 | PA5336 | PA5336 | N/A | PA5336 | N/A | N/A | N/A |
| PA5337 | PA5337 | PA5337 | PA5337 | PA5337 | PA5337 | PA5337 | PA5337 | N/A | PA5337 | PA5337 | PA5337 | PA5337 | PA5337 | PA5337 | PA5337 | N/A | N/A | N/A |
| PA5338 | PA5338 | PA5338 | PA5338 | PA5338 | PA5338 | PA5338 | N/A | PA5338 | PA5338 | PA5338 | PA5338 | PA5338 | PA5338 | PA5338 | N/A | N/A | PA5338 | N/A |
| PA5339 | PA5339 | PA5339 | PA5339 | PA5339 | PA5339 | PA5339 | PA5339 | N/A | PA5339 | PA5339 | PA5339 | PA5339 | PA5339 | PA5339 | PA5339 | PA5339 | PA5339 | N/A |
| PA5340 | PA5340 | PA5340 | N/A | PA5340 | N/A | N/A | PA5340 | N/A | PA5340 | PA5340 | PA5340 | PA5340 | N/A | PA5340 | PA5340 | PA5340 | N/A | N/A |
| N/A | N/A | N/A | N/A | N/A | PA5342 | N/A | N/A | N/A | N/A | N/A | N/A | N/A | N/A | N/A | N/A | N/A | N/A | N/A |
| PA5343 | PA5343 | PA5343 | PA5343 | PA5343 | PA5343 | N/A | PA5343 | PA5343 | PA5343 | PA5343 | PA5343 | PA5343 | PA5343 | N/A | PA5343 | N/A | N/A | N/A |
| PA5344 | PA5344 | PA5344 | PA5344 | PA5344 | PA5344 | PA5344 | PA5344 | PA5344 | PA5344 | PA5344 | PA5344 | PA5344 | PA5344 | PA5344 | N/A | N/A | PA5344 | PA5344 |
| PA5345 | PA5345 | PA5345 | N/A | PA5345 | PA5345 | N/A | N/A | PA5345 | PA5345 | PA5345 | PA5345 | PA5345 | PA5345 | PA5345 | N/A | N/A | N/A | N/A |
| PA5346 | PA5346 | N/A | PA5346 | PA5346 | PA5346 | N/A | N/A | PA5346 | PA5346 | PA5346 | PA5346 | N/A | PA5346 | N/A | PA5346 | N/A | N/A | N/A |
| PA5347 | N/A | N/A | N/A | PA5347 | PA5347 | N/A | PA5347 | N/A | PA5347 | N/A | PA5347 | PA5347 | N/A | N/A | N/A | N/A | PA5347 | PA5347 |

| | | | | | | | | | | | | | | | | | | |
|---|---|---|---|---|---|---|---|---|---|---|---|---|---|---|---|---|---|---|
| PA5348 | PA5348 | PA5348 | PA5348 | PA5348 | PA5348 | PA5348 | PA5348 | N/A | PA5348 | PA5348 | PA5348 | PA5348 | PA5348 | PA5348 | PA5348 | N/A | PA5348 | N/A |
| PA5349 | PA5349 | PA5349 | PA5349 | PA5349 | PA5349 | N/A | PA5349 | PA5349 | PA5349 | PA5349 | PA5349 | PA5349 | PA5349 | N/A | N/A | N/A | N/A | N/A |
| N/A | N/A | N/A | PA5351 | N/A | N/A | N/A | N/A | N/A | N/A | N/A | N/A | N/A | N/A | N/A | N/A | N/A | N/A | N/A |
| PA5353 | N/A | N/A | N/A | N/A | N/A | N/A | N/A | N/A | N/A | N/A | N/A | PA5353 | PA5353 | N/A | N/A | N/A | N/A | N/A |
| PA5354 | N/A | N/A | N/A | N/A | N/A | N/A | N/A | N/A | N/A | N/A | N/A | N/A | N/A | N/A | N/A | N/A | N/A | N/A |
| PA5356 | N/A | N/A | N/A | N/A | PA5356 | N/A | N/A | N/A | N/A | N/A | N/A | N/A | N/A | N/A | N/A | N/A | N/A | N/A |
| PA5357 | N/A | N/A | N/A | PA5357 | PA5357 | N/A | N/A | N/A | PA5357 | PA5357 | PA5357 | PA5357 | PA5357 | N/A | PA5357 | N/A | N/A | N/A |
| PA5358 | N/A | N/A | N/A | PA5358 | PA5358 | N/A | N/A | N/A | N/A | PA5358 | PA5358 | PA5358 | PA5358 | PA5358 | N/A | N/A | PA5358 | N/A |
| PA5359 | N/A | N/A | N/A | N/A | PA5359 | PA5359 | N/A | N/A | PA5359 | PA5359 | PA5359 | PA5359 | PA5359 | N/A | N/A | N/A | N/A | N/A |
| PA5360 | N/A | N/A | N/A | PA5360 | N/A | N/A | PA5360 | N/A | PA5360 | PA5360 | PA5360 | PA5360 | PA5360 | N/A | N/A | N/A | PA5360 | N/A |
| PA5361 | N/A | N/A | N/A | PA5361 | N/A | N/A | N/A | PA5361 | N/A | N/A | N/A | N/A | N/A | PA5361 | N/A | N/A | PA5361 | N/A |
| PA5362 | PA5362 | PA5362 | PA5362 | PA5362 | N/A | N/A | N/A | PA5362 | PA5362 | PA5362 | PA5362 | PA5362 | PA5362 | PA5362 | PA5362 | N/A | N/A | N/A |
| PA5363 | N/A | N/A | N/A | N/A | PA5363 | N/A | N/A | N/A | N/A | N/A | N/A | N/A | N/A | N/A | N/A | N/A | PA5363 | N/A |
| PA5364 | PA5364 | PA5364 | PA5364 | PA5364 | PA5364 | N/A | N/A | PA5364 | PA5364 | PA5364 | PA5364 | PA5364 | PA5364 | PA5364 | N/A | N/A | N/A | PA5364 |
| PA5365 | N/A | PA5365 | PA5365 | PA5365 | PA5365 | PA5365 | PA5365 | N/A | PA5365 | PA5365 | PA5365 | PA5365 | PA5365 | PA5365 | N/A | N/A | PA5365 | N/A |
| PA5366 | PA5366 | PA5366 | PA5366 | PA5366 | PA5366 | PA5366 | PA5366 | PA5366 | PA5366 | PA5366 | PA5366 | PA5366 | PA5366 | PA5366 | N/A | N/A | PA5366 | N/A |
| PA5367 | PA5367 | PA5367 | N/A | PA5367 | N/A | N/A | N/A | N/A | PA5367 | N/A | N/A | PA5367 | N/A | PA5367 | PA5367 | N/A | PA5367 | N/A |
| PA5368 | PA5368 | N/A | N/A | PA5368 | PA5368 | N/A | N/A | N/A | PA5368 | PA5368 | N/A | PA5368 | N/A | PA5368 | PA5368 | N/A | PA5368 | N/A |
| PA5369 | N/A | PA5369 | N/A | PA5369 | PA5369 | PA5369 | N/A | N/A | PA5369 | PA5369 | PA5369 | PA5369 | PA5369 | PA5369 | N/A | N/A | N/A | N/A |
| PA5370 | N/A | N/A | N/A | N/A | PA5370 | N/A | N/A | N/A | N/A | N/A | N/A | PA5370 | PA5370 | PA5370 | N/A | N/A | PA5370 | N/A |
| PA5371 | N/A | N/A | N/A | N/A | PA5371 | PA5371 | N/A | N/A | PA5371 | PA5371 | PA5371 | PA5371 | PA5371 | N/A | N/A | N/A | N/A | N/A |
| PA5372 | PA5372 | PA5372 | PA5372 | PA5372 | PA5372 | N/A | N/A | PA5372 | PA5372 | PA5372 | N/A | PA5372 | PA5372 | PA5372 | N/A | N/A | PA5372 | PA5372 |
| PA5373 | PA5373 | PA5373 | PA5373 | PA5373 | PA5373 | PA5373 | N/A | PA5373 | PA5373 | PA5373 | PA5373 | PA5373 | PA5373 | PA5373 | N/A | N/A | N/A | N/A |
| PA5374 | N/A | N/A | N/A | PA5374 | N/A | PA5374 | PA5374 | N/A | PA5374 | PA5374 | PA5374 | PA5374 | PA5374 | PA5374 | N/A | N/A | PA5374 | N/A |
| PA5375 | N/A | PA5375 | N/A | N/A | N/A | N/A | N/A | N/A | N/A | PA5375 | N/A | PA5375 | N/A | PA5375 | N/A | N/A | PA5375 | N/A |
| PA5376 | N/A | PA5376 | N/A | PA5376 | N/A | N/A | N/A | N/A | N/A | PA5376 | PA5376 | PA5376 | PA5376 | PA5376 | PA5376 | N/A | PA5376 | N/A |
| PA5377 | N/A | N/A | N/A | N/A | PA5377 | N/A | N/A | N/A | PA5377 | N/A | N/A | PA5377 | N/A | PA5377 | N/A | N/A | PA5377 | N/A |
| PA5378 | N/A | N/A | N/A | PA5378 | PA5378 | N/A | N/A | N/A | N/A | PA5378 | PA5378 | PA5378 | PA5378 | N/A | PA5378 | PA5378 | N/A | N/A |
| N/A | N/A | N/A | N/A | N/A | N/A | N/A | N/A | N/A | N/A | N/A | N/A | N/A | PA5379 | N/A | N/A | N/A | N/A | N/A |
| N/A | N/A | N/A | N/A | PA5381 | N/A | N/A | N/A | N/A | N/A | N/A | N/A | N/A | PA5381 | N/A | N/A | N/A | N/A | N/A |
| N/A | N/A | N/A | N/A | PA5387 | N/A | N/A | N/A | N/A | N/A | N/A | N/A | N/A | N/A | N/A | PA5387 | N/A | N/A | N/A |
| N/A | N/A | N/A | N/A | N/A | N/A | N/A | N/A | N/A | N/A | N/A | N/A | PA5389 | N/A | N/A | N/A | N/A | N/A | N/A |
| PA5390 | N/A | N/A | N/A | N/A | N/A | N/A | N/A | N/A | N/A | N/A | N/A | N/A | N/A | N/A | N/A | N/A | N/A | N/A |
| PA5394 | N/A | N/A | N/A | N/A | N/A | N/A | N/A | N/A | N/A | N/A | N/A | N/A | N/A | PA5394 | N/A | N/A | N/A | N/A |
| PA5395 | N/A | N/A | N/A | N/A | N/A | N/A | N/A | N/A | N/A | N/A | N/A | N/A | PA5395 | N/A | N/A | N/A | N/A | N/A |
| N/A | N/A | N/A | N/A | N/A | N/A | PA5396 | N/A | N/A | N/A | PA5396 | N/A | PA5396 | PA5396 | N/A | N/A | N/A | N/A | N/A |
| N/A | N/A | N/A | N/A | N/A | N/A | PA5397 | N/A | N/A | PA5397 | N/A | N/A | PA5397 | N/A | N/A | N/A | N/A | N/A | N/A |
| N/A | N/A | N/A | N/A | N/A | N/A | N/A | N/A | N/A | N/A | N/A | N/A | PA5398 | N/A | N/A | N/A | N/A | N/A | N/A |
| PA5399 | N/A | N/A | N/A | N/A | N/A | N/A | N/A | N/A | N/A | N/A | N/A | N/A | N/A | PA5399 | N/A | N/A | PA5399 | N/A |
| N/A | N/A | N/A | N/A | PA5403 | N/A | N/A | N/A | N/A | N/A | N/A | PA5403 | N/A | PA5403 | N/A | N/A | N/A | N/A | N/A |
| N/A | PA5406 | PA5406 | PA5406 | PA5406 | N/A | N/A | N/A | N/A | PA5406 | N/A | PA5406 | PA5406 | N/A | N/A | N/A | N/A | N/A | N/A |
| N/A | PA5407 | PA5407 | PA5407 | PA5407 | N/A | N/A | N/A | N/A | PA5407 | N/A | PA5407 | N/A | N/A | N/A | N/A | N/A | N/A | N/A |
| N/A | N/A | N/A | N/A | N/A | N/A | N/A | N/A | N/A | N/A | N/A | N/A | N/A | N/A | N/A | N/A | N/A | PA5409 | N/A |
| N/A | N/A | N/A | N/A | N/A | N/A | N/A | N/A | N/A | N/A | N/A | N/A | PA5410 | PA5410 | N/A | N/A | N/A | N/A | N/A |
| N/A | N/A | N/A | N/A | N/A | N/A | N/A | N/A | N/A | N/A | N/A | N/A | PA5411 | N/A | N/A | N/A | N/A | N/A | N/A |
| PA5412 | N/A | N/A | N/A | PA5412 | PA5412 | N/A | N/A | PA5412 | N/A | N/A | PA5412 | N/A | N/A | PA5412 | N/A | N/A | PA5412 | N/A |
| PA5413 | PA5413 | PA5413 | PA5413 | PA5413 | PA5413 | PA5413 | PA5413 | PA5413 | PA5413 | PA5413 | PA5413 | PA5413 | PA5413 | N/A | PA5413 | PA5413 | N/A | N/A |
| PA5414 | PA5414 | PA5414 | N/A | PA5414 | N/A | PA5414 | PA5414 | PA5414 | PA5414 | PA5414 | PA5414 | PA5414 | PA5414 | PA5414 | PA5414 | PA5414 | PA5414 | N/A |
| PA5415 | PA5415 | PA5415 | PA5415 | PA5415 | PA5415 | PA5415 | PA5415 | PA5415 | PA5415 | PA5415 | PA5415 | PA5415 | PA5415 | N/A | PA5415 | N/A | N/A | N/A |
| N/A | N/A | N/A | N/A | N/A | N/A | N/A | N/A | N/A | N/A | N/A | N/A | PA5417 | N/A | N/A | N/A | N/A | N/A | N/A |
| N/A | N/A | N/A | N/A | N/A | N/A | N/A | N/A | N/A | N/A | N/A | N/A | PA5418 | N/A | N/A | N/A | N/A | N/A | N/A |
| PA5421 | N/A | N/A | N/A | N/A | PA5421 | PA5421 | N/A | N/A | N/A | N/A | N/A | PA5421 | PA5421 | N/A | N/A | N/A | N/A | N/A |
| PA5422 | PA5422 | PA5422 | N/A | PA5422 | PA5422 | N/A | N/A | N/A | PA5422 | PA5422 | PA5422 | PA5422 | PA5422 | N/A | N/A | N/A | N/A | N/A |
| PA5423 | N/A | N/A | N/A | PA5423 | N/A | N/A | N/A | N/A | PA5423 | PA5423 | PA5423 | PA5423 | PA5423 | N/A | PA5423 | N/A | N/A | N/A |
| N/A | N/A | N/A | N/A | N/A | PA5424 | N/A | N/A | N/A | N/A | N/A | N/A | N/A | N/A | N/A | N/A | N/A | N/A | N/A |
| PA5425 | PA5425 | PA5425 | PA5425 | PA5425 | PA5425 | PA5425 | PA5425 | PA5425 | PA5425 | PA5425 | PA5425 | PA5425 | PA5425 | N/A | PA5425 | PA5425 | N/A | N/A |
| PA5426 | PA5426 | PA5426 | PA5426 | PA5426 | PA5426 | PA5426 | PA5426 | N/A | PA5426 | PA5426 | PA5426 | PA5426 | PA5426 | PA5426 | PA5426 | PA5426 | N/A | N/A |
| PA5427 | PA5427 | PA5427 | N/A | PA5427 | PA5427 | PA5427 | N/A | N/A | PA5427 | PA5427 | PA5427 | PA5427 | PA5427 | PA5427 | N/A | PA5427 | PA5427 | N/A |
| PA5428 | PA5428 | PA5428 | PA5428 | PA5428 | N/A | N/A | N/A | N/A | N/A | N/A | PA5428 | N/A | N/A | N/A | N/A | N/A | N/A | N/A |
| PA5429 | PA5429 | PA5429 | PA5429 | PA5429 | PA5429 | PA5429 | PA5429 | PA5429 | PA5429 | PA5429 | PA5429 | PA5429 | PA5429 | PA5429 | PA5429 | PA5429 | N/A | N/A |
| PA5430 | N/A | N/A | N/A | PA5430 | N/A | N/A | N/A | N/A | N/A | PA5430 | N/A | PA5430 | N/A | PA5430 | N/A | N/A | N/A | N/A |
| PA5432 | N/A | N/A | N/A | N/A | N/A | N/A | N/A | N/A | N/A | N/A | N/A | N/A | N/A | N/A | N/A | N/A | N/A | N/A |
| PA5433 | N/A | N/A | N/A | N/A | N/A | N/A | N/A | N/A | PA5433 | N/A | PA5433 | PA5433 | N/A | N/A | N/A | N/A | N/A | N/A |
| PA5434 | N/A | N/A | N/A | N/A | PA5434 | N/A | N/A | N/A | N/A | N/A | N/A | N/A | N/A | N/A | N/A | N/A | N/A | N/A |
| PA5435 | PA5435 | PA5435 | PA5435 | PA5435 | PA5435 | PA5435 | N/A | PA5435 | PA5435 | PA5435 | PA5435 | PA5435 | PA5435 | PA5435 | PA5435 | PA5435 | N/A | N/A |
| PA5436 | PA5436 | PA5436 | PA5436 | PA5436 | PA5436 | PA5436 | N/A | PA5436 | PA5436 | PA5436 | PA5436 | PA5436 | PA5436 | PA5436 | PA5436 | N/A | N/A | N/A |
| PA5437 | N/A | N/A | N/A | PA5437 | PA5437 | N/A | N/A | N/A | N/A | N/A | N/A | N/A | N/A | N/A | N/A | N/A | N/A | N/A |
| PA5438 | PA5438 | PA5438 | PA5438 | PA5438 | PA5438 | PA5438 | N/A | PA5438 | PA5438 | PA5438 | PA5438 | PA5438 | PA5438 | N/A | N/A | N/A | PA5438 | PA5438 |
| PA5439 | N/A | N/A | N/A | PA5439 | PA5439 | N/A | N/A | N/A | N/A | N/A | N/A | N/A | N/A | N/A | N/A | N/A | N/A | N/A |
| PA5440 | N/A | N/A | N/A | PA5440 | PA5440 | N/A | N/A | PA5440 | N/A | PA5440 | PA5440 | PA5440 | PA5440 | N/A | N/A | N/A | N/A | N/A |
| PA5441 | PA5441 | PA5441 | PA5441 | PA5441 | PA5441 | PA5441 | N/A | PA5441 | PA5441 | PA5441 | PA5441 | PA5441 | PA5441 | PA5441 | PA5441 | N/A | PA5441 | PA5441 |
| N/A | N/A | N/A | N/A | N/A | PA5442 | N/A | N/A | N/A | N/A | N/A | N/A | N/A | N/A | N/A | N/A | N/A | N/A | N/A |
| PA5443 | PA5443 | PA5443 | PA5443 | PA5443 | PA5443 | PA5443 | PA5443 | PA5443 | PA5443 | PA5443 | PA5443 | PA5443 | PA5443 | PA5443 | PA5443 | N/A | N/A | N/A |
| N/A | N/A | N/A | N/A | N/A | PA5444 | N/A | N/A | N/A | N/A | N/A | N/A | N/A | N/A | N/A | N/A | N/A | N/A | N/A |
| N/A | PA5445 | PA5445 | PA5445 | PA5445 | PA5445 | N/A | N/A | PA5445 | PA5445 | PA5445 | N/A | N/A | N/A | N/A | PA5445 | N/A | N/A | N/A |
| PA5446 | N/A | N/A | N/A | PA5446 | N/A | PA5446 | N/A | N/A | PA5446 | N/A | PA5446 | PA5446 | PA5446 | PA5446 | N/A | N/A | PA5446 | PA5446 |
| PA5447 | N/A | N/A | N/A | N/A | N/A | N/A | N/A | N/A | N/A | N/A | N/A | N/A | N/A | PA5447 | N/A | N/A | PA5447 | N/A |
| PA5448 | N/A | N/A | N/A | N/A | N/A | N/A | N/A | N/A | N/A | N/A | N/A | N/A | N/A | N/A | N/A | N/A | N/A | N/A |
| PA5449 | N/A | N/A | N/A | PA5449 | N/A | N/A | N/A | N/A | PA5449 | N/A | PA5449 | PA5449 | PA5449 | PA5449 | N/A | N/A | N/A | N/A |
| PA5450 | N/A | N/A | N/A | PA5450 | N/A | N/A | N/A | N/A | N/A | N/A | N/A | N/A | N/A | PA5450 | N/A | N/A | N/A | N/A |
| PA5451 | N/A | N/A | N/A | PA5451 | N/A | N/A | N/A | N/A | N/A | N/A | N/A | N/A | N/A | N/A | N/A | N/A | N/A | N/A |
| N/A | N/A | N/A | N/A | N/A | N/A | N/A | N/A | PA5452 | N/A | N/A | N/A | PA5452 | PA5452 | N/A | N/A | N/A | N/A | N/A |
| PA5453 | PA5453 | N/A | N/A | PA5453 | N/A | PA5453 | N/A | N/A | PA5453 | N/A | PA5453 | PA5453 | PA5453 | N/A | N/A | N/A | PA5453 | N/A |
| PA5454 | N/A | N/A | N/A | PA5454 | N/A | N/A | N/A | N/A | N/A | N/A | PA5454 | PA5454 | PA5454 | N/A | N/A | N/A | N/A | N/A |
| PA5455 | N/A | N/A | N/A | PA5455 | N/A | N/A | N/A | N/A | PA5455 | N/A | PA5455 | PA5455 | PA5455 | PA5455 | PA5455 | N/A | N/A | N/A |
| PA5456 | N/A | N/A | N/A | PA5456 | N/A | N/A | N/A | N/A | PA5456 | N/A | PA5456 | PA5456 | N/A | PA5456 | PA5456 | N/A | N/A | N/A |
| PA5457 | N/A | N/A | N/A | PA5457 | N/A | N/A | PA5457 | N/A | N/A | N/A | PA5457 | PA5457 | PA5457 | PA5457 | N/A | N/A | N/A | N/A |
| PA5459 | N/A | PA5459 | N/A | PA5459 | N/A | N/A | N/A | N/A | N/A | PA5459 | PA5459 | PA5459 | PA5459 | N/A | N/A | N/A | N/A | N/A |
| PA5464 | N/A | N/A | N/A | PA5464 | N/A | N/A | N/A | PA5464 | PA5464 | PA5464 | PA5464 | PA5464 | PA5464 | PA5464 | N/A | N/A | N/A | N/A |
| PA5465 | N/A | N/A | N/A | N/A | N/A | N/A | N/A | N/A | N/A | N/A | N/A | PA5465 | N/A | N/A | N/A | N/A | N/A | N/A |
| PA5466 | N/A | N/A | N/A | PA5466 | N/A | N/A | N/A | N/A | N/A | N/A | N/A | N/A | N/A | PA5466 | N/A | N/A | N/A | N/A |
| N/A | N/A | N/A | N/A | PA5468 | N/A | N/A | N/A | N/A | N/A | N/A | N/A | N/A | N/A | N/A | N/A | N/A | N/A | N/A |
| PA5469 | N/A | N/A | N/A | N/A | N/A | N/A | N/A | N/A | N/A | N/A | N/A | N/A | N/A | N/A | N/A | N/A | N/A | N/A |
| PA5470 | N/A | PA5470 | N/A | N/A | N/A | N/A | N/A | N/A | N/A | N/A | N/A | N/A | N/A | N/A | N/A | N/A | N/A | N/A |
| PA5471 | N/A | N/A | N/A | PA5471 | N/A | N/A | PA5471 | PA5471 | N/A | N/A | PA5471 | PA5471 | PA5471 | N/A | N/A | N/A | N/A | N/A |
| N/A | PA5472 | PA5472 | PA5472 | PA5472 | N/A | N/A | PA5472 | N/A | PA5472 | PA5472 | PA5472 | N/A | N/A | N/A | PA5472 | N/A | N/A | N/A |
| PA5473 | N/A | N/A | N/A | N/A | N/A | N/A | N/A | N/A | N/A | N/A | N/A | PA5473 | N/A | PA5473 | N/A | N/A | N/A | N/A |
| PA5474 | N/A | N/A | N/A | PA5474 | PA5474 | N/A | PA5474 | PA5474 | PA5474 | PA5474 | N/A | PA5474 | PA5474 | PA5474 | N/A | N/A | N/A | N/A |
| PA5475 | N/A | PA5475 | PA5475 | PA5475 | PA5475 | PA5475 | N/A | N/A | PA5475 | PA5475 | PA5475 | PA5475 | PA5475 | N/A | N/A | N/A | PA5475 | PA5475 |
| PA5477 | N/A | N/A | N/A | N/A | N/A | N/A | N/A | N/A | N/A | PA5477 | N/A | N/A | N/A | PA5477 | N/A | N/A | N/A | N/A |
| PA5478 | N/A | N/A | N/A | PA5478 | N/A | PA5478 | N/A | PA5478 | PA5478 | PA5478 | PA5478 | PA5478 | PA5478 | PA5478 | N/A | N/A | N/A | N/A |
| PA5479 | PA5479 | PA5479 | PA5479 | PA5479 | N/A | PA5479 | N/A | PA5479 | PA5479 | PA5479 | PA5479 | PA5479 | N/A | PA5479 | N/A | N/A | PA5479 | N/A |
| PA5481 | N/A | N/A | N/A | N/A | N/A | N/A | N/A | N/A | PA5481 | PA5481 | PA5481 | PA5481 | PA5481 | N/A | N/A | N/A | N/A | N/A |
| N/A | N/A | N/A | N/A | N/A | PA5482 | N/A | N/A | N/A | N/A | N/A | N/A | N/A | N/A | N/A | N/A | N/A | N/A | N/A |
| PA5483 | PA5483 | PA5483 | PA5483 | PA5483 | PA5483 | PA5483 | N/A | PA5483 | PA5483 | PA5483 | PA5483 | PA5483 | PA5483 | N/A | PA5483 | N/A | N/A | N/A |
| PA5484 | N/A | N/A | N/A | PA5484 | N/A | N/A | N/A | N/A | PA5484 | N/A | PA5484 | PA5484 | PA5484 | PA5484 | N/A | N/A | N/A | N/A |
| PA5485 | PA5485 | PA5485 | PA5485 | PA5485 | N/A | N/A | PA5485 | PA5485 | PA5485 | PA5485 | PA5485 | PA5485 | PA5485 | PA5485 | PA5485 | N/A | N/A | N/A |
| PA5487 | N/A | N/A | N/A | PA5487 | N/A | N/A | N/A | PA5487 | PA5487 | PA5487 | PA5487 | PA5487 | PA5487 | N/A | N/A | N/A | N/A | N/A |
| PA5488 | N/A | N/A | N/A | N/A | PA5488 | N/A | N/A | N/A | N/A | PA5488 | PA5488 | N/A | N/A | N/A | N/A | N/A | N/A | N/A |

| | | | | | | | | | | | | | | | | | | |
|---|---|---|---|---|---|---|---|---|---|---|---|---|---|---|---|---|---|---|
| PA5489 | PA5489 | PA5489 | PA5489 | PA5489 | PA5489 | PA5489 | PA5489 | N/A | PA5489 | PA5489 | PA5489 | PA5489 | PA5489 | N/A | PA5489 | PA5489 | N/A | N/A |
| PA5490 | PA5490 | PA5490 | PA5490 | PA5490 | N/A | PA5490 | PA5490 | PA5490 | PA5490 | PA5490 | PA5490 | PA5490 | PA5490 | PA5490 | PA5490 | PA5490 | PA5490 | PA5490 |
| N/A | N/A | N/A | N/A | N/A | PA5491 | N/A | N/A | N/A | N/A | N/A | PA5491 | N/A | N/A | N/A | N/A | N/A | N/A | N/A |
| PA5492 | N/A | PA5492 | PA5492 | PA5492 | N/A | PA5492 | PA5492 | N/A | PA5492 | PA5492 | PA5492 | PA5492 | PA5492 | PA5492 | N/A | N/A | N/A | N/A |
| PA5493 | PA5493 | PA5493 | PA5493 | PA5493 | PA5493 | PA5493 | N/A | PA5493 | PA5493 | PA5493 | PA5493 | PA5493 | PA5493 | PA5493 | PA5493 | N/A | PA5493 | N/A |
| PA5494 | N/A | N/A | N/A | PA5494 | PA5494 | PA5494 | N/A | N/A | PA5494 | PA5494 | PA5494 | PA5494 | PA5494 | N/A | N/A | N/A | N/A | N/A |
| PA5495 | PA5495 | PA5495 | PA5495 | PA5495 | PA5495 | N/A | PA5495 | PA5495 | PA5495 | PA5495 | PA5495 | PA5495 | PA5495 | N/A | PA5495 | N/A | N/A | N/A |
| PA5496 | N/A | N/A | N/A | N/A | PA5496 | PA5496 | PA5496 | N/A | PA5496 | PA5496 | PA5496 | PA5496 | PA5496 | PA5496 | N/A | N/A | PA5496 | N/A |
| PA5497 | PA5497 | N/A | N/A | PA5497 | PA5497 | PA5497 | N/A | N/A | PA5497 | PA5497 | PA5497 | PA5497 | PA5497 | PA5497 | N/A | N/A | N/A | N/A |
| PA5498 | PA5498 | N/A | N/A | PA5498 | PA5498 | N/A | N/A | N/A | PA5498 | PA5498 | N/A | N/A | N/A | N/A | PA5498 | N/A | N/A | N/A |
| PA5499 | PA5499 | N/A | N/A | PA5499 | N/A | N/A | N/A | N/A | PA5499 | PA5499 | PA5499 | PA5499 | N/A | N/A | N/A | N/A | N/A | N/A |
| PA5500 | N/A | N/A | N/A | PA5500 | N/A | N/A | N/A | N/A | N/A | N/A | N/A | N/A | N/A | PA5500 | N/A | N/A | N/A | N/A |
| PA5502 | PA5502 | PA5502 | N/A | PA5502 | N/A | N/A | PA5502 | N/A | N/A | PA5502 | PA5502 | PA5502 | N/A | N/A | PA5502 | N/A | N/A | N/A |
| PA5503 | PA5503 | PA5503 | PA5503 | PA5503 | N/A | N/A | PA5503 | N/A | PA5503 | PA5503 | N/A | PA5503 | PA5503 | PA5503 | PA5503 | N/A | PA5503 | N/A |
| PA5504 | N/A | PA5504 | N/A | PA5504 | PA5504 | N/A | N/A | N/A | PA5504 | N/A | N/A | N/A | N/A | PA5504 | N/A | N/A | PA5504 | N/A |
| PA5505 | PA5505 | PA5505 | PA5505 | PA5505 | PA5505 | PA5505 | PA5505 | N/A | PA5505 | PA5505 | PA5505 | PA5505 | PA5505 | N/A | PA5505 | PA5505 | N/A | N/A |
| PA5506 | N/A | N/A | N/A | PA5506 | PA5506 | N/A | N/A | PA5506 | PA5506 | PA5506 | PA5506 | PA5506 | PA5506 | PA5506 | N/A | N/A | N/A | N/A |
| PA5507 | PA5507 | PA5507 | PA5507 | PA5507 | PA5507 | PA5507 | PA5507 | N/A | PA5507 | PA5507 | PA5507 | PA5507 | PA5507 | PA5507 | PA5507 | PA5507 | PA5507 | N/A |
| PA5508 | PA5508 | PA5508 | N/A | PA5508 | PA5508 | N/A | PA5508 | PA5508 | PA5508 | N/A | PA5508 | PA5508 | PA5508 | PA5508 | N/A | N/A | N/A | N/A |
| PA5509 | N/A | N/A | N/A | PA5509 | PA5509 | N/A | N/A | N/A | PA5509 | PA5509 | PA5509 | PA5509 | PA5509 | N/A | PA5509 | N/A | N/A | N/A |
| PA5510 | N/A | N/A | N/A | PA5510 | N/A | N/A | N/A | PA5510 | N/A | N/A | PA5510 | PA5510 | N/A | PA5510 | N/A | N/A | N/A | N/A |
| PA5511 | N/A | N/A | N/A | PA5511 | N/A | N/A | N/A | PA5511 | PA5511 | PA5511 | PA5511 | PA5511 | PA5511 | N/A | N/A | N/A | N/A | N/A |
| PA5512 | N/A | N/A | N/A | N/A | N/A | N/A | N/A | N/A | N/A | N/A | N/A | N/A | N/A | PA5512 | N/A | N/A | N/A | N/A |
| PA5513 | N/A | N/A | N/A | N/A | N/A | N/A | N/A | N/A | N/A | N/A | N/A | N/A | N/A | N/A | N/A | N/A | N/A | N/A |
| PA5514 | N/A | N/A | N/A | N/A | N/A | N/A | N/A | N/A | N/A | N/A | N/A | N/A | N/A | N/A | N/A | N/A | N/A | N/A |
| PA5515 | N/A | N/A | N/A | PA5515 | PA5515 | N/A | N/A | N/A | N/A | PA5515 | N/A | PA5515 | N/A | PA5515 | N/A | N/A | N/A | N/A |
| PA5516 | N/A | PA5516 | N/A | PA5516 | PA5516 | N/A | N/A | N/A | PA5516 | PA5516 | PA5516 | PA5516 | PA5516 | N/A | N/A | N/A | N/A | N/A |
| PA5518 | N/A | N/A | N/A | PA5518 | N/A | N/A | N/A | PA5518 | N/A | N/A | N/A | N/A | N/A | PA5518 | N/A | N/A | N/A | N/A |
| PA5519 | N/A | PA5519 | N/A | N/A | N/A | N/A | N/A | N/A | PA5519 | PA5519 | PA5519 | PA5519 | PA5519 | N/A | N/A | N/A | N/A | N/A |
| PA5520 | N/A | N/A | N/A | N/A | PA5520 | N/A | N/A | N/A | N/A | PA5520 | N/A | N/A | N/A | N/A | N/A | N/A | N/A | N/A |
| PA5521 | PA5521 | PA5521 | PA5521 | PA5521 | PA5521 | PA5521 | N/A | N/A | PA5521 | PA5521 | PA5521 | PA5521 | PA5521 | N/A | N/A | N/A | N/A | N/A |
| PA5522 | N/A | N/A | N/A | PA5522 | PA5522 | N/A | PA5522 | PA5522 | N/A | PA5522 | N/A | N/A | PA5522 | N/A | N/A | N/A | N/A | N/A |
| PA5523 | PA5523 | PA5523 | PA5523 | PA5523 | PA5523 | PA5523 | N/A | PA5523 | N/A | PA5523 | PA5523 | PA5523 | PA5523 | N/A | N/A | N/A | N/A | N/A |
| PA5524 | N/A | N/A | N/A | PA5524 | N/A | N/A | N/A | N/A | N/A | PA5524 | N/A | N/A | N/A | N/A | N/A | N/A | N/A | N/A |
| PA5525 | N/A | N/A | N/A | PA5525 | N/A | N/A | N/A | N/A | N/A | N/A | N/A | N/A | N/A | N/A | N/A | N/A | N/A | N/A |
| PA5526 | N/A | N/A | N/A | N/A | N/A | N/A | N/A | N/A | N/A | N/A | N/A | PA5526 | PA5526 | N/A | N/A | N/A | N/A | N/A |
| PA5527 | N/A | PA5527 | N/A | N/A | PA5527 | N/A | N/A | N/A | N/A | N/A | N/A | PA5527 | N/A | N/A | N/A | N/A | N/A | N/A |
| PA5528 | PA5528 | PA5528 | PA5528 | PA5528 | PA5528 | PA5528 | PA5528 | PA5528 | PA5528 | PA5528 | PA5528 | PA5528 | PA5528 | PA5528 | PA5528 | PA5528 | PA5528 | PA5528 |
| N/A | N/A | N/A | N/A | N/A | N/A | N/A | N/A | N/A | N/A | N/A | N/A | N/A | N/A | PA5529 | N/A | N/A | N/A | N/A |
| PA5530 | PA5530 | N/A | N/A | PA5530 | N/A | N/A | N/A | PA5530 | N/A | N/A | N/A | PA5530 | N/A | PA5530 | PA5530 | N/A | PA5530 | N/A |
| PA5531 | N/A | N/A | PA5531 | PA5531 | N/A | N/A | N/A | N/A | N/A | N/A | N/A | N/A | PA5531 | PA5531 | PA5531 | N/A | PA5531 | N/A |
| PA5533 | N/A | N/A | N/A | N/A | N/A | N/A | N/A | N/A | N/A | N/A | N/A | N/A | N/A | N/A | N/A | N/A | N/A | N/A |
| N/A | N/A | N/A | N/A | N/A | N/A | N/A | N/A | N/A | N/A | N/A | PA5534 | N/A | N/A | N/A | N/A | N/A | N/A | N/A |
| N/A | N/A | N/A | N/A | PA5535 | N/A | N/A | N/A | N/A | N/A | N/A | N/A | N/A | N/A | N/A | N/A | N/A | N/A | N/A |
| PA5542 | N/A | N/A | N/A | N/A | N/A | N/A | N/A | N/A | N/A | N/A | PA5542 | PA5542 | PA5542 | N/A | N/A | N/A | N/A | N/A |
| N/A | N/A | N/A | N/A | N/A | PA5544 | N/A | N/A | N/A | N/A | N/A | N/A | N/A | N/A | N/A | N/A | N/A | N/A | N/A |
| PA5545 | N/A | N/A | N/A | PA5545 | PA5545 | PA5545 | N/A | N/A | PA5545 | N/A | PA5545 | PA5545 | PA5545 | N/A | PA5545 | PA5545 | N/A | N/A |
| PA5546 | N/A | N/A | N/A | PA5546 | PA5546 | PA5546 | N/A | N/A | N/A | PA5546 | PA5546 | PA5546 | PA5546 | N/A | N/A | N/A | N/A | N/A |
| PA5547 | N/A | N/A | N/A | N/A | N/A | N/A | N/A | N/A | N/A | N/A | N/A | N/A | N/A | PA5547 | N/A | N/A | N/A | N/A |
| N/A | N/A | N/A | N/A | N/A | PA5548 | N/A | N/A | N/A | N/A | N/A | N/A | N/A | N/A | N/A | N/A | N/A | N/A | N/A |
| PA5549 | PA5549 | PA5549 | PA5549 | PA5549 | PA5549 | PA5549 | N/A | N/A | PA5549 | PA5549 | PA5549 | PA5549 | PA5549 | N/A | PA5549 | N/A | N/A | N/A |
| N/A | N/A | N/A | N/A | PA5550 | N/A | N/A | N/A | N/A | N/A | N/A | N/A | N/A | N/A | N/A | N/A | N/A | N/A | N/A |
| PA5551 | N/A | N/A | N/A | N/A | PA5551 | N/A | N/A | N/A | N/A | N/A | N/A | N/A | N/A | N/A | N/A | N/A | N/A | N/A |
| PA5552 | PA5552 | PA5552 | PA5552 | PA5552 | PA5552 | PA5552 | N/A | PA5552 | PA5552 | PA5552 | PA5552 | PA5552 | PA5552 | PA5552 | PA5552 | N/A | N/A | N/A |
| PA5553 | PA5553 | PA5553 | PA5553 | PA5553 | PA5553 | PA5553 | PA5553 | N/A | PA5553 | PA5553 | PA5553 | PA5553 | PA5553 | PA5553 | PA5553 | N/A | PA5553 | PA5553 |
| PA5554 | PA5554 | PA5554 | PA5554 | PA5554 | PA5554 | PA5554 | PA5554 | PA5554 | PA5554 | PA5554 | PA5554 | PA5554 | PA5554 | PA5554 | PA5554 | PA5554 | PA5554 | PA5554 |
| PA5555 | PA5555 | PA5555 | PA5555 | PA5555 | PA5555 | PA5555 | PA5555 | PA5555 | PA5555 | PA5555 | PA5555 | PA5555 | PA5555 | PA5555 | PA5555 | PA5555 | PA5555 | PA5555 |
| PA5556 | PA5556 | PA5556 | PA5556 | PA5556 | PA5556 | PA5556 | PA5556 | PA5556 | PA5556 | PA5556 | PA5556 | PA5556 | PA5556 | PA5556 | PA5556 | PA5556 | PA5556 | PA5556 |
| PA5557 | PA5557 | PA5557 | PA5557 | PA5557 | PA5557 | PA5557 | PA5557 | PA5557 | PA5557 | PA5557 | PA5557 | PA5557 | PA5557 | PA5557 | PA5557 | N/A | PA5557 | PA5557 |
| PA5558 | PA5558 | PA5558 | PA5558 | PA5558 | PA5558 | PA5558 | PA5558 | PA5558 | PA5558 | PA5558 | PA5558 | PA5558 | PA5558 | PA5558 | PA5558 | PA5558 | PA5558 | PA5558 |
| PA5559 | PA5559 | PA5559 | PA5559 | N/A | PA5559 | PA5559 | N/A | PA5559 | PA5559 | PA5559 | PA5559 | PA5559 | PA5559 | PA5559 | N/A | N/A | PA5559 | N/A |
| PA5560 | PA5560 | PA5560 | PA5560 | PA5560 | PA5560 | PA5560 | N/A | PA5560 | PA5560 | PA5560 | PA5560 | PA5560 | PA5560 | PA5560 | PA5560 | N/A | PA5560 | N/A |
| N/A | N/A | N/A | N/A | N/A | PA5561 | N/A | N/A | N/A | N/A | N/A | N/A | N/A | N/A | N/A | N/A | N/A | N/A | N/A |
| PA5562 | PA5562 | PA5562 | PA5562 | PA5562 | PA5562 | PA5562 | PA5562 | PA5562 | PA5562 | PA5562 | PA5562 | PA5562 | PA5562 | PA5562 | PA5562 | N/A | N/A | N/A |
| PA5563 | PA5563 | PA5563 | PA5563 | PA5563 | PA5563 | PA5563 | PA5563 | PA5563 | PA5563 | PA5563 | PA5563 | PA5563 | PA5563 | PA5563 | N/A | N/A | PA5563 | N/A |
| PA5564 | PA5564 | PA5564 | PA5564 | PA5564 | PA5564 | N/A | PA5564 | N/A | PA5564 | PA5564 | PA5564 | PA5564 | PA5564 | N/A | PA5564 | N/A | N/A | N/A |
| PA5565 | PA5565 | PA5565 | PA5565 | PA5565 | N/A | N/A | N/A | PA5565 | PA5565 | PA5565 | PA5565 | PA5565 | PA5565 | N/A | PA5565 | N/A | N/A | N/A |
| N/A | N/A | N/A | N/A | N/A | PA5566 | N/A | N/A | N/A | N/A | N/A | N/A | N/A | N/A | N/A | N/A | N/A | N/A | N/A |
| PA5567 | PA5567 | PA5567 | PA5567 | PA5567 | PA5567 | N/A | N/A | N/A | PA5567 | PA5567 | N/A | N/A | PA5567 | N/A | N/A | N/A | N/A | N/A |
| PA5568 | PA5568 | PA5568 | PA5568 | PA5568 | PA5568 | PA5568 | PA5568 | PA5568 | PA5568 | PA5568 | PA5568 | PA5568 | PA5568 | PA5568 | PA5568 | N/A | PA5568 | PA5568 |
| PA5569 | PA5569 | PA5569 | N/A | PA5569 | N/A | N/A | PA5569 | N/A | PA5569 | PA5569 | PA5569 | PA5569 | N/A | PA5569 | PA5569 | N/A | N/A | N/A |
| PA5570 | PA5570 | PA5570 | PA5570 | PA5570 | PA5570 | PA5570 | PA5570 | N/A | PA5570 | PA5570 | PA5570 | PA5570 | PA5570 | N/A | N/A | N/A | N/A | N/A |
